# A Budding Yeast Model for Human Disease Mutations in the *EXOSC2* Cap Subunit of the RNA Exosome

**DOI:** 10.1101/2020.12.06.413658

**Authors:** Maria C. Sterrett, Liz Enyenihi, Sara W. Leung, Laurie Hess, Sarah E. Strassler, Daniela Farchi, Richard S. Lee, Elise S. Withers, Isaac Kremsky, Richard E. Baker, Munira A. Basrai, Ambro van Hoof, Milo B. Fasken, Anita H. Corbett

**Author notes:** Equal author contribution (Maria C. Sterrett, Liz Enyenihi). Contact information co-corresponding authors: Anita H. Corbett, Department of Biology, RRC 1021, Emory University, 1510 Clifton Road., NE, Atlanta, GA 30322, Milo B. Fasken, Department of Biology, RRC 1081, Emory University, 1510 Clifton Road., NE, Atlanta, GA 30322.

## Abstract

RNA exosomopathies, a growing family of tissue-specific diseases, are linked to missense mutations in genes encoding the structural subunits of the conserved 10-subunit exoribonuclease complex, the RNA exosome. Such mutations in the cap subunit gene *EXOSC2* cause the novel syndrome SHRF (<u>S</u>hort stature, <u>H</u>earing loss, <u>R</u>etinitis pigmentosa and distinctive <u>F</u>acies). In contrast, exosomopathy mutations in the cap subunit gene *EXOSC3* cause pontocerebellar hypoplasia type 1b (PCH1b). Though having strikingly different disease pathologies, *EXOSC2* and *EXOSC3* exosomopathy mutations result in amino acid substitutions in similar, conserved domains of the cap subunits, suggesting that these exosomopathy mutations have distinct consequences for RNA exosome function. We generated the first *in vivo* model of the SHRF pathogenic amino acid substitutions using budding yeast by introducing the *EXOSC2* mutations in the orthologous *S. cerevisiae* gene *RRP4*. The resulting *rrp4* mutant cells have defects in cell growth and RNA exosome function. We detect significant transcriptomic changes in both coding and non-coding RNAs in the *rrp4* variant, *rrp4-G226D*, which models *EXOSC2* p.Gly198Asp. Comparing this *rrp4-G226D* mutant to the previously studied *S. cerevisiae* model of *EXOSC3* PCH1b mutation, *rrp40-W195R*, reveals that these mutants have disparate effects on certain RNA targets, providing the first evidence for different mechanistic consequences of these exosomopathy mutations. Congruently, we detect specific negative genetic interactions between RNA exosome cofactor mutants and *rrp4-G226D* but not *rrp40-W195R*. These data provide insight into how SHRF mutations could alter the function of the RNA exosome and allow the first direct comparison of exosomopathy mutations that cause distinct pathologies.

## Introduction

The RNA exosome is a highly conserved, multi-subunit molecular machine responsible for processing and/or degradation of nearly every class of RNA. First identified in *Saccharomyces cerevisiae* in a screen for ribosomal RNA processing (*rrp*) mutants (Mitchell et al. 1996; Mitchell et al. 1997), the RNA exosome is essential in all systems studied thus far (Mitchell et al. 1997; Lorentzen et al. 2007; Hou et al. 2012; Lim et al. 2013; Pefanis et al. 2014). In addition to ribosomal RNA, the RNA exosome processes a variety of small ncRNAs, including small nuclear RNAs (snRNAs), small nucleolar RNAs (snoRNAs), and transfer RNAs (tRNAs), (Allmang et al. 1999; van Hoof et al. 2000; Kilchert et al. 2016; Fasken et al. 2020). Beyond processing numerous RNAs, the RNA exosome is also required for RNA decay and surveillance, such as the nuclear degradation of cryptic unstable transcripts (CUTs) that result from pervasive transcription (Wyers et al. 2005; Parker 2012; Schneider et al. 2012). This essential RNA processing/degradation machine is composed of nine structural subunits associated with a catalytic 3’-5’ exo/endoribonuclease (Mitchell et al. 1997; Makino et al. 2013). As illustrated in Figure 1A, the 9-subunit structural barrel is composed of an upper ring of three S1/KH cap subunits [(EXOSC1/2/3 (human); Csl4/Rrp4/Rrp40 (budding yeast)] and a lower ring of six PH-like subunits (EXOSC4/7/8/9/5/6; Rrp41/Rrp42/Rrp43/Rrp45/Rrp46/Mtr3). Structural studies in eukaryotic systems have revealed conservation in this structural organization of the complex (Figure 1B) (Liu et al. 2006; Bonneau et al. 2009; Makino et al. 2013; Wasmuth et al. 2014; Zinder et al. 2016), suggesting evolutionary conservation not just within subunit sequence but within overall complex structure and organization.

**Figure 1.**
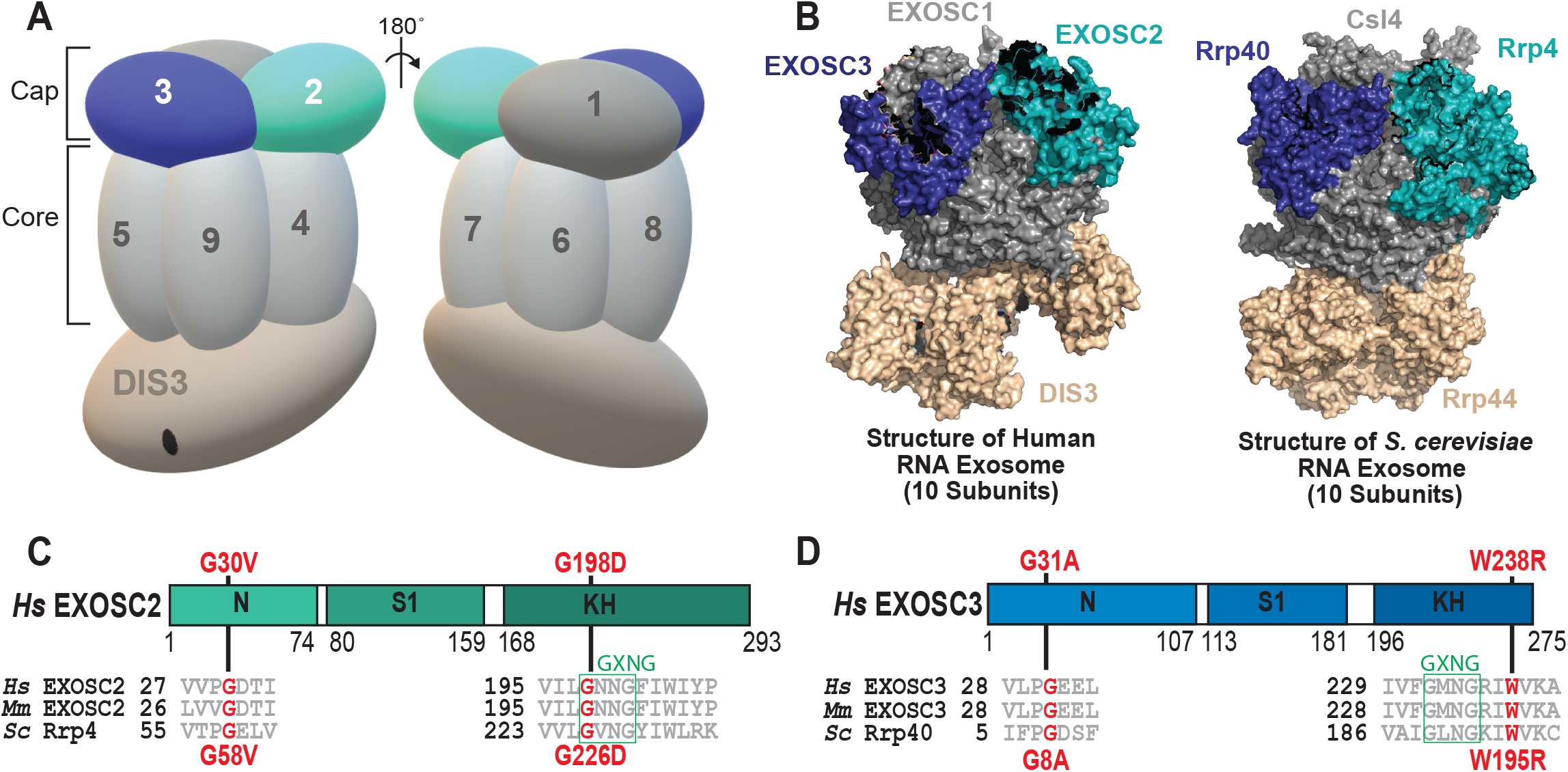
Overview of pathogenic amino acid substitutions in the human cap subunit EXOSC2 of the RNA exosome. (A) The RNA exosome is an evolutionary conserved ribonuclease complex composed of nine structural subunits (EXOSC1-9) and one catalytic subunit (DIS3) that form a “cap” and “core” ring-like structure. The 3-subunit cap at the top of the complex is composed of EXOSC1/Csl4 (Human/*S. cerevisiae*), EXOSC2/Rrp4, and EXOSC3/Rrp40. The 6-subunit core is composed of EXOSC4/Rrp41, EXOSC5/Rrp46, EXOSC6/Mtr3, EXOSC7/Rrp42, EXOSC8/Rrp43, and EXOSC9/Rrp45. The DIS3/Dis3/Rrp44 catalytic subunit is located at the bottom. Missense mutations in the gene encoding the EXOSC2 cap subunit (teal blue, labeled 2,) are linked to a novel syndrome termed SHRF (short stature, hearing loss, retinitis pigmentosa and distinctive facies) (Di Donato et al. 2016). In contrast, missense mutations in the gene encoding the EXOSC3 cap subunit (dark blue, labeled 3) cause PCH1b (pontocerebellar hypoplasia type 1b) (Wan et al. 2012; Biancheri et al. 2013; Eggens et al. 2014; Halevy et al. 2014; Schottmann et al. 2017). (B) The structure and organization of the RNA exosome is highly conserved across eukaryotes. A structural model of the human RNA exosome (left) [PDB 6D6R] (Weick et al. 2018) and the *S. cerevisiae* RNA exosome (right) [PDB 6FS7] (Schuller et al. 2018) are depicted with the cap subunits EXOSC1/Csl4 (Human/*S. cerevisiae*), EXOSC2/Rrp4, and EXOSC3/Rrp40 labeled. (C,D) Domain structures are shown for (C) EXOSC2/Rrp4 and (D) EXOSC3/Rrp40. Each of these cap subunits is composed of three different domains: an N-terminal domain, an S1 putative RNA binding domain, and a C-terminal putative RNA binding KH (K homology) domain. The “GxNG” motif identified in the KH domain of both cap subunits is boxed in green. The position of the disease-linked amino acid substitutions in human EXOSC2 and EXOSC3 are depicted above the domain structures in red. Sequence alignments of EXOSC2/Rrp4 and EXOSC3/Rrp40 orthologs from *Homo sapiens* (*Hs*), *Mus musculus* (*Mm*) and *S. cerevisiae* (*Sc*) below the domain structures show the highly conserved residues altered in disease in red and the conserved Sequences flanking these residues in gray. The amino acid substitutions in *S. cerevisiae* Rrp4 generated in this study and those in *S. cerevisiae* Rrp40, described previously (Fasken et al. 2017; Gillespie et al. 2017), that correspond to the disease-linked amino acid substitutions in human EXOSC2 and EXOSC3 are shown below the structures in red.

Recent studies have linked missense mutations in *EXOSC* genes encoding the structural subunits of the RNA exosome to various human pathologies, which comprise a growing family of diseases termed “RNA exosomopathies” (Wan et al. 2012; Biancheri et al. 2013; Boczonadi et al. 2014; Eggens et al. 2014; Di Donato et al. 2016; Schottmann et al. 2017; Burns et al. 2018; Fasken et al. 2020; Slavotinek et al. 2020). Intriguingly, these single amino acid substitutions often occur in highly conserved domains of the exosome subunits. Mutations in the cap subunit gene *EXOSC3* and the core subunit gene *EXOSC8* cause forms of pontocerebellar hypoplasia (PCH1b and PCH1c, respectively), neurological disorders characterized by atrophy of the pons and cerebellum (Wan et al. 2012; Biancheri et al. 2013; Boczonadi et al. 2014; Eggens et al. 2014; Schottmann et al. 2017; Morton et al. 2018), while mutations in the core subunit genes *EXOSC5* and *EXOSC9* have been linked to similar neurological defects including cerebellar degeneration, neuronopathy and neurodevelopmental delays (Burns et al. 2017; Burns et al. 2018; Slavotinek et al. 2020). In contrast to the other exosomopathy mutations described thus far, missense mutations in the cap subunit gene *EXOSC2* have been linked to a novel, complex disorder characterized by retinitis pigmentosa, progressive hearing loss, premature aging, short stature, mild intellectual disability and distinctive gestalt (Di Donato et al. 2016), later named SHRF (<u>S</u>hort stature, <u>H</u>earing loss, <u>R</u>etinitis pigmentosa and distinctive <u>F</u>acies) (OMIM #617763) (Yang et al. 2019). While SHRF patients do show some cerebellar atrophy (Di Donato et al. 2016), the disease phenotype is distinct from PCH as well as the other neurological deficits observed in patients with other exosomopathies, suggesting a unique molecular pathology linked to *EXSOC2* mutations.

Whole exosome sequencing of the three identified SHRF patients, representing two related patients and one unrelated patient, identified missense mutations in the *EXOSC2* gene that alter conserved amino acids in this cap subunit, shown in Figure 1C (Di Donato et al. 2016). The two related patients have a homozygous missense mutation *EXOSC2* p.Gly30Val (G30V) in the N-terminal domain of EXOSC2 (Di Donato et al. 2016). The other patient carries compound heterozygous missense mutations *EXOSC2* p.Gly30Val and *EXOSC2* p.Gly198Asp (G30V/G198D), the G198D missense mutation is located within the K-homology (KH) RNA binding domain (Di Donato et al. 2016). These amino acid substitutions occur in highly conserved residues of EXOSC2, which are conserved across EXOSC2/Rrp4 orthologs from different eukaryotic species and conserved between EXOSC2 and the EXOSC3/Rrp40 cap subunits of the eukaryotic RNA exosome (Figure S1). Notably, EXOSC2 Gly30 and EXOSC3 Gly31, an amino acid that is substituted in PCH1b patients (Wan et al. 2012), are conserved and in the same position in the two the cap subunits, falling within a conserved “VxPG” consensus sequence (Figure S1). EXOSC2 Gly198 and EXOSC3 Trp238, an amino acid that is also substituted in PCH1b patients (Wan et al. 2012), lie in the KH domains of the two cap subunits, falling within or adjacent to the conserved “GxNG” motif. The “GxNG” motif is unique to the KH domain of these RNA exosome cap subunits and is predicted to play a structural role (Oddone et al. 2007). Yet when EXOSC2 Gly30, EXOSC2 Gly198 and EXOSC3 Gly31, EXOSC3 Trp238 are substituted, they give rise to distinct diseases phenotypes, suggesting that similar missense mutations in *EXOSC2* and *EXOSC3* have different mechanist effects on the RNA exosome *in vivo*. Therefore, to better understand the molecular pathology of these exosomopathies, including SHRF, it is necessary to investigate the molecular and functional consequences of these pathogenic amino acid substitutions that underlie each disease.

A previous study provided some important insights into how the *EXOSC2* mutations that cause SHRF could contribute to pathology (Yang et al. 2019). This study employed several different approaches, including using patient B-lymphoblasts, *in vitro* cell culture and a *D. melanogaster* model depleted for the fly EXOSC2/Rrp4 ortholog. Taken together, results from this study suggest that EXOSC2 dysfunction could compromise downstream molecular pathways, including neurodevelopment and autophagy (Yang et al. 2019). While informative in probing the molecular pathology that may underlie the SHRF syndrome, a limitation of this study is that the authors did not examine known targets of the RNA exosome nor did they examine the *in vivo* consequences of the SHRF-linked EXOSC2 variants within a whole organism. Given that this diverse class of RNA exosomopathies arises from amino acid substitutions in structural subunits of a singular complex, assessing defects in RNA exosome function *in vivo* is critical for a holistic understanding of the molecular and functional consequences underlying each disease phenotype. Previous studies have assessed the functional and molecular consequences of exosomopathy-linked *EXOSC3* and *EXOSC5* mutations *in vivo* using yeast and fly genetic model systems (Fasken et al. 2017; Gillespie et al. 2017; de Amorim 2020; Morton et al. 2020; Slavotinek et al. 2020). Utilizing a genetic model system to explore the consequences of the specific amino acid changes that occur in SHRF can provide insight into how RNA exosome function is altered in disease.

To explore the functional consequences of the amino acid substitutions in EXOSC2 that occur in SHRF, we took advantage of the budding yeast model system. We generated variants of the *S. cerevisiae* EXOSC2 ortholog, Rrp4, that model the pathogenic amino acid substitutions and examined their function in budding yeast. Our results show that the yeast *rrp4-G58V* variant, corresponding to the *EXOSC2*-*G30V* variant, is not able to replace the function of the essential *RRP4* gene. In contrast, cells that express the *rrp4-G226D* variant, corresponding to the *EXOSC2*-*G198D* variant, show impaired cell growth and defects in RNA exosome function. Based on RNA-Seq analysis, the *rrp4-G226D* cells show transcriptomic changes that suggest defects in nuclear surveillance by the RNA exosome. A comparison of the RNA transcripts altered in *rrp4-G226D* cells with those in *rrp40-W195R* mutant cells, which models the *EXOSC3*-*W238R* variant identified in PCH1b patients, reveals that these two RNA exosome cap subunit mutants affect some overlapping and some distinct RNA targets. In addition, genetic assays demonstrate that the *rrp4-G226D* mutant exhibits distinct negative genetic interactions with RNA exosome cofactor mutants that are not shared by the *rrp40-W195R* mutant. Combined, these results suggest that amino acid changes in Rrp4 and Rrp40 that model those in EXOSC2 in SHRF and EXOSC3 in PCH1b, respectively, alter the overall function of the RNA exosome through different mechanisms. Taken more broadly, these data suggest that each exosomopathy mutation may cause distinct molecular and functional consequences for the RNA exosome which could underlie the diverse disease pathologies.

## Results

### EXOSC2 amino acid substitutions linked to SHRF are located in conserved domains

To explore how EXOSC2 G30V and EXOSC2 G198D variants could alter the structure of the EXOSC2 protein or the RNA exosome complex, we modeled these EXSOC2 amino acid substitutions using a recent structure of the human RNA exosome [PDB 6D6R (Weick et al. 2018)] (Figure 2A, 2B). Structural modeling shows that the EXOSC2 Gly30 residue is positioned at the interface with EXOSC4 towards the exterior of the RNA exosome complex in a region with little disorder (Figure 2A). The EXOSC2 Gly30 residue is located in a β-turn next to a highly conserved proline (Pro29) and is essential for the region to have the flexibility needed to make the sharp turn observed in the structure. EXOSC2 Gly30 is also adjacent to an aspartic acid (Asp31), which forms a salt bridge with Arg232 of EXOSC4, likely stabilizing the interaction between the two subunits (Figure S2). An amino acid substitution of Gly30 is predicted to alter the β-turn and position the Asp31 residue away from Arg232 such that the salt bridge would be disrupted. In addition, the EXOSC2 G30V substitution introduces a significantly larger valine residue which appears to clash with residues Asp154 and Ala191 in EXOSC4 (Figure 2A) which could also negatively impact the interactions between the cap EXOSC2 and core EXOSC4 subunits. In contrast, EXOSC2 Gly198 is positioned in a dense region of the subunit, surrounded by four β sheets (Figure 2B). The EXOSC2 G198D substitution introduces a significantly larger aspartic acid residue which appears to clash with neighboring residues Val85 and Asn200 (Figure 2B) and could impede the ability of EXOSC2 to fold properly. In addition, the EXOSC2 G198D substitution introduces a polar aspartic acid residue in place of glycine with an electronegative oxygen that would undergo repulsion with the oxygen of Asn200, making the folding and structure seen in Figure 2B extremely unlikely.

**Figure 2.**
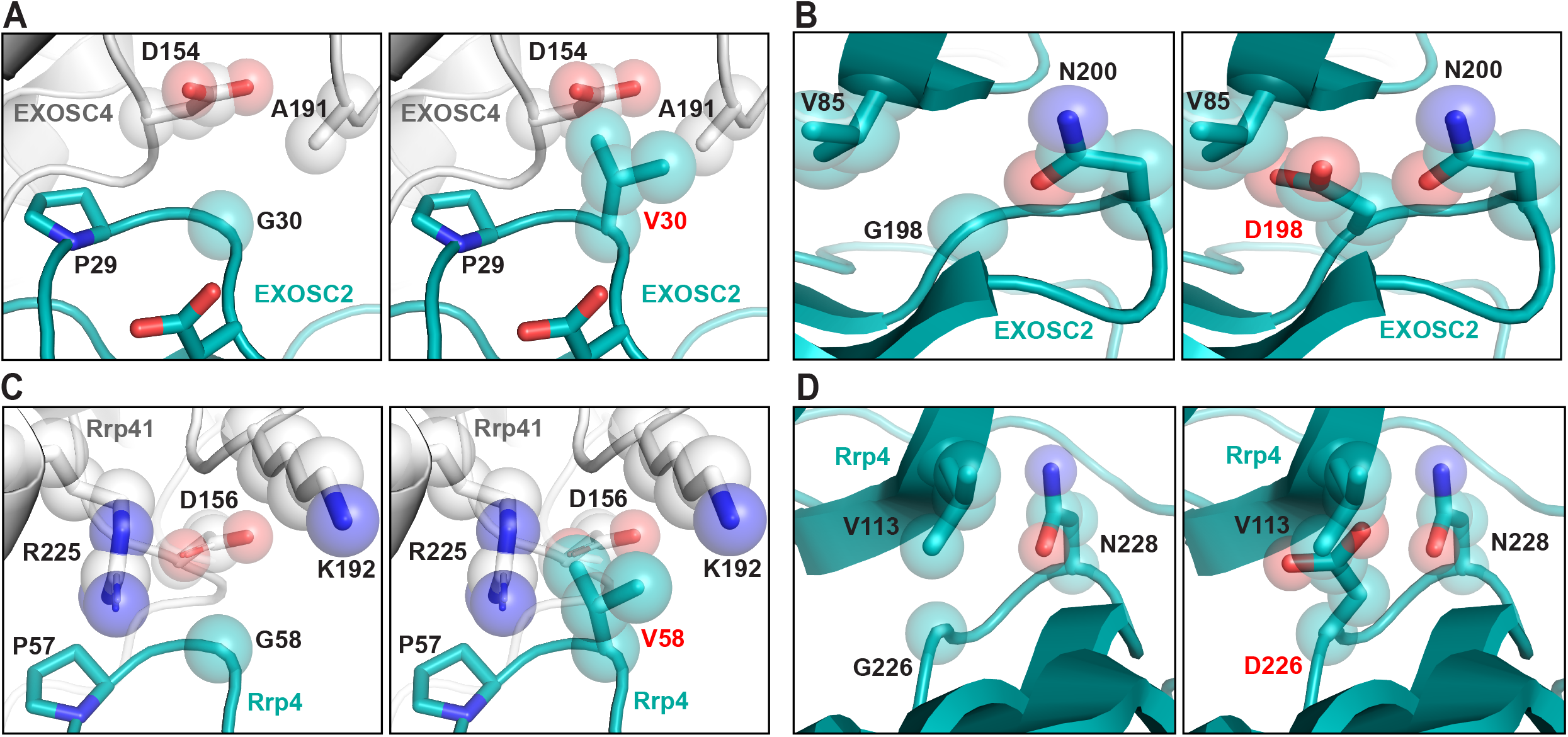
Modeling pathogenic amino acid substitutions in Human EXOSC2 and *S. cerevisiae* Rrp4. (A) Structural modeling of the EXOSC2 p.Gly30Val (G30V) amino acid substitution identified in patients with SHRF syndrome. Zoomed-in representations of the interface between EXOSC2 (teal blue) and EXOSC4 (light gray) modeling the native EXOSC2 Gly30 (G30) residue (left) or the pathogenic EXOSC2 Val30 (V30) residue (right) are depicted. The EXOSC2 Gly30 residue is located in the N-terminal domain of EXOSC2, near the interface of EXOSC2 with the core subunit, EXOSC4. (B) Structural modeling of EXOSC2 p.Gly30Val amino acid change in budding yeast Rrp4 (G58V). Zoomed-in representations of the interface between Rrp4 (teal blue) and the budding yeast EXOSC4 ortholog, Rrp41 (light gray), modeling the native Rrp4 Gly58 (G58) residue (left) or the modeled pathogenic Rrp4 Val58 (V58) residue (right) are shown. Rrp4 Gly58 residue is conserved between human and yeast and, similar to EXOSC2 Gly30, is located in the N-terminal domain of Rrp4, near the interface of Rrp4 with the core subunit, Rrp41. (C) Structural modeling of the EXOSC2 p.Gly198Asp (G198D) amino acid substitution. Zoomed-in representations of EXOSC2 modeling the native EXOSC2 Gly198 (G198) residue (left) or the pathogenic EXOSC2 Asp198 (D198) residue (right) are shown. The EXOSC2 Gly198 residue is located in the KH-domain of EXOSC2 within a dense region of the protein, surrounded by four β-sheets. (D) Structural modeling of the EXOSC2 p.Gly198Asp amino acid change in Rrp4 (G226D). Zoomed-in representations of Rrp4 modeling the native Rrp4 Gly226 (G226) residue (left) or the modeled pathogenic Rrp4 Asp226 (D226) residue (right) are shown. Rrp4 Gly226 residue, which is conserved between human and yeast, is located in the KH-domain of Rrp4 within a dense region of the protein, surrounded by four β-sheets Structural modeling in (A) and (C) was performed with the human RNA exosome structure (PDB 6D6R) (Weick et al. 2018) and in (B) and (D) with the yeast RNA exosome structure (PDB 6FSZ) (Schuller et al. 2018) using PyMOL (PyMOL).

The online server mCSM-PPI2 was used to calculate the change in Gibbs free energy (ΔΔG) to predict the effect of the EXOSC2 amino acid substitutions on protein-protein interactions. Consistent with observations from structural modeling, the software predicts destabilizing changes in the affinity of the protein-protein interactions for both EXOSC2 G30V (ΔΔG=-1.012 Kcal/mol) and EXOSC2 G198D (ΔΔG=-0.509 Kcal/mol). The EXOSC2 G198D substitution is also predicted to reduce protein stability (Score = 1.000 Polymorphism Phenotyping v2). These predictions are consistent with previous work showing EXOSC2 G198D has reduced stability compared to wild-type EXOSC2 (Yang et al. 2019). Furthermore, both substitutions are strongly predicted to have deleterious effects on EXOSC2 function (G30V score −7.938 and G198D score −6.35 calculated by PROVEAN; G30V score 91, G198D score 94 calculated by SNAP-2).

To model the pathogenic amino acid substitutions in the budding yeast EXOSC2 ortholog, Rrp4, we used a recent structure of the *S. cerevisiae* RNA exosome [PDB 6FSZ (Schuller et al. 2018)] (Figure 2C and 2D). Structural modeling shows that the Rrp4 Gly58 residue, corresponding to EXOSC2 Gly30, is positioned at the interface with Rrp41, the budding yeast EXOSC4 ortholog, and is located in a β-turn next to a highly conserved proline (Pro57) in Rrp4, similar to the human structural model (Figure 2C). Rrp4 Gly58 is situated next to a glutamic acid (Glu59) in Rrp4 that forms a salt bridge with Arg233 of Rrp41, similar to the EXSOC2-EXOSC4 interface (Figure S2). The Rrp4 Gly58 residue is also predicted to be essential for the flexibility of the region, facilitating the β-turn, stabilizing the Rrp4-Rrp41 interface. The budding yeast Rrp4-Rrp41 interface does differ from the orthologous human EXOSC2-EXOSC4 interface due to the highly charged nature of the Rrp41 residues Arg225, Asp156 and Lys192. Notably, Rrp41 Asp156 is conserved across eukaryotes and corresponds to EXOSC4 Asp154. However, Rrp41 Lys192 and Arg225 are not highly conserved, suggesting subunit interactions within the RNA exosome of different species vary biochemically. Structural modeling of the Rrp4 Gly226 residue, corresponding to EXOSC2 Gly198, shows that this residue is positioned in a dense region of Rrp4, surrounded by four β sheets (Figure 2D). The residues neighboring Rrp4 Gly226, Val113 and Asn228, are highly conserved and correspond to EXOSC2 Val85 and Asn200, suggesting that the budding yeast Rrp4 G226D substitution can accurately model the structural changes predicted for the human EXOSC2 G198D substitution.

Both Rrp4 G58V (which models EXOSC2 G30V) and Rrp4 G226D (which models EXOSC2 G198D) are predicted to have reduced protein stability (Score = 1.000 Polymorphism Phenotyping v2) as well as deleterious effects on function (G58V score −8.981 and G226D score −6.517 calculated by PROVEAN). Rrp4 G58V is likely to alter the native protein (score 63 calculated by SNAP2), though to a slightly lower degree than calculated for the human the EXOSC2 G30V variant. However, Rrp4 G226D likely results in change to the native protein (score 92 by SNAP2), mirroring the strong effect predicted for the human EXOSC2 G198D variant. In conclusion, these *in silico* predictions (summarized in Supplementary Table S3) suggest that the pathogenic amino acid substitutions have molecular consequences that could affect RNA exosome function in both human and budding yeast.

### *Saccharomyces cerevisiae* Rrp4 variants that model the pathogenic EXOSC2 variants impair its function

To assess the *in vivo* consequences of the pathogenic amino acid substitutions in EXOSC2, G30V and G198D, we generated the corresponding amino acid changes in the *S. cerevisiae* ortholog Rrp4, G58V and G226D. As all core RNA exosome subunits genes are essential in budding yeast (Allmang et al. 1999), we first assessed whether these *rrp4* mutant gene variants can replace the essential *RRP4* gene. In a plasmid shuffle assay, *rrp4Δ* cells containing a *RRP4* maintenance plasmid and *rrp4-G58V* or *rrp4-G226D* variant were serially diluted and spotted on 5-FOA plates to select for cells that harbor the *rrp4* variants as the sole copy of *RRP4* (Figure 3A). The *rrp4-G58V* mutant cells are not viable at any temperature tested, whereas the *rrp4-G226D* cells exhibit impaired growth defect at 37°C as compared to control *RRP4* cells (Figure 3A). The impaired growth of *rrp4-G226D* mutant cells was further analyzed by serial dilution and spotting on solid minimal media (Figure 3B) and in a liquid media growth assay (Figure 3C). On solid media and in liquid culture, the *rrp4-G226D* cells show impaired growth at 37°C compared to control *RRP4* cells (Figure 3B, 3C). For comparison, we also assessed the growth of the previously characterized *rrp40-W195R* cells (Fasken et al. 2017; Gillespie et al. 2017), which corresponds to *EXOSC3-W238R* variant linked to PCH1b. The *rrp4-G226D* cells exhibit a more profound growth defect than *rrp40-W195R* cells at 37°C (Figure 3B, 3C).

**Figure 3.**
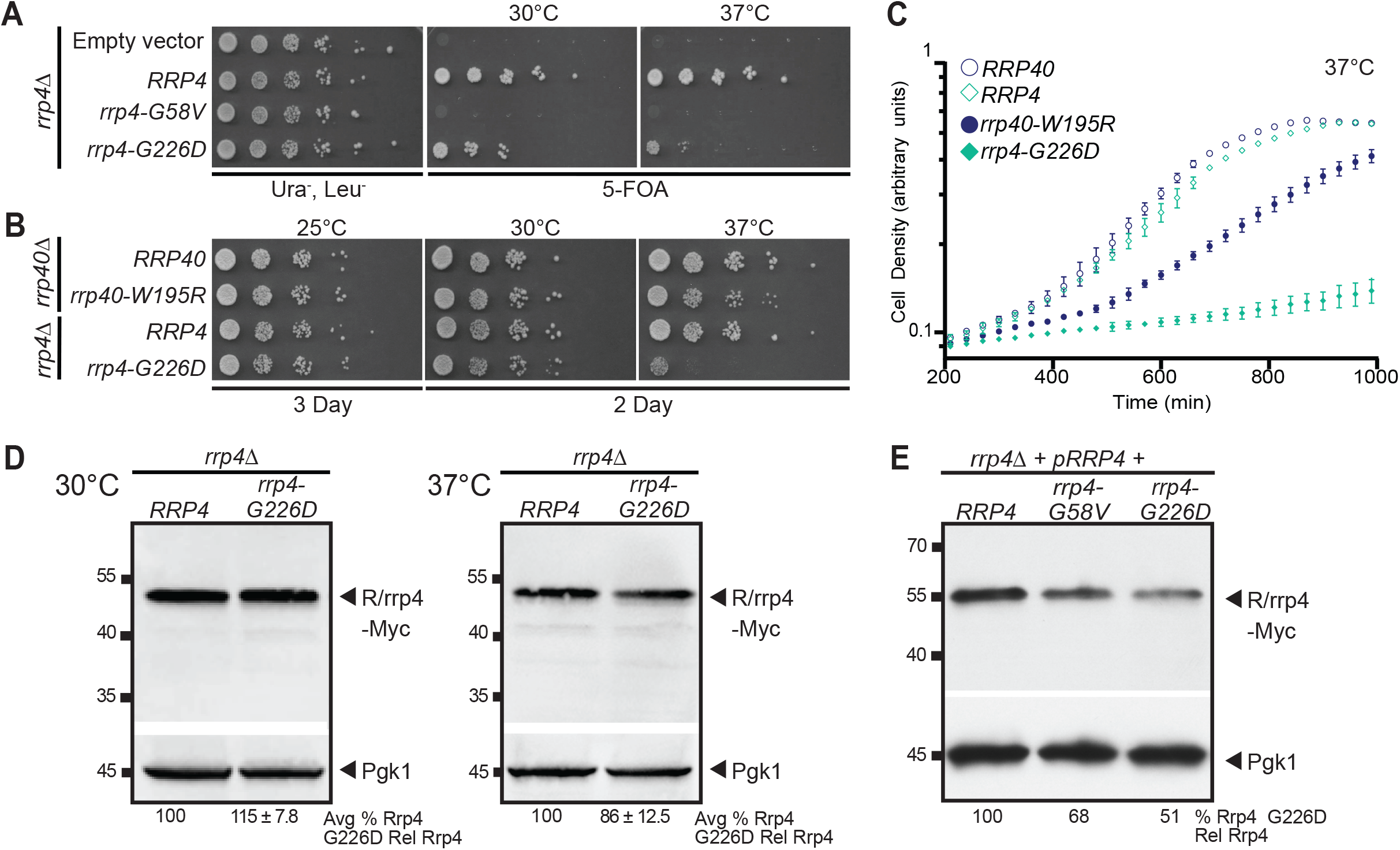
*S. cerevisiae* Rrp4 variants that model EXOSC2 variants identified in patients show impaired function. *S. cerevisiae* cells expressing Rrp4 variants that model pathogenic amino acid changes found in EXOSC2 were generated as described in *Materials and Methods*. (A) Although cells growth is comparable for all mutants that contain a wild-type *RRP4* maintenance plasmid (Ura^-^ Leu^-^), *rrp4-G58V* mutant cells are not viable on plates containing 5-FOA where the maintenance plasmid is not present. *rrp4-G226D* cells show temperature sensitive growth on 5-FOA relative to control *RRP4* cells. Cells were grown at the indicated temperatures. (B, C) The *rrp4-G226D* cells exhibit profoundly impaired growth compared to control *RRP4* cells at 37°C as assessed by (B) serial dilution growth assay on plates or (C) growth in liquid media. (B) *rrp4Δ* cells expressing *RRP4* or *rrp4-G226D* and *rrp40Δ* cells expressing *RRP40* or *rrp40-W195R* were serially diluted, spotted onto solid media grown at the indicated temperatures or (C) grown in liquid media at 37°C with optical density measurement used to assess cell density over time. The growth of *rrp40-W195R* cells, previously reported to be moderately impaired at 37°C (Fasken et al. 2017; Gillespie et al. 2017), was included as a comparative control. (D) The steady-state level of the Rrp4 G226D protein variant is modestly decreased at 37°C. Lysates of *rrp4Δ* cells solely expressing Myc-tagged wild-type Rrp4 or rrp4-G226D grown at 30°C or 37°C were analyzed by immunoblotting with an anti-Myc antibody to detect Rrp4-Myc and an anti-Pgk1 antibody to detect 3-phosphoglycerate kinase as a loading control. The value for the average percentage of rrp4-G226D or Rrp4 protein detected relative to wild-type Rrp4 with standard error from four experiments on different biological replicates (n=4) is shown below each lane. (E) The Rrp4-G58V protein variant is expressed and the steady-state level of the rrp4-G226D protein variant is decreased in cells co-expressing wild-type Rrp4. Lysates of *rrp4Δ* cells co-expressing untagged wild-type Rrp4 and Myc-tagged wild-type Rrp4, rrp4-G58V, or rrp4-G226D grown at 30°C were analyzed by immunoblotting with an anti-Myc antibody to detect Rrp4-Myc and anti-Pgk1 antibody to detect 3-phosphoglycerate kinase as loading control. The percentage of Myc-tagged rrp4-G58V, rrp4-G226D, or Rrp4 relative to Myc-tagged wild-type Rrp4 from a single experiment but representative of many independent experiments is quantitated below each lane. Quantitation of immunoblots in (D) and (E) was performed as described in *Materials and Methods*.

The growth defects associated with the *rrp4* mutant cells could be due to a decrease in the level of the essential Rrp4 protein, as shown for human EXOSC2 G198D (Yang et al. 2019). To explore this possibility, we examined the expression of Myc-tagged wild-type Rrp4 and Rrp4 G226D as sole copy of the Rrp4 protein in *rrp4Δ* cells grown at either 30°C or 37°C. Immunoblotting reveals that the steady-state level of Rrp4 G226D is comparable to wild-type Rrp4 at 30°C; however, at 37°C, the level of Rrp4 G226D is slightly decreased to ∼86% of that of wild-type Rrp4 (Figure 3D). As Rrp4 G58V does not support cell viability, we could not examine the expression of this variant as the sole copy of Rrp4 in cells. Thus, we examined the expression of Myc-tagged Rrp4, Rrp4 G58V, and Rrp4 G226D in *rrp4Δ* cells containing the *RRP4* maintenance plasmid. Under these conditions, where an untagged copy of Rrp4 is present, the steady-state level of Rrp4 G58V-Myc is decreased to 68% and Rrp4 G226D-Myc is decreased to is 51% of that of the wild-type Rrp4-Myc (Figure 3E). These data suggest that pathogenic amino acid substitutions in Rrp4 only modestly impact the level of the Rrp4 protein.

### The Rrp4 G226D variant impairs RNA exosome function

To assess the function of the RNA exosome in *rrp4-G226D* cells, we examined the steady-state level of several of well-defined RNA exosome target transcripts. The RNA exosome has a critical role in ribosomal RNA (rRNA) processing, specifically processing 7S pre-rRNA into mature 5.8S rRNA (Mitchell et al. 1996; Allmang et al. 1999). We analyzed the processing of 5.8S rRNA in *rrp4-G226D* cells using northern blotting. We also compared 5.8S rRNA processing in *rrp4-G226D* cells to yeast cells modeling *EXOSC3* PCH1b mutations, *rrp40-G8A* and *rrp40-W195R* (Fasken et al. 2017; Gillespie et al. 2017). As shown in Figure 4A, *rrp4-G226D* cells accumulate 7S pre-rRNA, a precursor of mature 5.8S rRNA. In addition, several intermediate precursors of 5.8S rRNA, indicated by asterisks, accumulate in *rrp4-G226D* cells. Despite the accumulation of precursors, the level of mature 5.8S rRNA does not appear to differ in *rrp4-G226D* cells compared to control *RRP4* cells. Interestingly, the accumulation of 7S pre-rRNA and other 5.8S rRNA precursors in *rrp4-G226D* cells is greater than that detected in *rrp40-W195R* cells (Figure 4A), which have been previously shown to have accumulation this rRNA precursor (Gillespie et al. 2017).

**Figure 4.**
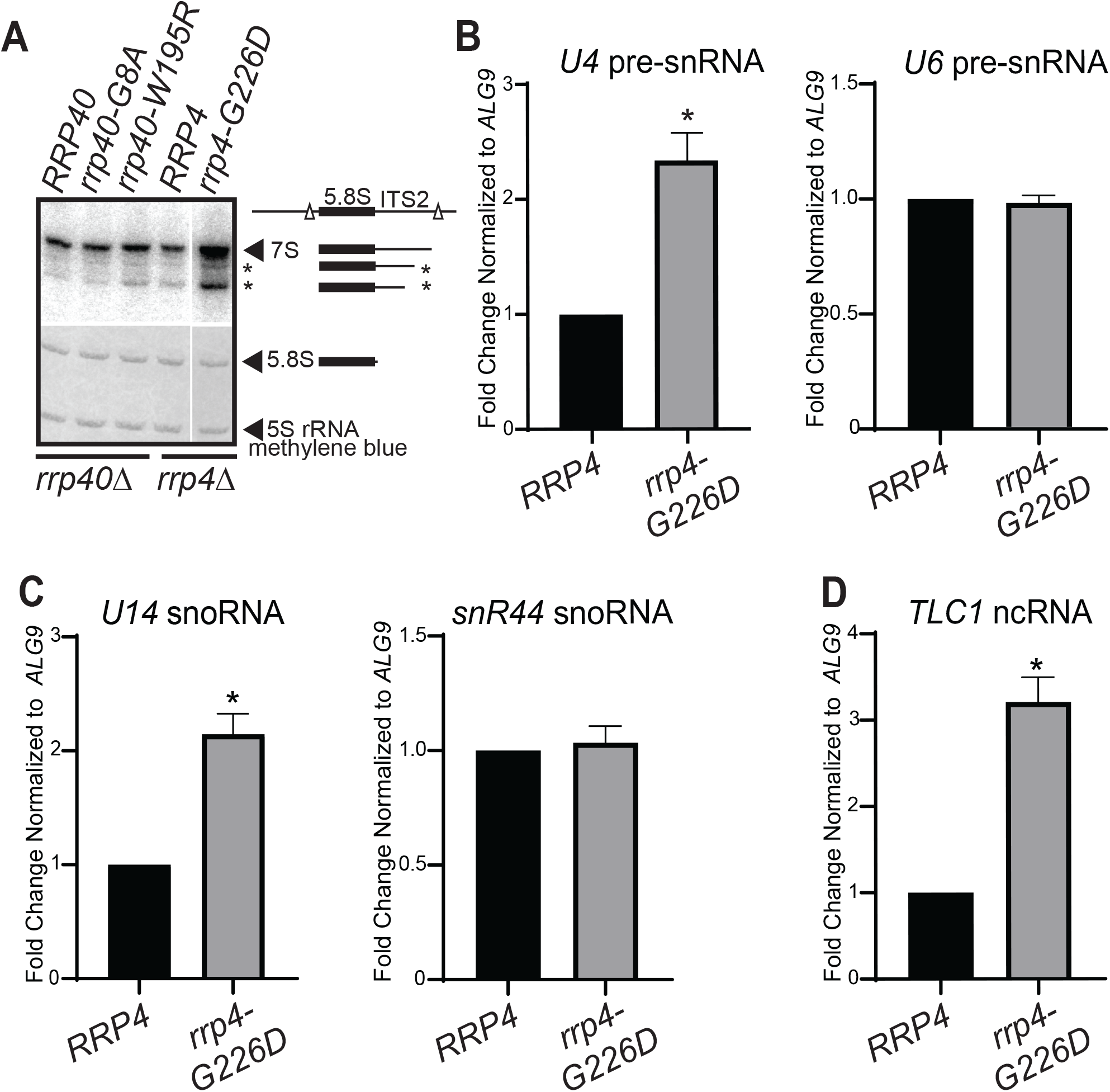
The *rrp4-G226D* variant cells show elevated levels of some but not all RNA exosome target transcripts. (A) The *rrp4-G226D* cells exhibit greater accumulation of 7S pre-RNA compared to *RRP4* and *rrp40-W195R* cells grown at 37°C. Total RNA from *RRP40, rrp40-G8A, rrp40-W195R, RRP4*, and *rrp4-G226D* cells grown at 37°C was analyzed by northern blotting with an 5.8S-ITS2 probe to detect 7S pre-rRNA. Mature 5.8S rRNA and 5S rRNA was detected by methylene blue staining as a loading control. The 7S pre-rRNA is normally processed to mature 5.8S rRNA by 3’-5’ decay of the internal transcribed spacer 2 (ITS2) via the nuclear RNA exosome (Mitchell et al. 1996; Allmang et al. 1999). All lanes imaged from same northern blot with a gap in the loading indicated by the white line. The simplified schematics to the right illustrate the processing steps of 7S precursor following endonucleolytic cleavage from larger 27S precursor (indicated by white triangles) (B) The *rrp4-G226D* cells show an elevated steady-state level of 3’-extended pre-*U4* snRNA but not 3’-extended pre-*U6* snRNA relative to *RRP4* cells at 37°C. (C) The *rrp4-G226D* cells exhibit an increased steady-state level of *U14* (*snR128*) snoRNA but not *snR44* snoRNA relative to *RRP4* cells at 37°C. (D) The *rrp4-G226D* cells show an elevated steady-state level of mature *TLC1* telomerase component ncRNA relative to *RRP4* cells at 37°C. In (B-D), total RNA was isolated from cells grown at 37°C and transcript levels were measured by RT-qPCR using gene specific primers (Table S2), normalized relative to *RRP4*, and graphed as described in *Materials and Methods*. Error bars represent standard error of the mean from three biological replicates. Statistical significance of the RNA levels in *rrp4-G226D* cells relative to *RRP4* cells is denoted by an asterisk (\**p*-value < 0.05).

We next analyzed the steady-state of levels of several RNA exosome target transcripts in *rrp4-G226D* cells using quantitative RT-PCR (Allmang et al. 1999). We measured the steady-state levels of 3’-extended *U4* and *U6* pre-snRNA as well as *U14* and *snR44* snoRNA. The *rrp4-G226D* cells exhibit a significant increase in the level of 3’-extended *U4* pre-snRNA compared to *RRP4* control cells, suggesting 3’-end processing of *U4* snRNA by the RNA exosome is impaired (Figure 4B). In contrast, *rrp4-G226D* cells do not show a significant change in the level of 3’-extended *U6* pre-snRNA (Figure 4B). Like the two snRNAs, the *rrp4-G226D* cells also show a differential effect on the steady-state levels of the two snoRNAs examined. The *rrp4-G226D* cells exhibit a significant increase in the level of *U14* box C/D snoRNA, whereas they show no significant difference in the level of the *snR44* box H/ACA snoRNA relative to *RRP4* cells. We also measured steady-state levels of telomerase component RNA *TLC1*, which is processed by the RNA exosome in a pathway similar to pre-snRNA processing (Coy et al. 2013). The *rrp4-G226D* cells exhibit a significant increase in the steady-state level of mature *TLC1* compared to *RRP4* cells. These data indicate that known RNA exosome target transcripts accumulate in *rrp4-G226D* cells and suggest that Rrp4 G226D impairs the function of the RNA exosome.

### The Rrp4 G226D variant causes broad transcriptomic changes

To further investigate the molecular consequences of the Rrp4 G226D substitution, we performed RNA-Seq analysis on three independent biological replicates of the *rrp4-G226D* and *RRP4* cells as described in *Materials and Methods* (full dataset available as Table S4). Unbiased principal component analysis (PCA) of the resulting RNA-Seq data produced two distinct clusters, indicating that the *rrp4* mutant transcriptome is distinct from the wild-type *RRP4* control (Figure 5A). This separation between the two genotypes and reproducibility amongst the RNA-Seq replicates allowed us to identify transcriptomic changes in *rrp4-G226D* cells (Figure 5B). From differential gene expression analysis, we detect 860 transcripts increased (≥+1.5 Fold Change [FC], p<0.05) and 802 transcripts decreased (≥-1.5 FC, p<0.05) in *rrp4-G226D* cells compared to the *RRP4* control (Figure 5B). Of the 860 transcripts increased, only a third are mRNAs (34.2%, 296 transcripts), with the majority being cryptic unstable transcripts (CUTs), stable unannotated transcripts (SUTs), and other ncRNAs (Figure 5C). Consistent with the role the RNA exosome plays in degradation of nascent ncRNA species, the CUTs and SUTs combined make up the majority (65%) of transcripts that show a steady-state increase in *rrp4-G226D* cells (Figure 5C). Of the 802 transcripts decreased, a majority are mRNAs (89.7%, 719 transcripts) (Figure 5C), with the most significantly decreased transcript (≥-4 FC) being *INO1*, an mRNA that encodes Inositol-3-phosphate synthetase (Donahue and Henry 1981; Klig and Henry 1984) and has previously been characterized as regulated directly by the RNA exosome (Delan-Forino et al. 2017).

**Figure 5.**
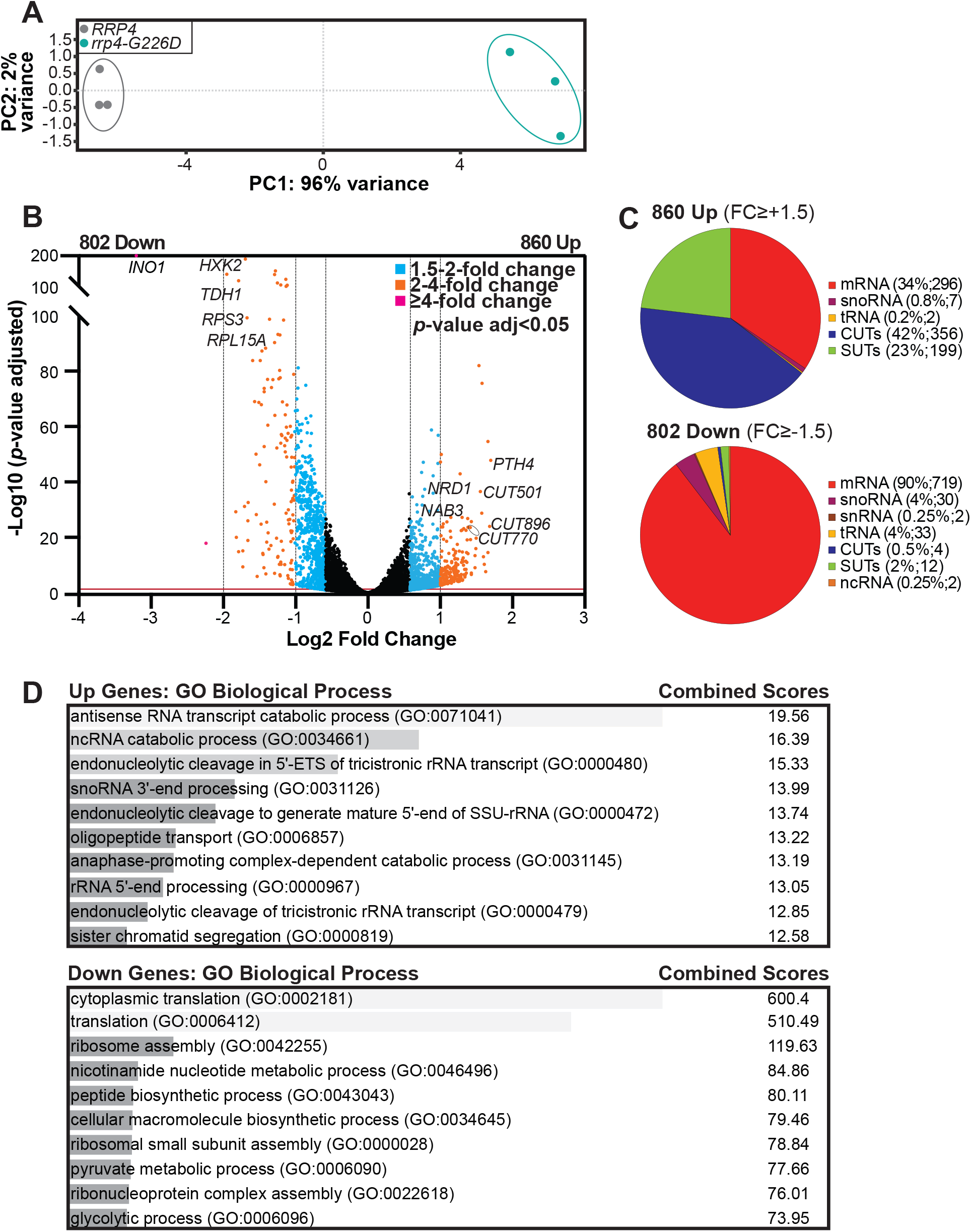
RNA-Seq analysis of *rrp4-G226D* cells reveal distinct transcriptomic changes compared to *RRP4* cells. (A) Principal component analysis (PCA) of RNA-seq data collected from triplicate *RRP4* and *rrp4-G226D* cell samples shows that the gene expression patterns from independent *rrp4-G226D* samples are similar and thus cluster together, but are distinct from *RRP4* samples, which also cluster together. (B) A volcano plot of the differentially expressed transcripts in *rrp4-G226D* cells compared to *RRP4* cells shows that 860 transcripts are significantly Up and 802 transcripts are Down by 1.5-fold or more in *rrp4-G226D* cells. Statistically significant fold changes in transcript levels (Down or Up) in *rrp4-G226D* cells relative to *RRP4* cells are color coded (1.5-2 FC (blue); 2-4 FC (orange); ≥ 4 FC (purple); *p*-value adjusted < 0.05). Transcripts that were subsequently validated by RT-qPCR are labeled. (C) Pie charts of the percentages of different RNA classes within the 860 Up and 802 Down transcripts in *rrp4-G226D* cells reveal that increased transcripts are predominantly ncRNAs (CUTs; SUTs) and decreased transcripts are predominantly mRNAs. The RNA classes identified include messenger RNA (mRNA), small nuclear RNA (snRNA), small nucleolar RNA (snoRNA), transfer RNA (tRNA), cryptic unstable transcripts (CUTs; small, non-coding RNA), stable unannotated transcripts (SUTs; small, non-coding RNA) and other non-coding RNA (ncRNA; e.g. *TLC1*), (D) Gene ontology (GO) analysis for biological process in the Up and Down transcripts in *rrp4-G226D* cells reveals that ncRNA processing is significantly represented in the Up transcripts and translation is significantly represented in the Down transcripts. GO analysis was performed on coding (mRNA) and non-coding RNA (tRNAs, snoRNAs, and snRNAs) classes using the YeastEnrichr web server (Chen et al. 2013; Kuleshov et al. 2016; Kuleshov et al. 2019). Gray bars represent the statistical significance of the biological process categories computed using combined score listed (log of the *p*-value from the Fisher exact test multiplied by the z-score of the deviation from the expected rank).

Gene Ontology (GO) analysis of the differentially expressed transcripts in *rrp4-G226D* cells using YeastEnrichr (Chen et al. 2013; Kuleshov et al. 2016; Kuleshov et al. 2019) reveals that ncRNA catabolic process is the most significant category for the increased transcripts (Combined score 19.56) and cytoplasmic translation is the most significant biological process category for the decreased transcripts (Combined score 600.4) (Figure 5D). These GO analyses align with the transcripts that are altered, as two significantly decreased mRNAs (≥-1.5 FC), *RPS3* and *RPL15A*, encode components of the ribosome, and two significantly increased mRNAs (≥+1.5 FC), *NRD1 NAB3*, are components of the Nrd1-Nab3-Sen1 (NNS) transcription termination complex. (Steinmetz and Brow 1998; Steinmetz et al. 2001; Wolin et al. 2012; Belair et al. 2018)

To both validate altered gene expression detected in the RNA-Seq analysis and compare the spectrum of transcripts altered in *rrp4*-*G226D* cells to *rrp40-W195R* cells, we measured the levels of a subset of transcripts altered in the RNA-Seq data in these mutant cells by RT-qPCR (Figure 6). We performed this analysis on a select number of coding and non-coding transcripts (labeled in Figure 5B). The steady-state levels of three non-coding CUT transcripts —*CUT501, CUT770, CUT896* (Figure 6A) *—* and three coding mRNAs —*PTH4, NAB3, NRD1* (Figure 6C, 6D)—that increased in the RNA-Seq analysis are significantly increased (p<0.05 at least) in *rrp4-G226D* cells compared to *RRP4* control cells. We also validated decreased steady-state levels of coding RNAs (*RPS3, RPL15A, INO1, HXK2, TDH1)* (p<0.01) in *rrp4-G226D* cells compared to control (Figure 6B, C).

**Figure 6.**
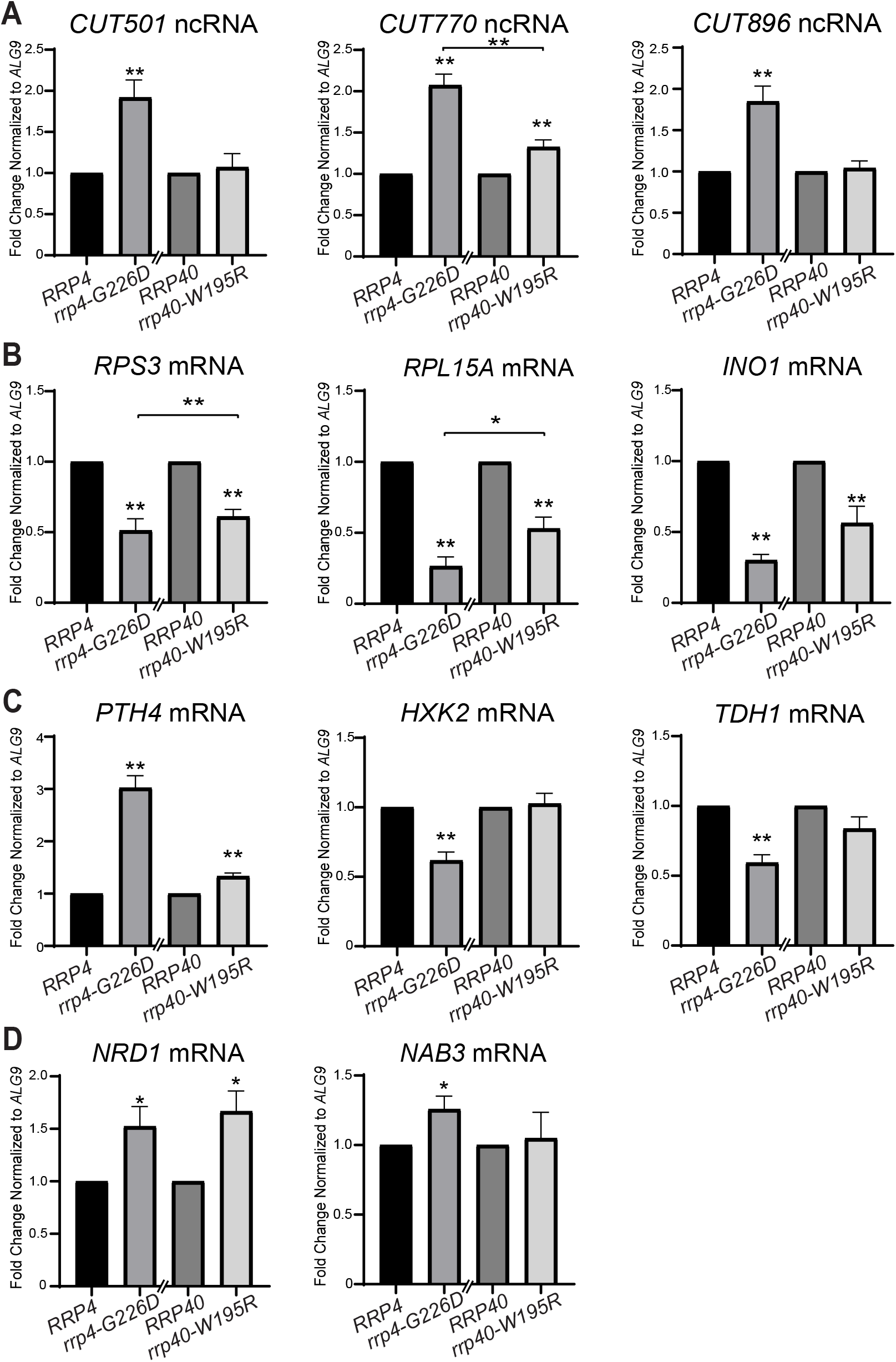
Validation of the differentially expressed transcripts identified in the RNA-Seq confirms that the levels of key mRNAs and CUTs are significantly altered in *rrp4-G226D* cells and reveals that some of these transcripts are not changed in *rrp40-W195R* cells. (A) The steady-state levels of non-coding, cryptic unstable transcripts, *CUT501, CUT770*, and *CUT896*, are significantly increased in *rrp4-G226D* cells compared to control. The *CUT770* level is also increased but the *CUT501* and *CUT896* levels are not altered in *rrp40-W195R* cells. (B) The steady-state levels of ribosomal protein gene mRNAs, *RPS3* and *RPL15A*, and inositol-3-phosphate synthase mRNA, *INO1*, are significantly decreased in *rrp4-G226D* and *rrp40-W195R* cells relative to control *RRP4/40* cells. (C) The steady-state level of peptidyl tRNA hydrolase 4 mRNA, *PTH4*, is significantly increased in *rrp4-G226D* and *rrp40-W195R* cells relative to controls, whereas the levels of hexokinase isoenzyme 2 mRNA, *HXK2*, and glyceraldehyde-3-phosphate dehydrogenase isozyme 1 mRNA, *TDH1*, are significantly decreased in *rrp4-G226D* compared to control. The *HXK2* and *TDH1* levels are not altered in *rrp40-W195R* cells. (D) The steady-state levels of RNA exosome/termination cofactor mRNAs, *NRD1* and *NAB3*, are significantly increased in *rrp4-G226D* cells compared to controls. The *NRD1* level is increased but the *NAB3* level is not altered in *rrp40-W195R* cells. In (A-D), total RNA was isolated from cells grown at 37°C and transcript levels were measured by RT-qPCR using gene specific primers (Table S2), normalized relative to *RRP4/40*, and graphed as described in *Materials and Methods*. Error bars represent standard error of the mean from three biological replicates. Statistical significance of the RNA levels in *rrp4-G226D* and *rrp40-W195R* cells relative to control *RRP4/40* cells or between *rrp4/40* mutants is denoted by asterisks (\**p*-value < 0.05; \*\**p*-value < 0.01).

To compare the molecular consequences resulting from the two pathogenic missense mutations in RNA exosome cap subunits (EXOSC2/Rrp4 and EXOSC3/Rrp40), we expanded the RT-qPCR analysis to include *rrp40-W195R* cells. Intriguingly, we found that some altered targets in *rrp4-G226D* cells were affected in both mutants, while others were significantly affected only in the *rrp4* variant. The steady state levels of *CUT501, CUT770*, and *CUT896* were only significantly increased in *rrp4-G226D* cells and not *rrp40-W195R* cells (Figure 6A). Steady-state levels of coding *RSP3, RPL15A*, and *INO1* mRNAs were significantly decreased in both *rrp-G226D* and *rrp40-W195R* cells compared to control cells (Figure 6B). In contrast, the decrease in steady-state levels of *HXK2* mRNA and *TDH1* mRNA is unique to the *rrp4-G226D* cells, as these coding RNAs were not affected in *rrp40-W195R* cells (Figure 6C). The coding mRNA *PTH4* was significantly increased in *rrp40-W19R5* cells compared to control *RRP40* cells, as observed in *rrp4-G226D* cells, however the magnitude of the change detected was quite different. With respect to the NNS components, *NRD1* steady-state levels change to a similar extent in both *rrp4-G226D* and *rrp40-W195R* cells compared to control; however, the significant increase in *NAB3* steady-state levels occurs only in *rrp4-G226D* cells (Figure 6D). Taken together, these results suggest that amino acid changes in the Rrp4 and Rrp40 cap subunits of the RNA exosome can differentially impact the function of the complex with respect to distinct target RNAs.

### The *rrp4-G226D* mutant shows genetic interactions with nuclear RNA exosome cofactors

The specificity of the RNA exosome for different RNA substrates is conferred by several interacting cofactors, which were first extensively characterized in budding yeast (Schneider and Tollervey 2013; Zinder and Lima 2017). The exonuclease Rrp6, dimerized to its stabilizing partner Rrp47, and Mpp6 are the only cofactors known to directly interact with the RNA exosome, as depicted in Figure 7A (Wasmuth et al. 2017). To determine whether the *rrp4-G226D* variant exhibits genetic interactions with RNA exosome cofactor mutants, we deleted the non-essential, nuclear exosome cofactor genes *MPP6, RRP47* and *RRP6* in combination with *rrp4-G226D*. For comparison, we also determined if the *rrp40-W195R* variant shows genetic interactions with these cofactor mutants by deleting them in combination with *rrp40-W195R*. We examined the growth of these double mutants relative to single mutants (*rrp4-G226D* and *rrp40-*W195R) in solid media growth assays (Figure 7). Interestingly, the *rrp4-G226D mpp6Δ, rrp4-G226D rrp6Δ*, and the *rrp4-G226D rrp47Δ* double mutant cells all exhibit impaired growth compared to *rrp4-G226D* and cofactor single mutants at 30°C (Figure 7A), indicating that deletion of *MPP6, RRP47* or *RRP6* exacerbates the growth defect of *rrp4-G226D* cells. The impaired growth of the *rrp4-G226D rrp6Δ* double mutant is particularly striking. In contrast, *rrp40-W195R mpp6Δ, rrp40-W195R rrp47Δ*, and *rrp40-W195R rrp6Δ* double mutant cells do not show altered growth compared to *rrp40-W195R* or cofactor single mutant cells at 30°C (Figure 7B).

**Figure 7.**
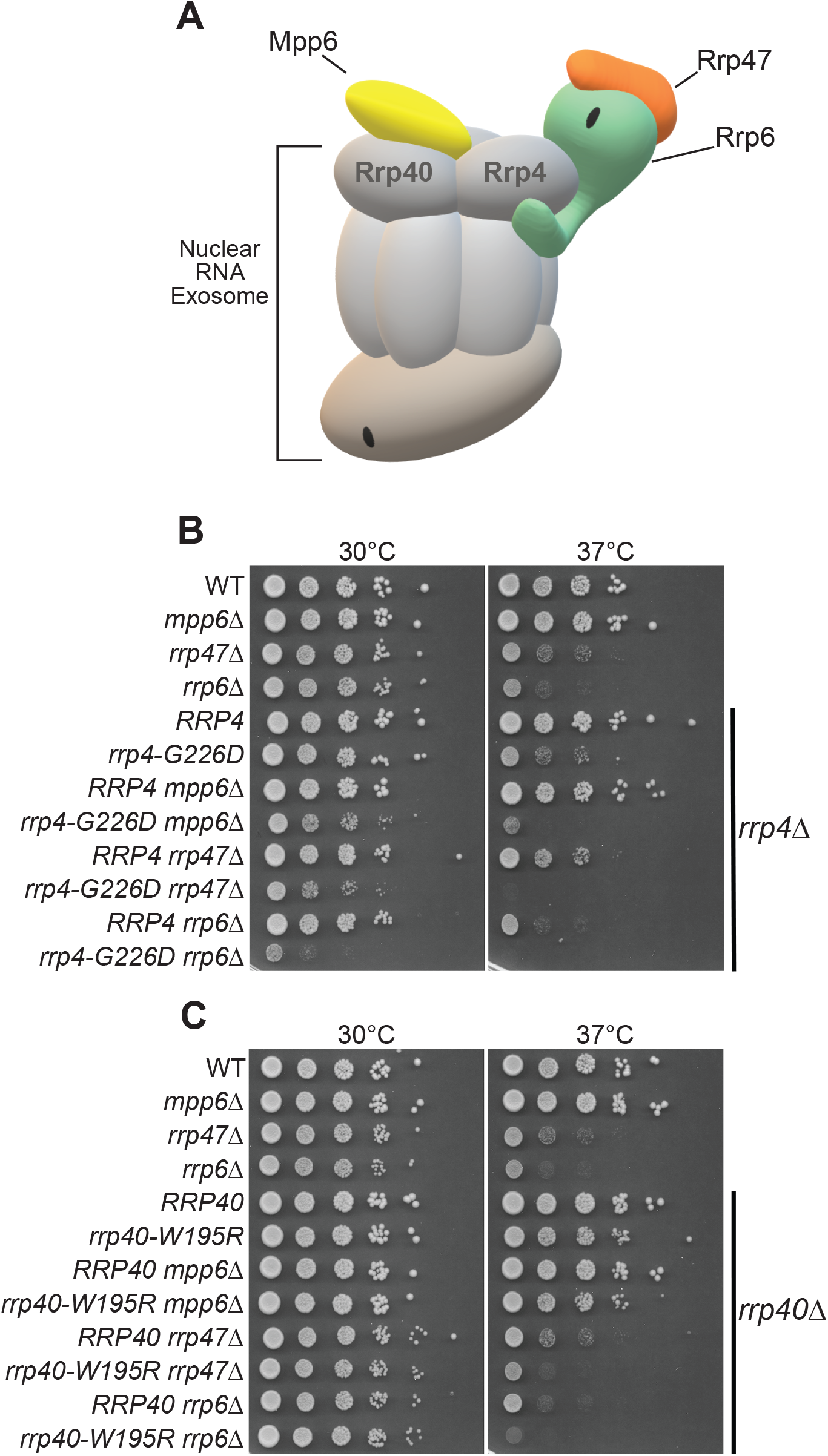
The *rrp4-G226D* mutant exhibits distinct negative genetic interactions with RNA exosome cofactor mutants that are not shared by the *rrp40-W195R* mutant. (A) Double mutant cells containing *rrp4-G226D* and *mpp6Δ, rrp47Δ*, or *rrp6Δ*show a impaired growth compared to single mutants at 30°C and 37°C. The double mutant cells (*rrp4Δ* with *mpp6Δ, rrp47Δ*, or *rrp6Δ*) containing control *RRP4* or *rrp4-G226D* plasmid were serially diluted, spotted onto solid media, and grown at the indicated temperatures for 3 days. (B) Double mutant cells containing *rrp40-W195R* and *mpp6Δ* do not exhibit a change in growth compared to single mutants, whereas double mutant cells containing *rrp40-W195R* and *rrp47Δ* or *rrp6Δ* show impaired growth compared to single mutants at 37°C. The double mutant cells (*rrp40Δ* with *mpp6Δ, rrp47Δ*, or *rrp6Δ*) containing control *RRP40* or *rrp40-W195R* plasmid were serially diluted, spotted onto solid media, and grown at indicated temperatures for 3 days.

The *rrp4-G226D* cofactor double mutants also exhibit enhanced growth defects relative to single mutants at 37°C. The impaired growth of the *rrp4-G226D mpp6Δ* double mutant at 37°C is particularly noteworthy as loss of *MPP6* does not alter cell growth at either 30°C or 37°C in single mutant cells or in double mutant *rrp40-W195R* cells (Figure 7A). The *rrp40-W195R rrp47Δ* and *rrp40-W195R rrp6Δ* double mutant cells do exhibit impaired growth at 37°C, though not substantially worse when compared to the impaired growth of single mutants *rrp47Δ* or *rrp6Δ* at 37°C, as has been previously reported (Briggs et al. 1998; Mitchell et al. 2003) (Figure 7B). These data indicate that the *rrp4-G226D* mutant has distinct negative genetic interactions with *MPP6, RRP47*, and *RRP6* cofactor mutants that are not shared by the *rrp40-W195R* mutant, demonstrating distinct molecular consequences caused by pathogenic amino acid substitutions modeled in Rrp4 as compared to Rrp40. Furthermore, these data suggest the Rrp4 G226D and Rrp40 W195R cap subunit variants could impair RNA exosome function by distinct mechanisms.

## Discussion

In this study, we modeled and analyzed pathogenic amino acid substitutions in the *S. cerevisiae* EXOSC2 ortholog, Rrp4. We generated *rrp4-G58V* and *rrp4-G226D* mutants, which correspond to the SHRF-linked mutations *EXOSC2-G30V* and *EXOSC2-G198D*, respectively. Analysis of the *rrp4-G58V* and *rrp4-G226D* cells reveals that these amino acid substitutions have distinct effects on RNA exosome function. The Rrp4-G58V variant is not able to function as the sole copy of the essential Rrp4 RNA exosome cap subunit as *rrp4-G58V* cells are not viable. In contrast, *rrp4-G226D* cells show a growth defect at 37°C that is more striking than the growth defect previously observed for the *rrp40-W195R* exosome cap subunit mutant, which models a PCH1b-linked *EXOSC3-W195R* mutation (Wan et al. 2012; Fasken et al. 2017; Gillespie et al. 2017). These *rrp4-G226D* cells show significant transcriptomic changes compared to wild-type cells, including increases in steady-state levels of known RNA exosome targets such as 5.8S ribosomal RNA (rRNA) precursors, cryptic unstable transcripts (CUTs) and stable unannotated transcripts (SUTs) (Mitchell et al. 1996; Allmang et al. 1999; Davis and Ares 2006; Xu et al. 2009; Marquardt et al. 2011; Parker 2012; Schneider et al. 2012). A comparison of the two models of RNA exosome cap subunit mutations, *rrp4-G226D* and *rrp40-W195R*, reveals that the two mutations affect some of the same RNA targets but also some distinct targets, suggesting these single amino acid changes in neighboring cap subunits alter the overall function of the RNA exosome through different molecular mechanisms. Genetic analyses support this model, as pathogenic missense mutations in RNA exosome cap subunits show differential genetic interactions with RNA exosome cofactors. These results provide the first *in vivo* model of pathogenic amino acid substitutions that occur in EXOSC2 and allow the first direct comparison between exosomopathy models to reveal that mutations in genes encoding RNA exosome cap subunits have different effects on RNA exosome function.

This study complements prior studies that employed SHRF patient cells and RNAi-mediated depletion of *RRP4* in flies (Yang et al. 2019). We focused our analysis here on the Rrp4-G226D variant, which models the G198D pathogenic missense mutation in *EXOSC2*. The *rrp4-G226D* cells show a growth defect at 37°C that is accompanied by only a modest decrease in steady-state protein levels. Consistent with the Rrp4 G226D substitution impacting the function of the RNA exosome, known RNA exosome targets show altered processing and/or accumulation. A puzzling result from our study is the finding that the *rrp4-G58V* cells are not viable. Of the three SHRF patients identified thus far, two are homozygous for the missense mutation *EXOSC2-G30V* (Di Donato et al. 2016), suggesting that this EXOSC2 variant can provide the function of this essential RNA subunit in humans. From our structural modeling, we do observe biochemical differences between the EXOSC2-EXOSC4 and Rrp4-Rrp41 interface (Figure 2A, 2C). Though the overall structure remains similar, and a stabilizing salt bridge between the two RNA exosome subunits is facilitated by the conserved Gly30 residue in EXOSC2 and Gly58 residue in Rrp4, the differences in charge at the EXOSC2-EXOSC4 and Rrp4-Rrp41 interfaces may be differentially impacted by the valine substitution in the two eukaryotic species. This could account for the difference in viability seen between *rrp4-G58V* budding yeast cells and *EXOSC2-G30V* homozygous human patients. Previous studies suggest the RNA exosome plays an important role in tissue development and human embryonic stem cell differentiation (Belair et al. 2019; Yatsuka et al. 2020), which may be a pathway disrupted by these pathogenic amino acid substitutions that underlie SHRF pathology given the complexity of tissues affected. Therefore, the differential effects observed between *rrp4-G58V* cells and human *EXOSC3-G30V* could be indicative of differences in developmental timepoints or requirements between the two eukaryotes. Integrating additional disease models across other systems will be required to define how pathogenic missense mutations differentially impact RNA exosome function in a tissue-specific manner, leading to diverse disease pathologies.

RNA-Seq analysis of the *rrp4-G226D* mutant cells revealed a broad spectrum of RNA classes that are altered in these mutant cells. The majority of the significantly increased transcripts are comprised of the non-coding RNAs, CUTs and SUTs (64% of all down transcripts with FC≥+1.5). As CUTs and SUTs have previously been shown to be stabilized in RNA exosome mutants and cross-link to the RNA exosome (Wyers et al. 2005; Davis and Ares 2006; Gudipati et al. 2012; Schneider et al. 2012), indicating that these transcripts are direct targets of the RNA exosome. The elevated CUTs and SUTs observed in *rrp4-G226D* cells are therefore consistent with a significant impairment of exosome function due to the Rrp4 G226D substitution. Based on the CUTs and SUTs, other increased transcripts in *rrp4-G226D* cells could thus be direct targets of the RNA exosome and therefore characterization of the elevated transcripts could shed light on the molecular consequences specific to Rrp4 G226D. In contrast, the overwhelming majority of the significantly decreased transcripts are mRNAs (90% of all up transcripts with FC≥-1.5), with the most significantly decreased transcript being *INO1* mRNA. Notably, many ribosomal protein gene (RPG) mRNAs are decreased in *rrp4-G226D* cells and GO analysis of the decreased transcripts revealed cytoplasmic translation to be the most significantly enriched biological process (Figure 5D). Consistent with these data, decreases in RPG mRNAs have also been observed *rrp6Δ* mutant cells (Fox et al. 2015). Many of the decreased mRNA transcripts could be indirect effects that reflect cellular changes that occur when the function of the RNA exosome is compromised, leading to numerous downstream changes. However, *INO1* has been previously shown to be a direct target of the RNA exosome (Delan-Forino et al. 2017), and therefore some of these mRNAs may be directly impacted by defects in RNA exosome function due to pathogenic amino acid substitutions. Previous work in *D. melanogaster* that employed RNAi to deplete Rrp4 identified decreased levels of several transcripts encoding autophagy proteins (Yang et al. 2019). The authors postulated that defective autophagy could contribute to SHRF pathology (Yang et al. 2019). In our RNA-Seq analysis of *rrp4-G226D* cells, we identified 16 autophagy genes that were decreased −1.5-fold (p<0.05) (Figure S3) which could be consistent with this previous study. Further studies will be required to assess whether *rrp4-G226D* cells have impaired autophagy as well to determine whether these transcripts are direct targets of the RNA exosome or due to downstream consequences.

While validating results of the RNA-Seq analysis, we compared effects on specific target RNAs in *rrp4-G226D* cells to the previously characterized *rrp40-W195R* mutant (Fasken et al. 2017; Gillespie et al. 2017). We observed differential effects on the steady-state levels of several CUTs, including *CUT501, CUT707* and *CUT896* in *rrp4-G226D* cells as compared to *rrp40-W195R* cells. The increase in CUTs specifically observed in *rrp4-G226D* cells may suggest defects in nuclear surveillance. We also found differences in transcripts encoding components of the Nrd1-Nab3-Sen1 (NNS) complex, suggesting a distinct difference in NNS complex regulation between the two exosomopathy mutant models. In addition to differences detected in RNA targets, our genetic data lends support to the idea that mutations in *RRP4* and *RRP40* have distinct effects on RNA exosome function. Deletion of the RNA exosome cofactor gene *MPP6* exacerbates the growth defect in *rrp4-G226D* cells (Figure 6A) with no similar genetic interaction detected for the *rrp40-W195R* mutant (Fig. 6B). The nuclear cofactor Mpp6 interacts with the cap subunit EXOSC3/Rrp40, helping to stabilize the interaction between the RNA exosome and the essential RNA helicase Mtr4 (Falk et al. 2017; Wasmuth et al. 2017; Schuller et al. 2018; Weick et al. 2018). Mtr4 aids the RNA exosome in targeting and processing target RNA, such as the 5.8S rRNA precursor (7S rRNA) and is a member of the TRAMP (Trf4/5-Air1/2-Mtr4 Polyadenylation) complex which helps facilitates RNA exosome nuclear surveillance (de la Cruz et al. 1998; LaCava et al. 2005; Vaňáčová et al. 2005; Stuparevic et al. 2013; Schuch et al. 2014; Rodríguez-Galán et al. 2015; Falk et al. 2017). Therefore, the distinct negative genetic interaction in *rrp4-G226D mpp6*Δ cells could suggest destabilization of this critical Mtr4-RNA exosome complex due to the G226D amino acid substitution. This idea is further supported by the significant accumulation of the 7S precursor rRNA observed in *rrp4-G226D* cells compared to the *rrp40-W195R* cells (Figure 4A). Furthermore, loss of *RRP6* and *RRP47* in both *rrp40-W195R* and *rrp4-G226D* cells results in impaired growth at 37°C, but the *rrp4-G226D rrp6Δ* and *rrp4-G226D rrp47Δ* cells also show impaired growth at 30°C relative to *rrp4-G226D* cells. While both exosomopathy mutant models exhibit genetic interactions with *RRP6* and the stabilizing partner *RRP47*, these data could suggest a specific relationship between the nuclear cofactor and the Rrp4 G226D substitution, as *rrp4-G226D* mutant cells are more adversely affected by loss of both genes. This specific relationship between the *RRP6* deletion mutant and the *rrp4-G226D* variant could be due to the loss of the catalytic activity of Rrp6, or the loss of other RNA exosome interactions mediated by Rrp6. Perturbance or destabilization of cofactor-RNA exosome interactions could lead to changes in target RNA levels as these cofactor interactions are imperative for proper targeting and processing/degradation by the complex. Thus, the differential effects on a subset of RNA targets and the differential genetic interactions observed in *rrp4-G226D* and *rrp40-W195R* cells support the conclusion that pathogenic missense mutations in RNA exosome cap subunit genes have distinct consequences for the function of this essential complex.

Utilizing the yeast genetic model system, we have begun to elucidate the distinct functional consequences that result from pathogenic exosomopathy mutations. By modeling these mutations in the corresponding *RRP4* gene, we have generated a system in which to assess the direct effects the pathogenic amino acid substitutions have on the function of the RNA exosome. This study also adds to the growing collection of *in vivo* RNA exosomopathy mutant models that can be compared to one another to catalog the *in vivo* consequences resulting from each mutation. For several RNA exosomopathies, including SHRF syndrome, the patient population is small in number, making analysis with patient tissue samples challenging. Our analyses of *rrp4-G226D* yeast cells and comparison with *rrp40-W195R* provide evidence that the defects in the function of the RNA exosome resulting from the EXOSC2 pathogenic amino acid substitution are distinct from those of the EXOSC3 pathogenic amino acid substitution. These findings can be integrated into the body of work describing the SHRF *EXOSC2* mutations, further expanding our understanding of the unique disease pathology. This study not only provides the first *in vivo* study that models SHRF identified mutations in *EXOSC2* but also provides the first direct comparison of the consequences of pathogenic missense mutations in genes encoding cap subunits of the RNA exosome.

## Materials and Methods

### Chemicals and media

All chemicals were obtained from Sigma-Aldrich (St. Louis, MO), United States Biological (Swampscott, MA), or Fisher Scientific (Pittsburgh, PA) unless otherwise noted. All media were prepared by standard procedures (Adams et al. 1997).

### Protein structure analysis

We used the cryo-EM structure (PDB 6D6R) of the human nuclear RNA exosome at 3.45Å resolution (Weick et al. 2018) and the cryo-EM structure (PDB 6FSZ) of the budding yeast nuclear RNA exosome at 4.6Å (Schuller et al. 2018). Structural modeling was performed using the PyMOL viewer (The PyMOL Molecular Graphics System, Version 2.0 Schrödinger, LLC) (PyMOL). The mCSM-PP12 (Rodrigues et al. 2019), Polymorphism Phenotyping V2 (PolyPhen-2) (Adzhubei et al. 2010), <u>Pro</u>tein <u>V</u>ariation <u>E</u>ffect <u>An</u>alyzer (PROVEAN) (Choi 2012; Choi et al. 2012) and SNAP-2 (Hecht et al. 2015) webservers were used for predicting the effects of the *EXOSC2* mutations on protein stability and function.

### *Saccharomyces cerevisiae* strains and plasmids

All DNA manipulations were performed according to standard procedures (Sambrook et al. 1989). *S. cerevisiae* strains and plasmids used in this study are listed in Table S1 and S2. *S. cerevisiae* strains and plasmids used in this study are listed in Table S1 and S2, respectively. The *rrp4*Δ (yAV1103) and *rrp40*Δ (yAV1107) strains were previously described (Schaeffer et al. 2009; Losh 2018). The *rrp4Δ mpp6Δ* (ACY2471), *rrp4Δ rrp47Δ* (ACY2474), and *rrp4Δ rrp6Δ* (ACY2478) strains and the *rrp40Δ mpp6Δ* (ACY2638), *rrp40Δ rrp47Δ* (ACY2462), *rrp40Δ rrp6Δ* (ACY2466) strains were constructed by deletion of the *MPP6, RRP47*, and *RRP6* ORF in the *rrp4Δ* (yAV1103) and *rrp40Δ* (yAV1107) strains by homologous recombination using *MPP6*-, *RRP47*-, or *RRP6-UTR natMX4* PCR products. Construction of *RRP40-2xMyc* and *rrp40-2xMyc* variant plasmids (pAC3161, pAC3162 and pAC3259) was reported previously (Fasken et al. 2017). The *RRP4-2xMyc LEU2 CEN6* (pAC3474) plasmid was constructed by PCR amplification of the endogenous promoter, 5’-UTR and ORF of the *RRP4* gene from *S. cerevisiae* genomic DNA and cloning into pRS315 plasmid containing a C-terminal 2xMyc tag and the *ADH1* 3’-UTR (Sikorski and Hieter 1989). The *rrp4-G58V-2xMyc* (pAC3476) and *rrp4-G226D-2xMyc* (pAC3477) plasmids were generated by site-directed mutagenesis of the *RRP4-2xMyc* (pAC3474) plasmid using oligonucleotides containing the SHRF syndrome-linked G58V and G226D missense mutations and the QuikChange Site-Directed Mutagenesis Kit (Stratagene). The untagged *RRP4/rrp4-G226D* (pAC3656, pAC3659) and *RRP40/rrp40-W195R* (pAC3652, pAC3655) plasmids and Myc-tagged *RRP4/rrp4-G226D* (pAC3669, pACY3672) plasmid containing native 3’-UTRs were generated by excision of the *2xMyc*-*ADH1* 3’-UTR from each *RRP4/40-Myc LEU2 CEN6* plasmid by restriction digestion and cloning of the native *RRP4* or *RRP40* 3’-UTR into each plasmid using NEBuilder HiFi Assembly (New England BioLabs).

### *S*. *cerevisiae* transformations and growth assays

All yeast transformations were performed according to the standard Lithium Acetate (LiOAc) protocol (Burke et al., 2000). Cells were grown overnight to saturation in a 30°C incubator in liquid YEPD (1% yeast extract, 2% peptone, 2% dextrose, in distilled water). Cell concentrations were normalized to OD_600_ = 0.4 in 10 mL YEPD then incubated at 30°C for 5 hours. The cells were washed with TE/LiOAc then resuspended in TE/LiOAc to a concentration of 2 × 10^9^ cells/mL. To these cells, plasmid DNA, single-stranded carrier DNA, and PEG/TE/LiOAc were added. The cells were agitated for 30 minutes at 30°C before adding DMSO. The cells were heat shocked at 42°C for 15 minutes, washed, and plated onto selective media.

To test the *in vivo* function of the *rrp4* variants that model the *EXOSC2* variants in SHRF syndrome, a standard plasmid shuffle assay was employed. The *rrp4Δ* (yAV1103) cells containing a *RRP4 URA3* covering plasmid and transformed with vector (pRS315), *RRP4-2xMyc* (pAC3474), *rrp4-G8A-2xMyc* (pAC3476), or *rrp4-G226D-2xMyc* (pAC3477) plasmid were grown overnight and serially diluted and spotted onto Ura^-^ Leu^-^ minimal media plates, which select for cells that contain both the covering *RRP4 URA3* plasmid and the *RRP4*/*rrp4 LEU2* plasmid, and 5-FOA Leu^-^ minimal media plates, which select for cells that lack the covering *RRP4 URA3* plasmid and contain only the *RRP4*/*rrp4 LEU2* plasmid. The plates were incubated at 30°C and 37°C for 2 days.

The *in vivo* function of the *rrp4-G226D* variant and the *rrp40-W195R* variant was assessed in growth assays on solid media and in a liquid culture. For growth on solid media, *rrp4Δ* (yAV1103) cells containing only *RRP4* (pAC3656) or *rrp4-G226D* (pAC3659) and *rrp40Δ* (yAV1107) cells containing only *RRP40* (pAC3652) or *rrp40-W195R* (pAC3655) were grown in 2mL Leu^-^ minimal media overnight at 30°C to saturation. Cell concentrations were normalized to OD_600_ = 0.5, serially diluted in 10-fold dilutions, spotted on Leu^-^ minimal media plates, and grown at 25°C, 30°C, and 37°C for 2-3 days. For growth in liquid culture, cells were grown in 2 mL Leu^-^ minimal media overnight at 30°C to saturation, diluted to an OD_600_ = 0.01 in Leu^-^ minimal media in a 24-well plate, and growth at 37°C was monitored and recorded at OD_600_ in a BioTek® SynergyMx microplate reader with Gen5™ v2.04 software over 24 hours. Technical triplicates of each strain were measured, and the average of these triplicates was calculated and graphed.

### Immunoblotting

For analysis of C-terminally Myc-tagged Rrp4 protein expression levels, *rrp4Δ* (yAV1103) cells expressing only Rrp4-2xMyc (pAC3669) or rrp4-G226D-2xMyc (pAC3672) were grown in 2 mL Leu^-^ minimal media overnight at 30°C to saturation and 10 mL cultures with an OD_600_ = 0.2 were prepared and grown at 30°C and 37°C for 5 hr. Additionally, *rrp4Δ* (yAV1103) cells containing *RRP4 URA3* covering plasmid and expressing Rrp4-2xMyc (pAC3474), rrp4-G58V (pAC3476), or rrp4-G226D-2xMyc (pAC3477) were grown in 2 mL Ura^-^ Leu^-^ minimal media overnight at 30°C and 10 mL cultures with an OD_600_ = 0.2 were prepared and grown at 30°C for 5 hr. Cell pellets were collected by centrifugation, transferred to 2 mL screw-cap tubes and stored at −80°C. Yeast cell lysates were prepared by resuspending cell pellets in 0.3 mL of RIPA-2 Buffer (50 mM Tris-HCl, pH 8; 150 mM NaCl; 0.5% sodium deoxycholate; 1% NP40; 0.1% SDS) supplemented with protease inhibitors [1 mM PMSF; Pierce™ Protease Inhibitors (Thermo Fisher Scientific)], and 300 µl of glass beads. Cells were disrupted in a Mini Bead Beater 16 Cell Disrupter (Biospec) for 4 × 1 min at 25°C with 1 min on ice between repetitions, and then centrifuged at 16,000 × *g* for 15 min at 4°C. Protein lysate concentration was determined by Pierce BCA Protein Assay Kit (Life Technologies). Whole cell lysate protein samples (40 µg) were resolved on Criterion 4–20% gradient denaturing gels (Bio-Rad), transferred to nitrocellulose membranes (Bio-Rad) and Myc-tagged Rrp4 proteins were detected with anti-Myc monoclonal antibody 9B11 (1:2000; Cell Signaling). 3-phosphoglycerate kinase (Pgk1) protein was detected using anti-Pgk1 monoclonal antibody (1:30,000; Invitrogen) as a loading control.

### Quantitation of immunoblotting

The protein band intensities/areas from immunoblots were quantitated using ImageJ v1.4 software (National Institute of Health, MD; http://rsb.info.nih.gov/ij/) and mean fold changes in protein levels were calculated in Microsoft Excel (Microsoft Corporation). To quantitate the mean fold change in rrp4-G226D-Myc variant level relative to wild-type Rrp4-Myc level in *rrp4Δ* cells grown at 30°C and 37°C from three immunoblots (Figure 3D) or the fold change in rrp4-G58V-Myc and rrp4-G226D-Myc level in rrp4Δ cells containing untagged *RRP4* from one immunoblot representative of several (Figure 3E), R/rrp4-Myc intensity was first normalized to loading control Pgk1 intensity and then normalized to wildtype Rrp4-Myc intensity at 30°C or 37°C for each immunoblot. The mean fold change in R/rrp4 −Myc level relative to Rrp4-Myc and standard error of the mean were calculated.

### Northern blotting

For analysis of 5.8S pre-rRNA processing - detection of 7S pre-rRNA and processing intermediates - in *rrp4* and *rrp40* mutant cells, *rrp4Δ* (yAV1103) cells containing *RRP4-2xMyc* (pAC3474) or *rrp4-G226D-2xMyc* (pAC3477) and *rrp40Δ* (yAV1107) cells containing *RRP40-2xMyc* (pAC3161), *rrp40-G8A-2xMyc* (pAC3162), or *rrp40-W195R-2xMyc* (pAC3259) were grown in 2 mL Leu-minimal media overnight at 30°C, 10 mL cultures with an OD_600_ = 0.4 were prepared and grown at 37°C for 5 hr. Cells were collected by centrifugation (2,163 x *g*), transferred to 2 mL screw cap tubes and stored at −80°C. Total RNA from cells was resolved on an Criterion TBE-Urea polyacrylamide gel (Bio-Rad), blotted to a nylon membrane and membrane was probed with radiolabeled 5.8S-ITS2 rRNA (boundary) oligonucleotide (AC4211/Probe 020-5’-TGAGAAGGAAATGACGCT) to detect 7S pre-rRNA and intermediates and stained with methylene blue stain to visualize 5.8S rRNA as a loading control. Total RNA (5µg) was mixed with equal volume of RNA loading dye (1xTBE; 12% Ficoll; 7M Urea; 0.01 bromophenol blue; 0.02% xylene cyanol) and resolved on 10% TBE-Urea polyacrylamide gel in 1xTBE at 200V for 1.5 hr. RNA was transferred to Hybond™-N+ nylon membrane (Amersham, GE Healthcare) at 15V for 100 min in 1xTBE and cross-linked to membrane with UV light (120,000 µJoules) using UV Stratalinker® 2400 (Stratagene). Membrane was incubated in Rapid-hyb hybridization buffer (Amersham, GE healthcare) at 37°C for 1 hr. DNA oligonucleotide (100 ng) was 5’-end labeled with [γ-P32]-ATP (PerkinElmer) using polynucleotide kinase (New England Biolabs) at 37°C for 30 min. [P32]-Labeled oligonucleotide probe was purified through G25 microspin column (GE Healthcare), heated at 100°C for 5 min, and added to hybridization buffer. Oligonucleotide probe was hybridized to membrane in hybridization buffer at 37°C overnight. Following removal of hybridization buffer, membrane was rinsed twice in 5 x SSPE; 0.1% SDS at 25°C and washed twice in 0.5 x SSPE; 0.1% SDS at 37°C for 20 min each. Membrane was exposed to phosphoscreen overnight and imaged using Typhoon FLA 7000 phosphoimager (GE Healthcare).

### Total RNA Isolation

Total RNA from *S. cerevisiae rrp4* and *rrp40* mutant cells was isolated using TRIzol (Invitrogen) for qRT-PCR and northern blotting and MasterPure™ Yeast RNA Purification Kit (Epicentre, Lucigen) for RNA-seq. *S. cerevisiae* cells were grown in 2 mL Leu^-^ minimal media overnight at 30°C to saturation. Cultures were diluted in 10 mL Leu^-^ minimal media to an OD_600_ = 0.2 and grown for 5 hours at 37°C. Cells were pelleted by centrifugation, transferred to in 2 mL screw cap tubes, and stored at −80°C. To prepare total RNA using TRIzol, cells were resuspended in 1 mL TRIzol (Invitrogen) with 300 µL of glass beads. Cell samples were disrupted in a Biospec Mini Bead Beater 16 Cell Disrupter for 2 min at 25°C. For each sample, 100 µL of 1-bromo-3-chloropropane (BCP) was added, sample was vortexed for 15 sec, and incubated at 25°C for 2 min. The sample was centrifuged at 16,300 x *g* for 8 min at 4°C, and the upper layer was transferred to a fresh microfuge tube. RNA was precipitated with 500 µL isopropanol and sample was vortexed for 10 sec to mix. Total RNA was pelleted by centrifugation at 16,300 × *g* for 8 min at 4°C. The RNA pellet was washed with 1 mL 75% ethanol, centrifuged at 16,300 × *g* for 5 min at 4°C, and air-dried for 15 min. Total RNA was resuspended in 50 µL diethylpyrocarbonate (DEPC, Sigma)-treated water and stored at −80°C. Total RNA was prepared using MasterPure™ Yeast RNA Purification Kit (Epicentre, Lucigen) according to manufacturer’s protocol. Total RNA was resuspended in in 50 µL DEPC-treated water and stored at −80°C.

### qRT-PCR

For analysis of steady-state RNA levels using quantitative PCR, three independent biological replicates of *rrp4Δ* (yAV1103) cells containing only *RRP4* (pAC3656) or *rrp4-G226D* (pAC3659) and *rrp40Δ* (yAV1107) cells containing only *RRP40* (pAC3652) or *rrp40-W195R* (pAC3655) were grown in 2mL Leu^-^ minimal media overnight at 30°C, 10 mL cultures with an OD_600_ = 0.2 were prepared and cells were grown at 37°C for 5 hr. Total RNA was isolated from cell pellets and 1 μg of total RNA was reverse transcribed to first strand cDNA using the M-MLV Reverse Transcriptase (Invitrogen) according to manufacturer’s protocol. Quantitative PCR was performed on technical triplicates of cDNA (10 ng) from three independent biological replicates using gene specific primers (0.5mM; Table S2), QuantiTect SYBR Green PCR master mix (Qiagen) on a StepOnePlus Real-Time PCR machine (Applied Biosystems; Tanneal=55°C, 44 cycles). *ALG9* was used as an internal control. The mean RNA levels were calculated by the ΔΔCt method (Livak and Schmittgen 2001). Mean levels of RNA calculated in mutant cells are normalized to mean levels in wild-type cells and converted and graphed as RNA fold change relative to wild-type. All primers used are summarized in Table S2.

### RNA-Seq analysis

RNA-Seq was performed on three independent biological replicates of *rrp4*Δ (yAV1103) cells containing *RRP4-2xMyc* (pAC3474) or *rrp4-G226D-2xMyc* (pAC3477) as the sole copy of *RRP4* grown at 37°C. Cells were grown in 2 mL Leu^-^ minimal media overnight at 30°C, diluted to an OD_600_ = 0.4 in 10 mL Leu-minimal media, grown at 37°C for 5 hr, and collected and stored at −80°C. Total RNA was isolated, rRNA was depleted, and stranded cDNA libraries were prepared using TruSeq Total RNA Stranded Library Prep kit (Illumina). Paired-end sequencing of the cDNA libraries was performed on a HiSeq4000 instrument (2 × 150 cycles) at Frederick National Laboratory for Cancer Research (FNLCR) at the CCR Sequencing Facility, NCI, NIH, Frederick, MD. The *RRP4* samples yielded an average of 28,890,739 pass filter reads and the *rrp4-G226D* samples yielded an average of 34,644,683 pass filter reads, with a base call quality of 94% of bases with Q30 and above. The reads were mapped to the *S. cerevisiae* S288C genome assembly R64-1-1, annotated with CUTs and SUTs (Xu et al. 2009), using the STAR RNA-seq aligner [v2.7.5b (Dobin et al. 2012)]. The reads were per gene feature were counted using featureCounts [v1.6.4+galaxy2 (Liao et al. 2014)]. Differential gene expression analysis on raw read counts was performed using DESeq2 [v2.11.40.6+galaxy1(Love et al. 2014)] to identify genes significantly changed (*p*-value<0.05, ≥1.5-fold change) in *rrp4-G226D* samples relative to *RRP4* samples. Principal component analysis (PCA) on raw read counts was also performed using DESeq2. Volcano plot of differential gene expression data was produced in Prism 8 (Graphpad Software). Piecharts of RNA class percentages in significantly altered genes were generated in Microsoft Excel for Mac (Microsoft Corp.). Gene Ontology (GO) analysis on significantly altered genes for Biological Process category was performed using the YeastEnrichr webserver [http://amp.pharm.mssm.edu/YeastEnrichr/ (Kuleshov et al. 2019)]. The full RNA-Seq dataset is compiled in Table S4.

### Genetic Interaction Analysis

To test genetic interactions between *rrp4-G226D* or *rrp40-W195R* and RNA exosome cofactor/subunit deletion mutants, *rrp4Δ mpp6Δ* (ACY2471), *rrp4Δ rrp47Δ* (ACY2474), and *rrp4Δ rrp6Δ* (ACY2478) cells containing only *RRP4* (pAC3656) or *rrp4-G226D* (pAC3659) and *rrp40Δ mpp6Δ* (ACY2638), *rrp40Δ rrp47Δ* (ACY2462), and *rrp40Δ rrp6Δ* (ACY2466) cells containing only *RRP40* (ACY3652) or *rrp40-W195R* (ACY3655) were grown in 2 mL Leu^-^ minimal media overnight at 30°C to saturation, serially diluted, and spotted on Leu^-^ minimal media plates. The plates were incubated at 30°C and 37°C for 3 days.

## Acknowledgements

We thank members of the Corbett and van Hoof laboratories for critical discussions and input. This work was supported by National Institutes of Health (NIH) R01 grants (GM058728) to A.H.C. and (GM099790) to A.v.H. S.B. was supported by NIHR25 GM099644. M.C.S. was supported by a National Institute of General Medical Sciences (NIGMS) F31 grant (GM134649-01). L.E. was supported by the NIH-funded Emory Initiative for Maximizing Student Development. S.E.S. was supported by the National Science Foundation (NSF) Graduate Research Fellowship (GRFP 1937971). M.A.B is supported by the NIH-funded Intramural Research Program of the National Cancer Institute. Lastly, we would like to thank the RNA society and Saccharomyces Genomic Database (SGD) (Cherry et al. 2012) for providing community resources and support for scientific discovery.

