## Supplemental Figures for "A Budding Yeast Model for Human Disease Mutations in the *EXOSC2* Cap Subunit of the RNA Exosome"

**Supplemental Material**

**Supplemental Figure S1. Protein Sequence alignment of human EXOSC2 and EXOSC3.** Protein Sequence alignment of the human RNA exosome cap subunits EXOSC2 and EXOSC3, *S. cerevisiae* Rrp4 and Rrp40, including other EXOSC2/Rrp4 and EXOSC3/Rrp40 orthologs. Conserved EXOSC2 amino acids substituted in patients with SHRF (DI DONATO *et al.* 2016) and EXOSC3 amino acids substituted in patients with PCH1b disease (WAN *et al.* 2012) are highlighted in green. Identical residues (in red) and similar residues (in blue) are indicated. The different species aligned are as follows: Hs: *Homo sapiens*, Mm: *Mus musculus*, Dr: *Danio rerio*, Dm: *Drosophila melanogaster*, Ce: *Caenorhabditis elegans*, Sc: *Saccharomyces cerevisiae*, Sp: *Schizosaccharomyces pombe*, Cn: *Cryptococcus neoformans*, At: *Arabidopsis thaliana*, Os: *Oryza sativa*, Ss: *Sulfolobus solfataricus* (archaea).

**Supplemental Figure S2. Modeling of the human EXOSC2-EXOSC4 and yeast Rrp4-Rrp41 interface show structural conservation.** (A) Zoomed-in representations of the interface between the RNA exosome cap subunit EXOSC2 (teal blue) and core subunit EXOSC4 (light gray). EXOSC2 G30 facilitates a β turn that positions EXOSC2 Asp31 (D31) near an arginine in EXOSC4, R232. Structural modeling shows a salt bridge that forms between EXOSC2 D31 and EXOSC4 R232, represented by the red dashed lines. (B) Zoomed-in representation of the interface between the yeast RNA exosome cap subunit Rrp4 (teal blue) and core subunit Rrp41 (light gray). Rrp4 G58, which corresponds to EXOSC2 G30, facilitates a β turn that positions Rrp4 Glu59 (E59) near an arginine in the EXOSC4 yeast ortholog, Rrp41 (R233). Structural modeling shows salt bridge forms between Rrp4 E59 and Rrp41 R233, represented by the red dashed lines. Structural modeling in (A) was performed with the human RNA exosome structure (PDB 6D6R) (WEICK *et al.* 2018) and in (B) with the yeast RNA exosome structure (PDB 6FSZ) (SCHULLER *et al.* 2018) using PyMOL (PYMOL).

**Supplemental Figure S3. Volcano plot of autophagy transcripts differentially expressed in the *rrp4-G226D* RNA-Seq**. A total of 18 autophagy transcripts show ±1.5-fold change (*p*-value adjusted < 0.05) in *rrp4-G226D* cells compared to *RRP4* cells. Of those, 2 transcripts are increased 1.5-2 fold (*NFT1* and *SNO4* (blue)) and 16 transcripts are decreased 1.5-2-fold (*TEF4, RPL14A, RAS2, EGD1 SEC4, TPS1, TMA19, PEP4, SEC14, MSS51, RPL15B, CAP1, FRQ1, PNP1* (blue)) with two transcripts decreased > 2-fold (*ASC1* and *SOD1* (orange)).

**Table S1**. Yeast Strains and Plasmids used in this study.

**Table S2**. DNA Oligonucleotides employed for RT-qPCR

**Table S3**. Summary of *in silico* predictions for pathogenic amino acid substitutions in EXOSC2 and Rrp4.

**Table S4.** Full RNA-Seq dataset.

### Supplemental Figure S1

**Sterrett Enyenihi *et al.***

```
HsEXOSC2   -----------------------------MAMEMRLPVARK--PL-SERLGR--DTKKHLVVPGDTITTD-------------------------TGFMRGHGTYMGEEK----------------LIASVAGSVERVNKL------ICVKAL
MmRrp4     -----------------------------MALEMRLPKARK--PL-SESLGR--DSKKHLVVPGDTITTD-------------------------TGFMRGHGTYMGEEK----------------LIASVAGSVERVNKL------ICVKAL
DrRrp4     -----------------------------MAVEMRLPAVHK--HV-SLKTGQSSREDKHLVVPGDVITSD-------------------------TGFMRGHGTYMDEDR----------------LTASVAGQVERVNKL------ICVKPL
DmRrp4     -----------------------------MSTNAAIDLALD--RVDWRDLAAQTEEQPRVYTPGEVLMPE-------------------------AGFMRGHGTFVEDEN----------------IKSSVAGVIQKVNKL------ISVRPL
CeRrp4     -------------------MSFEVTGPP----------LVSPWMMLSMSEARNDIDETKIVIPGHSVCDAP------------------------QQFMRGHGTYVRDGE----------------IVSSLSGVVQQLNRL------LMVKTI
ScRrp4     MSEVITITKRNGAFQNSSNLSYNNTGIS---DDENDEEDIY---MHDVNSASKSESDSQIVTPGELVTDD-------------------------PIWMRGHGTYFLDNM----------------TYSSVAGTVSRVNRL------LSVIPL
SpRrp4     ---MVTILKPEEFYVSSEADIVNDVSMTEMEDEIMEDEQMGLVDGEDVLEEFDKSIHQNLVTPGQLVTDD-------------------------PQFMRGHGTYFEDGG----------------IYASVAGSVQRVNKL------ISVKPL
CnRrp4     ---MFSVKAV----APSISQAFTEIVPSTHRK---HQAELDAMDLDEDEFGGAGPSKRSIVSPGEVITSS-------------------------KEYMRGHGTYVEESN----------------VVSSVAGTIERVNKL------ISVRPL
AtRrp4     ------------MVMRKLQLPLSQTQ------KVRFERAIE--RLQSLSS-SANSDASVIVTDSIPVNHD-------------------------DAFLKGHGTSEVDGE----------------LLATVCGVVERVDKL------VYVRTL
OsRrp4     --------------MRDLQLSLNQTQ------RVRLEAALH--ELQTVAP-----AAAVTVADTIPVNDE-------------------------DNILKGHGTSDQDGE----------------VVATLCGVVERVNKL------VYVRTL
HsEXSOC3   ---------------------------------MAEPASVAAESLAGSRARAARTVLGQVVLPGEELLLPEQEDAEGPGGAVERPLSLNARACSRVRVVCGPGLRRCGDR----------------LLVTKCGRLRHKEPGSGSGGGVYWVDS
MmRrp40    ---------------------------------MAEVLSAGPESVAGCRARAVHKVLNQVVLPGEELVLPDHEDVDGLGGAGEQPLRLNAGARPRLRVVCGPGLRRCGDR----------------LLVTKCGRLRHKEPS-GGGGGVYWVDS
DrRrp40    -------------------------------------------MDSSVHTSLLERIGDVVLPGDLLFSFSPPE-------------AGDANPKADRLICGPGLRRSGAE----------------IRVCRAGVLKHKQPN------MYWVNC
DmRrp40    ---------------------------------------------------------MSATSTIVMPGERIAA-------------------IEELAKSKRVILGPGLRRLDDT----------------VVASKAGPLRHKEPG------TFWVDN
CeRrp40    -------------------------------------------------------------MTVYLPGDVINEPSS----------------------SDSSIIGYGISVRGQT----------------RIATQPGAFH-------NDDGKVWLNV
ScRrp40    ------------------------------------------------------------MSTFIFPGDSFPVDP----------------------TTPVKLGPGIYCDPNTQE--------------IRPVNTGVLHVSAKG-KSGVQTAYIDY
SpRrp40    ----------------------------------------------------MAEVEEKGLLSYYFPGERIPDSIIS--------------------QDGSIRLGPGLVFQKKDRE------------EEIVVSKAGRLH-Q-----TGKNAMLIDS
CnRrp40    ------------------------------------------------------------MPTLILPGETVPIPS----------------------TSKAVVLGPGISQSTPTLQASSSTASGSSSAQGYIATRLGMLH-SGKG-KERSQKLWIEG
AtRrp40    -------------------------------------------MAAKVSVSPTSLLNQIVVPGDVVLDLSNM--------------------NNQTIKLGSGLLQDNDA----------------ISAMRAGKLSYSKPN------KYWLES
OsRrp40    --------------------------------------------MESRKPPPSALVDNHVVPGDVVLDLAEM--------------------TNQTIKLGAGLRQDCDT----------------IQATSAGRLRLSKPS------KYWVES
SsRrp4     -------------------------------------------MNMSQSQKIVLQPRSIVVPGELLAEGE-------------------------FQIPWSPYILKINSK----------------YYSTVVGLFDVKDTQ------FEVIPL
consensus                                                             pg  l                                g g                                G l

HsEXOSC2   KT-RYIGEVGDIVVGRITEVQQKRWKVETNSR-LDSVLLLSSMNLPGGELRRRSAEDELAMRGFLQEGDLISAEVQAVFSDGAVSL------HTRSLKYGKL-GQGVLVQ-VSPSLVK----RQKTHFHDLP-----CGASVILGNNGFIWIY
MmRrp4     KT-RYNGEVGDIVVGRITEVQQKRWKVETNSR-LDSVLLLSSMNLPGGELRRRSAEDELAMRGFLQEGDLISAEVQAVFSDGAVSL------HTRSLKYGKL-GQGVLVQ-VSPSLVK----RQKTHFHDLP-----CGASVILGNNGFIWIY
DrRrp4     KT-RYNGEVGDVVVGRITEVQQKRWKVETNSR-LDSVLLLSAVNLPGGELRRRSTQDELAMRDYLQEGDLISAEVQSIFSDGALSL------HTRSLKYGKL-GQGVLVY-VSPSLVK----RQKTHFHNLP-----CGASLILGNNGYVWLS
DmRrp4     KS-RYVGEIGDVVVARVSEVQQKRWRVDTNSR-LDSILLLSSVNLPGGELRRRSAEDEQMMRRYLDEGDLISAEVQNIFEEGSLSL------YTRSLKYGKL-SQGILVK-VFPALVK----RRKMHFHNLP-----CGASVILGNNGYIWIS
CeRrp4     KQ-RYAGEVGDVVVARVVEVQAKRWKCDVASR-LHANLPLGSVLLPGGDFRRKDVEDEEKMSEFLKNGELICAEVQQVQHDGTLML------HTRNNKYGKL-QQGILIK-VPPHLIK----KSKKHFHTLP-----YGMAVIIGCNGSVWVT
ScRrp4     KG-RYAPETGDHVVGRIAEVGNKRWKVDIGGK-QHAVLMLGSVNLPGGILRRKSESDELQMRSFLKEGDLLNAEVQSLFQDGSASL------HTRSLKYGKL-RNGMFCQ-VPSSLIV----RAKNHTHNLP-----GNITVVLGVNGYIWLR
SpRrp4     RS-KYVPEIGDLIIGKIAEVQPKRWKVDIGAK-QNAVLMLSSINLPGGIQRRKLETDELQMRSFFQEGDLLVAEVQQYFSDGSVSI------HTRSLKYGKL-RNGVFLK-VPPALVV----RSKSHAYALA-----GGVDIILSVNGYVWVS
CnRrp4     RS-RYTPEVGDLVIGRIVEVGAQRWRVDANGR-QDAVLMLSSVNLPGGVQRRKIESDALKMREFLAEGDLLVAEVQAFFGDGAMSL------HTRSLKYGKL-RNGFLLT-VPPQLIR----RLKSHFYHIPPPCGPTGVDVILGLNGYVWVS
AtRrp4     RA-RYKPEVGDIVVGRVIEVAQKRWRVELNFN-QDGVLMLSSMNMPDGIQRRRTSVDELNMRNIFVEHDVVCAEVRNFQHDGSLQL------QARSQKYGKL-EKGQLLK-VDPYLVK----RSKHHFHYVE----SLGIDLIIGCNGFIWVG
OsRrp4     RA-RYKPEVGDIIVGRVIEIAPKRWRLEINFS-QDAVLMLSSMNLPDGIQRRRTAVDELNMRTIFEENDVICAEVRGFQHDGSLHL------QARSQKYGKL-ERGQLLV-VPAYLVK----RRKQHFHHLE----QYDVDLILGCNGFIWVG
HsEXSOC3   QQKRYVPVKGDHVIGIVTAKSGDIFKVDVGGS-EPA--SLSYLSFEGATKRN---------RPNVQVGDLIYGQFVVANKDMEPEMVCIDS-CGRANGMGVIGQDGLLFK-VTLGLIRKLLA---PDCEIIQEVGKLYPLEIVFGMNGRIWVK
MmRrp40    QQKRYVPVKGDHVIGIVIAKSGDIFKVDVGGS-EPA--SLSYLAFEGATKRN---------RPNVQVGDLIYGQCVVANKDMEPEMVCIDS-CGRANGMGVIGQDGLLFK-VTLGLIRKLLA---PDCEIVQELGKLYPLEIVFGMNGRIWVK
DrRrp40    QQRRYVPAKGESVIGIVTAKSGDVFKVDVGGS-EQA--SLSYLAFEGATKRN---------RPNVQVGDLVFGQFTIANKDMEPELVCIDS-CGRANGMGVFGGDGLLFK-VSLGLVRRLLA---PQSDLVSDLEKMFPCEMVVGMNGRVWVK
DmRrp40    YQRRYIPARGDLILGIVRAKAGDLYRVDIGAT-DTA--SISYLAFEAASKKN---------RPDLIPGDLIYARVLNASADIEPELVCVNS-VGKSGKLGVL-TDGFFFK-CSLNLGRMLLR---ENCPVLAALTRELPYEIAVGVNGRIWLK
CeRrp40    HSKRYIPQEGDRVIAIVTSKTGDFFRLDIGTA-EYA--MINFTNFEGATKRN---------RPNLKTGDIIYATVFDTTPRTEAELTCVDD-EKRARGMGQL-NGGFMFK-VSLNHCRRLIN---PSCKILQTVGKFFKFEITVGMNGRIWIS
ScRrp40    SSKRYIPSVNDFVIGVIIGTFSDSYKVSLQNFSSSV--SLSYMAFPNASKKN---------RPTLQVGDLVYARVCTAEKELEAEIECFDSTTGRDAGFGIL-EDGMIID-VNLNFARQLLFN--NDFPLLKVLAAHTKFEVAIGLNGKIWVK
SpRrp40    RTKRYIPATNDQVIGQIISRFAEGYRVDIGSA-HIA--QLNALAFENVTRKS---------RPNLNVGSLVYARVSLADRDMEPELECFDATTGKAAGYGEL-KNGYMITGLSLSHCRKLILP--KN-TLLQTLGSYIPFEIAVGMNGRVWVN
CnRrp40    NSKRYIPAQKDIVLGIIIARHADGYRVDLGSS-QMA--QLDALAFEGATKRS---------KPNLKVGTLVFARVMSASRDMEPELECFDPNTGKSDGFGEL-KGGVMVN-ATLQLCRRLLDR--NH-IVLPTIASIFPFEIAIGLNGRVWMK
AtRrp40    SHKRYIPRPDDHVLGIVVDSRADNFWVDIKGP-QLA--LLPVLAFEGGTRRN---------IPKFEVGTLLYLRVVKTNIGMNPELSCTEA-SGKAALFGPL-RDGFMFE-TSTGLSRMLLNS--PTRPVIEALGKKLSFEIAVGLNGRVWVN
OsRrp40    SQKRYIPSVEDTVLGVVVDTKPDNFLVDIKGP-NLA--FLPVLAFEGGTRRN---------IPKFEIGTLIYARVVKANSIMNPELSCMDA-TGKAAEFGQL-KDGYMFD-TSTGLSRMLLSS--PTCPILEALGKKLSFEIAVGLNGRVWVN
SsRrp4     EGSFYYPKINDIVIGLVEDVEIYGWVVDIKAP-YKAYLPAS-------NLLGRSINVGEDLRRYLDVGDYVIARIENFDRSIDPVL------SVKGKDLGRV-SNGIVID-IMPVKVPRVIGKNKSMYETLT---SKSGCSIFVANNGRIWAT
consensus     rY     d vig v        f vd            l  l    a  r                 g li g            m        r    G i   G                            i          i vg NG iWv

HsEXOSC2   PTPEHKE--------------------------------EE-AGGFIANL-EPV-SLADREVISRLRNCIISLVTQRMMLYDTSILYCYEASLPHQIKDI------LKPEIME-EIVMETRQRL------------LEQEG--
MmRrp4     PTPEHKD--------------------------------ED-AGGFIANL-EPV-ALSDREVISRLRNCVVLLVTQRMMLFDTSILYCYEASLAHQIKDI------LKPEVME-EIMLETRQRL------------LDQEG--
DrRrp4     PTPALQE--------------------------------EQ-AGGFYTSM-EPV-SLSDREVVSRLRNCLLALANQKVLLFDTSVLYCYEASLQHQIKDI------LKPEIME-EIVLETRQRL------------MEHEG--
DmRrp4     PTKGQEE--------------------------------EGGEGGFAQNLNEHV-PRADREVIARLRNSILALAKCKLMIYDTSIQYAYEESLRYEAHEL------LQQNAIY-DIGQQTQARL------------RDADE--
CeRrp4     PSLPETTLE------------------------------EDGS---HVH-EFQIVPPDVRVTMVRIAACVRLLRDYSISIYLNSLTTCYEMSQPYEIKELAEQDTSSRLAYL---IAARLLQELQ-----------QQK----
ScRrp4     KTSQMDLARDTPSANNSSSIKSTG PTGAVSLNPSITRLEEESSWQIYSDENDPSISNNIRQAICRYANVIKALAFCEIGITQQRIVSAYEASMVY--SNVGEL---IEKNVM-ESIGSDILTAEK-----------MRGNGN-
SpRrp4     KHNENQH-----------------------SSVSITRLEEEASESIYSNENDEI-DGYTRLNISRVSICIKGLASRSLPLTQASITNFYESSLVF--SNLQDL---TVPKNM-DQIAMEAMQ---------------------
CnRrp4     NGTSQARRE---------------------------GGEGFDSEGVYSNQNDEI-PPQGREAISLVANIIKALASEGVPLTETLIGESYAWAEKNVPSGSGPFDSETERKMLNEIVGLEFLEFA-------------------
AtRrp4     EHVEVRDPMAIDD----------------------QKDEEMISSS-STGKEQSHIPLETRQTICRIGNAIRVLSNLGFTVTLEVIMETVNLSNSKNIDIH-------------DMLGSEFHVVVAENEAERRRTKRKK-----
OsRrp4     EHVVVGENANMME----------------------NKL------N-LSAEVENFTPLETRKHICRLANAVRVLSALGFTLTVELIIETAEASVSSNIEIN-------------NMLGAEFYVQTAEREVKRRADLLRKKSGAR
HsEXSOC3   AKTIQQTLILANILEACEHMTSDQRKQ----------------------------------IFSRLAES---------------------------------------------------------------------------
MmRrp40    AKTIQQTLILANVLEACEHMTTEQRKQ----------------------------------IFARLAES---------------------------------------------------------------------------
DrRrp40    ARTVQHTLILSNLLEACENMSTAQRHT----------------------------------LFRRVADGGI-------------------------------------------------------------------------
DmRrp40    AHSLKETVALANAISALEQSGCAEIDK----------------------------------ICGNLGDFLQA------------------------------------------------------------------------
CeRrp40    ASTSDDIIKIHDIINKSEFITDEDELIT----------------------------LVQNNYTRSVSD--------------------------------------------------------------------------
ScRrp40    CEELSNTLACYRTIMECCQKNDTAAFKD----------------------------IAKRQFKEILTVKEE-------------------------------------------------------------------------
SpRrp40    SENLSTTVLICTAIRNCEFMSDEEQIKY----------------------------------CKDLIKKL---------------------------------------------------------------------------
CnRrp40    AETVGETLAMKRLIESVDDHIIPVEEKA------------------------------IKEKLREYLS----------------------------------------------------------------------------
AtRrp40    ATAPRIVIIVANAIMNSESLSGTQQRI------------------------------MVEKLLAIISD---------------------------------------------------------------------------
OsRrp40    APSPSNVIVVSNAIIKSESLSGIGQRS------------------------------MVESLLERLS----------------------------------------------------------------------------
SsRrp4     CPSRFSEEILIEAI---------------------RKI-----------ENESH--------IKGLTDRIKQFIEEKLGERNASSGETKTNS------------------------------------------------------
consensus                                                                      l
```

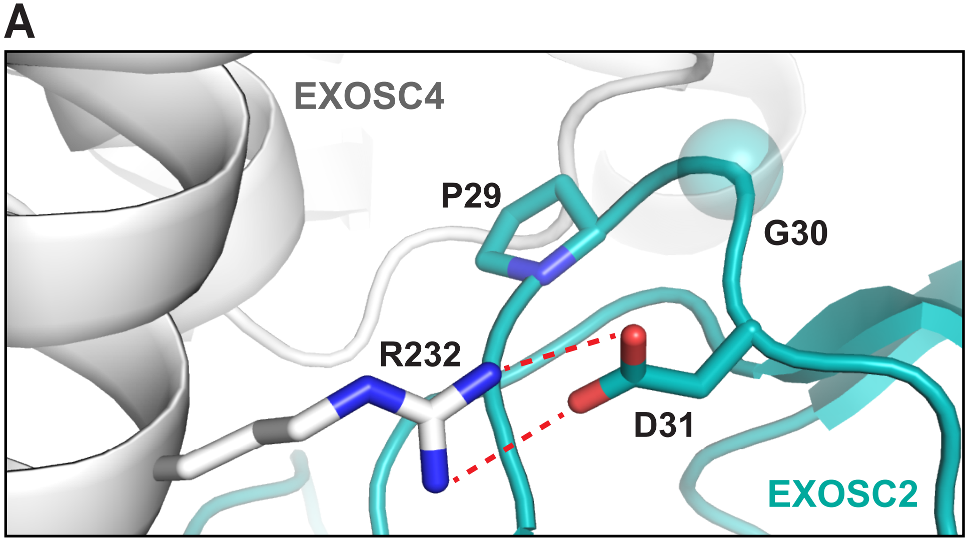
A
EXOSC4
P29
G30
R232
D31
EXOSC2

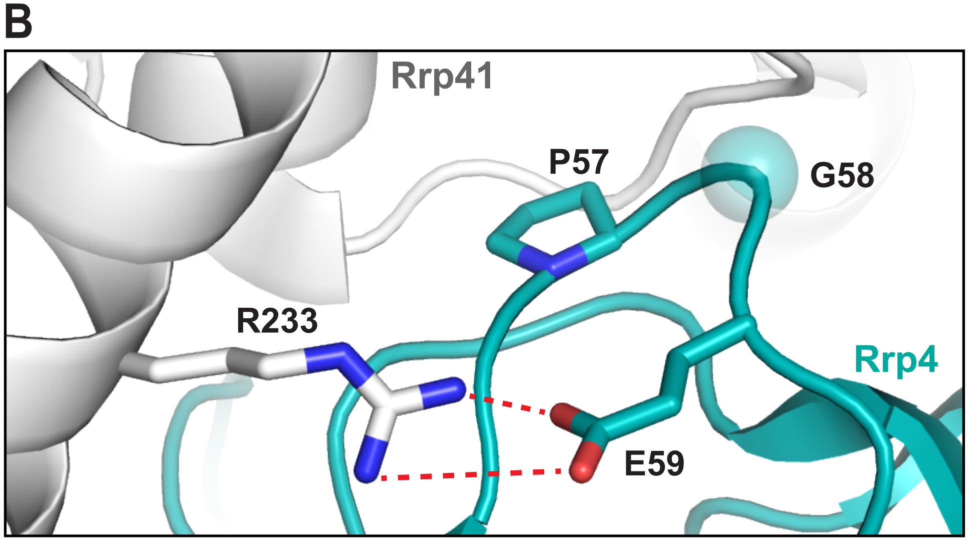
B
Rrp41
P57
G58
R233
Rrp4
E59

### Supplemental Figure S3 Sterrett Enyenihi *et al.*

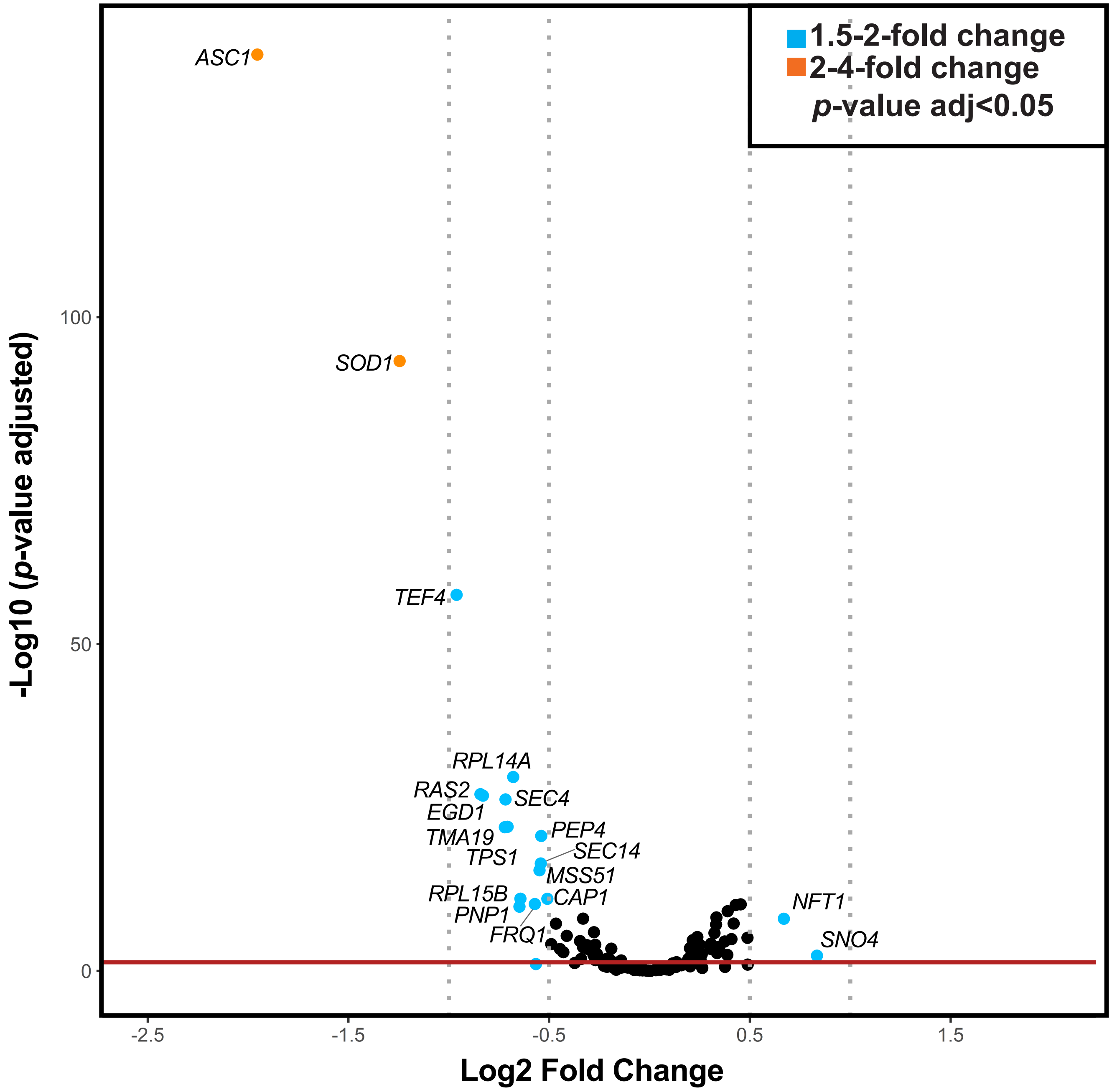

### Supplemental Table S1 — Sterrett Enyenihi *et al.*

| Strain/Plasmid | Description | Source |
|---|---|---|
| *rrp4Δ* (yAV1103) | MATa his3Δ1 leu2Δ0 ura3Δ0 lys2Δ0 rrp4Δ::NEO [RRP4,URA3] | (SCHAEFFER *et al.* 2009) |
| *rrp40Δ* (yAV1107) | MATa his3Δ1 leu2Δ0 ura3Δ0 rrp40Δ::NEO [RRP40,URA3] | (SCHAEFFER *et al.* 2009) |
| *rrp4Δ mpp6Δ* (ACY2471) | *MATα, ura3Δ0, leu2Δ0, his3Δ1, lys2Δ0, rrp4Δ::NEO, mpp6Δ::natMX4,* [*RRP4, URA3*] | This study |
| *rrp4Δ rrp47Δ* (ACY2474) | *MATα, ura3Δ0, leu2Δ0, his3Δ1, lys2Δ0, rrp4Δ::NEO, rrp47Δ::natMX4,* [*RRP4, URA3*] | This study |
| *rrp4Δ rrp6Δ* (ACY2478) | *MATα, ura3Δ0, leu2Δ0, his3Δ1, lys2Δ0, rrp4Δ::NEO, rrp6Δ::natMX4,* [*RRP4, URA3*] | This study |
| *rrp40Δ mpp6Δ* (ACY2638) | *MATa; ura3Δ0, leu2Δ0, his3Δ1, rrp40Δ::NEO, mpp6Δ::natMX4,* [*RRP40, URA3*] | This study |
| *rrp40Δ rrp47Δ* (ACY2462) | *MATa; ura3Δ0, leu2Δ0, his3Δ1, rrp40Δ::NEO, rrp47Δ::natMX4,* [*RRP40, URA3*] | This study |
| *rrp40Δ rrp6Δ* (ACY2466) | *MATa; ura3Δ0, leu2Δ0, his3Δ1, rrp40Δ::NEO, rrp6Δ::natMX4,* [*RRP40, URA3*] | This study |
| pRS315 | CEN6l LEU2, amp$^{R}$ | (SIKORSKI AND HIETER 1989) |
| pAC3161 | RRP40-2xMyc in pRS315, CEN6, LEU2, amp$^{R}$ | (FASKEN *et al.* 2017) |
| pAC3162 | rrp40-G8A-2xMyc in pRS315, CEN6, LEU2, amp$^{R}$ | (FASKEN *et al.* 2017) |
| pAC3259 | rrp40-W195R-2xMyc in pRS315, CEN6, LEU2, amp$^{R}$ | (FASKEN *et al.* 2017) |
| pAC3652 | RRP40-Native 3'UTR in pRS315, CEN6, LEU2, amp$^{R}$ | This study |
| pAC3655 | rrp40-W195R-Native 3'UTR in pRS315, CEN6, LEU2, amp$^{R}$ | This study |
| pAC3474 | RRP4-2xMyc in pRS315, CEN6, LEU2, amp$^{R}$ | This study |
| pAC3476 | rrp4-G58V-2xMyc in pRS315, CEN6, LEU2, amp$^{R}$ | This study |
| pAC3477 | rrp4-G226D-2xMyc in pRS315, CEN6, LEU2, amp$^{R}$ | This study |
| pAC3656 | RRP4-Native 3'UTR in pRS315, CEN6, LEU2, amp$^{R}$ | This study |
| pAC3659 | rrp4-G226D-Native 3'UTR in pRS315, CEN6, LEU2, amp$^{R}$ | This study |
| pAC3669 | RRP4-2xMyc-Native 3'UTR in pRS315, CEN6, LEU2, amp$^{R}$ | This study |
| pAC3672 | rrp4-G226D-2xMyc-Native 3'UTR in pRS315, CEN6, LEU2, amp$^{R}$ | This study |

# Supplemental Table S2      Sterrett Enyenihi *et al.*

| Description | Sequence (5'-3') | Name |
|---|---|---|
| *pre-U4 snRNA* Fwd | ATCCTTATGCACGGGAAATACG | AC5723 |
| *pre-U4 snRNA* Rev | AAAGAATGAATATCGGTAATG | AC5722 |
| *pre-U6 snRNA* Fwd | GAAGTATCGTGACTTGTACATTTG | AC7748 |
| *pre-U6 snRNA* Rev | TCTTCTATGTTCAACCGATAGC | AC7749 |
| *U14 snoRNA (snR128)* Fwd | GATCACGGTGATGAAAGACTGG | AC5397 |
| *U14 snoRNA (snR128)* Rev | CTACAGTATACGATCACTCAGACATCCTA | AC5398 |
| *snR44 snoRNA* Fwd | GCATTTCCACATGGGATTAAA | AC6272 |
| *snR44 snoRNA* Rev | ATGGTGTGATCGGGCAGTAT | AC6270 |
| *TLC1 ncRNA* Fwd | AAGGCAAGGGTGTCCTTTCT | AC6420 |
| *TLC1 ncRNA* Rev | TTCCGCTTGGAAAATAATGC | AC6421 |
| *RPS3 mRNA* Fwd | TCCAACCAAGACCGAAGTTATC | AC9226 |
| *RPS3 mRNA* Rev | GTACCTGGAGCGTACTTGAATC | AC9227 |
| *RPL15A mRNA* Fwd | CCAGACAAGGCTAGAAGATTGG | AC9309 |
| *RPL15A mRNA* Rev | CCGTAAGTAGCACCCTTTGG | AC9308 |
| *INO1 mRNA* Fwd | TTGGACTGCAAATACTGAGAGG | AC9303 |
| *INO1 mRNA* Rev | AAGATCGTGGAAGGAGCAATC | AC9302 |
| *PTH4 mRNA* Fwd | ACTGTGCTTGGATTCCTCAG | AC9248 |
| *PTH4 mRNA* Rev | CTATAGAATCGCTGCCCTTAGC | AC9249 |
| *HXK2 mRNA* Fwd | TACTGGTGTCAATGGTGCTTAC | AC9307 |
| *HXK2 mRNA* Rev | TTGGAGCAGATGGTGGAATG | AC9306 |
| *TDH1 mRNA* Fwd | GGTAGATACAAGGGTACTGTTTCC | AC9230 |
| *TDH1 mRNA* Rev | TGAGCGGTGTCCAATTCC | AC9232 |
| *CUT501* Fwd | GGTTCAACGTTGCAGGATCT | AC9254 |
| *CUT501* Rev | GCTAGCACCTGTTGCTGTAAT | AC9255 |
| *CUT770* Fwd | AAACAACCCGCTAGTGTGAC | AC9262 |
| *CUT770* Rev | AGAGCAACTCACTGCAAAGG | AC9263 |
| *CUT896* Fwd | ATCAGCAGGTGTCATGTTACAG | AC9256 |
| *CUT896* Rev | CCCAGAGGCAAAGATGTTAAGT | AC9257 |
| *NRD1 mRNA* Fwd | CAAGCAGAGGTCGAAACAAATC | AC9244 |
| *NRD1 mRNA* Rev | GCTGGATCTGTGGAAGTCAA | AC9245 |
| *NAB3 mRNA* Fwd | ACAGTTCGGTAGGCTCAGATAG | AC9246 |
| *NAB3 mRNA* Rev | GGCGAAGTTCGACCTCTTTATC | AC9247 |
| *ALG9 mRNA* Fwd | CACGGATAGTGGCTTTGGTGAACAATTAC | AC5067 |
| *ALG9 mRNA* Rev | TATGATTATCTGGCAGCAGGAAAGAACTTGGG | AC5068 |

| | *Hs* EXOSC2 | | *Sc* Rrp4 | |
|---|---|---|---|---|
| **Exosomopathy amino acid substitution** | G30V | G198D | G58V | G226D |
| **Missense 3D* (Phyre2)** | Neutral | Damaging | Neutral | Damaging |
| **PolyPhen-2** (HumDiv)** | Posibly Damaging (Score: 1.000) | Probably Damaging (Score: 1.000) | Probably Damaging (Score: 1.000) | Probably Damaging (Score: 1.000) |
| **Provean┼** | Deletrious (-7.938) | Deletrious (-6.35) | Deletrious (-8.981) | Deletrious (-6.517) |
| **SNAP2╪** | Effect (Score: 91) | Effect (Score: 94) | Effect (Score: 63 | Effect (Score: 92) |

***Missense 3D** part of tool suite **Phyre2** (**Protein H**omology/analog**Y Re**cognition Engine) v2.0; predicts structural changes introducted by an amino acid substitution through three-dimensional protein modeling.

****PolyPhen-2** (**Poly**morphism **Phen**otyping) v2 HumDiv trained model; Naïve Bayes posterior probability that mutation is damaging and qualitative classification as benign, possibly damaging, or probably damaging based on 5%/10% false positive rate (FPR) thresholds (FPR, the chance the mutation is classified as damaging when it is non-damaging).

┼**PROVEAN** (**Pro**tein **V**ariation **E**ffect **An**alyzer) v1.1; delta alignment scores equal to or below predefined threshold (-2.5), protein variant predicited to have "deleterious" effect on function.

╪**SNAP2** predicted score for functional effects of mutations; scores range from -100 strong neutral prediction to +100 strong effect prediction.

**Table S4.** Full RNA-Seq dataset.

| GeneName | GeneID | MeanNorm ( | log2 fold cha | SE of log2 fol | Wald Statisti | p-value |
|---|---|---|---|---|---|---|
| INO1 | YJL153C | 1935.65818 | -3.2048483 | 0.06816352 | -47.017064 | 0 |
| HXK2 | YGL253W | 5340.67745 | -1.6969303 | 0.05712811 | -29.703945 | 6.83E-194 |
| PSA1 | YDL055C | 10586.0216 | -1.2816199 | 0.04824151 | -26.566747 | 1.65E-155 |
| ASC1 | YMR116C | 7339.07856 | -1.9537701 | 0.07640558 | -25.571039 | 3.20E-144 |
| TSA1 | YML028W | 9515.48488 | -1.2901632 | 0.05061336 | -25.490563 | 2.51E-143 |
| SSC1 | YJR045C | 9123.05226 | -1.1370321 | 0.0474259 | -23.974918 | 5.08E-127 |
| TDH1 | YJL052W | 6737.93908 | -1.7904916 | 0.07600774 | -23.556702 | 1.07E-122 |
| PBI2 | YNL015W | 2950.84462 | -1.2532781 | 0.05483193 | -22.856722 | 1.25E-115 |
| SSA2 | YLL024C | 22760.1842 | -1.206077 | 0.05451542 | -22.123595 | 1.87E-108 |
| SSA1 | YAL005C | 31428.1277 | -1.1171251 | 0.05064013 | -22.060078 | 7.65E-108 |
| TUF1 | YOR187W | 3139.44249 | -1.132389 | 0.05233408 | -21.637697 | 7.94E-104 |
| RPS3 | YNL178W | 4045.47159 | -1.6739223 | 0.07774633 | -21.530562 | 8.06E-103 |
| GDH1 | YOR375C | 2194.39916 | -1.3601583 | 0.06333676 | -21.475023 | 2.67E-102 |
| TDH2 | YJR009C | 12642.8701 | -1.1669713 | 0.05442949 | -21.440056 | 5.65E-102 |
| SOD1 | YJR104C | 7200.72138 | -1.2448965 | 0.05963582 | -20.874978 | 9.04E-97 |
| GND1 | YHR183W | 4838.38733 | -1.2175525 | 0.05841607 | -20.842767 | 1.77E-96 |
| SEC53 | YFL045C | 2743.41074 | -1.2915176 | 0.06289754 | -20.533672 | 1.08E-93 |
| RPL15A | YLR029C | 6982.96388 | -1.4162457 | 0.06975403 | -20.303426 | 1.20E-91 |
| PGK1 | YCR012W | 22038.9432 | -1.4650081 | 0.07254387 | -20.194789 | 1.09E-90 |
| OPI3 | YJR073C | 2047.74909 | -1.5930327 | 0.08047386 | -19.795656 | 3.25E-87 |
| RPS31 | YLR167W | 14587.2667 | -1.4911109 | 0.07534647 | -19.790057 | 3.63E-87 |
| Xu_2009_CUTs67 | Xu_2009_CU | 858.118252 | 1.5342528 | 0.07838568 | 19.5731252 | 2.62E-85 |
| PDC1 | YLR044C | 83347.8231 | -0.9636194 | 0.04947481 | -19.476971 | 1.72E-84 |
| GPM1 | YKL152C | 23574.3926 | -1.219696 | 0.06296759 | -19.370221 | 1.38E-83 |
| KAR2 | YJL034W | 7878.68946 | -1.0278471 | 0.05354901 | -19.194514 | 4.11E-82 |
| TDH3 | YGR192C | 62146.6772 | -1.2137339 | 0.06370783 | -19.051566 | 6.38E-81 |
| HTB2 | YBL002W | 2288.66113 | -1.2336077 | 0.06495547 | -18.991591 | 2.00E-80 |
| ADH1 | YOL086C | 35417.1103 | -1.3897775 | 0.07326525 | -18.969122 | 3.07E-80 |
| PST2 | YDR032C | 2202.6492 | -1.3156879 | 0.06943347 | -18.948901 | 4.51E-80 |
| Xu_2009_SUTs39! | Xu_2009_SU | 644.965105 | 1.57888475 | 0.08402257 | 18.7911976 | 8.91E-79 |
| ITR1 | YDR497C | 10110.5914 | -0.8610989 | 0.046036 | -18.704902 | 4.52E-78 |
| SNR83 | snR83 | 22005.9 | -1.3330507 | 0.07166377 | -18.60146 | 3.13E-77 |
| FBA1 | YKL060C | 53396.0209 | -0.9674845 | 0.05207658 | -18.57811 | 4.83E-77 |
| APE1 | YKL103C | 1491.32174 | -1.4363876 | 0.07735974 | -18.567639 | 5.87E-77 |
| TOM5 | YPR133W-A | 1083.66051 | -1.435155 | 0.07796134 | -18.408548 | 1.12E-75 |
| CDC19 | YAL038W | 72340.2027 | -0.9838043 | 0.05443512 | -18.072969 | 5.20E-73 |
| HO | YDL227C | 590.145573 | -1.5675469 | 0.08726349 | -17.963376 | 3.77E-72 |
| SAM2 | YDR502C | 1494.89557 | -1.1815666 | 0.06585312 | -17.942454 | 5.50E-72 |
| RPL41B | YDL133C-A | 1440.22834 | -1.5107107 | 0.08431968 | -17.916466 | 8.77E-72 |
| SNR191 | snR191 | 2884.21452 | -1.4743932 | 0.08278506 | -17.809894 | 5.92E-71 |

| | | | | | | |
|---|---|---|---|---|---|---|
| RPL30 | YGL030W | 2482.69398 | -1.1211345 | 0.0629845 | -17.800165 | 7.05E-71 |
| BAT1 | YHR208W | 3872.16232 | -0.9754989 | 0.05561798 | -17.539271 | 7.18E-69 |
| HSP10 | YOR020C | 3380.7215 | -1.0052134 | 0.05737009 | -17.521559 | 9.81E-69 |
| RPL4B | YDR012W | 3016.01408 | -1.1418078 | 0.06587067 | -17.334085 | 2.60E-67 |
| ARG1 | YOL058W | 945.637718 | -1.2858195 | 0.07432488 | -17.299989 | 4.70E-67 |
| RNR4 | YGR180C | 7499.38304 | -0.7894159 | 0.04572188 | -17.265605 | 8.54E-67 |
| HSC82 | YMR186W | 35661.6012 | -0.8544828 | 0.04983813 | -17.14516 | 6.83E-66 |
| HSP60 | YLR259C | 11585.7422 | -0.9776621 | 0.0575347 | -16.992565 | 9.32E-65 |
| ALD6 | YPL061W | 9525.2983 | -0.8327932 | 0.04914217 | -16.946609 | 2.04E-64 |
| VMA2 | YBR127C | 6604.72363 | -0.9095794 | 0.05389275 | -16.877584 | 6.58E-64 |
| ENO2 | YHR174W | 30007.3853 | -0.9476258 | 0.05628538 | -16.83609 | 1.33E-63 |
| RPL41A | YDL184C | 1820.03171 | -1.3834106 | 0.08229629 | -16.810121 | 2.06E-63 |
| VMA16 | YHR026W | 2230.71491 | -1.0726624 | 0.06400844 | -16.758139 | 4.94E-63 |
| RPL4A | YBR031W | 6893.78181 | -1.1276486 | 0.06748703 | -16.709116 | 1.12E-62 |
| RPL11A | YPR102C | 3585.64424 | -0.9854321 | 0.05915671 | -16.657995 | 2.65E-62 |
| NUM1 | YDR150W | 3943.37022 | 0.87877119 | 0.05300572 | 16.5787996 | 9.92E-62 |
| RPL18A | YOL120C | 6658.77309 | -0.9709423 | 0.05869366 | -16.542542 | 1.81E-61 |
| RPS22A | YJL190C | 5401.54818 | -0.928634 | 0.05614618 | -16.539576 | 1.90E-61 |
| RPL40B | YKR094C | 4214.0406 | -1.0115714 | 0.06135957 | -16.485959 | 4.63E-61 |
| WTM1 | YOR230W | 4408.97852 | -0.8409289 | 0.05121222 | -16.420473 | 1.36E-60 |
| IPP1 | YBR011C | 12500.6559 | -0.78107 | 0.04764312 | -16.394182 | 2.10E-60 |
| HHF2 | YNL030W | 2922.17001 | -1.1038496 | 0.06733039 | -16.394524 | 2.09E-60 |
| TEF4 | YKL081W | 6010.59859 | -0.9611108 | 0.05864161 | -16.389571 | 2.27E-60 |
| RPL37A | YLR185W | 3959.11604 | -1.1989314 | 0.07339047 | -16.336337 | 5.44E-60 |
| TPI1 | YDR050C | 7812.78229 | -1.0528155 | 0.06459229 | -16.299398 | 9.97E-60 |
| DYN1 | YKR054C | 3624.88624 | 0.97057749 | 0.05957025 | 16.2929889 | 1.11E-59 |
| RPL22A | YLR061W | 3943.39155 | -1.1887131 | 0.0730147 | -16.280462 | 1.36E-59 |
| SAH1 | YER043C | 9081.65886 | -1.0663227 | 0.06577316 | -16.212126 | 4.14E-59 |
| RPP2A | YOL039W | 5038.72245 | -1.1429077 | 0.07079691 | -16.14347 | 1.26E-58 |
| SAR1 | YPL218W | 1289.22485 | -1.119367 | 0.06980784 | -16.034976 | 7.28E-58 |
| SNR71 | snR71 | 332.51104 | 1.6606144 | 0.10396333 | 15.9730777 | 1.97E-57 |
| TAL1 | YLR354C | 5054.43436 | -0.9235222 | 0.05784628 | -15.965111 | 2.24E-57 |
| FTR1 | YER145C | 1426.01052 | -1.2064533 | 0.07596558 | -15.881577 | 8.50E-57 |
| RPS6A | YPL090C | 5534.45606 | -0.9719034 | 0.06163829 | -15.767853 | 5.18E-56 |
| UGP1 | YKL035W | 3369.36978 | -1.1525595 | 0.07317915 | -15.749834 | 6.89E-56 |
| MET6 | YER091C | 11629.3284 | -0.7972661 | 0.0506322 | -15.746227 | 7.29E-56 |
| AYR1 | YIL124W | 608.9611 | -1.4370896 | 0.09154287 | -15.698542 | 1.55E-55 |
| ADK1 | YDR226W | 3453.9587 | -0.9606307 | 0.06134439 | -15.659634 | 2.86E-55 |
| ADE17 | YMR120C | 1517.25607 | -1.2589294 | 0.08056389 | -15.626472 | 4.81E-55 |
| DUG1 | YFR044C | 1824.92522 | -0.9960112 | 0.06400691 | -15.560994 | 1.34E-54 |
| BNA5 | YLR231C | 813.65003 | -1.1444969 | 0.07422253 | -15.419806 | 1.20E-53 |

| | | | | | | |
|---|---|---|---|---|---|---|
| TIS11 | YLR136C | 422.628651 | -1.5486021 | 0.10126995 | -15.291823 | 8.67E-53 |
| YBR238C | YBR238C | 2565.35428 | 1.01380537 | 0.0663321 | 15.2837823 | 9.81E-53 |
| HYP2 | YEL034W | 10220.3207 | -0.8013356 | 0.05271427 | -15.201494 | 3.46E-52 |
| COQ5 | YML110C | 1972.22225 | -0.901217 | 0.05934116 | -15.187048 | 4.31E-52 |
| PGI1 | YBR196C | 10384.2518 | -0.8189788 | 0.05408339 | -15.142889 | 8.44E-52 |
| HSP26 | YBR072W | 6164.15771 | -1.1238226 | 0.07426151 | -15.133311 | 9.77E-52 |
| NTF2 | YER009W | 1486.44397 | -1.0523549 | 0.06957671 | -15.125104 | 1.11E-51 |
| SPE3 | YPR069C | 1587.3484 | -1.0148921 | 0.06720537 | -15.101354 | 1.59E-51 |
| RPS14A | YCR031C | 7006.18541 | -0.8683362 | 0.05752273 | -15.095533 | 1.73E-51 |
| RPL3 | YOR063W | 15314.3956 | -0.7808551 | 0.05201687 | -15.011575 | 6.17E-51 |
| RPL6B | YLR448W | 5494.14836 | -0.7743602 | 0.05162807 | -14.998822 | 7.47E-51 |
| PTH4 | YOL114C | 277.755153 | 1.69834692 | 0.11360269 | 14.9498825 | 1.56E-50 |
| PMI40 | YER003C | 1298.58062 | -0.9563623 | 0.06418544 | -14.899987 | 3.30E-50 |
| RPS20 | YHL015W | 13621.4777 | -0.9548019 | 0.06418311 | -14.876218 | 4.70E-50 |
| ERG10 | YPL028W | 2403.44412 | -0.8651359 | 0.05818077 | -14.869791 | 5.18E-50 |
| RPS21B | YJL136C | 4075.22545 | -0.8026163 | 0.05398277 | -14.868008 | 5.32E-50 |
| SNT2 | YGL131C | 1414.35339 | 1.00420299 | 0.06761981 | 14.8507216 | 6.88E-50 |
| GLT1 | YDL171C | 7232.65545 | 0.76766975 | 0.05177652 | 14.8266002 | 9.86E-50 |
| SIM1 | YIL123W | 3227.99412 | -0.8255427 | 0.05578692 | -14.79814 | 1.51E-49 |
| REA1 | YLR106C | 8770.42843 | 0.96618791 | 0.06542918 | 14.7669265 | 2.39E-49 |
| RPL12B | YDR418W | 5939.82809 | -0.9898456 | 0.06711056 | -14.749476 | 3.10E-49 |
| CAR1 | YPL111W | 481.291115 | -1.4993243 | 0.10207243 | -14.688827 | 7.60E-49 |
| RPS11B | YBR048W | 3354.67589 | -0.7838084 | 0.05354961 | -14.637053 | 1.63E-48 |
| RPL37B | YDR500C | 3530.86507 | -0.9187959 | 0.06281568 | -14.626856 | 1.89E-48 |
| GPH1 | YPR160W | 3884.41262 | -0.9127204 | 0.0625963 | -14.58106 | 3.71E-48 |
| SNR19 | snR19 | 151403.531 | -1.016343 | 0.0697151 | -14.578521 | 3.85E-48 |
| RPS24A | YER074W | 3249.97888 | -0.9490596 | 0.06513019 | -14.571732 | 4.25E-48 |
| TEF1 | YPR080W | 24949.4479 | -1.0621071 | 0.07297908 | -14.553585 | 5.54E-48 |
| ARF1 | YDL192W | 2367.11739 | -0.9848229 | 0.0677107 | -14.544568 | 6.32E-48 |
| TEF2 | YBR118W | 19587.394 | -0.9317439 | 0.06427473 | -14.496271 | 1.28E-47 |
| RPL25 | YOL127W | 5730.21244 | -0.9712333 | 0.06706449 | -14.482081 | 1.57E-47 |
| AAT2 | YLR027C | 3280.71648 | -0.9848807 | 0.068164 | -14.448693 | 2.55E-47 |
| ADH3 | YMR083W | 4497.90816 | -0.7583657 | 0.05255613 | -14.429633 | 3.37E-47 |
| PDI1 | YCL043C | 12545.5406 | -0.7471726 | 0.05179224 | -14.426341 | 3.53E-47 |
| ILV3 | YJR016C | 4109.26988 | -0.7732793 | 0.05380005 | -14.373209 | 7.62E-47 |
| RPL40A | YIL148W | 6059.47475 | -0.9774382 | 0.06824481 | -14.322529 | 1.58E-46 |
| GLO1 | YML004C | 2115.77049 | -0.9791345 | 0.06869482 | -14.253396 | 4.27E-46 |
| TRA1 | YHR099W | 6262.85264 | 0.74277635 | 0.05214214 | 14.2452232 | 4.80E-46 |
| RPL24B | YGR148C | 5694.79184 | -0.8599466 | 0.06039695 | -14.238245 | 5.30E-46 |
| ACP1 | YKL192C | 1660.41882 | -0.9274216 | 0.06516868 | -14.231093 | 5.88E-46 |
| LEU1 | YGL009C | 5200.69993 | -0.8774052 | 0.06167012 | -14.227396 | 6.19E-46 |

| | | | | | | |
|---|---|---|---|---|---|---|
| SAC6 | YDR129C | 3893.68748 | -0.7374804 | 0.05209262 | -14.157099 | 1.69E-45 |
| Xu_2009_SUTs18( | Xu_2009_SU[illegible] | 528.470397 | 1.27712689 | 0.09022915 | 14.1542604 | 1.76E-45 |
| RPS26A | YGL189C | 9999.33136 | -0.8762817 | 0.0619538 | -14.144115 | 2.03E-45 |
| RPL1A | YPL220W | 2360.16589 | -0.9977787 | 0.07071231 | -14.110397 | 3.28E-45 |
| MBF1 | YOR298C-A | 6997.32612 | -0.9073544 | 0.06442266 | -14.084399 | 4.74E-45 |
| RPS16B | YDL083C | 3216.85316 | -0.8843495 | 0.06284244 | -14.072487 | 5.61E-45 |
| RPS15 | YOL040C | 11339.1799 | -0.8408772 | 0.05981434 | -14.058121 | 6.87E-45 |
| NRD1 | YNL251C | 3589.40992 | 0.900242 | 0.06413822 | 14.0359682 | 9.39E-45 |
| RPL23B | YER117W | 4823.7031 | -0.8430646 | 0.06015945 | -14.013834 | 1.28E-44 |
| RPS1A | YLR441C | 9086.50654 | -0.792808 | 0.05667819 | -13.987887 | 1.85E-44 |
| RPS5 | YJR123W | 10467.5102 | -0.891865 | 0.06382807 | -13.972929 | 2.28E-44 |
| EFB1 | YAL003W | 7399.62731 | -0.8663903 | 0.06217823 | -13.933983 | 3.94E-44 |
| SUN4 | YNL066W | 2943.92716 | -0.7971724 | 0.05737016 | -13.895245 | 6.77E-44 |
| ACS2 | YLR153C | 2253.91349 | -0.8716385 | 0.06273342 | -13.894324 | 6.86E-44 |
| RPS2 | YGL123W | 11169.3334 | -0.7909157 | 0.05711401 | -13.848015 | 1.31E-43 |
| HXT4 | YHR092C | 1497.22947 | -0.9350987 | 0.0678457 | -13.782726 | 3.24E-43 |
| ILV5 | YLR355C | 9223.43501 | -0.9077809 | 0.06605981 | -13.741804 | 5.70E-43 |
| RPL1B | YGL135W | 4998.21591 | -1.0399023 | 0.07579861 | -13.719279 | 7.78E-43 |
| SUB2 | YDL084W | 3420.87707 | -0.7786129 | 0.05686295 | -13.692797 | 1.12E-42 |
| NHP6A | YPR052C | 432.039104 | -1.4102996 | 0.10301043 | -13.690843 | 1.15E-42 |
| ECM33 | YBR078W | 6840.97752 | -0.8181242 | 0.05987725 | -13.663357 | 1.68E-42 |
| GAS3 | YMR215W | 884.227379 | -1.0415358 | 0.07631252 | -13.648295 | 2.07E-42 |
| RPL42B | YHR141C | 4336.76227 | -0.8669033 | 0.06354059 | -13.6433 | 2.21E-42 |
| STM1 | YLR150W | 7702.50428 | -0.9382287 | 0.0688908 | -13.619071 | 3.08E-42 |
| RPL17B | YJL177W | 4571.68333 | -0.7302376 | 0.05362374 | -13.617803 | 3.14E-42 |
| RPS9B | YBR189W | 7635.79913 | -0.8890329 | 0.06529099 | -13.616471 | 3.20E-42 |
| RPL8B | YLL045C | 10655.4401 | -0.7441924 | 0.05473961 | -13.595137 | 4.28E-42 |
| RPL2A | YFR031C-A | 5697.45793 | -0.8939518 | 0.06584094 | -13.577447 | 5.45E-42 |
| POR1 | YNL055C | 9169.66563 | -0.795014 | 0.05859613 | -13.567688 | 6.23E-42 |
| SHM2 | YLR058C | 4663.47111 | -0.8769617 | 0.06471115 | -13.551941 | 7.72E-42 |
| RPL16A | YIL133C | 3408.70846 | -0.8172854 | 0.06051362 | -13.505808 | 1.45E-41 |
| ADO1 | YJR105W | 2909.35532 | -0.7775504 | 0.05775318 | -13.463333 | 2.57E-41 |
| BNA1 | YJR025C | 1235.77439 | -0.9498 | 0.07076096 | -13.422656 | 4.45E-41 |
| RPL13B | YMR142C | 6371.10021 | -0.8304689 | 0.06198072 | -13.398825 | 6.14E-41 |
| RPL36B | YPL249C-A | 6324.1264 | -0.7664594 | 0.05747019 | -13.336644 | 1.42E-40 |
| FPR1 | YNL135C | 3397.0427 | -0.8394986 | 0.06300163 | -13.325029 | 1.66E-40 |
| RPS27B | YHR021C | 3354.41001 | -0.9197573 | 0.0691186 | -13.306945 | 2.11E-40 |
| RPL35B | YDL136W | 2768.10669 | -0.8326883 | 0.06263827 | -13.293602 | 2.52E-40 |
| RPS18A | YDR450W | 4321.50413 | -0.9474348 | 0.07133854 | -13.280827 | 2.99E-40 |
| RRP4 | YHR069C | 2038.47274 | 0.78493719 | 0.05942519 | 13.2088289 | 7.80E-40 |
| RPL43A | YPR043W | 5681.98207 | -0.8365117 | 0.06334106 | -13.206468 | 8.05E-40 |

| | | | | | | |
|---|---|---|---|---|---|---|
| BNA6 | YFR047C | 1148.24576 | -0.8938082 | 0.06768401 | -13.205604 | 8.14E-40 |
| RPL21B | YPL079W | 2929.89444 | -0.9026893 | 0.06838351 | -13.200395 | 8.73E-40 |
| EFT2 | YDR385W | 9712.47287 | -0.8085139 | 0.06133619 | -13.181678 | 1.12E-39 |
| SMT3 | YDR510W | 2951.03778 | -0.78222 | 0.05934444 | -13.181016 | 1.13E-39 |
| RPL6A | YML073C | 5331.01856 | -0.8411076 | 0.06383919 | -13.175413 | 1.22E-39 |
| LSP1 | YPL004C | 2823.8454 | -0.8878257 | 0.06743517 | -13.165618 | 1.38E-39 |
| VMA3 | YEL027W | 5268.7769 | -0.7186144 | 0.05462092 | -13.156396 | 1.56E-39 |
| RPL14B | YHL001W | 2241.55342 | -0.8312327 | 0.06322654 | -13.146894 | 1.77E-39 |
| RPS19B | YNL302C | 3115.26987 | -1.0564411 | 0.08036113 | -13.14617 | 1.79E-39 |
| PGM2 | YMR105C | 2093.75625 | -0.9826859 | 0.07480924 | -13.135889 | 2.05E-39 |
| RPL21A | YBR191W | 5897.85065 | -0.7401931 | 0.05636818 | -13.131398 | 2.18E-39 |
| LSR1 | LSR1 | 847856.996 | -0.9876674 | 0.07525548 | -13.124192 | 2.39E-39 |
| CPT1 | YNL130C | 1632.08114 | -0.7690203 | 0.05873069 | -13.094012 | 3.56E-39 |
| PFY1 | YOR122C | 2665.28018 | -0.9162926 | 0.06998217 | -13.093229 | 3.60E-39 |
| RPS6B | YBR181C | 7664.58669 | -0.7867238 | 0.06014328 | -13.080826 | 4.24E-39 |
| CDS1 | YBR029C | 1128.80737 | -0.9595129 | 0.07340847 | -13.070875 | 4.83E-39 |
| Xu_2009_CUTs50 | Xu_2009_CU | 217.944164 | 1.55450871 | 0.1190413 | 13.0585661 | 5.68E-39 |
| HOR7 | YMR251W-A | 4627.68035 | -0.8364208 | 0.06409027 | -13.050666 | 6.30E-39 |
| AHP1 | YLR109W | 10978.8948 | -0.7326133 | 0.05616066 | -13.044954 | 6.79E-39 |
| YDR524C-B | YDR524C-B | 5455.45766 | -0.8152072 | 0.06249649 | -13.044047 | 6.87E-39 |
| ALD4 | YOR374W | 3829.49666 | -0.7671836 | 0.05906424 | -12.988971 | 1.41E-38 |
| YMR315W | YMR315W | 2228.30173 | -0.7778334 | 0.05999045 | -12.965953 | 1.91E-38 |
| RPS7B | YNL096C | 3393.76397 | -0.6816518 | 0.05263213 | -12.951249 | 2.31E-38 |
| Xu_2009_SUTs22 | Xu_2009_SU | 1059.88182 | 0.90376029 | 0.06984676 | 12.9391864 | 2.70E-38 |
| TIF1 | YKR059W | 8648.36559 | -0.6679961 | 0.05166082 | -12.930421 | 3.03E-38 |
| PDR5 | YOR153W | 11850.0101 | 0.57322657 | 0.04441176 | 12.9070889 | 4.11E-38 |
| RPL33B | YOR234C | 2036.02587 | -0.9940888 | 0.07717442 | -12.881066 | 5.75E-38 |
| RPS0A | YGR214W | 7929.88576 | -0.7979275 | 0.06227319 | -12.813339 | 1.38E-37 |
| HRI1 | YLR301W | 944.321427 | -1.0205904 | 0.07976751 | -12.794562 | 1.76E-37 |
| RPL9A | YGL147C | 3437.3716 | -0.9404215 | 0.07360737 | -12.776187 | 2.23E-37 |
| SSB2 | YNL209W | 9519.92654 | -0.8913961 | 0.06976963 | -12.776277 | 2.22E-37 |
| THS1 | YIL078W | 5585.52314 | -0.6389099 | 0.05003445 | -12.769401 | 2.43E-37 |
| RGT1 | YKL038W | 1639.85758 | 0.75844827 | 0.05949811 | 12.7474346 | 3.22E-37 |
| YDL124W | YDL124W | 6294.48862 | -0.6996376 | 0.05490034 | -12.743775 | 3.38E-37 |
| IRA2 | YOL081W | 5716.17765 | 0.61544858 | 0.04841476 | 12.7120042 | 5.07E-37 |
| RPL31A | YDL075W | 5988.83729 | -0.6903049 | 0.05443689 | -12.68083 | 7.55E-37 |
| RPL7A | YGL076C | 7271.12525 | -0.9599413 | 0.07587462 | -12.651678 | 1.10E-36 |
| HHT1 | YBR010W | 1714.73735 | -0.9899552 | 0.07834023 | -12.636613 | 1.33E-36 |
| RPS17A | YML024W | 4104.14378 | -0.9569908 | 0.07583093 | -12.62006 | 1.64E-36 |
| RFS1 | YBR052C | 887.090137 | -1.0403483 | 0.08244055 | -12.619377 | 1.65E-36 |
| RPS0B | YLR048W | 9719.10219 | -0.703703 | 0.05583312 | -12.603683 | 2.02E-36 |

| | | | | | | |
|---|---|---|---|---|---|---|
| RPL32 | YBL092W | 5404.32672 | -0.8117801 | 0.06446968 | -12.591656 | 2.35E-36 |
| PRC1 | YMR297W | 5563.44344 | -0.732373 | 0.05817981 | -12.588095 | 2.46E-36 |
| CPR1 | YDR155C | 8175.69718 | -0.7313079 | 0.0581082 | -12.58528 | 2.54E-36 |
| TOM7 | YNL070W | 1157.5717 | -0.8647002 | 0.06895923 | -12.539297 | 4.55E-36 |
| RPP0 | YLR340W | 8248.34242 | -0.908192 | 0.07243442 | -12.538128 | 4.62E-36 |
| CWP2 | YKL096W-A | 25370.2375 | -0.8240831 | 0.06573253 | -12.536914 | 4.69E-36 |
| COF1 | YLL050C | 2068.12606 | -1.0440675 | 0.08336722 | -12.523717 | 5.54E-36 |
| RPS26B | YER131W | 5397.24247 | -0.8287566 | 0.06618623 | -12.521587 | 5.69E-36 |
| ADE2 | YOR128C | 965.765998 | -0.9239057 | 0.07379474 | -12.519939 | 5.81E-36 |
| RPS24B | YIL069C | 3459.14977 | -0.7999089 | 0.06398805 | -12.500909 | 7.38E-36 |
| HOM2 | YDR158W | 1770.07621 | -0.9418721 | 0.07538314 | -12.494466 | 8.00E-36 |
| RPL38 | YLR325C | 4801.32764 | -0.8355046 | 0.06689538 | -12.48972 | 8.50E-36 |
| GLK1 | YCL040W | 5188.0453 | -1.0236085 | 0.08201866 | -12.48019 | 9.58E-36 |
| TOS6 | YNL300W | 856.027702 | -0.9798187 | 0.07857664 | -12.469592 | 1.09E-35 |
| RPS8A | YBL072C | 3584.9181 | -0.9251079 | 0.07421925 | -12.464528 | 1.17E-35 |
| SOL3 | YHR163W | 1143.55808 | -0.8625773 | 0.06944393 | -12.421205 | 2.01E-35 |
| GUK1 | YDR454C | 1493.51209 | -0.8717595 | 0.07024816 | -12.409713 | 2.31E-35 |
| MNP1 | YGL068W | 753.175867 | -1.06054 | 0.08557933 | -12.392478 | 2.87E-35 |
| RPS17B | YDR447C | 2211.29615 | -0.8753451 | 0.07064207 | -12.391272 | 2.91E-35 |
| RPL17A | YKL180W | 5827.9625 | -0.81761 | 0.06606809 | -12.375262 | 3.56E-35 |
| RPS13 | YDR064W | 4182.50255 | -0.787685 | 0.06367887 | -12.369645 | 3.82E-35 |
| ERG26 | YGL001C | 970.981085 | -0.8962476 | 0.07247387 | -12.366493 | 3.97E-35 |
| CPS1 | YJL172W | 3811.87851 | -0.6848391 | 0.05540897 | -12.359715 | 4.32E-35 |
| EMI2 | YDR516C | 1351.65251 | -0.9519953 | 0.07704816 | -12.355847 | 4.53E-35 |
| RGI1 | YER067W | 270.19187 | -1.6587972 | 0.13449646 | -12.333389 | 5.99E-35 |
| ARC1 | YGL105W | 4004.03167 | -0.5994387 | 0.04863798 | -12.324499 | 6.69E-35 |
| DSE2 | YHR143W | 2871.87518 | -0.9194216 | 0.07466726 | -12.313584 | 7.65E-35 |
| TKL1 | YPR074C | 9843.35818 | -0.6150784 | 0.05013223 | -12.269121 | 1.33E-34 |
| RPL28 | YGL103W | 5661.08616 | -0.8364466 | 0.0682021 | -12.264236 | 1.41E-34 |
| HTA2 | YBL003C | 418.391484 | -1.2960388 | 0.10611357 | -12.213695 | 2.63E-34 |
| TOM1 | YDR457W | 6424.44591 | 0.67496869 | 0.05548192 | 12.1655602 | 4.74E-34 |
| SRV2 | YNL138W | 1933.58926 | -0.8283723 | 0.0681003 | -12.164003 | 4.83E-34 |
| MYO1 | YHR023W | 3017.96237 | 0.74857723 | 0.06158012 | 12.1561509 | 5.32E-34 |
| RPL10 | YLR075W | 8020.93948 | -0.6896611 | 0.05680335 | -12.141204 | 6.39E-34 |
| SOD2 | YHR008C | 1973.90244 | -0.7886577 | 0.06496963 | -12.138866 | 6.57E-34 |
| RPS4A | YJR145C | 6756.70076 | -0.718348 | 0.05929993 | -12.113808 | 8.93E-34 |
| YNK1 | YKL067W | 1546.08633 | -0.866021 | 0.07155767 | -12.102421 | 1.03E-33 |
| RPL16B | YNL069C | 8441.15646 | -0.7380613 | 0.06107168 | -12.085164 | 1.27E-33 |
| CHO1 | YER026C | 1442.14383 | -0.9256856 | 0.07667725 | -12.072493 | 1.48E-33 |
| HHF1 | YBR009C | 3099.80676 | -0.7299281 | 0.06047733 | -12.06945 | 1.53E-33 |
| GRS1 | YBR121C | 5034.14984 | -0.6786624 | 0.05624306 | -12.0666 | 1.59E-33 |

| | | | | | | |
|---|---|---|---|---|---|---|
| SNR17A | snR17a | 204973.957 | -1.7079392 | 0.1416259 | -12.059512 | 1.73E-33 |
| RPS4B | YHR203C | 8305.25961 | -0.8368749 | 0.06952473 | -12.037082 | 2.27E-33 |
| RPL5 | YPL131W | 15357.4531 | -0.7228027 | 0.06014427 | -12.017815 | 2.86E-33 |
| OLA1 | YBR025C | 4527.55803 | -0.6316512 | 0.05256149 | -12.017375 | 2.88E-33 |
| HYR1 | YIR037W | 1052.22403 | -0.928435 | 0.07736847 | -12.000172 | 3.55E-33 |
| RPL11B | YGR085C | 2980.42206 | -0.7965854 | 0.06640048 | -11.996681 | 3.70E-33 |
| RNR1 | YER070W | 3264.162 | -0.6338532 | 0.05285612 | -11.992048 | 3.91E-33 |
| SCS2 | YER120W | 2656.05342 | -0.7056158 | 0.05890063 | -11.979766 | 4.54E-33 |
| MRPS18 | YNL306W | 1160.82157 | -0.838445 | 0.07011689 | -11.957818 | 5.91E-33 |
| RIM1 | YCR028C-A | 859.975943 | -0.9264619 | 0.07777259 | -11.912448 | 1.02E-32 |
| RPL13A | YDL082W | 3863.16787 | -0.7981192 | 0.06718645 | -11.87917 | 1.52E-32 |
| RPS8B | YER102W | 3023.10115 | -0.801549 | 0.06747489 | -11.879219 | 1.52E-32 |
| YKL033W-A | YKL033W-A | 1299.98951 | -0.9039291 | 0.07622935 | -11.858019 | 1.96E-32 |
| OAC1 | YKL120W | 928.844968 | -0.8956347 | 0.07559171 | -11.848318 | 2.20E-32 |
| LAT1 | YNL071W | 2180.09409 | -0.6947538 | 0.05872395 | -11.830841 | 2.70E-32 |
| PMT2 | YAL023C | 2919.72802 | -0.6363607 | 0.05385834 | -11.815453 | 3.25E-32 |
| RPS7A | YOR096W | 2617.57317 | -0.7682953 | 0.06502757 | -11.814916 | 3.27E-32 |
| SNR44 | snR44 | 468.277793 | -1.1019625 | 0.09331941 | -11.808503 | 3.53E-32 |
| RPS1B | YML063W | 12048.9571 | -0.7704731 | 0.06529955 | -11.799057 | 3.95E-32 |
| FLX1 | YIL134W | 561.959972 | 0.96172586 | 0.08151317 | 11.7984103 | 3.98E-32 |
| RPS27A | YKL156W | 2519.65849 | -0.7908494 | 0.06714576 | -11.778099 | 5.06E-32 |
| RPL14A | YKL006W | 2659.47072 | -0.6792603 | 0.05779178 | -11.753578 | 6.77E-32 |
| TOM22 | YNL131W | 1433.99746 | -0.7883938 | 0.06720916 | -11.730452 | 8.90E-32 |
| MMF1 | YIL051C | 3247.66246 | -0.6976159 | 0.0594973 | -11.725169 | 9.47E-32 |
| YNR065C | YNR065C | 1322.44796 | 0.84712021 | 0.07231794 | 11.7138315 | 1.08E-31 |
| PDA1 | YER178W | 4502.24368 | -0.7125026 | 0.06088752 | -11.701947 | 1.25E-31 |
| ILV6 | YCL009C | 2092.71416 | -0.7804144 | 0.06669393 | -11.701431 | 1.25E-31 |
| ACT1 | YFL039C | 11021.1542 | -0.7455703 | 0.06383896 | -11.678923 | 1.63E-31 |
| GSP1 | YLR293C | 5307.63286 | -0.6657848 | 0.05701681 | -11.676992 | 1.67E-31 |
| SNR11 | snR11 | 13976.1231 | -1.8271622 | 0.1565773 | -11.669394 | 1.83E-31 |
| PMD1 | YER132C | 2626.517 | 0.68324097 | 0.05859645 | 11.6601078 | 2.04E-31 |
| SBP1 | YHL034C | 3551.45331 | -0.621335 | 0.053349 | -11.646611 | 2.39E-31 |
| SBH1 | YER087C-B | 1272.68666 | -0.8125313 | 0.07008181 | -11.59404 | 4.42E-31 |
| HAC1 | YFL031W | 18139.6757 | -0.8168828 | 0.07052624 | -11.582679 | 5.04E-31 |
| Xu_2009_CUTs27 | Xu_2009_CU | 580.651214 | -1.1329627 | 0.09784165 | -11.579555 | 5.23E-31 |
| Xu_2009_SUTs97 | Xu_2009_SU | 184.491378 | 1.57294334 | 0.13587221 | 11.5766378 | 5.41E-31 |
| ERG4 | YGL012W | 2600.78844 | -0.7013272 | 0.06075032 | -11.54442 | 7.88E-31 |
| ERP2 | YAL007C | 1262.47288 | -0.8158008 | 0.0708593 | -11.512967 | 1.14E-30 |
| GAS1 | YMR307W | 12141.8343 | -0.7001852 | 0.06085868 | -11.505101 | 1.24E-30 |
| TOM20 | YGR082W | 848.559201 | -0.8470321 | 0.07373359 | -11.487738 | 1.52E-30 |
| TRR1 | YDR353W | 3970.79442 | -0.5953505 | 0.05190594 | -11.469796 | 1.87E-30 |

| | | | | | | |
|---|---|---|---|---|---|---|
| TIM10 | YHR005C-A | 992.340549 | -0.8201837 | 0.07153379 | -11.465682 | 1.96E-30 |
| SNR30 | snR30 | 170105.748 | -1.5433384 | 0.13464427 | -11.46234 | 2.04E-30 |
| RPL20A | YMR242C | 5233.78024 | -0.7964689 | 0.0694996 | -11.46005 | 2.09E-30 |
| RPL36A | YMR194W | 1222.43382 | -1.0379103 | 0.09067829 | -11.446072 | 2.46E-30 |
| RPL19B | YBL027W | 8306.68959 | -0.7284507 | 0.06379573 | -11.418487 | 3.38E-30 |
| VMA5 | YKL080W | 1687.53153 | -0.7478259 | 0.06556028 | -11.406691 | 3.87E-30 |
| TUB2 | YFL037W | 3060.96838 | -0.6527412 | 0.05724607 | -11.402376 | 4.07E-30 |
| RPL35A | YDL191W | 3583.08197 | -0.8566536 | 0.07571045 | -11.314866 | 1.11E-29 |
| RPS18B | YML026C | 3676.69494 | -0.6661777 | 0.05897305 | -11.296306 | 1.37E-29 |
| ERP1 | YAR002C-A | 1942.25567 | -0.6985339 | 0.06190935 | -11.283173 | 1.59E-29 |
| ASN2 | YGR124W | 4164.43295 | -0.6499136 | 0.0576661 | -11.27029 | 1.84E-29 |
| Xu_2009_SUTs69( | Xu_2009_SU | 241.8132 | 1.30105746 | 0.11545322 | 11.2691306 | 1.86E-29 |
| Xu_2009_CUTs45 | Xu_2009_CU | 361.579241 | 1.14856485 | 0.10195851 | 11.2650221 | 1.95E-29 |
| RPL2B | YIL018W | 6341.30142 | -0.8066594 | 0.07183805 | -11.228859 | 2.94E-29 |
| SOP4 | YJL192C | 1401.93114 | -0.7641591 | 0.0680529 | -11.228897 | 2.94E-29 |
| RPL20B | YOR312C | 6159.28598 | -0.6986892 | 0.06227737 | -11.218991 | 3.29E-29 |
| RAS2 | YNL098C | 1243.87101 | -0.8424951 | 0.07509837 | -11.218554 | 3.31E-29 |
| ATP2 | YJR121W | 9161.50401 | -0.6109366 | 0.05452706 | -11.204284 | 3.88E-29 |
| CPR3 | YML078W | 672.552882 | -0.9091611 | 0.0812246 | -11.193174 | 4.40E-29 |
| ERG13 | YML126C | 2405.20844 | -0.7325391 | 0.06548689 | -11.186041 | 4.77E-29 |
| LEU2 | YCL018W | 17711.7459 | -0.5915422 | 0.05291728 | -11.178622 | 5.19E-29 |
| EGD1 | YPL037C | 2598.77539 | -0.8297994 | 0.07427654 | -11.171756 | 5.61E-29 |
| YHR069C-A | YHR069C-A | 211.283611 | 1.38093845 | 0.12361424 | 11.1713544 | 5.63E-29 |
| PCL1 | YNL289W | 2703.26072 | -0.6691457 | 0.05991074 | -11.169045 | 5.78E-29 |
| RPP1B | YDL130W | 3168.17487 | -0.834046 | 0.07477706 | -11.153768 | 6.86E-29 |
| THR4 | YCR053W | 2402.40708 | -0.6601252 | 0.05919403 | -11.151888 | 7.01E-29 |
| ARF2 | YDL137W | 2017.84697 | -0.698171 | 0.06263999 | -11.145771 | 7.51E-29 |
| BLM10 | YFL007W | 4446.23014 | 0.57414374 | 0.05153991 | 11.1397895 | 8.03E-29 |
| RPL33A | YPL143W | 7163.98563 | -0.5806042 | 0.05216975 | -11.129134 | 9.05E-29 |
| BPH1 | YCR032W | 1681.40942 | 0.7306865 | 0.06565707 | 11.1288322 | 9.08E-29 |
| SNR43 | snR43 | 651.180219 | -1.0831942 | 0.09756354 | -11.102449 | 1.22E-28 |
| RPL24A | YGL031C | 4201.04376 | -0.7000358 | 0.06309273 | -11.095348 | 1.32E-28 |
| PET9 | YBL030C | 2267.35376 | -0.7768361 | 0.07002316 | -11.093987 | 1.34E-28 |
| RPL34A | YER056C-A | 7875.35246 | -0.6282576 | 0.05673015 | -11.074493 | 1.67E-28 |
| PNC1 | YGL037C | 6908.54907 | -0.809882 | 0.07313605 | -11.073635 | 1.68E-28 |
| HTA1 | YDR225W | 2718.97183 | -0.8652945 | 0.07818184 | -11.067717 | 1.80E-28 |
| YKR075C | YKR075C | 823.940668 | -1.0995055 | 0.09934285 | -11.067787 | 1.80E-28 |
| ASP1 | YDR321W | 844.938651 | -0.8520276 | 0.07702371 | -11.061886 | 1.92E-28 |
| SEC4 | YFL005W | 2475.98268 | -0.7176701 | 0.06496301 | -11.047365 | 2.26E-28 |
| RPL22B | YFL034C-A | 542.520723 | -0.9605815 | 0.08700069 | -11.04108 | 2.42E-28 |
| IFH1 | YLR223C | 3912.6933 | 0.71800839 | 0.06511753 | 11.0263453 | 2.85E-28 |

| | | | | | | |
|---|---|---|---|---|---|---|
| RPP1A | YDL081C | 3720.27543 | -0.7278478 | 0.06601798 | -11.024993 | 2.90E-28 |
| GPD1 | YDL022W | 3960.93474 | -0.7264398 | 0.0659184 | -11.020288 | 3.05E-28 |
| SNR84 | snR84 | 6001.88304 | -1.6624752 | 0.15097196 | -11.011814 | 3.35E-28 |
| RPS29A | YLR388W | 1734.23424 | -0.8448003 | 0.07671728 | -11.011865 | 3.35E-28 |
| XPT1 | YJR133W | 485.781068 | -1.0285009 | 0.09350226 | -10.999743 | 3.83E-28 |
| SSZ1 | YHR064C | 4160.27806 | -0.6872378 | 0.06248807 | -10.997904 | 3.91E-28 |
| SEC61 | YLR378C | 3163.72918 | -0.5989957 | 0.05454049 | -10.982589 | 4.63E-28 |
| WBP1 | YEL002C | 1636.83877 | -0.7428639 | 0.06766312 | -10.97886 | 4.83E-28 |
| IRA1 | YBR140C | 4525.22177 | 0.59769187 | 0.05449466 | 10.9678974 | 5.45E-28 |
| IMD3 | YLR432W | 3683.00874 | -0.6796221 | 0.06209588 | -10.94472 | 7.04E-28 |
| CTS1 | YLR286C | 9859.27353 | -0.6072385 | 0.05548894 | -10.943416 | 7.15E-28 |
| Xu_2009_CUTs40 | Xu_2009_CU | 203.787059 | 1.43027611 | 0.13084135 | 10.9313768 | 8.16E-28 |
| NAB3 | YPL190C | 4343.21437 | 0.67610155 | 0.06196855 | 10.9103973 | 1.03E-27 |
| Xu_2009_CUTs48 | Xu_2009_CU | 443.060508 | 1.0941493 | 0.10030856 | 10.9078361 | 1.06E-27 |
| YML131W | YML131W | 602.759966 | -0.958879 | 0.08810113 | -10.883845 | 1.38E-27 |
| HOT13 | YKL084W | 202.146103 | -1.3983873 | 0.12849394 | -10.882905 | 1.39E-27 |
| CCW12 | YLR110C | 38059.3825 | -0.7500463 | 0.06895774 | -10.876898 | 1.49E-27 |
| SMC4 | YLR086W | 1980.22659 | 0.69927771 | 0.06430611 | 10.8742039 | 1.53E-27 |
| BDH1 | YAL060W | 1143.7836 | -0.8021928 | 0.07379757 | -10.87018 | 1.60E-27 |
| SEN1 | YLR430W | 2745.0281 | 0.5849373 | 0.05398306 | 10.8355715 | 2.34E-27 |
| ERV25 | YML012W | 2340.35103 | -0.7141464 | 0.06600763 | -10.819148 | 2.79E-27 |
| EGD2 | YHR193C | 3738.02364 | -0.6482822 | 0.05998151 | -10.808033 | 3.15E-27 |
| MOG1 | YJR074W | 1009.56133 | -0.9328337 | 0.08631638 | -10.807146 | 3.18E-27 |
| BCY1 | YIL033C | 2072.25949 | -0.6167516 | 0.05720405 | -10.781608 | 4.20E-27 |
| RPS19A | YOL121C | 1911.95316 | -0.7463274 | 0.06928207 | -10.772303 | 4.65E-27 |
| YEL077C | YEL077C | 439.064368 | 1.04300803 | 0.09682831 | 10.7717263 | 4.68E-27 |
| SSB1 | YDL229W | 11556.1808 | -0.7630233 | 0.07086242 | -10.767672 | 4.89E-27 |
| EGO4 | YNR034W-A | 651.332389 | -1.0926346 | 0.10163056 | -10.751044 | 5.86E-27 |
| RPL26A | YLR344W | 5572.76487 | -0.5744715 | 0.05343585 | -10.750676 | 5.88E-27 |
| ERC1 | YHR032W | 1643.91637 | 0.70867492 | 0.06596383 | 10.7433869 | 6.37E-27 |
| SNR63 | snR63 | 39132.3898 | -1.4772516 | 0.13760268 | -10.73563 | 6.92E-27 |
| Xu_2009_SUTs75 | Xu_2009_SU | 684.475635 | 0.89991969 | 0.08383337 | 10.7346232 | 7.00E-27 |
| RDL1 | YOR285W | 1275.1646 | -0.8325014 | 0.07755633 | -10.734151 | 7.04E-27 |
| SNR34 | snR34 | 1954.42853 | -0.9253463 | 0.08626192 | -10.727171 | 7.59E-27 |
| RIB4 | YOL143C | 1230.53411 | -0.7388957 | 0.0690359 | -10.703065 | 9.85E-27 |
| RPS10B | YMR230W | 3684.59427 | -0.6603497 | 0.06170709 | -10.701358 | 1.00E-26 |
| YML079W | YML079W | 474.00522 | -0.9405266 | 0.08797203 | -10.691201 | 1.12E-26 |
| RPS11A | YDR025W | 4150.63376 | -0.7286898 | 0.06836322 | -10.659091 | 1.58E-26 |
| PRB1 | YEL060C | 6878.56585 | -0.5263637 | 0.0493879 | -10.657746 | 1.60E-26 |
| ENO1 | YGR254W | 10643.5621 | -0.7083458 | 0.06649456 | -10.652688 | 1.69E-26 |
| RPO26 | YPR187W | 1292.40938 | -0.7695749 | 0.07228209 | -10.646827 | 1.80E-26 |

| | | | | | | |
|---|---|---|---|---|---|---|
| EFT1 | YOR133W | 5898.16431 | -0.7178874 | 0.06743653 | -10.645378 | 1.83E-26 |
| CMD1 | YBR109C | 1725.68919 | -0.628661 | 0.05909239 | -10.638612 | 1.97E-26 |
| UBR2 | YLR024C | 910.999288 | 0.7489886 | 0.07049844 | 10.6241862 | 2.30E-26 |
| YIP1 | YGR172C | 506.883856 | -0.9144415 | 0.08612436 | -10.617687 | 2.47E-26 |
| MDH1 | YKL085W | 1157.94444 | -0.828145 | 0.07801091 | -10.615758 | 2.52E-26 |
| YDL086W | YDL086W | 1011.43142 | -0.8328697 | 0.07880336 | -10.568961 | 4.15E-26 |
| Xu_2009_CUTs89 | Xu_2009_CU | 118.690695 | 1.68246082 | 0.15931862 | 10.5603527 | 4.55E-26 |
| SGT2 | YOR007C | 4420.31498 | -0.5173122 | 0.04898646 | -10.560311 | 4.55E-26 |
| NCW1 | YMR122W-A | 2943.12823 | -0.7415966 | 0.07032117 | -10.545852 | 5.31E-26 |
| RPL34B | YIL052C | 2994.38416 | -0.7010662 | 0.066498 | -10.542666 | 5.49E-26 |
| YLR256W-A | YLR256W-A | 2672.93517 | 0.56914008 | 0.05399143 | 10.5413035 | 5.57E-26 |
| RPL12A | YEL054C | 4270.57627 | -0.8021991 | 0.07617356 | -10.5312 | 6.20E-26 |
| CHO2 | YGR157W | 3255.39695 | -0.5803823 | 0.05514071 | -10.525478 | 6.59E-26 |
| Xu_2009_SUTs51 | Xu_2009_SU | 340.935603 | 1.03569763 | 0.09841706 | 10.5235574 | 6.73E-26 |
| BRN1 | YBL097W | 479.655729 | 1.01362407 | 0.09633281 | 10.5221068 | 6.83E-26 |
| RPL19A | YBR084C-A | 6569.08048 | -0.7086083 | 0.06734798 | -10.521597 | 6.87E-26 |
| CAR2 | YLR438W | 462.6274 | -0.9680717 | 0.09208331 | -10.512998 | 7.53E-26 |
| RPS25A | YGR027C | 7581.24375 | -0.5822848 | 0.05541442 | -10.507821 | 7.95E-26 |
| RBD2 | YPL246C | 730.758188 | -0.8201667 | 0.07806308 | -10.506461 | 8.07E-26 |
| YPL222C-A | YPL222C-A | 200.74682 | 1.38484801 | 0.13193809 | 10.4961954 | 8.99E-26 |
| YDC1 | YPL087W | 673.967819 | -0.934007 | 0.08900991 | -10.493292 | 9.27E-26 |
| POL2 | YNL262W | 1726.66756 | 0.6283381 | 0.05988141 | 10.493041 | 9.30E-26 |
| RER1 | YCL001W | 737.265466 | -0.8275871 | 0.07889292 | -10.490005 | 9.60E-26 |
| ARO2 | YGL148W | 3138.65571 | -0.6463056 | 0.06161588 | -10.48927 | 9.68E-26 |
| ABP1 | YCR088W | 2569.62294 | -0.6029738 | 0.05751163 | -10.484381 | 1.02E-25 |
| YLR227W-B | YLR227W-B | 2545.25639 | 0.5948079 | 0.0567732 | 10.4769139 | 1.10E-25 |
| TIF2 | YJL138C | 5954.075 | -0.584192 | 0.05578698 | -10.471835 | 1.16E-25 |
| Xu_2009_CUTs77 | Xu_2009_CU | 187.306928 | 1.41687633 | 0.13531823 | 10.470698 | 1.18E-25 |
| RPS12 | YOR369C | 3706.24578 | -0.8021167 | 0.07665091 | -10.464542 | 1.26E-25 |
| YNL054W-B | YNL054W-B | 2267.69387 | 0.65533374 | 0.0626393 | 10.4620216 | 1.29E-25 |
| RPS29B | YDL061C | 1314.70408 | -0.8693692 | 0.08334066 | -10.431513 | 1.78E-25 |
| RPS23B | YPR132W | 4075.86079 | -0.7089004 | 0.06797927 | -10.428186 | 1.84E-25 |
| RPL26B | YGR034W | 3682.87642 | -0.695766 | 0.06672533 | -10.427315 | 1.86E-25 |
| RPL23A | YBL087C | 2675.03514 | -0.7555477 | 0.07271025 | -10.391213 | 2.72E-25 |
| TOA2 | YKL058W | 514.053385 | -0.9258563 | 0.08914123 | -10.386398 | 2.86E-25 |
| RPS16A | YMR143W | 6318.82258 | -0.7345973 | 0.07074986 | -10.38302 | 2.96E-25 |
| COX2 | Q0250 | 770.681109 | -0.9522636 | 0.09180265 | -10.372942 | 3.29E-25 |
| Xu_2009_CUTs84 | Xu_2009_CU | 206.118704 | 1.3242029 | 0.12766894 | 10.3721614 | 3.32E-25 |
| FLO5 | YHR211W | 505.666953 | 0.9632776 | 0.09298102 | 10.3599379 | 3.77E-25 |
| MRPL20 | YKR085C | 496.752811 | -0.971552 | 0.09390128 | -10.346526 | 4.34E-25 |
| RPC40 | YPR110C | 2348.05748 | -0.7231282 | 0.06994798 | -10.338085 | 4.74E-25 |

| | | | | | | |
|---|---|---|---|---|---|---|
| AIP1 | YMR092C | 1868.68224 | -0.6320121 | 0.06120999 | -10.325311 | 5.41E-25 |
| PUP2 | YGR253C | 1567.35333 | -0.6959532 | 0.06742617 | -10.321706 | 5.62E-25 |
| ARO4 | YBR249C | 1673.74642 | -0.7077532 | 0.06862 | -10.314096 | 6.09E-25 |
| RPL39 | YJL189W | 1962.7427 | -0.8392155 | 0.0815159 | -10.295115 | 7.41E-25 |
| Xu_2009_SUTs67! | Xu_2009_SU | 423.93156 | 0.95982142 | 0.09324964 | 10.2930311 | 7.58E-25 |
| SFM1 | YOR021C | 1092.89839 | -0.7701486 | 0.07485627 | -10.288365 | 7.95E-25 |
| PRX1 | YBL064C | 861.992192 | -0.7806144 | 0.07603553 | -10.266443 | 9.98E-25 |
| Xu_2009_CUTs77 | Xu_2009_CU | 239.78874 | 1.30087178 | 0.12674763 | 10.2634798 | 1.03E-24 |
| RPL43B | YJR094W-A | 1604.20325 | -0.8798562 | 0.08581155 | -10.253353 | 1.14E-24 |
| RPS25B | YLR333C | 1189.032 | -0.9143926 | 0.08920355 | -10.250631 | 1.18E-24 |
| RPS22B | YLR367W | 1730.34878 | -0.6405604 | 0.06249056 | -10.250514 | 1.18E-24 |
| RPL27B | YDR471W | 2903.61315 | -0.741571 | 0.07235457 | -10.249124 | 1.19E-24 |
| Xu_2009_SUTs39 | Xu_2009_SU | 221.017553 | 1.35690447 | 0.13251401 | 10.2397059 | 1.32E-24 |
| CDC16 | YKL022C | 897.913125 | 0.71055642 | 0.06944482 | 10.2319577 | 1.43E-24 |
| UTR2 | YEL040W | 1009.15921 | -0.7711762 | 0.07539236 | -10.228837 | 1.47E-24 |
| Xu_2009_CUTs28 | Xu_2009_CU | 198.202503 | 1.28955925 | 0.12615281 | 10.2222002 | 1.58E-24 |
| UTP20 | YBL004W | 5006.0223 | 0.61880865 | 0.06055686 | 10.2186377 | 1.64E-24 |
| VPS13 | YLL040C | 4350.48742 | 0.53332045 | 0.05226763 | 10.2036468 | 1.91E-24 |
| YBL005W-B | YBL005W-B | 2512.80726 | 0.60398951 | 0.05920715 | 10.2012943 | 1.96E-24 |
| YPT1 | YFL038C | 4307.03282 | -0.5267531 | 0.05167189 | -10.19419 | 2.10E-24 |
| EUG1 | YDR518W | 830.276029 | -0.7654796 | 0.07510182 | -10.192557 | 2.14E-24 |
| PRO2 | YOR323C | 1565.28231 | -0.666777 | 0.06543075 | -10.190575 | 2.18E-24 |
| VMA11 | YPL234C | 653.378328 | -0.9797046 | 0.09614165 | -10.19022 | 2.19E-24 |
| TIF11 | YMR260C | 2739.70846 | -0.7054914 | 0.06926754 | -10.185021 | 2.31E-24 |
| THO1 | YER063W | 658.376709 | -0.8415425 | 0.08264284 | -10.182885 | 2.36E-24 |
| MRP49 | YKL167C | 309.882414 | -1.0644881 | 0.1045599 | -10.180653 | 2.42E-24 |
| FLO1 | YAR050W | 292.389379 | 1.13786369 | 0.11194579 | 10.1644166 | 2.86E-24 |
| RTC3 | YHR087W | 602.666608 | -1.0498046 | 0.10334103 | -10.158643 | 3.03E-24 |
| BNA2 | YJR078W | 625.777797 | -0.8696061 | 0.08570274 | -10.146772 | 3.43E-24 |
| VHT1 | YGR065C | 1242.88585 | -0.6804682 | 0.06714272 | -10.134653 | 3.88E-24 |
| IMD4 | YML056C | 2709.48828 | -0.8089349 | 0.07982691 | -10.133612 | 3.92E-24 |
| MIX17 | YMR002W | 2055.33982 | -0.5794405 | 0.05719491 | -10.130981 | 4.03E-24 |
| PYP1 | YNL010W | 1006.91961 | -0.8912518 | 0.08809389 | -10.117067 | 4.64E-24 |
| TPS1 | YBR126C | 5361.59202 | -0.7074951 | 0.06993554 | -10.116389 | 4.67E-24 |
| LYS9 | YNR050C | 714.237502 | -0.8181154 | 0.08099026 | -10.101405 | 5.45E-24 |
| TMA19 | YKL056C | 4287.78271 | -0.7208853 | 0.07138536 | -10.098504 | 5.61E-24 |
| HOM6 | YJR139C | 2104.98155 | -0.6361476 | 0.0630469 | -10.090069 | 6.11E-24 |
| ALG7 | YBR243C | 436.532018 | -0.9079361 | 0.09009054 | -10.078041 | 6.91E-24 |
| FRS2 | YFL022C | 2161.85107 | -0.6536182 | 0.06488792 | -10.073033 | 7.27E-24 |
| IMG1 | YCR046C | 359.992175 | -1.0583032 | 0.10528897 | -10.051416 | 9.06E-24 |
| RPL27A | YHR010W | 5433.00941 | -0.6458537 | 0.06428144 | -10.047282 | 9.44E-24 |

| | | | | | | |
|---|---|---|---|---|---|---|
| NSG1 | YHR133C | 549.565065 | -0.8589107 | 0.08552377 | -10.042947 | 9.87E-24 |
| PIL1 | YGR086C | 4764.90709 | -0.5587795 | 0.05564958 | -10.041038 | 1.01E-23 |
| YTP1 | YNL237W | 175.487394 | -1.3820077 | 0.13776611 | -10.03155 | 1.11E-23 |
| GVP36 | YIL041W | 2519.66339 | -0.5606527 | 0.0560079 | -10.010242 | 1.37E-23 |
| TSC13 | YDL015C | 1766.48005 | -0.72789 | 0.07279208 | -9.9995778 | 1.53E-23 |
| SCP1 | YOR367W | 454.072931 | -0.8814931 | 0.08822106 | -9.9918663 | 1.65E-23 |
| IDP1 | YDL066W | 2163.2439 | -0.5830422 | 0.05838319 | -9.986474 | 1.75E-23 |
| RPS21A | YKR057W | 874.138927 | -0.9263036 | 0.09277766 | -9.9841241 | 1.79E-23 |
| PIR1 | YKL164C | 6643.48576 | -0.5611335 | 0.05624819 | -9.9760274 | 1.94E-23 |
| THO2 | YNL139C | 1801.79152 | 0.59838761 | 0.05999031 | 9.97473802 | 1.97E-23 |
| UTP14 | YML093W | 2642.51842 | 0.63756563 | 0.06400591 | 9.96104303 | 2.26E-23 |
| PRO3 | YER023W | 1629.42394 | -0.660985 | 0.06636176 | -9.9603287 | 2.27E-23 |
| MET13 | YGL125W | 2302.38173 | -0.5832617 | 0.05856703 | -9.958873 | 2.31E-23 |
| HMO1 | YDR174W | 1731.8717 | -0.6573135 | 0.06611591 | -9.9418344 | 2.74E-23 |
| YMR045C | YMR045C | 1533.34919 | 0.72873767 | 0.073761 | 9.87971469 | 5.10E-23 |
| YCP4 | YCR004C | 2285.02496 | -0.5844656 | 0.05932039 | -9.8526939 | 6.67E-23 |
| BRR2 | YER172C | 2397.30027 | 0.60139234 | 0.06108571 | 9.84505741 | 7.20E-23 |
| HFA1 | YMR207C | 1061.52935 | 0.7450709 | 0.07568182 | 9.84478093 | 7.22E-23 |
| YDR341C | YDR341C | 2866.8988 | -0.5873475 | 0.05980452 | -9.8211216 | 9.13E-23 |
| CDC33 | YOL139C | 2774.88267 | -0.5736124 | 0.05850784 | -9.8040258 | 1.08E-22 |
| RRP14 | YKL082C | 924.153564 | 1.00894263 | 0.10303455 | 9.79227484 | 1.22E-22 |
| PEP4 | YPL154C | 7952.21062 | -0.5396533 | 0.05512837 | -9.7890323 | 1.25E-22 |
| GET4 | YOR164C | 1107.99308 | -0.7753973 | 0.07965423 | -9.7345401 | 2.15E-22 |
| YBL113C | YBL113C | 385.386448 | 0.9449885 | 0.09741185 | 9.70096079 | 2.99E-22 |
| RPL29 | YFR032C-A | 1122.18702 | -0.9452865 | 0.09744031 | -9.7011856 | 2.98E-22 |
| CSF1 | YLR087C | 2022.50907 | 0.57301844 | 0.05906721 | 9.70112629 | 2.98E-22 |
| INA1 | YLR413W | 1844.23633 | 0.9128372 | 0.09436344 | 9.67363188 | 3.90E-22 |
| MOT1 | YPL082C | 2379.33133 | 0.54040883 | 0.05589674 | 9.66798546 | 4.12E-22 |
| FET3 | YMR058W | 2425.89255 | -1.8195597 | 0.18853858 | -9.6508612 | 4.87E-22 |
| HTB1 | YDR224C | 2353.85127 | -0.636015 | 0.06591256 | -9.6493752 | 4.95E-22 |
| RNR2 | YJL026W | 3743.65463 | -0.6172574 | 0.06399611 | -9.6452328 | 5.15E-22 |
| Xu_2009_SUTs21 | Xu_2009_SU | 523.962202 | 0.79552672 | 0.08249917 | 9.64284509 | 5.27E-22 |
| PDB1 | YBR221C | 2107.41213 | -0.6256847 | 0.06490189 | -9.6404701 | 5.39E-22 |
| Xu_2009_CUTs43 | Xu_2009_CU | 245.845662 | 1.14830187 | 0.1191611 | 9.6365497 | 5.60E-22 |
| SBA1 | YKL117W | 2940.65558 | -0.5646142 | 0.05861099 | -9.6332488 | 5.79E-22 |
| YPP1 | YGR198W | 1604.35874 | 0.6126702 | 0.06361319 | 9.63118204 | 5.90E-22 |
| HSP82 | YPL240C | 21950.7478 | -0.7249922 | 0.07533989 | -9.6229526 | 6.40E-22 |
| RPS30B | YOR182C | 1272.31866 | -0.7385602 | 0.07681638 | -9.6146192 | 6.94E-22 |
| APC1 | YNL172W | 1162.54074 | 0.64028531 | 0.06659802 | 9.61417897 | 6.97E-22 |
| OPT1 | YJL212C | 405.828777 | 0.9070609 | 0.09444427 | 9.60419189 | 7.68E-22 |
| TOR2 | YKL203C | 3188.68789 | 0.5909769 | 0.06154567 | 9.60224968 | 7.82E-22 |

| | | | | | | |
|---|---|---|---|---|---|---|
| IRC22 | YEL001C | 1852.90547 | -0.5645112 | 0.05886298 | -9.5902581 | 8.79E-22 |
| Xu_2009_SUTs36 | Xu_2009_SU | 352.496298 | 0.95455528 | 0.09956337 | 9.58741454 | 9.03E-22 |
| IQG1 | YPL242C | 2309.67611 | 0.53994519 | 0.05634774 | 9.5823754 | 9.48E-22 |
| Xu_2009_CUTs42 | Xu_2009_CU | 166.837682 | 1.49614396 | 0.15617893 | 9.57967857 | 9.73E-22 |
| ELO3 | YLR372W | 1933.75116 | -0.6501079 | 0.06789877 | -9.5746651 | 1.02E-21 |
| SNR189 | snR189 | 10192.8532 | -1.2659618 | 0.13232328 | -9.5671886 | 1.10E-21 |
| MEC1 | YBR136W | 1601.46512 | 0.59728568 | 0.06254526 | 9.54965547 | 1.30E-21 |
| YJL068C | YJL068C | 842.13974 | -0.7368753 | 0.0772612 | -9.5374568 | 1.46E-21 |
| GUS1 | YGL245W | 6599.0406 | -0.5019186 | 0.05262812 | -9.5370806 | 1.47E-21 |
| HXT7 | YDR342C | 500.581315 | -0.9768128 | 0.10256534 | -9.5238096 | 1.67E-21 |
| GET3 | YDL100C | 2682.06806 | -0.5206441 | 0.05467566 | -9.5224104 | 1.69E-21 |
| MRPL17 | YNL252C | 371.832376 | -0.9552797 | 0.10031788 | -9.5225273 | 1.69E-21 |
| MRPS12 | YNR036C | 667.716833 | -1.0330697 | 0.10864753 | -9.5084508 | 1.94E-21 |
| TAF2 | YCR042C | 1372.04857 | 0.66216412 | 0.06969198 | 9.50129551 | 2.07E-21 |
| APA1 | YCL050C | 1718.84535 | -0.5931944 | 0.06251116 | -9.4894157 | 2.32E-21 |
| PHO5 | YBR093C | 757.795506 | -0.7553254 | 0.07962629 | -9.4858804 | 2.40E-21 |
| PAB1 | YER165W | 7965.76864 | -0.5730479 | 0.06051959 | -9.4688005 | 2.83E-21 |
| DIC1 | YLR348C | 1207.78802 | -0.7762376 | 0.08204136 | -9.4615402 | 3.03E-21 |
| RDL2 | YOR286W | 582.394507 | -0.804272 | 0.08501942 | -9.4598625 | 3.08E-21 |
| Xu_2009_SUTs46 | Xu_2009_SU | 254.141805 | 1.0453653 | 0.11051818 | 9.45876301 | 3.12E-21 |
| CYS3 | YAL012W | 2225.75809 | -0.5747793 | 0.06077805 | -9.4570216 | 3.17E-21 |
| SHM1 | YBR263W | 2273.89719 | -0.5210346 | 0.05512994 | -9.451029 | 3.36E-21 |
| Xu_2009_SUTs37 | Xu_2009_SU | 128.176674 | 1.41955521 | 0.15033844 | 9.44239671 | 3.64E-21 |
| ERG9 | YHR190W | 2296.58197 | -0.544506 | 0.05766695 | -9.4422552 | 3.65E-21 |
| PUP3 | YER094C | 2013.91094 | -0.5806175 | 0.06153255 | -9.4359417 | 3.87E-21 |
| Xu_2009_SUTs40 | Xu_2009_SU | 133.231349 | 1.37425478 | 0.14606286 | 9.40865295 | 5.03E-21 |
| TIM23 | YNR017W | 1108.85789 | -0.6337163 | 0.06735674 | -9.4083568 | 5.04E-21 |
| Xu_2009_SUTs68 | Xu_2009_SU | 508.762414 | 0.7945159 | 0.08449987 | 9.40256913 | 5.32E-21 |
| VMA4 | YOR332W | 5283.99055 | -0.5062033 | 0.05394994 | -9.3828337 | 6.42E-21 |
| TSC10 | YBR265W | 696.303502 | -0.7827696 | 0.08351237 | -9.3730977 | 7.04E-21 |
| TRP5 | YGL026C | 3292.33694 | -0.4985745 | 0.05320165 | -9.3714096 | 7.16E-21 |
| STE24 | YJR117W | 2095.09894 | -0.5494093 | 0.05862799 | -9.3711096 | 7.18E-21 |
| HXT6 | YDR343C | 96.4991923 | -1.6608463 | 0.17728724 | -9.3681095 | 7.38E-21 |
| GRE2 | YOL151W | 688.354082 | -0.742884 | 0.07941295 | -9.354696 | 8.38E-21 |
| ASK10 | YGR097W | 3196.42198 | 0.5020805 | 0.05371381 | 9.34732561 | 8.99E-21 |
| MIR1 | YJR077C | 2044.7845 | -0.6189547 | 0.06639617 | -9.3221444 | 1.14E-20 |
| SML1 | YML058W | 1085.49519 | -0.8042735 | 0.08633523 | -9.3157036 | 1.21E-20 |
| Xu_2009_SUTs58 | Xu_2009_SU | 276.202303 | 1.08683533 | 0.11681362 | 9.30401171 | 1.35E-20 |
| QCR8 | YJL166W | 934.27728 | -0.704403 | 0.07571517 | -9.3033265 | 1.36E-20 |
| SCD6 | YPR129W | 1408.45899 | -0.6172995 | 0.06644484 | -9.2904055 | 1.54E-20 |
| RET2 | YFR051C | 1341.24231 | -0.6857442 | 0.07383184 | -9.2879194 | 1.57E-20 |

| | | | | | | |
|---|---|---|---|---|---|---|
| ADH2 | YMR303C | 283.864012 | -1.1977794 | 0.12899996 | -9.2851147 | 1.62E-20 |
| YPR117W | YPR117W | 2413.6346 | 0.50028279 | 0.05387917 | 9.28527331 | 1.61E-20 |
| SND1 | YDR186C | 3412.22225 | 0.55829132 | 0.06013331 | 9.28422813 | 1.63E-20 |
| RSM19 | YNR037C | 438.811476 | -0.8979275 | 0.0968216 | -9.2740417 | 1.79E-20 |
| MRPL39 | YML009C | 250.313555 | -1.0677612 | 0.11521096 | -9.2678787 | 1.90E-20 |
| ECM32 | YER176W | 1457.05561 | 0.5744843 | 0.0619915 | 9.26714604 | 1.91E-20 |
| GSY2 | YLR258W | 4245.54154 | -0.6241874 | 0.06740292 | -9.2605395 | 2.03E-20 |
| NCA3 | YJL116C | 1631.89506 | 0.7541913 | 0.08146722 | 9.25760396 | 2.09E-20 |
| DNA2 | YHR164C | 1003.11704 | 0.66544022 | 0.07199127 | 9.24334567 | 2.39E-20 |
| THR1 | YHR025W | 2406.67208 | -0.5549873 | 0.06018402 | -9.2215057 | 2.93E-20 |
| HXK1 | YFR053C | 4535.3635 | -1.4001246 | 0.15195443 | -9.2141082 | 3.14E-20 |
| YJL107C | YJL107C | 421.136514 | 0.86624089 | 0.09422327 | 9.19349248 | 3.80E-20 |
| AVT7 | YIL088C | 1025.74701 | -0.6283132 | 0.06838376 | -9.1880477 | 4.00E-20 |
| IRC20 | YLR247C | 1922.70346 | 0.54046043 | 0.0588369 | 9.18573986 | 4.09E-20 |
| ARG4 | YHR018C | 1311.9536 | -0.6212138 | 0.06766425 | -9.1808274 | 4.28E-20 |
| ALD5 | YER073W | 2243.51658 | -0.6197874 | 0.06753192 | -9.1776956 | 4.40E-20 |
| NST1 | YNL091W | 3475.39662 | 0.56258102 | 0.0613526 | 9.16963656 | 4.75E-20 |
| SES1 | YDR023W | 3627.1373 | -0.5224141 | 0.05703882 | -9.1589215 | 5.24E-20 |
| MBA1 | YBR185C | 285.580751 | -1.0088253 | 0.11046364 | -9.1326452 | 6.68E-20 |
| ELO2 | YCR034W | 628.819333 | -0.8254994 | 0.09049022 | -9.1225256 | 7.34E-20 |
| MNL2 | YLR057W | 1124.43105 | 0.60604581 | 0.0664667 | 9.11803642 | 7.65E-20 |
| RPS10A | YOR293W | 6856.74027 | -0.5666842 | 0.06218234 | -9.113266 | 7.99E-20 |
| SNR190 | snR190 | 20609.3624 | -2.2404888 | 0.24603729 | -9.106298 | 8.52E-20 |
| YNL208W | YNL208W | 4753.55238 | -0.6558269 | 0.07213142 | -9.092112 | 9.71E-20 |
| TRX2 | YGR209C | 848.392615 | -0.8174124 | 0.09006541 | -9.0757644 | 1.13E-19 |
| TUB1 | YML085C | 2172.55438 | -0.5913831 | 0.06516211 | -9.075567 | 1.13E-19 |
| CCS1 | YMR038C | 767.329736 | -0.67439 | 0.07439642 | -9.0648179 | 1.25E-19 |
| RIE1 | YGR250C | 3268.64682 | 0.57331256 | 0.06325062 | 9.06414128 | 1.26E-19 |
| RPB8 | YOR224C | 1030.92726 | -0.6970615 | 0.07709178 | -9.0419691 | 1.54E-19 |
| RRN3 | YKL125W | 1615.70824 | 0.58910665 | 0.06520983 | 9.03401579 | 1.65E-19 |
| WWM1 | YFL010C | 2710.49396 | -0.572364 | 0.06336792 | -9.0323947 | 1.68E-19 |
| THP3 | YPR045C | 1462.73046 | 0.53421759 | 0.0592182 | 9.02117205 | 1.86E-19 |
| Xu_2009_SUTs14 | Xu_2009_SU | 155.345935 | 1.23818487 | 0.13751576 | 9.00394882 | 2.18E-19 |
| CLN1 | YMR199W | 1701.90176 | -0.5293494 | 0.0589517 | -8.9793752 | 2.72E-19 |
| SLH1 | YGR271W | 1826.4844 | 0.54175559 | 0.06035911 | 8.97553913 | 2.82E-19 |
| DSE1 | YER124C | 1877.33513 | -0.5936606 | 0.06617466 | -8.971116 | 2.94E-19 |
| HES1 | YOR237W | 427.003236 | 0.85337101 | 0.09525124 | 8.95915928 | 3.27E-19 |
| TOM40 | YMR203W | 1337.05268 | -0.6425502 | 0.07176855 | -8.9530888 | 3.46E-19 |
| BMS1 | YPL217C | 2194.18106 | 0.63104838 | 0.07049923 | 8.95113812 | 3.52E-19 |
| YBR137W | YBR137W | 540.742558 | -0.9788182 | 0.10938625 | -8.9482748 | 3.61E-19 |
| RPL9B | YNL067W | 2155.92885 | -0.7014018 | 0.07840744 | -8.9456026 | 3.70E-19 |

| | | | | | | |
|---|---|---|---|---|---|---|
| HSL1 | YKL101W | 4206.05922 | 0.58412616 | 0.06531307 | 8.94348016 | 3.77E-19 |
| DYS1 | YHR068W | 1411.8416 | -0.5598633 | 0.06265038 | -8.936312 | 4.02E-19 |
| POL30 | YBR088C | 573.901294 | -0.7984405 | 0.08935957 | -8.9351433 | 4.07E-19 |
| SAS10 | YDL153C | 751.361351 | 0.7284278 | 0.08154024 | 8.93335354 | 4.13E-19 |
| VAS1 | YGR094W | 5868.67694 | -0.5331168 | 0.05986338 | -8.905559 | 5.31E-19 |
| TOM70 | YNL121C | 3572.96583 | -0.4552452 | 0.05117185 | -8.8963993 | 5.77E-19 |
| SNR17B | snR17b | 84949.5747 | -1.3420209 | 0.15092077 | -8.8922217 | 5.99E-19 |
| FAB1 | YFR019W | 2909.39528 | 0.59435483 | 0.06685096 | 8.89074429 | 6.07E-19 |
| SSS1 | YDR086C | 1594.51061 | -0.5464499 | 0.06150097 | -8.8852239 | 6.38E-19 |
| EMC2 | YJR088C | 894.12181 | -0.7066241 | 0.07961021 | -8.8760485 | 6.93E-19 |
| YNL284C-B | YNL284C-B | 878.234775 | 0.6243988 | 0.07039434 | 8.87001405 | 7.31E-19 |
| HKR1 | YDR420W | 2492.39459 | 0.51702567 | 0.05830099 | 8.86821384 | 7.43E-19 |
| CPR6 | YLR216C | 3738.12283 | -0.5812819 | 0.06558037 | -8.8636571 | 7.74E-19 |
| CHS1 | YNL192W | 5338.78614 | 0.50487394 | 0.05697356 | 8.86154815 | 7.89E-19 |
| YPR1 | YDR368W | 1102.88752 | -0.6554701 | 0.07402511 | -8.8546993 | 8.39E-19 |
| YIP3 | YNL044W | 985.929417 | -0.7225624 | 0.08169821 | -8.8442878 | 9.21E-19 |
| ARP3 | YJR065C | 1682.77584 | -0.5447112 | 0.06159307 | -8.8437095 | 9.26E-19 |
| UFD4 | YKL010C | 2647.07495 | 0.54303982 | 0.06148415 | 8.83219266 | 1.03E-18 |
| PUS9 | YDL036C | 552.577858 | 0.7566976 | 0.08577933 | 8.82144477 | 1.13E-18 |
| SNU13 | YEL026W | 1273.68586 | -0.6822289 | 0.0774583 | -8.8076934 | 1.28E-18 |
| CAM1 | YPL048W | 4870.14511 | -0.4660857 | 0.05295994 | -8.8007203 | 1.36E-18 |
| HTZ1 | YOL012C | 349.494989 | -0.8847995 | 0.10068535 | -8.7877682 | 1.53E-18 |
| DTD1 | YDL219W | 396.551145 | -0.8477594 | 0.09659096 | -8.7767981 | 1.68E-18 |
| PCA1 | YBR295W | 772.300597 | 0.70458506 | 0.08028745 | 8.7757802 | 1.70E-18 |
| MLP2 | YIL149C | 2415.17588 | 0.52210704 | 0.05956376 | 8.7655157 | 1.86E-18 |
| GDE1 | YPL110C | 1121.75233 | 0.66972846 | 0.07659001 | 8.74433186 | 2.24E-18 |
| YDL157C | YDL157C | 474.385115 | -0.8387104 | 0.09595186 | -8.7409504 | 2.31E-18 |
| PEX1 | YKL197C | 1220.0229 | 0.61911744 | 0.07084954 | 8.73848216 | 2.36E-18 |
| PDR1 | YGL013C | 1502.68824 | 0.55990114 | 0.06407697 | 8.73794674 | 2.37E-18 |
| ARP2 | YDL029W | 1056.01358 | -0.680089 | 0.07787216 | -8.7334036 | 2.47E-18 |
| SNR41 | snR41 | 408.620706 | -0.8327795 | 0.09540519 | -8.72887 | 2.57E-18 |
| CCW22 | YDR134C | 1468.73059 | -0.5937549 | 0.06803025 | -8.7278068 | 2.60E-18 |
| SEC14 | YMR079W | 1774.21271 | -0.5418571 | 0.06209209 | -8.7266692 | 2.62E-18 |
| VMA7 | YGR020C | 1444.97906 | -0.597041 | 0.06844284 | -8.7232064 | 2.70E-18 |
| ATG42 | YBR139W | 2073.15862 | -0.6461848 | 0.07413184 | -8.7166965 | 2.86E-18 |
| PRE3 | YJL001W | 1121.52725 | -0.5978138 | 0.06862813 | -8.7109153 | 3.01E-18 |
| POM33 | YLL023C | 320.88164 | -0.8796193 | 0.10129151 | -8.6840378 | 3.82E-18 |
| TIM50 | YPL063W | 1998.99521 | -0.4989885 | 0.05754492 | -8.6712867 | 4.27E-18 |
| ARO10 | YDR380W | 305.127264 | -0.9487155 | 0.10942099 | -8.6703242 | 4.31E-18 |
| VMA8 | YEL051W | 1500.83045 | -0.6276401 | 0.07275436 | -8.6268379 | 6.31E-18 |
| YTA7 | YGR270W | 3085.04725 | 0.50890601 | 0.05903702 | 8.62011738 | 6.69E-18 |

| | | | | | | |
|---|---|---|---|---|---|---|
| LAS17 | YOR181W | 406.58499 | -0.8028508 | 0.09319065 | -8.6151428 | 6.99E-18 |
| ARA1 | YBR149W | 2893.10705 | -0.5208528 | 0.06047868 | -8.6121728 | 7.17E-18 |
| CIN1 | YOR349W | 624.434518 | 0.71285373 | 0.08279711 | 8.60964529 | 7.33E-18 |
| Xu_2009_CUTs56 | Xu_2009_CU | 90.3466962 | 1.49270741 | 0.17348509 | 8.60424042 | 7.68E-18 |
| CPR5 | YDR304C | 2320.7055 | -0.4826005 | 0.05612399 | -8.5988285 | 8.05E-18 |
| YKR070W | YKR070W | 643.383497 | -0.6735541 | 0.07838834 | -8.5925299 | 8.51E-18 |
| MLP1 | YKR095W | 3385.61234 | 0.46192763 | 0.05380086 | 8.58587762 | 9.01E-18 |
| YML6 | YML025C | 775.369951 | -0.6367564 | 0.07422011 | -8.5792979 | 9.55E-18 |
| SCP160 | YJL080C | 3960.06138 | -0.4220388 | 0.04919938 | -8.5781335 | 9.64E-18 |
| CYS4 | YGR155W | 7083.54908 | -0.6113221 | 0.07132894 | -8.570464 | 1.03E-17 |
| SAG1 | YJR004C | 506.44454 | 0.73827387 | 0.08614838 | 8.5697944 | 1.04E-17 |
| EHT1 | YBR177C | 1220.01263 | -0.607976 | 0.07097652 | -8.5658741 | 1.07E-17 |
| SCR1 | SCR1 | 585448.854 | -1.3690577 | 0.15990373 | -8.5617621 | 1.11E-17 |
| MSY1 | YPL097W | 731.55586 | -0.6506158 | 0.07618803 | -8.5396064 | 1.35E-17 |
| HMF1 | YER057C | 664.318173 | -0.716532 | 0.08392909 | -8.5373493 | 1.37E-17 |
| RPS30A | YLR287C-A | 6682.55578 | -0.5045951 | 0.0591494 | -8.5308568 | 1.45E-17 |
| YJL047C-A | YJL047C-A | 142.567941 | 1.18309492 | 0.13872284 | 8.52847961 | 1.48E-17 |
| PSY2 | YNL201C | 2365.49363 | 0.5170713 | 0.06064961 | 8.52555027 | 1.52E-17 |
| CBR1 | YIL043C | 1927.22558 | -0.5293276 | 0.06209119 | -8.5250032 | 1.53E-17 |
| YNL040W | YNL040W | 624.768873 | -0.6950312 | 0.08153768 | -8.5240495 | 1.54E-17 |
| CWP1 | YKL096W | 1931.45203 | -0.5626722 | 0.06601314 | -8.5236395 | 1.55E-17 |
| KRI1 | YNL308C | 1449.64382 | 0.75033926 | 0.0880578 | 8.52098623 | 1.58E-17 |
| MGE1 | YOR232W | 1671.42475 | -0.5923518 | 0.06952925 | -8.5194616 | 1.60E-17 |
| ERP6 | YGL002W | 355.89407 | -0.8794664 | 0.10359247 | -8.4896751 | 2.07E-17 |
| YOX1 | YML027W | 1174.71518 | -0.6271513 | 0.07387981 | -8.488804 | 2.09E-17 |
| TOP1 | YOL006C | 2324.94329 | 0.47396905 | 0.05583795 | 8.48829641 | 2.10E-17 |
| YJR027W | YJR027W | 1367.59567 | 0.60629303 | 0.07143292 | 8.48758584 | 2.11E-17 |
| KTR1 | YOR099W | 2080.24847 | -0.4689461 | 0.05525308 | -8.4872406 | 2.12E-17 |
| FDC1 | YDR539W | 440.731048 | -0.7734529 | 0.09145183 | -8.4574895 | 2.73E-17 |
| AIM45 | YPR004C | 910.817471 | -0.6340414 | 0.07499103 | -8.4548966 | 2.79E-17 |
| MCK1 | YNL307C | 1279.43793 | -0.5927685 | 0.07015409 | -8.4495224 | 2.92E-17 |
| MSS51 | YLR203C | 1223.90665 | -0.5483682 | 0.06495061 | -8.442848 | 3.10E-17 |
| GET1 | YGL020C | 410.414509 | -0.8353803 | 0.09909839 | -8.4298073 | 3.46E-17 |
| TOM6 | YOR045W | 175.846888 | -1.1734932 | 0.13926758 | -8.426176 | 3.57E-17 |
| RPL42A | YNL162W | 1700.05392 | -0.6603551 | 0.07838357 | -8.4246622 | 3.62E-17 |
| ADE13 | YLR359W | 1294.0729 | -0.6536708 | 0.07761646 | -8.4218073 | 3.71E-17 |
| TPD3 | YAL016W | 1523.81694 | -0.582173 | 0.06913737 | -8.4205256 | 3.75E-17 |
| AGP1 | YCL025C | 2009.78632 | -0.5549274 | 0.06599315 | -8.4088646 | 4.14E-17 |
| HHO1 | YPL127C | 543.056391 | -0.7263353 | 0.08640575 | -8.4060995 | 4.24E-17 |
| CIS3 | YJL158C | 7708.10676 | -0.5486259 | 0.0652779 | -8.4044657 | 4.30E-17 |
| TMH11 | YJR085C | 1132.4929 | -0.6575965 | 0.07845008 | -8.3823556 | 5.19E-17 |

| | | | | | | |
|---|---|---|---|---|---|---|
| SNR42 | snR42 | 7987.7952 | -1.7297132 | 0.20647599 | -8.377309 | 5.42E-17 |
| SAP185 | YJL098W | 1981.53452 | 0.52622485 | 0.06282439 | 8.37612341 | 5.47E-17 |
| MNT3 | YIL014W | 1220.72096 | 0.54285665 | 0.06489262 | 8.36546062 | 5.99E-17 |
| ERG20 | YJL167W | 5048.46064 | -0.4000425 | 0.04785167 | -8.3600535 | 6.27E-17 |
| YER097W | YER097W | 129.347266 | 1.2641414 | 0.15123408 | 8.35883925 | 6.33E-17 |
| TEL1 | YBL088C | 1409.21452 | 0.52877361 | 0.06333743 | 8.34851737 | 6.91E-17 |
| Xu_2009_SUTs28 | Xu_2009_SU | 226.662204 | 1.02345237 | 0.12270939 | 8.34045676 | 7.40E-17 |
| ASN1 | YPR145W | 2792.10997 | -0.4756627 | 0.05703258 | -8.3401935 | 7.42E-17 |
| TOP2 | YNL088W | 2629.38338 | 0.476938 | 0.05718979 | 8.33956511 | 7.46E-17 |
| HXT3 | YDR345C | 8783.47092 | -0.5685458 | 0.06823677 | -8.331957 | 7.95E-17 |
| LSB3 | YFR024C-A | 1459.05889 | -0.5768742 | 0.06924261 | -8.3312022 | 8.00E-17 |
| ECM15 | YBL001C | 813.884669 | -0.6567105 | 0.07895731 | -8.317285 | 9.00E-17 |
| Xu_2009_SUTs17 | Xu_2009_SU | 131.83577 | 1.21139866 | 0.14570679 | 8.31394808 | 9.26E-17 |
| Xu_2009_SUTs63 | Xu_2009_SU | 370.598115 | 0.92231826 | 0.11095119 | 8.31282866 | 9.34E-17 |
| HAP1 | YLR256W | 2116.35271 | 0.52447453 | 0.06312199 | 8.3089032 | 9.66E-17 |
| SWP1 | YMR149W | 1049.46 | -0.5869952 | 0.07069724 | -8.3029426 | 1.02E-16 |
| UGA1 | YGR019W | 977.527103 | -0.639541 | 0.07703214 | -8.3022622 | 1.02E-16 |
| tI(UAU)D | tI(UAU)D | 67.5468879 | -1.7900341 | 0.21567887 | -8.2995342 | 1.05E-16 |
| LEU4 | YNL104C | 3880.46119 | -0.5253196 | 0.06334701 | -8.2927289 | 1.11E-16 |
| GGC1 | YDL198C | 389.090154 | -0.8405249 | 0.10142667 | -8.2870205 | 1.16E-16 |
| CIT1 | YNR001C | 1750.57066 | -0.6836145 | 0.08249843 | -8.2863945 | 1.17E-16 |
| MDG1 | YNL173C | 669.706862 | -0.7532186 | 0.09104188 | -8.2733198 | 1.30E-16 |
| MRP8 | YKL142W | 1499.11028 | -0.6116463 | 0.07397813 | -8.2679335 | 1.36E-16 |
| PFK2 | YMR205C | 7691.49744 | -0.4538046 | 0.05489553 | -8.2666961 | 1.38E-16 |
| Xu_2009_SUTs48 | Xu_2009_SU | 111.214814 | 1.30515091 | 0.1580995 | 8.25524992 | 1.52E-16 |
| PMT1 | YDL095W | 4305.35658 | -0.4255716 | 0.05156769 | -8.2526782 | 1.55E-16 |
| Xu_2009_SUTs43 | Xu_2009_SU | 238.454639 | 0.92046578 | 0.11158296 | 8.24916057 | 1.60E-16 |
| OYE2 | YHR179W | 6364.0074 | -0.4004607 | 0.04856617 | -8.2456719 | 1.64E-16 |
| ABF2 | YMR072W | 1084.02791 | -0.6714567 | 0.08149157 | -8.2395847 | 1.73E-16 |
| YDJ1 | YNL064C | 3730.1892 | -0.4006612 | 0.04872246 | -8.2233358 | 1.98E-16 |
| CSS1 | YIL169C | 116.503485 | 1.36562489 | 0.16619172 | 8.21716568 | 2.08E-16 |
| CTI6 | YPL181W | 1853.85502 | 0.55279441 | 0.06732599 | 8.21071353 | 2.20E-16 |
| PER33 | YLR064W | 658.905428 | -0.6655907 | 0.08107582 | -8.2094848 | 2.22E-16 |
| PPA2 | YMR267W | 660.795297 | -0.7650779 | 0.09321957 | -8.2072669 | 2.26E-16 |
| ZUO1 | YGR285C | 6255.55634 | -0.406783 | 0.04957261 | -8.2058016 | 2.29E-16 |
| TPO1 | YLL028W | 1276.94328 | 0.53686964 | 0.06544731 | 8.20308149 | 2.34E-16 |
| AMN1 | YBR158W | 2306.40515 | -0.4490797 | 0.05476353 | -8.2003417 | 2.40E-16 |
| SWH1 | YAR042W | 2514.99729 | 0.51324905 | 0.06261646 | 8.19671172 | 2.47E-16 |
| Xu_2009_CUTs64 | Xu_2009_CU | 72.8395786 | 1.6619551 | 0.20278925 | 8.19547911 | 2.50E-16 |
| CAP2 | YIL034C | 1412.18501 | -0.5613225 | 0.06853484 | -8.1903237 | 2.61E-16 |
| GLR1 | YPL091W | 1254.66286 | -0.5555021 | 0.06785006 | -8.1872015 | 2.67E-16 |

| | | | | | | |
|---|---|---|---|---|---|---|
| HNM1 | YGL077C | 2209.80259 | -0.5165652 | 0.06313624 | -8.1817552 | 2.80E-16 |
| DPS1 | YLL018C | 5253.7356 | -0.4429843 | 0.05418233 | -8.175806 | 2.94E-16 |
| UPS1 | YLR193C | 340.014559 | -0.8379262 | 0.10255026 | -8.1708834 | 3.06E-16 |
| NNR1 | YNL200C | 848.91371 | -0.742731 | 0.09094362 | -8.1669394 | 3.16E-16 |
| Xu_2009_SUTs22 | Xu_2009_SU | 378.9789 | 0.80539816 | 0.09866492 | 8.16296368 | 3.27E-16 |
| Xu_2009_SUTs32 | Xu_2009_SU | 137.822151 | 1.20868819 | 0.14809436 | 8.16160858 | 3.31E-16 |
| MPC1 | YGL080W | 821.493725 | -0.5943617 | 0.07294342 | -8.1482563 | 3.69E-16 |
| GRX5 | YPL059W | 332.798536 | -0.8446835 | 0.10382063 | -8.1359891 | 4.09E-16 |
| CTR2 | YHR175W | 454.977513 | -0.716203 | 0.08803899 | -8.1350655 | 4.12E-16 |
| RPS28B | YLR264W | 525.955759 | -0.7127471 | 0.08761723 | -8.1347821 | 4.13E-16 |
| YLR177W | YLR177W | 612.050726 | -0.7906444 | 0.09720278 | -8.1339684 | 4.15E-16 |
| RPL31B | YLR406C | 2109.49491 | -0.5169238 | 0.06356158 | -8.1326459 | 4.20E-16 |
| MDH3 | YDL078C | 927.226288 | -0.6020303 | 0.07403317 | -8.1318986 | 4.23E-16 |
| MCD4 | YKL165C | 805.779483 | -0.6053705 | 0.07455772 | -8.1194874 | 4.68E-16 |
| FMP27 | YLR454W | 2806.64277 | 0.45696122 | 0.05630144 | 8.11633307 | 4.80E-16 |
| UTP25 | YIL091C | 1784.40028 | 0.5873141 | 0.07240047 | 8.11202038 | 4.98E-16 |
| FIT2 | YOR382W | 423.826715 | -1.4536316 | 0.1792632 | -8.1089238 | 5.11E-16 |
| SNR47 | snR47 | 250.693496 | 1.11875485 | 0.13801678 | 8.10593357 | 5.23E-16 |
| PMT4 | YJR143C | 1896.17908 | -0.5370492 | 0.06627554 | -8.1032798 | 5.35E-16 |
| BGL2 | YGR282C | 4013.28626 | -0.5418902 | 0.06687836 | -8.1026244 | 5.38E-16 |
| IRC24 | YIR036C | 455.182638 | -0.742711 | 0.09170617 | -8.0988118 | 5.55E-16 |
| SPB1 | YCL054W | 2281.81003 | 0.5449118 | 0.06731306 | 8.09518722 | 5.72E-16 |
| AIM6 | YDL237W | 721.405792 | -0.6702325 | 0.0828012 | -8.0944771 | 5.75E-16 |
| PIS1 | YPR113W | 1115.45381 | -0.5693568 | 0.0703469 | -8.0935595 | 5.79E-16 |
| HEK2 | YBL032W | 1114.64548 | -0.5285607 | 0.06536697 | -8.0860513 | 6.16E-16 |
| CAF20 | YOR276W | 2217.88324 | -0.4991748 | 0.06174556 | -8.0843833 | 6.25E-16 |
| Xu_2009_CUTs68 | Xu_2009_CU | 87.5989834 | 1.38514326 | 0.17147122 | 8.07799274 | 6.58E-16 |
| YDR034C-D | YDR034C-D | 771.238212 | 0.66015661 | 0.08177096 | 8.07324006 | 6.85E-16 |
| FPR2 | YDR519W | 502.596738 | -0.6947595 | 0.08620068 | -8.0597911 | 7.64E-16 |
| Xu_2009_SUTs25 | Xu_2009_SU | 468.2282 | 0.7439571 | 0.09231806 | 8.05863028 | 7.72E-16 |
| YAR009C | YAR009C | 671.25678 | 0.70228699 | 0.08735527 | 8.03943433 | 9.03E-16 |
| Xu_2009_SUTs63 | Xu_2009_SU | 264.389639 | 0.89537652 | 0.11142439 | 8.03573221 | 9.30E-16 |
| DSE4 | YNR067C | 3622.57454 | -0.5920266 | 0.07379921 | -8.0221262 | 1.04E-15 |
| BUD9 | YGR041W | 919.68524 | -0.5972728 | 0.07453661 | -8.0131461 | 1.12E-15 |
| BUR6 | YER159C | 972.647996 | 0.64092764 | 0.0799922 | 8.0123764 | 1.13E-15 |
| NME1 | NME1 | 12496.7796 | -1.4505085 | 0.18109283 | -8.0097511 | 1.15E-15 |
| PIR3 | YKL163W | 1143.8455 | 0.61471853 | 0.0768566 | 7.99825297 | 1.26E-15 |
| COQ9 | YLR201C | 460.244055 | -0.7245728 | 0.09061384 | -7.9962709 | 1.28E-15 |
| FAA1 | YOR317W | 3251.14364 | -0.5439506 | 0.06803739 | -7.9948787 | 1.30E-15 |
| ARG3 | YJL088W | 305.723173 | -0.8920559 | 0.1116914 | -7.9867912 | 1.38E-15 |
| MSF1 | YPR047W | 475.256043 | -0.7373781 | 0.09240493 | -7.9798566 | 1.47E-15 |

| | | | | | | |
|---|---|---|---|---|---|---|
| LSC1 | YOR142W | 1385.84685 | -0.5106507 | 0.06402989 | -7.9751927 | 1.52E-15 |
| DIP2 | YLR129W | 2847.75863 | 0.53589673 | 0.06724282 | 7.96957486 | 1.59E-15 |
| ARG8 | YOL140W | 601.443394 | -0.6768625 | 0.0849978 | -7.9632945 | 1.68E-15 |
| SCW11 | YGL028C | 3766.01188 | -0.5985329 | 0.07522656 | -7.956404 | 1.77E-15 |
| VID24 | YBR105C | 3916.99524 | 0.61140013 | 0.07690627 | 7.94993916 | 1.87E-15 |
| SEC22 | YLR268W | 1179.49434 | -0.5668053 | 0.0713025 | -7.9493049 | 1.88E-15 |
| NHA1 | YLR138W | 2367.56009 | 0.44970443 | 0.05660629 | 7.94442468 | 1.95E-15 |
| MTH1 | YDR277C | 582.505285 | -0.7054297 | 0.08883751 | -7.9406742 | 2.01E-15 |
| BSC1 | YDL037C | 1990.87608 | 0.59496186 | 0.07499988 | 7.93283713 | 2.14E-15 |
| OST1 | YJL002C | 2056.3618 | -0.4709455 | 0.05939275 | -7.9293439 | 2.20E-15 |
| OST5 | YGL226C-A | 299.222463 | -0.8308279 | 0.10482882 | -7.925567 | 2.27E-15 |
| Xu_2009_SUTs45( | Xu_2009_SU | 380.513217 | 0.77169303 | 0.09738326 | 7.92428835 | 2.29E-15 |
| Xu_2009_CUTs51 | Xu_2009_CU | 125.669468 | 1.18872262 | 0.15041541 | 7.90293108 | 2.72E-15 |
| MRH1 | YDR033W | 5530.25218 | -0.4939526 | 0.06250742 | -7.9023032 | 2.74E-15 |
| YSA1 | YBR111C | 1306.54471 | -0.5684568 | 0.07194648 | -7.9011063 | 2.76E-15 |
| ILS1 | YBL076C | 6014.58239 | -0.3975737 | 0.05032361 | -7.9003404 | 2.78E-15 |
| PER1 | YCR044C | 443.072696 | -0.7546779 | 0.09555422 | -7.8979026 | 2.84E-15 |
| PSD1 | YNL169C | 1298.95333 | -0.5086379 | 0.06444306 | -7.8928263 | 2.95E-15 |
| SUI1 | YNL244C | 2035.07017 | -0.4665368 | 0.0591259 | -7.8905668 | 3.01E-15 |
| SCS3 | YGL126W | 649.612187 | -0.7488632 | 0.09491935 | -7.8894681 | 3.03E-15 |
| Xu_2009_CUTs23 | Xu_2009_CU | 79.4916308 | 1.48073225 | 0.18790072 | 7.8803969 | 3.26E-15 |
| NCP1 | YHR042W | 2965.33245 | -0.4834859 | 0.06135274 | -7.880429 | 3.26E-15 |
| MRP4 | YHL004W | 1141.27378 | -0.5678932 | 0.07223111 | -7.8621692 | 3.78E-15 |
| FLO10 | YKR102W | 253.899782 | 0.9076275 | 0.11569379 | 7.84508367 | 4.33E-15 |
| RAD2 | YGR258C | 1255.2924 | 0.52965261 | 0.06766666 | 7.82737974 | 4.98E-15 |
| TPM1 | YNL079C | 6923.26547 | -0.4102835 | 0.05255502 | -7.806742 | 5.87E-15 |
| POP1 | YNL221C | 2078.18483 | 0.4797715 | 0.06146816 | 7.80520338 | 5.94E-15 |
| YPR127W | YPR127W | 452.192596 | -0.7088029 | 0.09081526 | -7.8048872 | 5.96E-15 |
| ARO9 | YHR137W | 426.683614 | -0.8102495 | 0.10387646 | -7.8001265 | 6.18E-15 |
| MRPL36 | YBR122C | 1008.63806 | -0.5836999 | 0.07484476 | -7.7988072 | 6.25E-15 |
| COA1 | YIL157C | 845.560876 | -0.574451 | 0.0736667 | -7.7979729 | 6.29E-15 |
| YNR066C | YNR066C | 369.497097 | 0.8127372 | 0.10423342 | 7.79728055 | 6.33E-15 |
| KGD2 | YDR148C | 1098.62433 | -0.5487167 | 0.07039075 | -7.7952953 | 6.43E-15 |
| ARG7 | YMR062C | 811.124524 | -0.5800316 | 0.07441368 | -7.79469 | 6.46E-15 |
| FUN30 | YAL019W | 2374.88782 | 0.4684038 | 0.06011139 | 7.7922641 | 6.58E-15 |
| LOG1 | YJL055W | 1969.52393 | -0.5234961 | 0.06721458 | -7.7884309 | 6.78E-15 |
| SNR4 | snR4 | 50525.5914 | -1.2440195 | 0.15978283 | -7.7856894 | 6.93E-15 |
| LAP3 | YNL239W | 801.826746 | -0.5837536 | 0.07497826 | -7.7856378 | 6.94E-15 |
| PRM10 | YJL108C | 287.747179 | 0.83136061 | 0.10681123 | 7.78345658 | 7.06E-15 |
| ACA1 | YER045C | 729.080537 | 0.64957555 | 0.08351175 | 7.77825305 | 7.35E-15 |
| MRPS9 | YBR146W | 970.586728 | -0.5765714 | 0.07419569 | -7.7709556 | 7.79E-15 |

| | | | | | | |
|---|---|---|---|---|---|---|
| PDX3 | YBR035C | 772.656499 | -0.6076647 | 0.07820359 | -7.7702923 | 7.83E-15 |
| ZWF1 | YNL241C | 2664.91136 | -0.4480946 | 0.05767431 | -7.7693965 | 7.89E-15 |
| Xu_2009_CUTs33 | Xu_2009_CU | 65.7022922 | 1.59707351 | 0.20562202 | 7.76703549 | 8.03E-15 |
| HBT1 | YDL223C | 1512.6863 | 0.56221742 | 0.07240985 | 7.76437753 | 8.20E-15 |
| DBP5 | YOR046C | 1602.63408 | -0.4977212 | 0.06413526 | -7.7604911 | 8.46E-15 |
| MIG1 | YGL035C | 1132.76892 | -0.5625429 | 0.07251111 | -7.7580243 | 8.63E-15 |
| RPS23A | YGR118W | 3399.51199 | -0.613863 | 0.07925127 | -7.7457816 | 9.50E-15 |
| Xu_2009_SUTs26 | Xu_2009_SU | 172.314187 | 1.0147667 | 0.13103901 | 7.74400441 | 9.63E-15 |
| CYB2 | YML054C | 253.040848 | -0.9339789 | 0.12065786 | -7.7407219 | 9.89E-15 |
| BNA4 | YBL098W | 418.478514 | -0.717045 | 0.09280244 | -7.7265749 | 1.10E-14 |
| TIM13 | YGR181W | 360.683226 | -0.8589019 | 0.11118685 | -7.724851 | 1.12E-14 |
| SVP26 | YHR181W | 342.814492 | -0.8150996 | 0.10560745 | -7.7182014 | 1.18E-14 |
| FIG2 | YCR089W | 787.338637 | 0.583983 | 0.07566631 | 7.71787324 | 1.18E-14 |
| OLI1 | Q0130 | 2569.02825 | -1.2503845 | 0.16211062 | -7.7131563 | 1.23E-14 |
| Xu_2009_SUTs72 | Xu_2009_SU | 190.633949 | 0.95811757 | 0.1243193 | 7.70690912 | 1.29E-14 |
| YPF1 | YKL100C | 1369.09914 | -0.5063876 | 0.06570492 | -7.7069968 | 1.29E-14 |
| RPT4 | YOR259C | 1740.59295 | -0.448909 | 0.0582552 | -7.7059041 | 1.30E-14 |
| MPM1 | YJL066C | 611.208339 | -0.6718231 | 0.08719813 | -7.7045586 | 1.31E-14 |
| MRC1 | YCL061C | 1222.50563 | 0.54009594 | 0.07023486 | 7.68985532 | 1.47E-14 |
| LAP2 | YNL045W | 1323.03774 | -0.499783 | 0.0650216 | -7.6864147 | 1.51E-14 |
| SEC11 | YIR022W | 476.471326 | -0.6677713 | 0.08689978 | -7.6843836 | 1.54E-14 |
| PRP8 | YHR165C | 4078.57877 | 0.40783981 | 0.05310816 | 7.67941952 | 1.60E-14 |
| ARC35 | YNR035C | 1161.30758 | -0.5515216 | 0.07184508 | -7.6765404 | 1.63E-14 |
| ATP3 | YBR039W | 2047.76975 | -0.4823521 | 0.06288795 | -7.6700241 | 1.72E-14 |
| YHR214C-B | YHR214C-B | 1564.37213 | 0.54613463 | 0.07120274 | 7.67013487 | 1.72E-14 |
| UPC2 | YDR213W | 796.37387 | 0.60656794 | 0.07913532 | 7.66494591 | 1.79E-14 |
| YGR109W-B | YGR109W-B | 644.750579 | 0.69471139 | 0.09064926 | 7.66372926 | 1.81E-14 |
| TOR1 | YJR066W | 2532.53488 | 0.44118556 | 0.05758042 | 7.66207581 | 1.83E-14 |
| SER33 | YIL074C | 641.740821 | -0.6925906 | 0.09046953 | -7.6555122 | 1.93E-14 |
| Xu_2009_CUTs53 | Xu_2009_CU | 226.838559 | 0.9343637 | 0.12212736 | 7.65073197 | 2.00E-14 |
| Xu_2009_CUTs42 | Xu_2009_CU | 92.3803189 | 1.31717639 | 0.17217386 | 7.65026947 | 2.01E-14 |
| MCO8 | YIL156W-B | 266.047443 | -0.8702637 | 0.11375437 | -7.6503762 | 2.00E-14 |
| MRPL19 | YNL185C | 407.684998 | -0.7520673 | 0.09833779 | -7.6477955 | 2.04E-14 |
| LPL1 | YOR059C | 755.84468 | 0.59413629 | 0.07772803 | 7.64378461 | 2.11E-14 |
| AIP5 | YFR016C | 1961.88068 | 0.64211493 | 0.08404198 | 7.64040723 | 2.17E-14 |
| ATP16 | YDL004W | 1967.76219 | -0.4685627 | 0.06133768 | -7.639069 | 2.19E-14 |
| URA2 | YJL130C | 8635.37062 | 0.42097727 | 0.05520371 | 7.62588762 | 2.42E-14 |
| EMC6 | YLL014W | 352.837168 | -0.8064695 | 0.10575927 | -7.6255214 | 2.43E-14 |
| PXR1 | YGR280C | 471.450967 | 0.81504453 | 0.10689835 | 7.62448163 | 2.45E-14 |
| DUT1 | YBR252W | 399.992747 | -0.7300922 | 0.0957653 | -7.6237655 | 2.46E-14 |
| RPE1 | YJL121C | 1646.90076 | -0.4768445 | 0.0625666 | -7.6213905 | 2.51E-14 |

| | | | | | | |
|---|---|---|---|---|---|---|
| Xu_2009_SUTs4 | Xu_2009_SU | 303.767035 | 0.81023272 | 0.10639681 | 7.61519749 | 2.63E-14 |
| YGR026W | YGR026W | 914.558799 | -0.5861885 | 0.07701729 | -7.6111286 | 2.72E-14 |
| PTR2 | YKR093W | 2138.60668 | -0.4794351 | 0.06299294 | -7.610935 | 2.72E-14 |
| RDS1 | YCR106W | 277.07659 | 0.88076453 | 0.11573496 | 7.61018561 | 2.74E-14 |
| TSC11 | YER093C | 1412.39221 | 0.47391012 | 0.06231054 | 7.60561712 | 2.84E-14 |
| YLR419W | YLR419W | 1863.92724 | 0.46395009 | 0.06104738 | 7.59983602 | 2.97E-14 |
| USO1 | YDL058W | 2335.77837 | 0.47884769 | 0.06301309 | 7.59917861 | 2.98E-14 |
| PHM8 | YER037W | 1611.17195 | 0.57316006 | 0.07555391 | 7.58610732 | 3.30E-14 |
| YOP1 | YPR028W | 1816.5058 | -0.5917674 | 0.07804989 | -7.581912 | 3.40E-14 |
| YDR042C | YDR042C | 215.314993 | 0.95592633 | 0.12614072 | 7.57825346 | 3.50E-14 |
| RET3 | YPL010W | 685.938493 | -0.6312304 | 0.08331107 | -7.5767891 | 3.54E-14 |
| MRPL1 | YDR116C | 494.497616 | -0.6891016 | 0.09098472 | -7.5738177 | 3.62E-14 |
| PUP1 | YOR157C | 941.754261 | -0.5234845 | 0.06911886 | -7.5736853 | 3.63E-14 |
| STF2 | YGR008C | 1351.81831 | -0.605644 | 0.08000324 | -7.5702432 | 3.73E-14 |
| Xu_2009_CUTs49 | Xu_2009_CU | 130.299079 | 1.20914885 | 0.15994004 | 7.56001366 | 4.03E-14 |
| ORM2 | YLR350W | 2186.69062 | -0.4740558 | 0.06272492 | -7.5576956 | 4.10E-14 |
| OM14 | YBR230C | 1247.22397 | -0.4970682 | 0.06581318 | -7.5527144 | 4.26E-14 |
| SCS7 | YMR272C | 1659.7057 | -0.4600118 | 0.06091656 | -7.5515056 | 4.30E-14 |
| SND3 | YBR106W | 1991.39138 | -0.5110094 | 0.06787869 | -7.5282747 | 5.14E-14 |
| GLC3 | YEL011W | 612.343772 | -0.7603966 | 0.10103913 | -7.5257632 | 5.24E-14 |
| ACB1 | YGR037C | 1791.53371 | -0.573553 | 0.07625504 | -7.5215098 | 5.41E-14 |
| MRPL35 | YDR322W | 459.99601 | -0.6967658 | 0.09265094 | -7.5203312 | 5.46E-14 |
| VTC2 | YFL004W | 1832.0543 | -0.447747 | 0.05958849 | -7.5139836 | 5.74E-14 |
| Xu_2009_SUTs44 | Xu_2009_SU | 665.562655 | 0.6005977 | 0.07994611 | 7.51253195 | 5.80E-14 |
| BAT2 | YJR148W | 1520.51442 | -0.4851529 | 0.06461866 | -7.5079388 | 6.01E-14 |
| Xu_2009_CUTs41 | Xu_2009_CU | 96.7132443 | 1.23935212 | 0.16511978 | 7.50577643 | 6.11E-14 |
| Xu_2009_CUTs51 | Xu_2009_CU | 120.238798 | 1.20239729 | 0.16027257 | 7.50220241 | 6.28E-14 |
| PAM18 | YLR008C | 870.284141 | -0.5664977 | 0.07552967 | -7.5003325 | 6.37E-14 |
| NUP116 | YMR047C | 2359.46717 | 0.42103687 | 0.05614705 | 7.49882418 | 6.44E-14 |
| MYO4 | YAL029C | 2652.84919 | 0.41459303 | 0.05529503 | 7.4978348 | 6.49E-14 |
| YOR192C-B | YOR192C-B | 558.223686 | 0.64747375 | 0.08637318 | 7.49623578 | 6.57E-14 |
| COX17 | YLL009C | 452.22044 | -0.7270557 | 0.09714549 | -7.4841943 | 7.20E-14 |
| ADR1 | YDR216W | 1697.74216 | 0.44931885 | 0.06004749 | 7.48272471 | 7.28E-14 |
| STT3 | YGL022W | 4172.65287 | -0.3917123 | 0.05234876 | -7.4827429 | 7.28E-14 |
| CDC21 | YOR074C | 504.018814 | -0.6760242 | 0.0903801 | -7.4797895 | 7.44E-14 |
| MGM101 | YJR144W | 734.357821 | -0.5938769 | 0.07957124 | -7.4634627 | 8.43E-14 |
| MRPL16 | YBL038W | 664.68448 | -0.5691976 | 0.07632821 | -7.4572378 | 8.84E-14 |
| EBS1 | YDR206W | 1189.31617 | 0.48410752 | 0.06505055 | 7.44202033 | 9.92E-14 |
| MAK16 | YAL025C | 1423.13716 | 0.64257798 | 0.08634783 | 7.44173843 | 9.94E-14 |
| ERT1 | YBR239C | 644.564208 | 0.5848245 | 0.07863044 | 7.43763449 | 1.03E-13 |
| Xu_2009_CUTs17 | Xu_2009_CU | 87.381431 | 1.30506247 | 0.17568831 | 7.42828301 | 1.10E-13 |

| | | | | | | |
|---|---|---|---|---|---|---|
| DOG2 | YHR043C | 490.430483 | -0.7174844 | 0.09661968 | -7.4258618 | 1.12E-13 |
| MEF1 | YLR069C | 1404.0332 | -0.4959159 | 0.06678805 | -7.425219 | 1.13E-13 |
| JHD2 | YJR119C | 604.464703 | 0.63712957 | 0.08582832 | 7.42330251 | 1.14E-13 |
| NIT3 | YLR351C | 862.568901 | -0.6152782 | 0.08295201 | -7.4172795 | 1.20E-13 |
| YDR170W-A | YDR170W-A | 458.221275 | 0.70073167 | 0.09451133 | 7.41426089 | 1.22E-13 |
| MRPL23 | YOR150W | 370.278461 | -0.8350308 | 0.11266008 | -7.4119491 | 1.24E-13 |
| RPN10 | YHR200W | 1270.26317 | -0.4558675 | 0.06152166 | -7.409869 | 1.26E-13 |
| SSK22 | YCR073C | 630.050623 | 0.60362283 | 0.08147063 | 7.40908505 | 1.27E-13 |
| ILM1 | YJR118C | 386.581141 | -0.6977399 | 0.09417745 | -7.4087797 | 1.27E-13 |
| TCP1 | YDR212W | 2212.39479 | -0.4707986 | 0.06358953 | -7.4037128 | 1.32E-13 |
| NOP10 | YHR072W-A | 1135.86973 | -0.5723809 | 0.0773363 | -7.401194 | 1.35E-13 |
| MAL33 | YBR297W | 343.706977 | 0.7929912 | 0.1071896 | 7.39802372 | 1.38E-13 |
| ARO1 | YDR127W | 3416.46797 | -0.4265429 | 0.05768459 | -7.3943987 | 1.42E-13 |
| YSC84 | YHR016C | 463.028965 | -0.7100192 | 0.09604292 | -7.3927284 | 1.44E-13 |
| YCL049C | YCL049C | 387.930082 | 0.71518379 | 0.09674529 | 7.39244019 | 1.44E-13 |
| YPL216W | YPL216W | 455.319366 | 0.6771931 | 0.0917737 | 7.37894525 | 1.60E-13 |
| THG1 | YGR024C | 509.636345 | -0.6381403 | 0.08651429 | -7.3761252 | 1.63E-13 |
| ATP15 | YPL271W | 1335.68886 | -0.5409757 | 0.07340575 | -7.3696634 | 1.71E-13 |
| EMP24 | YGL200C | 2262.08107 | -0.4309108 | 0.05849591 | -7.366511 | 1.75E-13 |
| YNL134C | YNL134C | 1706.99268 | -0.4677144 | 0.06349574 | -7.366075 | 1.76E-13 |
| MCM3 | YEL032W | 2596.40983 | 0.42247037 | 0.05735544 | 7.36582863 | 1.76E-13 |
| GCN1 | YGL195W | 8325.7611 | 0.37315773 | 0.05068479 | 7.3623223 | 1.81E-13 |
| SUI2 | YJR007W | 1919.98349 | -0.5712548 | 0.07759508 | -7.3619974 | 1.81E-13 |
| RPS9A | YPL081W | 1784.61755 | -0.5428519 | 0.07375311 | -7.360393 | 1.83E-13 |
| ARC15 | YIL062C | 678.157095 | -0.5906249 | 0.08043534 | -7.3428532 | 2.09E-13 |
| DPH5 | YLR172C | 1625.7232 | -0.4322821 | 0.05888261 | -7.3414228 | 2.11E-13 |
| URA10 | YMR271C | 152.642696 | -1.0542429 | 0.14360023 | -7.3415122 | 2.11E-13 |
| MSH5 | YDL154W | 288.860323 | 0.82564276 | 0.11247274 | 7.3408253 | 2.12E-13 |
| URA5 | YML106W | 1591.34027 | -0.4678673 | 0.06373695 | -7.3405982 | 2.13E-13 |
| AIM9 | YER080W | 1445.52738 | -0.4676694 | 0.0637672 | -7.3340125 | 2.23E-13 |
| COX15 | YER141W | 1095.30237 | -0.5496 | 0.07502551 | -7.3255083 | 2.38E-13 |
| SNQ2 | YDR011W | 2442.03389 | 0.44263372 | 0.06043268 | 7.32441021 | 2.40E-13 |
| MIM1 | YOL026C | 499.122126 | -0.6199616 | 0.08464158 | -7.3245518 | 2.40E-13 |
| ERV41 | YML067C | 855.72028 | -0.5523063 | 0.07549202 | -7.3160882 | 2.55E-13 |
| MNN5 | YJL186W | 1376.41062 | -0.4715596 | 0.06454013 | -7.3064556 | 2.74E-13 |
| ALR2 | YFL050C | 584.419833 | 0.61166969 | 0.08376548 | 7.30216907 | 2.83E-13 |
| Xu_2009_SUTs58 | Xu_2009_SU | 126.038369 | 1.06628676 | 0.1461292 | 7.29687689 | 2.95E-13 |
| Xu_2009_SUTs55 | Xu_2009_SU | 294.675875 | 0.83897557 | 0.11501245 | 7.29465004 | 2.99E-13 |
| Xu_2009_SUTs43 | Xu_2009_SU | 295.009552 | 0.7768859 | 0.10672424 | 7.2793765 | 3.35E-13 |
| TRX3 | YCR083W | 557.834529 | -0.7099855 | 0.09759947 | -7.2744808 | 3.48E-13 |
| MNN11 | YJL183W | 693.104406 | -0.5461363 | 0.07510586 | -7.2715539 | 3.55E-13 |

| | | | | | | |
|---|---|---|---|---|---|---|
| SER2 | YGR208W | 537.28703 | -0.6722879 | 0.09250981 | -7.2672058 | 3.67E-13 |
| IRC13 | YOR235W | 237.944811 | -0.9042081 | 0.12443951 | -7.2662464 | 3.70E-13 |
| BUD14 | YAR014C | 1798.12864 | 0.48287088 | 0.06645739 | 7.26587188 | 3.71E-13 |
| Xu_2009_SUTs51 | Xu_2009_SU | 123.747429 | 1.08833724 | 0.14980404 | 7.26507272 | 3.73E-13 |
| MRP1 | YDR347W | 880.884739 | -0.5508256 | 0.07581758 | -7.2651429 | 3.73E-13 |
| Xu_2009_SUTs17 | Xu_2009_SU | 72.2504683 | 1.35159029 | 0.18607342 | 7.2637474 | 3.77E-13 |
| GRX2 | YDR513W | 948.172604 | -0.6038442 | 0.08314596 | -7.2624597 | 3.80E-13 |
| DBP2 | YNL112W | 6124.0866 | 0.59266963 | 0.08165399 | 7.25830617 | 3.92E-13 |
| PHO86 | YJL117W | 1459.805 | -0.4497667 | 0.06196973 | -7.2578447 | 3.93E-13 |
| Xu_2009_SUTs89 | Xu_2009_SU | 296.567451 | 0.83217771 | 0.11466908 | 7.25721081 | 3.95E-13 |
| Xu_2009_CUTs28 | Xu_2009_CU | 199.729239 | 0.98545403 | 0.13606481 | 7.24253437 | 4.40E-13 |
| PCL9 | YDL179W | 666.146023 | -0.6073578 | 0.08393123 | -7.2363742 | 4.61E-13 |
| UTP21 | YLR409C | 2217.90514 | 0.46631736 | 0.06445338 | 7.23495549 | 4.66E-13 |
| HFD1 | YMR110C | 917.188295 | -0.5772194 | 0.07982378 | -7.2311701 | 4.79E-13 |
| Xu_2009_SUTs58 | Xu_2009_SU | 114.07395 | 1.1575881 | 0.16012273 | 7.22938037 | 4.85E-13 |
| ZRG8 | YER033C | 1277.29123 | 0.49764459 | 0.06893533 | 7.21900642 | 5.24E-13 |
| Xu_2009_SUTs25 | Xu_2009_SU | 93.2551852 | 1.31688859 | 0.18249802 | 7.21590601 | 5.36E-13 |
| GLN4 | YOR168W | 2365.13032 | -0.4502145 | 0.06241753 | -7.2129503 | 5.48E-13 |
| SPT7 | YBR081C | 1485.17528 | 0.5070003 | 0.07030287 | 7.21165871 | 5.53E-13 |
| UBC7 | YMR022W | 660.364068 | -0.587406 | 0.08145326 | -7.2115717 | 5.53E-13 |
| GLC7 | YER133W | 2582.0346 | -0.4082292 | 0.05661741 | -7.2103116 | 5.58E-13 |
| YPL247C | YPL247C | 506.002475 | -0.6240212 | 0.0866014 | -7.2056712 | 5.78E-13 |
| TDA3 | YHR009C | 2663.99582 | -0.3774506 | 0.05239042 | -7.2045723 | 5.82E-13 |
| SNU66 | YOR308C | 747.033861 | 0.6063268 | 0.08417582 | 7.20309909 | 5.89E-13 |
| Xu_2009_CUTs36 | Xu_2009_CU | 74.9876216 | 1.31091951 | 0.18201704 | 7.20218019 | 5.93E-13 |
| SNT1 | YCR033W | 2879.17716 | 0.40863384 | 0.05673758 | 7.20217267 | 5.93E-13 |
| DUS3 | YLR401C | 1878.86854 | 0.54053849 | 0.07511205 | 7.19642799 | 6.18E-13 |
| MLC1 | YGL106W | 1398.68134 | -0.4971928 | 0.06909497 | -7.1957884 | 6.21E-13 |
| ELO1 | YJL196C | 634.474425 | -0.5761032 | 0.08014003 | -7.1887073 | 6.54E-13 |
| MPD1 | YOR288C | 399.886887 | -0.77704 | 0.10811952 | -7.1868608 | 6.63E-13 |
| SWC5 | YBR231C | 463.465157 | 0.67040156 | 0.09342755 | 7.17563069 | 7.20E-13 |
| COX4 | YGL187C | 600.802065 | -0.6380299 | 0.08896737 | -7.1715042 | 7.42E-13 |
| Xu_2009_SUTs55 | Xu_2009_SU | 220.925294 | 0.85076437 | 0.11866087 | 7.16971312 | 7.52E-13 |
| WRS1 | YOL097C | 1057.83391 | -0.4741346 | 0.06620902 | -7.1611782 | 8.00E-13 |
| BUD27 | YFL023W | 1163.16104 | 0.50129054 | 0.07007922 | 7.15319848 | 8.48E-13 |
| FHL1 | YPR104C | 1175.68806 | 0.47903757 | 0.06701904 | 7.1477833 | 8.82E-13 |
| MRPL50 | YNR022C | 514.36507 | -0.6613733 | 0.09257323 | -7.1443251 | 9.04E-13 |
| YPL114W | YPL114W | 231.837274 | 0.88500838 | 0.12389215 | 7.14337755 | 9.11E-13 |
| APM1 | YPL259C | 897.243458 | -0.4863458 | 0.06810214 | -7.1414169 | 9.24E-13 |
| MTC7 | YEL033W | 122.438587 | -1.204002 | 0.16870566 | -7.1367018 | 9.56E-13 |
| FLO9 | YAL063C | 86.4423057 | 1.29457537 | 0.18149303 | 7.13292059 | 9.83E-13 |

| | | | | | | |
|---|---|---|---|---|---|---|
| RPL15B | YMR121C | 674.073959 | -0.6430442 | 0.09017681 | -7.1309266 | 9.97E-13 |
| RLF2 | YPR018W | 973.77065 | 0.48948329 | 0.06869716 | 7.12523293 | 1.04E-12 |
| CAP1 | YKL007W | 1131.38625 | -0.5089438 | 0.07145943 | -7.1221354 | 1.06E-12 |
| IDH2 | YOR136W | 1658.51431 | -0.4743257 | 0.06667919 | -7.1135486 | 1.13E-12 |
| SOK2 | YMR016C | 2026.68629 | 0.49573661 | 0.06970415 | 7.11200944 | 1.14E-12 |
| ERG25 | YGR060W | 2795.18483 | -0.4468828 | 0.0628406 | -7.1113709 | 1.15E-12 |
| SNR82 | snR82 | 3733.70834 | -0.6301132 | 0.08869879 | -7.1039659 | 1.21E-12 |
| YMR102C | YMR102C | 812.693505 | 0.56058146 | 0.07890959 | 7.10409802 | 1.21E-12 |
| YGP1 | YNL160W | 12531.1601 | 0.37776595 | 0.0531746 | 7.10425504 | 1.21E-12 |
| Xu_2009_CUTs22 | Xu_2009_CU | 64.6090383 | 1.36963379 | 0.1928074 | 7.10363714 | 1.22E-12 |
| MRPL15 | YLR312W-A | 486.342568 | -0.6133636 | 0.08636891 | -7.1016713 | 1.23E-12 |
| ALG9 | YNL219C | 1169.65321 | -0.4907948 | 0.06912629 | -7.0999731 | 1.25E-12 |
| YMR099C | YMR099C | 1454.14719 | -0.4868497 | 0.06858808 | -7.0981685 | 1.26E-12 |
| ICP55 | YER078C | 396.123857 | -0.6918834 | 0.0974795 | -7.0977324 | 1.27E-12 |
| LHP1 | YDL051W | 1482.92128 | -0.5174887 | 0.07298021 | -7.0908092 | 1.33E-12 |
| AGA1 | YNR044W | 595.205401 | 0.63589392 | 0.08970881 | 7.08842239 | 1.36E-12 |
| Xu_2009_SUTs21 | Xu_2009_SU | 686.339386 | 0.58549237 | 0.08264321 | 7.0845792 | 1.39E-12 |
| RPO21 | YDL140C | 7425.9574 | 0.38941341 | 0.05500225 | 7.07995414 | 1.44E-12 |
| YPR063C | YPR063C | 1149.68067 | -0.5397419 | 0.07625658 | -7.0779711 | 1.46E-12 |
| Xu_2009_SUTs76 | Xu_2009_SU | 102.896764 | 1.15729244 | 0.16369851 | 7.0696578 | 1.55E-12 |
| RIF1 | YBR275C | 1443.60349 | 0.49113568 | 0.06949717 | 7.066988 | 1.58E-12 |
| MAP2 | YBL091C | 1479.00442 | -0.4990865 | 0.070629 | -7.066311 | 1.59E-12 |
| SEC28 | YIL076W | 1177.36271 | -0.4650816 | 0.06581828 | -7.0661461 | 1.59E-12 |
| YKL066W | YKL066W | 121.334727 | -1.0664923 | 0.15096482 | -7.0645086 | 1.61E-12 |
| Xu_2009_CUTs90 | Xu_2009_CU | 84.6577042 | 1.21628713 | 0.17220807 | 7.06289277 | 1.63E-12 |
| SPP41 | YDR464W | 1694.46087 | 0.43285141 | 0.06130274 | 7.06088194 | 1.65E-12 |
| ARC18 | YLR370C | 789.467485 | -0.5774874 | 0.08179932 | -7.0598068 | 1.67E-12 |
| APE2 | YKL157W | 2720.84423 | -0.3996893 | 0.05662201 | -7.058904 | 1.68E-12 |
| SNR40 | snR40 | 160.97795 | -0.993926 | 0.14087095 | -7.0555784 | 1.72E-12 |
| PET123 | YOR158W | 692.33575 | -0.5443057 | 0.07716049 | -7.0542025 | 1.74E-12 |
| CLU1 | YMR012W | 7069.36222 | -0.343097 | 0.04866071 | -7.0508011 | 1.78E-12 |
| EMP47 | YFL048C | 1026.38118 | -0.5406605 | 0.07685744 | -7.0345887 | 2.00E-12 |
| SYF2 | YGR129W | 455.49027 | 0.65451315 | 0.0932094 | 7.02196465 | 2.19E-12 |
| EPT1 | YHR123W | 754.401508 | -0.5518005 | 0.07859502 | -7.0208069 | 2.21E-12 |
| MAS2 | YHR024C | 897.049647 | -0.5202058 | 0.07411546 | -7.0188567 | 2.24E-12 |
| PDR15 | YDR406W | 926.088405 | 0.57146007 | 0.08143325 | 7.01752753 | 2.26E-12 |
| ATP18 | YML081C-A | 956.61715 | -0.523558 | 0.07473102 | -7.0058997 | 2.45E-12 |
| ERG2 | YMR202W | 1461.41588 | -0.4240008 | 0.0605226 | -7.0056613 | 2.46E-12 |
| EGT2 | YNL327W | 4826.91959 | -0.36327 | 0.05188433 | -7.001536 | 2.53E-12 |
| PMA1 | YGL008C | 38982.3187 | -0.3745232 | 0.05350657 | -6.9995735 | 2.57E-12 |
| COX14 | YML129C | 402.266146 | -0.7126865 | 0.1018202 | -6.999461 | 2.57E-12 |

| | | | | | | |
|---|---|---|---|---|---|---|
| VMA6 | YLR447C | 2392.56493 | -0.4273258 | 0.06119191 | -6.9833712 | 2.88E-12 |
| VHR1 | YIL056W | 1667.05588 | 0.48607092 | 0.06968068 | 6.97569127 | 3.04E-12 |
| FRD1 | YEL047C | 1231.28802 | -0.4675206 | 0.06703103 | -6.9746893 | 3.07E-12 |
| ETR1 | YBR026C | 1185.26024 | -0.5157241 | 0.07406747 | -6.9628963 | 3.33E-12 |
| GIC1 | YHR061C | 402.594142 | -0.6462226 | 0.09281235 | -6.9626795 | 3.34E-12 |
| MDM35 | YKL053C-A | 152.056656 | -0.9994176 | 0.14366163 | -6.9567471 | 3.48E-12 |
| AFG1 | YEL052W | 776.387666 | -0.6035145 | 0.08683637 | -6.9500197 | 3.65E-12 |
| SNR48 | snR48 | 181.108099 | 0.9271834 | 0.13366166 | 6.93679383 | 4.01E-12 |
| SKN1 | YGR143W | 1658.73583 | 0.43640984 | 0.062965 | 6.93099136 | 4.18E-12 |
| ARO80 | YDR421W | 664.322847 | 0.57874201 | 0.08356311 | 6.9258072 | 4.33E-12 |
| NOP1 | YDL014W | 2520.72278 | -0.5599192 | 0.08088343 | -6.9225457 | 4.44E-12 |
| MRP10 | YDL045W-A | 447.405733 | -0.6565577 | 0.09485271 | -6.9218661 | 4.46E-12 |
| DDR48 | YMR173W | 1629.01285 | -0.5252665 | 0.07589982 | -6.9205234 | 4.50E-12 |
| COA3 | YJL062W-A | 458.184534 | -0.647524 | 0.09362157 | -6.9163975 | 4.63E-12 |
| Xu_2009_SUTs42 | Xu_2009_SU | 193.028749 | 0.84642711 | 0.12241819 | 6.91422674 | 4.70E-12 |
| HEM15 | YOR176W | 1117.23833 | -0.4535901 | 0.06565324 | -6.9088764 | 4.89E-12 |
| RPN8 | YOR261C | 1315.27253 | -0.463127 | 0.06705677 | -6.9064908 | 4.97E-12 |
| BNI1 | YNL271C | 3223.31032 | 0.42281842 | 0.06122795 | 6.90564383 | 5.00E-12 |
| YNR021W | YNR021W | 920.161643 | -0.5326283 | 0.07713826 | -6.9048518 | 5.03E-12 |
| AIM24 | YJR080C | 653.617834 | -0.6156072 | 0.08917915 | -6.9030392 | 5.09E-12 |
| GSP2 | YOR185C | 459.987971 | -0.6758463 | 0.09797226 | -6.8983428 | 5.26E-12 |
| YNL108C | YNL108C | 482.60969 | -0.6228042 | 0.09030416 | -6.8967389 | 5.32E-12 |
| PAM17 | YKR065C | 586.510869 | -0.5581276 | 0.08096788 | -6.8931977 | 5.46E-12 |
| YMR173W-A | YMR173W-A | 91.0246373 | -1.202413 | 0.17460359 | -6.8865308 | 5.72E-12 |
| VMA1 | YDL185W | 10822.111 | -0.3142948 | 0.04564949 | -6.8849576 | 5.78E-12 |
| YHL050C | YHL050C | 228.313847 | 0.90117228 | 0.1309461 | 6.88200945 | 5.90E-12 |
| SNR74 | snR74 | 89.8155418 | -1.2249596 | 0.17811836 | -6.8772227 | 6.10E-12 |
| CLN2 | YPL256C | 2280.71689 | -0.4054688 | 0.05896575 | -6.8763446 | 6.14E-12 |
| GLN1 | YPR035W | 12563.7772 | -0.4762755 | 0.06932308 | -6.8703738 | 6.40E-12 |
| NOP4 | YPL043W | 3006.92034 | 0.53452477 | 0.07781944 | 6.86878228 | 6.48E-12 |
| YDR544C | YDR544C | 45.1841705 | 1.60943453 | 0.2343244 | 6.86840354 | 6.49E-12 |
| ECM29 | YHL030W | 2767.74221 | 0.3861191 | 0.05626075 | 6.86302739 | 6.74E-12 |
| UBC9 | YDL064W | 514.696953 | -0.6133328 | 0.08940442 | -6.8602061 | 6.88E-12 |
| MRX3 | YBL095W | 318.565143 | -0.6982009 | 0.10183111 | -6.8564596 | 7.06E-12 |
| FRQ1 | YDR373W | 538.944724 | -0.5713232 | 0.08333537 | -6.855711 | 7.10E-12 |
| SIS1 | YNL007C | 3187.73379 | -0.492302 | 0.07188421 | -6.8485408 | 7.46E-12 |
| CMC4 | YMR194C-B | 95.4359792 | -1.1525621 | 0.16840705 | -6.8439062 | 7.71E-12 |
| Xu_2009_SUTs56 | Xu_2009_SU | 392.084119 | 0.71256037 | 0.10420225 | 6.83824327 | 8.02E-12 |
| PIK1 | YNL267W | 1676.66566 | 0.4541376 | 0.06643482 | 6.83583679 | 8.15E-12 |
| Xu_2009_SUTs56 | Xu_2009_SU | 204.816737 | 0.91632074 | 0.13405002 | 6.83566304 | 8.16E-12 |
| Xu_2009_CUTs64 | Xu_2009_CU | 76.6495249 | 1.24248882 | 0.18177599 | 6.83527462 | 8.18E-12 |

| | | | | | | |
|---|---|---|---|---|---|---|
| SER1 | YOR184W | 1218.7338 | -0.5190621 | 0.07594169 | -6.8350086 | 8.20E-12 |
| CTR1 | YPR124W | 263.547665 | -0.7815361 | 0.11450295 | -6.8254667 | 8.76E-12 |
| FUM1 | YPL262W | 2314.22636 | -0.3685781 | 0.05402859 | -6.8219093 | 8.98E-12 |
| CSR1 | YLR380W | 1412.38717 | -0.4567483 | 0.0669656 | -6.8206408 | 9.06E-12 |
| SLT2 | YHR030C | 2402.9803 | 0.42943215 | 0.06312498 | 6.80288802 | 1.03E-11 |
| AVL9 | YLR114C | 669.273416 | 0.60920375 | 0.08955226 | 6.80277343 | 1.03E-11 |
| Xu_2009_SUTs6 | Xu_2009_SU | 238.254034 | 0.76144865 | 0.11204027 | 6.79620493 | 1.07E-11 |
| VMA13 | YPR036W | 2274.28705 | -0.4884658 | 0.07188355 | -6.795238 | 1.08E-11 |
| APT1 | YML022W | 494.883749 | -0.7135169 | 0.10504627 | -6.7924053 | 1.10E-11 |
| AKR2 | YOR034C | 621.36077 | 0.54151528 | 0.07988111 | 6.77901551 | 1.21E-11 |
| RIB3 | YDR487C | 1745.79821 | -0.4744873 | 0.07003293 | -6.7752024 | 1.24E-11 |
| ARG81 | YML099C | 1260.00874 | 0.43194136 | 0.06376849 | 6.77358618 | 1.26E-11 |
| TAO3 | YIL129C | 2822.63992 | 0.36925338 | 0.05451615 | 6.77328465 | 1.26E-11 |
| Xu_2009_CUTs48 | Xu_2009_CU | 67.9079326 | 1.32014231 | 0.19493501 | 6.77221762 | 1.27E-11 |
| VPS1 | YKR001C | 2317.75284 | -0.4027826 | 0.05949878 | -6.7695954 | 1.29E-11 |
| DCS1 | YLR270W | 918.86596 | -0.6529731 | 0.09648746 | -6.7674401 | 1.31E-11 |
| YER079W | YER079W | 419.667593 | -0.6610234 | 0.09777943 | -6.7603528 | 1.38E-11 |
| RGM1 | YMR182C | 227.190216 | 0.81710011 | 0.12087973 | 6.75961228 | 1.38E-11 |
| SNF2 | YOR290C | 2459.83594 | 0.41072863 | 0.06083481 | 6.75153948 | 1.46E-11 |
| NHP2 | YDL208W | 1644.82265 | -0.5492984 | 0.08136307 | -6.7512006 | 1.47E-11 |
| FTH1 | YBR207W | 1150.33841 | -0.4452212 | 0.06596191 | -6.749671 | 1.48E-11 |
| PEP1 | YBL017C | 4395.29571 | 0.3294099 | 0.04886995 | 6.74054169 | 1.58E-11 |
| CBF2 | YGR140W | 1042.64563 | 0.48691291 | 0.07223845 | 6.74035681 | 1.58E-11 |
| LEU3 | YLR451W | 864.846809 | 0.47600751 | 0.0706679 | 6.73583766 | 1.63E-11 |
| URC2 | YDR520C | 577.225768 | 0.55243153 | 0.08205385 | 6.73254895 | 1.67E-11 |
| SMC2 | YFR031C | 1210.40894 | 0.45143975 | 0.06707256 | 6.73061741 | 1.69E-11 |
| MMS2 | YGL087C | 1074.13286 | -0.4442582 | 0.06604128 | -6.7269781 | 1.73E-11 |
| RPT6 | YGL048C | 1756.16426 | -0.4166619 | 0.06197865 | -6.7226682 | 1.78E-11 |
| PNP1 | YLR209C | 429.118293 | -0.6481176 | 0.09649078 | -6.7168861 | 1.86E-11 |
| SMX2 | YFL017W-A | 159.576986 | -0.9622195 | 0.14333958 | -6.7128665 | 1.91E-11 |
| PMU1 | YKL128C | 970.449942 | -0.5334569 | 0.07947292 | -6.7124354 | 1.91E-11 |
| PCS60 | YBR222C | 1152.85493 | -0.4833294 | 0.07201029 | -6.7119486 | 1.92E-11 |
| HEM2 | YGL040C | 1013.31478 | -0.4518641 | 0.06741479 | -6.7027438 | 2.05E-11 |
| Xu_2009_CUTs59 | Xu_2009_CU | 78.6534091 | 1.20887652 | 0.18038471 | 6.70165743 | 2.06E-11 |
| SLN1 | YIL147C | 1496.03603 | 0.45125602 | 0.06733794 | 6.70136403 | 2.06E-11 |
| MIN8 | YPR010C-A | 787.020494 | -0.6263158 | 0.09353087 | -6.6963535 | 2.14E-11 |
| ECM4 | YKR076W | 371.09496 | -0.6974112 | 0.10419141 | -6.6935579 | 2.18E-11 |
| HOC1 | YJR075W | 1309.55923 | -0.4209675 | 0.06294773 | -6.6875719 | 2.27E-11 |
| TIF35 | YDR429C | 1916.89729 | -0.4493117 | 0.06719305 | -6.6868783 | 2.28E-11 |
| ALE1 | YOR175C | 1351.15263 | -0.4506331 | 0.06740773 | -6.6851835 | 2.31E-11 |
| Xu_2009_SUTs53 | Xu_2009_SU | 145.631741 | 0.97210659 | 0.14545346 | 6.68328254 | 2.34E-11 |

| | | | | | | |
|---|---|---|---|---|---|---|
| Xu_2009_SUTs36 | Xu_2009_SU | 101.241006 | 1.10437177 | 0.16534879 | 6.67904363 | 2.41E-11 |
| FLO11 | YIR019C | 184.003316 | 0.92526659 | 0.13866352 | 6.67274676 | 2.51E-11 |
| ADH5 | YBR145W | 174.481866 | -0.9327475 | 0.13979509 | -6.6722482 | 2.52E-11 |
| SAD1 | YFR005C | 351.196653 | -0.6731021 | 0.10112959 | -6.6558369 | 2.82E-11 |
| ERV29 | YGR284C | 2000.9314 | -0.4183768 | 0.06288007 | -6.6535678 | 2.86E-11 |
| CMK1 | YFR014C | 431.08277 | -0.6317806 | 0.09496318 | -6.6529009 | 2.87E-11 |
| NAB2 | YGL122C | 3695.62665 | 0.47213506 | 0.07098643 | 6.65106056 | 2.91E-11 |
| AIM2 | YAL049C | 766.18343 | -0.5364881 | 0.08067023 | -6.6503853 | 2.92E-11 |
| Xu_2009_SUTs27 | Xu_2009_SU | 322.67488 | 0.73382541 | 0.11042292 | 6.64558981 | 3.02E-11 |
| MSC1 | YML128C | 2411.23284 | -0.5304763 | 0.0798233 | -6.6456322 | 3.02E-11 |
| MRP17 | YKL003C | 516.979325 | -0.5728992 | 0.08621239 | -6.6452078 | 3.03E-11 |
| YOR343W-B | YOR343W-B | 768.529849 | 0.52151359 | 0.07848012 | 6.64516799 | 3.03E-11 |
| Xu_2009_SUTs45 | Xu_2009_SU | 361.322448 | 0.64814716 | 0.09754232 | 6.64477908 | 3.04E-11 |
| CDC39 | YCR093W | 6189.93668 | 0.33535463 | 0.05049083 | 6.64189181 | 3.10E-11 |
| FAP1 | YNL023C | 836.217964 | 0.54061026 | 0.08146267 | 6.63629442 | 3.22E-11 |
| KNS1 | YLL019C | 2224.36625 | 0.37001218 | 0.05576474 | 6.63523579 | 3.24E-11 |
| YSC83 | YHR017W | 363.784904 | -0.6508619 | 0.09809551 | -6.6349816 | 3.25E-11 |
| NRT1 | YOR071C | 555.093269 | -0.5541178 | 0.0835202 | -6.634536 | 3.26E-11 |
| LPD1 | YFL018C | 2445.64701 | -0.4511775 | 0.06802589 | -6.6324378 | 3.30E-11 |
| YAH1 | YPL252C | 415.679435 | -0.6370275 | 0.09615398 | -6.625077 | 3.47E-11 |
| RAD34 | YDR314C | 379.454136 | 0.6898477 | 0.10421842 | 6.61924892 | 3.61E-11 |
| APE4 | YHR113W | 819.705652 | -0.4978921 | 0.07551582 | -6.5932152 | 4.30E-11 |
| NPP1 | YCR026C | 1432.18677 | 0.45692024 | 0.06940552 | 6.58334193 | 4.60E-11 |
| ECM16 | YMR128W | 1944.30109 | 0.45774046 | 0.0695693 | 6.57963323 | 4.72E-11 |
| MIC10 | YCL057C-A | 516.51969 | -0.58158 | 0.08849851 | -6.571636 | 4.98E-11 |
| tI(UAU)L | tI(UAU)L | 29.8473433 | -1.7321084 | 0.26360836 | -6.5707643 | 5.01E-11 |
| Xu_2009_CUTs85 | Xu_2009_CU | 75.4690111 | 1.21670177 | 0.1852796 | 6.56684138 | 5.14E-11 |
| MRD1 | YPR112C | 2507.31633 | 0.46818402 | 0.07131736 | 6.56479746 | 5.21E-11 |
| MRPL9 | YGR220C | 541.299713 | -0.5341192 | 0.08136958 | -6.5641138 | 5.23E-11 |
| RPR1 | RPR1 | 25867.4288 | -0.9902489 | 0.15103785 | -6.5562965 | 5.52E-11 |
| NIP100 | YPL174C | 735.712774 | 0.5109138 | 0.07804854 | 6.54610321 | 5.91E-11 |
| GCN3 | YKR026C | 779.760398 | -0.4698664 | 0.07178334 | -6.5456184 | 5.92E-11 |
| YOL057W | YOL057W | 1421.03242 | -0.3890414 | 0.0594716 | -6.5416337 | 6.09E-11 |
| VOA1 | YGR106C | 1742.90702 | -0.4257746 | 0.06510019 | -6.5402979 | 6.14E-11 |
| HEF3 | YNL014W | 326.575309 | 0.68931639 | 0.10542533 | 6.53843234 | 6.22E-11 |
| Xu_2009_SUTs56 | Xu_2009_SU | 77.9526103 | 1.18926567 | 0.18189665 | 6.53813965 | 6.23E-11 |
| RPT5 | YOR117W | 1726.41555 | -0.4690703 | 0.0717542 | -6.5371817 | 6.27E-11 |
| SSK2 | YNR031C | 1801.18683 | 0.40737063 | 0.06235296 | 6.53330074 | 6.43E-11 |
| BAG7 | YOR134W | 1099.1558 | 0.50517798 | 0.07745933 | 6.52184779 | 6.94E-11 |
| LIA1 | YJR070C | 1922.33661 | -0.5221024 | 0.08006812 | -6.5207273 | 7.00E-11 |
| Xu_2009_SUTs13 | Xu_2009_SU | 86.4631593 | 1.0965002 | 0.16819305 | 6.51929536 | 7.06E-11 |

| | | | | | | |
|---|---|---|---|---|---|---|
| SNA3 | YJL151C | 1734.79419 | -0.4370292 | 0.06705464 | -6.5175077 | 7.15E-11 |
| MSB4 | YOL112W | 787.609293 | 0.49857872 | 0.0766667 | 6.50319811 | 7.86E-11 |
| CUE1 | YMR264W | 580.852484 | -0.5532535 | 0.08510915 | -6.5005169 | 8.00E-11 |
| Xu_2009_CUTs91 | Xu_2009_CU | 102.825013 | 1.12769039 | 0.17349175 | 6.49996543 | 8.03E-11 |
| NRG1 | YDR043C | 834.627353 | 0.51152365 | 0.07871687 | 6.49827238 | 8.12E-11 |
| Xu_2009_CUTs34 | Xu_2009_CU | 103.105227 | 1.07031589 | 0.16475988 | 6.49621671 | 8.24E-11 |
| YMR226C | YMR226C | 2624.95548 | -0.4538652 | 0.06988404 | -6.4945465 | 8.33E-11 |
| Xu_2009_SUTs35 | Xu_2009_SU | 492.090064 | 0.61369301 | 0.09457627 | 6.48886895 | 8.65E-11 |
| Xu_2009_SUTs25 | Xu_2009_SU | 97.354849 | 1.08481363 | 0.16733508 | 6.48288216 | 9.00E-11 |
| SDA1 | YGR245C | 2323.98386 | 0.51242629 | 0.07904859 | 6.48242151 | 9.03E-11 |
| MDY2 | YOL111C | 467.204483 | -0.6069003 | 0.09366885 | -6.4792108 | 9.22E-11 |
| LSM5 | YER146W | 441.290153 | -0.6473297 | 0.10004128 | -6.4706261 | 9.76E-11 |
| MZM1 | YDR493W | 216.375463 | -0.8104833 | 0.12527528 | -6.4696192 | 9.83E-11 |
| INO80 | YGL150C | 2675.02742 | 0.36495968 | 0.05642054 | 6.46856043 | 9.89E-11 |
| RIM101 | YHL027W | 2095.83831 | 0.38933734 | 0.0601988 | 6.46752651 | 9.96E-11 |
| tV(AAC)O | tV(AAC)O | 48.9986387 | -1.4372691 | 0.22249196 | -6.4598697 | 1.05E-10 |
| ATP5 | YDR298C | 1552.17623 | -0.4200944 | 0.06509134 | -6.4539214 | 1.09E-10 |
| YET3 | YDL072C | 1135.80411 | -0.5332326 | 0.08263948 | -6.4525155 | 1.10E-10 |
| OST4 | YDL232W | 395.117154 | -0.6851682 | 0.10626239 | -6.4478903 | 1.13E-10 |
| TIF34 | YMR146C | 3131.32307 | -0.3870737 | 0.06003642 | -6.4473148 | 1.14E-10 |
| Xu_2009_CUTs73 | Xu_2009_CU | 39.8637266 | 1.48128372 | 0.22986683 | 6.44409508 | 1.16E-10 |
| Xu_2009_SUTs59 | Xu_2009_SU | 159.400321 | -0.8964344 | 0.13914008 | -6.4426758 | 1.17E-10 |
| RTN1 | YDR233C | 4466.24787 | -0.4006725 | 0.06220999 | -6.4406457 | 1.19E-10 |
| ILV2 | YMR108W | 4264.73729 | -0.3174143 | 0.04929598 | -6.4389479 | 1.20E-10 |
| GUP1 | YGL084C | 733.492447 | -0.5186631 | 0.08056383 | -6.4379147 | 1.21E-10 |
| XBP1 | YIL101C | 864.907058 | 0.47003842 | 0.07304275 | 6.43511383 | 1.23E-10 |
| CDC5 | YMR001C | 1345.78009 | 0.44485412 | 0.06914763 | 6.4333966 | 1.25E-10 |
| REV1 | YOR346W | 993.314729 | 0.48518018 | 0.07541847 | 6.43317489 | 1.25E-10 |
| VCX1 | YDL128W | 1523.29626 | -0.4334385 | 0.06739739 | -6.4310878 | 1.27E-10 |
| LAG1 | YHL003C | 893.087479 | -0.514779 | 0.08015817 | -6.4220396 | 1.34E-10 |
| TAF7 | YMR227C | 306.16938 | 0.69640284 | 0.10844183 | 6.42190207 | 1.35E-10 |
| TVP38 | YKR088C | 356.280538 | -0.645502 | 0.10055464 | -6.419415 | 1.37E-10 |
| DOP1 | YDR141C | 2312.63449 | 0.40731994 | 0.0634582 | 6.41871248 | 1.37E-10 |
| AIM44 | YPL158C | 1129.3682 | -0.449337 | 0.07002898 | -6.4164444 | 1.39E-10 |
| DAK1 | YML070W | 2047.86421 | -0.428981 | 0.06689299 | -6.4129436 | 1.43E-10 |
| SPT10 | YJL127C | 715.45656 | 0.47867266 | 0.07466202 | 6.41119326 | 1.44E-10 |
| MAP1 | YLR244C | 2555.6517 | -0.3612374 | 0.05635495 | -6.410038 | 1.45E-10 |
| CPR4 | YCR069W | 1316.71693 | -0.4239135 | 0.06614514 | -6.4088376 | 1.47E-10 |
| TMA20 | YER007C-A | 713.422649 | -0.5189027 | 0.08100666 | -6.4056799 | 1.50E-10 |
| HOS4 | YIL112W | 1063.90725 | 0.47794063 | 0.0746466 | 6.40271169 | 1.53E-10 |
| MPC2 | YHR162W | 1465.31214 | -0.4513139 | 0.07070309 | -6.3832268 | 1.73E-10 |

| | | | | | | |
|---|---|---|---|---|---|---|
| Xu_2009_SUTs13 | Xu_2009_SU | 244.722195 | 0.72701763 | 0.11390835 | 6.38247906 | 1.74E-10 |
| BCD1 | YHR040W | 472.033519 | 0.61266863 | 0.09599579 | 6.38224504 | 1.75E-10 |
| Xu_2009_CUTs31 | Xu_2009_CU | 83.424976 | 1.17375627 | 0.18394091 | 6.38115934 | 1.76E-10 |
| MTD1 | YKR080W | 1163.65729 | -0.4478715 | 0.07026155 | -6.3743474 | 1.84E-10 |
| KAP120 | YPL125W | 2045.60968 | 0.36590429 | 0.05742767 | 6.3715673 | 1.87E-10 |
| Xu_2009_CUTs45 | Xu_2009_CU | 100.108712 | 1.02063532 | 0.16019738 | 6.37111139 | 1.88E-10 |
| URA4 | YLR420W | 1027.83785 | -0.4840856 | 0.07598182 | -6.3710715 | 1.88E-10 |
| NUS1 | YDL193W | 369.795192 | -0.6385441 | 0.10025497 | -6.3692019 | 1.90E-10 |
| HBN1 | YCL026C-B | 295.844905 | -0.8050238 | 0.12641665 | -6.3680204 | 1.91E-10 |
| YKL050C | YKL050C | 409.439594 | 0.65123101 | 0.10226573 | 6.36802777 | 1.91E-10 |
| UBP5 | YER144C | 1540.00027 | 0.42554622 | 0.06686018 | 6.36471814 | 1.96E-10 |
| CBC2 | YPL178W | 565.202072 | -0.6071981 | 0.09546808 | -6.3602205 | 2.01E-10 |
| MCA1 | YOR197W | 2428.53385 | -0.363685 | 0.0572026 | -6.3578411 | 2.05E-10 |
| ERV46 | YAL042W | 2119.41902 | -0.4262619 | 0.06707439 | -6.3550614 | 2.08E-10 |
| CAF16 | YFL028C | 473.594681 | -0.5717703 | 0.08998974 | -6.3537281 | 2.10E-10 |
| TSC3 | YBR058C-A | 330.542063 | -0.6465465 | 0.10176543 | -6.3533019 | 2.11E-10 |
| AIM7 | YDR063W | 446.445716 | -0.5707757 | 0.08984479 | -6.3529076 | 2.11E-10 |
| OCH1 | YGL038C | 1505.6348 | -0.3754517 | 0.05912447 | -6.3501904 | 2.15E-10 |
| MMS1 | YPR164W | 1762.09988 | 0.41793766 | 0.06585307 | 6.34651727 | 2.20E-10 |
| TOS1 | YBR162C | 3733.89213 | -0.4047131 | 0.06381036 | -6.3424357 | 2.26E-10 |
| YBR053C | YBR053C | 1072.57053 | -0.4859685 | 0.07664408 | -6.3405869 | 2.29E-10 |
| YGR153W | YGR153W | 405.256166 | 0.63067571 | 0.09949709 | 6.33863466 | 2.32E-10 |
| RAD54 | YGL163C | 776.793122 | 0.4691117 | 0.07403609 | 6.33625674 | 2.35E-10 |
| MDS3 | YGL197W | 2559.5217 | 0.35393109 | 0.05589609 | 6.33194767 | 2.42E-10 |
| BUD22 | YMR014W | 1181.22299 | 0.50185664 | 0.07925887 | 6.33186745 | 2.42E-10 |
| Xu_2009_SUTs25 | Xu_2009_SU | 138.878561 | 0.93763516 | 0.14823098 | 6.3255008 | 2.52E-10 |
| EPO1 | YMR124W | 1582.41488 | 0.38491323 | 0.06089665 | 6.32076225 | 2.60E-10 |
| YGR027W-B | YGR027W-B | 1093.46601 | 0.4108007 | 0.06501388 | 6.318661 | 2.64E-10 |
| SSE1 | YPL106C | 19497.9689 | -0.3001161 | 0.04751135 | -6.316724 | 2.67E-10 |
| SUI3 | YPL237W | 1652.65657 | -0.4592663 | 0.07272588 | -6.3150325 | 2.70E-10 |
| STF1 | YDL130W-A | 1624.58028 | -0.4198674 | 0.06649993 | -6.3138027 | 2.72E-10 |
| PRE8 | YML092C | 991.031712 | -0.4688961 | 0.07431737 | -6.309374 | 2.80E-10 |
| Xu_2009_CUTs37 | Xu_2009_CU | 108.915527 | 0.98274882 | 0.15581539 | 6.30713573 | 2.84E-10 |
| CCT8 | YJL008C | 2240.98133 | -0.3881367 | 0.0615551 | -6.3055161 | 2.87E-10 |
| RFA2 | YNL312W | 419.801651 | -0.5948925 | 0.09436026 | -6.3044821 | 2.89E-10 |
| TUB3 | YML124C | 2186.62465 | -0.36142 | 0.05739711 | -6.2968323 | 3.04E-10 |
| Xu_2009_CUTs90 | Xu_2009_CU | 81.8338206 | 1.13743755 | 0.1806461 | 6.29649675 | 3.04E-10 |
| GGA2 | YHR108W | 1870.16489 | -0.3905307 | 0.06203491 | -6.2953381 | 3.07E-10 |
| SIL1 | YOL031C | 597.097244 | -0.534171 | 0.08486108 | -6.2946527 | 3.08E-10 |
| IMP2 | YMR035W | 140.718902 | -0.9229913 | 0.14665506 | -6.2936205 | 3.10E-10 |
| MIT1 | YEL007W | 836.978965 | -0.5421838 | 0.08618539 | -6.2909012 | 3.16E-10 |

| | | | | | | |
|---|---|---|---|---|---|---|
| YPT7 | YML001W | 414.231208 | -0.6283179 | 0.0998923 | -6.2899533 | 3.18E-10 |
| MET22 | YOL064C | 433.424211 | -0.6430745 | 0.10232522 | -6.2846145 | 3.29E-10 |
| MAM33 | YIL070C | 600.238543 | -0.576532 | 0.0917435 | -6.2841729 | 3.30E-10 |
| REE1 | YJL217W | 374.226782 | -0.7227319 | 0.11501811 | -6.2836353 | 3.31E-10 |
| LIP1 | YMR298W | 744.938643 | -0.509873 | 0.08115522 | -6.2826887 | 3.33E-10 |
| PMP2 | YEL017C-A | 435.254283 | -0.655219 | 0.10434602 | -6.2792911 | 3.40E-10 |
| PLB2 | YMR006C | 566.630321 | -0.5177389 | 0.08250953 | -6.274898 | 3.50E-10 |
| Xu_2009_CUTs18 | Xu_2009_CU | 79.316628 | 1.17047473 | 0.18658639 | 6.27309806 | 3.54E-10 |
| YJR096W | YJR096W | 945.125524 | -0.4896774 | 0.07807441 | -6.271932 | 3.57E-10 |
| Xu_2009_SUTs80 | Xu_2009_SU | 116.372095 | 0.96119678 | 0.15330492 | 6.26983641 | 3.61E-10 |
| YLR255C | YLR255C | 86.9958669 | 1.05827747 | 0.16881173 | 6.26898064 | 3.63E-10 |
| DEP1 | YAL013W | 928.644904 | 0.48017634 | 0.07660393 | 6.26829896 | 3.65E-10 |
| YNL122C | YNL122C | 198.163016 | -0.7911868 | 0.12622906 | -6.2678658 | 3.66E-10 |
| SCS22 | YBL091C-A | 114.644977 | -0.992191 | 0.15832564 | -6.2667736 | 3.69E-10 |
| URH1 | YDR400W | 439.290258 | -0.5747678 | 0.09191939 | -6.2529544 | 4.03E-10 |
| LCB3 | YJL134W | 373.972873 | -0.6021304 | 0.09635076 | -6.2493575 | 4.12E-10 |
| DAN4 | YJR151C | 231.82417 | 0.76495207 | 0.12253232 | 6.24285954 | 4.30E-10 |
| ENT1 | YDL161W | 1660.68345 | 0.41806671 | 0.06697044 | 6.24255569 | 4.30E-10 |
| ARC40 | YBR234C | 1821.23956 | -0.3981149 | 0.06382336 | -6.2377619 | 4.44E-10 |
| TOA1 | YOR194C | 553.697841 | 0.58745107 | 0.0942052 | 6.23586688 | 4.49E-10 |
| VMA9 | YCL005W-A | 1169.47245 | -0.4636655 | 0.07437228 | -6.2343856 | 4.54E-10 |
| Xu_2009_CUTs53 | Xu_2009_CU | 60.7175351 | 1.21656093 | 0.19521169 | 6.23200867 | 4.60E-10 |
| YAL064W-B | YAL064W-B | 48.2807294 | 1.35772753 | 0.21811283 | 6.22488611 | 4.82E-10 |
| DBP10 | YDL031W | 2608.50649 | 0.40736186 | 0.06544841 | 6.22416711 | 4.84E-10 |
| RPT3 | YDR394W | 2935.53511 | -0.3235573 | 0.05198757 | -6.2237434 | 4.85E-10 |
| Xu_2009_CUTs56 | Xu_2009_CU | 47.1959756 | 1.33625898 | 0.21476607 | 6.22192785 | 4.91E-10 |
| Xu_2009_SUTs68 | Xu_2009_SU | 114.94858 | 0.99319335 | 0.15970003 | 6.21911808 | 5.00E-10 |
| ATF2 | YGR177C | 689.512268 | 0.54045512 | 0.08705132 | 6.2084655 | 5.35E-10 |
| SCEI | Q0160 | 94.8608591 | -1.0752197 | 0.17320642 | -6.2077355 | 5.38E-10 |
| PKC1 | YBL105C | 2272.46309 | 0.38521845 | 0.06207377 | 6.20581716 | 5.44E-10 |
| FSF1 | YOR271C | 1408.16274 | -0.4504683 | 0.0726561 | -6.2000058 | 5.65E-10 |
| YER010C | YER010C | 432.462664 | -0.6459264 | 0.10423911 | -6.1965837 | 5.77E-10 |
| SMF3 | YLR034C | 1060.20639 | -0.4975205 | 0.08035254 | -6.1917211 | 5.95E-10 |
| TPN1 | YGL186C | 756.056885 | -0.4783688 | 0.07730845 | -6.1877949 | 6.10E-10 |
| GPI12 | YMR281W | 310.240551 | -0.6356785 | 0.10274564 | -6.1869139 | 6.14E-10 |
| TMA7 | YLR262C-A | 644.180725 | -0.6029569 | 0.0974922 | -6.1846684 | 6.22E-10 |
| AVO1 | YOL078W | 831.632994 | 0.45650655 | 0.07386609 | 6.18019087 | 6.40E-10 |
| SKP1 | YDR328C | 1339.1937 | -0.3975965 | 0.06436272 | -6.1774337 | 6.52E-10 |
| MKT1 | YNL085W | 4282.68477 | -0.3095933 | 0.05015756 | -6.1724158 | 6.73E-10 |
| TOS4 | YLR183C | 352.00578 | -0.6281411 | 0.10177955 | -6.1715848 | 6.76E-10 |
| SNR37 | snR37 | 29234.3026 | -1.0476915 | 0.16985903 | -6.1680063 | 6.92E-10 |

| | | | | | | |
|---|---|---|---|---|---|---|
| ARD1 | YHR013C | 730.966399 | -0.4779896 | 0.07749772 | -6.1677893 | 6.93E-10 |
| YHB1 | YGR234W | 721.899594 | -0.5688781 | 0.0923201 | -6.1620183 | 7.18E-10 |
| 21S_RRNA | 21S_rRNA | 3027.51625 | -0.8784875 | 0.14260335 | -6.1603565 | 7.26E-10 |
| MGA2 | YIR033W | 2377.17652 | 0.38592385 | 0.06269036 | 6.1560314 | 7.46E-10 |
| SEC3 | YER008C | 2390.80916 | 0.38124976 | 0.06194513 | 6.15463639 | 7.52E-10 |
| SEC13 | YLR208W | 1255.03273 | -0.4186861 | 0.06807985 | -6.1499259 | 7.75E-10 |
| SDS22 | YKL193C | 648.162308 | -0.5821246 | 0.09469963 | -6.1470634 | 7.89E-10 |
| GTT1 | YIR038C | 423.994822 | -0.709509 | 0.11544079 | -6.1460863 | 7.94E-10 |
| TUP1 | YCR084C | 3302.13043 | -0.3270829 | 0.0532834 | -6.1385509 | 8.33E-10 |
| BOL2 | YGL220W | 884.198547 | -0.4580398 | 0.07462043 | -6.1382626 | 8.34E-10 |
| UBR1 | YGR184C | 3920.28263 | 0.32940199 | 0.05368572 | 6.13574675 | 8.48E-10 |
| YDR365W-B | YDR365W-B | 475.250245 | 0.60687418 | 0.09893609 | 6.13400219 | 8.57E-10 |
| Xu_2009_CUTs58 | Xu_2009_CU | 103.808981 | 1.00933093 | 0.16460013 | 6.13201798 | 8.68E-10 |
| MRPL28 | YDR462W | 334.473247 | -0.6113719 | 0.09973035 | -6.1302487 | 8.77E-10 |
| SEC66 | YBR171W | 823.729687 | -0.4419379 | 0.07212946 | -6.1270103 | 8.95E-10 |
| TPS3 | YMR261C | 1662.95229 | -0.3988312 | 0.06510949 | -6.1255467 | 9.04E-10 |
| MRPL13 | YKR006C | 527.303281 | -0.5157024 | 0.08422606 | -6.122837 | 9.19E-10 |
| TMA10 | YLR327C | 1402.00389 | -0.5555014 | 0.0907354 | -6.1222125 | 9.23E-10 |
| Xu_2009_SUTs31 | Xu_2009_SU | 85.527539 | 1.06700655 | 0.17434877 | 6.11995469 | 9.36E-10 |
| RCN2 | YOR220W | 873.068526 | -0.472646 | 0.07723492 | -6.1195898 | 9.38E-10 |
| IRC15 | YPL017C | 291.306236 | 0.68798469 | 0.11242387 | 6.11956053 | 9.38E-10 |
| BAS1 | YKR099W | 1070.11878 | 0.48095805 | 0.0786105 | 6.11824193 | 9.46E-10 |
| SNR86 | snR86 | 62847.6186 | -0.3939659 | 0.06440384 | -6.1171185 | 9.53E-10 |
| SSD1 | YDR293C | 5879.76689 | 0.2861006 | 0.04678724 | 6.11492809 | 9.66E-10 |
| SEC16 | YPL085W | 4051.79456 | 0.3330043 | 0.05446118 | 6.11452588 | 9.68E-10 |
| RTC6 | YPL183W-A | 219.527557 | -0.735681 | 0.12031956 | -6.1143921 | 9.69E-10 |
| KRE2 | YDR483W | 3454.51789 | -0.3900776 | 0.0638505 | -6.109233 | 1.00E-09 |
| DFM1 | YDR411C | 672.998682 | -0.4987329 | 0.0816531 | -6.107948 | 1.01E-09 |
| SNR69 | snR69 | 131.526157 | 0.87715204 | 0.1438231 | 6.098826 | 1.07E-09 |
| VPS55 | YJR044C | 739.022991 | -0.5181139 | 0.08499318 | -6.0959465 | 1.09E-09 |
| SFP1 | YLR403W | 1680.14295 | 0.37423228 | 0.06145136 | 6.08989416 | 1.13E-09 |
| TEA1 | YOR337W | 1756.30682 | 0.40199091 | 0.06602801 | 6.08818741 | 1.14E-09 |
| PDR10 | YOR328W | 703.416966 | 0.53225579 | 0.08746154 | 6.0855979 | 1.16E-09 |
| COT1 | YOR316C | 903.386109 | -0.4681442 | 0.07697673 | -6.0816329 | 1.19E-09 |
| ERG6 | YML008C | 2624.04204 | -0.3317107 | 0.0545452 | -6.0813923 | 1.19E-09 |
| Xu_2009_CUTs81 | Xu_2009_CU | 90.7173405 | 1.04210998 | 0.17178858 | 6.0662356 | 1.31E-09 |
| PIN2 | YOR104W | 495.08435 | 0.54576861 | 0.089981 | 6.06537589 | 1.32E-09 |
| RSF2 | YJR127C | 2348.14847 | 0.38326542 | 0.06319339 | 6.06496015 | 1.32E-09 |
| LST4 | YKL176C | 1319.33872 | 0.39219443 | 0.06469533 | 6.06217519 | 1.34E-09 |
| PGM1 | YKL127W | 811.917514 | -0.4419307 | 0.07292133 | -6.0603761 | 1.36E-09 |
| YPR172W | YPR172W | 495.798893 | -0.5461166 | 0.09015939 | -6.0572347 | 1.38E-09 |

| | | | | | | |
|---|---|---|---|---|---|---|
| NAN1 | YPL126W | 3672.3426 | 0.38336157 | 0.06329341 | 6.05689538 | 1.39E-09 |
| RSM25 | YIL093C | 596.719961 | -0.5046393 | 0.08333584 | -6.0554892 | 1.40E-09 |
| AXL2 | YIL140W | 747.56015 | -0.4496882 | 0.0743146 | -6.0511427 | 1.44E-09 |
| SWI1 | YPL016W | 857.459098 | 0.50001393 | 0.0826307 | 6.05118835 | 1.44E-09 |
| POF1 | YCL047C | 365.554075 | -0.6900278 | 0.11410078 | -6.0475291 | 1.47E-09 |
| PSG1 | YKL077W | 1310.77427 | -0.4344549 | 0.07185674 | -6.0461255 | 1.48E-09 |
| SHP1 | YBL058W | 2237.05742 | -0.3312468 | 0.05481799 | -6.0426657 | 1.52E-09 |
| Xu_2009_CUTs35 | Xu_2009_CU | 58.848004 | 1.18418731 | 0.196023 | 6.0410631 | 1.53E-09 |
| PRP3 | YDR473C | 459.949554 | 0.53744303 | 0.08902353 | 6.03708979 | 1.57E-09 |
| STU2 | YLR045C | 1425.10099 | 0.44397705 | 0.07355531 | 6.03596202 | 1.58E-09 |
| NFT1 | YKR103W | 304.165478 | 0.66908854 | 0.11087558 | 6.03458884 | 1.59E-09 |
| DBF2 | YGR092W | 985.890721 | 0.41036918 | 0.06800813 | 6.03411969 | 1.60E-09 |
| ADY3 | YDL239C | 427.78992 | 0.60201744 | 0.09983629 | 6.0300463 | 1.64E-09 |
| ISF1 | YMR081C | 290.779935 | -0.6538982 | 0.10846071 | -6.0288943 | 1.65E-09 |
| CLB2 | YPR119W | 1505.11769 | 0.38019576 | 0.063066 | 6.02853765 | 1.65E-09 |
| Xu_2009_CUTs12 | Xu_2009_CU | 83.9876501 | 1.07734345 | 0.17887652 | 6.02283331 | 1.71E-09 |
| NMD2 | YHR077C | 1771.53445 | 0.40469825 | 0.06721329 | 6.02110464 | 1.73E-09 |
| EOS1 | YNL080C | 967.097161 | -0.4252057 | 0.07067414 | -6.0164258 | 1.78E-09 |
| Xu_2009_CUTs26 | Xu_2009_CU | 97.0814276 | 1.02969851 | 0.17132797 | 6.01010151 | 1.85E-09 |
| LOA1 | YPR139C | 936.946541 | -0.4201983 | 0.06993219 | -6.0086539 | 1.87E-09 |
| GNA1 | YFL017C | 303.470743 | -0.6839707 | 0.11388399 | -6.0058547 | 1.90E-09 |
| YDR261W-B | YDR261W-B | 848.768762 | 0.44967615 | 0.07490308 | 6.00343991 | 1.93E-09 |
| RAD50 | YNL250W | 723.164793 | 0.46378911 | 0.0772613 | 6.0028646 | 1.94E-09 |
| DCS2 | YOR173W | 723.79021 | -0.532579 | 0.08880165 | -5.9973994 | 2.01E-09 |
| SAK1 | YER129W | 1827.52064 | 0.39241126 | 0.06547972 | 5.99286746 | 2.06E-09 |
| KIP2 | YPL155C | 638.81017 | 0.46684525 | 0.07792242 | 5.99115412 | 2.08E-09 |
| IES2 | YNL215W | 101.892415 | 1.02282108 | 0.17073945 | 5.99053765 | 2.09E-09 |
| CAX4 | YGR036C | 615.619571 | 0.51072472 | 0.08534073 | 5.98453686 | 2.17E-09 |
| Xu_2009_CUTs59 | Xu_2009_CU | 75.6329236 | 1.09572175 | 0.18318523 | 5.9814963 | 2.21E-09 |
| YDR210W | YDR210W | 212.179455 | -0.7532002 | 0.12596867 | -5.9792662 | 2.24E-09 |
| Xu_2009_SUTs10 | Xu_2009_SU | 101.468899 | 0.97441358 | 0.16313359 | 5.97310218 | 2.33E-09 |
| ERG3 | YLR056W | 2818.03935 | -0.3469995 | 0.05809654 | -5.9728085 | 2.33E-09 |
| YGR126W | YGR126W | 288.244967 | 0.66049739 | 0.11075954 | 5.96334546 | 2.47E-09 |
| YEL008C-A | YEL008C-A | 52.8289441 | 1.24719604 | 0.20929315 | 5.95908683 | 2.54E-09 |
| RGC1 | YPR115W | 2051.55235 | 0.32754338 | 0.05498187 | 5.95729805 | 2.56E-09 |
| Xu_2009_SUTs79 | Xu_2009_SU | 122.10641 | 0.92838524 | 0.15584787 | 5.95699648 | 2.57E-09 |
| GAT1 | YFL021W | 662.565317 | -0.4625141 | 0.07767715 | -5.9543138 | 2.61E-09 |
| BNA3 | YJL060W | 512.887785 | -0.5362422 | 0.09008525 | -5.9526078 | 2.64E-09 |
| MRP51 | YPL118W | 1234.44472 | -0.441151 | 0.07413348 | -5.9507654 | 2.67E-09 |
| CTF19 | YPL018W | 297.132425 | 0.69943795 | 0.11754936 | 5.95016375 | 2.68E-09 |
| VPS68 | YOL129W | 1014.06083 | -0.4413069 | 0.07418725 | -5.9485549 | 2.71E-09 |

| | | | | | | |
|---|---|---|---|---|---|---|
| ATP17 | YDR377W | 950.859413 | -0.5001169 | 0.08407669 | -5.9483414 | 2.71E-09 |
| Xu_2009_SUTs49: | Xu_2009_SU | 319.434851 | 0.5975985 | 0.10053313 | 5.94429413 | 2.78E-09 |
| YGL199C | YGL199C | 121.89598 | 0.88119216 | 0.14856388 | 5.9314025 | 3.00E-09 |
| NRM1 | YNR009W | 398.890551 | -0.5706892 | 0.09622896 | -5.9305352 | 3.02E-09 |
| YHR217C | YHR217C | 23.6963807 | 1.62759592 | 0.2744944 | 5.92943219 | 3.04E-09 |
| CSI2 | YOL007C | 1088.30428 | -0.4131261 | 0.06970976 | -5.9263745 | 3.10E-09 |
| YMR262W | YMR262W | 295.890073 | -0.6690997 | 0.11291629 | -5.9256263 | 3.11E-09 |
| GRX1 | YCL035C | 1773.55872 | -0.4579886 | 0.07742723 | -5.915084 | 3.32E-09 |
| MRPL38 | YKL170W | 394.014209 | -0.5614683 | 0.09509986 | -5.9039859 | 3.55E-09 |
| TAF12 | YDR145W | 1228.14143 | 0.37290884 | 0.06321103 | 5.89942683 | 3.65E-09 |
| PUL4 | YNR063W | 235.545509 | 0.67619593 | 0.11464174 | 5.89833951 | 3.67E-09 |
| FCY1 | YPR062W | 803.415591 | 0.53800641 | 0.09122083 | 5.89784617 | 3.68E-09 |
| RAD9 | YDR217C | 1536.83625 | 0.39338622 | 0.06670394 | 5.89749555 | 3.69E-09 |
| YOL075C | YOL075C | 713.99031 | 0.45810102 | 0.07768465 | 5.89693105 | 3.70E-09 |
| CMR3 | YPR013C | 549.541015 | 0.52495754 | 0.08903258 | 5.89624063 | 3.72E-09 |
| RPA190 | YOR341W | 6293.76712 | 0.36075839 | 0.06121045 | 5.89373836 | 3.78E-09 |
| POP6 | YGR030C | 348.677217 | -0.604675 | 0.10259871 | -5.8935923 | 3.78E-09 |
| RRG1 | YDR065W | 376.29308 | 0.56625303 | 0.0961464 | 5.88948764 | 3.87E-09 |
| YHI9 | YHR029C | 216.717534 | -0.6992257 | 0.11872724 | -5.8893451 | 3.88E-09 |
| Xu_2009_CUTs45 | Xu_2009_CU | 59.2001925 | 1.19771489 | 0.20345092 | 5.88699679 | 3.93E-09 |
| Xu_2009_SUTs44! | Xu_2009_SU | 90.8837456 | 0.9976835 | 0.1694864 | 5.88651054 | 3.94E-09 |
| UBC4 | YBR082C | 7299.67948 | -0.3234669 | 0.054957 | -5.8858173 | 3.96E-09 |
| GEF1 | YJR040W | 561.538091 | -0.5153796 | 0.08759256 | -5.8838287 | 4.01E-09 |
| ICL2 | YPR006C | 243.922741 | -0.7087825 | 0.12054544 | -5.8797953 | 4.11E-09 |
| ASF1 | YJL115W | 414.772799 | 0.64604167 | 0.10991015 | 5.8779071 | 4.15E-09 |
| SLK19 | YOR195W | 815.162348 | 0.44442071 | 0.07568735 | 5.87179615 | 4.31E-09 |
| ALG12 | YNR030W | 1357.70841 | -0.3844095 | 0.06548254 | -5.8704113 | 4.35E-09 |
| MXR2 | YCL033C | 325.347786 | -0.6192742 | 0.10549772 | -5.8700243 | 4.36E-09 |
| NRP1 | YDL167C | 1337.20838 | 0.41541258 | 0.07078217 | 5.86888713 | 4.39E-09 |
| ELF1 | YKL160W | 931.876693 | -0.4461112 | 0.07601458 | -5.8687585 | 4.39E-09 |
| Xu_2009_CUTs33 | Xu_2009_CU | 128.501401 | 0.90015757 | 0.15340957 | 5.86767554 | 4.42E-09 |
| MSW1 | YDR268W | 381.075896 | -0.5478137 | 0.09348151 | -5.8601286 | 4.63E-09 |
| Xu_2009_CUTs88 | Xu_2009_CU | 49.9352219 | 1.23155966 | 0.21025079 | 5.85757455 | 4.70E-09 |
| 15S_RRNA | 15S_rRNA | 828.194719 | -0.9586997 | 0.16370836 | -5.8561436 | 4.74E-09 |
| COQ11 | YLR290C | 287.361944 | -0.6192583 | 0.10580284 | -5.8529456 | 4.83E-09 |
| Xu_2009_CUTs60 | Xu_2009_CU | 63.4589311 | 1.14123693 | 0.1951652 | 5.84754318 | 4.99E-09 |
| ALG2 | YGL065C | 389.850772 | -0.5812259 | 0.09941251 | -5.846607 | 5.02E-09 |
| SNR3 | snR3 | 2152.1635 | -0.7834111 | 0.13408028 | -5.8428508 | 5.13E-09 |
| Xu_2009_SUTs17 | Xu_2009_SU | 215.26129 | 0.69105488 | 0.11835991 | 5.83858933 | 5.26E-09 |
| TRX1 | YLR043C | 727.585046 | -0.9386616 | 0.16086734 | -5.8350041 | 5.38E-09 |
| ARN1 | YHL040C | 592.038878 | -0.5627067 | 0.09644974 | -5.834196 | 5.41E-09 |

| | | | | | | |
|---|---|---|---|---|---|---|
| YOR1 | YGR281W | 3210.05315 | 0.29393904 | 0.05040477 | 5.83157182 | 5.49E-09 |
| MOD5 | YOR274W | 931.110417 | 0.41699109 | 0.07157422 | 5.82599517 | 5.68E-09 |
| CIA1 | YDR267C | 375.50836 | 0.55999005 | 0.09615571 | 5.82378389 | 5.75E-09 |
| RIO2 | YNL207W | 1262.97848 | 0.41616784 | 0.07159452 | 5.81284472 | 6.14E-09 |
| MNN2 | YBR015C | 2307.62918 | -0.3099842 | 0.05336889 | -5.808332 | 6.31E-09 |
| UBC1 | YDR177W | 1455.68705 | -0.3668829 | 0.06318301 | -5.8066701 | 6.37E-09 |
| Xu_2009_SUTs24 | Xu_2009_SU | 108.02066 | 0.9177502 | 0.15809358 | 5.80510715 | 6.43E-09 |
| SSU1 | YPL092W | 433.472587 | -0.5875347 | 0.10128607 | -5.8007448 | 6.60E-09 |
| Xu_2009_CUTs97 | Xu_2009_CU | 100.091522 | 1.01175725 | 0.17448201 | 5.79863347 | 6.69E-09 |
| SNC2 | YOR327C | 572.037163 | -0.5087689 | 0.08773839 | -5.7987031 | 6.68E-09 |
| RGT2 | YDL138W | 900.538406 | 0.39564837 | 0.06825416 | 5.79669235 | 6.76E-09 |
| HRK1 | YOR267C | 2261.32526 | 0.37415251 | 0.06454731 | 5.79656202 | 6.77E-09 |
| NEL1 | YHR035W | 358.212377 | 0.57510711 | 0.09923666 | 5.79530904 | 6.82E-09 |
| CHD1 | YER164W | 2487.56327 | 0.31473839 | 0.05432175 | 5.79396596 | 6.87E-09 |
| COQ1 | YBR003W | 669.149617 | -0.4439153 | 0.0766223 | -5.7935528 | 6.89E-09 |
| YOR111W | YOR111W | 166.191663 | -0.7608685 | 0.13135972 | -5.7922511 | 6.94E-09 |
| BMH1 | YER177W | 17004.3435 | -0.3246311 | 0.05605043 | -5.7917684 | 6.96E-09 |
| YEF3 | YLR249W | 33720.6538 | -0.3736881 | 0.06453066 | -5.7908616 | 7.00E-09 |
| Xu_2009_CUTs10 | Xu_2009_CU | 100.985243 | 0.96776735 | 0.16722981 | 5.78705062 | 7.16E-09 |
| DPC29 | YGR021W | 455.636084 | -0.6072437 | 0.10495558 | -5.7857215 | 7.22E-09 |
| KTR2 | YKR061W | 1967.71936 | 0.37556383 | 0.06494291 | 5.78298449 | 7.34E-09 |
| COQ4 | YDR204W | 549.818653 | -0.5166675 | 0.08946368 | -5.7751651 | 7.69E-09 |
| TRP4 | YDR354W | 764.186515 | -0.4377372 | 0.07582476 | -5.7730113 | 7.79E-09 |
| YML119W | YML119W | 593.809407 | 0.45941417 | 0.07966446 | 5.76686479 | 8.08E-09 |
| Xu_2009_SUTs16 | Xu_2009_SU | 65.7677821 | 1.11686396 | 0.1936986 | 5.76598869 | 8.12E-09 |
| IMH1 | YLR309C | 1044.10882 | 0.46621443 | 0.08086427 | 5.76539452 | 8.15E-09 |
| YMD8 | YML038C | 1389.02326 | -0.3649928 | 0.06339318 | -5.7576031 | 8.53E-09 |
| SRY1 | YKL218C | 251.640686 | -0.6608091 | 0.11479113 | -5.7566216 | 8.58E-09 |
| EHD3 | YDR036C | 632.528378 | -0.4658227 | 0.08094734 | -5.7546394 | 8.68E-09 |
| BSC3 | YLR465C | 84.0946054 | 0.97973737 | 0.170273 | 5.7539208 | 8.72E-09 |
| YBL100W-B | YBL100W-B | 429.893387 | 0.5547652 | 0.09646962 | 5.75067243 | 8.89E-09 |
| GSY1 | YFR015C | 394.951092 | -0.6323266 | 0.10999563 | -5.7486519 | 9.00E-09 |
| Xu_2009_CUTs25 | Xu_2009_CU | 54.3393105 | 1.1935133 | 0.20765125 | 5.74768187 | 9.05E-09 |
| RAD16 | YBR114W | 1704.07454 | 0.39351337 | 0.06848718 | 5.74579585 | 9.15E-09 |
| MRPS28 | YDR337W | 541.486663 | -0.5076719 | 0.08840403 | -5.7426333 | 9.32E-09 |
| Xu_2009_SUTs18 | Xu_2009_SU | 273.907265 | 0.60723866 | 0.10575863 | 5.74174086 | 9.37E-09 |
| RPH1 | YER169W | 2103.51563 | 0.41890574 | 0.07296239 | 5.7413926 | 9.39E-09 |
| TIF6 | YPR016C | 1284.75955 | -0.4341214 | 0.07563399 | -5.7397664 | 9.48E-09 |
| SPA2 | YLL021W | 2335.99579 | 0.38382306 | 0.06689523 | 5.73767463 | 9.60E-09 |
| ARL1 | YBR164C | 563.51879 | -0.4662449 | 0.08127495 | -5.7366369 | 9.66E-09 |
| DOT5 | YIL010W | 672.359451 | -0.4577261 | 0.07982561 | -5.7340751 | 9.80E-09 |

| | | | | | | |
|---|---|---|---|---|---|---|
| NMT1 | YLR195C | 1114.43172 | -0.3950711 | 0.06890246 | -5.7337732 | 9.82E-09 |
| NDD1 | YOR372C | 1680.17037 | 0.35382753 | 0.06173166 | 5.73170301 | 9.94E-09 |
| BEM2 | YER155C | 5022.49138 | 0.28133081 | 0.04910318 | 5.7293808 | 1.01E-08 |
| YDR210C-D | YDR210C-D | 732.021665 | 0.51238928 | 0.08943495 | 5.7291837 | 1.01E-08 |
| Xu_2009_CUTs35 | Xu_2009_CU | 67.2760093 | 1.11953562 | 0.19553156 | 5.72560048 | 1.03E-08 |
| MAS1 | YLR163C | 666.886639 | -0.458592 | 0.08013514 | -5.7227334 | 1.05E-08 |
| PSK2 | YOL045W | 1406.80757 | 0.4235003 | 0.07405439 | 5.71877404 | 1.07E-08 |
| FRK1 | YPL141C | 1523.49716 | 0.46477071 | 0.08129101 | 5.71736909 | 1.08E-08 |
| OXA1 | YER154W | 943.742709 | -0.471832 | 0.08256711 | -5.7145279 | 1.10E-08 |
| CHA1 | YCL064C | 297.77736 | 0.63367469 | 0.11095664 | 5.71101217 | 1.12E-08 |
| Xu_2009_CUTs5 | Xu_2009_CU | 80.0828031 | 1.07869098 | 0.18891637 | 5.7098861 | 1.13E-08 |
| MRPL33 | YMR286W | 258.10048 | -0.6425994 | 0.11255406 | -5.7092509 | 1.13E-08 |
| Xu_2009_CUTs23 | Xu_2009_CU | 38.409898 | 1.36006305 | 0.23837707 | 5.70551132 | 1.16E-08 |
| YKL030W | YKL030W | 28.6407078 | 1.43134319 | 0.25095542 | 5.7035755 | 1.17E-08 |
| MET5 | YJR137C | 996.69254 | 0.44607788 | 0.07825579 | 5.7002536 | 1.20E-08 |
| RLM1 | YPL089C | 2786.93005 | 0.31825788 | 0.05586675 | 5.69673113 | 1.22E-08 |
| ASE1 | YOR058C | 727.541425 | 0.42535557 | 0.07467134 | 5.69636961 | 1.22E-08 |
| IST3 | YIR005W | 117.20683 | -0.9555922 | 0.1678867 | -5.6918875 | 1.26E-08 |
| AIM20 | YIL158W | 414.040697 | -0.5329043 | 0.0936581 | -5.68989 | 1.27E-08 |
| KIC1 | YHR102W | 1562.13637 | 0.36072413 | 0.06345473 | 5.68474765 | 1.31E-08 |
| CHS3 | YBR023C | 2588.25164 | 0.31975641 | 0.0562612 | 5.68342644 | 1.32E-08 |
| HCR1 | YLR192C | 1517.55295 | -0.3652221 | 0.0642596 | -5.6835416 | 1.32E-08 |
| RIM15 | YFL033C | 2249.82807 | 0.33171022 | 0.05836849 | 5.68303566 | 1.32E-08 |
| NPL3 | YDR432W | 5656.952 | 0.33717154 | 0.0593356 | 5.68244893 | 1.33E-08 |
| Xu_2009_SUTs31 | Xu_2009_SU | 205.022526 | 0.71433663 | 0.12578623 | 5.67897343 | 1.36E-08 |
| BNI4 | YNL233W | 1413.09765 | 0.36117033 | 0.06361453 | 5.67748128 | 1.37E-08 |
| YPL113C | YPL113C | 43.4196185 | -1.2894741 | 0.2272586 | -5.6740386 | 1.39E-08 |
| DED81 | YHR019C | 4939.42432 | -0.282271 | 0.0497535 | -5.6733901 | 1.40E-08 |
| AST1 | YBL069W | 829.291125 | 0.46491032 | 0.08196422 | 5.67211295 | 1.41E-08 |
| HSP78 | YDR258C | 2902.50438 | -0.4391231 | 0.0774163 | -5.6722303 | 1.41E-08 |
| DBP7 | YKR024C | 1622.46536 | 0.40115783 | 0.07072735 | 5.67189093 | 1.41E-08 |
| RAD27 | YKL113C | 690.341509 | -0.4303943 | 0.07596619 | -5.6656028 | 1.47E-08 |
| OPT2 | YPR194C | 374.078844 | -0.5488415 | 0.09693437 | -5.6619904 | 1.50E-08 |
| NAT4 | YMR069W | 119.901291 | 0.86038241 | 0.15224544 | 5.65128536 | 1.59E-08 |
| PUT2 | YHR037W | 974.042782 | -0.405942 | 0.07184967 | -5.6498793 | 1.61E-08 |
| DPP1 | YDR284C | 439.566623 | -0.5196854 | 0.09198859 | -5.6494554 | 1.61E-08 |
| ARO8 | YGL202W | 3927.10372 | -0.2927218 | 0.05184889 | -5.6456713 | 1.65E-08 |
| IGO2 | YHR132W-A | 403.42412 | -0.5606587 | 0.09931722 | -5.6451312 | 1.65E-08 |
| CYB5 | YNL111C | 708.757092 | -0.4527454 | 0.0802534 | -5.6414476 | 1.69E-08 |
| MRPL49 | YJL096W | 407.264588 | -0.5264116 | 0.09336322 | -5.6383186 | 1.72E-08 |
| TOH1 | YJL171C | 3141.06612 | -0.379481 | 0.06733444 | -5.635764 | 1.74E-08 |

| | | | | | | |
|---|---|---|---|---|---|---|
| BUD4 | YJR092W | 1763.2031 | 0.38577124 | 0.06845793 | 5.63515822 | 1.75E-08 |
| YNL144C | YNL144C | 375.520498 | -0.5354289 | 0.09506035 | -5.6325151 | 1.78E-08 |
| EFM5 | YGR001C | 474.325994 | -0.5476113 | 0.09723416 | -5.6318819 | 1.78E-08 |
| AIM36 | YMR157C | 345.364474 | -0.614665 | 0.1091797 | -5.6298469 | 1.80E-08 |
| ARR1 | YPR199C | 474.27691 | -0.4922744 | 0.0875093 | -5.6253952 | 1.85E-08 |
| PPM1 | YDR435C | 174.809183 | -0.7272797 | 0.12932362 | -5.6237191 | 1.87E-08 |
| ITC1 | YGL133W | 1202.10741 | 0.42221739 | 0.0750798 | 5.62358185 | 1.87E-08 |
| ACM1 | YPL267W | 188.184259 | -0.718764 | 0.12782273 | -5.6231309 | 1.88E-08 |
| Xu_2009_SUTs48 | Xu_2009_SU | 171.638893 | 0.72972299 | 0.12979947 | 5.62192559 | 1.89E-08 |
| RAT1 | YOR048C | 2262.29952 | 0.35379702 | 0.06297907 | 5.61769211 | 1.94E-08 |
| BUD3 | YCL014W | 2001.71504 | 0.35765014 | 0.06368627 | 5.61581267 | 1.96E-08 |
| SGD1 | YLR336C | 1332.53153 | 0.41561081 | 0.07400923 | 5.61566184 | 1.96E-08 |
| Xu_2009_CUTs91 | Xu_2009_CU | 85.9973663 | 0.99141734 | 0.17663798 | 5.61270751 | 1.99E-08 |
| MRPS5 | YBR251W | 402.693234 | -0.5423578 | 0.09682945 | -5.6011656 | 2.13E-08 |
| DPM1 | YPR183W | 808.820303 | -0.4842539 | 0.08652233 | -5.5968658 | 2.18E-08 |
| ADE4 | YMR300C | 1002.57299 | -0.4219395 | 0.07539144 | -5.5966499 | 2.19E-08 |
| ALG6 | YOR002W | 942.96614 | -0.4223737 | 0.07559138 | -5.5875901 | 2.30E-08 |
| YFR006W | YFR006W | 1250.65446 | -0.3819399 | 0.06838453 | -5.5851803 | 2.33E-08 |
| MND2 | YIR025W | 259.253808 | 0.67043334 | 0.1200487 | 5.58467826 | 2.34E-08 |
| TDA10 | YGR205W | 266.937472 | -0.6836748 | 0.12244967 | -5.5833129 | 2.36E-08 |
| VPS72 | YDR485C | 1323.96797 | 0.35208388 | 0.06307309 | 5.58215713 | 2.38E-08 |
| XYL2 | YLR070C | 100.839774 | -0.9720393 | 0.17414361 | -5.581826 | 2.38E-08 |
| Xu_2009_SUTs26 | Xu_2009_SU | 171.010379 | 0.74539953 | 0.13356792 | 5.58067795 | 2.40E-08 |
| DRS1 | YLL008W | 2772.99924 | 0.39888966 | 0.0715058 | 5.57842408 | 2.43E-08 |
| Xu_2009_SUTs18 | Xu_2009_SU | 45.2634812 | 1.20670172 | 0.21632665 | 5.57814646 | 2.43E-08 |
| PRE7 | YBL041W | 1183.05196 | -0.3868947 | 0.06938277 | -5.5762354 | 2.46E-08 |
| Q0182 | Q0182 | 31.4585803 | -1.4104853 | 0.25307668 | -5.5733516 | 2.50E-08 |
| GAL83 | YER027C | 1057.71193 | -0.4171166 | 0.07484734 | -5.572898 | 2.51E-08 |
| YJR015W | YJR015W | 910.33924 | -0.4090337 | 0.07342313 | -5.5709101 | 2.53E-08 |
| RPF1 | YHR088W | 922.621189 | 0.45216208 | 0.08135453 | 5.55792122 | 2.73E-08 |
| LAM4 | YHR080C | 1709.19985 | 0.35958922 | 0.0647311 | 5.55512302 | 2.77E-08 |
| COX23 | YHR116W | 253.034425 | -0.6587305 | 0.11861589 | -5.5534765 | 2.80E-08 |
| Xu_2009_CUTs52 | Xu_2009_CU | 66.5549653 | 1.05629663 | 0.19022696 | 5.55282305 | 2.81E-08 |
| CDC3 | YLR314C | 2150.77122 | -0.3041899 | 0.05478861 | -5.5520645 | 2.82E-08 |
| ESF1 | YDR365C | 3023.99891 | 0.37472138 | 0.06757788 | 5.54502983 | 2.94E-08 |
| Xu_2009_CUTs70 | Xu_2009_CU | 130.83201 | 0.95900746 | 0.1730073 | 5.54316184 | 2.97E-08 |
| SFB3 | YHR098C | 1734.46205 | -0.3271022 | 0.05901438 | -5.5427537 | 2.98E-08 |
| YLR173W | YLR173W | 489.094797 | 0.47106128 | 0.08498932 | 5.54259378 | 2.98E-08 |
| COG8 | YML071C | 916.239801 | 0.38979951 | 0.07035002 | 5.54085881 | 3.01E-08 |
| HIS7 | YBR248C | 1111.44248 | -0.3997043 | 0.07214091 | -5.5406053 | 3.01E-08 |
| Xu_2009_CUTs19 | Xu_2009_CU | 45.4992869 | 1.20524383 | 0.21758143 | 5.5392772 | 3.04E-08 |

| | | | | | | |
|---|---|---|---|---|---|---|
| Xu_2009_SUTs82 | Xu_2009_SU | 290.335822 | 0.59668509 | 0.10777259 | 5.53651975 | 3.09E-08 |
| MRPL25 | YGR076C | 231.658011 | -0.6665359 | 0.12039076 | -5.5364371 | 3.09E-08 |
| Xu_2009_SUTs33 | Xu_2009_SU | 98.8325637 | 0.94247588 | 0.17028434 | 5.53471846 | 3.12E-08 |
| Xu_2009_SUTs26 | Xu_2009_SU | 112.839391 | 0.86419855 | 0.15621539 | 5.5320961 | 3.16E-08 |
| Xu_2009_SUTs10 | Xu_2009_SU | 57.5868523 | 1.12246244 | 0.2032652 | 5.52215741 | 3.35E-08 |
| YIF1 | YNL263C | 1117.87962 | -0.36863 | 0.06682917 | -5.5160051 | 3.47E-08 |
| Xu_2009_CUTs22 | Xu_2009_CU | 231.043388 | 0.68719581 | 0.12467138 | 5.51205761 | 3.55E-08 |
| Xu_2009_SUTs36 | Xu_2009_SU | 83.4743447 | 1.00549381 | 0.18241988 | 5.51197509 | 3.55E-08 |
| SDD2 | YMR074C | 395.414761 | -0.5566088 | 0.10102037 | -5.5098671 | 3.59E-08 |
| MRPL10 | YNL284C | 812.677003 | -0.4105514 | 0.07454708 | -5.5072767 | 3.64E-08 |
| SFA1 | YDL168W | 1097.02045 | -0.3828229 | 0.06951389 | -5.5071417 | 3.65E-08 |
| Xu_2009_CUTs78 | Xu_2009_CU | 29.7184453 | 1.43099101 | 0.26017395 | 5.50013177 | 3.80E-08 |
| Xu_2009_CUTs13 | Xu_2009_CU | 35.297343 | 1.29007887 | 0.23471248 | 5.49642219 | 3.88E-08 |
| MIA40 | YKL195W | 2008.67202 | -0.3838899 | 0.06984513 | -5.4963012 | 3.88E-08 |
| TVP18 | YMR071C | 401.560675 | -0.5793125 | 0.10540038 | -5.4963038 | 3.88E-08 |
| FMS1 | YMR020W | 767.888235 | 0.51296076 | 0.09335143 | 5.49494288 | 3.91E-08 |
| FCF1 | YDR339C | 702.958422 | -0.4579518 | 0.08336756 | -5.4931651 | 3.95E-08 |
| HNT1 | YDL125C | 1719.61207 | -0.3236843 | 0.05894722 | -5.4910876 | 3.99E-08 |
| PHO3 | YBR092C | 4166.75318 | -0.2678275 | 0.04880442 | -5.4877702 | 4.07E-08 |
| UPS2 | YLR168C | 505.130262 | -0.5163206 | 0.09408902 | -5.4875748 | 4.07E-08 |
| Xu_2009_CUTs27 | Xu_2009_CU | 46.1410331 | 1.20442224 | 0.21961945 | 5.48413282 | 4.16E-08 |
| YNL114C | YNL114C | 46.510596 | 1.23704715 | 0.22572105 | 5.48042444 | 4.24E-08 |
| HRT1 | YOL133W | 411.824137 | -0.5348777 | 0.09759678 | -5.4804847 | 4.24E-08 |
| MDM20 | YOL076W | 1135.10868 | 0.38856074 | 0.07092875 | 5.47818381 | 4.30E-08 |
| SEC9 | YGR009C | 938.603073 | 0.42112695 | 0.07688468 | 5.47738475 | 4.32E-08 |
| COQ21 | YBR230W-A | 146.644097 | -0.8198563 | 0.1497246 | -5.4757622 | 4.36E-08 |
| Xu_2009_SUTs39 | Xu_2009_SU | 105.111328 | -0.9321403 | 0.17028004 | -5.4741605 | 4.40E-08 |
| OSH3 | YHR073W | 3309.30867 | 0.33801048 | 0.06175141 | 5.47372902 | 4.41E-08 |
| STE3 | YKL178C | 1396.96629 | -0.3610339 | 0.06598602 | -5.4713699 | 4.47E-08 |
| ERJ5 | YFR041C | 401.545987 | -0.5290646 | 0.09669998 | -5.4711967 | 4.47E-08 |
| Xu_2009_CUTs42 | Xu_2009_CU | 38.2719062 | 1.32882355 | 0.24294191 | 5.46971732 | 4.51E-08 |
| Xu_2009_SUTs80 | Xu_2009_SU | 211.014266 | 0.65164895 | 0.1191454 | 5.46935882 | 4.52E-08 |
| EGO2 | YCR075W-A | 150.410972 | -0.7524693 | 0.13763472 | -5.4671474 | 4.57E-08 |
| SNR49 | snR49 | 2850.17735 | -1.2185874 | 0.22306146 | -5.4630118 | 4.68E-08 |
| RIM21 | YNL294C | 1540.63449 | 0.32882563 | 0.06027508 | 5.45541628 | 4.89E-08 |
| GCN2 | YDR283C | 1784.35081 | 0.34844137 | 0.06390396 | 5.45257897 | 4.96E-08 |
| HEM4 | YOR278W | 332.255236 | -0.6073399 | 0.11138534 | -5.4526019 | 4.96E-08 |
| COQ6 | YGR255C | 1315.87211 | -0.344055 | 0.06313603 | -5.4494249 | 5.05E-08 |
| LYS12 | YIL094C | 1167.23093 | -0.4077852 | 0.0748692 | -5.446635 | 5.13E-08 |
| LSB1 | YGR136W | 834.947716 | -0.4568424 | 0.08389883 | -5.4451578 | 5.18E-08 |
| KCS1 | YDR017C | 1975.89272 | 0.33504999 | 0.06155132 | 5.44342454 | 5.23E-08 |

| | | | | | | |
|---|---|---|---|---|---|---|
| Xu_2009_CUTs71 | Xu_2009_CU | 74.9835561 | 0.98727794 | 0.18141524 | 5.44208929 | 5.27E-08 |
| APE3 | YBR286W | 9541.40956 | -0.2817358 | 0.0517843 | -5.440565 | 5.31E-08 |
| KOG1 | YHR186C | 1732.32509 | 0.33249771 | 0.06115182 | 5.43724999 | 5.41E-08 |
| Xu_2009_CUTs87 | Xu_2009_CU | 69.5172466 | 1.03518627 | 0.19050648 | 5.43386391 | 5.51E-08 |
| MIG2 | YGL209W | 1597.4218 | -0.3504356 | 0.06450418 | -5.4327579 | 5.55E-08 |
| PRE2 | YPR103W | 1654.15327 | -0.3184595 | 0.05866425 | -5.4285095 | 5.68E-08 |
| ENT3 | YJR125C | 1275.91886 | -0.4203763 | 0.07744846 | -5.4278196 | 5.70E-08 |
| COX3 | Q0275 | 2158.79824 | -0.4315143 | 0.07957542 | -5.4227078 | 5.87E-08 |
| PRS1 | YKL181W | 2598.49829 | 0.33329219 | 0.06147172 | 5.42187799 | 5.90E-08 |
| MNN9 | YPL050C | 1658.33171 | -0.3359904 | 0.06200113 | -5.419101 | 5.99E-08 |
| SSY5 | YJL156C | 801.255812 | -0.3900519 | 0.07200468 | -5.4170353 | 6.06E-08 |
| KSH1 | YNL024C-A | 271.069226 | -0.5844331 | 0.10791447 | -5.4157068 | 6.10E-08 |
| CAB4 | YGR277C | 490.713516 | -0.4543881 | 0.08390378 | -5.4155853 | 6.11E-08 |
| SDH6 | YDR379C-A | 348.213545 | -0.5461663 | 0.10092294 | -5.411716 | 6.24E-08 |
| ESC1 | YMR219W | 1876.93083 | 0.37737647 | 0.06973289 | 5.41174295 | 6.24E-08 |
| MCH4 | YOL119C | 636.687926 | 0.47566437 | 0.08789288 | 5.41186497 | 6.24E-08 |
| YPR204W | YPR204W | 133.562259 | 0.81469799 | 0.15057194 | 5.41068923 | 6.28E-08 |
| PHS1 | YJL097W | 370.377019 | -0.5128879 | 0.09479696 | -5.4103836 | 6.29E-08 |
| BFR2 | YDR299W | 1224.15697 | 0.39050707 | 0.07218769 | 5.40960767 | 6.32E-08 |
| MSK1 | YNL073W | 726.723758 | -0.4354181 | 0.08055342 | -5.4053332 | 6.47E-08 |
| YRB30 | YGL164C | 256.24871 | 0.61737377 | 0.11429797 | 5.40144122 | 6.61E-08 |
| SUC2 | YIL162W | 879.473808 | -0.3798149 | 0.07032164 | -5.4011098 | 6.62E-08 |
| Xu_2009_CUTs60 | Xu_2009_CU | 941.478373 | 0.36275938 | 0.06716996 | 5.40061888 | 6.64E-08 |
| DIP5 | YPL265W | 993.836548 | -0.4571615 | 0.08468007 | -5.3986906 | 6.71E-08 |
| COX5B | YIL111W | 1145.68303 | -0.3678003 | 0.06813893 | -5.3977995 | 6.75E-08 |
| SPO71 | YDR104C | 693.805537 | 0.40224575 | 0.07452382 | 5.39754566 | 6.76E-08 |
| PAN5 | YHR063C | 916.865121 | -0.4051401 | 0.07510848 | -5.3940668 | 6.89E-08 |
| MRP7 | YNL005C | 819.731768 | -0.3990981 | 0.07399107 | -5.3938687 | 6.90E-08 |
| UBI4 | YLL039C | 4934.09767 | -0.3429012 | 0.0635851 | -5.3927919 | 6.94E-08 |
| PRY2 | YKR013W | 4598.4821 | -0.3239413 | 0.0600883 | -5.3910875 | 7.00E-08 |
| Xu_2009_SUTs56 | Xu_2009_SU | 133.546443 | 0.77559838 | 0.14388949 | 5.39023642 | 7.04E-08 |
| MTM1 | YGR257C | 552.00027 | -0.4749684 | 0.08814096 | -5.3887361 | 7.10E-08 |
| EAF7 | YNL136W | 514.04369 | 0.48235486 | 0.08951888 | 5.38830315 | 7.11E-08 |
| GPT2 | YKR067W | 731.380263 | -0.515208 | 0.09562453 | -5.3878224 | 7.13E-08 |
| RPL18B | YNL301C | 2148.08832 | 0.34661055 | 0.06434725 | 5.38656325 | 7.18E-08 |
| NTE1 | YML059C | 2098.96923 | 0.30527651 | 0.05667548 | 5.38639446 | 7.19E-08 |
| PRE1 | YER012W | 1546.62207 | -0.3671769 | 0.06816992 | -5.3862015 | 7.20E-08 |
| SNR80 | snR80 | 34.4867167 | 1.34675069 | 0.25005501 | 5.38581768 | 7.21E-08 |
| DSK2 | YMR276W | 1623.12814 | -0.327299 | 0.06077338 | -5.3855655 | 7.22E-08 |
| Xu_2009_SUTs38 | Xu_2009_SU | 99.1123461 | 0.88337802 | 0.16408764 | 5.38357454 | 7.30E-08 |
| RPS14B | YJL191W | 3286.05455 | -0.3253124 | 0.06042621 | -5.38363 | 7.30E-08 |

| | | | | | | |
|---|---|---|---|---|---|---|
| CRS1 | YNL247W | 3257.92511 | -0.3135911 | 0.05826396 | -5.3822484 | 7.36E-08 |
| Xu_2009_SUTs67( | Xu_2009_SU | 163.55282 | 0.72225825 | 0.13419641 | 5.38209809 | 7.36E-08 |
| SGE1 | YPR198W | 467.722823 | 0.46835474 | 0.08702808 | 5.38165061 | 7.38E-08 |
| SNU114 | YKL173W | 1267.71412 | 0.34078846 | 0.06333503 | 5.3807263 | 7.42E-08 |
| FMP49 | YER038W-A | 78.918016 | 0.95931171 | 0.17842043 | 5.37669195 | 7.59E-08 |
| TRK1 | YJL129C | 2187.62903 | 0.30523518 | 0.05682051 | 5.37191924 | 7.79E-08 |
| TOM71 | YHR117W | 1158.05247 | -0.3484629 | 0.06487396 | -5.3713835 | 7.81E-08 |
| OST2 | YOR103C | 976.116169 | -0.3690985 | 0.06872492 | -5.3706643 | 7.84E-08 |
| YMR181C | YMR181C | 938.528592 | 0.36643249 | 0.06828222 | 5.36644113 | 8.03E-08 |
| ZRC1 | YMR243C | 2305.29477 | -0.3730103 | 0.06953126 | -5.3646414 | 8.11E-08 |
| MRPL8 | YJL063C | 592.271209 | -0.4375126 | 0.08157577 | -5.3632667 | 8.17E-08 |
| Xu_2009_SUTs11 | Xu_2009_SU | 60.8139182 | 1.0756484 | 0.20057835 | 5.36273429 | 8.20E-08 |
| Xu_2009_CUTs79 | Xu_2009_CU | 90.7805328 | 0.90041197 | 0.16794266 | 5.36142494 | 8.26E-08 |
| CEP3 | YMR168C | 343.738983 | 0.53695366 | 0.10027821 | 5.35463968 | 8.57E-08 |
| Xu_2009_CUTs50 | Xu_2009_CU | 32.8752022 | 1.28377028 | 0.23985935 | 5.35217944 | 8.69E-08 |
| POX1 | YGL205W | 64.5721674 | 1.04716297 | 0.19565053 | 5.35221125 | 8.69E-08 |
| TIM17 | YJL143W | 1160.02257 | -0.3643541 | 0.0680747 | -5.3522692 | 8.69E-08 |
| SNR70 | snR70 | 231.865507 | -0.6333974 | 0.11837144 | -5.3509308 | 8.75E-08 |
| PYC1 | YGL062W | 4971.73763 | -0.2907955 | 0.05436991 | -5.3484643 | 8.87E-08 |
| Xu_2009_CUTs38 | Xu_2009_CU | 69.5138334 | 1.07932821 | 0.20184873 | 5.34721321 | 8.93E-08 |
| YCG1 | YDR325W | 864.935188 | 0.40517357 | 0.07587548 | 5.33998059 | 9.30E-08 |
| PEX3 | YDR329C | 749.321391 | -0.3938942 | 0.07378181 | -5.3386361 | 9.36E-08 |
| FUB1 | YCR076C | 517.434778 | -0.4609664 | 0.08635788 | -5.3378618 | 9.40E-08 |
| KEX1 | YGL203C | 2642.21272 | 0.37645784 | 0.07054718 | 5.33625622 | 9.49E-08 |
| MEF2 | YJL102W | 474.453288 | 0.47292736 | 0.08864773 | 5.33490662 | 9.56E-08 |
| STI1 | YOR027W | 7746.3594 | -0.3809005 | 0.07139956 | -5.334774 | 9.57E-08 |
| BET5 | YML077W | 426.049944 | -0.4790307 | 0.08979817 | -5.3345265 | 9.58E-08 |
| COQ3 | YOL096C | 207.909261 | -0.6309832 | 0.11829175 | -5.3341269 | 9.60E-08 |
| EBP2 | YKL172W | 2692.62209 | 0.40881229 | 0.07666748 | 5.33227752 | 9.70E-08 |
| Xu_2009_CUTs41 | Xu_2009_CU | 82.0506094 | 0.95762956 | 0.17987835 | 5.32376228 | 1.02E-07 |
| CLC1 | YGR167W | 1715.23023 | -0.3324774 | 0.06245839 | -5.3231832 | 1.02E-07 |
| PGA3 | YML125C | 1290.09969 | -0.4120948 | 0.07742687 | -5.3223746 | 1.02E-07 |
| YDL086C-A | YDL086C-A | 107.033566 | 0.84730732 | 0.15923254 | 5.32119457 | 1.03E-07 |
| CIN2 | YPL241C | 52.5083521 | -1.1087572 | 0.2083799 | -5.320845 | 1.03E-07 |
| SRL1 | YOR247W | 800.102335 | -0.4099387 | 0.07705015 | -5.3204147 | 1.04E-07 |
| GAP1 | YKR039W | 471.024825 | -0.4603262 | 0.08654396 | -5.3189874 | 1.04E-07 |
| SNR79 | snR79 | 230.462292 | -0.7123053 | 0.13392289 | -5.3187717 | 1.04E-07 |
| CCT5 | YJR064W | 3309.83043 | -0.2870141 | 0.05399083 | -5.3159776 | 1.06E-07 |
| YLR363W-A | YLR363W-A | 455.959922 | -0.5315017 | 0.10006184 | -5.3117327 | 1.09E-07 |
| NUP192 | YJL039C | 1919.22057 | 0.34081975 | 0.06417051 | 5.31115818 | 1.09E-07 |
| Xu_2009_CUTs53 | Xu_2009_CU | 36.6422177 | 1.24748248 | 0.23499229 | 5.30861033 | 1.10E-07 |

| | | | | | | |
|---|---|---|---|---|---|---|
| Xu_2009_SUTs69: | Xu_2009_SU | 149.751917 | 0.75702304 | 0.14262789 | 5.30767887 | 1.11E-07 |
| NOP8 | YOL144W | 656.778624 | 0.50845588 | 0.09584689 | 5.30487599 | 1.13E-07 |
| MCR1 | YKL150W | 2318.3966 | -0.3910031 | 0.07373591 | -5.3027503 | 1.14E-07 |
| YPL250W-A | YPL250W-A | 37.2123854 | 1.22038969 | 0.23020202 | 5.30138568 | 1.15E-07 |
| YBR056W | YBR056W | 973.742384 | -0.4571738 | 0.08624096 | -5.3011216 | 1.15E-07 |
| Xu_2009_CUTs29 | Xu_2009_CU | 34.8515353 | 1.26908158 | 0.23943711 | 5.30027093 | 1.16E-07 |
| HAM1 | YJR069C | 418.344887 | -0.5427003 | 0.10241592 | -5.2989835 | 1.16E-07 |
| SHE4 | YOR035C | 883.192843 | 0.39871336 | 0.07532106 | 5.29351767 | 1.20E-07 |
| MEP2 | YNL142W | 302.228224 | -0.6165744 | 0.11658093 | -5.2888102 | 1.23E-07 |
| NMA1 | YLR328W | 501.057067 | -0.5250636 | 0.09928435 | -5.2884827 | 1.23E-07 |
| Xu_2009_SUTs13( | Xu_2009_SU | 130.523837 | 0.77436995 | 0.14651807 | 5.2851499 | 1.26E-07 |
| SKT5 | YBL061C | 1282.78116 | 0.32709091 | 0.0619589 | 5.27915971 | 1.30E-07 |
| OSH6 | YKR003W | 729.42782 | -0.4130639 | 0.07824879 | -5.2788538 | 1.30E-07 |
| Xu_2009_CUTs27 | Xu_2009_CU | 43.8024328 | -1.2136606 | 0.22992694 | -5.2784618 | 1.30E-07 |
| LSB5 | YCL034W | 545.436132 | -0.4347396 | 0.08237704 | -5.2774369 | 1.31E-07 |
| MEP1 | YGR121C | 182.69978 | -0.6539617 | 0.12392854 | -5.2769256 | 1.31E-07 |
| BCS1 | YDR375C | 435.571454 | -0.509542 | 0.09659543 | -5.2750121 | 1.33E-07 |
| YML108W | YML108W | 404.785722 | 0.56109421 | 0.1063761 | 5.27462655 | 1.33E-07 |
| YEL025C | YEL025C | 938.861726 | 0.40731799 | 0.07723077 | 5.27403771 | 1.33E-07 |
| Xu_2009_CUTs86 | Xu_2009_CU | 35.2044381 | 1.24634092 | 0.2363593 | 5.27307761 | 1.34E-07 |
| QRI1 | YDL103C | 895.507853 | -0.3749492 | 0.0711237 | -5.2717899 | 1.35E-07 |
| NPT1 | YOR209C | 2849.19441 | -0.3689626 | 0.07002473 | -5.2690331 | 1.37E-07 |
| NGL3 | YML118W | 342.85955 | 0.5291998 | 0.10044972 | 5.26830548 | 1.38E-07 |
| Xu_2009_CUTs42 | Xu_2009_CU | 40.680154 | 1.20270849 | 0.22853429 | 5.26270471 | 1.42E-07 |
| YCR051W | YCR051W | 733.908051 | -0.5179292 | 0.09841321 | -5.262801 | 1.42E-07 |
| DSD1 | YGL196W | 588.695276 | -0.4459651 | 0.08474558 | -5.2623997 | 1.42E-07 |
| NUT1 | YGL151W | 1333.15821 | 0.33498535 | 0.06371374 | 5.25766244 | 1.46E-07 |
| VMA10 | YHR039C-A | 1258.37703 | -0.3462441 | 0.06586078 | -5.2572126 | 1.46E-07 |
| DIA2 | YOR080W | 665.709819 | 0.42273605 | 0.080415 | 5.25693034 | 1.46E-07 |
| LTP1 | YPR073C | 667.105496 | -0.4246918 | 0.08079677 | -5.2562965 | 1.47E-07 |
| ODC1 | YPL134C | 198.729363 | -0.6423619 | 0.12229693 | -5.2524776 | 1.50E-07 |
| SHR3 | YDL212W | 1218.10964 | -0.330592 | 0.06300266 | -5.2472714 | 1.54E-07 |
| STO1 | YMR125W | 1791.50429 | -0.3156876 | 0.0602339 | -5.2410281 | 1.60E-07 |
| IZH3 | YLR023C | 1713.684 | 0.36488081 | 0.06964795 | 5.23893066 | 1.62E-07 |
| HAP4 | YKL109W | 806.306377 | -0.3792889 | 0.07244927 | -5.2352337 | 1.65E-07 |
| Xu_2009_CUTs85 | Xu_2009_CU | 81.8380641 | 0.95272079 | 0.18204149 | 5.23353647 | 1.66E-07 |
| HST3 | YOR025W | 830.060252 | 0.41472835 | 0.07927168 | 5.231734 | 1.68E-07 |
| DEF1 | YKL054C | 7393.28587 | -0.2910276 | 0.05563498 | -5.2310186 | 1.69E-07 |
| HSE1 | YHL002W | 697.913152 | -0.3977234 | 0.07603835 | -5.2305628 | 1.69E-07 |
| RRT5 | YFR032C | 329.394355 | -0.5861079 | 0.11214317 | -5.2264247 | 1.73E-07 |
| Xu_2009_CUTs38 | Xu_2009_CU | 57.6302441 | 1.14556402 | 0.21924791 | 5.22497128 | 1.74E-07 |

| | | | | | | |
|---|---|---|---|---|---|---|
| Xu_2009_CUTs47 | Xu_2009_CU | 61.6222469 | 1.03349779 | 0.19783384 | 5.22406989 | 1.75E-07 |
| Xu_2009_SUTs33 | Xu_2009_SU | 198.502086 | 0.63812247 | 0.12221034 | 5.22150948 | 1.77E-07 |
| CPA2 | YJR109C | 1818.12901 | -0.3138813 | 0.06011285 | -5.2215347 | 1.77E-07 |
| TYS1 | YGR185C | 3240.31985 | -0.2799363 | 0.05363313 | -5.2194667 | 1.79E-07 |
| SHE3 | YBR130C | 827.082873 | -0.3624672 | 0.06948258 | -5.2166633 | 1.82E-07 |
| SKS1 | YPL026C | 1224.61231 | 0.35262665 | 0.06760512 | 5.2159756 | 1.83E-07 |
| Xu_2009_CUTs18 | Xu_2009_CU | 136.742641 | 0.76750756 | 0.14721529 | 5.21350444 | 1.85E-07 |
| YER152C | YER152C | 917.136619 | -0.449363 | 0.08628354 | -5.2079802 | 1.91E-07 |
| YNL058C | YNL058C | 812.791063 | 0.44016533 | 0.08452624 | 5.20744011 | 1.91E-07 |
| CBP6 | YBR120C | 143.832525 | -0.7157234 | 0.13747586 | -5.2061749 | 1.93E-07 |
| CSE4 | YKL049C | 532.688293 | 0.53810304 | 0.10337191 | 5.20550534 | 1.93E-07 |
| BOL3 | YAL046C | 186.544578 | -0.6958078 | 0.13367398 | -5.2052598 | 1.94E-07 |
| SDH8 | YBR269C | 490.653728 | -0.4775682 | 0.09179378 | -5.2026199 | 1.96E-07 |
| HCA4 | YJL033W | 1982.58363 | 0.37736902 | 0.0726324 | 5.1956017 | 2.04E-07 |
| ERG28 | YER044C | 798.037694 | -0.4011547 | 0.07721689 | -5.1951684 | 2.05E-07 |
| YFL012W | YFL012W | 12.3992544 | -1.532489 | 0.29501505 | -5.1946129 | 2.05E-07 |
| PSR2 | YLR019W | 1049.2921 | -0.3656669 | 0.07039741 | -5.1943236 | 2.05E-07 |
| LAC1 | YKL008C | 835.043169 | -0.4788685 | 0.09219972 | -5.1938176 | 2.06E-07 |
| CTH1 | YDR151C | 1034.06783 | 0.35463116 | 0.06829107 | 5.19293643 | 2.07E-07 |
| YLL037W | YLL037W | 52.5209639 | 1.15041994 | 0.22166032 | 5.19001309 | 2.10E-07 |
| RAD23 | YEL037C | 2504.78611 | -0.3100971 | 0.05975988 | -5.1890506 | 2.11E-07 |
| CMC2 | YBL059C-A | 409.558464 | -0.5291906 | 0.10200337 | -5.1879712 | 2.13E-07 |
| GRR1 | YJR090C | 1825.17719 | 0.33074684 | 0.06378922 | 5.18499593 | 2.16E-07 |
| RGL1 | YPL066W | 1843.4608 | -0.329712 | 0.0636162 | -5.1828318 | 2.19E-07 |
| SAM4 | YPL273W | 976.279188 | -0.3778842 | 0.07291625 | -5.1824412 | 2.19E-07 |
| YJL225C | YJL225C | 72.1169236 | 0.94222704 | 0.18183191 | 5.18185743 | 2.20E-07 |
| PIB2 | YGL023C | 1157.95717 | 0.35062928 | 0.06771247 | 5.17820879 | 2.24E-07 |
| TOP3 | YLR234W | 296.588957 | 0.54190137 | 0.1046583 | 5.17781547 | 2.24E-07 |
| CFT2 | YLR115W | 1015.30999 | 0.3872634 | 0.07486242 | 5.17300132 | 2.30E-07 |
| YOR225W | YOR225W | 29.2699256 | 1.2709868 | 0.24600521 | 5.16650357 | 2.39E-07 |
| SEC23 | YPR181C | 3830.05126 | -0.2767408 | 0.05357793 | -5.1652009 | 2.40E-07 |
| GSF2 | YML048W | 2249.46281 | -0.3233514 | 0.06261142 | -5.1644158 | 2.41E-07 |
| NOP14 | YDL148C | 1719.30037 | 0.3286997 | 0.06366318 | 5.1631052 | 2.43E-07 |
| ESC2 | YDR363W | 536.537128 | 0.44150188 | 0.08555472 | 5.16046209 | 2.46E-07 |
| SLC1 | YDL052C | 968.147001 | -0.3629584 | 0.07040199 | -5.1555132 | 2.53E-07 |
| SKI2 | YLR398C | 1587.6913 | 0.31836873 | 0.06177212 | 5.15392255 | 2.55E-07 |
| SSO2 | YMR183C | 989.89262 | -0.3678418 | 0.07137808 | -5.1534283 | 2.56E-07 |
| SMC1 | YFL008W | 1030.0565 | 0.36958097 | 0.07173084 | 5.15233047 | 2.57E-07 |
| AGE2 | YIL044C | 679.727724 | -0.3957159 | 0.07681555 | -5.1515076 | 2.58E-07 |
| Xu_2009_CUTs76 | Xu_2009_CU | 56.9327372 | 1.03746867 | 0.20149987 | 5.1487313 | 2.62E-07 |
| CDC60 | YPL160W | 5526.8397 | -0.2609131 | 0.05070823 | -5.1453802 | 2.67E-07 |

| | | | | | | |
|---|---|---|---|---|---|---|
| ACH1 | YBL015W | 716.5441 | -0.4172745 | 0.08112499 | -5.1436 | 2.70E-07 |
| YRB1 | YDR002W | 1777.60003 | -0.3898302 | 0.07581544 | -5.1418314 | 2.72E-07 |
| MEH1 | YKR007W | 297.605038 | -0.5469067 | 0.10651055 | -5.1347654 | 2.82E-07 |
| MRPL31 | YKL138C | 535.742111 | -0.4525315 | 0.08814149 | -5.1341488 | 2.83E-07 |
| ROG3 | YFR022W | 773.331061 | 0.42384128 | 0.08256838 | 5.13321525 | 2.85E-07 |
| RTF1 | YGL244W | 676.445663 | 0.50319603 | 0.09804958 | 5.13205706 | 2.87E-07 |
| CCT4 | YDL143W | 2150.09799 | -0.3245448 | 0.0633118 | -5.1261337 | 2.96E-07 |
| MNN4 | YKL201C | 1030.55221 | 0.43491802 | 0.08484527 | 5.12601383 | 2.96E-07 |
| Xu_2009_CUTs29 | Xu_2009_CU | 107.989005 | 0.93483229 | 0.18261529 | 5.11913481 | 3.07E-07 |
| JJJ1 | YNL227C | 820.65609 | 0.40759549 | 0.07962738 | 5.11878536 | 3.08E-07 |
| ARG5,6 | YER069W | 1296.91794 | -0.3328857 | 0.06503706 | -5.1183998 | 3.08E-07 |
| BOP3 | YNL042W | 863.549416 | 0.37843812 | 0.07395861 | 5.11689044 | 3.11E-07 |
| RTG1 | YOL067C | 288.786652 | 0.5439265 | 0.10631724 | 5.11607049 | 3.12E-07 |
| Xu_2009_CUTs86 | Xu_2009_CU | 32.5746067 | 1.28179825 | 0.25072182 | 5.11243197 | 3.18E-07 |
| POL5 | YEL055C | 2114.0913 | 0.32349912 | 0.06330771 | 5.10994837 | 3.22E-07 |
| ULS1 | YOR191W | 2152.78265 | 0.30276656 | 0.05925608 | 5.10946007 | 3.23E-07 |
| Xu_2009_CUTs12 | Xu_2009_CU | 47.921157 | 1.14037651 | 0.22321417 | 5.1088893 | 3.24E-07 |
| BAP2 | YBR068C | 5536.82416 | 0.28572554 | 0.05594347 | 5.10739767 | 3.27E-07 |
| Xu_2009_SUTs63 | Xu_2009_SU | 413.488451 | 0.47590229 | 0.093207 | 5.10586407 | 3.29E-07 |
| CAT5 | YOR125C | 415.63058 | -0.5184988 | 0.1015565 | -5.1055203 | 3.30E-07 |
| Xu_2009_SUTs13 | Xu_2009_SU | 189.330209 | 0.64949346 | 0.12726675 | 5.10340243 | 3.34E-07 |
| TIF3 | YPR163C | 3277.15301 | -0.3504719 | 0.06873022 | -5.0992399 | 3.41E-07 |
| RPP2B | YDR382W | 2865.10901 | -0.4089706 | 0.08021054 | -5.0987145 | 3.42E-07 |
| CCC2 | YDR270W | 558.119115 | -0.4517435 | 0.08866987 | -5.0946669 | 3.49E-07 |
| ADE5,7 | YGL234W | 2388.40904 | -0.3367029 | 0.06609387 | -5.0943143 | 3.50E-07 |
| SCD5 | YOR329C | 1368.64814 | 0.35494339 | 0.06968483 | 5.09355325 | 3.51E-07 |
| UBP10 | YNL186W | 2673.73764 | 0.3072081 | 0.06032102 | 5.09288644 | 3.53E-07 |
| DCP1 | YOL149W | 386.145833 | -0.5033795 | 0.09886193 | -5.0917425 | 3.55E-07 |
| AVO2 | YMR068W | 211.369966 | 0.63748946 | 0.12532522 | 5.08668145 | 3.64E-07 |
| GOT1 | YMR292W | 136.181158 | -0.7701417 | 0.15140475 | -5.0866413 | 3.64E-07 |
| ZRT2 | YLR130C | 1432.91307 | -0.3072825 | 0.06042271 | -5.0855467 | 3.67E-07 |
| SCL1 | YGL011C | 1634.003 | -0.3254957 | 0.0640058 | -5.0854104 | 3.67E-07 |
| YNL089C | YNL089C | 25.4890093 | 1.34156272 | 0.26387329 | 5.08411716 | 3.69E-07 |
| ATS1 | YAL020C | 390.085344 | 0.50422042 | 0.09917951 | 5.08391722 | 3.70E-07 |
| IMD2 | YHR216W | 231.336907 | 0.63307231 | 0.12461687 | 5.08014916 | 3.77E-07 |
| TLG1 | YDR468C | 800.230518 | 0.46306905 | 0.09120654 | 5.07714733 | 3.83E-07 |
| AMD1 | YML035C | 2046.79031 | 0.34122565 | 0.06723525 | 5.0751007 | 3.87E-07 |
| PST1 | YDR055W | 7265.50738 | 0.32926864 | 0.06492951 | 5.07117125 | 3.95E-07 |
| TRS20 | YBR254C | 295.450483 | -0.5384173 | 0.10617969 | -5.0708125 | 3.96E-07 |
| SSH1 | YBR283C | 2422.46869 | -0.3113036 | 0.06142759 | -5.0678141 | 4.02E-07 |
| ERS1 | YCR075C | 128.207553 | -0.8254972 | 0.16290538 | -5.0673412 | 4.03E-07 |

| | | | | | | |
|---|---|---|---|---|---|---|
| PGA2 | YNL149C | 1034.42795 | -0.3568693 | 0.07044213 | -5.0661342 | 4.06E-07 |
| BRR1 | YPR057W | 375.572752 | -0.4909257 | 0.09695173 | -5.0636091 | 4.11E-07 |
| SAM35 | YHR083W | 865.793913 | -0.3989673 | 0.07879405 | -5.0634195 | 4.12E-07 |
| COX11 | YPL132W | 329.679426 | -0.4970245 | 0.09819243 | -5.0617391 | 4.15E-07 |
| FMP10 | YER182W | 413.254357 | -0.4689068 | 0.09264338 | -5.0614177 | 4.16E-07 |
| BRR6 | YGL247W | 306.79942 | -0.5531183 | 0.10928606 | -5.0611972 | 4.17E-07 |
| SPI1 | YER150W | 2328.56289 | -0.3466955 | 0.06854209 | -5.0581399 | 4.23E-07 |
| SNR161 | snR161 | 234.548172 | 0.61707342 | 0.12209464 | 5.05405822 | 4.33E-07 |
| DFG5 | YMR238W | 947.508502 | 0.35373167 | 0.07002304 | 5.05164689 | 4.38E-07 |
| Xu_2009_CUTs79 | Xu_2009_CU | 47.9621565 | 1.08898306 | 0.21565582 | 5.0496345 | 4.43E-07 |
| YOS1 | YER074W-A | 686.021907 | -0.4193945 | 0.08306474 | -5.0490082 | 4.44E-07 |
| IPA1 | YJR141W | 178.959902 | -0.6584837 | 0.13059836 | -5.042052 | 4.61E-07 |
| DNF2 | YDR093W | 2097.64456 | 0.31330846 | 0.06221399 | 5.03598065 | 4.75E-07 |
| YMR050C | YMR050C | 240.169106 | 0.60149771 | 0.11949826 | 5.03352711 | 4.82E-07 |
| PRP5 | YBR237W | 883.210255 | 0.36890761 | 0.07331743 | 5.0316496 | 4.86E-07 |
| NUP188 | YML103C | 3764.60717 | 0.30278401 | 0.0601879 | 5.03064567 | 4.89E-07 |
| Xu_2009_SUTs77 | Xu_2009_SU | 70.3926605 | 0.95107732 | 0.18909971 | 5.02950178 | 4.92E-07 |
| HTS1 | YPR033C | 2088.79961 | -0.2912071 | 0.05790153 | -5.029351 | 4.92E-07 |
| CMI7 | YIL002W-A | 297.334196 | -0.5774375 | 0.11481967 | -5.0290818 | 4.93E-07 |
| Xu_2009_CUTs51 | Xu_2009_CU | 67.723308 | 0.98900925 | 0.19671987 | 5.02750052 | 4.97E-07 |
| TYE7 | YOR344C | 1664.39846 | 0.320733 | 0.06381123 | 5.02627798 | 5.00E-07 |
| AAT1 | YKL106W | 713.867643 | -0.3718448 | 0.07401692 | -5.0237812 | 5.07E-07 |
| Xu_2009_SUTs62 | Xu_2009_SU | 117.381648 | 0.77122387 | 0.15355026 | 5.02261511 | 5.10E-07 |
| DAL5 | YJR152W | 160.943895 | 0.69644182 | 0.13872411 | 5.02033716 | 5.16E-07 |
| YHR219W | YHR219W | 32.1841435 | 1.28748427 | 0.25666427 | 5.01621939 | 5.27E-07 |
| NAS6 | YGR232W | 239.704287 | -0.6014355 | 0.11991949 | -5.0153275 | 5.29E-07 |
| ISN1 | YOR155C | 650.821493 | -0.3982079 | 0.07940831 | -5.0146877 | 5.31E-07 |
| GPB1 | YOR371C | 917.024328 | 0.35820845 | 0.07143496 | 5.01447013 | 5.32E-07 |
| TVP23 | YDR084C | 208.075018 | -0.6260678 | 0.1249271 | -5.011465 | 5.40E-07 |
| VRG4 | YGL225W | 1488.43379 | -0.3122611 | 0.06234028 | -5.0089789 | 5.47E-07 |
| SST2 | YLR452C | 942.731369 | 0.3987429 | 0.07960738 | 5.00886848 | 5.48E-07 |
| MAK21 | YDR060W | 3902.99678 | 0.31777775 | 0.06346548 | 5.00709581 | 5.53E-07 |
| Xu_2009_SUTs31( | Xu_2009_SU | 70.177858 | 0.94388307 | 0.18853089 | 5.00651673 | 5.54E-07 |
| CKB1 | YGL019W | 1369.57811 | -0.3129111 | 0.06250847 | -5.0058987 | 5.56E-07 |
| END3 | YNL084C | 717.386733 | -0.3914887 | 0.07824285 | -5.003507 | 5.63E-07 |
| MES1 | YGR264C | 2359.38369 | -0.3444919 | 0.06895546 | -4.9958609 | 5.86E-07 |
| REH1 | YLR387C | 1359.27671 | 0.31366344 | 0.06278547 | 4.99579696 | 5.86E-07 |
| AHC2 | YCR082W | 1214.68467 | -0.3204693 | 0.06417032 | -4.9940425 | 5.91E-07 |
| RVS167 | YDR388W | 2963.59705 | -0.2667258 | 0.05340813 | -4.9941044 | 5.91E-07 |
| Xu_2009_CUTs31 | Xu_2009_CU | 91.4720096 | 0.82826976 | 0.16586934 | 4.9935073 | 5.93E-07 |
| SNR39B | snR39B | 1035.50162 | 0.3883706 | 0.07779565 | 4.9921889 | 5.97E-07 |

| | | | | | | |
|---|---|---|---|---|---|---|
| Xu_2009_SUTs31 | Xu_2009_SU | 95.874694 | 0.82697027 | 0.16581176 | 4.98740428 | 6.12E-07 |
| NOP19 | YGR251W | 448.143457 | 0.44846111 | 0.09002907 | 4.98129209 | 6.32E-07 |
| NAT1 | YDL040C | 1987.31184 | -0.2994664 | 0.06013651 | -4.9797775 | 6.37E-07 |
| Xu_2009_CUTs84 | Xu_2009_CU | 53.2665432 | 1.10117241 | 0.22139842 | 4.97371385 | 6.57E-07 |
| FCY21 | YER060W | 237.014858 | -0.6034333 | 0.1213544 | -4.972488 | 6.61E-07 |
| TRM732 | YMR259C | 1321.1427 | 0.37323947 | 0.07508349 | 4.9709926 | 6.66E-07 |
| GAD1 | YMR250W | 1744.74682 | -0.4027466 | 0.08103175 | -4.9702313 | 6.69E-07 |
| NPR1 | YNL183C | 881.343178 | -0.3534312 | 0.07112446 | -4.9691936 | 6.72E-07 |
| MGS1 | YNL218W | 496.110629 | 0.4234981 | 0.08529455 | 4.9651247 | 6.87E-07 |
| Xu_2009_CUTs47 | Xu_2009_CU | 64.3269882 | 0.9738519 | 0.1961501 | 4.96483 | 6.88E-07 |
| PEX17 | YNL214W | 126.846163 | -0.732437 | 0.147533 | -4.9645643 | 6.89E-07 |
| HOT1 | YMR172W | 848.371883 | 0.37170076 | 0.07489398 | 4.96302584 | 6.94E-07 |
| YPC1 | YBR183W | 402.248929 | -0.5036761 | 0.10155072 | -4.9598473 | 7.05E-07 |
| RSM10 | YDR041W | 422.28153 | -0.4641384 | 0.09361348 | -4.9580293 | 7.12E-07 |
| YPR145C-A | YPR145C-A | 135.218178 | 0.81417527 | 0.16437368 | 4.95319731 | 7.30E-07 |
| SOL1 | YNR034W | 129.035482 | -0.7274035 | 0.1468878 | -4.9521032 | 7.34E-07 |
| SPF1 | YEL031W | 3833.20389 | -0.2899307 | 0.05856639 | -4.9504628 | 7.40E-07 |
| NUP159 | YIL115C | 2176.5245 | 0.291903 | 0.05899729 | 4.94773542 | 7.51E-07 |
| Xu_2009_CUTs31 | Xu_2009_CU | 99.2733208 | 0.80900805 | 0.16351462 | 4.94761909 | 7.51E-07 |
| IOC2 | YLR095C | 1854.12228 | 0.28832459 | 0.05828151 | 4.94710241 | 7.53E-07 |
| HIR1 | YBL008W | 841.282966 | 0.36763402 | 0.07433846 | 4.94540799 | 7.60E-07 |
| TYW1 | YPL207W | 1877.5221 | 0.34281601 | 0.06933769 | 4.94415095 | 7.65E-07 |
| YMR306C-A | YMR306C-A | 82.3427032 | 0.93210828 | 0.18855214 | 4.94350414 | 7.67E-07 |
| Xu_2009_CUTs76 | Xu_2009_CU | 160.156553 | 0.67182657 | 0.13596576 | 4.94114528 | 7.77E-07 |
| MSE1 | YOL033W | 285.916097 | -0.5733924 | 0.11604876 | -4.9409609 | 7.77E-07 |
| Xu_2009_CUTs6 | Xu_2009_CU | 29.6871502 | 1.20734082 | 0.24451816 | 4.93763248 | 7.91E-07 |
| YPQ1 | YOL092W | 613.22457 | -0.3892449 | 0.07884022 | -4.9371361 | 7.93E-07 |
| Xu_2009_SUTs63 | Xu_2009_SU | 67.5302464 | 0.92178369 | 0.18671091 | 4.93695666 | 7.94E-07 |
| Xu_2009_CUTs79 | Xu_2009_CU | 34.8583639 | 1.17943164 | 0.23895061 | 4.93588045 | 7.98E-07 |
| ENP2 | YGR145W | 1239.28837 | 0.416124 | 0.08432503 | 4.93476276 | 8.02E-07 |
| SKY1 | YMR216C | 3226.46103 | 0.27861174 | 0.05646055 | 4.93462715 | 8.03E-07 |
| Xu_2009_CUTs39 | Xu_2009_CU | 32.4053918 | 1.19917391 | 0.24328811 | 4.92902809 | 8.26E-07 |
| TAF9 | YMR236W | 512.260831 | -0.4098974 | 0.08316289 | -4.9288503 | 8.27E-07 |
| MAK5 | YBR142W | 1936.63691 | 0.39360469 | 0.07992915 | 4.92441956 | 8.46E-07 |
| YKL091C | YKL091C | 425.824679 | -0.574601 | 0.11672665 | -4.9226204 | 8.54E-07 |
| LYS1 | YIR034C | 397.864597 | -0.4918545 | 0.09997356 | -4.919846 | 8.66E-07 |
| GPM2 | YDL021W | 499.972143 | -0.5702358 | 0.11591437 | -4.9194569 | 8.68E-07 |
| OPI1 | YHL020C | 1292.57537 | -0.3271592 | 0.06653245 | -4.9172884 | 8.78E-07 |
| IBD2 | YNL164C | 537.801184 | 0.4427874 | 0.09006123 | 4.91651528 | 8.81E-07 |
| RSN1 | YMR266W | 2281.88895 | -0.3581695 | 0.07286106 | -4.9157868 | 8.84E-07 |
| WSC2 | YNL283C | 1257.30065 | 0.3195034 | 0.06500392 | 4.91514048 | 8.87E-07 |

| | | | | | | |
|---|---|---|---|---|---|---|
| PAA1 | YDR071C | 1250.38446 | -0.3102406 | 0.06313621 | -4.9138293 | 8.93E-07 |
| Xu_2009_SUTs41: | Xu_2009_SU | 85.7781137 | 0.85857746 | 0.17475242 | 4.91310771 | 8.96E-07 |
| RGA1 | YOR127W | 1567.60108 | 0.32249485 | 0.06566337 | 4.91133601 | 9.05E-07 |
| RIM8 | YGL045W | 1350.69863 | 0.33392736 | 0.06805274 | 4.90689058 | 9.25E-07 |
| TPS2 | YDR074W | 2899.952 | -0.4124975 | 0.08407577 | -4.906259 | 9.28E-07 |
| YIL092W | YIL092W | 305.27573 | 0.50757985 | 0.10347301 | 4.90543218 | 9.32E-07 |
| HAL5 | YJL165C | 1519.58766 | 0.35463865 | 0.07229988 | 4.90510731 | 9.34E-07 |
| Xu_2009_CUTs48 | Xu_2009_CU | 29.2940041 | 1.19264295 | 0.24327516 | 4.90244443 | 9.47E-07 |
| MET4 | YNL103W | 2232.71259 | 0.31932669 | 0.06516411 | 4.90034576 | 9.57E-07 |
| SFH5 | YJL145W | 647.218646 | -0.3859262 | 0.07875729 | -4.9001964 | 9.57E-07 |
| PAT1 | YCR077C | 1518.97507 | -0.3199849 | 0.0653292 | -4.8980376 | 9.68E-07 |
| SVS1 | YPL163C | 2305.03868 | -0.3201651 | 0.06544265 | -4.8922998 | 9.97E-07 |
| MIM2 | YLR099W-A | 302.05987 | -0.5362112 | 0.10967102 | -4.8892692 | 1.01E-06 |
| SNR46 | snR46 | 2009.90242 | -0.8861056 | 0.18135211 | -4.886106 | 1.03E-06 |
| Xu_2009_SUTs65 | Xu_2009_SU | 192.262041 | 0.61717322 | 0.12641884 | 4.8819717 | 1.05E-06 |
| FRS1 | YLR060W | 3370.20717 | -0.2534941 | 0.0519366 | -4.880837 | 1.06E-06 |
| MRX1 | YER077C | 1355.81304 | 0.36707411 | 0.07521348 | 4.8804295 | 1.06E-06 |
| YCL012C | YCL012C | 157.353309 | -0.7243872 | 0.14859259 | -4.8749884 | 1.09E-06 |
| MSS116 | YDR194C | 2358.42456 | -0.2675114 | 0.05490842 | -4.8719557 | 1.10E-06 |
| TOD6 | YBL054W | 1578.07997 | 0.40734157 | 0.08370297 | 4.86651303 | 1.14E-06 |
| Xu_2009_SUTs14 | Xu_2009_SU | 33.1973407 | 1.17735949 | 0.242187 | 4.86136527 | 1.17E-06 |
| RDI1 | YDL135C | 1292.61333 | -0.3316837 | 0.0682342 | -4.8609597 | 1.17E-06 |
| Xu_2009_SUTs55 | Xu_2009_SU | 148.764851 | 0.68634355 | 0.14121774 | 4.86017931 | 1.17E-06 |
| FKH2 | YNL068C | 1367.10256 | 0.30697713 | 0.06316168 | 4.86018013 | 1.17E-06 |
| HCH1 | YNL281W | 2717.84281 | -0.2607083 | 0.05365939 | -4.8585781 | 1.18E-06 |
| IST2 | YBR086C | 3838.50626 | -0.2496241 | 0.05147646 | -4.8492858 | 1.24E-06 |
| TIR4 | YOR009W | 374.191053 | 0.54435875 | 0.11231054 | 4.84690706 | 1.25E-06 |
| GEX2 | YKR106W | 173.622676 | 0.67258855 | 0.1387762 | 4.84656996 | 1.26E-06 |
| Xu_2009_CUTs16 | Xu_2009_CU | 48.5788105 | 1.00744413 | 0.20790189 | 4.84576699 | 1.26E-06 |
| SBE2 | YDR351W | 1264.20782 | 0.29881471 | 0.06166466 | 4.84580195 | 1.26E-06 |
| Xu_2009_SUTs40 | Xu_2009_SU | 95.8215415 | 0.79970709 | 0.16505417 | 4.8451189 | 1.27E-06 |
| Xu_2009_CUTs30 | Xu_2009_CU | 48.0061276 | 1.03714394 | 0.2141433 | 4.8432238 | 1.28E-06 |
| RPA14 | YDR156W | 570.170512 | -0.4323113 | 0.08926475 | -4.8430232 | 1.28E-06 |
| DSL1 | YNL258C | 714.469539 | 0.38973283 | 0.08047525 | 4.8428903 | 1.28E-06 |
| DFR1 | YOR236W | 202.60024 | -0.6290514 | 0.12990766 | -4.8422965 | 1.28E-06 |
| MPT5 | YGL178W | 2622.97089 | 0.28025251 | 0.05788193 | 4.84179627 | 1.29E-06 |
| HLJ1 | YMR161W | 650.309799 | -0.3873868 | 0.08000733 | -4.8418913 | 1.29E-06 |
| GCG1 | YER163C | 907.033068 | -0.3538132 | 0.07312767 | -4.8382946 | 1.31E-06 |
| Xu_2009_CUTs18 | Xu_2009_CU | 50.7166483 | 1.05125654 | 0.21743756 | 4.83475134 | 1.33E-06 |
| YDL187C | YDL187C | 85.433138 | 0.82947333 | 0.17164932 | 4.83237181 | 1.35E-06 |
| TIM18 | YOR297C | 216.98728 | -0.5956371 | 0.12329103 | -4.8311469 | 1.36E-06 |

| | | | | | | |
|---|---|---|---|---|---|---|
| LAS21 | YJL062W | 700.623283 | -0.3598246 | 0.07448537 | -4.8308092 | 1.36E-06 |
| ERR3 | YMR323W | 47.1670257 | 1.05042876 | 0.21745793 | 4.83049179 | 1.36E-06 |
| ZDS1 | YMR273C | 1710.36319 | 0.31780189 | 0.06579591 | 4.83011625 | 1.36E-06 |
| tI(AAU)P1 | tI(AAU)P1 | 40.3898754 | -1.1011596 | 0.22799597 | -4.8297329 | 1.37E-06 |
| MRPL7 | YDR237W | 398.248638 | -0.4431694 | 0.09183551 | -4.8256869 | 1.40E-06 |
| Xu_2009_CUTs90 | Xu_2009_CU | 61.1322336 | 1.00786436 | 0.2089391 | 4.82372313 | 1.41E-06 |
| XRN1 | YGL173C | 6290.24488 | 0.23809182 | 0.04937736 | 4.82188216 | 1.42E-06 |
| MGR2 | YPL098C | 482.706343 | -0.4119297 | 0.08562495 | -4.8108599 | 1.50E-06 |
| BPT1 | YLL015W | 1263.16152 | 0.34646101 | 0.07205528 | 4.80826702 | 1.52E-06 |
| ADE12 | YNL220W | 1517.34968 | -0.3608153 | 0.07504553 | -4.8079521 | 1.52E-06 |
| Xu_2009_SUTs73 | Xu_2009_SU | 43.5247776 | 1.07317202 | 0.223287 | 4.80624492 | 1.54E-06 |
| TMN2 | YDR107C | 409.86775 | -0.4638004 | 0.09655087 | -4.80369 | 1.56E-06 |
| Xu_2009_CUTs98 | Xu_2009_CU | 56.7874661 | 0.96596022 | 0.20128514 | 4.79896442 | 1.59E-06 |
| RPO41 | YFL036W | 2548.24639 | -0.2782258 | 0.05799238 | -4.797627 | 1.61E-06 |
| SLI15 | YBR156C | 434.703567 | 0.44994928 | 0.09379621 | 4.79709427 | 1.61E-06 |
| BNR1 | YIL159W | 834.995979 | 0.35321006 | 0.07365092 | 4.79573191 | 1.62E-06 |
| NHP6B | YBR089C-A | 93.7681649 | -0.8364872 | 0.17444325 | -4.7951821 | 1.63E-06 |
| LAA1 | YJL207C | 1686.96606 | 0.28370082 | 0.05916904 | 4.79475155 | 1.63E-06 |
| RMI1 | YPL024W | 744.947828 | 0.39030777 | 0.08140312 | 4.79475195 | 1.63E-06 |
| FCY2 | YER056C | 4185.53995 | -0.3457951 | 0.0721609 | -4.7920014 | 1.65E-06 |
| Xu_2009_SUTs44 | Xu_2009_SU | 109.244837 | 0.80501131 | 0.16800213 | 4.79167313 | 1.65E-06 |
| OM45 | YIL136W | 1012.02712 | 0.48748779 | 0.10176999 | 4.79009346 | 1.67E-06 |
| HIS1 | YER055C | 1499.01559 | -0.3215567 | 0.0671357 | -4.7896524 | 1.67E-06 |
| MCM7 | YBR202W | 1904.27009 | 0.30652733 | 0.06402645 | 4.78751089 | 1.69E-06 |
| DPI8 | YJL133C-A | 266.985429 | -0.575046 | 0.12016256 | -4.7855671 | 1.71E-06 |
| KIN3 | YAR018C | 771.059993 | 0.34945448 | 0.07302444 | 4.78544525 | 1.71E-06 |
| YGR201C | YGR201C | 199.472197 | -0.6694656 | 0.13993013 | -4.7842848 | 1.72E-06 |
| RRP12 | YPL012W | 3184.65381 | 0.3556244 | 0.07433496 | 4.78408035 | 1.72E-06 |
| YNL143C | YNL143C | 86.4982601 | 0.82997699 | 0.17356157 | 4.78203208 | 1.74E-06 |
| YET1 | YKL065C | 1315.63078 | -0.3108645 | 0.06506802 | -4.7775322 | 1.77E-06 |
| PSO2 | YMR137C | 272.644364 | 0.50972024 | 0.10672684 | 4.77593319 | 1.79E-06 |
| GRH1 | YDR517W | 770.613057 | -0.3798938 | 0.07958222 | -4.7736019 | 1.81E-06 |
| Xu_2009_SUTs20 | Xu_2009_SU | 105.395885 | 0.74383208 | 0.15586176 | 4.77238351 | 1.82E-06 |
| AVT6 | YER119C | 796.010153 | -0.3711202 | 0.07777437 | -4.7717544 | 1.83E-06 |
| CMR1 | YDL156W | 294.451124 | 0.49477745 | 0.10373814 | 4.76948431 | 1.85E-06 |
| TMS1 | YDR105C | 559.285948 | -0.3881346 | 0.08139906 | -4.7682936 | 1.86E-06 |
| BFR1 | YOR198C | 3703.37957 | -0.2485142 | 0.05214665 | -4.7656784 | 1.88E-06 |
| ATG39 | YLR312C | 356.888956 | 0.48781015 | 0.10239051 | 4.76421263 | 1.90E-06 |
| COX16 | YJL003W | 96.020583 | -0.7854536 | 0.16491265 | -4.7628463 | 1.91E-06 |
| HIS3 | YOR202W | 742.586905 | -0.4249678 | 0.08922912 | -4.762658 | 1.91E-06 |
| CYC7 | YEL039C | 250.2562 | 0.64081476 | 0.13459953 | 4.760899 | 1.93E-06 |

| | | | | | | |
|---|---|---|---|---|---|---|
| Xu_2009_CUTs48 | Xu_2009_CU | 117.100156 | 0.73184326 | 0.15374393 | 4.76014395 | 1.93E-06 |
| CMI8 | YDR461C-A | 126.66436 | -0.7408473 | 0.15565569 | -4.7595262 | 1.94E-06 |
| Xu_2009_CUTs40 | Xu_2009_CU | 24.3225305 | 1.24258981 | 0.26114448 | 4.7582465 | 1.95E-06 |
| PCF11 | YDR228C | 1214.72545 | 0.31524876 | 0.06625972 | 4.75777397 | 1.96E-06 |
| TRP2 | YER090W | 1243.14785 | -0.2957321 | 0.06216519 | -4.7571965 | 1.96E-06 |
| YLR400W | YLR400W | 26.5996992 | 1.20077708 | 0.25241984 | 4.75706304 | 1.96E-06 |
| Xu_2009_CUTs27 | Xu_2009_CU | 103.956428 | 0.74101716 | 0.15578864 | 4.75655453 | 1.97E-06 |
| BOL1 | YAL044W-A | 333.845473 | -0.5125934 | 0.10786703 | -4.7520861 | 2.01E-06 |
| RSC9 | YML127W | 867.342199 | -0.3490974 | 0.07346182 | -4.7520923 | 2.01E-06 |
| PCL8 | YPL219W | 490.563481 | -0.4045987 | 0.08514346 | -4.751965 | 2.01E-06 |
| YCS4 | YLR272C | 1043.22887 | 0.34228166 | 0.07208648 | 4.74820903 | 2.05E-06 |
| PFA4 | YOL003C | 298.82356 | -0.5365062 | 0.11301819 | -4.7470787 | 2.06E-06 |
| PET100 | YDR079W | 244.418628 | -0.6053838 | 0.12759496 | -4.7445742 | 2.09E-06 |
| GPI16 | YHR188C | 1617.11388 | -0.2996578 | 0.06320028 | -4.7413999 | 2.12E-06 |
| RPN12 | YFR052W | 1097.0341 | -0.3503879 | 0.07397336 | -4.7366771 | 2.17E-06 |
| Xu_2009_SUTs68 | Xu_2009_SU | 272.871796 | 0.49954944 | 0.10551331 | 4.73446864 | 2.20E-06 |
| RTG3 | YBL103C | 1024.50705 | 0.3311992 | 0.06996056 | 4.73408412 | 2.20E-06 |
| VTC1 | YER072W | 672.338594 | -0.4189053 | 0.08854526 | -4.7309735 | 2.23E-06 |
| Xu_2009_SUTs81 | Xu_2009_SU | 152.622557 | 0.66528928 | 0.14064743 | 4.73019142 | 2.24E-06 |
| UBP1 | YDL122W | 3131.21293 | 0.2864153 | 0.06061285 | 4.72532327 | 2.30E-06 |
| CNA1 | YLR433C | 926.450915 | 0.32509622 | 0.06884535 | 4.72212319 | 2.33E-06 |
| SNR14 | snR14 | 600.715357 | -0.836713 | 0.17721319 | -4.7215052 | 2.34E-06 |
| Xu_2009_SUTs84 | Xu_2009_SU | 180.135512 | 0.62832393 | 0.13308537 | 4.72120954 | 2.34E-06 |
| YJL213W | YJL213W | 239.741238 | 0.54464761 | 0.11537015 | 4.72087099 | 2.35E-06 |
| TNA1 | YGR260W | 2499.37396 | -0.3292488 | 0.06975993 | -4.7197412 | 2.36E-06 |
| UBA1 | YKL210W | 6021.3312 | -0.2301654 | 0.04881532 | -4.7150235 | 2.42E-06 |
| CDC42 | YLR229C | 868.301433 | -0.3363064 | 0.07136542 | -4.7124566 | 2.45E-06 |
| YPR158C-D | YPR158C-D | 1891.12825 | 0.29579715 | 0.06277352 | 4.71213258 | 2.45E-06 |
| NAB6 | YML117W | 3545.40984 | 0.26018006 | 0.05523159 | 4.71071079 | 2.47E-06 |
| LSM4 | YER112W | 964.239037 | -0.3167345 | 0.06732716 | -4.7044086 | 2.55E-06 |
| DSC2 | YOL073C | 306.093197 | -0.4939234 | 0.10501604 | -4.7033138 | 2.56E-06 |
| Xu_2009_CUTs73 | Xu_2009_CU | 42.2716962 | 1.07303532 | 0.22825742 | 4.70098764 | 2.59E-06 |
| THI4 | YGR144W | 146.62495 | -0.6643367 | 0.14141348 | -4.6978314 | 2.63E-06 |
| Xu_2009_SUTs26 | Xu_2009_SU | 281.016529 | 0.51103511 | 0.10879822 | 4.69709068 | 2.64E-06 |
| MED7 | YOL135C | 302.467634 | -0.5234562 | 0.11144396 | -4.6970351 | 2.64E-06 |
| BNA7 | YDR428C | 136.184576 | -0.6669305 | 0.14200772 | -4.6964384 | 2.65E-06 |
| Xu_2009_SUTs53 | Xu_2009_SU | 95.946511 | 0.74274326 | 0.15819444 | 4.69512881 | 2.66E-06 |
| YOR041C | YOR041C | 27.6059294 | 1.18827591 | 0.25309237 | 4.69502862 | 2.67E-06 |
| Xu_2009_SUTs33 | Xu_2009_SU | 109.462638 | 0.70512276 | 0.15020097 | 4.69452871 | 2.67E-06 |
| Xu_2009_CUTs74 | Xu_2009_CU | 29.0728632 | 1.1550502 | 0.24604938 | 4.69438374 | 2.67E-06 |
| COX5A | YNL052W | 1486.78196 | -0.3228873 | 0.06878523 | -4.694137 | 2.68E-06 |

| | | | | | | |
|---|---|---|---|---|---|---|
| YJL113W | YJL113W | 309.322603 | 0.51108386 | 0.10892344 | 4.69213854 | 2.70E-06 |
| CCT6 | YDR188W | 1861.06012 | -0.3031557 | 0.06461145 | -4.6919806 | 2.71E-06 |
| SLD7 | YOR060C | 547.084971 | 0.42091168 | 0.08972078 | 4.69135118 | 2.71E-06 |
| YAR1 | YPL239W | 1230.92415 | 0.39201417 | 0.08359293 | 4.68956104 | 2.74E-06 |
| YHR054C | YHR054C | 250.404284 | -0.5412673 | 0.11552221 | -4.6853958 | 2.79E-06 |
| BXI1 | YNL305C | 461.229692 | -0.4387886 | 0.09365769 | -4.6850245 | 2.80E-06 |
| BET3 | YKR068C | 778.463145 | -0.407681 | 0.08703181 | -4.6842763 | 2.81E-06 |
| KDX1 | YKL161C | 569.625282 | 0.38279822 | 0.0817751 | 4.68110969 | 2.85E-06 |
| PRE4 | YFR050C | 1395.56171 | -0.2909042 | 0.06216594 | -4.6794784 | 2.88E-06 |
| RAX2 | YLR084C | 1848.17149 | 0.28359045 | 0.06060966 | 4.6789645 | 2.88E-06 |
| WSS1 | YHR134W | 391.144795 | 0.4776741 | 0.10209224 | 4.67884849 | 2.88E-06 |
| SFI1 | YLL003W | 600.839797 | 0.37679982 | 0.08055539 | 4.67752473 | 2.90E-06 |
| COX19 | YLL018C-A | 269.636117 | 0.51691019 | 0.11059421 | 4.67393538 | 2.95E-06 |
| VPS75 | YNL246W | 505.858129 | -0.4193982 | 0.08974831 | -4.6730488 | 2.97E-06 |
| Xu_2009_SUTs43 | Xu_2009_SU | 153.081532 | 0.64128784 | 0.13730227 | 4.67062792 | 3.00E-06 |
| BUD5 | YCR038C | 375.325422 | 0.45330917 | 0.09708621 | 4.66914083 | 3.02E-06 |
| GIM4 | YEL003W | 297.700734 | -0.490099 | 0.10498052 | -4.6684755 | 3.03E-06 |
| MTG2 | YHR168W | 299.648933 | -0.4751337 | 0.10178398 | -4.6680599 | 3.04E-06 |
| RRP40 | YOL142W | 641.281586 | -0.411978 | 0.08826422 | -4.6675546 | 3.05E-06 |
| SEC65 | YML105C | 556.969494 | -0.4214267 | 0.09030461 | -4.6667245 | 3.06E-06 |
| Xu_2009_SUTs67 | Xu_2009_SU | 165.819544 | 0.6183245 | 0.13252517 | 4.66571369 | 3.08E-06 |
| ECM2 | YBR065C | 569.287011 | 0.41413457 | 0.08877447 | 4.66501893 | 3.09E-06 |
| SEC12 | YNR026C | 588.297336 | 0.40809932 | 0.08756115 | 4.66073524 | 3.15E-06 |
| ESL2 | YKR096W | 1996.50457 | 0.27451573 | 0.05892638 | 4.65862221 | 3.18E-06 |
| PPX1 | YHR201C | 1123.85346 | 0.32059078 | 0.06882415 | 4.65811482 | 3.19E-06 |
| STE6 | YKL209C | 370.865209 | 0.46711932 | 0.10028688 | 4.65783101 | 3.20E-06 |
| YPK3 | YBR028C | 953.671477 | -0.3262907 | 0.07010798 | -4.6541164 | 3.25E-06 |
| SDH5 | YOL071W | 415.020014 | -0.4369533 | 0.09389035 | -4.6538683 | 3.26E-06 |
| YIL161W | YIL161W | 453.483869 | 0.43417608 | 0.09334537 | 4.65128695 | 3.30E-06 |
| YNL097W-A | YNL097W-A | 37.9313353 | 1.0581691 | 0.22755219 | 4.65022605 | 3.32E-06 |
| CYC3 | YAL039C | 1069.89894 | -0.3978039 | 0.08554871 | -4.6500278 | 3.32E-06 |
| URA1 | YKL216W | 2591.15277 | -0.3510821 | 0.0755021 | -4.6499642 | 3.32E-06 |
| GPI11 | YDR302W | 415.326209 | -0.4454285 | 0.09581552 | -4.6488141 | 3.34E-06 |
| YIL080W | YIL080W | 128.335143 | 0.70415377 | 0.15158487 | 4.64527735 | 3.40E-06 |
| LIF1 | YGL090W | 403.009384 | 0.45077725 | 0.09704645 | 4.64496362 | 3.40E-06 |
| YBR096W | YBR096W | 937.670385 | -0.3563137 | 0.07672274 | -4.6441739 | 3.41E-06 |
| ATP4 | YPL078C | 2708.25518 | -0.3207186 | 0.06906472 | -4.6437392 | 3.42E-06 |
| RPC19 | YNL113W | 963.281352 | -0.3259511 | 0.07020412 | -4.6429052 | 3.44E-06 |
| FAF1 | YIL019W | 459.895525 | 0.45272673 | 0.09751433 | 4.64266855 | 3.44E-06 |
| HIR3 | YJR140C | 1073.46486 | 0.32748553 | 0.07055815 | 4.6413563 | 3.46E-06 |
| YFL066C | YFL066C | 54.2915932 | 0.92525283 | 0.19957614 | 4.63608949 | 3.55E-06 |

| | | | | | | |
|---|---|---|---|---|---|---|
| Xu_2009_SUTs51 | Xu_2009_SU | 68.0204694 | 0.93316882 | 0.20129767 | 4.63576554 | 3.56E-06 |
| PRM5 | YIL117C | 616.880031 | 0.39946634 | 0.08616993 | 4.63579769 | 3.56E-06 |
| SEC63 | YOR254C | 2395.04275 | -0.2593014 | 0.0559796 | -4.6320689 | 3.62E-06 |
| SKA1 | YKL023W | 461.382743 | 0.40172522 | 0.08676175 | 4.63021108 | 3.65E-06 |
| Xu_2009_SUTs72 | Xu_2009_SU | 68.6617344 | 0.89216639 | 0.19289715 | 4.62508859 | 3.74E-06 |
| Xu_2009_CUTs84 | Xu_2009_CU | 122.272793 | 0.66782516 | 0.14440426 | 4.62469148 | 3.75E-06 |
| TIP1 | YBR067C | 4085.73425 | -0.3039206 | 0.06574751 | -4.6225419 | 3.79E-06 |
| SPR6 | YER115C | 161.013514 | -0.698531 | 0.15111373 | -4.6225518 | 3.79E-06 |
| AIM29 | YKR074W | 369.631274 | -0.4747222 | 0.10272648 | -4.6212251 | 3.81E-06 |
| FYV7 | YLR068W | 375.822943 | 0.57182963 | 0.123831 | 4.61782299 | 3.88E-06 |
| HFL1 | YKR051W | 476.283669 | -0.3970398 | 0.08602635 | -4.6153273 | 3.92E-06 |
| Xu_2009_CUTs92 | Xu_2009_CU | 67.8306078 | 0.95348757 | 0.20660569 | 4.61501119 | 3.93E-06 |
| Xu_2009_CUTs71 | Xu_2009_CU | 44.5599158 | 1.00122733 | 0.21706569 | 4.6125546 | 3.98E-06 |
| MRPS35 | YGR165W | 677.412836 | -0.3622362 | 0.07853808 | -4.612237 | 3.98E-06 |
| Xu_2009_CUTs11 | Xu_2009_CU | 19.552408 | 1.27083376 | 0.27567642 | 4.60987474 | 4.03E-06 |
| SDS24 | YBR214W | 903.708363 | -0.3824585 | 0.08296935 | -4.6096364 | 4.03E-06 |
| COX10 | YPL172C | 457.353154 | -0.4383007 | 0.09510725 | -4.6084891 | 4.06E-06 |
| Xu_2009_CUTs66 | Xu_2009_CU | 21.0768375 | 1.310032 | 0.28437074 | 4.60677488 | 4.09E-06 |
| Xu_2009_CUTs81 | Xu_2009_CU | 101.897086 | 0.77416959 | 0.16816494 | 4.60363242 | 4.15E-06 |
| PUF3 | YLL013C | 2803.44495 | 0.30767231 | 0.06683083 | 4.60374837 | 4.15E-06 |
| BTS1 | YPL069C | 366.965255 | -0.4515766 | 0.09809212 | -4.6035969 | 4.15E-06 |
| Xu_2009_SUTs15 | Xu_2009_SU | 139.85746 | 0.64507115 | 0.14013745 | 4.60313166 | 4.16E-06 |
| Xu_2009_SUTs42 | Xu_2009_SU | 101.720952 | 0.71968764 | 0.15637441 | 4.60233639 | 4.18E-06 |
| SAM50 | YNL026W | 874.377002 | -0.3578752 | 0.07784411 | -4.5973321 | 4.28E-06 |
| RCR1 | YBR005W | 396.849363 | -0.440345 | 0.09581713 | -4.5956812 | 4.31E-06 |
| SEC21 | YNL287W | 3248.87904 | -0.2470684 | 0.05379567 | -4.5927196 | 4.38E-06 |
| Xu_2009_CUTs73 | Xu_2009_CU | 27.1169073 | 1.18489952 | 0.25800608 | 4.59252551 | 4.38E-06 |
| MCO76 | YPL109C | 353.686963 | 0.43370323 | 0.09444927 | 4.591917 | 4.39E-06 |
| YEL1 | YBL060W | 500.567414 | 0.40135201 | 0.08742998 | 4.59055385 | 4.42E-06 |
| DPC25 | YPL107W | 355.333551 | 0.48175073 | 0.10495562 | 4.59004235 | 4.43E-06 |
| YGR266W | YGR266W | 558.734144 | 0.38360612 | 0.083589 | 4.58919373 | 4.45E-06 |
| ARB1 | YER036C | 6862.13063 | -0.3060448 | 0.06670565 | -4.5879893 | 4.48E-06 |
| Xu_2009_CUTs81 | Xu_2009_CU | 36.5320389 | 1.07521689 | 0.23458952 | 4.58339695 | 4.57E-06 |
| PFU1 | YDR306C | 480.297506 | -0.4113642 | 0.08977305 | -4.5822679 | 4.60E-06 |
| Xu_2009_CUTs68 | Xu_2009_CU | 141.433379 | 0.68137911 | 0.14876188 | 4.580334 | 4.64E-06 |
| TOG1 | YER184C | 450.98221 | 0.41206959 | 0.08999302 | 4.5789061 | 4.67E-06 |
| LNP1 | YHR192W | 317.867837 | -0.4658425 | 0.10178774 | -4.5766075 | 4.73E-06 |
| MRS4 | YKR052C | 265.907379 | -0.4991373 | 0.10910741 | -4.5747336 | 4.77E-06 |
| SEC7 | YDR170C | 6107.4004 | 0.21517421 | 0.04704295 | 4.57399498 | 4.79E-06 |
| MCP2 | YLR253W | 753.547472 | -0.3542464 | 0.0774669 | -4.572874 | 4.81E-06 |
| YGK1 | YGL101W | 333.881311 | -0.4798366 | 0.10493552 | -4.5726806 | 4.82E-06 |

| | | | | | | |
|---|---|---|---|---|---|---|
| RAD30 | YDR419W | 569.228912 | 0.37944312 | 0.08307842 | 4.5672883 | 4.94E-06 |
| MIC27 | YNL100W | 406.967578 | -0.4311438 | 0.09443039 | -4.5657313 | 4.98E-06 |
| Xu_2009_CUTs66 | Xu_2009_CU | 57.7764224 | 0.92145316 | 0.20202535 | 4.56107701 | 5.09E-06 |
| Xu_2009_CUTs49 | Xu_2009_CU | 68.0247147 | 0.85066281 | 0.18662138 | 4.55822809 | 5.16E-06 |
| Xu_2009_SUTs63 | Xu_2009_SU | 58.166455 | 0.89246573 | 0.19586286 | 4.55658491 | 5.20E-06 |
| SCW10 | YMR305C | 4886.7853 | -0.3378629 | 0.07415018 | -4.5564676 | 5.20E-06 |
| TAD2 | YJL035C | 347.569297 | 0.46594988 | 0.10227533 | 4.55583851 | 5.22E-06 |
| PTH2 | YBL057C | 616.262114 | -0.3824927 | 0.08405787 | -4.5503496 | 5.36E-06 |
| ERG1 | YGR175C | 4293.20723 | -0.2556491 | 0.05618485 | -4.5501428 | 5.36E-06 |
| MET17 | YLR303W | 586.266798 | -0.3706737 | 0.08147131 | -4.5497451 | 5.37E-06 |
| YPR089W | YPR089W | 1569.32078 | 0.26997184 | 0.05938235 | 4.54633152 | 5.46E-06 |
| AAD3 | YCR107W | 120.201313 | 0.7042626 | 0.15493152 | 4.54563791 | 5.48E-06 |
| Xu_2009_CUTs46 | Xu_2009_CU | 79.5345371 | 0.85884357 | 0.18898715 | 4.54445491 | 5.51E-06 |
| Xu_2009_SUTs81 | Xu_2009_SU | 101.332005 | 0.70612035 | 0.15540194 | 4.54383228 | 5.52E-06 |
| Xu_2009_CUTs43 | Xu_2009_CU | 112.972576 | 0.72809035 | 0.16036609 | 4.54017637 | 5.62E-06 |
| SLX5 | YDL013W | 905.763761 | 0.34679818 | 0.07641124 | 4.53857573 | 5.66E-06 |
| YPT6 | YLR262C | 207.435948 | -0.544494 | 0.11998968 | -4.5378404 | 5.68E-06 |
| Xu_2009_SUTs43 | Xu_2009_SU | 46.1090887 | 0.96735811 | 0.21322744 | 4.53674299 | 5.71E-06 |
| tK(UUU)D | tK(UUU)D | 46.3100883 | -1.0592582 | 0.23351173 | -4.5362099 | 5.73E-06 |
| DDP1 | YOR163W | 653.625455 | -0.3571086 | 0.07873828 | -4.5353878 | 5.75E-06 |
| NUP2 | YLR335W | 1501.22762 | -0.2925081 | 0.06450001 | -4.5350088 | 5.76E-06 |
| ASI2 | YNL159C | 460.741273 | -0.3946908 | 0.0870345 | -4.5348777 | 5.76E-06 |
| SPC34 | YKR037C | 282.626292 | -0.549315 | 0.12121231 | -4.5318417 | 5.85E-06 |
| VPS71 | YML041C | 309.363449 | -0.4757273 | 0.10508911 | -4.5268946 | 5.99E-06 |
| YPR097W | YPR097W | 1925.29021 | 0.25829756 | 0.05705948 | 4.52681209 | 5.99E-06 |
| NUP100 | YKL068W | 2261.60814 | 0.25999898 | 0.05744398 | 4.52613071 | 6.01E-06 |
| Xu_2009_CUTs13 | Xu_2009_CU | 107.694517 | 0.69682651 | 0.15401355 | 4.5244495 | 6.06E-06 |
| YLL066C | YLL066C | 54.5453147 | 0.91409506 | 0.20213573 | 4.52218447 | 6.12E-06 |
| GAA1 | YLR088W | 782.748142 | -0.3329965 | 0.07363817 | -4.5220646 | 6.12E-06 |
| AGP3 | YFL055W | 228.038994 | 0.53444174 | 0.11819567 | 4.52166937 | 6.14E-06 |
| DAK2 | YFL053W | 27.037165 | 1.15348039 | 0.25511664 | 4.52138443 | 6.14E-06 |
| TRZ1 | YKR079C | 1077.74055 | 0.30888716 | 0.0683486 | 4.5192901 | 6.20E-06 |
| MRPL11 | YDL202W | 372.713095 | -0.4395769 | 0.0973694 | -4.5145284 | 6.35E-06 |
| LDO45 | YMR147W | 365.728901 | 0.43793377 | 0.09701694 | 4.51399292 | 6.36E-06 |
| TOS3 | YGL179C | 1196.00801 | 0.31910887 | 0.07072733 | 4.5118184 | 6.43E-06 |
| PRT1 | YOR361C | 3620.87129 | -0.2514166 | 0.05572946 | -4.5113772 | 6.44E-06 |
| ATG19 | YOL082W | 895.548348 | -0.3464641 | 0.07683716 | -4.5090694 | 6.51E-06 |
| RFC5 | YBR087W | 494.275435 | -0.397968 | 0.08826748 | -4.5086595 | 6.52E-06 |
| YOR015W | YOR015W | 107.108058 | -0.7088011 | 0.1572345 | -4.5079235 | 6.55E-06 |
| MRX14 | YDR115W | 336.387081 | -0.4655191 | 0.1034845 | -4.4984425 | 6.85E-06 |
| CSH1 | YBR161W | 412.842138 | -0.467977 | 0.10407095 | -4.4967108 | 6.90E-06 |

| | | | | | | |
|---|---|---|---|---|---|---|
| LSM6 | YDR378C | 483.844793 | -0.3890407 | 0.08652057 | -4.4965105 | 6.91E-06 |
| Xu_2009_CUTs44 | Xu_2009_CU | 119.922973 | 0.69711504 | 0.15508475 | 4.4950586 | 6.96E-06 |
| MCD1 | YDL003W | 1247.36339 | -0.3390726 | 0.07544656 | -4.4942089 | 6.98E-06 |
| Xu_2009_CUTs35 | Xu_2009_CU | 19.8596337 | 1.24453075 | 0.27697748 | 4.49325607 | 7.01E-06 |
| ATX1 | YNL259C | 76.101112 | -0.8397362 | 0.18691571 | -4.4925927 | 7.04E-06 |
| Xu_2009_CUTs65 | Xu_2009_CU | 13.1997698 | 1.30797449 | 0.29124386 | 4.49099425 | 7.09E-06 |
| SEC62 | YPL094C | 2102.2464 | -0.2602741 | 0.05797345 | -4.4895401 | 7.14E-06 |
| SNU71 | YGR013W | 807.169144 | 0.33857514 | 0.07542 | 4.4891958 | 7.15E-06 |
| QRI5 | YLR204W | 698.506012 | -0.3634474 | 0.08097139 | -4.4885901 | 7.17E-06 |
| ADD66 | YKL206C | 426.109404 | -0.4148802 | 0.09246083 | -4.4870909 | 7.22E-06 |
| Xu_2009_CUTs76 | Xu_2009_CU | 24.8822541 | 1.17403861 | 0.2617956 | 4.48456203 | 7.31E-06 |
| NUP157 | YER105C | 2499.9066 | 0.24413774 | 0.05446254 | 4.48267271 | 7.37E-06 |
| IRC5 | YFR038W | 954.161387 | 0.33314467 | 0.07435812 | 4.48027306 | 7.45E-06 |
| Xu_2009_CUTs12 | Xu_2009_CU | 71.9817854 | 0.82429801 | 0.18399353 | 4.48003809 | 7.46E-06 |
| tF(GAA)P1 | tF(GAA)P1 | 80.5093456 | -0.7924782 | 0.17699712 | -4.4773505 | 7.56E-06 |
| MHO1 | YJR008W | 533.037284 | -0.4574657 | 0.10219457 | -4.476419 | 7.59E-06 |
| CAB3 | YKL088W | 554.389717 | 0.39224207 | 0.08764178 | 4.47551468 | 7.62E-06 |
| SPT15 | YER148W | 1173.2242 | -0.287592 | 0.06434368 | -4.4696236 | 7.84E-06 |
| TRS130 | YMR218C | 905.526748 | 0.37451345 | 0.08379943 | 4.46916434 | 7.85E-06 |
| YIP4 | YGL198W | 292.209738 | -0.4750303 | 0.10644439 | -4.4627085 | 8.09E-06 |
| GRE1 | YPL223C | 170.661287 | 0.57919646 | 0.12983936 | 4.46086946 | 8.16E-06 |
| Xu_2009_SUTs30 | Xu_2009_SU | 291.051778 | 0.48067877 | 0.10779914 | 4.45902225 | 8.23E-06 |
| ATP11 | YNL315C | 600.381521 | -0.3690624 | 0.08278412 | -4.4581298 | 8.27E-06 |
| YGL109W | YGL109W | 19.8450887 | 1.23651638 | 0.27752543 | 4.45550662 | 8.37E-06 |
| Xu_2009_SUTs83 | Xu_2009_SU | 102.290535 | 0.71302163 | 0.16007869 | 4.45419469 | 8.42E-06 |
| Xu_2009_CUTs38 | Xu_2009_CU | 32.545634 | 1.07820893 | 0.24208758 | 4.4537969 | 8.44E-06 |
| YEL045C | YEL045C | 29.2627941 | 1.09750903 | 0.24641624 | 4.45388271 | 8.43E-06 |
| NSA1 | YGL111W | 1535.09543 | 0.32809417 | 0.07367002 | 4.45356427 | 8.45E-06 |
| PSF3 | YOL146W | 271.893805 | -0.5269247 | 0.11834118 | -4.452589 | 8.48E-06 |
| CYT1 | YOR065W | 792.770632 | -0.3230984 | 0.07259399 | -4.4507594 | 8.56E-06 |
| MRPL32 | YCR003W | 331.098785 | -0.4473796 | 0.10052441 | -4.4504572 | 8.57E-06 |
| Xu_2009_CUTs33 | Xu_2009_CU | 105.51163 | 0.72284944 | 0.16252927 | 4.44750325 | 8.69E-06 |
| Xu_2009_CUTs82 | Xu_2009_CU | 46.8499631 | 0.97763924 | 0.21986035 | 4.44663731 | 8.72E-06 |
| RPS28A | YOR167C | 2784.21079 | -0.2816497 | 0.0634677 | -4.4376866 | 9.09E-06 |
| Xu_2009_SUTs66 | Xu_2009_SU | 136.701133 | 0.6582518 | 0.14837747 | 4.43633249 | 9.15E-06 |
| LHS1 | YKL073W | 1163.96127 | -0.2809262 | 0.06333459 | -4.4355884 | 9.18E-06 |
| HSP104 | YLL026W | 7811.51826 | -0.5999344 | 0.13525474 | -4.4355889 | 9.18E-06 |
| ADE1 | YAR015W | 827.374781 | -0.3931714 | 0.08865426 | -4.4348842 | 9.21E-06 |
| NRE1 | YIR035C | 847.492845 | -0.3999796 | 0.09019152 | -4.4347807 | 9.22E-06 |
| SPC1 | YJR010C-A | 130.601551 | -0.6502272 | 0.14670088 | -4.4323333 | 9.32E-06 |
| AQR1 | YNL065W | 2641.36411 | 0.39780778 | 0.08978487 | 4.43067717 | 9.39E-06 |

| | | | | | | |
|---|---|---|---|---|---|---|
| YBR016W | YBR016W | 1167.04134 | -0.3323529 | 0.07504717 | -4.4285863 | 9.49E-06 |
| YHR182C-A | YHR182C-A | 15.2475808 | 1.26076298 | 0.28473294 | 4.42787894 | 9.52E-06 |
| YJL193W | YJL193W | 341.705775 | -0.4528437 | 0.10239861 | -4.4223623 | 9.76E-06 |
| UIP3 | YAR027W | 559.841068 | -0.390227 | 0.08824263 | -4.4222055 | 9.77E-06 |
| SPO14 | YKR031C | 2060.46969 | 0.25645407 | 0.05799308 | 4.42214975 | 9.77E-06 |
| TSA2 | YDR453C | 691.655701 | -0.3714482 | 0.08401634 | -4.4211425 | 9.82E-06 |
| PTR3 | YFR029W | 801.091878 | 0.34133851 | 0.07720835 | 4.42100495 | 9.82E-06 |
| CHS6 | YJL099W | 670.445362 | 0.34450047 | 0.07793353 | 4.42043971 | 9.85E-06 |
| FPS1 | YLL043W | 1501.05062 | -0.2910658 | 0.06589765 | -4.4169379 | 1.00E-05 |
| YMR027W | YMR027W | 1541.18593 | -0.2745206 | 0.06215731 | -4.4165464 | 1.00E-05 |
| UGA2 | YBR006W | 471.77868 | -0.3850657 | 0.08723952 | -4.4138904 | 1.02E-05 |
| SDH1 | YKL148C | 1101.01267 | -0.3368846 | 0.07632306 | -4.4139286 | 1.02E-05 |
| SEC5 | YDR166C | 815.262525 | 0.31670051 | 0.0717591 | 4.41338489 | 1.02E-05 |
| SNR67 | snR67 | 435.707619 | -0.5045952 | 0.11437658 | -4.4117007 | 1.03E-05 |
| URB1 | YKL014C | 3157.63935 | 0.26951343 | 0.0610933 | 4.41150583 | 1.03E-05 |
| PCL5 | YHR071W | 597.291341 | -0.4416 | 0.10011306 | -4.4110131 | 1.03E-05 |
| HLR1 | YDR528W | 396.675132 | -0.4291059 | 0.09730239 | -4.4100238 | 1.03E-05 |
| FMN1 | YDR236C | 209.692477 | -0.5378259 | 0.12195836 | -4.4099139 | 1.03E-05 |
| Xu_2009_SUTs65: | Xu_2009_SU | 63.1967235 | 0.84093688 | 0.19079921 | 4.40744421 | 1.05E-05 |
| PUT4 | YOR348C | 62.2720001 | -0.8958783 | 0.20330046 | -4.4066715 | 1.05E-05 |
| SIW14 | YNL032W | 698.051239 | 0.36867396 | 0.08366771 | 4.4064065 | 1.05E-05 |
| RPN7 | YPR108W | 2178.28172 | -0.2400165 | 0.05447198 | -4.4062389 | 1.05E-05 |
| TIM11 | YDR322C-A | 300.437699 | -0.5134117 | 0.11653647 | -4.4055884 | 1.05E-05 |
| COX8 | YLR395C | 565.414959 | -0.3754976 | 0.08523161 | -4.4056144 | 1.05E-05 |
| TRM8 | YDL201W | 840.725216 | -0.3515553 | 0.07980807 | -4.4050098 | 1.06E-05 |
| SPC97 | YHR172W | 418.72731 | 0.41149227 | 0.09345216 | 4.40323978 | 1.07E-05 |
| Xu_2009_CUTs58 | Xu_2009_CU | 46.4255566 | 0.99423802 | 0.22586306 | 4.40195046 | 1.07E-05 |
| YJR020W | YJR020W | 59.9779884 | 0.8930674 | 0.202914 | 4.40121144 | 1.08E-05 |
| TTI1 | YKL033W | 553.920618 | 0.37530573 | 0.08527136 | 4.40131047 | 1.08E-05 |
| SKI8 | YGL213C | 463.191712 | -0.3794208 | 0.08625232 | -4.398963 | 1.09E-05 |
| TLC1 | TLC1 | 4173.4021 | -0.2703596 | 0.06146284 | -4.3987485 | 1.09E-05 |
| GON7 | YJL184W | 327.017797 | -0.4737294 | 0.10771365 | -4.3980443 | 1.09E-05 |
| ISA2 | YPR067W | 324.807748 | -0.4475274 | 0.10182028 | -4.3952673 | 1.11E-05 |
| YIL012W | YIL012W | 35.8294134 | 1.01022809 | 0.22996643 | 4.3929373 | 1.12E-05 |
| MOB2 | YFL034C-B | 693.519275 | -0.3407402 | 0.07757439 | -4.3924309 | 1.12E-05 |
| FKS1 | YLR342W | 16599.7805 | 0.2281618 | 0.0519436 | 4.39249093 | 1.12E-05 |
| HHT2 | YNL031C | 9730.22671 | -0.2746805 | 0.06253751 | -4.3922517 | 1.12E-05 |
| PFK1 | YGR240C | 8526.25322 | -0.2190861 | 0.04988487 | -4.3918354 | 1.12E-05 |
| TFS1 | YLR178C | 1732.53827 | -0.2993083 | 0.06817891 | -4.3900418 | 1.13E-05 |
| Xu_2009_SUTs7 | Xu_2009_SU | 93.2472951 | 0.75725392 | 0.17251802 | 4.38941933 | 1.14E-05 |
| PRE9 | YGR135W | 1182.27864 | -0.3187599 | 0.07263113 | -4.3887509 | 1.14E-05 |

| | | | | | | |
|---|---|---|---|---|---|---|
| HUG1 | YML058W-A | 63.9073758 | -0.875742 | 0.19971989 | -4.3848512 | 1.16E-05 |
| NSE1 | YLR007W | 725.63257 | 0.34699154 | 0.07914072 | 4.38448789 | 1.16E-05 |
| GAS4 | YOL132W | 125.086062 | -0.6790649 | 0.15487712 | -4.3845399 | 1.16E-05 |
| STE23 | YLR389C | 1992.23765 | 0.25303238 | 0.05772289 | 4.38357101 | 1.17E-05 |
| YJR056C | YJR056C | 514.481963 | 0.38424931 | 0.08766579 | 4.38311576 | 1.17E-05 |
| Xu_2009_SUTs76 | Xu_2009_SU | 66.5850492 | 0.80820568 | 0.18450956 | 4.38029169 | 1.19E-05 |
| SEC15 | YGL233W | 822.481754 | -0.3563636 | 0.08137066 | -4.3795103 | 1.19E-05 |
| ECM5 | YMR176W | 1223.37955 | 0.30229002 | 0.06902676 | 4.37931649 | 1.19E-05 |
| ERG11 | YHR007C | 3801.69373 | -0.2994045 | 0.06839099 | -4.3778348 | 1.20E-05 |
| SWI5 | YDR146C | 1336.76343 | 0.33801722 | 0.07723757 | 4.37633177 | 1.21E-05 |
| APT2 | YDR441C | 388.803488 | -0.4127431 | 0.09431708 | -4.3761226 | 1.21E-05 |
| Xu_2009_SUTs35 | Xu_2009_SU | 77.7006139 | 0.80538011 | 0.18412526 | 4.37408803 | 1.22E-05 |
| Xu_2009_SUTs68 | Xu_2009_SU | 228.60482 | 0.51888972 | 0.11862623 | 4.37415674 | 1.22E-05 |
| SFG1 | YOR315W | 571.90895 | -0.3598221 | 0.08228328 | -4.3729672 | 1.23E-05 |
| HNT2 | YDR305C | 274.617402 | -0.4671168 | 0.10684525 | -4.3719007 | 1.23E-05 |
| YLR252W | YLR252W | 96.4707017 | -0.7599172 | 0.17388424 | -4.370248 | 1.24E-05 |
| RTC1 | YOL138C | 1183.63086 | 0.32393606 | 0.07412473 | 4.37014825 | 1.24E-05 |
| SAY1 | YGR263C | 374.616417 | -0.4175011 | 0.09558201 | -4.3679884 | 1.25E-05 |
| YLR179C | YLR179C | 1490.65308 | -0.3046897 | 0.06977451 | -4.3667773 | 1.26E-05 |
| LYS21 | YDL131W | 1889.98621 | -0.2460353 | 0.05634877 | -4.3662934 | 1.26E-05 |
| VPS24 | YKL041W | 47.773706 | 0.98101905 | 0.22471687 | 4.36557811 | 1.27E-05 |
| Xu_2009_CUTs68 | Xu_2009_CU | 22.4628398 | 1.14689355 | 0.26275698 | 4.36484519 | 1.27E-05 |
| CPA1 | YOR303W | 3747.98041 | 0.28260891 | 0.06475829 | 4.36405775 | 1.28E-05 |
| POM34 | YLR018C | 762.747307 | -0.3196838 | 0.07325726 | -4.3638513 | 1.28E-05 |
| CMR2 | YOR093C | 1182.37747 | 0.30829763 | 0.0706521 | 4.36360194 | 1.28E-05 |
| Xu_2009_CUTs77 | Xu_2009_CU | 27.6097216 | 1.14007863 | 0.26130486 | 4.36302113 | 1.28E-05 |
| YHM2 | YMR241W | 1317.97993 | -0.3183888 | 0.07302187 | -4.3601841 | 1.30E-05 |
| PEX30 | YLR324W | 660.884468 | -0.3646298 | 0.08363622 | -4.3597112 | 1.30E-05 |
| ALY2 | YJL084C | 1859.84531 | 0.26260473 | 0.06028809 | 4.35583135 | 1.33E-05 |
| MRPS8 | YMR158W | 491.498513 | -0.388996 | 0.08931693 | -4.355233 | 1.33E-05 |
| NVJ2 | YPR091C | 1283.97241 | -0.2813379 | 0.06465457 | -4.3514004 | 1.35E-05 |
| YLR126C | YLR126C | 101.128119 | -0.7906919 | 0.18172778 | -4.3509686 | 1.36E-05 |
| FIT3 | YOR383C | 341.363078 | -0.5478428 | 0.12591717 | -4.3508194 | 1.36E-05 |
| MRP13 | YGR084C | 869.946909 | -0.3098961 | 0.07125568 | -4.3490725 | 1.37E-05 |
| RRT14 | YIL127C | 930.19806 | 0.42508892 | 0.09778708 | 4.34708666 | 1.38E-05 |
| LEO1 | YOR123C | 761.636024 | 0.43174636 | 0.0993486 | 4.34577215 | 1.39E-05 |
| Xu_2009_CUTs89 | Xu_2009_CU | 49.3763424 | 0.90977404 | 0.20939658 | 4.34474161 | 1.39E-05 |
| Xu_2009_CUTs27 | Xu_2009_CU | 30.5267998 | 1.05558268 | 0.24296217 | 4.34463806 | 1.40E-05 |
| Xu_2009_CUTs53 | Xu_2009_CU | 79.6150109 | 0.80881723 | 0.18617622 | 4.34436382 | 1.40E-05 |
| CDC28 | YBR160W | 812.008862 | -0.3319339 | 0.0764491 | -4.3418943 | 1.41E-05 |
| OMA1 | YKR087C | 607.86989 | -0.3500258 | 0.08068399 | -4.3382308 | 1.44E-05 |

| | | | | | | |
|---|---|---|---|---|---|---|
| Xu_2009_CUTs29 | Xu_2009_CU | 72.3604946 | 0.82837315 | 0.19100661 | 4.33688205 | 1.45E-05 |
| MRP2 | YPR166C | 154.823526 | -0.6261832 | 0.14449792 | -4.3335102 | 1.47E-05 |
| TIM21 | YGR033C | 959.246012 | -0.2969717 | 0.06854743 | -4.3323542 | 1.48E-05 |
| MLH1 | YMR167W | 484.027523 | 0.38573258 | 0.08903398 | 4.33242002 | 1.47E-05 |
| AFB1 | YLR040C | 1570.1739 | -0.3219064 | 0.07433835 | -4.330287 | 1.49E-05 |
| DCK1 | YLR422W | 1849.10256 | 0.26215159 | 0.06058537 | 4.32697825 | 1.51E-05 |
| YKR104W | YKR104W | 92.0478971 | 0.74355135 | 0.17184701 | 4.32682144 | 1.51E-05 |
| NOP7 | YGR103W | 4672.6669 | 0.30117762 | 0.06961084 | 4.32659098 | 1.51E-05 |
| YBR285W | YBR285W | 50.2207162 | -0.9014659 | 0.20841211 | -4.3254006 | 1.52E-05 |
| Xu_2009_CUTs36 | Xu_2009_CU | 42.148448 | 0.96833155 | 0.22391321 | 4.32458422 | 1.53E-05 |
| Xu_2009_CUTs11 | Xu_2009_CU | 24.0984095 | 1.12305336 | 0.25999198 | 4.31956931 | 1.56E-05 |
| TRL1 | YJL087C | 853.995401 | 0.34106725 | 0.07896834 | 4.31903796 | 1.57E-05 |
| EMC3 | YKL207W | 1125.97272 | -0.327533 | 0.07588969 | -4.3159088 | 1.59E-05 |
| MEU1 | YLR017W | 1141.5653 | -0.3148495 | 0.07295433 | -4.3157063 | 1.59E-05 |
| LYS14 | YDR034C | 1022.68812 | 0.33495707 | 0.07761974 | 4.31535942 | 1.59E-05 |
| Xu_2009_CUTs63 | Xu_2009_CU | 46.5290533 | 0.92829318 | 0.21516861 | 4.31425926 | 1.60E-05 |
| Xu_2009_SUTs50 | Xu_2009_SU | 37.3137611 | 0.9776513 | 0.22671222 | 4.3123008 | 1.62E-05 |
| Xu_2009_CUTs90 | Xu_2009_CU | 99.6894671 | 0.69277663 | 0.16065759 | 4.31213146 | 1.62E-05 |
| YOR072W-A | YOR072W-A | 28.3806603 | 1.07200563 | 0.2486116 | 4.31196942 | 1.62E-05 |
| RAD61 | YDR014W | 453.885491 | 0.39195014 | 0.09091263 | 4.31128364 | 1.62E-05 |
| REG1 | YDR028C | 2987.88118 | 0.25127992 | 0.05828967 | 4.31088275 | 1.63E-05 |
| GPI14 | YJR013W | 447.503931 | -0.3789825 | 0.08791342 | -4.3108603 | 1.63E-05 |
| Xu_2009_CUTs46 | Xu_2009_CU | 26.6643455 | 1.10575131 | 0.25674115 | 4.30687212 | 1.66E-05 |
| PHO87 | YCR037C | 954.203682 | 0.30949222 | 0.07186857 | 4.30636382 | 1.66E-05 |
| ASR1 | YPR093C | 311.131778 | 0.45146702 | 0.10485057 | 4.30581377 | 1.66E-05 |
| SCJ1 | YMR214W | 747.608585 | -0.3387159 | 0.07867697 | -4.3051468 | 1.67E-05 |
| YHR071C-A | YHR071C-A | 78.6087487 | 0.78089085 | 0.18146493 | 4.30326056 | 1.68E-05 |
| PPH21 | YDL134C | 1356.45412 | -0.2976226 | 0.06917168 | -4.3026663 | 1.69E-05 |
| ICR1 | ICR1 | 347.042824 | 0.4467494 | 0.10383882 | 4.30233498 | 1.69E-05 |
| YBR072C-A | YBR072C-A | 13.7477537 | 1.24192764 | 0.2886621 | 4.30235778 | 1.69E-05 |
| BSC2 | YDR275W | 529.863626 | 0.39088021 | 0.09086664 | 4.30169096 | 1.69E-05 |
| TAF6 | YGL112C | 1002.29672 | -0.2874357 | 0.06682429 | -4.3013657 | 1.70E-05 |
| FSH3 | YOR280C | 466.519233 | -0.3868769 | 0.08994458 | -4.3012807 | 1.70E-05 |
| Xu_2009_CUTs23 | Xu_2009_CU | 41.2340152 | 0.99507486 | 0.23143021 | 4.29967569 | 1.71E-05 |
| HMX1 | YLR205C | 315.884041 | 0.42985287 | 0.10000091 | 4.29848978 | 1.72E-05 |
| Xu_2009_CUTs18 | Xu_2009_CU | 65.408081 | 0.82965819 | 0.19306221 | 4.29736191 | 1.73E-05 |
| PSY4 | YBL046W | 452.747548 | 0.41323821 | 0.09615963 | 4.29741876 | 1.73E-05 |
| AIM17 | YHL021C | 805.405982 | -0.6523366 | 0.15180592 | -4.2971747 | 1.73E-05 |
| SUR2 | YDR297W | 1972.38954 | -0.2841675 | 0.06613769 | -4.2966054 | 1.73E-05 |
| PIN3 | YPR154W | 1136.91311 | -0.2935233 | 0.06834054 | -4.2950093 | 1.75E-05 |
| CNS1 | YBR155W | 930.174446 | 0.33307592 | 0.07757148 | 4.29379349 | 1.76E-05 |

| | | | | | | |
|---|---|---|---|---|---|---|
| Xu_2009_CUTs52 | Xu_2009_CU | 8.73437357 | 1.26729053 | 0.29516056 | 4.29356321 | 1.76E-05 |
| YLR120W-A | YLR120W-A | 35.7013215 | 1.0105645 | 0.2353659 | 4.29358931 | 1.76E-05 |
| IVY1 | YDR229W | 1011.88656 | -0.2976614 | 0.06934631 | -4.2923899 | 1.77E-05 |
| SRP54 | YPR088C | 1236.72824 | -0.2809 | 0.06548786 | -4.2893442 | 1.79E-05 |
| VPS41 | YDR080W | 814.308235 | 0.30592119 | 0.07136674 | 4.28660767 | 1.81E-05 |
| COS10 | YNR075W | 439.574831 | 0.50001199 | 0.11666687 | 4.2858096 | 1.82E-05 |
| Xu_2009_CUTs47 | Xu_2009_CU | 35.8341662 | 0.98513225 | 0.22997004 | 4.28374164 | 1.84E-05 |
| DNF3 | YMR162C | 1675.09455 | 0.25704602 | 0.06000454 | 4.28377598 | 1.84E-05 |
| FYV8 | YGR196C | 792.992464 | 0.35916275 | 0.08385782 | 4.28299655 | 1.84E-05 |
| ATP1 | YBL099W | 3712.77132 | -0.2756784 | 0.06437964 | -4.2820748 | 1.85E-05 |
| NIS1 | YNL078W | 971.162502 | -0.3778607 | 0.08828575 | -4.2799738 | 1.87E-05 |
| Xu_2009_SUTs25 | Xu_2009_SU | 123.957003 | 0.66196073 | 0.15490919 | 4.27321805 | 1.93E-05 |
| GLE2 | YER107C | 378.595 | -0.4220181 | 0.09875784 | -4.273262 | 1.93E-05 |
| YHR022C | YHR022C | 151.406142 | 0.68112151 | 0.15952176 | 4.26977171 | 1.96E-05 |
| OPY1 | YBR129C | 329.494289 | -0.4730795 | 0.11087819 | -4.2666601 | 1.98E-05 |
| UTP4 | YDR324C | 2049.30288 | 0.32654245 | 0.07657213 | 4.26450791 | 2.00E-05 |
| DNL4 | YOR005C | 410.049171 | 0.39125992 | 0.09175771 | 4.26405484 | 2.01E-05 |
| DER1 | YBR201W | 334.172182 | -0.4977905 | 0.11682068 | -4.2611504 | 2.03E-05 |
| SIT1 | YEL065W | 274.242209 | -0.5266849 | 0.12363659 | -4.2599434 | 2.04E-05 |
| PKH3 | YDR466W | 1532.48727 | 0.26363465 | 0.0618899 | 4.25973624 | 2.05E-05 |
| SRP102 | YKL154W | 360.240944 | -0.4231435 | 0.099339 | -4.2595905 | 2.05E-05 |
| SRP72 | YPL210C | 1851.20745 | -0.2575468 | 0.06047169 | -4.2589646 | 2.05E-05 |
| Xu_2009_CUTs25 | Xu_2009_CU | 235.599518 | 0.51626895 | 0.12123647 | 4.25836346 | 2.06E-05 |
| MSB3 | YNL293W | 474.327829 | 0.39780413 | 0.09342147 | 4.25816622 | 2.06E-05 |
| Xu_2009_SUTs35 | Xu_2009_SU | 17.3738492 | 1.18933558 | 0.27935897 | 4.25737381 | 2.07E-05 |
| YLR463C | YLR463C | 40.2372149 | 0.99478398 | 0.23370906 | 4.25650581 | 2.08E-05 |
| THI72 | YOR192C | 132.32517 | 0.61180798 | 0.14382373 | 4.25387381 | 2.10E-05 |
| Xu_2009_SUTs15 | Xu_2009_SU | 170.130243 | 0.53569083 | 0.12593567 | 4.25368639 | 2.10E-05 |
| ATG7 | YHR171W | 256.074585 | -0.4897529 | 0.11521028 | -4.2509473 | 2.13E-05 |
| GCD7 | YLR291C | 888.149329 | -0.3224278 | 0.07590138 | -4.2479838 | 2.16E-05 |
| Xu_2009_SUTs18 | Xu_2009_SU | 293.247065 | 0.42714637 | 0.10056254 | 4.24756949 | 2.16E-05 |
| SNF3 | YDL194W | 368.459578 | -0.4005983 | 0.09431742 | -4.2473416 | 2.16E-05 |
| Xu_2009_CUTs75 | Xu_2009_CU | 10.8814097 | 1.25195844 | 0.2948516 | 4.24606291 | 2.18E-05 |
| RTP1 | YMR185W | 660.809773 | 0.32849375 | 0.07737265 | 4.24560585 | 2.18E-05 |
| SSN2 | YDR443C | 1204.9546 | 0.27564355 | 0.06496495 | 4.24295806 | 2.21E-05 |
| ENT2 | YLR206W | 2653.32982 | 0.30658638 | 0.07229135 | 4.24098298 | 2.23E-05 |
| tV(AAC)K2 | tV(AAC)K2 | 68.366141 | 0.84657433 | 0.19963702 | 4.24056775 | 2.23E-05 |
| BUD7 | YOR299W | 1563.93739 | 0.25292947 | 0.05972064 | 4.23520999 | 2.28E-05 |
| Xu_2009_CUTs42 | Xu_2009_CU | 22.4646432 | 1.1371108 | 0.26860252 | 4.23343314 | 2.30E-05 |
| RFC2 | YJR068W | 907.94291 | -0.2883521 | 0.06812577 | -4.2326433 | 2.31E-05 |
| Xu_2009_CUTs49 | Xu_2009_CU | 66.134084 | 0.80338671 | 0.18981706 | 4.23242626 | 2.31E-05 |

| | | | | | | |
|---|---|---|---|---|---|---|
| ADE3 | YGR204W | 4584.48016 | -0.2252515 | 0.05323608 | -4.231181 | 2.32E-05 |
| PRE10 | YOR362C | 2133.01122 | -0.2390574 | 0.05651058 | -4.2303126 | 2.33E-05 |
| DCW1 | YKL046C | 861.092815 | -0.3064166 | 0.07243583 | -4.2301793 | 2.34E-05 |
| ODC2 | YOR222W | 1020.5762 | -0.3092615 | 0.0731173 | -4.2296616 | 2.34E-05 |
| QCR10 | YHR001W-A | 484.66493 | -0.4082907 | 0.09657269 | -4.2278073 | 2.36E-05 |
| MLO1 | YMR252C | 236.054141 | -0.4828331 | 0.1142835 | -4.2248712 | 2.39E-05 |
| CMP2 | YML057W | 754.197563 | -0.3190496 | 0.07562599 | -4.2187825 | 2.46E-05 |
| SSM4 | YIL030C | 2457.25172 | 0.26860633 | 0.06367379 | 4.21847544 | 2.46E-05 |
| CDC23 | YHR166C | 327.55139 | 0.43241734 | 0.10251943 | 4.21790614 | 2.47E-05 |
| SMF1 | YOL122C | 496.088896 | -0.4262838 | 0.10106724 | -4.217824 | 2.47E-05 |
| RPR2 | YIR015W | 319.0295 | 0.43931857 | 0.10432859 | 4.21091245 | 2.54E-05 |
| Xu_2009_CUTs69 | Xu_2009_CU | 21.6704494 | 1.14380901 | 0.27168709 | 4.21002336 | 2.55E-05 |
| MCH1 | YDL054C | 446.237173 | -0.4111938 | 0.0976876 | -4.209273 | 2.56E-05 |
| RCR2 | YDR003W | 323.77251 | -0.4499807 | 0.10691104 | -4.2089263 | 2.57E-05 |
| COG2 | YGR120C | 265.759951 | -0.483115 | 0.11479326 | -4.2085657 | 2.57E-05 |
| ASI1 | YMR119W | 554.318187 | -0.3493339 | 0.08302012 | -4.2078228 | 2.58E-05 |
| Xu_2009_SUTs57 | Xu_2009_SU | 110.35617 | 0.65155729 | 0.15486409 | 4.20728455 | 2.58E-05 |
| FMP41 | YNL168C | 1060.10317 | -0.306216 | 0.07283638 | -4.204163 | 2.62E-05 |
| CTL1 | YMR180C | 400.75397 | 0.39595613 | 0.09420826 | 4.2029876 | 2.63E-05 |
| YDL183C | YDL183C | 264.949481 | -0.4503393 | 0.10716343 | -4.2023599 | 2.64E-05 |
| STT4 | YLR305C | 1929.26401 | 0.25017312 | 0.05954835 | 4.20117614 | 2.66E-05 |
| HPA2 | YPR193C | 53.0732878 | 0.86122975 | 0.20519403 | 4.19714828 | 2.70E-05 |
| YCR087C-A | YCR087C-A | 595.427023 | 0.45036491 | 0.10732838 | 4.19614017 | 2.72E-05 |
| TRM10 | YOL093W | 472.534824 | 0.39916177 | 0.09512459 | 4.19619952 | 2.71E-05 |
| TAE1 | YBR261C | 568.366865 | -0.3984563 | 0.09497993 | -4.1951635 | 2.73E-05 |
| SNF1 | YDR477W | 1344.15076 | -0.2702919 | 0.0644491 | -4.1938815 | 2.74E-05 |
| DGK1 | YOR311C | 540.728007 | -0.3615956 | 0.08636813 | -4.1866792 | 2.83E-05 |
| ISD11 | YER048W-A | 846.324275 | -0.3285327 | 0.07849923 | -4.1851703 | 2.85E-05 |
| OCA5 | YHL029C | 1677.7592 | 0.27694078 | 0.06618612 | 4.18427276 | 2.86E-05 |
| MST1 | YKL194C | 400.457665 | -0.4362549 | 0.10430763 | -4.1823873 | 2.88E-05 |
| AYT1 | YLL063C | 161.854735 | 0.6042179 | 0.1445355 | 4.18041159 | 2.91E-05 |
| RNT1 | YMR239C | 549.040193 | 0.39833611 | 0.0953272 | 4.17861966 | 2.93E-05 |
| YNR014W | YNR014W | 278.843211 | 0.46545635 | 0.11142152 | 4.17743681 | 2.95E-05 |
| VPS54 | YDR027C | 833.333875 | 0.33560637 | 0.08036014 | 4.17627914 | 2.96E-05 |
| RNA15 | YGL044C | 729.52029 | 0.33191792 | 0.07949746 | 4.17520148 | 2.98E-05 |
| EMA19 | YLR050C | 135.937481 | -0.6188106 | 0.14823226 | -4.1746015 | 2.99E-05 |
| Xu_2009_SUTs35 | Xu_2009_SU | 18.7657865 | 1.14550807 | 0.27445953 | 4.1736866 | 3.00E-05 |
| RTC2 | YBR147W | 94.7655579 | -0.6789167 | 0.16266701 | -4.1736594 | 3.00E-05 |
| EMG1 | YLR186W | 868.485482 | -0.3767015 | 0.09025783 | -4.1736155 | 3.00E-05 |
| tR(UCU)G2 | tR(UCU)G2 | 77.7912996 | -0.7618844 | 0.18273327 | -4.1693798 | 3.05E-05 |
| SUB1 | YMR039C | 1121.8119 | 0.29533359 | 0.07085268 | 4.16827699 | 3.07E-05 |

| | | | | | | |
|---|---|---|---|---|---|---|
| YGR161C-D | YGR161C-D | 419.12918 | 0.40428053 | 0.09699357 | 4.16811698 | 3.07E-05 |
| DBF4 | YDR052C | 1451.20834 | 0.26918638 | 0.06460255 | 4.16680724 | 3.09E-05 |
| PRE6 | YOL038W | 2694.75436 | -0.2369178 | 0.05686155 | -4.1665736 | 3.09E-05 |
| SIT4 | YDL047W | 1078.1288 | -0.27384 | 0.06572863 | -4.1662216 | 3.10E-05 |
| Xu_2009_CUTs38 | Xu_2009_CU | 44.9556666 | 0.90894667 | 0.21822107 | 4.1652563 | 3.11E-05 |
| Xu_2009_SUTs52 | Xu_2009_SU | 390.298711 | 0.45129349 | 0.10840006 | 4.16322159 | 3.14E-05 |
| NOP16 | YER002W | 818.281928 | 0.36787172 | 0.08835824 | 4.16341163 | 3.14E-05 |
| ESS1 | YJR017C | 747.607563 | -0.3134115 | 0.07528049 | -4.1632494 | 3.14E-05 |
| SGS1 | YMR190C | 1838.85449 | 0.25462263 | 0.06115569 | 4.16351487 | 3.13E-05 |
| RFA1 | YAR007C | 886.881584 | -0.3012352 | 0.07236521 | -4.1627071 | 3.14E-05 |
| GCN4 | YEL009C | 2806.63848 | -0.3113861 | 0.07480837 | -4.1624501 | 3.15E-05 |
| MVB12 | YGR206W | 132.597008 | -0.5966625 | 0.14336489 | -4.1618453 | 3.16E-05 |
| TML25 | YLR118C | 492.485947 | -0.4036166 | 0.09698033 | -4.1618401 | 3.16E-05 |
| CMC1 | YKL137W | 242.994125 | -0.4761501 | 0.11445099 | -4.1602965 | 3.18E-05 |
| BYE1 | YKL005C | 995.57601 | 0.29297562 | 0.07042748 | 4.15996193 | 3.18E-05 |
| REI1 | YBR267W | 1697.29697 | 0.35688382 | 0.08581295 | 4.15885752 | 3.20E-05 |
| BOP2 | YLR267W | 128.960039 | -0.6398229 | 0.15385405 | -4.1586357 | 3.20E-05 |
| YDL221W | YDL221W | 38.5152184 | 0.94079463 | 0.22638715 | 4.15568913 | 3.24E-05 |
| ARC19 | YKL013C | 823.887444 | -0.3601566 | 0.08666877 | -4.1555518 | 3.25E-05 |
| YAT2 | YER024W | 544.654875 | 0.37859855 | 0.09114266 | 4.15391172 | 3.27E-05 |
| Xu_2009_SUTs78 | Xu_2009_SU | 37.1695181 | 0.97841773 | 0.23560752 | 4.15274413 | 3.29E-05 |
| ERV14 | YGL054C | 789.284325 | -0.3231699 | 0.07783452 | -4.1520129 | 3.30E-05 |
| ARP8 | YOR141C | 1066.47491 | 0.33163648 | 0.07995928 | 4.1475671 | 3.36E-05 |
| SET1 | YHR119W | 1342.33991 | 0.27046676 | 0.065258 | 4.14457622 | 3.40E-05 |
| PPH3 | YDR075W | 963.998989 | 0.31293683 | 0.07556117 | 4.14150325 | 3.45E-05 |
| UTR4 | YEL038W | 306.606841 | -0.4974721 | 0.12019595 | -4.1388425 | 3.49E-05 |
| COX18 | YGR062C | 202.255012 | -0.5180528 | 0.12518939 | -4.1381521 | 3.50E-05 |
| Xu_2009_SUTs32 | Xu_2009_SU | 81.9771803 | 0.72896652 | 0.17620726 | 4.13698354 | 3.52E-05 |
| RSM7 | YJR113C | 623.119904 | -0.3843053 | 0.0928993 | -4.1367943 | 3.52E-05 |
| RAD26 | YJR035W | 760.974423 | 0.32283367 | 0.0780691 | 4.13523011 | 3.55E-05 |
| KRE9 | YJL174W | 2017.8929 | -0.2365414 | 0.05721377 | -4.1343448 | 3.56E-05 |
| TAF8 | YML114C | 1312.36449 | 0.25699325 | 0.06217015 | 4.13370787 | 3.57E-05 |
| RAD7 | YJR052W | 575.502866 | 0.35130739 | 0.08505648 | 4.13028349 | 3.62E-05 |
| IES1 | YFL013C | 1277.42764 | 0.25319741 | 0.06134121 | 4.12768887 | 3.66E-05 |
| NIP7 | YPL211W | 393.788299 | -0.4101514 | 0.09936625 | -4.1276734 | 3.66E-05 |
| YKL202W | YKL202W | 226.416313 | 0.4966522 | 0.12034605 | 4.12686743 | 3.68E-05 |
| SPC2 | YML055W | 801.670044 | -0.3267768 | 0.07921206 | -4.1253416 | 3.70E-05 |
| YEN1 | YER041W | 629.87044 | 0.33960319 | 0.08240583 | 4.12110643 | 3.77E-05 |
| YIR043C | YIR043C | 19.894487 | 1.11763089 | 0.2715019 | 4.11647537 | 3.85E-05 |
| GDA1 | YEL042W | 1683.64493 | -0.2924893 | 0.07105857 | -4.1161716 | 3.85E-05 |
| MSP1 | YGR028W | 356.217958 | -0.4201331 | 0.10210931 | -4.1145422 | 3.88E-05 |

| | | | | | | |
|---|---|---|---|---|---|---|
| Xu_2009_CUTs72 | Xu_2009_CU | 31.3078886 | 1.00680299 | 0.24475491 | 4.11351498 | 3.90E-05 |
| YOL163W | YOL163W | 60.9614261 | 0.80776231 | 0.19644273 | 4.11194799 | 3.92E-05 |
| MRP21 | YBL090W | 224.95248 | -0.5059376 | 0.12309357 | -4.1101869 | 3.95E-05 |
| YLR278C | YLR278C | 1380.98962 | 0.25778905 | 0.0627387 | 4.10893183 | 3.97E-05 |
| KEI1 | YDR367W | 185.856547 | -0.5260827 | 0.12806844 | -4.107825 | 3.99E-05 |
| YDL085C-A | YDL085C-A | 280.564456 | -0.4576244 | 0.11146177 | -4.1056623 | 4.03E-05 |
| SRB2 | YHR041C | 207.717616 | -0.4977677 | 0.12124684 | -4.1054077 | 4.04E-05 |
| SSF2 | YDR312W | 861.617575 | 0.28726361 | 0.07002347 | 4.10239005 | 4.09E-05 |
| CDC12 | YHR107C | 1347.84309 | -0.2621275 | 0.06391609 | -4.1011196 | 4.11E-05 |
| GCN20 | YFR009W | 2528.01123 | -0.2164612 | 0.05278724 | -4.1006347 | 4.12E-05 |
| RBG2 | YGR173W | 613.249899 | -0.3486256 | 0.08501824 | -4.1005985 | 4.12E-05 |
| SLD5 | YDR489W | 429.378882 | 0.40210548 | 0.09810052 | 4.09891278 | 4.15E-05 |
| Xu_2009_SUTs64 | Xu_2009_SU | 133.584741 | 0.60822578 | 0.14846268 | 4.09682606 | 4.19E-05 |
| LIP2 | YLR239C | 471.318742 | -0.3660716 | 0.08937203 | -4.0960421 | 4.20E-05 |
| MGP12 | YDR286C | 91.5900262 | -0.7134972 | 0.17421569 | -4.095482 | 4.21E-05 |
| YNL190W | YNL190W | 9637.55125 | -0.1851179 | 0.0452172 | -4.0939723 | 4.24E-05 |
| IRC7 | YFR055W | 799.814814 | 0.33346213 | 0.08151085 | 4.09101549 | 4.29E-05 |
| ANB1 | YJR047C | 39.8919373 | -0.9363422 | 0.22891092 | -4.0904216 | 4.31E-05 |
| RPN3 | YER021W | 2524.25668 | -0.2313352 | 0.05665822 | -4.0829942 | 4.45E-05 |
| Xu_2009_SUTs70 | Xu_2009_SU | 184.030397 | 0.57823909 | 0.14163408 | 4.0826268 | 4.45E-05 |
| PMP1 | YCR024C-A | 2221.82887 | -0.238837 | 0.05851318 | -4.0817641 | 4.47E-05 |
| ZTA1 | YBR046C | 560.71949 | -0.3503261 | 0.08585951 | -4.0802252 | 4.50E-05 |
| AIM46 | YHR199C | 476.531413 | -0.359944 | 0.08821893 | -4.0801214 | 4.50E-05 |
| BIO4 | YNR057C | 249.300822 | -0.5126728 | 0.12573082 | -4.0775433 | 4.55E-05 |
| Xu_2009_SUTs65 | Xu_2009_SU | 144.031582 | 0.58552221 | 0.14365589 | 4.07586646 | 4.58E-05 |
| MIN6 | YBL039W-B | 98.1431273 | -0.6852446 | 0.16816375 | -4.0748651 | 4.60E-05 |
| tL(CAA)L | tL(CAA)L | 26.2091937 | -1.0440657 | 0.25627182 | -4.0740559 | 4.62E-05 |
| RIA1 | YNL163C | 1979.19136 | 0.27025349 | 0.06637426 | 4.07166092 | 4.67E-05 |
| BUL1 | YMR275C | 1368.2487 | 0.27869781 | 0.06848941 | 4.06921006 | 4.72E-05 |
| MKK2 | YPL140C | 497.51087 | 0.34731381 | 0.08537321 | 4.06818238 | 4.74E-05 |
| GLD1 | YPR109W | 243.529732 | -0.499756 | 0.1230263 | -4.0621881 | 4.86E-05 |
| YMR245W | YMR245W | 66.3957541 | 0.78317078 | 0.19285224 | 4.06098888 | 4.89E-05 |
| RFA3 | YJL173C | 318.743914 | -0.4628766 | 0.11399145 | -4.0606251 | 4.89E-05 |
| RTS2 | YOR077W | 228.470728 | 0.52299011 | 0.12884018 | 4.0592159 | 4.92E-05 |
| DSS4 | YPR017C | 192.433955 | -0.5199773 | 0.12813601 | -4.0580105 | 4.95E-05 |
| GTO1 | YGR154C | 186.746054 | 0.5295752 | 0.13050575 | 4.05786874 | 4.95E-05 |
| SEC26 | YDR238C | 3337.90415 | -0.2033478 | 0.05011787 | -4.0573911 | 4.96E-05 |
| YJU2 | YKL095W | 454.688672 | 0.35375437 | 0.08723387 | 4.05524117 | 5.01E-05 |
| CIP1 | YPL014W | 941.243167 | 0.34419993 | 0.08491026 | 4.05369084 | 5.04E-05 |
| LGE1 | YPL055C | 583.458837 | 0.34948151 | 0.08626171 | 4.05140952 | 5.09E-05 |
| SAP4 | YGL229C | 214.088193 | -0.4860683 | 0.12000262 | -4.0504804 | 5.11E-05 |

| | | | | | | |
|---|---|---|---|---|---|---|
| PET54 | YGR222W | 552.204246 | 0.34479949 | 0.08513327 | 4.05011457 | 5.12E-05 |
| CIR1 | YGR207C | 546.311539 | -0.3333085 | 0.08231152 | -4.0493548 | 5.14E-05 |
| SRO7 | YPR032W | 1267.96454 | 0.26557047 | 0.06571333 | 4.0413487 | 5.31E-05 |
| SRB8 | YCR081W | 976.614493 | 0.28664089 | 0.07098587 | 4.03799945 | 5.39E-05 |
| YPT32 | YGL210W | 424.971123 | -0.3707012 | 0.09182407 | -4.0370807 | 5.41E-05 |
| CHZ1 | YER030W | 1323.2203 | -0.3163341 | 0.07837048 | -4.0363935 | 5.43E-05 |
| GLG1 | YKR058W | 525.542822 | -0.3519507 | 0.0872681 | -4.0329827 | 5.51E-05 |
| AAD15 | YOL165C | 17.4821171 | 1.12213087 | 0.2782907 | 4.03222561 | 5.53E-05 |
| CYC1 | YJR048W | 359.411405 | -0.4902092 | 0.12157871 | -4.0320314 | 5.53E-05 |
| MRPL6 | YHR147C | 380.265489 | -0.4185249 | 0.1038265 | -4.031003 | 5.55E-05 |
| Xu_2009_SUTs66 | Xu_2009_SU | 577.52014 | 0.37569819 | 0.09321311 | 4.03052934 | 5.57E-05 |
| Xu_2009_CUTs65 | Xu_2009_CU | 23.0705 | 1.06184064 | 0.26357387 | 4.02862635 | 5.61E-05 |
| Xu_2009_CUTs79 | Xu_2009_CU | 20.1074671 | 1.12526054 | 0.27931113 | 4.02869926 | 5.61E-05 |
| PRE5 | YMR314W | 1281.23351 | -0.2667675 | 0.06621501 | -4.0288064 | 5.61E-05 |
| RPN6 | YDL097C | 1876.53512 | -0.2478601 | 0.06154625 | -4.0272171 | 5.64E-05 |
| DFG16 | YOR030W | 866.231097 | 0.29509617 | 0.07332052 | 4.02474215 | 5.70E-05 |
| Xu_2009_SUTs29 | Xu_2009_SU | 390.477078 | 0.39195913 | 0.09741885 | 4.0234425 | 5.74E-05 |
| WHI2 | YOR043W | 1448.59705 | -0.2771684 | 0.0689129 | -4.0220105 | 5.77E-05 |
| CCZ1 | YBR131W | 700.923155 | 0.34710029 | 0.08635351 | 4.01952711 | 5.83E-05 |
| Xu_2009_CUTs61 | Xu_2009_CU | 54.0425269 | 0.82908922 | 0.2063057 | 4.01874111 | 5.85E-05 |
| Xu_2009_CUTs45 | Xu_2009_CU | 40.7450958 | 0.90338642 | 0.22504088 | 4.01432134 | 5.96E-05 |
| Xu_2009_CUTs62 | Xu_2009_CU | 35.9225405 | 0.93495446 | 0.23290188 | 4.01437061 | 5.96E-05 |
| Xu_2009_CUTs11 | Xu_2009_CU | 53.2551306 | 0.8389777 | 0.20909841 | 4.01235804 | 6.01E-05 |
| KRS1 | YDR037W | 5284.46653 | -0.2634656 | 0.06568371 | -4.0111248 | 6.04E-05 |
| PPQ1 | YPL179W | 3768.93618 | 0.27330806 | 0.06821416 | 4.00661754 | 6.16E-05 |
| COA4 | YLR218C | 93.5739518 | -0.6717886 | 0.1678297 | -4.0027991 | 6.26E-05 |
| BRF1 | YGR246C | 612.272111 | 0.31731865 | 0.07927969 | 4.00252107 | 6.27E-05 |
| Xu_2009_CUTs10 | Xu_2009_CU | 57.2498045 | -0.8230249 | 0.20575719 | -3.9999814 | 6.33E-05 |
| YER134C | YER134C | 368.435187 | -0.3953154 | 0.09884601 | -3.9993062 | 6.35E-05 |
| AFT2 | YPL202C | 868.134014 | 0.30597926 | 0.07655286 | 3.99696714 | 6.42E-05 |
| YBR287W | YBR287W | 1793.94964 | -0.3010716 | 0.07534089 | -3.9961247 | 6.44E-05 |
| THI22 | YPR121W | 440.715996 | 0.3601677 | 0.09013279 | 3.99596736 | 6.44E-05 |
| PSD2 | YGR170W | 1736.01494 | 0.25710127 | 0.06436084 | 3.99468491 | 6.48E-05 |
| TIM9 | YEL020W-A | 186.617111 | -0.5038148 | 0.12615995 | -3.9934607 | 6.51E-05 |
| RAD14 | YMR201C | 465.517522 | 0.40863189 | 0.10237208 | 3.99163419 | 6.56E-05 |
| Xu_2009_SUTs25 | Xu_2009_SU | 56.3581674 | 0.7869367 | 0.19721401 | 3.99026768 | 6.60E-05 |
| LPX1 | YOR084W | 174.466853 | -0.5542812 | 0.13893472 | -3.9895083 | 6.62E-05 |
| MNR2 | YKL064W | 2252.95047 | 0.21884302 | 0.0548624 | 3.9889435 | 6.64E-05 |
| YJR124C | YJR124C | 480.791687 | -0.3563167 | 0.08933577 | -3.9885107 | 6.65E-05 |
| YNL234W | YNL234W | 741.95273 | 0.32611905 | 0.08177825 | 3.98784584 | 6.67E-05 |
| Xu_2009_SUTs63 | Xu_2009_SU | 32.6963685 | 0.95621587 | 0.23982325 | 3.9871691 | 6.69E-05 |

| | | | | | | |
|---|---|---|---|---|---|---|
| IRC4 | YDR540C | 91.6111561 | -0.6587227 | 0.16524807 | -3.9862659 | 6.71E-05 |
| ATG27 | YJL178C | 954.848395 | -0.2933036 | 0.07357979 | -3.9861979 | 6.71E-05 |
| Xu_2009_CUTs62 | Xu_2009_CU | 80.3926257 | 0.71294274 | 0.17900063 | 3.98290639 | 6.81E-05 |
| SAP155 | YFR040W | 2294.0296 | 0.2447842 | 0.06146376 | 3.98257745 | 6.82E-05 |
| Xu_2009_SUTs55 | Xu_2009_SU | 126.44667 | 0.61929023 | 0.15550609 | 3.9824179 | 6.82E-05 |
| Xu_2009_CUTs53 | Xu_2009_CU | 35.3197809 | 0.92345899 | 0.2320063 | 3.98031868 | 6.88E-05 |
| NMD5 | YJR132W | 1045.48293 | 0.28816039 | 0.07239466 | 3.9804096 | 6.88E-05 |
| YAP6 | YDR259C | 285.061737 | 0.42634505 | 0.10714855 | 3.97900919 | 6.92E-05 |
| Xu_2009_SUTs30 | Xu_2009_SU | 251.943412 | 0.47497253 | 0.11939669 | 3.97810459 | 6.95E-05 |
| Xu_2009_CUTs22 | Xu_2009_CU | 42.4952499 | 0.90727557 | 0.22814759 | 3.97670461 | 6.99E-05 |
| ATG26 | YLR189C | 981.415452 | 0.30623241 | 0.07700779 | 3.97664216 | 6.99E-05 |
| PKH1 | YDR490C | 1304.97518 | 0.28070205 | 0.07059632 | 3.97615709 | 7.00E-05 |
| RPD3 | YNL330C | 607.128144 | -0.3312305 | 0.08335297 | -3.9738298 | 7.07E-05 |
| SRD1 | YCR018C | 490.256249 | 0.36112875 | 0.09088679 | 3.97339115 | 7.09E-05 |
| Xu_2009_SUTs30 | Xu_2009_SU | 37.5646622 | -0.8988528 | 0.22640618 | -3.9700892 | 7.18E-05 |
| GAC1 | YOR178C | 815.718121 | -0.3349444 | 0.08439101 | -3.9689588 | 7.22E-05 |
| RAD5 | YLR032W | 641.087358 | 0.34988742 | 0.08817691 | 3.96801633 | 7.25E-05 |
| OXR1 | YPL196W | 376.521364 | -0.4140557 | 0.10441135 | -3.9656197 | 7.32E-05 |
| YMR295C | YMR295C | 2256.10956 | -0.2921053 | 0.07368408 | -3.9642933 | 7.36E-05 |
| MCY1 | YGR012W | 573.315332 | -0.3214792 | 0.08112104 | -3.9629567 | 7.40E-05 |
| CPR8 | YNR028W | 469.181671 | -0.368917 | 0.09318528 | -3.9589619 | 7.53E-05 |
| Xu_2009_SUTs52 | Xu_2009_SU | 123.279393 | 0.61366633 | 0.15512403 | 3.95597208 | 7.62E-05 |
| Xu_2009_CUTs21 | Xu_2009_CU | 44.5767253 | 0.89165747 | 0.22558177 | 3.9527018 | 7.73E-05 |
| YAP3 | YHL009C | 401.261124 | 0.37831568 | 0.09572774 | 3.95199626 | 7.75E-05 |
| BIO2 | YGR286C | 1584.46104 | -0.3334612 | 0.08443747 | -3.9492093 | 7.84E-05 |
| YEL073C | YEL073C | 130.537566 | -0.618601 | 0.15675395 | -3.9463187 | 7.94E-05 |
| YDR248C | YDR248C | 242.040983 | -0.4539654 | 0.11505005 | -3.945808 | 7.95E-05 |
| Xu_2009_CUTs40 | Xu_2009_CU | 61.3040333 | 0.83830486 | 0.21258932 | 3.94330658 | 8.04E-05 |
| PRP22 | YER013W | 999.941255 | 0.27318501 | 0.06931016 | 3.94148571 | 8.10E-05 |
| ARI1 | YGL157W | 792.689626 | -0.2900116 | 0.0735945 | -3.9406695 | 8.13E-05 |
| PRP2 | YNR011C | 735.498924 | 0.31189143 | 0.07915698 | 3.94016311 | 8.14E-05 |
| Xu_2009_SUTs19 | Xu_2009_SU | 127.079856 | 0.57715164 | 0.1465192 | 3.93908527 | 8.18E-05 |
| Xu_2009_CUTs39 | Xu_2009_CU | 77.9409254 | 0.68867933 | 0.17490293 | 3.93749449 | 8.23E-05 |
| VAC7 | YNL054W | 1536.01111 | 0.23184921 | 0.05888883 | 3.93706588 | 8.25E-05 |
| Xu_2009_SUTs57 | Xu_2009_SU | 225.998974 | 0.45909908 | 0.11662547 | 3.93652515 | 8.27E-05 |
| TIM8 | YJR135W-A | 178.250947 | -0.5216812 | 0.13257138 | -3.9350963 | 8.32E-05 |
| TVP15 | YDR100W | 550.038776 | -0.3916052 | 0.09952475 | -3.934752 | 8.33E-05 |
| JHD1 | YER051W | 277.148675 | 0.43692532 | 0.11105148 | 3.93443937 | 8.34E-05 |
| Xu_2009_CUTs13 | Xu_2009_CU | 178.026152 | 0.52565094 | 0.1336285 | 3.9336738 | 8.37E-05 |
| YGR269W | YGR269W | 44.7814661 | 0.84940202 | 0.21592913 | 3.93370741 | 8.36E-05 |
| ENB1 | YOL158C | 311.678543 | -0.478109 | 0.12160913 | -3.9315227 | 8.44E-05 |

| | | | | | | |
|---|---|---|---|---|---|---|
| Xu_2009_SUTs71( | Xu_2009_SU | 33.4545733 | 0.93483719 | 0.23792437 | 3.92913588 | 8.53E-05 |
| CPD1 | YGR247W | 162.350018 | -0.5291512 | 0.13474229 | -3.927135 | 8.60E-05 |
| COX20 | YDR231C | 385.309804 | -0.3714137 | 0.09458575 | -3.9267407 | 8.61E-05 |
| Xu_2009_SUTs74 | Xu_2009_SU | 101.348459 | 0.63133131 | 0.16090675 | 3.92358504 | 8.72E-05 |
| SHE1 | YBL031W | 168.596489 | -0.5186992 | 0.1322456 | -3.9222414 | 8.77E-05 |
| Xu_2009_SUTs36 | Xu_2009_SU | 118.272686 | 0.58940086 | 0.15034324 | 3.92036823 | 8.84E-05 |
| RRP45 | YDR280W | 794.957446 | 0.34446373 | 0.08786299 | 3.92046423 | 8.84E-05 |
| ELP4 | YPL101W | 894.816475 | -0.2772402 | 0.07072716 | -3.9198548 | 8.86E-05 |
| PEX6 | YNL329C | 1028.885 | 0.28505705 | 0.07273328 | 3.91921079 | 8.88E-05 |
| NOG2 | YNR053C | 4434.06972 | 0.28716874 | 0.07331746 | 3.91678507 | 8.97E-05 |
| Xu_2009_CUTs72 | Xu_2009_CU | 47.0962799 | 0.81641607 | 0.20844974 | 3.91660867 | 8.98E-05 |
| Xu_2009_SUTs34 | Xu_2009_SU | 55.6778855 | 0.79752423 | 0.20378125 | 3.91362905 | 9.09E-05 |
| RKM4 | YDR257C | 289.253931 | 0.43593032 | 0.11139019 | 3.9135432 | 9.10E-05 |
| ADK2 | YER170W | 114.406188 | -0.6024353 | 0.15396575 | -3.9127876 | 9.12E-05 |
| COS12 | YGL263W | 8.18951673 | 1.15349355 | 0.29493437 | 3.91101779 | 9.19E-05 |
| MCT1 | YOR221C | 748.276804 | -0.2982586 | 0.07626399 | -3.9108713 | 9.20E-05 |
| LSC2 | YGR244C | 2048.74498 | -0.2559018 | 0.06544511 | -3.9101738 | 9.22E-05 |
| SRG1 | SRG1 | 684.64699 | 0.34698529 | 0.08880287 | 3.90736591 | 9.33E-05 |
| FET5 | YFL041W | 858.923974 | -0.289531 | 0.07411282 | -3.9066245 | 9.36E-05 |
| Xu_2009_CUTs34 | Xu_2009_CU | 58.6378755 | 0.79189974 | 0.20271733 | 3.90642343 | 9.37E-05 |
| YHR131C | YHR131C | 849.526586 | 0.32014615 | 0.08197751 | 3.90529233 | 9.41E-05 |
| PSH1 | YOL054W | 636.82267 | 0.32113651 | 0.08223075 | 3.90530907 | 9.41E-05 |
| YMR316C-B | YMR316C-B | 64.7804651 | 0.76470496 | 0.19584601 | 3.90462364 | 9.44E-05 |
| RSR1 | YGR152C | 713.096387 | 0.32720125 | 0.08380844 | 3.90415635 | 9.46E-05 |
| YJR120W | YJR120W | 95.7438231 | -0.6501029 | 0.16654682 | -3.9034245 | 9.48E-05 |
| ZPR1 | YGR211W | 1948.08352 | -0.2515215 | 0.0644391 | -3.903243 | 9.49E-05 |
| ARK1 | YNL020C | 484.246551 | 0.33649614 | 0.08621644 | 3.90292332 | 9.50E-05 |
| CIK1 | YMR198W | 405.105925 | 0.44208186 | 0.11327595 | 3.90269843 | 9.51E-05 |
| OST6 | YML019W | 479.91788 | -0.3478006 | 0.08921853 | -3.8983007 | 9.69E-05 |
| NHX1 | YDR456W | 775.684523 | -0.278648 | 0.07148767 | -3.8978474 | 9.71E-05 |
| LSO2 | YGR169C-A | 234.204608 | 0.50417976 | 0.12938848 | 3.89663575 | 9.75E-05 |
| ISM1 | YPL040C | 632.719019 | 0.32227438 | 0.0827538 | 3.89437566 | 9.85E-05 |
| SWT21 | YNL187W | 552.380695 | 0.32504074 | 0.08347222 | 3.8939989 | 9.86E-05 |
| CTK2 | YJL006C | 302.799124 | -0.4085068 | 0.10493682 | -3.8928837 | 9.91E-05 |
| GPN2 | YOR262W | 518.787852 | 0.33844454 | 0.08700301 | 3.89003268 | 0.00010023 |
| tS(AGA)D2 | tS(AGA)D2 | 2.10591528 | -0.8964811 | 0.23049188 | -3.8894261 | 0.00010048 |
| UGO1 | YDR470C | 381.47286 | -0.4048004 | 0.10408042 | -3.8893043 | 0.00010053 |
| Xu_2009_CUTs89 | Xu_2009_CU | 31.0228451 | 1.00318335 | 0.25799012 | 3.88845645 | 0.00010088 |
| YJL211C | YJL211C | 32.4189673 | 0.95857586 | 0.24654204 | 3.88808269 | 0.00010104 |
| RRN6 | YBL014C | 1410.81608 | 0.26406615 | 0.06792817 | 3.8874319 | 0.00010131 |
| YTA6 | YPL074W | 725.503736 | 0.3316986 | 0.08534276 | 3.88666355 | 0.00010163 |

| | | | | | | |
|---|---|---|---|---|---|---|
| KTR3 | YBR205W | 625.390569 | -0.3302502 | 0.08498171 | -3.8861319 | 0.00010185 |
| CKB2 | YOR039W | 780.005928 | -0.3239139 | 0.08335524 | -3.8859448 | 0.00010193 |
| Xu_2009_CUTs30 | Xu_2009_CU | 59.3545442 | 0.79424731 | 0.20450562 | 3.88374329 | 0.00010286 |
| NET1 | YJL076W | 3824.81649 | 0.20117426 | 0.05182263 | 3.88197709 | 0.00010361 |
| SRP1 | YNL189W | 2895.85329 | -0.2006972 | 0.05171662 | -3.8807105 | 0.00010415 |
| RHO3 | YIL118W | 2104.01 | 0.27878255 | 0.0718904 | 3.87788299 | 0.00010537 |
| Xu_2009_SUTs23 | Xu_2009_SU | 256.328238 | -0.4493618 | 0.11589024 | -3.877477 | 0.00010555 |
| PRM9 | YAR031W | 114.432395 | -0.5982429 | 0.15433117 | -3.8763583 | 0.00010603 |
| TPT1 | YOL102C | 275.978988 | -0.4247096 | 0.10957743 | -3.8758856 | 0.00010624 |
| RPT1 | YKL145W | 2891.97463 | -0.2200696 | 0.05680166 | -3.874352 | 0.00010691 |
| PTM1 | YKL039W | 1745.94016 | -0.2447023 | 0.06316488 | -3.8740242 | 0.00010705 |
| Xu_2009_SUTs26 | Xu_2009_SU | 133.210482 | 0.55139589 | 0.14234777 | 3.87358288 | 0.00010725 |
| SBH2 | YER019C-A | 712.41488 | -0.3955665 | 0.10213561 | -3.8729541 | 0.00010752 |
| ROX1 | YPR065W | 1493.53332 | 0.24721396 | 0.06383646 | 3.87261361 | 0.00010767 |
| CRF1 | YDR223W | 186.402542 | 0.48251256 | 0.12473184 | 3.86839923 | 0.00010955 |
| ORC6 | YHR118C | 314.974446 | 0.40737288 | 0.10536152 | 3.86642929 | 0.00011044 |
| YNL046W | YNL046W | 329.092272 | -0.4107227 | 0.10626942 | -3.8649188 | 0.00011113 |
| YRA2 | YKL214C | 215.299367 | -0.481069 | 0.12448944 | -3.864336 | 0.00011139 |
| YLL066W-B | YLL066W-B | 17.5443606 | 1.10008297 | 0.28473102 | 3.86358664 | 0.00011173 |
| ORC1 | YML065W | 719.969083 | 0.29872057 | 0.07733535 | 3.86266548 | 0.00011216 |
| TAZ1 | YPR140W | 421.004186 | -0.3431069 | 0.08891275 | -3.8589169 | 0.00011389 |
| RQC2 | YPL009C | 1502.78069 | 0.26666548 | 0.06913602 | 3.8571137 | 0.00011473 |
| SMX3 | YPR182W | 291.846477 | -0.4189792 | 0.10862799 | -3.8570097 | 0.00011478 |
| Xu_2009_CUTs39 | Xu_2009_CU | 20.6245514 | 1.02874514 | 0.26682179 | 3.85555151 | 0.00011547 |
| Xu_2009_SUTs60 | Xu_2009_SU | 36.6959087 | 0.94002434 | 0.24390633 | 3.85403834 | 0.00011619 |
| Xu_2009_CUTs83 | Xu_2009_CU | 68.3699203 | 0.7332805 | 0.19032926 | 3.85269456 | 0.00011683 |
| BOS1 | YLR078C | 317.970967 | -0.3957032 | 0.1027253 | -3.8520519 | 0.00011713 |
| SOK1 | YDR006C | 2973.27198 | 0.24635605 | 0.0639586 | 3.85180504 | 0.00011725 |
| Xu_2009_SUTs11 | Xu_2009_SU | 189.366445 | 0.49791008 | 0.12935127 | 3.84928628 | 0.00011846 |
| Xu_2009_CUTs74 | Xu_2009_CU | 18.9765906 | 1.08601828 | 0.28225941 | 3.84758924 | 0.00011929 |
| SEC31 | YDL195W | 4341.64157 | -0.1904607 | 0.04950627 | -3.8472027 | 0.00011947 |
| FIT1 | YDR534C | 91.5881027 | 0.64607368 | 0.16794595 | 3.8469144 | 0.00011961 |
| SLX9 | YGR081C | 630.396016 | 0.35087591 | 0.0912216 | 3.84641241 | 0.00011986 |
| VPS28 | YPL065W | 220.66144 | -0.4841211 | 0.12591863 | -3.844714 | 0.00012069 |
| YCR090C | YCR090C | 516.161385 | -0.3197362 | 0.08319858 | -3.8430489 | 0.00012152 |
| YLR410W-B | YLR410W-B | 159.945199 | 0.5349038 | 0.13918416 | 3.84313698 | 0.00012147 |
| COG4 | YPR105C | 946.376572 | -0.2705591 | 0.07041111 | -3.8425623 | 0.00012176 |
| CBP4 | YGR174C | 692.139854 | -0.3563698 | 0.09276248 | -3.841745 | 0.00012216 |
| SNR5 | snR5 | 503.70495 | -0.7880622 | 0.20524353 | -3.8396443 | 0.00012321 |
| Xu_2009_CUTs57 | Xu_2009_CU | 22.9928244 | 1.0267807 | 0.26751138 | 3.83826918 | 0.0001239 |
| Xu_2009_CUTs42 | Xu_2009_CU | 33.6322879 | 0.91469555 | 0.23836017 | 3.83745128 | 0.00012432 |

| | | | | | | |
|---|---|---|---|---|---|---|
| ATG15 | YCR068W | 706.470969 | 0.30453139 | 0.07940198 | 3.83531242 | 0.0001254 |
| VTC4 | YJL012C | 1297.87019 | -0.2864959 | 0.07470072 | -3.8352497 | 0.00012544 |
| TIF4632 | YGL049C | 2812.0419 | 0.2141145 | 0.0558426 | 3.8342502 | 0.00012595 |
| Xu_2009_SUTs45 | Xu_2009_SU | 35.4955635 | -0.9018132 | 0.235228 | -3.8337835 | 0.00012619 |
| PBN1 | YCL052C | 1157.91977 | -0.2529332 | 0.0659759 | -3.8337213 | 0.00012622 |
| SGN1 | YIR001C | 573.132486 | -0.3046962 | 0.07952542 | -3.8314311 | 0.0001274 |
| MRPL24 | YMR193W | 393.088294 | -0.3767556 | 0.09834419 | -3.83099 | 0.00012763 |
| SNR35 | snR35 | 906.147828 | -0.5874362 | 0.15344658 | -3.8282784 | 0.00012904 |
| COM2 | YER130C | 1228.99261 | 0.26734989 | 0.06984085 | 3.82798739 | 0.0001292 |
| YPL225W | YPL225W | 2456.89578 | -0.2435465 | 0.06363411 | -3.8272956 | 0.00012956 |
| Xu_2009_CUTs36 | Xu_2009_CU | 395.473915 | -0.4127468 | 0.10784904 | -3.8270793 | 0.00012967 |
| Xu_2009_SUTs83 | Xu_2009_SU | 74.9013518 | 0.68169115 | 0.17813446 | 3.8268349 | 0.0001298 |
| TES1 | YJR019C | 281.744865 | -0.4217816 | 0.11021996 | -3.8267256 | 0.00012986 |
| ATG33 | YLR356W | 631.723263 | -0.3066917 | 0.08017612 | -3.8252246 | 0.00013065 |
| ALG8 | YOR067C | 682.243011 | -0.3003444 | 0.07852082 | -3.8250287 | 0.00013076 |
| COP1 | YDL145C | 5348.63067 | -0.1811122 | 0.04735497 | -3.8245647 | 0.000131 |
| YGL140C | YGL140C | 2024.02799 | 0.23277976 | 0.06087433 | 3.82393935 | 0.00013134 |
| YUH1 | YJR099W | 524.599341 | -0.3360178 | 0.08787644 | -3.8237528 | 0.00013144 |
| SGT1 | YOR057W | 638.113686 | -0.2965958 | 0.07761058 | -3.8215894 | 0.00013259 |
| Xu_2009_CUTs61 | Xu_2009_CU | 64.6918108 | 0.72344976 | 0.18934658 | 3.8207702 | 0.00013304 |
| SPR28 | YDR218C | 67.4590114 | -0.7211704 | 0.18877022 | -3.8203609 | 0.00013326 |
| TRK2 | YKR050W | 850.674477 | 0.271573 | 0.07109605 | 3.81980425 | 0.00013356 |
| BUD31 | YCR063W | 243.421107 | 0.43301535 | 0.11338941 | 3.81883405 | 0.00013408 |
| MCM2 | YBL023C | 1336.02555 | 0.25284572 | 0.0662313 | 3.81761658 | 0.00013475 |
| RKM2 | YDR198C | 557.281241 | 0.32176896 | 0.08430179 | 3.81686982 | 0.00013516 |
| PBI1 | YPL272C | 276.380948 | 0.40526363 | 0.10619219 | 3.81632229 | 0.00013546 |
| Xu_2009_CUTs83 | Xu_2009_CU | 86.9418316 | 0.70636406 | 0.18518364 | 3.81439778 | 0.00013652 |
| RTK1 | YDL025C | 682.848237 | -0.3159118 | 0.08287577 | -3.811872 | 0.00013792 |
| KES1 | YPL145C | 2257.55019 | -0.2496849 | 0.06550027 | -3.8119669 | 0.00013787 |
| ATP25 | YMR098C | 846.58678 | -0.2879078 | 0.07554287 | -3.8111845 | 0.0001383 |
| FMC1 | YIL098C | 260.165488 | -0.4464409 | 0.11714778 | -3.8109211 | 0.00013845 |
| AHK1 | YDL073W | 949.441529 | 0.25957461 | 0.0681288 | 3.81005702 | 0.00013893 |
| MPD2 | YOL088C | 737.169032 | -0.3083728 | 0.08101046 | -3.8065799 | 0.0001409 |
| BEM4 | YPL161C | 362.568171 | 0.38750893 | 0.10181678 | 3.8059437 | 0.00014126 |
| HDA3 | YPR179C | 548.094427 | 0.32619228 | 0.08574667 | 3.80413944 | 0.0001423 |
| RAD3 | YER171W | 1094.85903 | 0.26996784 | 0.07102621 | 3.80096085 | 0.00014414 |
| CDC13 | YDL220C | 446.689315 | 0.35537843 | 0.09352476 | 3.79983256 | 0.00014479 |
| PPM2 | YOL141W | 342.607058 | 0.40179734 | 0.10574768 | 3.79958542 | 0.00014494 |
| Xu_2009_SUTs16 | Xu_2009_SU | 258.241905 | 0.4525271 | 0.11910239 | 3.79947956 | 0.000145 |
| RNA1 | YMR235C | 1937.90118 | -0.2342364 | 0.06165187 | -3.7993401 | 0.00014508 |
| YMC1 | YPR058W | 302.566342 | -0.3907488 | 0.10285241 | -3.7991212 | 0.00014521 |

| | | | | | | |
|---|---|---|---|---|---|---|
| BCH1 | YMR237W | 1256.26645 | -0.2581029 | 0.06793969 | -3.7990001 | 0.00014528 |
| YLL053C | YLL053C | 34.5803425 | -0.9075742 | 0.23890793 | -3.798845 | 0.00014537 |
| MPH1 | YIR002C | 843.374077 | 0.32481472 | 0.0855139 | 3.79838494 | 0.00014564 |
| ISW2 | YOR304W | 1724.94948 | 0.2620764 | 0.06900392 | 3.79799286 | 0.00014587 |
| YGL036W | YGL036W | 1190.64898 | 0.26977978 | 0.07107979 | 3.79544999 | 0.00014738 |
| Xu_2009_CUTs79 | Xu_2009_CU | 90.1623524 | 0.62693236 | 0.1652063 | 3.79484533 | 0.00014774 |
| ERG5 | YMR015C | 949.085949 | -0.3016399 | 0.07949201 | -3.7945934 | 0.00014789 |
| SSA3 | YBL075C | 897.694087 | 0.31787302 | 0.08379915 | 3.79327261 | 0.00014867 |
| Xu_2009_CUTs16 | Xu_2009_CU | 64.6214336 | 0.7542119 | 0.19883472 | 3.79315994 | 0.00014874 |
| RTC5 | YOR118W | 627.857733 | 0.28974158 | 0.07644981 | 3.78995822 | 0.00015067 |
| LTV1 | YKL143W | 1017.53274 | 0.34301055 | 0.09052572 | 3.78909481 | 0.0001512 |
| Xu_2009_CUTs86 | Xu_2009_CU | 27.654145 | 0.96611512 | 0.25500668 | 3.78858753 | 0.00015151 |
| SPO24 | YPR036W-A | 8117.81902 | -0.3110384 | 0.08212909 | -3.787189 | 0.00015236 |
| NNF2 | YGR089W | 1263.53929 | 0.28259802 | 0.07463421 | 3.78644102 | 0.00015282 |
| Xu_2009_SUTs18 | Xu_2009_SU | 207.636235 | 0.47983377 | 0.12672812 | 3.7863244 | 0.00015289 |
| YJR011C | YJR011C | 97.9248005 | -0.6789493 | 0.17938632 | -3.7848445 | 0.0001538 |
| PRM4 | YPL156C | 247.582169 | -0.4562411 | 0.12058221 | -3.7836519 | 0.00015454 |
| YIL055C | YIL055C | 439.052718 | 0.34164022 | 0.09030042 | 3.78337372 | 0.00015472 |
| Xu_2009_SUTs14 | Xu_2009_SU | 22.1011389 | 1.02649883 | 0.27134687 | 3.78297654 | 0.00015496 |
| CTR9 | YOL145C | 2048.45623 | 0.2276509 | 0.06022228 | 3.78017748 | 0.00015672 |
| MRS2 | YOR334W | 392.384223 | -0.354043 | 0.09365725 | -3.7801987 | 0.0001567 |
| Xu_2009_CUTs43 | Xu_2009_CU | 56.6463497 | 0.77363044 | 0.20472647 | 3.77884916 | 0.00015755 |
| YOR008W-B | YOR008W-B | 103.822887 | 0.64552359 | 0.17092475 | 3.77665372 | 0.00015895 |
| Xu_2009_CUTs21 | Xu_2009_CU | 63.0758821 | 0.74266064 | 0.19665399 | 3.77648393 | 0.00015906 |
| YKR015C | YKR015C | 47.0911368 | 0.79188294 | 0.20973173 | 3.77569461 | 0.00015956 |
| Xu_2009_CUTs75 | Xu_2009_CU | 63.6005888 | 0.74132606 | 0.19638793 | 3.77480469 | 0.00016013 |
| PSF1 | YDR013W | 144.839768 | -0.5348768 | 0.14172843 | -3.7739557 | 0.00016068 |
| Xu_2009_SUTs79 | Xu_2009_SU | 27.2024396 | 0.97041829 | 0.25723385 | 3.77251399 | 0.00016161 |
| Xu_2009_CUTs58 | Xu_2009_CU | 78.9213873 | 0.66524365 | 0.17636934 | 3.77187819 | 0.00016202 |
| VRP1 | YLR337C | 831.677618 | -0.2743984 | 0.07278637 | -3.7699145 | 0.0001633 |
| MIC26 | YGR235C | 862.063417 | -0.2613178 | 0.06935669 | -3.7677369 | 0.00016473 |
| MMT2 | YPL224C | 685.603245 | -0.2897601 | 0.07691052 | -3.7674965 | 0.00016489 |
| Xu_2009_CUTs91 | Xu_2009_CU | 42.5436681 | 0.84603593 | 0.2247836 | 3.76377955 | 0.00016736 |
| SUR7 | YML052W | 1275.30362 | -0.2478218 | 0.06593677 | -3.758476 | 0.00017095 |
| HER1 | YOR227W | 1622.61533 | 0.23114705 | 0.0615058 | 3.75813417 | 0.00017119 |
| tM(CAU)M/EMT4 | tM(CAU)M | 16.5698306 | -1.0744298 | 0.28592163 | -3.7577771 | 0.00017143 |
| IMD1 | YAR073W | 13.0512751 | 1.10284445 | 0.29352811 | 3.75720215 | 0.00017182 |
| Xu_2009_CUTs84 | Xu_2009_CU | 48.3679574 | 0.80951047 | 0.21558065 | 3.75502373 | 0.00017333 |
| MCO12 | YKL018C-A | 60.3685981 | 0.73565988 | 0.19591419 | 3.75501079 | 0.00017333 |
| SDH7 | YDR511W | 177.767455 | -0.476342 | 0.12686291 | -3.7547778 | 0.0001735 |
| MRK1 | YDL079C | 331.575956 | 0.39148162 | 0.10428403 | 3.75399392 | 0.00017404 |

| | | | | | | |
|---|---|---|---|---|---|---|
| ADD37 | YMR184W | 357.922353 | -0.3861947 | 0.10287996 | -3.753838 | 0.00017415 |
| CLN3 | YAL040C | 3040.06244 | 0.24041235 | 0.06407181 | 3.75223308 | 0.00017527 |
| SYT1 | YPR095C | 1137.92668 | 0.26812304 | 0.07147726 | 3.75116591 | 0.00017601 |
| MRPS16 | YPL013C | 582.907988 | -0.3533567 | 0.09423146 | -3.7498807 | 0.00017692 |
| YGL204C | YGL204C | 106.709768 | -0.6124194 | 0.16336663 | -3.7487423 | 0.00017772 |
| Xu_2009_CUTs54 | Xu_2009_CU | 99.2911189 | 0.60808386 | 0.16223527 | 3.74816072 | 0.00017814 |
| CAJ1 | YER048C | 1376.1214 | -0.2397506 | 0.06396686 | -3.7480436 | 0.00017822 |
| Xu_2009_SUTs43 | Xu_2009_SU | 658.464563 | 0.29628681 | 0.07909799 | 3.74581963 | 0.00017981 |
| NUP170 | YBL079W | 3669.27382 | 0.18979382 | 0.05066782 | 3.74584553 | 0.00017979 |
| Xu_2009_SUTs40 | Xu_2009_SU | 58.0904863 | 0.80455537 | 0.21488873 | 3.74405568 | 0.00018107 |
| HXT2 | YMR011W | 1768.5171 | -0.7198857 | 0.19240416 | -3.7415284 | 0.0001829 |
| Xu_2009_CUTs41 | Xu_2009_CU | 47.7383299 | 0.8234502 | 0.2200944 | 3.74135007 | 0.00018303 |
| MYO2 | YOR326W | 3836.92398 | 0.18636592 | 0.04982365 | 3.74051126 | 0.00018365 |
| YBR085C-A | YBR085C-A | 5081.97879 | 0.26827805 | 0.07173995 | 3.73959059 | 0.00018432 |
| CTA1 | YDR256C | 99.5850418 | 0.60065058 | 0.16075329 | 3.73647446 | 0.00018662 |
| FMP46 | YKR049C | 695.760401 | -0.2792448 | 0.07481961 | -3.7322413 | 0.00018978 |
| ROM1 | YGR070W | 1077.58187 | 0.3237355 | 0.08678152 | 3.7304659 | 0.00019113 |
| RUP1 | YOR138C | 1443.23698 | 0.24949732 | 0.06688639 | 3.7301656 | 0.00019135 |
| Xu_2009_CUTs51 | Xu_2009_CU | 33.6277099 | 0.9157308 | 0.24551461 | 3.7298423 | 0.0001916 |
| Xu_2009_SUTs61 | Xu_2009_SU | 121.187306 | 0.5737123 | 0.15381738 | 3.72982754 | 0.00019161 |
| YER188C-A | YER188C-A | 14.7682583 | 1.06322411 | 0.28528363 | 3.72690194 | 0.00019385 |
| Xu_2009_CUTs73 | Xu_2009_CU | 39.1239491 | 0.87004669 | 0.23351431 | 3.72588172 | 0.00019463 |
| GRX4 | YER174C | 268.268162 | -0.4473851 | 0.12012146 | -3.7244398 | 0.00019575 |
| FMP52 | YER004W | 721.681349 | -0.2860074 | 0.07679997 | -3.7240562 | 0.00019605 |
| Xu_2009_SUTs20 | Xu_2009_SU | 69.3901584 | 0.73120626 | 0.1963746 | 3.72352771 | 0.00019646 |
| RVB1 | YDR190C | 1841.28673 | -0.2221733 | 0.0596851 | -3.7224246 | 0.00019732 |
| RPL7B | YPL198W | 2140.52934 | -0.238907 | 0.06419633 | -3.721505 | 0.00019804 |
| YIL014C-A | YIL014C-A | 294.371126 | -0.3911557 | 0.10513196 | -3.720616 | 0.00019874 |
| SKI6 | YGR195W | 351.210661 | -0.3571671 | 0.09611471 | -3.7160508 | 0.00020236 |
| AAP1 | YHR047C | 1323.831 | -0.2570988 | 0.06920996 | -3.7147655 | 0.00020339 |
| SEG2 | YKL105C | 1037.04678 | 0.28938328 | 0.07790283 | 3.71466991 | 0.00020347 |
| YNR048W | YNR048W | 414.266229 | -0.3589201 | 0.09664676 | -3.7137313 | 0.00020423 |
| IDP2 | YLR174W | 170.251805 | 0.48514117 | 0.13065014 | 3.71328462 | 0.00020459 |
| MCH5 | YOR306C | 1586.73576 | 0.30755548 | 0.0828484 | 3.7122682 | 0.00020541 |
| Xu_2009_SUTs32 | Xu_2009_SU | 70.1973901 | 0.74307458 | 0.20021027 | 3.71147084 | 0.00020606 |
| Xu_2009_CUTs49 | Xu_2009_CU | 83.0818053 | 0.65808651 | 0.17734018 | 3.71087088 | 0.00020655 |
| YER160C | YER160C | 364.049765 | 0.35443767 | 0.09551187 | 3.71092799 | 0.0002065 |
| ALT1 | YLR089C | 2136.52751 | -0.2372976 | 0.06395339 | -3.7104776 | 0.00020687 |
| Xu_2009_CUTs56 | Xu_2009_CU | 33.0567783 | 0.92122192 | 0.24851734 | 3.70687183 | 0.00020984 |
| Xu_2009_SUTs24 | Xu_2009_SU | 115.635958 | 0.63650295 | 0.17180333 | 3.70483471 | 0.00021153 |
| YMR086C-A | YMR086C-A | 31.8666388 | 0.92659077 | 0.25011391 | 3.70467506 | 0.00021166 |

| | | | | | | |
|---|---|---|---|---|---|---|
| STP4 | YDL048C | 2768.71107 | 0.22351734 | 0.06035533 | 3.70335695 | 0.00021277 |
| Xu_2009_SUTs71: | Xu_2009_SU | 385.69801 | 0.35825763 | 0.09677206 | 3.702077 | 0.00021384 |
| ORT1 | YOR130C | 166.89499 | -0.5069299 | 0.13695788 | -3.7013559 | 0.00021445 |
| RQT4 | YKR023W | 506.409976 | 0.32427966 | 0.08762195 | 3.7008953 | 0.00021484 |
| TAR1 | YLR154W-C | 472.160866 | -1.0385807 | 0.28071477 | -3.6997723 | 0.00021579 |
| Xu_2009_CUTs47 | Xu_2009_CU | 33.4661424 | 0.87159822 | 0.23562806 | 3.6990426 | 0.00021641 |
| NSG2 | YNL156C | 648.520575 | -0.2912819 | 0.07876666 | -3.6980352 | 0.00021727 |
| Xu_2009_SUTs61! | Xu_2009_SU | 46.6124443 | -0.8150728 | 0.22045446 | -3.6972391 | 0.00021796 |
| YBL071C | YBL071C | 17.5913015 | 1.03297357 | 0.27944929 | 3.69646154 | 0.00021863 |
| Xu_2009_SUTs64 | Xu_2009_SU | 202.150608 | 0.45783515 | 0.12386745 | 3.69617005 | 0.00021888 |
| TKL2 | YBR117C | 355.467287 | 0.39626865 | 0.10724146 | 3.69510679 | 0.00021979 |
| YJR112W-A | YJR112W-A | 275.620595 | -0.4020271 | 0.10880045 | -3.6950869 | 0.00021981 |
| Xu_2009_SUTs52: | Xu_2009_SU | 42.4215776 | 0.84538026 | 0.22882618 | 3.69442101 | 0.00022039 |
| DAD1 | YDR016C | 69.7801285 | -0.672116 | 0.18198504 | -3.6932487 | 0.00022141 |
| GET2 | YER083C | 703.032837 | -0.2932934 | 0.0794202 | -3.6929322 | 0.00022168 |
| YLR154W-F | YLR154W-F | 124.754379 | -0.966978 | 0.26189767 | -3.6921978 | 0.00022232 |
| DUG3 | YNL191W | 540.981203 | 0.29739187 | 0.08054846 | 3.69208663 | 0.00022242 |
| PBP1 | YGR178C | 2868.02687 | -0.2012264 | 0.0545045 | -3.6919227 | 0.00022257 |
| CLA4 | YNL298W | 1430.95278 | 0.23589413 | 0.06391116 | 3.69096931 | 0.0002234 |
| PMA2 | YPL036W | 486.376729 | 0.32323033 | 0.08760523 | 3.68962368 | 0.00022459 |
| SHH3 | YMR118C | 21.0434996 | -0.9862006 | 0.26734279 | -3.6888992 | 0.00022523 |
| PPR1 | YLR014C | 395.13262 | 0.34415344 | 0.09332115 | 3.68783982 | 0.00022617 |
| YGR111W | YGR111W | 584.622936 | -0.3200279 | 0.08681487 | -3.6863263 | 0.00022751 |
| SNC1 | YAL030W | 770.245366 | -0.3036024 | 0.08237649 | -3.6855464 | 0.00022821 |
| MCM6 | YGL201C | 1153.20943 | 0.2467623 | 0.06698438 | 3.68387797 | 0.00022971 |
| YJL132W | YJL132W | 377.575922 | 0.42635945 | 0.11591017 | 3.67836086 | 0.00023474 |
| TSR3 | YOR006C | 540.169024 | 0.38376061 | 0.10436896 | 3.67696109 | 0.00023603 |
| YKR011C | YKR011C | 603.436613 | -0.3393479 | 0.09229821 | -3.6766464 | 0.00023632 |
| SWR1 | YDR334W | 2190.91359 | 0.22143452 | 0.06024898 | 3.67532375 | 0.00023755 |
| RPN4 | YDL020C | 2699.25741 | 0.24245722 | 0.06603159 | 3.67183693 | 0.00024081 |
| YRA1 | YDR381W | 2537.12365 | -0.1983031 | 0.0540093 | -3.6716478 | 0.00024099 |
| tV(AAC)L | tV(AAC)L | 20.9961222 | -0.9864796 | 0.26874456 | -3.6706962 | 0.00024189 |
| Xu_2009_CUTs32 | Xu_2009_CU | 50.8380457 | 0.76593156 | 0.20866772 | 3.67057994 | 0.000242 |
| Xu_2009_CUTs76 | Xu_2009_CU | 21.0178647 | 1.02647946 | 0.27974083 | 3.66939443 | 0.00024313 |
| YPL060C-A | YPL060C-A | 63.6968429 | 0.728336 | 0.19849311 | 3.66932644 | 0.00024319 |
| Xu_2009_CUTs88 | Xu_2009_CU | 25.5775051 | 0.94400185 | 0.25739135 | 3.66757409 | 0.00024486 |
| Xu_2009_CUTs72 | Xu_2009_CU | 31.113023 | 0.89248757 | 0.24370861 | 3.66210927 | 0.00025015 |
| HUL5 | YGL141W | 1176.08404 | 0.28013297 | 0.07652305 | 3.66076607 | 0.00025146 |
| YRO2 | YBR054W | 3618.66783 | -0.2885708 | 0.07884655 | -3.6599032 | 0.00025231 |
| YBR051W | YBR051W | 15.5974299 | 1.04793941 | 0.28637027 | 3.65938613 | 0.00025282 |
| SYH1 | YPL105C | 2467.36057 | 0.24574271 | 0.06721154 | 3.65625766 | 0.00025592 |

| | | | | | | |
|---|---|---|---|---|---|---|
| YEL020C-B | YEL020C-B | 30.2426741 | 0.89574984 | 0.24506528 | 3.65514784 | 0.00025703 |
| APM2 | YHL019C | 682.981215 | 0.31301237 | 0.08570699 | 3.65212168 | 0.00026008 |
| Xu_2009_SUTs17( | Xu_2009_SU | 68.1526302 | -0.6817393 | 0.18669697 | -3.6515819 | 0.00026063 |
| RSP5 | YER125W | 2666.12912 | -0.20347 | 0.05572228 | -3.651502 | 0.00026071 |
| MAL31 | YBR298C | 155.157448 | -0.4875431 | 0.13353473 | -3.6510585 | 0.00026116 |
| Xu_2009_CUTs44 | Xu_2009_CU | 50.0872314 | 0.75238423 | 0.20610732 | 3.65044878 | 0.00026178 |
| NSA2 | YER126C | 1543.21726 | -0.2611737 | 0.07155741 | -3.649848 | 0.0002624 |
| MDE1 | YJR024C | 788.197465 | -0.2801196 | 0.07675615 | -3.6494745 | 0.00026278 |
| SPE2 | YOL052C | 818.019508 | -0.2708042 | 0.07421195 | -3.6490647 | 0.0002632 |
| KIN2 | YLR096W | 1680.37308 | 0.22434242 | 0.06148561 | 3.64869778 | 0.00026357 |
| Xu_2009_CUTs25 | Xu_2009_CU | 203.029773 | -0.4501867 | 0.12344609 | -3.6468283 | 0.0002655 |
| NVJ1 | YHR195W | 824.900521 | -0.2896318 | 0.07942158 | -3.6467644 | 0.00026556 |
| Xu_2009_SUTs47: | Xu_2009_SU | 127.02805 | 0.55110014 | 0.15113652 | 3.64637318 | 0.00026597 |
| CAN1 | YEL063C | 1987.06416 | -0.2192704 | 0.0601366 | -3.6462062 | 0.00026614 |
| SLX4 | YLR135W | 424.167938 | 0.34480652 | 0.0945657 | 3.64621113 | 0.00026614 |
| YMR320W | YMR320W | 103.177374 | 0.57662311 | 0.15819498 | 3.64501526 | 0.00026738 |
| RPB10 | YOR210W | 507.773677 | -0.3373227 | 0.092596 | -3.642951 | 0.00026953 |
| tW(CCA)P | tW(CCA)P | 122.317074 | 0.59469665 | 0.16327395 | 3.64232423 | 0.00027019 |
| SHH4 | YLR164W | 53.6001875 | -0.7334522 | 0.20137031 | -3.6423055 | 0.00027021 |
| Xu_2009_SUTs16! | Xu_2009_SU | 109.594957 | 0.5737532 | 0.15754413 | 3.64185717 | 0.00027068 |
| LAS1 | YKR063C | 774.733919 | 0.27258849 | 0.074879 | 3.6403864 | 0.00027223 |
| ECM19 | YLR390W | 510.50386 | -0.3074581 | 0.08447117 | -3.6397987 | 0.00027285 |
| YGR122W | YGR122W | 375.643003 | -0.3506161 | 0.09633551 | -3.6395312 | 0.00027313 |
| Xu_2009_SUTs14( | Xu_2009_SU | 51.0089835 | 0.75551056 | 0.20761613 | 3.63897816 | 0.00027372 |
| RSM26 | YJR101W | 392.400839 | -0.3854444 | 0.10593355 | -3.6385487 | 0.00027418 |
| KAP95 | YLR347C | 2731.33031 | 0.18724323 | 0.05148322 | 3.63697588 | 0.00027586 |
| Xu_2009_CUTs54 | Xu_2009_CU | 70.9789627 | 0.69456656 | 0.19103135 | 3.63587741 | 0.00027704 |
| YLR154W-A | YLR154W-A | 196.216724 | -1.0634068 | 0.29248789 | -3.6357293 | 0.0002772 |
| ERV1 | YGR029W | 319.29387 | -0.3704969 | 0.10191886 | -3.6352144 | 0.00027775 |
| POA1 | YBR022W | 230.152219 | -0.4475814 | 0.12315151 | -3.6343965 | 0.00027863 |
| TIP41 | YPR040W | 658.25775 | -0.2878574 | 0.07920291 | -3.6344299 | 0.0002786 |
| ALP1 | YNL270C | 179.477104 | -0.4874913 | 0.13418111 | -3.6330842 | 0.00028005 |
| GSC2 | YGR032W | 7681.88044 | -0.191958 | 0.05284226 | -3.6326602 | 0.00028051 |
| KAR9 | YPL269W | 188.748289 | -0.4721606 | 0.12999888 | -3.632036 | 0.00028119 |
| DOA4 | YDR069C | 498.424638 | -0.34417 | 0.09476448 | -3.6318464 | 0.0002814 |
| AQY3 | YFL054C | 317.767598 | 0.39817892 | 0.10964022 | 3.63168646 | 0.00028158 |
| MSM1 | YGR171C | 470.250731 | -0.3565607 | 0.09818745 | -3.6314281 | 0.00028186 |
| YAR028W | YAR028W | 451.689994 | 0.32691286 | 0.09006358 | 3.62980106 | 0.00028364 |
| MCO10 | YOR020W-A | 368.156362 | -0.3841295 | 0.10585399 | -3.6288623 | 0.00028467 |
| Xu_2009_SUTs69( | Xu_2009_SU | 35.044449 | -0.8497935 | 0.23420136 | -3.6284738 | 0.0002851 |
| SUS1 | YBR111W-A | 236.085691 | -0.4548038 | 0.12536435 | -3.6278562 | 0.00028578 |

| | | | | | | |
|---|---|---|---|---|---|---|
| VMS1 | YDR049W | 918.54021 | 0.25434973 | 0.07012141 | 3.62727608 | 0.00028643 |
| Xu_2009_SUTs79 | Xu_2009_SU | 53.355983 | 0.78393087 | 0.21613943 | 3.6269683 | 0.00028677 |
| SRP14 | YDL092W | 1029.2174 | -0.2488632 | 0.06864417 | -3.6254089 | 0.0002885 |
| BIR1 | YJR089W | 744.304271 | 0.29519347 | 0.08149244 | 3.62234194 | 0.00029195 |
| SCC2 | YDR180W | 819.584677 | 0.2836447 | 0.0783084 | 3.62214918 | 0.00029217 |
| Xu_2009_CUTs31 | Xu_2009_CU | 57.6427214 | 0.74721535 | 0.20631058 | 3.62179858 | 0.00029256 |
| ERG27 | YLR100W | 1598.37456 | -0.2458268 | 0.06787854 | -3.6215696 | 0.00029282 |
| FUI1 | YBL042C | 1423.13107 | 0.33136772 | 0.09160251 | 3.61745231 | 0.00029752 |
| TFG1 | YGR186W | 1882.78214 | 0.21654576 | 0.05986166 | 3.61743656 | 0.00029754 |
| EAR1 | YMR171C | 1136.17465 | 0.25039875 | 0.06928546 | 3.61401573 | 0.00030149 |
| GPX1 | YKL026C | 100.372202 | -0.5753925 | 0.15935355 | -3.6107922 | 0.00030526 |
| STE20 | YHL007C | 2715.34113 | 0.22097414 | 0.06120252 | 3.61053992 | 0.00030556 |
| Xu_2009_SUTs17 | Xu_2009_SU | 98.1009675 | 0.57538062 | 0.15941743 | 3.60927041 | 0.00030706 |
| Xu_2009_CUTs24 | Xu_2009_CU | 42.5092311 | 0.79395431 | 0.22001066 | 3.6087083 | 0.00030773 |
| AI5_BETA | Q0075 | 125.102623 | -0.582162 | 0.16140519 | -3.6068355 | 0.00030995 |
| MSC6 | YOR354C | 831.030868 | -0.2642568 | 0.07327824 | -3.606211 | 0.0003107 |
| BUD6 | YLR319C | 614.783136 | 0.28342824 | 0.07862239 | 3.60493041 | 0.00031224 |
| Xu_2009_CUTs29 | Xu_2009_CU | 45.1445365 | 0.78966697 | 0.21908176 | 3.60443965 | 0.00031283 |
| tK(UUU)P | tK(UUU)P | 5.47503273 | -1.0379723 | 0.28799136 | -3.6041786 | 0.00031314 |
| SLM3 | YDL033C | 520.763472 | 0.33671285 | 0.0934386 | 3.60357347 | 0.00031387 |
| PIB1 | YDR313C | 260.08667 | -0.4148996 | 0.11517929 | -3.6022065 | 0.00031553 |
| YJL119C | YJL119C | 57.0907249 | 0.73455261 | 0.20393915 | 3.60182237 | 0.00031599 |
| Xu_2009_CUTs49 | Xu_2009_CU | 57.9316087 | 0.70925978 | 0.19693255 | 3.60153654 | 0.00031634 |
| MNN14 | YJR061W | 747.69017 | -0.2821791 | 0.07836294 | -3.6009259 | 0.00031709 |
| MIF2 | YKL089W | 395.448791 | 0.33034348 | 0.09176124 | 3.60003286 | 0.00031818 |
| FAS2 | YPL231W | 17901.2752 | -0.2025292 | 0.05631541 | -3.5963372 | 0.00032273 |
| Xu_2009_SUTs23 | Xu_2009_SU | 55.5355812 | 0.731744 | 0.20352605 | 3.59533333 | 0.00032398 |
| YML020W | YML020W | 746.945563 | 0.26915649 | 0.07489475 | 3.59379671 | 0.00032589 |
| YBL028C | YBL028C | 935.874811 | -0.3290903 | 0.0916537 | -3.5905839 | 0.00032994 |
| COS6 | YGR295C | 929.342017 | 0.27625819 | 0.07694747 | 3.59021821 | 0.0003304 |
| GAT4 | YIR013C | 18.1792223 | 0.99892075 | 0.27824278 | 3.59010488 | 0.00033054 |
| Xu_2009_SUTs74 | Xu_2009_SU | 82.992485 | 0.64912836 | 0.18093349 | 3.58766287 | 0.00033366 |
| LMO1 | YLL007C | 330.033755 | 0.36703378 | 0.10240509 | 3.584136 | 0.0003382 |
| SOG2 | YOR353C | 885.081512 | 0.27965107 | 0.07802427 | 3.58415502 | 0.00033817 |
| RBH1 | YJL181W | 294.371964 | -0.3734474 | 0.10420054 | -3.5839294 | 0.00033846 |
| CWC21 | YDR482C | 612.78449 | 0.28414528 | 0.07929044 | 3.58360081 | 0.00033889 |
| Xu_2009_CUTs83 | Xu_2009_CU | 61.8801388 | 0.68077777 | 0.19006913 | 3.58173762 | 0.00034132 |
| RPB5 | YBR154C | 1661.91619 | -0.2214643 | 0.06183045 | -3.5817998 | 0.00034124 |
| YPR148C | YPR148C | 1396.01039 | 0.22127036 | 0.06179746 | 3.58057365 | 0.00034284 |
| TGL5 | YOR081C | 952.156086 | -0.2513136 | 0.0702013 | -3.5798992 | 0.00034373 |
| AIM4 | YBR194W | 194.158547 | 0.48079386 | 0.1343435 | 3.57883964 | 0.00034512 |

| | | | | | | |
|---|---|---|---|---|---|---|
| PUN1 | YLR414C | 1581.51691 | -0.2288222 | 0.06394967 | -3.5781615 | 0.00034602 |
| Xu_2009_CUTs11 | Xu_2009_CU | 32.4280134 | 0.86002178 | 0.24037733 | 3.57779902 | 0.0003465 |
| RPN11 | YFR004W | 1295.52779 | -0.2306252 | 0.06450004 | -3.5755826 | 0.00034945 |
| YPR170W-B | YPR170W-B | 37.8134317 | -0.833956 | 0.23323361 | -3.5756254 | 0.00034939 |
| MPO1 | YGL010W | 240.864926 | -0.4146508 | 0.11601136 | -3.5742258 | 0.00035127 |
| Xu_2009_CUTs79 | Xu_2009_CU | 33.286443 | 0.86614237 | 0.24235756 | 3.5738203 | 0.00035181 |
| BUD8 | YLR353W | 908.895077 | 0.265619 | 0.07436823 | 3.57167313 | 0.00035471 |
| GPP1 | YIL053W | 6504.01783 | -0.2826704 | 0.07915706 | -3.5710067 | 0.00035561 |
| Xu_2009_SUTs22 | Xu_2009_SU | 51.2721086 | 0.75804749 | 0.21229198 | 3.57077772 | 0.00035592 |
| RLI1 | YDR091C | 1797.55216 | -0.2589779 | 0.07253776 | -3.5702498 | 0.00035664 |
| YLR157C-B | YLR157C-B | 61.4683997 | 0.69260865 | 0.19404867 | 3.56925218 | 0.000358 |
| YIL046W-A | YIL046W-A | 41.4527021 | 0.79649624 | 0.22318831 | 3.56871844 | 0.00035873 |
| CUE3 | YGL110C | 599.973203 | 0.29833953 | 0.08361051 | 3.56820604 | 0.00035943 |
| OSH7 | YHR001W | 639.190554 | -0.3556304 | 0.09982618 | -3.5624966 | 0.00036734 |
| ERI1 | YPL096C-A | 89.952717 | -0.6234198 | 0.17504297 | -3.5615245 | 0.00036871 |
| Xu_2009_CUTs48 | Xu_2009_CU | 46.5380909 | 0.77834512 | 0.21862486 | 3.56018579 | 0.00037059 |
| PRM7 | YDL039C | 3448.51516 | 0.56349119 | 0.15836262 | 3.55823359 | 0.00037336 |
| Xu_2009_SUTs61 | Xu_2009_SU | 92.1686486 | 0.58925085 | 0.16565069 | 3.55718917 | 0.00037484 |
| Xu_2009_CUTs68 | Xu_2009_CU | 37.0713367 | 0.80247684 | 0.22563624 | 3.55650689 | 0.00037582 |
| YOR238W | YOR238W | 446.42331 | -0.3411583 | 0.09595391 | -3.5554389 | 0.00037735 |
| PMP3 | YDR276C | 1477.79841 | -0.2873442 | 0.08083057 | -3.5548948 | 0.00037813 |
| MMS22 | YLR320W | 720.669619 | 0.27382965 | 0.07703036 | 3.55482766 | 0.00037823 |
| HIR2 | YOR038C | 573.196738 | 0.29713009 | 0.08358878 | 3.55466467 | 0.00037846 |
| NNF1 | YJR112W | 209.492025 | -0.4423036 | 0.12444779 | -3.55413 | 0.00037923 |
| PRP16 | YKR086W | 1156.36188 | 0.22798907 | 0.06417488 | 3.55262186 | 0.00038141 |
| URA6 | YKL024C | 502.224676 | -0.3347814 | 0.09424151 | -3.5523773 | 0.00038177 |
| Xu_2009_CUTs4 | Xu_2009_CU | 31.7053927 | 0.84447758 | 0.23774819 | 3.55198315 | 0.00038234 |
| TAT1 | YBR069C | 1944.81632 | -0.2604841 | 0.07333604 | -3.5519243 | 0.00038242 |
| Xu_2009_SUTs23 | Xu_2009_SU | 53.1697903 | 0.75097498 | 0.21161645 | 3.54875525 | 0.00038706 |
| FKH1 | YIL131C | 722.141073 | -0.3208919 | 0.09042445 | -3.5487292 | 0.00038709 |
| GYP5 | YPL249C | 1009.10137 | -0.2672575 | 0.07531206 | -3.5486681 | 0.00038718 |
| Xu_2009_CUTs83 | Xu_2009_CU | 117.583793 | 0.51804884 | 0.14600385 | 3.54818614 | 0.00038789 |
| SMY2 | YBR172C | 2823.49725 | 0.2283195 | 0.0643617 | 3.54744339 | 0.00038899 |
| SEH1 | YGL100W | 776.287176 | -0.2669304 | 0.07525613 | -3.5469587 | 0.00038971 |
| PRS5 | YOL061W | 1213.74345 | -0.2768723 | 0.07807604 | -3.5461873 | 0.00039085 |
| SGA1 | YIL099W | 124.095363 | 0.52853485 | 0.14907029 | 3.54554125 | 0.00039181 |
| GPM3 | YOL056W | 283.912236 | -0.3722766 | 0.10507481 | -3.5429671 | 0.00039565 |
| TAF1 | YGR274C | 1355.97955 | 0.24257859 | 0.06850158 | 3.54121186 | 0.00039829 |
| GPI17 | YDR434W | 914.121893 | -0.250901 | 0.07089156 | -3.5392221 | 0.00040131 |
| PGC1 | YPL206C | 491.237158 | -0.339216 | 0.09587456 | -3.5381236 | 0.00040298 |
| AVT3 | YKL146W | 954.482138 | -0.2454149 | 0.06941835 | -3.5353038 | 0.00040731 |

| | | | | | | |
|---|---|---|---|---|---|---|
| YER076C | YER076C | 93.6933185 | -0.5896166 | 0.16678449 | -3.5352002 | 0.00040747 |
| Xu_2009_CUTs78 | Xu_2009_CU | 54.8256866 | 0.71128643 | 0.20134476 | 3.53267914 | 0.00041137 |
| RIO1 | YOR119C | 971.163069 | 0.29561065 | 0.08371368 | 3.53121075 | 0.00041366 |
| MTC5 | YDR128W | 1353.39739 | 0.22871417 | 0.06482426 | 3.52821901 | 0.00041837 |
| ATO3 | YDR384C | 198.445019 | -0.4841044 | 0.13721035 | -3.5281915 | 0.00041841 |
| BMT5 | YIL096C | 416.394988 | 0.38904682 | 0.11028815 | 3.52754865 | 0.00041943 |
| LSM1 | YJL124C | 900.647415 | -0.2828973 | 0.08026988 | -3.524327 | 0.00042456 |
| DIS3 | YOL021C | 2603.90808 | 0.22293129 | 0.06326697 | 3.5236597 | 0.00042563 |
| SPO11 | YHL022C | 42.447486 | -0.7850751 | 0.22286597 | -3.5226336 | 0.00042728 |
| YBR062C | YBR062C | 221.581447 | -0.4057931 | 0.11522989 | -3.5215957 | 0.00042896 |
| CBK1 | YNL161W | 1950.07493 | 0.24155237 | 0.06859639 | 3.52135678 | 0.00042934 |
| Xu_2009_CUTs13 | Xu_2009_CU | 30.8433416 | 0.87658505 | 0.24896346 | 3.52093856 | 0.00043002 |
| YCR024C-B | YCR024C-B | 918.640849 | -0.3057589 | 0.08696938 | -3.5157078 | 0.00043858 |
| Xu_2009_CUTs77 | Xu_2009_CU | 14.7516583 | 1.01515929 | 0.28880034 | 3.51509034 | 0.0004396 |
| SIA1 | YOR137C | 896.628044 | -0.2692565 | 0.076609 | -3.5146854 | 0.00044028 |
| YLR299C-A | YLR299C-A | 11.3350578 | 1.03102064 | 0.29339769 | 3.51407211 | 0.00044129 |
| HSP12 | YFL014W | 18228.0431 | -0.2310224 | 0.06577109 | -3.5125222 | 0.00044387 |
| GEA1 | YJR031C | 1108.10169 | 0.25345646 | 0.07216037 | 3.5124052 | 0.00044407 |
| TOF1 | YNL273W | 514.298758 | 0.3047165 | 0.08679263 | 3.51085665 | 0.00044667 |
| AGP2 | YBR132C | 538.298597 | -0.3005623 | 0.08563652 | -3.509745 | 0.00044854 |
| EMI1 | YDR512C | 415.285601 | -0.3371831 | 0.09606865 | -3.5098137 | 0.00044842 |
| RRF1 | YHR038W | 137.096076 | -0.5021935 | 0.14314 | -3.5084079 | 0.0004508 |
| RNH203 | YLR154C | 294.549399 | -0.3752479 | 0.10699243 | -3.5072377 | 0.00045278 |
| ATG12 | YBR217W | 197.310335 | -0.4293598 | 0.12250397 | -3.5048643 | 0.00045684 |
| Xu_2009_CUTs46 | Xu_2009_CU | 12.5912783 | 1.02603905 | 0.29290047 | 3.50302967 | 0.00046 |
| CSG2 | YBR036C | 1749.31655 | -0.2157479 | 0.06160369 | -3.5021908 | 0.00046145 |
| Xu_2009_CUTs58 | Xu_2009_CU | 13.1965887 | 1.00426816 | 0.28676962 | 3.50200329 | 0.00046177 |
| Xu_2009_CUTs67 | Xu_2009_CU | 39.925092 | 0.79230067 | 0.22634975 | 3.50033821 | 0.00046467 |
| OXP1 | YKL215C | 1289.00318 | 0.27170541 | 0.07762171 | 3.50037915 | 0.0004646 |
| INM1 | YHR046C | 312.696462 | 0.39103619 | 0.11172355 | 3.50003363 | 0.0004652 |
| YPS1 | YLR120C | 6329.34845 | -0.2009484 | 0.05742259 | -3.4994654 | 0.00046619 |
| FMP23 | YBR047W | 35.5064687 | -0.8516773 | 0.24339802 | -3.4991137 | 0.00046681 |
| SDH3 | YKL141W | 1532.93311 | -0.22506 | 0.0643503 | -3.49742 | 0.00046978 |
| HSP30 | YCR021C | 190.601545 | -0.4719028 | 0.13495656 | -3.4967015 | 0.00047105 |
| SSN3 | YPL042C | 828.401888 | -0.2487074 | 0.07112906 | -3.4965653 | 0.00047129 |
| REX3 | YLR107W | 966.08924 | 0.24163859 | 0.06911034 | 3.49641726 | 0.00047155 |
| Xu_2009_SUTs54 | Xu_2009_SU | 120.891867 | 0.54911944 | 0.15706279 | 3.49617783 | 0.00047197 |
| RUB1 | YDR139C | 441.324939 | -0.3714649 | 0.106256 | -3.4959429 | 0.00047239 |
| PRI1 | YIR008C | 365.446796 | -0.3394277 | 0.09710074 | -3.4956245 | 0.00047295 |
| Xu_2009_SUTs69 | Xu_2009_SU | 53.9271866 | 0.71855496 | 0.2055793 | 3.49526906 | 0.00047358 |
| CHS5 | YLR330W | 2886.00174 | 0.21063378 | 0.0602908 | 3.49363075 | 0.0004765 |

| | | | | | | |
|---|---|---|---|---|---|---|
| YGL152C | YGL152C | 25.9108794 | 0.88947855 | 0.25464768 | 3.49297729 | 0.00047767 |
| COX7 | YMR256C | 288.626914 | -0.3700108 | 0.10595708 | -3.4920816 | 0.00047927 |
| SPG5 | YMR191W | 987.1272 | -0.2529872 | 0.07246112 | -3.4913514 | 0.00048058 |
| PDR3 | YBL005W | 620.906948 | 0.28989729 | 0.08308226 | 3.48928019 | 0.00048432 |
| CSE2 | YNR010W | 151.709032 | -0.4686244 | 0.13431344 | -3.489036 | 0.00048477 |
| POL4 | YCR014C | 410.209137 | 0.33091507 | 0.09493298 | 3.48577556 | 0.00049071 |
| Xu_2009_SUTs36 | Xu_2009_SU | 170.244053 | 0.43960607 | 0.12612119 | 3.48558456 | 0.00049106 |
| ETP1 | YHL010C | 498.921598 | 0.33406001 | 0.09588086 | 3.48411592 | 0.00049377 |
| ARP9 | YMR033W | 1186.92276 | -0.2399401 | 0.06891263 | -3.4818009 | 0.00049805 |
| Xu_2009_SUTs64 | Xu_2009_SU | 333.453601 | 0.33591178 | 0.09648596 | 3.48145761 | 0.00049869 |
| Xu_2009_SUTs80 | Xu_2009_SU | 60.0741882 | 0.69065797 | 0.1984168 | 3.48084416 | 0.00049984 |
| EFM6 | YNL024C | 378.355595 | 0.40384797 | 0.11604685 | 3.48004254 | 0.00050133 |
| EPL1 | YFL024C | 1214.62557 | 0.24920708 | 0.0716169 | 3.4797247 | 0.00050193 |
| CDC1 | YDR182W | 1061.35296 | -0.2286892 | 0.06572672 | -3.4793948 | 0.00050255 |
| Xu_2009_CUTs54 | Xu_2009_CU | 102.478545 | 0.59488558 | 0.1710103 | 3.47865351 | 0.00050394 |
| MAK31 | YCR020C-A | 91.9235893 | -0.6146701 | 0.17669775 | -3.4786527 | 0.00050394 |
| YEA6 | YEL006W | 423.581206 | -0.3279559 | 0.09429032 | -3.4781505 | 0.00050489 |
| MF(ALPHA)1 | YPL187W | 13022.5019 | -0.2408285 | 0.06926714 | -3.476808 | 0.00050742 |
| Xu_2009_CUTs87 | Xu_2009_CU | 52.3248218 | 0.74458512 | 0.21424381 | 3.47541019 | 0.00051007 |
| SCO1 | YBR037C | 283.891418 | -0.3806251 | 0.1095283 | -3.4751305 | 0.0005106 |
| YLR302C | YLR302C | 20.0901884 | 0.93937268 | 0.27034931 | 3.47466271 | 0.0005115 |
| SAS3 | YBL052C | 751.424998 | 0.27999656 | 0.0805848 | 3.47455803 | 0.0005117 |
| DIA1 | YMR316W | 1367.43095 | 0.29577818 | 0.08513093 | 3.47439147 | 0.00051201 |
| IMA5 | YJL216C | 112.413797 | 0.56256606 | 0.16194269 | 3.47385884 | 0.00051303 |
| IRC8 | YJL051W | 748.351307 | 0.26039985 | 0.07497033 | 3.47337202 | 0.00051396 |
| Xu_2009_CUTs84 | Xu_2009_CU | 65.0867784 | 0.67180219 | 0.19353324 | 3.47124957 | 0.00051804 |
| PPN2 | YNL217W | 990.926909 | -0.2804627 | 0.08080226 | -3.4709756 | 0.00051857 |
| SUR1 | YPL057C | 2030.34342 | -0.2340231 | 0.06742136 | -3.4710525 | 0.00051842 |
| LST8 | YNL006W | 753.905545 | 0.28762732 | 0.08299318 | 3.46567412 | 0.0005289 |
| GAL11 | YOL051W | 2460.00305 | 0.21931592 | 0.06333878 | 3.46258507 | 0.00053501 |
| GCV3 | YAL044C | 1861.18761 | -0.2299145 | 0.06642929 | -3.4610408 | 0.00053809 |
| YNL228W | YNL228W | 22.7224452 | 0.90425638 | 0.26133044 | 3.46020308 | 0.00053977 |
| YOR131C | YOR131C | 414.017161 | -0.355551 | 0.10277493 | -3.4595111 | 0.00054116 |
| NSL1 | YPL233W | 623.950103 | 0.28374802 | 0.08201902 | 3.45953913 | 0.0005411 |
| SIR2 | YDL042C | 535.123656 | -0.2909864 | 0.08412052 | -3.4591613 | 0.00054186 |
| Xu_2009_SUTs75 | Xu_2009_SU | 66.3142513 | 0.69260944 | 0.20028512 | 3.45811727 | 0.00054396 |
| GIP3 | YPL137C | 2938.59833 | 0.20698061 | 0.05985869 | 3.45782069 | 0.00054456 |
| RAD1 | YPL022W | 1359.98054 | 0.23221204 | 0.06717228 | 3.45696242 | 0.0005463 |
| YFL064C | YFL064C | 29.6559902 | 0.83988442 | 0.24324739 | 3.45279936 | 0.0005548 |
| Xu_2009_SUTs79 | Xu_2009_SU | 225.074857 | 0.471826 | 0.13666474 | 3.45243399 | 0.00055555 |
| NCS6 | YGL211W | 738.642194 | 0.29441806 | 0.08530599 | 3.45131734 | 0.00055786 |

| | | | | | | |
|---|---|---|---|---|---|---|
| Xu_2009_CUTs84 | Xu_2009_CU | 15.7835424 | 0.96420687 | 0.27939828 | 3.45101216 | 0.00055849 |
| YIL001W | YIL001W | 344.736651 | 0.34296868 | 0.09939441 | 3.45058328 | 0.00055938 |
| Xu_2009_CUTs22 | Xu_2009_CU | 6.90584691 | 1.01627652 | 0.29461371 | 3.4495222 | 0.00056158 |
| Xu_2009_SUTs46 | Xu_2009_SU | 122.350415 | 0.50383018 | 0.14605773 | 3.44952773 | 0.00056157 |
| SFL1 | YOR140W | 1337.42418 | 0.23844907 | 0.06914187 | 3.44869283 | 0.00056331 |
| TRE1 | YPL176C | 927.015047 | 0.30928018 | 0.08973993 | 3.44640536 | 0.0005681 |
| CHC1 | YGL206C | 7772.76593 | -0.1859878 | 0.05401093 | -3.4435224 | 0.00057419 |
| ECM14 | YHR132C | 1129.88913 | -0.2312931 | 0.0671694 | -3.4434289 | 0.00057439 |
| MRX9 | YDL027C | 380.949444 | -0.3310133 | 0.09621927 | -3.4401977 | 0.00058129 |
| UBP14 | YBR058C | 1126.83338 | -0.230712 | 0.06706614 | -3.4400665 | 0.00058157 |
| TRM3 | YDL112W | 1908.69275 | 0.23779111 | 0.06917065 | 3.43774599 | 0.00058658 |
| DID2 | YKR035W-A | 767.577773 | -0.2574025 | 0.07487699 | -3.4376721 | 0.00058674 |
| YPR196W | YPR196W | 140.985818 | 0.48641108 | 0.14151402 | 3.43719345 | 0.00058778 |
| Xu_2009_CUTs37 | Xu_2009_CU | 50.3647018 | 0.77293734 | 0.22488491 | 3.43703516 | 0.00058812 |
| CSM2 | YIL132C | 82.4753652 | -0.6243687 | 0.18171758 | -3.4359293 | 0.00059053 |
| SHE2 | YKL130C | 318.479595 | -0.3877393 | 0.11287083 | -3.4352482 | 0.00059201 |
| RTT10 | YPL183C | 1997.70281 | 0.25594539 | 0.07450535 | 3.43526198 | 0.00059198 |
| YHR020W | YHR020W | 3038.96935 | -0.2181655 | 0.06353677 | -3.4336896 | 0.00059543 |
| YML133C | YML133C | 14.4460454 | 0.96766836 | 0.28180881 | 3.43377613 | 0.00059524 |
| PDR12 | YPL058C | 961.620903 | 0.24988558 | 0.07277391 | 3.43372463 | 0.00059535 |
| PCP1 | YGR101W | 540.289367 | -0.281093 | 0.08187864 | -3.4330441 | 0.00059685 |
| RIX1 | YHR197W | 1653.79199 | 0.27683355 | 0.08066163 | 3.43203503 | 0.00059907 |
| YDL109C | YDL109C | 370.775449 | 0.31812668 | 0.09277885 | 3.4288706 | 0.0006061 |
| Xu_2009_SUTs49 | Xu_2009_SU | 127.198568 | 0.51483008 | 0.15015616 | 3.42863109 | 0.00060663 |
| CBF1 | YJR060W | 656.359445 | 0.28833763 | 0.08411755 | 3.42779384 | 0.00060851 |
| SMF2 | YHR050W | 965.349421 | -0.2288819 | 0.06677791 | -3.4275094 | 0.00060915 |
| MCP1 | YOR228C | 211.085314 | -0.4004023 | 0.11682212 | -3.4274526 | 0.00060927 |
| SSA4 | YER103W | 1478.04158 | -0.2926394 | 0.08541977 | -3.4258977 | 0.00061277 |
| YEL068C | YEL068C | 182.382506 | -0.435096 | 0.12704555 | -3.4247247 | 0.00061542 |
| MPA43 | YNL249C | 167.94386 | 0.47428813 | 0.13849888 | 3.42449078 | 0.00061595 |
| TRS65 | YGR166W | 666.669984 | 0.27294492 | 0.07970913 | 3.42426163 | 0.00061647 |
| tK(CUU)M | tK(CUU)M | 18.9595333 | -0.9547504 | 0.27886102 | -3.42375 | 0.00061763 |
| VBA1 | YMR088C | 504.442386 | -0.2976029 | 0.0869489 | -3.4227337 | 0.00061995 |
| Xu_2009_SUTs87 | Xu_2009_SU | 67.1772826 | 0.64030675 | 0.18708159 | 3.42260695 | 0.00062024 |
| TEP1 | YNL128W | 92.1442982 | -0.5697194 | 0.16655349 | -3.4206394 | 0.00062474 |
| RTS3 | YGR161C | 2836.48099 | -0.2365988 | 0.06917234 | -3.4204252 | 0.00062523 |
| INO2 | YDR123C | 1402.60546 | 0.29971261 | 0.08771868 | 3.41674769 | 0.00063374 |
| Xu_2009_SUTs54 | Xu_2009_SU | 61.1044115 | -0.6604856 | 0.19332363 | -3.4164762 | 0.00063437 |
| DJP1 | YIR004W | 1024.9699 | 0.23499549 | 0.06879647 | 3.41580762 | 0.00063593 |
| TPM2 | YIL138C | 1494.83962 | -0.2144469 | 0.06281428 | -3.4139833 | 0.0006402 |
| ATG22 | YCL038C | 363.706651 | 0.33663727 | 0.09861997 | 3.41347979 | 0.00064139 |

| | | | | | | |
|---|---|---|---|---|---|---|
| Xu_2009_SUTs63! | Xu_2009_SU | 105.236524 | 0.53191229 | 0.15584961 | 3.41298429 | 0.00064256 |
| PEX9 | YMR018W | 102.081399 | 0.53680999 | 0.15730622 | 3.41251597 | 0.00064366 |
| TFB5 | YDR079C-A | 181.44819 | 0.44866751 | 0.13157566 | 3.40995827 | 0.00064973 |
| ECM7 | YLR443W | 557.87964 | 0.28333591 | 0.08316461 | 3.4069287 | 0.00065698 |
| Xu_2009_CUTs69 | Xu_2009_CU | 25.1385404 | 0.92096547 | 0.27040984 | 3.40581347 | 0.00065967 |
| BLI1 | YKL061W | 225.020635 | -0.4161254 | 0.12220395 | -3.405171 | 0.00066123 |
| Xu_2009_SUTs40( | Xu_2009_SU | 50.9919388 | 0.73161946 | 0.21490484 | 3.40438803 | 0.00066312 |
| DPH1 | YIL103W | 505.859243 | -0.3559704 | 0.10462197 | -3.4024441 | 0.00066786 |
| Xu_2009_CUTs59 | Xu_2009_CU | 37.8174228 | 0.76072553 | 0.22359444 | 3.40225594 | 0.00066832 |
| Xu_2009_CUTs87 | Xu_2009_CU | 14.9137563 | 0.98657715 | 0.28997945 | 3.40223131 | 0.00066838 |
| RPA12 | YJR063W | 581.183731 | -0.316719 | 0.09311654 | -3.4013187 | 0.00067062 |
| YMC2 | YBR104W | 546.093582 | 0.29329563 | 0.08623814 | 3.40099683 | 0.00067141 |
| DED1 | YOR204W | 5454.92395 | -0.2622876 | 0.07717643 | -3.3985449 | 0.00067745 |
| YET2 | YMR040W | 167.577852 | -0.4545329 | 0.13377964 | -3.3976239 | 0.00067974 |
| Xu_2009_CUTs43 | Xu_2009_CU | 52.5067792 | 0.67812161 | 0.19968919 | 3.39588534 | 0.00068407 |
| Xu_2009_SUTs68( | Xu_2009_SU | 17.8912325 | 0.95548321 | 0.28148972 | 3.39438043 | 0.00068784 |
| LTE1 | YAL024C | 872.80281 | 0.25295725 | 0.0745505 | 3.39309932 | 0.00069107 |
| URN1 | YPR152C | 393.929648 | 0.32283107 | 0.09516918 | 3.39218093 | 0.00069339 |
| Xu_2009_CUTs41 | Xu_2009_CU | 45.8915894 | 0.73068693 | 0.215453 | 3.39139833 | 0.00069537 |
| RPA34 | YJL148W | 1135.37464 | 0.27786906 | 0.08193572 | 3.39130547 | 0.00069561 |
| GUT1 | YHL032C | 549.066801 | -0.2900517 | 0.08553597 | -3.3909917 | 0.0006964 |
| Xu_2009_CUTs21 | Xu_2009_CU | 29.8255407 | 0.84762197 | 0.25003415 | 3.39002486 | 0.00069886 |
| YPT31 | YER031C | 526.680999 | -0.2879649 | 0.0849542 | -3.3896487 | 0.00069982 |
| FDO1 | YMR144W | 334.856411 | -0.3563145 | 0.105121 | -3.3895659 | 0.00070003 |
| PSF2 | YJL072C | 239.053913 | -0.3867873 | 0.11412436 | -3.3891743 | 0.00070103 |
| BNS1 | YGR230W | 242.850784 | 0.41048757 | 0.12112422 | 3.3889801 | 0.00070153 |
| RPG1 | YBR079C | 4260.21721 | -0.201265 | 0.0594058 | -3.3879682 | 0.00070412 |
| MRS6 | YOR370C | 1648.57735 | -0.2117297 | 0.06250145 | -3.3875974 | 0.00070508 |
| YPK2 | YMR104C | 835.037831 | 0.35877657 | 0.10594807 | 3.38634351 | 0.00070831 |
| Xu_2009_SUTs34: | Xu_2009_SU | 130.817145 | 0.52071013 | 0.1538256 | 3.38506805 | 0.00071161 |
| YER158C | YER158C | 893.364496 | 0.27762751 | 0.08201307 | 3.3851617 | 0.00071136 |
| CDA1 | YLR307W | 134.547573 | -0.5144372 | 0.15197218 | -3.3850746 | 0.00071159 |
| SYS1 | YJL004C | 284.591025 | -0.3640407 | 0.10758162 | -3.3838565 | 0.00071475 |
| YDR210W-B | YDR210W-B | 143.488841 | 0.49118941 | 0.14530284 | 3.38045303 | 0.00072366 |
| NRK1 | YNL129W | 190.019998 | -0.4271905 | 0.12636867 | -3.3805094 | 0.00072352 |
| UBP3 | YER151C | 2806.30148 | 0.19434055 | 0.05749662 | 3.3800342 | 0.00072477 |
| MHT1 | YLL062C | 65.0272602 | 0.6470566 | 0.19144384 | 3.37987692 | 0.00072518 |
| SIC1 | YLR079W | 1734.39501 | -0.217587 | 0.06437609 | -3.3799342 | 0.00072503 |
| YBL111C | YBL111C | 14.1811699 | 0.96745686 | 0.28631449 | 3.37900062 | 0.0007275 |
| TRP1 | YDR007W | 444.746619 | 0.30501632 | 0.0902685 | 3.37898973 | 0.00072753 |
| RCO1 | YMR075W | 868.91304 | -0.234095 | 0.06928825 | -3.3785668 | 0.00072865 |

| | | | | | | |
|---|---|---|---|---|---|---|
| Xu_2009_SUTs15! | Xu_2009_SU' | 144.137463 | 0.45390965 | 0.13440901 | 3.37707753 | 0.0007326 |
| Xu_2009_CUTs56 | Xu_2009_CU | 15.6311787 | 0.95531163 | 0.2832892 | 3.3722134 | 0.00074567 |
| VPS20 | YMR077C | 441.764746 | 0.31004849 | 0.09199751 | 3.37018331 | 0.00075118 |
| Xu_2009_SUTs69: | Xu_2009_SU' | 53.983576 | 0.71209873 | 0.21130951 | 3.36993219 | 0.00075187 |
| Xu_2009_CUTs47 | Xu_2009_CU | 50.6450018 | 0.69256396 | 0.20558087 | 3.36881518 | 0.00075492 |
| CFT1 | YDR301W | 1538.11509 | 0.22584774 | 0.06707137 | 3.36727472 | 0.00075915 |
| YCR064C | YCR064C | 37.9214101 | 0.77295134 | 0.22987 | 3.36255865 | 0.00077224 |
| Xu_2009_CUTs44 | Xu_2009_CU | 36.2798453 | 0.81827794 | 0.2433802 | 3.36213849 | 0.00077341 |
| YFR045W | YFR045W | 154.662155 | -0.461836 | 0.13743322 | -3.3604394 | 0.00077819 |
| Xu_2009_CUTs34 | Xu_2009_CU | 49.7031639 | 0.74405658 | 0.22142454 | 3.36031664 | 0.00077853 |
| Xu_2009_CUTs24 | Xu_2009_CU | 13.8396746 | 0.97265878 | 0.2894674 | 3.36016688 | 0.00077895 |
| YHR033W | YHR033W | 297.762501 | 0.3457968 | 0.10292673 | 3.35964055 | 0.00078044 |
| Xu_2009_SUTs68: | Xu_2009_SU' | 47.3826584 | 0.70765297 | 0.21067425 | 3.35899129 | 0.00078228 |
| PRS4 | YBL068W | 565.877488 | -0.2832099 | 0.08431637 | -3.3588962 | 0.00078254 |
| TMT1 | YER175C | 245.598745 | -0.375281 | 0.11173539 | -3.3586579 | 0.00078322 |
| RSM24 | YDR175C | 622.505389 | -0.2904878 | 0.08655449 | -3.3561261 | 0.00079043 |
| LRE1 | YCL051W | 699.676719 | -0.2789934 | 0.08313196 | -3.3560305 | 0.0007907 |
| NAM9 | YNL137C | 1538.87506 | -0.1997114 | 0.05952304 | -3.355195 | 0.00079309 |
| YGR146C-A | YGR146C-A | 40.104317 | 0.75493928 | 0.22504816 | 3.35456765 | 0.00079489 |
| NSI1 | YDR026C | 944.575898 | 0.23148729 | 0.0690164 | 3.35409116 | 0.00079626 |
| ILV1 | YER086W | 2281.40978 | -0.2180567 | 0.06506564 | -3.3513344 | 0.00080423 |
| RRP42 | YDL111C | 1200.59924 | -0.25175 | 0.07518918 | -3.3482218 | 0.00081332 |
| Xu_2009_SUTs34: | Xu_2009_SU' | 103.391227 | 0.5740205 | 0.17157161 | 3.34566136 | 0.00082087 |
| YHK8 | YHR048W | 119.192075 | 0.50729415 | 0.15166374 | 3.3448612 | 0.00082324 |
| LYP1 | YNL268W | 2279.28155 | -0.2214315 | 0.06620036 | -3.3448678 | 0.00082322 |
| TRM112 | YNR046W | 805.324807 | -0.250718 | 0.07498789 | -3.3434468 | 0.00082745 |
| LAM1 | YHR155W | 1566.07827 | 0.20192685 | 0.06043541 | 3.3412008 | 0.00083417 |
| Xu_2009_SUTs21 | Xu_2009_SU' | 56.6059514 | 0.65981495 | 0.19751512 | 3.3405795 | 0.00083604 |
| Xu_2009_SUTs36! | Xu_2009_SU' | 226.181939 | 0.40834831 | 0.1222627 | 3.33992562 | 0.00083801 |
| YLL067C | YLL067C | 16.1597591 | 0.94152191 | 0.28190612 | 3.33984203 | 0.00083826 |
| VPS17 | YOR132W | 799.166502 | -0.2540092 | 0.07608736 | -3.3383891 | 0.00084266 |
| SEC17 | YBL050W | 817.908979 | -0.2540244 | 0.07609919 | -3.3380696 | 0.00084363 |
| HMS1 | YOR032C | 386.519545 | 0.31147593 | 0.09331199 | 3.33800543 | 0.00084382 |
| RSC4 | YKR008W | 771.388437 | -0.2386379 | 0.07149508 | -3.3378223 | 0.00084438 |
| SCW4 | YGR279C | 5926.86436 | -0.193212 | 0.05794523 | -3.3343891 | 0.00085487 |
| Xu_2009_SUTs40! | Xu_2009_SU' | 73.4280076 | 0.6059762 | 0.1817809 | 3.3335526 | 0.00085744 |
| RRN11 | YML043C | 931.029136 | 0.24497585 | 0.0734974 | 3.33312276 | 0.00085877 |
| Xu_2009_CUTs91 | Xu_2009_CU | 111.669521 | 0.51715874 | 0.15519048 | 3.33241286 | 0.00086096 |
| YHR127W | YHR127W | 343.342857 | -0.3359949 | 0.10082825 | -3.3323492 | 0.00086116 |
| APM3 | YBR288C | 402.059004 | -0.3257265 | 0.09776442 | -3.3317487 | 0.00086302 |
| PMT3 | YOR321W | 459.862808 | -0.2983112 | 0.08954066 | -3.3315728 | 0.00086357 |

| | | | | | | |
|---|---|---|---|---|---|---|
| SPC19 | YDR201W | 122.701395 | -0.4924912 | 0.14786283 | -3.3307299 | 0.00086619 |
| SYO1 | YDL063C | 988.38675 | 0.32053125 | 0.0962588 | 3.32989037 | 0.0008688 |
| NCB2 | YDR397C | 531.472867 | -0.2881273 | 0.0865265 | -3.3299313 | 0.00086867 |
| IMA1 | YGR287C | 115.209431 | -0.5031448 | 0.15111774 | -3.3294888 | 0.00087006 |
| GIS2 | YNL255C | 4303.47076 | -0.1832014 | 0.05503733 | -3.3286759 | 0.0008726 |
| MSH1 | YHR120W | 488.159118 | 0.28990284 | 0.08710802 | 3.32808441 | 0.00087445 |
| Xu_2009_CUTs43 | Xu_2009_CU | 43.7939431 | 0.73537654 | 0.22097866 | 3.32781703 | 0.00087529 |
| PPN1 | YDR452W | 2109.23169 | 0.22275735 | 0.06694849 | 3.32729451 | 0.00087694 |
| KEG1 | YFR042W | 361.542157 | -0.3423259 | 0.10295423 | -3.3250296 | 0.00088409 |
| CRN1 | YLR429W | 2134.97315 | -0.2033804 | 0.06117099 | -3.3247854 | 0.00088487 |
| YAF9 | YNL107W | 288.42647 | -0.3413947 | 0.10268837 | -3.3245698 | 0.00088555 |
| Xu_2009_SUTs53 | Xu_2009_SU | 89.9108604 | -0.5683535 | 0.17097361 | -3.3242177 | 0.00088667 |
| TPO3 | YPR156C | 1415.19012 | 0.20675723 | 0.0622132 | 3.3233657 | 0.00088938 |
| FAT1 | YBR041W | 923.775261 | -0.2316607 | 0.06971029 | -3.3231922 | 0.00088994 |
| SPS19 | YNL202W | 80.3864348 | -0.5889556 | 0.17729899 | -3.3218213 | 0.00089432 |
| MAF1 | YDR005C | 430.177676 | -0.3135838 | 0.09444038 | -3.320442 | 0.00089875 |
| PHO8 | YDR481C | 1654.60245 | -0.2050381 | 0.06177293 | -3.3192227 | 0.00090268 |
| YLR046C | YLR046C | 216.474399 | -0.3943945 | 0.11882838 | -3.3190256 | 0.00090332 |
| RIM13 | YMR154C | 183.148968 | 0.43550328 | 0.13121177 | 3.31908702 | 0.00090312 |
| TOS8 | YGL096W | 246.383332 | 0.38903063 | 0.11733383 | 3.31558795 | 0.00091451 |
| Xu_2009_SUTs65 | Xu_2009_SU | 194.925211 | 0.40502688 | 0.12219014 | 3.31472649 | 0.00091733 |
| DPL1 | YDR294C | 1453.65378 | -0.2265355 | 0.06835113 | -3.3142897 | 0.00091876 |
| RRP7 | YCL031C | 1397.76831 | -0.2330379 | 0.07031551 | -3.3141752 | 0.00091914 |
| UTP22 | YGR090W | 3896.4082 | 0.21802548 | 0.06579203 | 3.31385847 | 0.00092018 |
| Xu_2009_CUTs34 | Xu_2009_CU | 45.760691 | 0.71831495 | 0.21677855 | 3.31358871 | 0.00092107 |
| Xu_2009_CUTs76 | Xu_2009_CU | 20.764734 | 0.89386807 | 0.26980177 | 3.31305485 | 0.00092283 |
| YLR108C | YLR108C | 354.728114 | -0.3549677 | 0.10718644 | -3.3116852 | 0.00092736 |
| PSE1 | YMR308C | 3250.80163 | 0.19843748 | 0.05992656 | 3.31134431 | 0.00092849 |
| CLB6 | YGR109C | 90.0085007 | -0.5580675 | 0.16855213 | -3.3109487 | 0.0009298 |
| MNE1 | YOR350C | 435.868626 | 0.29819347 | 0.09006433 | 3.31089417 | 0.00092998 |
| SYM1 | YLR251W | 252.668353 | -0.4159899 | 0.12569269 | -3.309579 | 0.00093436 |
| SNR8 | snR8 | 570.274561 | -0.3639856 | 0.11014417 | -3.3046285 | 0.00095102 |
| SNR85 | snR85 | 271.8412 | 0.36311954 | 0.10989 | 3.30439126 | 0.00095183 |
| TFB6 | YOR352W | 258.427936 | -0.3623902 | 0.10970107 | -3.3034337 | 0.00095509 |
| Xu_2009_SUTs57 | Xu_2009_SU | 320.419141 | -0.3392268 | 0.10269807 | -3.3031467 | 0.00095606 |
| ZRT3 | YKL175W | 1490.53353 | -0.2425457 | 0.07343691 | -3.3027763 | 0.00095733 |
| VPS45 | YGL095C | 833.215837 | 0.27814254 | 0.08426254 | 3.300904 | 0.00096374 |
| PHO85 | YPL031C | 777.211606 | -0.2624615 | 0.07955619 | -3.2990704 | 0.00097006 |
| MRPL40 | YPL173W | 789.391713 | -0.2577024 | 0.07812951 | -3.2984008 | 0.00097237 |
| Xu_2009_CUTs86 | Xu_2009_CU | 39.1829423 | 0.74328854 | 0.2253969 | 3.29768747 | 0.00097485 |
| tT(AGU)C | tT(AGU)C | 14.8175612 | -0.9464572 | 0.28705755 | -3.2970991 | 0.00097689 |

| | | | | | | |
|---|---|---|---|---|---|---|
| IRC11 | YOR013W | 61.8973493 | -0.6440403 | 0.19549382 | -3.2944279 | 0.00098622 |
| COS4 | YFL062W | 31.1351576 | 0.80221046 | 0.2436344 | 3.29268139 | 0.00099237 |
| CAF40 | YNL288W | 1044.47171 | 0.22118 | 0.06717269 | 3.29270697 | 0.00099228 |
| Xu_2009_SUTs59 | Xu_2009_SU | 249.588869 | 0.36671817 | 0.11143699 | 3.29081193 | 0.00099899 |
| MDJ1 | YFL016C | 1349.58459 | -0.266264 | 0.08092364 | -3.2903113 | 0.00100077 |
| YER156C | YER156C | 828.460091 | -0.2481089 | 0.07541602 | -3.2898697 | 0.00100234 |
| SUP45 | YBR143C | 3537.27846 | -0.1777591 | 0.05403565 | -3.289664 | 0.00100307 |
| LCB4 | YOR171C | 886.548753 | 0.2368076 | 0.07206473 | 3.28604024 | 0.00101606 |
| MRX8 | YDR336W | 230.910372 | -0.370759 | 0.11283773 | -3.2857715 | 0.00101703 |
| PDR17 | YNL264C | 451.260102 | -0.2890682 | 0.08797762 | -3.285702 | 0.00101729 |
| SIP1 | YDR422C | 1224.40735 | 0.21176639 | 0.06446394 | 3.28503634 | 0.00101969 |
| USE1 | YGL098W | 463.521169 | 0.28530058 | 0.08685524 | 3.28478265 | 0.00102061 |
| Xu_2009_SUTs59 | Xu_2009_SU | 114.647147 | 0.51450216 | 0.15669478 | 3.28346712 | 0.00102539 |
| GCV2 | YMR189W | 1577.53367 | 0.20118599 | 0.06127938 | 3.28309438 | 0.00102674 |
| Xu_2009_SUTs40 | Xu_2009_SU | 93.5924908 | 0.588629 | 0.17936341 | 3.28176736 | 0.00103159 |
| GUT2 | YIL155C | 1149.9457 | 0.237412 | 0.07234468 | 3.28167888 | 0.00103191 |
| YIR042C | YIR042C | 67.2688688 | 0.62747026 | 0.19120359 | 3.28168658 | 0.00103188 |
| Xu_2009_SUTs32 | Xu_2009_SU | 409.401397 | 0.33203307 | 0.10122937 | 3.2800074 | 0.00103804 |
| Xu_2009_CUTs35 | Xu_2009_CU | 66.1921858 | 0.66560277 | 0.20293429 | 3.27989307 | 0.00103846 |
| Xu_2009_CUTs66 | Xu_2009_CU | 20.3575922 | 0.88732615 | 0.27058577 | 3.27927873 | 0.00104073 |
| Xu_2009_SUTs36 | Xu_2009_SU | 68.8251028 | 0.61917611 | 0.18881771 | 3.27922681 | 0.00104092 |
| UGA4 | YDL210W | 50.7812327 | -0.689545 | 0.21032702 | -3.2784423 | 0.00104382 |
| RDH54 | YBR073W | 2033.49676 | 0.18231277 | 0.05563035 | 3.27721749 | 0.00104836 |
| KIP3 | YGL216W | 987.01576 | 0.21926793 | 0.06690736 | 3.27718683 | 0.00104847 |
| Xu_2009_CUTs67 | Xu_2009_CU | 21.753647 | 0.86690456 | 0.26456394 | 3.27672991 | 0.00105017 |
| RRI2 | YOL117W | 369.733523 | 0.31148607 | 0.09510858 | 3.27505752 | 0.0010564 |
| UBC11 | YOR339C | 79.1685322 | -0.6403372 | 0.19554739 | -3.2745884 | 0.00105816 |
| Xu_2009_CUTs78 | Xu_2009_CU | 54.7372468 | 0.68262965 | 0.20847154 | 3.27445009 | 0.00105868 |
| TOK1 | YJL093C | 707.515777 | -0.2744705 | 0.08384652 | -3.2734873 | 0.00106229 |
| CWC15 | YDR163W | 465.22333 | 0.29733447 | 0.09084555 | 3.27296675 | 0.00106425 |
| ENT5 | YDR153C | 1758.28788 | 0.2002445 | 0.06120056 | 3.27193901 | 0.00106813 |
| KAP114 | YGL241W | 620.958457 | 0.26728142 | 0.08169122 | 3.27185002 | 0.00106846 |
| Xu_2009_SUTs34 | Xu_2009_SU | 193.048568 | 0.42763903 | 0.1307268 | 3.27124225 | 0.00107076 |
| PDS1 | YDR113C | 305.340211 | -0.3825897 | 0.1170751 | -3.2679002 | 0.00108349 |
| PEP12 | YOR036W | 693.521967 | 0.2597904 | 0.07952329 | 3.26684672 | 0.00108753 |
| LAM5 | YFL042C | 796.680036 | 0.25407876 | 0.07781137 | 3.26531652 | 0.00109342 |
| YNL338W | YNL338W | 7.61082215 | 0.96464914 | 0.29579253 | 3.26123562 | 0.00110928 |
| YML090W | YML090W | 3.05339006 | 0.85131532 | 0.26107127 | 3.26085406 | 0.00111077 |
| Xu_2009_SUTs18 | Xu_2009_SU | 190.213168 | 0.4216299 | 0.12945662 | 3.25692037 | 0.00112628 |
| NIT2 | YJL126W | 194.192284 | -0.4066257 | 0.12485795 | -3.2567063 | 0.00112713 |
| Xu_2009_SUTs40 | Xu_2009_SU | 84.1598815 | 0.56860566 | 0.17471123 | 3.25454557 | 0.00113574 |

| | | | | | | |
|---|---|---|---|---|---|---|
| IMA2 | YOL157C | 121.557623 | -0.4884314 | 0.15020976 | -3.2516621 | 0.00114732 |
| Xu_2009_CUTs88 | Xu_2009_CU | 49.875426 | 0.69764526 | 0.21459619 | 3.25096758 | 0.00115013 |
| RBK1 | YCR036W | 1107.85914 | -0.2195054 | 0.06753366 | -3.2503119 | 0.00115278 |
| ENV9 | YOR246C | 984.721841 | -0.2343183 | 0.07210886 | -3.2495082 | 0.00115605 |
| tK(UUU)K | tK(UUU)K | 72.1896274 | -0.6033233 | 0.18568005 | -3.249263 | 0.00115704 |
| Xu_2009_CUTs90 | Xu_2009_CU | 36.1108628 | 0.77324012 | 0.23804355 | 3.24831374 | 0.00116091 |
| GUA1 | YMR217W | 3845.543 | -0.2715715 | 0.08363218 | -3.2472123 | 0.00116541 |
| DIA4 | YHR011W | 328.010544 | -0.3952092 | 0.12176491 | -3.2456739 | 0.00117173 |
| WIP1 | YDR374W-A | 131.342698 | -0.4776436 | 0.14718272 | -3.2452421 | 0.00117351 |
| MRPL4 | YLR439W | 358.829129 | -0.3204194 | 0.09876413 | -3.2442889 | 0.00117744 |
| PAP1 | YKR002W | 879.953367 | -0.238998 | 0.07372463 | -3.2417653 | 0.00118792 |
| Xu_2009_CUTs74 | Xu_2009_CU | 53.0881862 | 0.67795471 | 0.20926864 | 3.23963826 | 0.00119681 |
| Xu_2009_CUTs49 | Xu_2009_CU | 27.3161308 | 0.83488894 | 0.2578798 | 3.23751195 | 0.00120577 |
| YHR175W-A | YHR175W-A | 36.4862345 | -0.8434783 | 0.26065174 | -3.2360356 | 0.00121202 |
| ERP3 | YDL018C | 254.851981 | -0.3624809 | 0.11204069 | -3.2352615 | 0.00121531 |
| ERV2 | YPR037C | 472.262795 | -0.3360476 | 0.10388105 | -3.2349268 | 0.00121674 |
| YIL067C | YIL067C | 681.248144 | -0.2522449 | 0.07798428 | -3.2345608 | 0.0012183 |
| SNX41 | YDR425W | 520.553359 | -0.2991616 | 0.09250871 | -3.233875 | 0.00122123 |
| YOR186C-A | YOR186C-A | 26.4176512 | 0.83373652 | 0.25797606 | 3.23183679 | 0.00122997 |
| COS7 | YDL248W | 202.027646 | 0.38711083 | 0.11980594 | 3.23114893 | 0.00123294 |
| COR1 | YBL045C | 2417.45477 | -0.1988448 | 0.06155574 | -3.2303216 | 0.00123651 |
| Xu_2009_CUTs78 | Xu_2009_CU | 21.6099579 | 0.88399284 | 0.27367213 | 3.2301164 | 0.0012374 |
| ATG2 | YNL242W | 1217.55448 | 0.21817755 | 0.0675642 | 3.2291885 | 0.00124142 |
| TDA9 | YML081W | 2383.96206 | 0.17965958 | 0.05566114 | 3.22773823 | 0.00124773 |
| PUF6 | YDR496C | 2947.2518 | 0.21858581 | 0.06774122 | 3.2267768 | 0.00125193 |
| REG2 | YBR050C | 80.1465955 | -0.5601704 | 0.17365144 | -3.2258324 | 0.00125607 |
| RAD57 | YDR004W | 371.685069 | -0.3468519 | 0.10757028 | -3.2244215 | 0.00126227 |
| SRO77 | YBL106C | 394.857043 | 0.34932793 | 0.10836887 | 3.22350811 | 0.00126631 |
| PUF4 | YGL014W | 2164.71284 | -0.2104064 | 0.0653372 | -3.2203159 | 0.00128049 |
| YMR1 | YJR110W | 378.059813 | -0.3188868 | 0.09902343 | -3.2203162 | 0.00128049 |
| INM2 | YDR287W | 343.388372 | -0.3537592 | 0.10992244 | -3.2182622 | 0.0012897 |
| LDB18 | YLL049W | 184.065017 | -0.4609015 | 0.14325465 | -3.2173582 | 0.00129377 |
| PMT7 | YDR307W | 586.257378 | -0.2872335 | 0.08928117 | -3.2171793 | 0.00129458 |
| Xu_2009_SUTs26[illegible] | Xu_2009_SU[illegible] | 124.082666 | 0.50061608 | 0.15565664 | 3.2161562 | 0.0012992 |
| MVD1 | YNR043W | 1531.40943 | -0.2627773 | 0.08171648 | -3.2157192 | 0.00130118 |
| DOA1 | YKL213C | 2127.10612 | -0.1772779 | 0.05513471 | -3.2153584 | 0.00130282 |
| Xu_2009_SUTs82[illegible] | Xu_2009_SU[illegible] | 21.9425546 | 0.85127485 | 0.26481732 | 3.21457391 | 0.00130638 |
| tS(AGA)G | tS(AGA)G | 6.36843829 | -0.9432219 | 0.29351134 | -3.213579 | 0.00131092 |
| ATG18 | YFR021W | 524.830062 | -0.2770278 | 0.08622159 | -3.2129751 | 0.00131368 |
| YGL118C | YGL118C | 26.5399323 | 0.81060727 | 0.25244787 | 3.21098871 | 0.00132279 |
| RHB1 | YCR027C | 464.03769 | -0.2795894 | 0.08710786 | -3.2096924 | 0.00132877 |

| | | | | | | |
|---|---|---|---|---|---|---|
| Xu_2009_CUTs32 | Xu_2009_CU | 160.672372 | 0.44365984 | 0.13840909 | 3.20542423 | 0.00134864 |
| CDC15 | YAR019C | 632.340202 | 0.25171912 | 0.07853808 | 3.20505831 | 0.00135035 |
| VAM7 | YGL212W | 217.116396 | 0.38730331 | 0.12085912 | 3.20458491 | 0.00135257 |
| RPP1 | YHR062C | 321.135575 | -0.3357652 | 0.1048373 | -3.2027264 | 0.00136133 |
| Xu_2009_CUTs16 | Xu_2009_CU | 51.6549088 | 0.66234132 | 0.206844 | 3.20212967 | 0.00136416 |
| Xu_2009_SUTs10 | Xu_2009_SU | 88.1058645 | 0.55828681 | 0.17435711 | 3.20197338 | 0.0013649 |
| SNF4 | YGL115W | 695.918133 | -0.2572215 | 0.08033515 | -3.2018554 | 0.00136546 |
| Xu_2009_SUTs72 | Xu_2009_SU | 47.5488468 | 0.68001939 | 0.21242276 | 3.20125486 | 0.0013683 |
| Xu_2009_CUTs71 | Xu_2009_CU | 63.7707801 | 0.61125252 | 0.19103367 | 3.19971102 | 0.00137565 |
| DLD1 | YDL174C | 1418.2156 | -0.1979795 | 0.06189312 | -3.1987317 | 0.00138034 |
| DAP1 | YPL170W | 537.876896 | 0.30756942 | 0.09615896 | 3.19855194 | 0.0013812 |
| COX6 | YHR051W | 250.972636 | -0.3548253 | 0.11096029 | -3.1977686 | 0.00138495 |
| PCM1 | YEL058W | 1358.32121 | -0.2338566 | 0.07314014 | -3.1973766 | 0.00138684 |
| MLO50 | YLR049C | 365.869013 | -0.3109965 | 0.09740047 | -3.1929677 | 0.00140819 |
| MHF1 | YOL086W-A | 300.797889 | -0.351043 | 0.10998539 | -3.1917244 | 0.00141426 |
| VPS25 | YJR102C | 343.637028 | -0.353368 | 0.11076681 | -3.1901976 | 0.00142176 |
| CCW14 | YLR390W-A | 5563.44606 | -0.1605405 | 0.05033305 | -3.1895644 | 0.00142487 |
| BSC6 | YOL137W | 572.750715 | 0.26521242 | 0.08317394 | 3.18864804 | 0.0014294 |
| PYC2 | YBR218C | 3755.03859 | -0.1679898 | 0.05269164 | -3.188168 | 0.00143177 |
| MMR1 | YLR190W | 1865.62246 | 0.19627854 | 0.06158271 | 3.18723436 | 0.0014364 |
| MYO5 | YMR109W | 2315.79374 | -0.1749546 | 0.05489708 | -3.1869565 | 0.00143778 |
| CUS2 | YNL286W | 442.067046 | 0.2943475 | 0.09240928 | 3.18525913 | 0.00144624 |
| ELM1 | YKL048C | 521.948668 | -0.2647698 | 0.08314578 | -3.1844042 | 0.00145052 |
| Xu_2009_SUTs60 | Xu_2009_SU | 42.4541415 | 0.68983693 | 0.21664734 | 3.18414683 | 0.00145181 |
| RRP5 | YMR229C | 5747.35849 | 0.20146567 | 0.06328019 | 3.18370833 | 0.00145401 |
| YML096W | YML096W | 339.251511 | 0.32311993 | 0.10153433 | 3.18237109 | 0.00146075 |
| Xu_2009_SUTs27 | Xu_2009_SU | 10.7413341 | 0.93379539 | 0.2936091 | 3.18040345 | 0.0014707 |
| Xu_2009_CUTs70 | Xu_2009_CU | 54.7401536 | 0.64491253 | 0.20279671 | 3.18009362 | 0.00147227 |
| YHP1 | YDR451C | 437.722411 | -0.3024184 | 0.09511842 | -3.1793886 | 0.00147586 |
| GTT3 | YEL017W | 397.371815 | -0.3378098 | 0.10626852 | -3.1788323 | 0.0014787 |
| MBP1 | YDL056W | 1100.74549 | 0.21043983 | 0.06621459 | 3.17814875 | 0.00148219 |
| YGL182C | YGL182C | 35.1551331 | 0.76546574 | 0.24091137 | 3.17737497 | 0.00148615 |
| YBL081W | YBL081W | 1005.01998 | 0.31022814 | 0.09769275 | 3.17554917 | 0.00149553 |
| IWR1 | YDL115C | 634.095261 | 0.27477049 | 0.08653411 | 3.17528519 | 0.00149689 |
| Xu_2009_CUTs31 | Xu_2009_CU | 65.5614075 | 0.62472631 | 0.19679529 | 3.17449821 | 0.00150096 |
| Xu_2009_SUTs44 | Xu_2009_SU | 133.801979 | 0.49347483 | 0.1554713 | 3.17405741 | 0.00150324 |
| ISU2 | YOR226C | 274.386339 | -0.3712464 | 0.11699324 | -3.1732292 | 0.00150753 |
| AAH1 | YNL141W | 1410.8098 | -0.2911812 | 0.09183166 | -3.1708147 | 0.00152012 |
| Xu_2009_SUTs11 | Xu_2009_SU | 28.8070594 | 0.832815 | 0.2627475 | 3.16964008 | 0.00152628 |
| TMA64 | YDR117C | 537.850252 | -0.293212 | 0.09255231 | -3.1680683 | 0.00153455 |
| PHB2 | YGR231C | 1931.39938 | -0.2385338 | 0.07529431 | -3.1680193 | 0.00153481 |

| | | | | | | |
|---|---|---|---|---|---|---|
| DPH6 | YLR143W | 755.776926 | 0.24345642 | 0.07685487 | 3.16774226 | 0.00153628 |
| MRPL44 | YMR225C | 503.459329 | -0.2718543 | 0.08584337 | -3.1668645 | 0.00154092 |
| PCH2 | YBR186W | 60.3133218 | 0.6263206 | 0.19784397 | 3.16573009 | 0.00154694 |
| PFD1 | YJL179W | 475.589683 | -0.2849418 | 0.09009532 | -3.1626702 | 0.00156329 |
| TAF13 | YML098W | 337.882935 | 0.33495771 | 0.1059366 | 3.16186949 | 0.0015676 |
| ROF1 | YHR177W | 194.981614 | 0.39266615 | 0.12421068 | 3.1612913 | 0.00157071 |
| SMC6 | YLR383W | 536.625485 | 0.27757104 | 0.08781937 | 3.16070414 | 0.00157388 |
| PHO23 | YNL097C | 427.159465 | -0.2958848 | 0.09361344 | -3.160709 | 0.00157386 |
| Xu_2009_CUTs26 | Xu_2009_CU | 9.97996433 | 0.93543363 | 0.29606042 | 3.15960378 | 0.00157984 |
| CKS1 | YBR135W | 999.317979 | -0.2689936 | 0.0851553 | -3.1588588 | 0.00158388 |
| YEL077W-A | YEL077W-A | 3.64488434 | 0.85857322 | 0.27184392 | 3.15833158 | 0.00158675 |
| YLL059C | YLL059C | 91.2103491 | 0.56624481 | 0.17935465 | 3.15712361 | 0.00159334 |
| PUS7 | YOR243C | 1427.62266 | 0.24878304 | 0.07880712 | 3.1568599 | 0.00159478 |
| DST1 | YGL043W | 685.899521 | -0.2470661 | 0.07828739 | -3.1558858 | 0.00160011 |
| TAD1 | YGL243W | 482.95116 | 0.2882339 | 0.09141166 | 3.15314136 | 0.00161524 |
| ACO1 | YLR304C | 2801.10691 | -0.4096925 | 0.12993084 | -3.1531579 | 0.00161514 |
| Xu_2009_CUTs81 | Xu_2009_CU | 30.7085652 | 0.79822664 | 0.25316643 | 3.15297192 | 0.00161617 |
| TRS23 | YDR246W | 535.018015 | -0.2652716 | 0.08418306 | -3.1511276 | 0.00162641 |
| Xu_2009_CUTs43 | Xu_2009_CU | 14.1571854 | 0.90011669 | 0.28569485 | 3.15062276 | 0.00162923 |
| Xu_2009_CUTs50 | Xu_2009_CU | 52.4847986 | 0.64096518 | 0.2034454 | 3.15055134 | 0.00162963 |
| YDL158C | YDL158C | 28.0502065 | -0.796598 | 0.25289138 | -3.149961 | 0.00163292 |
| MET10 | YFR030W | 740.126376 | 0.24306676 | 0.07717507 | 3.14955026 | 0.00163522 |
| YOR292C | YOR292C | 264.316004 | -0.3452717 | 0.10970287 | -3.1473348 | 0.00164766 |
| MSO1 | YNR049C | 353.130834 | -0.3081412 | 0.09791213 | -3.1471195 | 0.00164888 |
| NTA1 | YJR062C | 472.401513 | 0.26834028 | 0.08533895 | 3.1444058 | 0.00166424 |
| Xu_2009_CUTs32 | Xu_2009_CU | 34.1411201 | 0.79104326 | 0.25159724 | 3.14408563 | 0.00166607 |
| UTP8 | YGR128C | 1951.88695 | 0.24086307 | 0.07664283 | 3.14266943 | 0.00167415 |
| YDR269C | YDR269C | 6.20367856 | 0.91933388 | 0.2927229 | 3.14062848 | 0.00168586 |
| IMP1 | YMR150C | 291.299679 | -0.3347411 | 0.10658126 | -3.1407124 | 0.00168537 |
| GDT1 | YBR187W | 467.858546 | -0.2768914 | 0.08817406 | -3.1402816 | 0.00168785 |
| AUR1 | YKL004W | 1110.61948 | -0.2130729 | 0.06786315 | -3.139744 | 0.00169096 |
| HIS4 | YCL030C | 673.134992 | -0.2503327 | 0.0797638 | -3.138425 | 0.00169858 |
| Xu_2009_CUTs58 | Xu_2009_CU | 4.66777107 | 0.89859861 | 0.28645499 | 3.13696268 | 0.00170708 |
| CLF1 | YLR117C | 846.921184 | 0.22806769 | 0.07270146 | 3.13704422 | 0.0017066 |
| Xu_2009_SUTs84 | Xu_2009_SU | 121.251388 | 0.46337803 | 0.14774693 | 3.13629551 | 0.00171097 |
| Xu_2009_CUTs27 | Xu_2009_CU | 44.0093418 | 0.69058635 | 0.22021838 | 3.13591599 | 0.00171318 |
| Xu_2009_SUTs25 | Xu_2009_SU | 139.660144 | 0.44947483 | 0.14335478 | 3.13540175 | 0.00171619 |
| YGL262W | YGL262W | 9.40810741 | 0.92692155 | 0.29565611 | 3.13513412 | 0.00171776 |
| PEX18 | YHR160C | 79.7345195 | -0.5554405 | 0.17716956 | -3.1350784 | 0.00171808 |
| Xu_2009_CUTs85 | Xu_2009_CU | 95.0918675 | 0.51732859 | 0.16501965 | 3.13495138 | 0.00171883 |
| MFB1 | YDR219C | 533.398767 | -0.2999462 | 0.09571672 | -3.133687 | 0.00172625 |

| | | | | | | |
|---|---|---|---|---|---|---|
| YKL071W | YKL071W | 58.6920946 | -0.6541333 | 0.20876191 | -3.1333942 | 0.00172797 |
| tY(GUA)J2/SUP5 | tY(GUA)M1 | 17.6162831 | -0.8785894 | 0.28070033 | -3.1299908 | 0.00174812 |
| YGR069W | YGR069W | 28.7147207 | 0.78657242 | 0.25152067 | 3.12726747 | 0.00176439 |
| YBR298C-A | YBR298C-A | 4.27784031 | 0.87808817 | 0.28080304 | 3.12706071 | 0.00176563 |
| URB2 | YJR041C | 1219.5375 | 0.24362204 | 0.07791267 | 3.12686023 | 0.00176684 |
| JJJ3 | YJR097W | 379.794387 | 0.33937998 | 0.10853849 | 3.12681687 | 0.0017671 |
| SNO4 | YMR322C | 20.4599306 | 0.83428761 | 0.26697112 | 3.12501076 | 0.00177799 |
| YHL019W-A | YHL019W-A | 29.363894 | 0.77650224 | 0.24854075 | 3.12424513 | 0.00178262 |
| SPO20 | YMR017W | 44.2553827 | 0.70756637 | 0.22649888 | 3.12392877 | 0.00178454 |
| Xu_2009_CUTs15 | Xu_2009_CU | 116.965606 | 0.50109321 | 0.1604557 | 3.12293804 | 0.00179055 |
| UTP5 | YDR398W | 2357.72228 | 0.22576451 | 0.07229938 | 3.12263413 | 0.0017924 |
| CRR1 | YLR213C | 113.932047 | -0.4791637 | 0.15345869 | -3.1224282 | 0.00179366 |
| Xu_2009_SUTs82 | Xu_2009_SU | 108.002958 | 0.51526047 | 0.16505979 | 3.12165962 | 0.00179835 |
| GCD10 | YNL062C | 783.840015 | 0.25612775 | 0.08204843 | 3.12166543 | 0.00179831 |
| Xu_2009_CUTs40 | Xu_2009_CU | 29.1593448 | 0.79652894 | 0.25524125 | 3.12069047 | 0.00180428 |
| Xu_2009_SUTs32 | Xu_2009_SU | 270.761703 | -0.3693839 | 0.11836723 | -3.1206604 | 0.00180446 |
| GCR2 | YNL199C | 1258.9585 | 0.20458203 | 0.06556116 | 3.12047621 | 0.00180559 |
| APL6 | YGR261C | 1470.3425 | -0.2111401 | 0.0676846 | -3.1194708 | 0.00181176 |
| UPF3 | YGR072W | 359.134918 | 0.30161445 | 0.09674538 | 3.11761097 | 0.00182323 |
| NMD3 | YHR170W | 4248.93613 | 0.22939355 | 0.07359735 | 3.11687262 | 0.00182781 |
| FZF1 | YGL254W | 334.372988 | 0.30644988 | 0.09833526 | 3.11637841 | 0.00183087 |
| ATP12 | YJL180C | 231.262488 | -0.370232 | 0.11883469 | -3.1155216 | 0.0018362 |
| PEP5 | YMR231W | 953.449203 | 0.25096238 | 0.08058835 | 3.11412716 | 0.0018449 |
| PEX2 | YJL210W | 292.010235 | -0.3293523 | 0.10578652 | -3.1133678 | 0.00184965 |
| YDR271C | YDR271C | 21.4689727 | 0.83622544 | 0.26870625 | 3.11204315 | 0.00185797 |
| MRPL27 | YBR282W | 295.506738 | -0.3209438 | 0.10315299 | -3.111338 | 0.00186242 |
| YJU3 | YKL094W | 1615.64268 | -0.2043555 | 0.06569173 | -3.1108248 | 0.00186566 |
| MRI1 | YPR118W | 766.134718 | -0.2321493 | 0.07478371 | -3.1042765 | 0.00190745 |
| YIL100C-A | YIL100C-A | 34.8332253 | 0.72825941 | 0.23466002 | 3.10346609 | 0.00191268 |
| SNR52 | snR52 | 22.5314087 | -0.8140337 | 0.26232989 | -3.1030918 | 0.0019151 |
| MDV1 | YJL112W | 783.818288 | 0.2390041 | 0.07702514 | 3.10293641 | 0.00191611 |
| MRS1 | YIR021W | 394.264815 | -0.3001863 | 0.09676149 | -3.1023325 | 0.00192002 |
| Xu_2009_CUTs24 | Xu_2009_CU | 65.5578297 | 0.58668805 | 0.18914215 | 3.10183667 | 0.00192324 |
| Xu_2009_CUTs34 | Xu_2009_CU | 50.2806009 | 0.70739058 | 0.22828847 | 3.09866982 | 0.00194392 |
| PEX21 | YGR239C | 311.233816 | 0.33571372 | 0.10834816 | 3.09847174 | 0.00194522 |
| YJR154W | YJR154W | 446.620663 | 0.27740031 | 0.08953896 | 3.09809635 | 0.00194768 |
| YHC3 | YJL059W | 108.874399 | 0.4810893 | 0.15534415 | 3.09692581 | 0.00195539 |
| PSY3 | YLR376C | 309.012148 | 0.31897052 | 0.1030216 | 3.09615181 | 0.0019605 |
| Xu_2009_SUTs57 | Xu_2009_SU | 54.2355509 | 0.65062471 | 0.2102701 | 3.09423317 | 0.00197322 |
| Xu_2009_CUTs82 | Xu_2009_CU | 17.0892884 | 0.86976224 | 0.28117784 | 3.09328162 | 0.00197956 |
| NFS1 | YCL017C | 968.202774 | -0.2214579 | 0.07162149 | -3.0920594 | 0.00198773 |

| | | | | | | |
|---|---|---|---|---|---|---|
| HDA1 | YNL021W | 1244.96005 | 0.1980943 | 0.06408393 | 3.09116955 | 0.0019937 |
| YJR018W | YJR018W | 25.6306881 | 0.81659836 | 0.26426023 | 3.09012963 | 0.00200069 |
| FOL1 | YNL256W | 1131.86568 | -0.2591238 | 0.08386727 | -3.089689 | 0.00200366 |
| YKL053W | YKL053W | 22.213924 | 0.80793864 | 0.26156066 | 3.08891501 | 0.00200889 |
| Xu_2009_CUTs37 | Xu_2009_CU | 39.4133467 | 0.71091682 | 0.23017297 | 3.08861996 | 0.00201088 |
| STE12 | YHR084W | 1389.88547 | 0.2291347 | 0.07423561 | 3.08658743 | 0.00202468 |
| Xu_2009_SUTs48 | Xu_2009_SU | 196.896574 | 0.3718555 | 0.12051524 | 3.08554767 | 0.00203178 |
| EXO5 | YBR163W | 691.028747 | -0.2548201 | 0.0825984 | -3.0850483 | 0.00203519 |
| Xu_2009_SUTs76 | Xu_2009_SU | 40.5234767 | 0.68621077 | 0.22245084 | 3.08477482 | 0.00203706 |
| GRE3 | YHR104W | 3212.43226 | -0.4123492 | 0.13373504 | -3.0833296 | 0.00204698 |
| INP51 | YIL002C | 988.024444 | 0.27727738 | 0.08994523 | 3.08273569 | 0.00205107 |
| Xu_2009_CUTs87 | Xu_2009_CU | 38.7666365 | 0.686865 | 0.22282158 | 3.08257848 | 0.00205216 |
| Xu_2009_SUTs22 | Xu_2009_SU | 91.5184836 | 0.50956395 | 0.16541637 | 3.08049288 | 0.00206658 |
| YPD1 | YDL235C | 953.576212 | -0.2133096 | 0.06926179 | -3.0797586 | 0.00207168 |
| HGH1 | YGR187C | 432.173682 | 0.30585106 | 0.09936045 | 3.07819725 | 0.00208257 |
| YDR444W | YDR444W | 555.160167 | 0.29290482 | 0.09518781 | 3.07712538 | 0.00209007 |
| SLA1 | YBL007C | 4191.75197 | -0.1514671 | 0.04924697 | -3.0756629 | 0.00210035 |
| SAP1 | YER047C | 819.543054 | 0.22871394 | 0.07437391 | 3.07519037 | 0.00210368 |
| Xu_2009_SUTs55 | Xu_2009_SU | 103.112065 | -0.5250764 | 0.17077895 | -3.0745966 | 0.00210787 |
| GCS1 | YDL226C | 1108.60676 | -0.1989402 | 0.0647341 | -3.0731893 | 0.00211784 |
| ISY1 | YJR050W | 327.549887 | -0.3369635 | 0.10974617 | -3.0703898 | 0.0021378 |
| ECM31 | YBR176W | 273.149976 | -0.3423261 | 0.1115074 | -3.0699857 | 0.00214069 |
| Xu_2009_CUTs63 | Xu_2009_CU | 27.1729639 | 0.76438341 | 0.24900984 | 3.0696916 | 0.0021428 |
| Xu_2009_CUTs18 | Xu_2009_CU | 4.69376119 | 0.86306929 | 0.28119415 | 3.06930027 | 0.00214561 |
| Xu_2009_CUTs13 | Xu_2009_CU | 24.1811545 | 0.79033725 | 0.25757025 | 3.06843378 | 0.00215184 |
| RSM23 | YGL129C | 833.012804 | -0.2217089 | 0.07226667 | -3.0679274 | 0.00215549 |
| YBR232C | YBR232C | 8.8022061 | 0.9077936 | 0.29597988 | 3.06707878 | 0.00216162 |
| UBA4 | YHR111W | 354.333744 | -0.3050732 | 0.09950896 | -3.065786 | 0.00217099 |
| TDA2 | YER071C | 183.995248 | -0.3899567 | 0.12722132 | -3.0651839 | 0.00217536 |
| PTC3 | YBL056W | 1341.54196 | -0.1929188 | 0.06294836 | -3.0647156 | 0.00217877 |
| KRE1 | YNL322C | 1991.78929 | -0.1880652 | 0.06137544 | -3.0641775 | 0.00218269 |
| PRM8 | YGL053W | 441.3814 | -0.2823233 | 0.09214221 | -3.0639956 | 0.00218402 |
| YCL007C | YCL007C | 41.3738812 | 0.70492976 | 0.23025351 | 3.0615375 | 0.00220203 |
| Xu_2009_CUTs56 | Xu_2009_CU | 27.2941851 | 0.79197577 | 0.25872689 | 3.06104935 | 0.00220563 |
| TDA1 | YMR291W | 2016.26061 | -0.1897471 | 0.06206754 | -3.057107 | 0.00223485 |
| SNR39 | snR39 | 101.75774 | 0.51024351 | 0.16691292 | 3.05694426 | 0.00223606 |
| Xu_2009_SUTs15 | Xu_2009_SU | 118.725101 | 0.46267511 | 0.15137025 | 3.05657897 | 0.00223879 |
| BI3 | Q0115 | 11.8256178 | -0.9057722 | 0.29637369 | -3.0561828 | 0.00224175 |
| SIR3 | YLR442C | 560.914511 | 0.27804714 | 0.09098328 | 3.05602462 | 0.00224293 |
| SEF1 | YBL066C | 946.427637 | 0.23122159 | 0.07566625 | 3.05580899 | 0.00224454 |
| SER3 | YER081W | 836.009621 | -0.2340895 | 0.07664263 | -3.0542984 | 0.00225588 |

| | | | | | | |
|---|---|---|---|---|---|---|
| EMW1 | YNL313C | 1723.3836 | 0.19521776 | 0.06391678 | 3.05424872 | 0.00225625 |
| Xu_2009_SUTs32 | Xu_2009_SU | 175.961516 | -0.3841919 | 0.12588467 | -3.0519358 | 0.00227371 |
| Xu_2009_CUTs22 | Xu_2009_CU | 46.1021419 | 0.64812041 | 0.21248097 | 3.05025165 | 0.0022865 |
| Xu_2009_SUTs65 | Xu_2009_SU | 25.5658652 | 0.80039028 | 0.26241507 | 3.05009261 | 0.00228771 |
| Xu_2009_CUTs13 | Xu_2009_CU | 22.1284147 | 0.80363844 | 0.26350718 | 3.04977809 | 0.00229011 |
| YKL222C | YKL222C | 444.452089 | 0.27823167 | 0.09124108 | 3.04941235 | 0.0022929 |
| Xu_2009_CUTs35 | Xu_2009_CU | 53.3193925 | 0.62189401 | 0.20396139 | 3.0490771 | 0.00229546 |
| Xu_2009_CUTs17 | Xu_2009_CU | 35.9350669 | 0.70710883 | 0.23193662 | 3.04871579 | 0.00229822 |
| IDI1 | YPL117C | 2767.33625 | -0.1608308 | 0.05277267 | -3.047615 | 0.00230665 |
| MTC4 | YBR255W | 640.164146 | 0.25224808 | 0.08278326 | 3.04709041 | 0.00231068 |
| JIP4 | YDR475C | 1008.07184 | 0.22158239 | 0.07272087 | 3.04702598 | 0.00231118 |
| YJL156W-A | YJL156W-A | 25.8015617 | 0.77059976 | 0.25297248 | 3.04618019 | 0.00231769 |
| Xu_2009_CUTs51 | Xu_2009_CU | 56.7627916 | 0.64976289 | 0.21335146 | 3.04550475 | 0.0023229 |
| SFT2 | YBL102W | 424.68498 | -0.2713675 | 0.08913881 | -3.0443247 | 0.00233203 |
| Xu_2009_SUTs54 | Xu_2009_SU | 19.4512823 | 0.83511272 | 0.27433714 | 3.04411108 | 0.00233369 |
| TSR1 | YDL060W | 2061.98863 | 0.23420464 | 0.07709117 | 3.0380215 | 0.00238137 |
| Xu_2009_CUTs82 | Xu_2009_CU | 36.825529 | 0.70644495 | 0.23277139 | 3.03493028 | 0.00240591 |
| SPL2 | YHR136C | 10.7030696 | 0.89290628 | 0.29430104 | 3.03398951 | 0.00241343 |
| SYF1 | YDR416W | 572.53719 | 0.26874835 | 0.0885984 | 3.03333171 | 0.0024187 |
| YGL082W | YGL082W | 786.377615 | -0.2308191 | 0.07609848 | -3.0331633 | 0.00242005 |
| UTH1 | YKR042W | 7421.13348 | -0.2425887 | 0.07998314 | -3.032998 | 0.00242137 |
| GAS5 | YOL030W | 2437.45503 | -0.1609502 | 0.05308933 | -3.031686 | 0.00243192 |
| VHC1 | YBR235W | 815.215147 | 0.22397753 | 0.07390991 | 3.03041272 | 0.0024422 |
| PHR1 | YOR386W | 410.918664 | 0.30016467 | 0.09906045 | 3.03011624 | 0.0024446 |
| TIF4631 | YGR162W | 5878.62475 | 0.16784225 | 0.05542523 | 3.0282642 | 0.00245963 |
| YPL245W | YPL245W | 782.207881 | 0.24297347 | 0.08032236 | 3.02497927 | 0.0024865 |
| Xu_2009_CUTs40 | Xu_2009_CU | 4.6131518 | 0.86350055 | 0.28547451 | 3.02479041 | 0.00248805 |
| Xu_2009_SUTs38 | Xu_2009_SU | 79.0613232 | 0.54313239 | 0.17959032 | 3.02428546 | 0.00249221 |
| AAR2 | YBL074C | 309.264192 | 0.31353529 | 0.10375455 | 3.02189438 | 0.00251198 |
| URA7 | YBL039C | 2920.95673 | -0.2283253 | 0.0756062 | -3.0199288 | 0.00252834 |
| Xu_2009_SUTs62 | Xu_2009_SU | 35.3413074 | -0.732052 | 0.24265096 | -3.0168929 | 0.0025538 |
| Xu_2009_SUTs62 | Xu_2009_SU | 57.3161053 | -0.5981843 | 0.19829481 | -3.016641 | 0.00255592 |
| Xu_2009_SUTs60 | Xu_2009_SU | 51.8745938 | 0.64798636 | 0.21482606 | 3.01633034 | 0.00255854 |
| Xu_2009_CUTs92 | Xu_2009_CU | 53.3362038 | 0.62926217 | 0.2087811 | 3.01398049 | 0.00257844 |
| EST1 | YLR233C | 263.381331 | 0.33026835 | 0.10958863 | 3.01370998 | 0.00258074 |
| Xu_2009_SUTs52 | Xu_2009_SU | 17.5670046 | 0.82891912 | 0.2753207 | 3.01074024 | 0.00260612 |
| MRPL37 | YBR268W | 302.401408 | -0.3199556 | 0.1062863 | -3.0103183 | 0.00260974 |
| Xu_2009_CUTs52 | Xu_2009_CU | 43.3340605 | 0.64474541 | 0.21438694 | 3.00739127 | 0.002635 |
| CDC20 | YGL116W | 692.22897 | 0.2380373 | 0.07915958 | 3.007056 | 0.00263791 |
| UTP10 | YJL109C | 4105.88785 | 0.22620203 | 0.07522481 | 3.00701361 | 0.00263828 |
| NAS2 | YIL007C | 338.076019 | -0.3247542 | 0.10804355 | -3.0057713 | 0.00264908 |

| | | | | | | |
|---|---|---|---|---|---|---|
| DHH1 | YDL160C | 2436.0805 | -0.1634976 | 0.05443809 | -3.0033683 | 0.00267009 |
| Xu_2009_SUTs11 | Xu_2009_SU | 90.0167904 | 0.52745343 | 0.17562864 | 3.00323138 | 0.00267129 |
| MRPL3 | YMR024W | 965.499333 | -0.2067635 | 0.06885094 | -3.0030592 | 0.0026728 |
| CYR1 | YJL005W | 2844.55746 | 0.22553183 | 0.07510746 | 3.00278857 | 0.00267518 |
| HOS1 | YPR068C | 231.343759 | -0.3541609 | 0.11794448 | -3.0027761 | 0.00267529 |
| SQS1 | YNL224C | 800.637864 | 0.22932465 | 0.07640663 | 3.00137097 | 0.00268767 |
| REX2 | YLR059C | 704.939045 | -0.2370773 | 0.07901841 | -3.0002788 | 0.00269733 |
| Xu_2009_SUTs53 | Xu_2009_SU | 29.5595483 | 0.74430064 | 0.24813881 | 2.99953331 | 0.00270394 |
| YHL050W-A | YHL050W-A | 19.3813572 | 0.83287365 | 0.27768861 | 2.99930797 | 0.00270594 |
| OPY2 | YPR075C | 1425.09456 | 0.21829238 | 0.07277935 | 2.9993725 | 0.00270536 |
| Xu_2009_CUTs59 | Xu_2009_CU | 19.2860977 | 0.83201783 | 0.27741926 | 2.99913508 | 0.00270747 |
| WSC4 | YHL028W | 473.503705 | -0.349174 | 0.11643985 | -2.9987504 | 0.00271089 |
| YLR125W | YLR125W | 92.7561105 | 0.51100325 | 0.17041129 | 2.99864669 | 0.00271182 |
| HCM1 | YCR065W | 956.612841 | -0.2309819 | 0.07703667 | -2.9983376 | 0.00271457 |
| TYW3 | YGL050W | 577.237094 | 0.28151624 | 0.09389632 | 2.99816048 | 0.00271615 |
| NOP6 | YDL213C | 921.981089 | 0.24899828 | 0.08308449 | 2.99692864 | 0.00272715 |
| AIM19 | YIL087C | 111.219086 | -0.4565508 | 0.15234315 | -2.9968583 | 0.00272777 |
| Xu_2009_CUTs73 | Xu_2009_CU | 32.1762118 | 0.72835772 | 0.24318105 | 2.99512537 | 0.00274332 |
| YDR316W-B | YDR316W-B | 12.3402043 | 0.8734392 | 0.29191239 | 2.99212788 | 0.0027704 |
| MSD1 | YPL104W | 428.334101 | -0.2734166 | 0.09139718 | -2.9915209 | 0.00277592 |
| DDI1 | YER143W | 601.87546 | -0.2585876 | 0.08648152 | -2.9900905 | 0.00278895 |
| GCY1 | YOR120W | 691.276853 | -0.2561164 | 0.08567594 | -2.9893616 | 0.00279561 |
| RAD33 | YML011C | 221.586008 | -0.3562371 | 0.11926581 | -2.9869174 | 0.00281806 |
| LCL2 | YLR104W | 445.263277 | -0.2846843 | 0.09546106 | -2.9822038 | 0.00286181 |
| ISR1 | YPR106W | 222.648037 | -0.3510881 | 0.11774091 | -2.9818701 | 0.00286494 |
| Xu_2009_CUTs99 | Xu_2009_CU | 14.3102742 | 0.86133711 | 0.28904441 | 2.97994727 | 0.00288298 |
| NAT3 | YPR131C | 436.223725 | -0.2806009 | 0.09417013 | -2.9797229 | 0.00288509 |
| YAP7 | YOL028C | 454.721473 | 0.25707827 | 0.08630231 | 2.97881093 | 0.00289369 |
| YDR344C | YDR344C | 102.836115 | -0.5158328 | 0.17322664 | -2.9777916 | 0.00290333 |
| BCH2 | YKR027W | 629.157054 | 0.23984864 | 0.08059066 | 2.97613446 | 0.00291907 |
| POP5 | YAL033W | 590.41534 | 0.23685166 | 0.07962534 | 2.97457657 | 0.00293393 |
| Xu_2009_CUTs21 | Xu_2009_CU | 59.2367303 | 0.56708906 | 0.19075966 | 2.97279337 | 0.00295103 |
| YER175W-A | YER175W-A | 57.4852948 | 0.58125305 | 0.19555073 | 2.97239006 | 0.00295491 |
| ROM2 | YLR371W | 1830.01967 | 0.18109717 | 0.06092571 | 2.97242598 | 0.00295456 |
| YER189W | YER189W | 19.6970146 | 0.81202189 | 0.27326534 | 2.97155096 | 0.002963 |
| PTC7 | YHR076W | 431.238433 | -0.2819445 | 0.09490636 | -2.970765 | 0.00297059 |
| RBA50 | YDR527W | 2164.43095 | 0.19276638 | 0.06490412 | 2.97001781 | 0.00297782 |
| USV1 | YPL230W | 348.277034 | 0.3314183 | 0.11159972 | 2.96970557 | 0.00298085 |
| HSL7 | YBR133C | 1347.3378 | 0.18507921 | 0.06232738 | 2.96946878 | 0.00298315 |
| EMA35 | YCR100C | 73.0587844 | 0.53305145 | 0.17951094 | 2.96946501 | 0.00298319 |
| HAT2 | YEL056W | 550.590263 | -0.2558784 | 0.08617754 | -2.969201 | 0.00298575 |

| | | | | | | |
|---|---|---|---|---|---|---|
| YGR107W | YGR107W | 33.6081184 | 0.7132737 | 0.24022698 | 2.9691657 | 0.0029861 |
| CCT2 | YIL142W | 2653.16796 | -0.1662916 | 0.05601041 | -2.9689404 | 0.00298829 |
| Xu_2009_SUTs52 | Xu_2009_SU | 21.7787657 | -0.7865762 | 0.26497463 | -2.9684964 | 0.00299261 |
| Xu_2009_CUTs21 | Xu_2009_CU | 37.8008604 | 0.68617749 | 0.23116382 | 2.96836017 | 0.00299393 |
| APL5 | YPL195W | 1863.48036 | -0.2030099 | 0.0684154 | -2.9673131 | 0.00300415 |
| IMG2 | YCR071C | 362.997581 | -0.3091044 | 0.10417692 | -2.9671098 | 0.00300614 |
| CAB1 | YDR531W | 368.081222 | -0.2839575 | 0.09570148 | -2.967117 | 0.00300607 |
| YOR366W | YOR366W | 42.6961065 | 0.66524055 | 0.22426503 | 2.96631418 | 0.00301392 |
| Xu_2009_CUTs69 | Xu_2009_CU | 26.6752516 | 0.73937325 | 0.24927376 | 2.96610942 | 0.00301593 |
| MRX15 | YNR040W | 388.798916 | -0.2945002 | 0.09934287 | -2.9644829 | 0.00303192 |
| YCR043C | YCR043C | 548.347683 | -0.2753131 | 0.0928884 | -2.9639123 | 0.00303755 |
| BRE5 | YNR051C | 1944.75365 | 0.21790958 | 0.07352686 | 2.96367315 | 0.00303991 |
| SEO1 | YAL067C | 68.7165401 | 0.55013544 | 0.18565992 | 2.96313527 | 0.00304523 |
| BCK2 | YER167W | 1689.41353 | 0.17634401 | 0.05952209 | 2.962665 | 0.00304988 |
| NOP58 | YOR310C | 2276.6505 | -0.204699 | 0.06909234 | -2.9626878 | 0.00304966 |
| YEL008W | YEL008W | 14.4274383 | 0.84521547 | 0.28533792 | 2.96215618 | 0.00305493 |
| RPI1 | YIL119C | 325.824998 | -0.2894072 | 0.09771987 | -2.9615999 | 0.00306045 |
| YLL047W | YLL047W | 12.6628392 | 0.8683704 | 0.29325474 | 2.96114696 | 0.00306496 |
| PAR32 | YDL173W | 1264.32282 | -0.2023741 | 0.06841737 | -2.9579349 | 0.00309707 |
| IML2 | YJL082W | 1899.06899 | -0.2158893 | 0.07299876 | -2.9574381 | 0.00310207 |
| IML1 | YJR138W | 1171.53556 | 0.20267537 | 0.06854548 | 2.9568016 | 0.00310848 |
| SWS2 | YNL081C | 582.444762 | -0.2508384 | 0.08484609 | -2.9563928 | 0.0031126 |
| GOR1 | YNL274C | 1013.70874 | -0.2113502 | 0.07150687 | -2.9556626 | 0.00311998 |
| YOR289W | YOR289W | 490.422031 | 0.26174339 | 0.08855624 | 2.95567404 | 0.00311987 |
| Xu_2009_SUTs33 | Xu_2009_SU | 7.4382974 | 0.87371137 | 0.29566692 | 2.95505285 | 0.00312615 |
| RET1 | YOR207C | 2099.65481 | 0.18191153 | 0.06156066 | 2.95499636 | 0.00312673 |
| ERB1 | YMR049C | 2436.90346 | 0.21894624 | 0.07413208 | 2.95346127 | 0.00314232 |
| MCM1 | YMR043W | 3177.27266 | 0.19931979 | 0.06750899 | 2.95249268 | 0.0031522 |
| MET12 | YPL023C | 1345.26154 | -0.1997133 | 0.06764661 | -2.9523038 | 0.00315412 |
| MIN9 | YKL023C-A | 497.680495 | -0.2811612 | 0.09524378 | -2.9520166 | 0.00315706 |
| Xu_2009_SUTs70 | Xu_2009_SU | 90.9470818 | 0.48309831 | 0.16365561 | 2.95192034 | 0.00315804 |
| YGR011W | YGR011W | 40.330212 | -0.6667816 | 0.22597937 | -2.9506305 | 0.00317126 |
| GIC2 | YDR309C | 1653.78336 | 0.19892723 | 0.06742459 | 2.95036615 | 0.00317398 |
| HRT3 | YLR097C | 319.647296 | -0.3089228 | 0.10483987 | -2.9466156 | 0.00321272 |
| NAT2 | YGR147C | 1170.63097 | -0.1977942 | 0.0671359 | -2.9461762 | 0.00321729 |
| Xu_2009_CUTs19 | Xu_2009_CU | 31.6807455 | 0.70734951 | 0.24016727 | 2.94523693 | 0.00322707 |
| VTI1 | YMR197C | 577.989534 | 0.24157257 | 0.08205576 | 2.94400489 | 0.00323995 |
| PRM1 | YNL279W | 106.098883 | 0.46097567 | 0.15657814 | 2.94406155 | 0.00323936 |
| Xu_2009_CUTs34 | Xu_2009_CU | 49.3631275 | 0.6499547 | 0.22089041 | 2.94243057 | 0.00325647 |
| Xu_2009_CUTs66 | Xu_2009_CU | 29.0571207 | 0.71742019 | 0.24385293 | 2.94201998 | 0.00326079 |
| YIL168W | YIL168W | 83.8309874 | 0.49407295 | 0.16796983 | 2.94143859 | 0.00326692 |

| | | | | | | |
|---|---|---|---|---|---|---|
| IRS4 | YKR019C | 410.317498 | 0.29916824 | 0.10173278 | 2.94072621 | 0.00327444 |
| Xu_2009_CUTs43 | Xu_2009_CU | 26.3822028 | 0.74255496 | 0.25252034 | 2.94057483 | 0.00327604 |
| SPC105 | YGL093W | 897.918494 | 0.22193363 | 0.07548216 | 2.94021304 | 0.00327987 |
| NUD1 | YOR373W | 557.31826 | 0.23648032 | 0.08044539 | 2.93963789 | 0.00328596 |
| TRF5 | YNL299W | 857.008524 | 0.22115948 | 0.07528851 | 2.93749295 | 0.00330878 |
| RCI37 | YIL077C | 350.428851 | -0.2936783 | 0.09999675 | -2.9368781 | 0.00331534 |
| YAP1 | YML007W | 2285.52955 | 0.16473303 | 0.0560899 | 2.93694609 | 0.00331462 |
| YCR016W | YCR016W | 1009.78001 | 0.21175171 | 0.07218469 | 2.9334713 | 0.00335195 |
| Xu_2009_CUTs74 | Xu_2009_CU | 23.9856879 | 0.77273029 | 0.26344562 | 2.93316816 | 0.00335522 |
| AGE1 | YDR524C | 978.038124 | 0.21286433 | 0.07257959 | 2.93284019 | 0.00335877 |
| HXT1 | YHR094C | 1203.49349 | -0.2256467 | 0.07694927 | -2.9324091 | 0.00336343 |
| OSH2 | YDL019C | 1846.91987 | 0.20300354 | 0.0692428 | 2.93176385 | 0.00337043 |
| Xu_2009_CUTs79 | Xu_2009_CU | 28.1623383 | 0.78386474 | 0.26744271 | 2.93096319 | 0.00337913 |
| THI2 | YBR240C | 243.916705 | -0.3508117 | 0.11971079 | -2.9304932 | 0.00338424 |
| Xu_2009_CUTs68 | Xu_2009_CU | 92.2004044 | 0.48438147 | 0.16549195 | 2.92691853 | 0.00342339 |
| Xu_2009_SUTs83 | Xu_2009_SU | 13.2374895 | 0.84352378 | 0.2882129 | 2.92673848 | 0.00342537 |
| SAP190 | YKR028W | 2122.69582 | 0.18124293 | 0.06194813 | 2.92572092 | 0.00343659 |
| SOL4 | YGR248W | 526.649515 | -0.4978973 | 0.17018725 | -2.9255851 | 0.00343809 |
| TIR2 | YOR010C | 80.0857182 | -0.506494 | 0.17313871 | -2.9253652 | 0.00344052 |
| Xu_2009_CUTs66 | Xu_2009_CU | 40.8721477 | 0.70028289 | 0.23948338 | 2.92413979 | 0.0034541 |
| CCC1 | YLR220W | 481.254617 | -0.2661322 | 0.09106318 | -2.9225001 | 0.00347234 |
| tT(AGU)D | tT(AGU)D | 18.0610823 | -0.8056109 | 0.27567363 | -2.9223357 | 0.00347417 |
| UBC8 | YEL012W | 349.464078 | -0.3241233 | 0.11091385 | -2.9222981 | 0.00347459 |
| tR(ACG)E | tR(ACG)E | 3.4951981 | -0.7752375 | 0.26530315 | -2.9220817 | 0.003477 |
| Xu_2009_SUTs27 | Xu_2009_SU | 110.596453 | 0.45056148 | 0.15420283 | 2.92187563 | 0.0034793 |
| NYV1 | YLR093C | 908.273635 | -0.224472 | 0.07683738 | -2.9213907 | 0.00348473 |
| Xu_2009_CUTs16 | Xu_2009_CU | 31.3805766 | 0.68732754 | 0.23533292 | 2.92066038 | 0.0034929 |
| HTC1 | YPL067C | 329.417419 | -0.326203 | 0.11173536 | -2.9194254 | 0.00350677 |
| SRC1 | YML034W | 1166.87696 | 0.21197416 | 0.07261122 | 2.91930322 | 0.00350815 |
| YLR297W | YLR297W | 250.4445 | 0.357901 | 0.12260231 | 2.9192027 | 0.00350928 |
| SDO1 | YLR022C | 598.962031 | -0.2437223 | 0.08351533 | -2.9182948 | 0.00351951 |
| IES3 | YLR052W | 936.970099 | -0.2060825 | 0.07062236 | -2.9180919 | 0.00352181 |
| NUF2 | YOL069W | 500.371024 | 0.26425462 | 0.09059232 | 2.91696504 | 0.00353455 |
| YKR018C | YKR018C | 2049.31493 | -0.1680385 | 0.05763521 | -2.9155529 | 0.00355059 |
| Xu_2009_CUTs22 | Xu_2009_CU | 29.2162076 | 0.70432676 | 0.24165004 | 2.91465608 | 0.00356081 |
| CDC25 | YLR310C | 2494.65769 | 0.16890165 | 0.0579477 | 2.91472568 | 0.00356001 |
| DUG2 | YBR281C | 726.591765 | 0.2211392 | 0.07587628 | 2.91447063 | 0.00356292 |
| Xu_2009_CUTs58 | Xu_2009_CU | 41.4699295 | 0.64578562 | 0.22165279 | 2.91350104 | 0.00357401 |
| GLO2 | YDR272W | 767.924782 | -0.2390199 | 0.08205049 | -2.9130827 | 0.0035788 |
| GEP7 | YGL057C | 180.952145 | -0.3785377 | 0.12994086 | -2.913154 | 0.00357798 |
| TIM54 | YJL054W | 1140.39325 | -0.2259042 | 0.07754978 | -2.9130217 | 0.0035795 |

| | | | | | | |
|---|---|---|---|---|---|---|
| tV(AAC)E2 | tV(AAC)E2 | 20.7691116 | -0.791104 | 0.27170077 | -2.9116737 | 0.00359498 |
| ADE16 | YLR028C | 2906.64073 | -0.1936358 | 0.06654999 | -2.9096289 | 0.00361858 |
| Xu_2009_CUTs18 | Xu_2009_CU | 7.2994151 | 0.85683255 | 0.29451305 | 2.90931948 | 0.00362216 |
| HPA3 | YEL066W | 193.520942 | -0.3802517 | 0.13080369 | -2.9070411 | 0.00364865 |
| Xu_2009_CUTs22 | Xu_2009_CU | 31.7549033 | 0.74318635 | 0.25573869 | 2.906038 | 0.00366037 |
| YJR149W | YJR149W | 298.567015 | -0.2939518 | 0.10114911 | -2.9061239 | 0.00365937 |
| REC8 | YPR007C | 187.823427 | 0.38040338 | 0.13091861 | 2.90564795 | 0.00366494 |
| Xu_2009_CUTs77 | Xu_2009_CU | 8.9381543 | 0.85904056 | 0.29568945 | 2.90521208 | 0.00367004 |
| ATG5 | YPL149W | 532.012153 | -0.2529288 | 0.08714399 | -2.9024242 | 0.00370287 |
| YBL029C-A | YBL029C-A | 457.934024 | -0.2571965 | 0.08861734 | -2.9023271 | 0.00370402 |
| PRO1 | YDR300C | 784.565976 | -0.2254783 | 0.07769933 | -2.9019335 | 0.00370867 |
| NOP53 | YPL146C | 1356.4974 | 0.21369565 | 0.07367697 | 2.90044034 | 0.00372639 |
| PAM16 | YJL104W | 735.109418 | -0.2669089 | 0.09203037 | -2.9002265 | 0.00372893 |
| APJ1 | YNL077W | 292.881809 | -0.3075202 | 0.10605432 | -2.8996482 | 0.00373582 |
| Xu_2009_CUTs53 | Xu_2009_CU | 54.2226361 | 0.60502682 | 0.20873757 | 2.89850465 | 0.00374947 |
| EST2 | YLR318W | 258.677196 | 0.33506448 | 0.11562395 | 2.89788122 | 0.00375693 |
| COS5 | YJR161C | 153.869954 | 0.4072576 | 0.14054986 | 2.89760231 | 0.00376027 |
| BUB2 | YMR055C | 281.649682 | 0.30555798 | 0.10551465 | 2.89588198 | 0.00378095 |
| SNR36 | snR36 | 189.467112 | -0.3889179 | 0.13435355 | -2.8947353 | 0.00379478 |
| PEP8 | YJL053W | 324.760281 | -0.3306352 | 0.11430704 | -2.8925179 | 0.00382167 |
| RAD4 | YER162C | 464.164382 | 0.26987883 | 0.09332607 | 2.89178407 | 0.00383061 |
| YML012C-A | YML012C-A | 25.616161 | 0.74425334 | 0.257369 | 2.89177536 | 0.00383072 |
| tE(UUC)K | tE(UUC)K | 15.9946007 | -0.8404035 | 0.29064997 | -2.8914626 | 0.00383453 |
| ZEO1 | YOL109W | 1447.93241 | -0.264524 | 0.0914886 | -2.8913327 | 0.00383612 |
| Xu_2009_CUTs58 | Xu_2009_CU | 13.5999217 | 0.82533598 | 0.28559779 | 2.88985426 | 0.0038542 |
| Xu_2009_CUTs90 | Xu_2009_CU | 13.6409785 | 0.82418659 | 0.28530683 | 2.88877274 | 0.00386748 |
| Xu_2009_CUTs50 | Xu_2009_CU | 30.9197251 | 0.70817947 | 0.24533169 | 2.88662046 | 0.00389404 |
| IOC3 | YFR013W | 981.304103 | 0.20516553 | 0.07107619 | 2.88655786 | 0.00389481 |
| PIG1 | YLR273C | 396.179608 | 0.28312822 | 0.0981341 | 2.8851154 | 0.0039127 |
| MCX1 | YBR227C | 618.368844 | -0.2340759 | 0.08114527 | -2.8846529 | 0.00391845 |
| YDR008C | YDR008C | 36.3440097 | 0.66018418 | 0.22888329 | 2.88437035 | 0.00392197 |
| CEF1 | YMR213W | 655.164414 | 0.25886964 | 0.08975055 | 2.88432361 | 0.00392255 |
| YDL025W-A | YDL025W-A | 11.0770294 | 0.84837562 | 0.29415145 | 2.88414569 | 0.00392477 |
| RGP1 | YDR137W | 858.312237 | 0.21714085 | 0.07529065 | 2.88403462 | 0.00392616 |
| HPR1 | YDR138W | 1196.79939 | 0.19460923 | 0.06748059 | 2.8839291 | 0.00392747 |
| TPO2 | YGR138C | 2414.21946 | 0.17599153 | 0.06103401 | 2.88349934 | 0.00393283 |
| Xu_2009_SUTs40 | Xu_2009_SU | 54.2695109 | 0.66502754 | 0.2307377 | 2.88217978 | 0.00394934 |
| RIX7 | YLL034C | 993.769637 | 0.23042667 | 0.07995026 | 2.88212525 | 0.00395003 |
| tL(CAA)C/SUP53 | tL(CAA)C | 299.216342 | 0.29784068 | 0.10340109 | 2.88044059 | 0.0039712 |
| STH1 | YIL126W | 2514.35415 | 0.15821692 | 0.05493599 | 2.88002316 | 0.00397646 |
| CUL3 | YGR003W | 769.323815 | 0.21850762 | 0.07590832 | 2.87857277 | 0.00399479 |

| | | | | | | |
|---|---|---|---|---|---|---|
| UTP15 | YMR093W | 1680.13042 | -0.1973334 | 0.06862785 | -2.8754127 | 0.004035 |
| ORC4 | YPR162C | 480.908226 | 0.24762844 | 0.08613121 | 2.87501391 | 0.0040401 |
| PCL2 | YDL127W | 540.654834 | -0.2643127 | 0.09200029 | -2.872955 | 0.00406652 |
| ARO3 | YDR035W | 1653.25432 | -0.1645844 | 0.05730962 | -2.8718468 | 0.00408081 |
| TAM41 | YGR046W | 1196.04813 | 0.185193 | 0.06448524 | 2.8718666 | 0.00408055 |
| URA8 | YJR103W | 1700.84629 | 0.17456521 | 0.06080726 | 2.87079544 | 0.0040944 |
| Xu_2009_CUTs9 | Xu_2009_CU | 10.3868134 | 0.84576071 | 0.29467084 | 2.87018801 | 0.00410228 |
| Xu_2009_SUTs22 | Xu_2009_SU | 24.4853161 | 0.76924187 | 0.26807647 | 2.86948668 | 0.00411139 |
| SLM4 | YBR077C | 419.870697 | 0.26393981 | 0.09204661 | 2.86745824 | 0.00413783 |
| CDC24 | YAL041W | 1597.88768 | 0.17287134 | 0.06031767 | 2.86601482 | 0.00415675 |
| Xu_2009_SUTs42 | Xu_2009_SU | 43.1798615 | 0.643122 | 0.22447359 | 2.86502296 | 0.00416979 |
| RPC82 | YPR190C | 1326.14483 | 0.21064648 | 0.07359746 | 2.86214334 | 0.00420787 |
| HEM13 | YDR044W | 832.351515 | -0.2266613 | 0.07921751 | -2.8612524 | 0.00421971 |
| NGL1 | YOL042W | 307.030044 | 0.28773189 | 0.10060854 | 2.85991518 | 0.00423754 |
| Xu_2009_CUTs20 | Xu_2009_CU | 13.5186893 | 0.83138513 | 0.29081015 | 2.85885877 | 0.00425168 |
| MED11 | YMR112C | 266.133834 | -0.312795 | 0.10943378 | -2.8583038 | 0.00425912 |
| BUD17 | YNR027W | 515.989257 | 0.24301877 | 0.08502534 | 2.85819229 | 0.00426062 |
| RAD28 | YDR030C | 231.730196 | 0.34365999 | 0.12033207 | 2.85593019 | 0.00429109 |
| MRX16 | YDR132C | 263.861894 | 0.31203186 | 0.1092592 | 2.85588627 | 0.00429169 |
| BPL1 | YDL141W | 649.527406 | -0.2262284 | 0.0792578 | -2.854336 | 0.00431269 |
| CUS1 | YMR240C | 259.974894 | 0.34579336 | 0.12119227 | 2.85326243 | 0.00432729 |
| Xu_2009_SUTs67 | Xu_2009_SU | 79.4627927 | 0.50317281 | 0.17647165 | 2.85129543 | 0.00435415 |
| CDD1 | YLR245C | 75.8678677 | -0.5294531 | 0.18579045 | -2.8497327 | 0.0043756 |
| YMR254C | YMR254C | 16.4920007 | 0.80579184 | 0.28288783 | 2.84844997 | 0.00439328 |
| CBS2 | YDR197W | 390.935812 | -0.2692636 | 0.09457901 | -2.8469696 | 0.00441376 |
| HXT17 | YNR072W | 12.20374 | 0.82766588 | 0.2907642 | 2.84651918 | 0.00442001 |
| JJJ2 | YJL162C | 432.411839 | 0.26670423 | 0.0937 | 2.84636312 | 0.00442217 |
| EMC10 | YDR056C | 534.679498 | -0.2384375 | 0.08377987 | -2.8459998 | 0.00442722 |
| Xu_2009_SUTs29 | Xu_2009_SU | 127.855144 | 0.47713675 | 0.167683 | 2.84546879 | 0.00443461 |
| Xu_2009_CUTs75 | Xu_2009_CU | 73.9234947 | 0.50192673 | 0.17678951 | 2.83912056 | 0.00452381 |
| SFT1 | YKL006C-A | 305.524864 | 0.3112927 | 0.10966393 | 2.83860621 | 0.0045311 |
| YOL097W-A | YOL097W-A | 38.5308861 | 0.6353094 | 0.22383592 | 2.8382817 | 0.00453571 |
| YOL085W-A | YOL085W-A | 76.3180106 | 0.5254439 | 0.18517093 | 2.8376155 | 0.00454519 |
| HUA1 | YGR268C | 675.686069 | -0.2508004 | 0.08848681 | -2.8343254 | 0.00459226 |
| Xu_2009_CUTs55 | Xu_2009_CU | 39.4965225 | 0.63862076 | 0.22534182 | 2.83400909 | 0.0045968 |
| LCL3 | YGL085W | 274.457003 | -0.3034452 | 0.10712633 | -2.8325927 | 0.00461722 |
| ECT1 | YGR007W | 729.514646 | -0.2295774 | 0.08105026 | -2.8325315 | 0.0046181 |
| NHP10 | YDL002C | 427.68373 | -0.2654008 | 0.09372529 | -2.8316877 | 0.0046303 |
| Xu_2009_CUTs17 | Xu_2009_CU | 16.1781668 | 0.79187532 | 0.27966571 | 2.83150671 | 0.00463293 |
| Xu_2009_SUTs61 | Xu_2009_SU | 21.5006009 | -0.7592048 | 0.26826014 | -2.8301066 | 0.00465325 |
| Xu_2009_SUTs43 | Xu_2009_SU | 64.3105451 | -0.5607423 | 0.19829717 | -2.8277876 | 0.00468709 |

| | | | | | | |
|---|---|---|---|---|---|---|
| Xu_2009_CUTs23 | Xu_2009_CU | 36.4390382 | 0.66376611 | 0.23474303 | 2.82762864 | 0.00468942 |
| BST1 | YFL025C | 949.530529 | 0.19379466 | 0.06854044 | 2.82745007 | 0.00469203 |
| SPH1 | YLR313C | 184.297881 | 0.3526629 | 0.12487008 | 2.82423868 | 0.00473931 |
| Xu_2009_CUTs34 | Xu_2009_CU | 45.7850544 | 0.60082457 | 0.21276239 | 2.82392281 | 0.00474398 |
| CDH1 | YGL003C | 783.571918 | -0.2060148 | 0.07296721 | -2.8233887 | 0.00475189 |
| Xu_2009_CUTs39 | Xu_2009_CU | 35.3478517 | 0.6664939 | 0.2361576 | 2.82224197 | 0.00476892 |
| RFC1 | YOR217W | 1116.44281 | 0.19643113 | 0.06960241 | 2.82218861 | 0.00476971 |
| Xu_2009_CUTs77 | Xu_2009_CU | 130.422933 | 0.42213717 | 0.1496183 | 2.82142739 | 0.00478105 |
| CCP1 | YKR066C | 1797.3318 | -0.1828766 | 0.06484528 | -2.8201983 | 0.0047994 |
| Xu_2009_CUTs14 | Xu_2009_CU | 30.7148126 | 0.6875882 | 0.24386123 | 2.81958797 | 0.00480853 |
| SNA4 | YDL123W | 443.285958 | -0.2538509 | 0.09003227 | -2.8195547 | 0.00480903 |
| MSH4 | YFL003C | 98.5308436 | 0.48415831 | 0.1717989 | 2.81816895 | 0.00482984 |
| Xu_2009_CUTs15 | Xu_2009_CU | 30.7024895 | 0.72083606 | 0.25583837 | 2.81754482 | 0.00483924 |
| Xu_2009_CUTs64 | Xu_2009_CU | 87.5160277 | 0.49875792 | 0.1770552 | 2.81696287 | 0.00484801 |
| Xu_2009_SUTs38 | Xu_2009_SU | 153.610798 | 0.38163021 | 0.13549098 | 2.81664656 | 0.00485279 |
| Xu_2009_CUTs17 | Xu_2009_CU | 4.10876129 | 0.78300197 | 0.27813118 | 2.81522543 | 0.0048743 |
| YJL045W | YJL045W | 121.705274 | 0.4191807 | 0.14894873 | 2.81426165 | 0.00488894 |
| APL2 | YKL135C | 1080.37759 | -0.1947818 | 0.06922333 | -2.8138167 | 0.00489571 |
| EDC3 | YEL015W | 1324.0475 | 0.18104357 | 0.06435954 | 2.81300283 | 0.00490812 |
| Xu_2009_CUTs61 | Xu_2009_CU | 47.3400917 | 0.60062507 | 0.21369501 | 2.81066491 | 0.00494392 |
| SPC98 | YNL126W | 444.478958 | -0.2617631 | 0.09313648 | -2.8105324 | 0.00494596 |
| YBR225W | YBR225W | 953.67597 | 0.20896703 | 0.07436407 | 2.81005383 | 0.00495332 |
| MLO127 | YJR039W | 378.000663 | 0.29331121 | 0.104377 | 2.81011331 | 0.00495241 |
| ICT1 | YLR099C | 1534.07004 | -0.2220419 | 0.07902266 | -2.8098513 | 0.00495644 |
| NCE102 | YPR149W | 3129.84348 | -0.2506543 | 0.08926828 | -2.8078764 | 0.00498694 |
| YGR017W | YGR017W | 1079.97807 | -0.217736 | 0.07755999 | -2.8073241 | 0.0049955 |
| AFT1 | YGL071W | 1556.34332 | -0.1816922 | 0.06472406 | -2.8071819 | 0.0049977 |
| DIT2 | YDR402C | 53.3077632 | 0.63185479 | 0.22512432 | 2.80669274 | 0.0050053 |
| tH(GUG)G2 | tH(GUG)G2 | 181.726187 | -0.3743729 | 0.13360241 | -2.8021416 | 0.00507646 |
| RSF1 | YMR030W | 283.020597 | -0.296457 | 0.10583879 | -2.8010237 | 0.00509408 |
| ARP6 | YLR085C | 623.757423 | -0.2398633 | 0.08566485 | -2.8000199 | 0.00510995 |
| YGR182C | YGR182C | 94.2300446 | -0.4702342 | 0.16837069 | -2.7928506 | 0.00522458 |
| RCK2 | YLR248W | 2176.19177 | -0.1597487 | 0.05721669 | -2.7919954 | 0.00523841 |
| LDH1 | YBR204C | 230.424516 | -0.3497015 | 0.12526689 | -2.7916515 | 0.00524398 |
| EXO1 | YOR033C | 821.32045 | 0.20806873 | 0.07453499 | 2.79155762 | 0.0052455 |
| Xu_2009_SUTs33 | Xu_2009_SU | 369.285038 | 0.28694993 | 0.10281258 | 2.79100018 | 0.00525454 |
| DPB11 | YJL090C | 752.340361 | 0.20956659 | 0.07508417 | 2.79108899 | 0.0052531 |
| PKH2 | YOL100W | 932.538099 | 0.19614143 | 0.07027235 | 2.7911607 | 0.00525194 |
| CIN10 | YDR387C | 549.198385 | -0.2476293 | 0.0887609 | -2.7898467 | 0.0052733 |
| Xu_2009_SUTs91 | Xu_2009_SU | 26.5863149 | 0.70558022 | 0.2529643 | 2.78924825 | 0.00528306 |
| CTR86 | YCR054C | 625.020131 | 0.21736144 | 0.07793543 | 2.78899376 | 0.00528721 |

| | | | | | | |
|---|---|---|---|---|---|---|
| CSM4 | YPL200W | 15.8986894 | 0.78587259 | 0.28185482 | 2.78821771 | 0.00529989 |
| EAF3 | YPR023C | 927.345867 | -0.1987288 | 0.07128095 | -2.7879652 | 0.00530402 |
| YFR018C | YFR018C | 197.223756 | -0.3736526 | 0.13408035 | -2.7867812 | 0.00532344 |
| ARG80 | YMR042W | 322.704068 | 0.28918569 | 0.10379818 | 2.78603804 | 0.00533566 |
| MAM3 | YOL060C | 1503.51723 | 0.18203546 | 0.0653414 | 2.78591306 | 0.00533772 |
| INP52 | YNL106C | 777.490628 | 0.23298176 | 0.08364837 | 2.78525153 | 0.00534862 |
| LIP5 | YOR196C | 977.02583 | 0.2156874 | 0.07744408 | 2.78507275 | 0.00535157 |
| YOS9 | YDR057W | 784.650354 | 0.20352509 | 0.07309895 | 2.78424104 | 0.00536531 |
| Xu_2009_SUTs84 | Xu_2009_SU | 14.4620174 | 0.79550926 | 0.28575953 | 2.78384158 | 0.00537193 |
| MSS18 | YPR134W | 309.209054 | -0.2881868 | 0.10354044 | -2.7833261 | 0.00538047 |
| DSE3 | YOR264W | 830.864396 | -0.2138161 | 0.07683476 | -2.7828043 | 0.00538913 |
| YBR220C | YBR220C | 331.473077 | -0.3117797 | 0.11209223 | -2.7814573 | 0.00541155 |
| YBR076C-A | YBR076C-A | 168.374239 | 0.36015617 | 0.1294948 | 2.78124032 | 0.00541516 |
| GEA2 | YEL022W | 2051.97903 | 0.16655748 | 0.05989638 | 2.78076026 | 0.00542318 |
| Xu_2009_CUTs19 | Xu_2009_CU | 23.8341777 | 0.72530534 | 0.26093185 | 2.77967345 | 0.00544136 |
| YDR514C | YDR514C | 671.719165 | 0.25380308 | 0.0913602 | 2.77804853 | 0.00546864 |
| YBT1 | YLL048C | 3202.48097 | 0.15352561 | 0.05526753 | 2.77786262 | 0.00547177 |
| Xu_2009_SUTs84 | Xu_2009_SU | 51.9314142 | 0.56393045 | 0.20304889 | 2.77731362 | 0.00548103 |
| ALO1 | YML086C | 1019.77352 | -0.2074942 | 0.07477508 | -2.7749108 | 0.00552169 |
| YDR476C | YDR476C | 410.568003 | -0.2748561 | 0.09909007 | -2.7738006 | 0.00554056 |
| YBR056C-B | YBR056C-B | 25.7380798 | 0.69786932 | 0.25172065 | 2.77239604 | 0.00556453 |
| SNR32 | snR32 | 3504.13665 | -0.3787266 | 0.13666905 | -2.7711221 | 0.00558635 |
| SEC18 | YBR080C | 1650.91088 | -0.1709152 | 0.06170544 | -2.7698559 | 0.00560811 |
| MDM34 | YGL219C | 942.840079 | -0.1900567 | 0.06866436 | -2.7679085 | 0.00564173 |
| SOM1 | YEL059C-A | 328.375579 | -0.273592 | 0.09888274 | -2.7668329 | 0.00566038 |
| YKU80 | YMR106C | 314.809534 | 0.33650884 | 0.12163987 | 2.76643537 | 0.00566728 |
| TFB3 | YDR460W | 536.063659 | 0.24927591 | 0.09013193 | 2.76567829 | 0.00568045 |
| CBP3 | YPL215W | 709.148658 | -0.2217067 | 0.08017112 | -2.7654191 | 0.00568497 |
| Xu_2009_CUTs76 | Xu_2009_CU | 20.6939195 | 0.76139479 | 0.2754068 | 2.76461871 | 0.00569894 |
| Xu_2009_SUTs60 | Xu_2009_SU | 574.143177 | -0.2640778 | 0.09557296 | -2.7631015 | 0.0057255 |
| Xu_2009_CUTs65 | Xu_2009_CU | 30.3897273 | 0.67168389 | 0.24311044 | 2.76287554 | 0.00572946 |
| Xu_2009_SUTs47 | Xu_2009_SU | 14.7020117 | 0.78345365 | 0.28382038 | 2.76038545 | 0.00577332 |
| ATP7 | YKL016C | 2063.27013 | -0.1605302 | 0.05815412 | -2.760427 | 0.00577259 |
| COQ2 | YNR041C | 342.395326 | -0.2909824 | 0.10540776 | -2.7605408 | 0.00577058 |
| Xu_2009_CUTs63 | Xu_2009_CU | 17.7393629 | 0.77935123 | 0.28236751 | 2.76005992 | 0.00577908 |
| RAD52 | YML032C | 1665.811 | 0.17167605 | 0.06219875 | 2.76012076 | 0.005778 |
| YOR097C | YOR097C | 319.500521 | -0.2817873 | 0.10212273 | -2.7593008 | 0.00579252 |
| RPN1 | YHR027C | 5361.11069 | 0.1489655 | 0.05399464 | 2.75889393 | 0.00579974 |
| ESA1 | YOR244W | 936.294006 | -0.2113481 | 0.0766174 | -2.7584873 | 0.00580696 |
| YBR090C | YBR090C | 170.761112 | 0.3618952 | 0.13122408 | 2.75784135 | 0.00581844 |
| PEX34 | YCL056C | 168.259327 | 0.37642505 | 0.1365869 | 2.75593812 | 0.00585241 |

| | | | | | | |
|---|---|---|---|---|---|---|
| NGL2 | YMR285C | 610.249107 | -0.2354916 | 0.08546991 | -2.7552578 | 0.00586459 |
| NMD4 | YLR363C | 268.139312 | -0.3268682 | 0.11869208 | -2.7539178 | 0.00588866 |
| SLY1 | YDR189W | 1748.53592 | -0.1606118 | 0.05836044 | -2.752067 | 0.00592204 |
| CLB5 | YPR120C | 407.655595 | -0.2500567 | 0.09086362 | -2.7519997 | 0.00592326 |
| Xu_2009_SUTs90 | Xu_2009_SU | 45.3128712 | 0.65284396 | 0.23734855 | 2.75057065 | 0.00594916 |
| Xu_2009_SUTs37 | Xu_2009_SU | 16.6934191 | 0.76588256 | 0.27854755 | 2.74955769 | 0.00596758 |
| HEM12 | YDR047W | 550.434728 | -0.2303519 | 0.08379288 | -2.7490634 | 0.00597658 |
| LSM7 | YNL147W | 349.783022 | -0.2678001 | 0.09746792 | -2.7475719 | 0.00600383 |
| SIR1 | YKR101W | 200.371274 | 0.40370261 | 0.14702546 | 2.7458007 | 0.00603634 |
| tH(GUG)E1 | tH(GUG)E1 | 22.8320456 | -0.7327957 | 0.26691708 | -2.7454058 | 0.00604361 |
| YIR017W-A | YIR017W-A | 10.5198838 | 0.80342952 | 0.29269859 | 2.74490397 | 0.00605286 |
| Xu_2009_CUTs88 | Xu_2009_CU | 54.3163553 | 0.54498611 | 0.19859142 | 2.744258 | 0.00606479 |
| Xu_2009_CUTs91 | Xu_2009_CU | 36.5236811 | 0.6417476 | 0.23402848 | 2.7421773 | 0.00610334 |
| TDA4 | YJR116W | 376.689524 | 0.26425484 | 0.0964038 | 2.7411248 | 0.00612292 |
| Xu_2009_SUTs15 | Xu_2009_SU | 40.2319573 | 0.63139864 | 0.23049345 | 2.73933446 | 0.00615637 |
| Xu_2009_CUTs85 | Xu_2009_CU | 16.3640947 | 0.7735752 | 0.28241382 | 2.7391549 | 0.00615973 |
| Xu_2009_SUTs74 | Xu_2009_SU | 237.312864 | 0.31644886 | 0.11558972 | 2.73769025 | 0.00618723 |
| NEO1 | YIL048W | 1129.81788 | 0.19084609 | 0.06971449 | 2.7375382 | 0.00619009 |
| tL(CAA)D | tL(CAA)D | 20.9874244 | -0.7636985 | 0.27902592 | -2.7370164 | 0.00619992 |
| Xu_2009_SUTs71 | Xu_2009_SU | 12.5985374 | 0.79470806 | 0.29040026 | 2.73659553 | 0.00620786 |
| Xu_2009_CUTs15 | Xu_2009_CU | 59.7821526 | 0.54507121 | 0.19920723 | 2.73620186 | 0.00621529 |
| Xu_2009_CUTs36 | Xu_2009_CU | 21.7837105 | 0.71049465 | 0.25971861 | 2.7356324 | 0.00622605 |
| PEX19 | YDL065C | 440.699263 | 0.26222318 | 0.09588341 | 2.73481271 | 0.00624158 |
| DCG1 | YIR030C | 94.5267118 | -0.45721 | 0.16726422 | -2.7334598 | 0.00626728 |
| DAL1 | YIR027C | 102.897653 | -0.4469356 | 0.16354501 | -2.732799 | 0.00627986 |
| SNR55 | snR55 | 50.9994891 | -0.598798 | 0.21916517 | -2.7321768 | 0.00629174 |
| FLC1 | YPL221W | 3781.02311 | 0.17904903 | 0.06554784 | 2.73157775 | 0.00630319 |
| Xu_2009_CUTs33 | Xu_2009_CU | 14.1432826 | 0.77892993 | 0.2853146 | 2.73007381 | 0.00633201 |
| NUP49 | YGL172W | 566.559127 | -0.2280273 | 0.08352828 | -2.7299413 | 0.00633456 |
| FOL2 | YGR267C | 1182.84247 | -0.1887867 | 0.06918437 | -2.7287481 | 0.00635752 |
| BOI2 | YER114C | 1784.40727 | 0.15510577 | 0.0568487 | 2.72839623 | 0.00636431 |
| RNY1 | YPL123C | 273.191294 | -0.5361951 | 0.19667217 | -2.7263395 | 0.00640411 |
| PEX10 | YDR265W | 521.298811 | -0.2438624 | 0.08952092 | -2.7240823 | 0.00644804 |
| Xu_2009_SUTs21 | Xu_2009_SU | 102.516635 | 0.43591227 | 0.16003375 | 2.72387706 | 0.00645205 |
| ADH7 | YCR105W | 17.3545379 | -0.761361 | 0.27977667 | -2.7213168 | 0.00650224 |
| UNG1 | YML021C | 563.837774 | -0.2258978 | 0.08301532 | -2.7211578 | 0.00650537 |
| ATG34 | YOL083W | 226.868711 | -0.341019 | 0.12534691 | -2.7206014 | 0.00651633 |
| CCH1 | YGR217W | 1460.02823 | 0.17203823 | 0.06324236 | 2.72030046 | 0.00652226 |
| Xu_2009_CUTs57 | Xu_2009_CU | 27.8203951 | 0.67139346 | 0.2468858 | 2.71944953 | 0.00653907 |
| Xu_2009_CUTs16 | Xu_2009_CU | 19.9154033 | 0.74173185 | 0.27276272 | 2.71933001 | 0.00654143 |
| FAA4 | YMR246W | 7040.01977 | 0.21018202 | 0.07731456 | 2.71853073 | 0.00655726 |

| | | | | | | |
|---|---|---|---|---|---|---|
| CRG1 | YHR209W | 408.984107 | 0.27958829 | 0.10287225 | 2.71782038 | 0.00657135 |
| YBR221W-A | YBR221W-A | 2.43170257 | -0.6706035 | 0.24696273 | -2.7154037 | 0.0066195 |
| tP(AGG)C/SUF2 | tP(AGG)C | 68.1958493 | 0.50486655 | 0.18602218 | 2.71401264 | 0.00664736 |
| QCR6 | YFR033C | 3474.41773 | 0.17458535 | 0.06433079 | 2.71386922 | 0.00665024 |
| ALY1 | YKR021W | 861.183821 | 0.19881516 | 0.0732896 | 2.71273372 | 0.00667307 |
| PDE1 | YGL248W | 1488.45915 | 0.21182929 | 0.07811581 | 2.71173396 | 0.00669323 |
| CTS2 | YDR371W | 374.537883 | 0.25174726 | 0.09287855 | 2.71049952 | 0.00671819 |
| THI3 | YDL080C | 709.961124 | 0.22371328 | 0.08256994 | 2.70937915 | 0.00674093 |
| ATG11 | YPR049C | 936.254086 | 0.1929078 | 0.07120109 | 2.7093378 | 0.00674177 |
| tV(AAC)M2 | tV(AAC)M2 | 114.431862 | -0.4272116 | 0.15769894 | -2.7090328 | 0.00674797 |
| MET8 | YBR213W | 88.3232623 | -0.471842 | 0.17417326 | -2.7090383 | 0.00674785 |
| YDL218W | YDL218W | 38.47071 | -0.6294474 | 0.23235128 | -2.7090333 | 0.00674796 |
| BTT1 | YDR252W | 65.8084097 | 0.50828784 | 0.18764136 | 2.70882626 | 0.00675217 |
| TUL1 | YKL034W | 596.342911 | -0.239516 | 0.08842308 | -2.7087499 | 0.00675372 |
| MLH2 | YLR035C | 351.357355 | 0.27690759 | 0.10226075 | 2.70785803 | 0.0067719 |
| YEL075C | YEL075C | 9.41886161 | 0.80149091 | 0.29614475 | 2.70641612 | 0.00680138 |
| MGR1 | YCL044C | 665.077254 | -0.2408309 | 0.08899568 | -2.7060959 | 0.00680794 |
| Xu_2009_CUTs83 | Xu_2009_CU | 15.5013517 | 0.76893138 | 0.28416571 | 2.70592597 | 0.00681142 |
| TAH18 | YPR048W | 610.289699 | 0.23388645 | 0.08651683 | 2.70336345 | 0.00686416 |
| BMH2 | YDR099W | 5438.04964 | -0.1613406 | 0.05968677 | -2.7031213 | 0.00686917 |
| Xu_2009_SUTs41 | Xu_2009_SU | 109.848929 | 0.41951717 | 0.15521569 | 2.70280137 | 0.00687578 |
| Xu_2009_CUTs74 | Xu_2009_CU | 19.5594645 | 0.73740581 | 0.27315563 | 2.69958126 | 0.00694268 |
| tS(AGA)J | tS(AGA)J | 7.5894611 | -0.799134 | 0.29613587 | -2.6985384 | 0.00696447 |
| SNR78 | snR78 | 10.3509672 | -0.7996414 | 0.29638564 | -2.6979763 | 0.00697624 |
| YPK9 | YOR291W | 1242.2723 | 0.18217418 | 0.06753836 | 2.69734389 | 0.0069895 |
| SNR56 | snR56 | 21.3161213 | 0.74551454 | 0.27648637 | 2.69638802 | 0.0070096 |
| YPL278C | YPL278C | 107.33151 | -0.4228977 | 0.15683445 | -2.6964591 | 0.0070081 |
| VPS63 | YLR261C | 4.98148514 | -0.7630541 | 0.28323327 | -2.6940835 | 0.00705825 |
| Xu_2009_SUTs68 | Xu_2009_SU | 36.3397939 | 0.6462871 | 0.23991638 | 2.69380143 | 0.00706422 |
| TRM9 | YML014W | 509.140054 | -0.2436585 | 0.09049583 | -2.6924832 | 0.00709221 |
| DPC13 | YGL041W-A | 269.705639 | -0.294551 | 0.10944355 | -2.6913512 | 0.00711632 |
| Xu_2009_CUTs82 | Xu_2009_CU | 25.2526313 | 0.68020584 | 0.25277503 | 2.69095351 | 0.00712481 |
| SET3 | YKR029C | 721.819647 | 0.208069 | 0.0773235 | 2.69088951 | 0.00712618 |
| Xu_2009_SUTs21 | Xu_2009_SU | 92.9135437 | 0.44053221 | 0.16372484 | 2.6906864 | 0.00713052 |
| Xu_2009_CUTs36 | Xu_2009_CU | 42.5894986 | 0.63067634 | 0.23445146 | 2.69000811 | 0.00714503 |
| Xu_2009_SUTs24 | Xu_2009_SU | 86.6553483 | 0.48473378 | 0.1801937 | 2.69007067 | 0.00714369 |
| TGL4 | YKR089C | 1621.58238 | 0.18263076 | 0.06789709 | 2.68981699 | 0.00714912 |
| PRP40 | YKL012W | 1111.69324 | 0.18509844 | 0.06887491 | 2.68745823 | 0.00719981 |
| LDB7 | YBL006C | 864.865503 | -0.2324761 | 0.0865141 | -2.6871464 | 0.00720654 |
| PAN3 | YKL025C | 838.839726 | -0.1981745 | 0.07379716 | -2.6853945 | 0.00724442 |
| SLY41 | YOR307C | 603.451941 | 0.22249588 | 0.08286957 | 2.6848924 | 0.00725532 |

| | | | | | | |
|---|---|---|---|---|---|---|
| YNL092W | YNL092W | 102.319576 | -0.4165498 | 0.15520178 | -2.6839239 | 0.00727637 |
| PRP46 | YPL151C | 513.365707 | 0.24941511 | 0.09297834 | 2.6825077 | 0.00730725 |
| VPS38 | YLR360W | 429.539431 | 0.25417594 | 0.0947571 | 2.68239473 | 0.00730972 |
| POS5 | YPL188W | 613.731254 | -0.2207574 | 0.0823069 | -2.682125 | 0.00731561 |
| YOL038C-A | YOL038C-A | 187.324589 | -0.3414952 | 0.12736166 | -2.6813026 | 0.00733362 |
| YOR053W | YOR053W | 18.6108918 | 0.74631671 | 0.27835869 | 2.68113317 | 0.00733733 |
| YGR050C | YGR050C | 76.5216169 | -0.4902825 | 0.18287307 | -2.6809991 | 0.00734027 |
| YNL174W | YNL174W | 8.16969622 | 0.79356267 | 0.29600536 | 2.68090646 | 0.0073423 |
| YIM1 | YMR152W | 1102.67945 | -0.1954489 | 0.07292177 | -2.6802543 | 0.00735662 |
| RLP7 | YNL002C | 2116.09737 | -0.1964681 | 0.07332335 | -2.6794749 | 0.00737377 |
| YDR222W | YDR222W | 428.796629 | 0.24518476 | 0.0915352 | 2.6785845 | 0.00739341 |
| MPP10 | YJR002W | 1553.17047 | 0.18728326 | 0.06992802 | 2.67822914 | 0.00740126 |
| Xu_2009_CUTs29 | Xu_2009_CU | 43.1573343 | 0.57450375 | 0.21458901 | 2.67722828 | 0.0074234 |
| PXA2 | YKL188C | 204.07908 | 0.33977923 | 0.12692939 | 2.67691536 | 0.00743034 |
| BSC5 | YNR069C | 183.307707 | 0.36774559 | 0.13738656 | 2.67672176 | 0.00743463 |
| RMR1 | YGL250W | 68.9076968 | -0.5015956 | 0.18761146 | -2.6735873 | 0.00750448 |
| PHO13 | YDL236W | 958.829646 | -0.2020665 | 0.07558631 | -2.6733213 | 0.00751043 |
| YBR012W-B | YBR012W-B | 256.731805 | 0.32195611 | 0.12051811 | 2.67143341 | 0.00755281 |
| YTH1 | YPR107C | 458.927296 | -0.2364719 | 0.08851726 | -2.6714774 | 0.00755181 |
| Xu_2009_CUTs59 | Xu_2009_CU | 54.5747863 | 0.54027059 | 0.20249235 | 2.66810369 | 0.00762807 |
| YHR173C | YHR173C | 10.3272697 | 0.787161 | 0.29501674 | 2.66819097 | 0.00762609 |
| BLS1 | YLR408C | 85.2994001 | -0.4812805 | 0.18039097 | -2.6679853 | 0.00763076 |
| Xu_2009_SUTs74 | Xu_2009_SU | 22.840427 | 0.7026319 | 0.26353974 | 2.66613263 | 0.00767294 |
| YOL153C | YOL153C | 151.218063 | -0.399828 | 0.15005335 | -2.6645724 | 0.00770863 |
| KTR4 | YBR199W | 2023.46948 | -0.1739496 | 0.06529452 | -2.6640765 | 0.00772 |
| FAU1 | YER183C | 221.836366 | -0.3308753 | 0.12426993 | -2.6625531 | 0.00775503 |
| YOR338W | YOR338W | 1830.50233 | 0.2332668 | 0.08761159 | 2.6625108 | 0.00775601 |
| CDC11 | YJR076C | 1437.24611 | -0.1744187 | 0.06554429 | -2.6610819 | 0.007789 |
| Xu_2009_CUTs69 | Xu_2009_CU | 70.0784964 | 0.50474362 | 0.18968913 | 2.66089899 | 0.00779323 |
| RAM2 | YKL019W | 636.911953 | -0.2258276 | 0.08492793 | -2.6590495 | 0.00783614 |
| TRM13 | YOL125W | 836.529492 | 0.19630562 | 0.07382379 | 2.65911037 | 0.00783473 |
| MET31 | YPL038W | 299.095686 | -0.2939478 | 0.11057041 | -2.6584668 | 0.00784971 |
| SRS2 | YJL092W | 1419.51906 | 0.17690687 | 0.06655991 | 2.65785919 | 0.00786387 |
| YGL194C-A | YGL194C-A | 225.826334 | -0.3202244 | 0.12049424 | -2.6575908 | 0.00787014 |
| Xu_2009_CUTs26 | Xu_2009_CU | 7.71387888 | 0.78600685 | 0.29582215 | 2.65702501 | 0.00788336 |
| PUB1 | YNL016W | 4454.83927 | -0.1364829 | 0.05136759 | -2.6569843 | 0.00788431 |
| ARP5 | YNL059C | 1513.42593 | 0.16062222 | 0.06045341 | 2.6569589 | 0.00788491 |
| RIF2 | YLR453C | 124.448682 | 0.40087327 | 0.15093923 | 2.65585875 | 0.00791067 |
| ROX3 | YBL093C | 3514.18701 | 0.15741523 | 0.05929448 | 2.65480412 | 0.00793545 |
| SHE10 | YGL228W | 1706.59388 | -0.1554281 | 0.05861452 | -2.6516996 | 0.00800878 |
| YMR253C | YMR253C | 339.08151 | -0.2653233 | 0.1001987 | -2.6479714 | 0.00809764 |

| | | | | | | |
|---|---|---|---|---|---|---|
| IBA57 | YJR122W | 360.253368 | -0.2557196 | 0.0965871 | -2.647554 | 0.00810764 |
| CRZ1 | YNL027W | 1195.20103 | 0.18400872 | 0.0695621 | 2.64524405 | 0.0081632 |
| YPR158W-B | YPR158W-B | 100.630404 | 0.45561778 | 0.17224213 | 2.64521684 | 0.00816386 |
| Xu_2009_CUTs14 | Xu_2009_CU | 79.0462135 | 0.49381608 | 0.18672395 | 2.64463167 | 0.00817799 |
| RAD6 | YGL058W | 1007.81138 | -0.1821179 | 0.06887593 | -2.6441445 | 0.00818977 |
| FYV5 | YCL058C | 27.3919071 | 0.66444238 | 0.2513877 | 2.64309819 | 0.00821512 |
| Xu_2009_CUTs40 | Xu_2009_CU | 22.6042012 | 0.69882243 | 0.26441445 | 2.64290556 | 0.0082198 |
| GTR1 | YML121W | 734.150256 | -0.1979514 | 0.07495667 | -2.6408775 | 0.00826916 |
| YCL001W-B | YCL001W-B | 40.7473449 | 0.61412281 | 0.23262712 | 2.63994507 | 0.00829195 |
| PEX31 | YGR004W | 346.680775 | -0.2773392 | 0.10506016 | -2.6398134 | 0.00829517 |
| ALK1 | YGL021W | 1453.08607 | 0.25830174 | 0.09786796 | 2.63928812 | 0.00830803 |
| YOR392W | YOR392W | 12.5626584 | 0.76345438 | 0.28934916 | 2.63852293 | 0.00832681 |
| SAM37 | YMR060C | 176.42044 | -0.3466197 | 0.1314494 | -2.6369056 | 0.00836661 |
| YAP1802 | YGR241C | 1356.2485 | 0.18533417 | 0.07029039 | 2.63669293 | 0.00837186 |
| STE5 | YDR103W | 1328.76277 | 0.16959269 | 0.06434627 | 2.63562575 | 0.00839823 |
| DCC1 | YCL016C | 225.978449 | -0.3030338 | 0.11502478 | -2.6345091 | 0.00842591 |
| YBR182C-A | YBR182C-A | 19.6009414 | 0.70685474 | 0.26839404 | 2.63364546 | 0.00844736 |
| YMR31 | YFR049W | 215.535833 | -0.3151471 | 0.11965934 | -2.6337026 | 0.00844594 |
| Xu_2009_CUTs73 | Xu_2009_CU | 15.1337724 | 0.75399043 | 0.28642958 | 2.63237629 | 0.00847899 |
| SPT2 | YER161C | 1064.38557 | 0.17393509 | 0.06608048 | 2.63217069 | 0.00848412 |
| PDC2 | YDR081C | 1000.15517 | 0.20005638 | 0.07600963 | 2.63198723 | 0.00848871 |
| Xu_2009_CUTs55 | Xu_2009_CU | 25.9725595 | 0.67113887 | 0.25507779 | 2.63111448 | 0.00851054 |
| AGC1 | YPR021C | 614.878171 | -0.2089416 | 0.07943256 | -2.6304272 | 0.00852776 |
| TDA8 | YAL064C-A | 8.73319973 | 0.777579 | 0.29588728 | 2.62795685 | 0.00858994 |
| DBR1 | YKL149C | 361.965005 | -0.2589082 | 0.09859077 | -2.6260897 | 0.00863721 |
| TPK2 | YPL203W | 647.474745 | -0.2728461 | 0.10392262 | -2.6254736 | 0.00865285 |
| PEP7 | YDR323C | 212.76768 | 0.31152634 | 0.11866833 | 2.62518511 | 0.00866019 |
| Xu_2009_CUTs12 | Xu_2009_CU | 16.7644426 | 0.73354232 | 0.27948232 | 2.62464661 | 0.00867389 |
| YML053C | YML053C | 334.3179 | -0.2566883 | 0.09786491 | -2.622884 | 0.00871889 |
| QDR2 | YIL121W | 404.141493 | -0.2607394 | 0.09946465 | -2.6214281 | 0.00875622 |
| DAP2 | YHR028C | 2197.76658 | -0.1616454 | 0.06167023 | -2.621125 | 0.00876401 |
| COQ8 | YGL119W | 721.421543 | -0.2028723 | 0.07740665 | -2.6208636 | 0.00877074 |
| Xu_2009_CUTs30 | Xu_2009_CU | 36.441651 | 0.61784658 | 0.23582397 | 2.61994815 | 0.00879431 |
| AIM41 | YOR215C | 737.882203 | -0.2450221 | 0.09354705 | -2.6192395 | 0.0088126 |
| Xu_2009_SUTs21 | Xu_2009_SU | 109.151351 | 0.41365025 | 0.15808808 | 2.61658084 | 0.00888153 |
| Xu_2009_CUTs37 | Xu_2009_CU | 50.5761316 | 0.58349547 | 0.22302188 | 2.61631488 | 0.00888845 |
| NSE3 | YDR288W | 383.938431 | -0.2641125 | 0.10095 | -2.6162702 | 0.00888962 |
| Xu_2009_SUTs75 | Xu_2009_SU | 220.758054 | 0.32643965 | 0.12478608 | 2.61599406 | 0.00889681 |
| IPT1 | YDR072C | 1977.66267 | -0.2044412 | 0.07817448 | -2.6151915 | 0.00891774 |
| FAS1 | YKL182W | 22144.1235 | -0.1447276 | 0.05541106 | -2.6118906 | 0.00900431 |
| YMR316C-A | YMR316C-A | 12.9558607 | 0.7652719 | 0.29313799 | 2.61061998 | 0.00903783 |

| | | | | | | |
|---|---|---|---|---|---|---|
| STB2 | YMR053C | 344.473973 | 0.27273426 | 0.10451809 | 2.60944548 | 0.00906891 |
| YHL009W-B | YHL009W-B | 45.1553731 | 0.56311109 | 0.21595846 | 2.60749721 | 0.00912068 |
| AXL1 | YPR122W | 1648.03302 | 0.1591041 | 0.06102253 | 2.60730104 | 0.00912591 |
| EAT1 | YGR015C | 186.219275 | -0.3446418 | 0.13219943 | -2.6069836 | 0.00913437 |
| FEN2 | YCR028C | 412.660816 | 0.24268409 | 0.09315669 | 2.60511719 | 0.00918428 |
| PRR1 | YKL116C | 732.481777 | -0.2028109 | 0.07789132 | -2.6037678 | 0.00922052 |
| PUS5 | YLR165C | 268.618284 | 0.31126365 | 0.11955452 | 2.60352886 | 0.00922695 |
| UFD2 | YDL190C | 1845.66237 | 0.16693668 | 0.06412481 | 2.60330871 | 0.00923288 |
| Xu_2009_SUTs68! | Xu_2009_SU | 116.894212 | 0.37896712 | 0.14563509 | 2.60216902 | 0.00926362 |
| Xu_2009_CUTs54 | Xu_2009_CU | 11.2338141 | 0.75872574 | 0.29163664 | 2.60161328 | 0.00927864 |
| PAI3 | YMR174C | 364.751442 | 0.25175627 | 0.09680117 | 2.60075643 | 0.00930185 |
| VPS52 | YDR484W | 375.792968 | -0.2524 | 0.09716533 | -2.5976347 | 0.00938683 |
| RPB7 | YDR404C | 564.054 | -0.2316327 | 0.08918781 | -2.5971339 | 0.00940053 |
| CCT7 | YJL111W | 1635.33512 | -0.1983724 | 0.07639193 | -2.5967724 | 0.00941043 |
| FOX2 | YKR009C | 215.788367 | -0.3050871 | 0.11748519 | -2.5968134 | 0.0094093 |
| Xu_2009_SUTs72( | Xu_2009_SU | 34.4930378 | 0.61442234 | 0.23668164 | 2.59598645 | 0.00943198 |
| IFM1 | YOL023W | 467.362654 | -0.2386617 | 0.09195364 | -2.5954565 | 0.00944653 |
| FPY1 | YMR178W | 406.797232 | -0.2374909 | 0.09151138 | -2.5952064 | 0.00945341 |
| Xu_2009_CUTs68 | Xu_2009_CU | 60.8405526 | 0.51435731 | 0.19828613 | 2.59401561 | 0.00948622 |
| YOR343C | YOR343C | 69.6346367 | 0.47918882 | 0.18485483 | 2.59224398 | 0.00953521 |
| Xu_2009_SUTs16 | Xu_2009_SU | 60.83573 | 0.50311735 | 0.19413023 | 2.59164872 | 0.00955172 |
| ESF2 | YNR054C | 762.031877 | 0.21911346 | 0.08455238 | 2.59145228 | 0.00955718 |
| GFD2 | YCL036W | 796.336444 | -0.2600484 | 0.10040909 | -2.5898888 | 0.00960069 |
| THI20 | YOL055C | 337.412998 | -0.2543121 | 0.09820592 | -2.5895805 | 0.00960929 |
| SMD1 | YGR074W | 360.63341 | -0.2616328 | 0.10105864 | -2.5889211 | 0.00962772 |
| SNR51 | snR51 | 246.759792 | 0.30948236 | 0.11955004 | 2.5887266 | 0.00963315 |
| tM(CAU)J3/IMT3 | tM(CAU)J3 | 2.36371008 | -0.6398043 | 0.24717974 | -2.5884172 | 0.00964181 |
| CSE1 | YGL238W | 2191.55251 | -0.1662176 | 0.0642889 | -2.585479 | 0.00972438 |
| ARP7 | YPR034W | 984.15574 | -0.181144 | 0.07006904 | -2.5852219 | 0.00973163 |
| SNF7 | YLR025W | 505.785769 | 0.24807989 | 0.09597015 | 2.58496921 | 0.00973877 |
| STP3 | YLR375W | 1002.56566 | -0.1735928 | 0.06715502 | -2.584957 | 0.00973911 |
| Xu_2009_CUTs12 | Xu_2009_CU | 80.9843693 | 0.44544308 | 0.17237933 | 2.58408646 | 0.00976373 |
| YGR054W | YGR054W | 3894.03832 | -0.1580286 | 0.06116046 | -2.5838354 | 0.00977084 |
| REC102 | YLR329W | 54.0795823 | -0.546874 | 0.21167954 | -2.5834997 | 0.00978036 |
| MYO3 | YKL129C | 2312.23143 | 0.2037014 | 0.07888525 | 2.58224962 | 0.00981586 |
| AI4 | Q0065 | 67.7794048 | -0.4825348 | 0.18688688 | -2.581962 | 0.00982404 |
| SNR66 | snR66 | 269.775404 | -0.3213939 | 0.12454178 | -2.5806114 | 0.00986255 |
| TGL1 | YKL140W | 826.616135 | -0.1925272 | 0.07462733 | -2.5798488 | 0.00988436 |
| RRP46 | YGR095C | 449.089845 | -0.2381563 | 0.0923524 | -2.5787777 | 0.00991506 |
| FUS2 | YMR232W | 236.057036 | 0.31883892 | 0.12368628 | 2.57780351 | 0.00994305 |
| RCK1 | YGL158W | 99.9253457 | 0.44490196 | 0.17269469 | 2.57623415 | 0.0099883 |

| | | | | | | |
|---|---|---|---|---|---|---|
| YKL068W-A | YKL068W-A | 767.936474 | -0.2153304 | 0.08362457 | -2.5749657 | 0.010025 |
| GTS1 | YGL181W | 676.533694 | -0.2070532 | 0.08042304 | -2.5745506 | 0.01003704 |
| HBS1 | YKR084C | 1074.93407 | -0.1732295 | 0.06731636 | -2.5733646 | 0.0100715 |
| Xu_2009_SUTs31 | Xu_2009_SU | 40.8537426 | -0.5683919 | 0.22103181 | -2.571539 | 0.01012476 |
| CRD1 | YDL142C | 246.311042 | -0.3063693 | 0.11914505 | -2.5713979 | 0.01012889 |
| Xu_2009_SUTs66 | Xu_2009_SU | 148.796527 | 0.34569309 | 0.13444325 | 2.57129379 | 0.01013193 |
| Xu_2009_CUTs73 | Xu_2009_CU | 35.0985554 | 0.60013166 | 0.23355915 | 2.56950606 | 0.01018436 |
| Xu_2009_CUTs19 | Xu_2009_CU | 85.5925923 | 0.43970962 | 0.17116923 | 2.56885903 | 0.0102034 |
| LDB19 | YOR322C | 1287.23191 | 0.16662659 | 0.06486421 | 2.56885232 | 0.01020359 |
| Xu_2009_CUTs20 | Xu_2009_CU | 13.3782428 | 0.73799708 | 0.28751295 | 2.56683071 | 0.01026327 |
| KIN28 | YDL108W | 308.245204 | -0.2697461 | 0.10513577 | -2.565693 | 0.01029699 |
| ECM3 | YOR092W | 1023.10147 | -0.1991977 | 0.07765368 | -2.5652056 | 0.01031147 |
| MBR1 | YKL093W | 316.833964 | -0.2753294 | 0.10737271 | -2.5642401 | 0.0103402 |
| FPR3 | YML074C | 3370.3663 | -0.1423097 | 0.05552625 | -2.5629268 | 0.01037939 |
| RIM9 | YMR063W | 120.850108 | -0.3897499 | 0.15207014 | -2.5629617 | 0.01037835 |
| Xu_2009_SUTs42 | Xu_2009_SU | 19.6019333 | 0.69624358 | 0.27172285 | 2.56232991 | 0.01039725 |
| NOC3 | YLR002C | 1413.30686 | 0.17670433 | 0.06896195 | 2.56234532 | 0.01039679 |
| VPS73 | YGL104C | 199.663834 | -0.3128153 | 0.12218135 | -2.5602542 | 0.01045956 |
| LCP5 | YER127W | 1166.40373 | 0.20032054 | 0.07826321 | 2.55957484 | 0.01048003 |
| RHO4 | YKR055W | 166.91775 | -0.3341517 | 0.13057632 | -2.5590525 | 0.01049579 |
| SET6 | YPL165C | 44.0691937 | 0.55377513 | 0.21642206 | 2.55877398 | 0.0105042 |
| MXR1 | YER042W | 224.411774 | -0.3058897 | 0.11956027 | -2.5584564 | 0.0105138 |
| LYS5 | YGL154C | 210.871106 | 0.30982732 | 0.12115239 | 2.55733571 | 0.01054774 |
| GPA1 | YHR005C | 1080.80463 | 0.17721654 | 0.06929837 | 2.55729722 | 0.0105489 |
| CMK2 | YOL016C | 2067.33657 | 0.22470336 | 0.08787194 | 2.55716872 | 0.0105528 |
| YRR1 | YOR162C | 397.165921 | 0.24289576 | 0.09500472 | 2.55667044 | 0.01056793 |
| VPS61 | YDR136C | 21.9945843 | 0.666732 | 0.26079254 | 2.55656088 | 0.01057126 |
| RBL2 | YOR265W | 791.958378 | 0.20242394 | 0.07918484 | 2.55634704 | 0.01057776 |
| ALF1 | YNL148C | 173.50477 | 0.34808235 | 0.13618624 | 2.55592898 | 0.01059047 |
| RIC1 | YLR039C | 692.311191 | 0.21165272 | 0.08282499 | 2.55542098 | 0.01060594 |
| Xu_2009_CUTs21 | Xu_2009_CU | 7.02906016 | 0.75114978 | 0.29397062 | 2.55518652 | 0.01061309 |
| SYP1 | YCR030C | 1029.0151 | -0.1829983 | 0.07162962 | -2.554785 | 0.01062534 |
| YDR338C | YDR338C | 439.379904 | -0.2333974 | 0.09137632 | -2.5542439 | 0.01064187 |
| ASA1 | YPR085C | 359.172236 | 0.27709303 | 0.10849605 | 2.55394584 | 0.01065098 |
| MRP20 | YDR405W | 622.307756 | -0.2116211 | 0.08294817 | -2.5512449 | 0.01073389 |
| ECM22 | YLR228C | 564.679922 | -0.2149964 | 0.08428713 | -2.5507619 | 0.01074877 |
| TAF14 | YPL129W | 1045.57604 | -0.1713125 | 0.06715916 | -2.5508432 | 0.01074627 |
| YIR044C | YIR044C | 2.9095905 | 0.67067133 | 0.26310631 | 2.54905072 | 0.01080166 |
| SAW1 | YAL027W | 185.120192 | -0.3125346 | 0.12268467 | -2.5474628 | 0.01085094 |
| tH(GUG)E2 | tH(GUG)E2 | 82.584064 | -0.4574981 | 0.17970476 | -2.5458319 | 0.01090177 |
| TUM1 | YOR251C | 481.066245 | -0.2619758 | 0.1029129 | -2.545607 | 0.01090879 |

| | | | | | | |
|---|---|---|---|---|---|---|
| Xu_2009_SUTs28( | Xu_2009_SU | 22.7598892 | 0.66984763 | 0.26344955 | 2.54260306 | 0.01100301 |
| HSM3 | YBR272C | 502.452911 | -0.2259526 | 0.08886698 | -2.542593 | 0.01100333 |
| Xu_2009_SUTs84( | Xu_2009_SU | 148.235652 | -0.3471218 | 0.136557 | -2.5419556 | 0.01102342 |
| SNA2 | YDR525W-A | 127.324569 | -0.4173993 | 0.16424314 | -2.5413498 | 0.01104254 |
| COX12 | YLR038C | 376.181079 | -0.2776785 | 0.10926098 | -2.5414242 | 0.01104019 |
| Xu_2009_SUTs5 | Xu_2009_SU | 90.0702598 | 0.42317331 | 0.166545 | 2.54089469 | 0.01105692 |
| YDR262W | YDR262W | 1339.42071 | -0.162295 | 0.06389763 | -2.5399223 | 0.01108771 |
| SPT8 | YLR055C | 544.902098 | 0.22186892 | 0.08739361 | 2.53873144 | 0.01112552 |
| TAF4 | YMR005W | 1012.23953 | -0.1739233 | 0.06850988 | -2.5386605 | 0.01112778 |
| MDR1 | YGR100W | 884.4349 | 0.18998755 | 0.07488998 | 2.53688884 | 0.01118425 |
| NBL1 | YHR199C-A | 138.989874 | 0.35540931 | 0.14015028 | 2.53591586 | 0.01121537 |
| tR(UCU)M1 | tR(UCU)M1 | 4.84128667 | -0.727874 | 0.28707444 | -2.5354888 | 0.01122905 |
| Xu_2009_SUTs22 | Xu_2009_SU | 20.3671986 | 0.68326445 | 0.26966702 | 2.53373384 | 0.01128544 |
| RRN5 | YLR141W | 194.449757 | -0.3071587 | 0.1212476 | -2.5333179 | 0.01129884 |
| IRC18 | YJL037W | 47.8518504 | -0.542438 | 0.21419375 | -2.5324643 | 0.01132639 |
| YLR154W-B | YLR154W-B | 164.587109 | -0.7463915 | 0.29485981 | -2.5313436 | 0.01136265 |
| Xu_2009_SUTs22( | Xu_2009_SU | 62.2284972 | 0.47829545 | 0.18896841 | 2.53108686 | 0.01137097 |
| Xu_2009_SUTs16 | Xu_2009_SU | 110.78161 | 0.39076696 | 0.15441342 | 2.53065417 | 0.011385 |
| Xu_2009_CUTs50 | Xu_2009_CU | 14.8524453 | 0.7161482 | 0.28302839 | 2.53030519 | 0.01139633 |
| CAF120 | YNL278W | 1411.09266 | 0.17895565 | 0.07076038 | 2.52903757 | 0.01143758 |
| CWH43 | YCR017C | 1708.06474 | -0.1557003 | 0.06156806 | -2.5289138 | 0.01144161 |
| ARE1 | YCR048W | 494.3916 | -0.2270561 | 0.08981769 | -2.527966 | 0.01147255 |
| AIM32 | YML050W | 132.736009 | -0.3916697 | 0.15513452 | -2.5247101 | 0.01157938 |
| ATE1 | YGL017W | 434.791787 | -0.228284 | 0.09042511 | -2.5245638 | 0.0115842 |
| SAC1 | YKL212W | 2425.94523 | -0.1414233 | 0.05603077 | -2.5240285 | 0.01160185 |
| GAL4 | YPL248C | 194.261294 | -0.3166478 | 0.12545037 | -2.5240879 | 0.01159989 |
| DUR1,2 | YBR208C | 976.729541 | 0.17634309 | 0.06990522 | 2.52260267 | 0.01164899 |
| ART10 | YLR392C | 389.476645 | -0.253557 | 0.10053448 | -2.5220898 | 0.01166599 |
| Xu_2009_SUTs69 | Xu_2009_SU | 269.730617 | 0.26804027 | 0.10637433 | 2.51978346 | 0.0117427 |
| OKP1 | YGR179C | 425.330069 | -0.2323267 | 0.09222063 | -2.5192482 | 0.01176057 |
| NUP145 | YGL092W | 2793.07921 | 0.14162814 | 0.05622851 | 2.51879575 | 0.01177569 |
| AIF1 | YNR074C | 456.187756 | 0.22418067 | 0.08901215 | 2.51854021 | 0.01178424 |
| ATO2 | YNR002C | 151.471703 | -0.3340552 | 0.13270806 | -2.5172187 | 0.01182854 |
| TIM12 | YBR091C | 104.392954 | -0.45928 | 0.18249474 | -2.5166754 | 0.01184679 |
| Xu_2009_SUTs31 | Xu_2009_SU | 140.089795 | -0.3400455 | 0.13515592 | -2.5159496 | 0.01187121 |
| FMP40 | YPL222W | 441.205491 | -0.2297472 | 0.09132885 | -2.5156036 | 0.01188288 |
| VAC17 | YCL063W | 586.793317 | 0.20530269 | 0.08169724 | 2.51296976 | 0.01197196 |
| PBP4 | YDL053C | 774.877743 | -0.2006956 | 0.07988647 | -2.5122602 | 0.01199606 |
| Xu_2009_CUTs38 | Xu_2009_CU | 11.2654052 | 0.73745351 | 0.29365378 | 2.51130263 | 0.01202865 |
| YOR283W | YOR283W | 1232.00473 | -0.1780677 | 0.07094028 | -2.5101067 | 0.01206947 |
| Xu_2009_SUTs11( | Xu_2009_SU | 118.979017 | 0.38469625 | 0.15340649 | 2.50769215 | 0.01215225 |

| | | | | | | |
|---|---|---|---|---|---|---|
| Xu_2009_SUTs82 | Xu_2009_SU | 33.5194503 | 0.62914089 | 0.25093898 | 2.5071469 | 0.01217101 |
| CLB3 | YDL155W | 815.61579 | 0.19151799 | 0.07642725 | 2.50588604 | 0.0122145 |
| tG(GCC)G1 | tG(GCC)G1 | 24.6926058 | -0.6557707 | 0.26176632 | -2.5051761 | 0.01223904 |
| MCH2 | YKL221W | 58.3775093 | 0.51236529 | 0.20451837 | 2.50522863 | 0.01223723 |
| Xu_2009_SUTs81 | Xu_2009_SU | 61.1667676 | 0.49609305 | 0.19804552 | 2.50494454 | 0.01224706 |
| Xu_2009_SUTs24 | Xu_2009_SU | 11.8109504 | 0.72646817 | 0.29007613 | 2.50440516 | 0.01226575 |
| TGL3 | YMR313C | 874.963768 | -0.1868306 | 0.07463466 | -2.5032689 | 0.0123052 |
| SLI1 | YGR212W | 225.211608 | 0.29821973 | 0.11915372 | 2.50281522 | 0.01232099 |
| YHR056W-A | YHR056W-A | 11.7920921 | 0.73231259 | 0.29278114 | 2.50122868 | 0.01237632 |
| UBP7 | YIL156W | 775.663816 | 0.18608287 | 0.07442759 | 2.50018678 | 0.01241278 |
| RKM1 | YPL208W | 250.409961 | -0.3227923 | 0.12916545 | -2.4990602 | 0.01245231 |
| YOL118C | YOL118C | 27.3213109 | 0.65663312 | 0.26287704 | 2.49787171 | 0.01249414 |
| CUE5 | YOR042W | 1620.32064 | -0.1810463 | 0.07249505 | -2.4973613 | 0.01251214 |
| YDR133C | YDR133C | 21.2358583 | -0.6734752 | 0.26984265 | -2.4958071 | 0.01256709 |
| ROY1 | YMR258C | 487.487923 | -0.2250314 | 0.0901697 | -2.4956437 | 0.01257288 |
| YKT6 | YKL196C | 2317.86988 | -0.1403745 | 0.05625369 | -2.495383 | 0.01258213 |
| Xu_2009_CUTs91 | Xu_2009_CU | 28.6383484 | 0.60764874 | 0.24353812 | 2.49508677 | 0.01259263 |
| ECL1 | YGR146C | 992.799791 | -0.1790968 | 0.07178538 | -2.4948924 | 0.01259953 |
| ICY1 | YMR195W | 743.200247 | -0.2026584 | 0.081247 | -2.4943488 | 0.01261885 |
| SLU7 | YDR088C | 637.345949 | 0.20166113 | 0.08087473 | 2.49349977 | 0.01264907 |
| ORC3 | YLL004W | 411.381224 | 0.22278022 | 0.08936863 | 2.49282357 | 0.01267318 |
| Xu_2009_SUTs48 | Xu_2009_SU | 26.493645 | 0.64562435 | 0.25909012 | 2.491891 | 0.0127065 |
| ARV1 | YLR242C | 179.740938 | 0.33615043 | 0.13490029 | 2.49184362 | 0.0127082 |
| IRC6 | YFR043C | 211.065019 | -0.3078027 | 0.12353525 | -2.4916181 | 0.01271627 |
| STU1 | YBL034C | 935.956756 | 0.18467338 | 0.07414568 | 2.49068284 | 0.01274979 |
| YLR035C-A | YLR035C-A | 104.978669 | 0.40137719 | 0.16115741 | 2.49059102 | 0.01275308 |
| FRE5 | YOR384W | 25.1385575 | -0.6530046 | 0.26220451 | -2.4904401 | 0.0127585 |
| Xu_2009_CUTs56 | Xu_2009_CU | 12.0205316 | 0.7270223 | 0.29199627 | 2.48983421 | 0.01278027 |
| ELC1 | YPL046C | 159.177457 | -0.3562415 | 0.14307775 | -2.4898456 | 0.01277986 |
| EEB1 | YPL095C | 258.637217 | -0.2837716 | 0.11398108 | -2.4896379 | 0.01278733 |
| ALG11 | YNL048W | 544.790655 | -0.2019587 | 0.08117224 | -2.4880269 | 0.0128454 |
| MAK11 | YKL021C | 588.936977 | 0.21670956 | 0.08712187 | 2.48742999 | 0.01286698 |
| tD(GUC)G1 | tD(GUC)G1 | 18.5098128 | -0.7001743 | 0.28159661 | -2.4864444 | 0.01290267 |
| TPC1 | YGR096W | 94.2258207 | 0.41050157 | 0.16527284 | 2.4837812 | 0.01299957 |
| MET30 | YIL046W | 1127.06464 | 0.19893233 | 0.08009709 | 2.48363998 | 0.01300472 |
| AIM18 | YHR198C | 236.288547 | -0.2974886 | 0.11986059 | -2.4819552 | 0.01306637 |
| REV3 | YPL167C | 571.122905 | 0.20236797 | 0.08156798 | 2.48097305 | 0.01310243 |
| ADE6 | YGR061C | 3556.4724 | -0.1326452 | 0.05349646 | -2.4795141 | 0.01315615 |
| SNR75 | snR75 | 47.4236207 | -0.541014 | 0.21822283 | -2.4791815 | 0.01316843 |
| Xu_2009_SUTs34 | Xu_2009_SU | 45.5980361 | 0.53230278 | 0.21475775 | 2.4786196 | 0.01318919 |
| Xu_2009_SUTs20 | Xu_2009_SU | 81.4308262 | 0.42617545 | 0.17196485 | 2.47827069 | 0.01320209 |

| | | | | | | |
|---|---|---|---|---|---|---|
| Xu_2009_SUTs64( | Xu_2009_SU | 48.8798076 | 0.54606569 | 0.22036464 | 2.47800957 | 0.01321176 |
| TSR2 | YLR435W | 723.459652 | -0.1879004 | 0.07585484 | -2.4771049 | 0.0132453 |
| FAT3 | YKL187C | 69.8661389 | 0.4661844 | 0.18837827 | 2.47472501 | 0.01333388 |
| SIP18 | YMR175W | 33.9021155 | 0.59808278 | 0.24169744 | 2.47451019 | 0.0133419 |
| IZH1 | YDR492W | 2811.05447 | 0.19316202 | 0.07808359 | 2.47378502 | 0.01336901 |
| Xu_2009_CUTs30 | Xu_2009_CU | 22.2097429 | -0.6800471 | 0.27492367 | -2.4735849 | 0.0133765 |
| YCR101C | YCR101C | 45.5297134 | 0.53748582 | 0.21732612 | 2.47317634 | 0.0133918 |
| YGL188C | YGL188C | 19.0327916 | -0.6881655 | 0.27826591 | -2.4730498 | 0.01339655 |
| Xu_2009_CUTs10 | Xu_2009_CU | 17.2652991 | 0.69019985 | 0.27914878 | 2.47251611 | 0.01341657 |
| MFA2 | YNL145W | 16.0717726 | -0.7029933 | 0.28436938 | -2.4721132 | 0.0134317 |
| THI80 | YOR143C | 305.888013 | -0.2700464 | 0.10924322 | -2.4719743 | 0.01343692 |
| Xu_2009_CUTs32 | Xu_2009_CU | 22.3574592 | 0.67032425 | 0.2711833 | 2.47184927 | 0.01344162 |
| Xu_2009_CUTs82 | Xu_2009_CU | 5.09231331 | 0.70632402 | 0.28592302 | 2.47032933 | 0.01349887 |
| YPT11 | YNL304W | 170.516066 | -0.3197159 | 0.12941961 | -2.4703824 | 0.01349687 |
| NFU1 | YKL040C | 695.011455 | -0.18421 | 0.07459386 | -2.4695058 | 0.01352998 |
| RSC6 | YCR052W | 1047.16382 | -0.1629113 | 0.06599197 | -2.4686539 | 0.01356223 |
| YLH47 | YPR125W | 2234.9675 | 0.169858 | 0.06880512 | 2.46868244 | 0.01356115 |
| RNH201 | YNL072W | 273.120345 | -0.2783808 | 0.11281327 | -2.4676248 | 0.01360128 |
| RPC25 | YKL144C | 361.946568 | -0.2450678 | 0.09933515 | -2.4670808 | 0.01362196 |
| YMR265C | YMR265C | 647.314046 | 0.23106029 | 0.09373575 | 2.46501785 | 0.01370064 |
| Xu_2009_CUTs39 | Xu_2009_CU | 8.97360414 | 0.72987345 | 0.2961035 | 2.46492679 | 0.01370413 |
| PWP2 | YCR057C | 2658.99186 | 0.18242282 | 0.07405632 | 2.46329855 | 0.01376652 |
| YHR214W-A | YHR214W-A | 21.804085 | 0.6587788 | 0.26758726 | 2.46192138 | 0.0138195 |
| tV(CAC)H | tV(CAC)H | 21.3888444 | -0.6894084 | 0.28006236 | -2.4616247 | 0.01383093 |
| ADE8 | YDR408C | 411.252246 | -0.2393617 | 0.09725485 | -2.4611796 | 0.0138481 |
| PFS1 | YHR185C | 10.6094342 | -0.7290684 | 0.29624927 | -2.4609966 | 0.01385517 |
| RCE1 | YMR274C | 133.616921 | -0.3747564 | 0.15228715 | -2.4608535 | 0.0138607 |
| YPS7 | YDR349C | 1110.70271 | 0.16783802 | 0.06820798 | 2.46068 | 0.0138674 |
| RPO31 | YOR116C | 2658.56582 | 0.14531716 | 0.05906405 | 2.46033199 | 0.01388085 |
| Xu_2009_SUTs12! | Xu_2009_SU | 38.3170989 | 0.56045326 | 0.22793619 | 2.45881649 | 0.01393959 |
| SPT5 | YML010W | 2149.62168 | 0.13977982 | 0.05685911 | 2.45835387 | 0.01395756 |
| SGV1 | YPR161C | 1206.4119 | 0.16100568 | 0.0654982 | 2.45816965 | 0.01396472 |
| Xu_2009_SUTs63 | Xu_2009_SU | 233.166851 | 0.28842092 | 0.11737897 | 2.45717706 | 0.01400336 |
| PRP28 | YDR243C | 407.47771 | 0.2255311 | 0.0917839 | 2.45719666 | 0.0140026 |
| IPI1 | YHR085W | 393.280428 | -0.2409966 | 0.09809512 | -2.4567639 | 0.01401948 |
| YGR117C | YGR117C | 491.21082 | -0.2276795 | 0.09271189 | -2.4557743 | 0.01405814 |
| MNN1 | YER001W | 1533.20212 | -0.1585794 | 0.06460764 | -2.4544991 | 0.0141081 |
| YLR466C-B | YLR466C-B | 53.9056219 | 0.48901585 | 0.19922519 | 2.45458839 | 0.0141046 |
| YBL112C | YBL112C | 2.67662561 | 0.62592375 | 0.25508098 | 2.45382369 | 0.01413463 |
| Xu_2009_SUTs79 | Xu_2009_SU | 54.0408196 | 0.50361384 | 0.2052445 | 2.45372634 | 0.01413845 |
| DRE2 | YKR071C | 1126.90358 | -0.1650057 | 0.06726801 | -2.4529598 | 0.01416862 |

| | | | | | | |
|---|---|---|---|---|---|---|
| YFR035C | YFR035C | 3.73929902 | 0.67035312 | 0.27341596 | 2.45177024 | 0.01421554 |
| ERG12 | YMR208W | 616.27575 | -0.2028631 | 0.08275273 | -2.4514369 | 0.01422871 |
| MFM1 | YPL060W | 227.062864 | 0.29959845 | 0.12224483 | 2.45080659 | 0.01425365 |
| YHL012W | YHL012W | 23.0533933 | -0.6581306 | 0.26861941 | -2.4500487 | 0.01428369 |
| CUE2 | YKL090W | 205.057057 | 0.29407623 | 0.12004973 | 2.44962009 | 0.0143007 |
| RXT3 | YDL076C | 538.276827 | -0.2061756 | 0.0841825 | -2.4491498 | 0.01431939 |
| BIG1 | YHR101C | 249.59004 | -0.2726407 | 0.11133465 | -2.4488399 | 0.01433171 |
| AHC1 | YOR023C | 1743.4358 | 0.14685192 | 0.06003282 | 2.44619408 | 0.01443733 |
| YLR279W | YLR279W | 8.18875929 | 0.72346369 | 0.29578623 | 2.44590048 | 0.01444909 |
| OCA1 | YNL099C | 484.998339 | -0.2082286 | 0.08514094 | -2.4456932 | 0.0144574 |
| CWC2 | YDL209C | 377.85446 | -0.2291987 | 0.09371884 | -2.4455991 | 0.01446117 |
| YLR402W | YLR402W | 99.9141779 | 0.38971919 | 0.15941325 | 2.44471024 | 0.01449686 |
| YDR124W | YDR124W | 409.597595 | 0.23736126 | 0.0971524 | 2.44318461 | 0.01455829 |
| DGA1 | YOR245C | 756.694167 | 0.19229431 | 0.07874492 | 2.4419901 | 0.01460655 |
| YBR138C | YBR138C | 344.621169 | 0.27997228 | 0.11474247 | 2.44000578 | 0.01468703 |
| Xu_2009_CUTs71 | Xu_2009_CU | 8.63663113 | 0.72144915 | 0.2958858 | 2.43826889 | 0.01475779 |
| UME6 | YDR207C | 805.727299 | 0.18715693 | 0.07676 | 2.43820911 | 0.01476023 |
| AIM34 | YMR003W | 145.207433 | -0.3544613 | 0.14537954 | -2.4381788 | 0.01476147 |
| DIG1 | YPL049C | 2701.97349 | 0.15775598 | 0.0647013 | 2.43821961 | 0.0147598 |
| NAM2 | YLR382C | 579.413126 | -0.2013127 | 0.08257639 | -2.4378966 | 0.014773 |
| CRH1 | YGR189C | 7604.59875 | -0.1119997 | 0.04595895 | -2.4369503 | 0.01481172 |
| NEM1 | YHR004C | 850.833488 | -0.1970543 | 0.08085686 | -2.4370761 | 0.01480656 |
| YHR112C | YHR112C | 739.452128 | 0.19583742 | 0.08036234 | 2.43693036 | 0.01481253 |
| YDL012C | YDL012C | 1229.33914 | -0.1736529 | 0.07130195 | -2.4354583 | 0.01487294 |
| ERG24 | YNL280C | 625.021886 | -0.1965165 | 0.08072141 | -2.4345023 | 0.01491228 |
| MHR1 | YDR296W | 790.724844 | -0.1914472 | 0.07866385 | -2.433738 | 0.01494381 |
| YJR098C | YJR098C | 549.848587 | 0.20693669 | 0.0850835 | 2.43216012 | 0.01500907 |
| YLR257W | YLR257W | 4884.14935 | -0.1702824 | 0.07005368 | -2.4307416 | 0.01506796 |
| RBG1 | YAL036C | 1333.41782 | -0.177886 | 0.07319961 | -2.4301499 | 0.01509258 |
| RNQ1 | YCL028W | 1465.95136 | -0.1448932 | 0.05962225 | -2.4301863 | 0.01509106 |
| HUT1 | YPL244C | 721.689565 | -0.1943426 | 0.08001597 | -2.428798 | 0.01514897 |
| YFL051C | YFL051C | 12.7481482 | 0.70683638 | 0.29130776 | 2.42642478 | 0.01524841 |
| Xu_2009_CUTs58 | Xu_2009_CU | 17.9235878 | 0.68868228 | 0.28391968 | 2.42562363 | 0.01528211 |
| Xu_2009_SUTs16 | Xu_2009_SU | 92.3783366 | 0.42472937 | 0.17509656 | 2.42568658 | 0.01527946 |
| tL(CAA)K | tL(CAA)K | 3.24926115 | -0.638758 | 0.26350399 | -2.4240925 | 0.0153467 |
| MDM30 | YLR368W | 732.529582 | -0.1760352 | 0.07262224 | -2.4239849 | 0.01535125 |
| Xu_2009_SUTs99 | Xu_2009_SU | 30.7410041 | 0.58005945 | 0.23940406 | 2.42293074 | 0.01539586 |
| Xu_2009_SUTs59 | Xu_2009_SU | 106.018414 | 0.3772139 | 0.15573334 | 2.42217822 | 0.01542778 |
| RCF1 | YML030W | 552.971294 | 0.2247648 | 0.09281925 | 2.4215322 | 0.01545523 |
| HIS2 | YFR025C | 482.816962 | -0.2310629 | 0.09546731 | -2.4203353 | 0.0155062 |
| AVT5 | YBL089W | 403.956462 | 0.22366664 | 0.0924297 | 2.41985693 | 0.01552661 |

| | | | | | | |
|---|---|---|---|---|---|---|
| RSM18 | YER050C | 695.344032 | -0.1970081 | 0.08142964 | -2.4193657 | 0.0155476 |
| Xu_2009_CUTs61 | Xu_2009_CU | 39.0614717 | 0.53627526 | 0.22171819 | 2.41872464 | 0.01557503 |
| SOH1 | YGL127C | 543.659438 | -0.2227189 | 0.09211716 | -2.417778 | 0.0156156 |
| Xu_2009_CUTs33 | Xu_2009_CU | 13.3854695 | 0.70056589 | 0.28984456 | 2.41703997 | 0.0156473 |
| Xu_2009_CUTs24 | Xu_2009_CU | 61.5987803 | -0.4672731 | 0.1934807 | -2.4150888 | 0.01573137 |
| Xu_2009_SUTs83 | Xu_2009_SU | 40.5135559 | 0.53529351 | 0.22167909 | 2.4147226 | 0.0157472 |
| Xu_2009_CUTs65 | Xu_2009_CU | 117.26564 | 0.3689548 | 0.1528019 | 2.41459565 | 0.01575269 |
| YGL242C | YGL242C | 721.285976 | -0.1929384 | 0.07992731 | -2.4139231 | 0.01578179 |
| MTO1 | YGL236C | 698.775505 | -0.2014592 | 0.08346396 | -2.4137264 | 0.01579031 |
| RPC17 | YJL011C | 207.613437 | -0.320477 | 0.13281272 | -2.4129995 | 0.01582184 |
| YBR259W | YBR259W | 451.152901 | 0.21548727 | 0.08931487 | 2.41266958 | 0.01583617 |
| RSC3 | YDR303C | 919.799139 | -0.174319 | 0.07236741 | -2.4088056 | 0.01600482 |
| ARH1 | YDR376W | 842.162299 | -0.1698059 | 0.07054639 | -2.4070107 | 0.0160837 |
| Xu_2009_CUTs17 | Xu_2009_CU | 9.32799064 | 0.7101683 | 0.29511731 | 2.40639327 | 0.01611091 |
| TMH18 | YPR098C | 353.762061 | -0.2397265 | 0.09969665 | -2.4045596 | 0.01619197 |
| SRN2 | YLR119W | 142.628087 | -0.3532637 | 0.14694072 | -2.4041238 | 0.01621128 |
| Xu_2009_CUTs23 | Xu_2009_CU | 33.0020003 | 0.587911 | 0.24460205 | 2.40354078 | 0.01623716 |
| DOC1 | YGL240W | 162.050841 | 0.33710906 | 0.14031245 | 2.40255984 | 0.01628077 |
| PTC2 | YER089C | 1284.40469 | -0.1892684 | 0.07884067 | -2.4006441 | 0.01636625 |
| SHS1 | YDL225W | 1665.68714 | 0.14879755 | 0.06199825 | 2.40002818 | 0.01639381 |
| MIS1 | YBR084W | 3534.20524 | 0.13614848 | 0.05676171 | 2.39859713 | 0.01645801 |
| PCI8 | YIL071C | 473.846317 | 0.22539821 | 0.09398429 | 2.39825419 | 0.01647343 |
| YCR041W | YCR041W | 16.74494 | 0.69623084 | 0.29036365 | 2.39778925 | 0.01649435 |
| DAL2 | YIR029W | 162.340462 | 0.32072667 | 0.13379613 | 2.39712965 | 0.01652408 |
| EAF1 | YDR359C | 1146.0496 | 0.17938165 | 0.07484703 | 2.39664359 | 0.01654601 |
| Xu_2009_CUTs43 | Xu_2009_CU | 8.70682863 | 0.70904532 | 0.29605069 | 2.3950132 | 0.01661977 |
| Xu_2009_CUTs24 | Xu_2009_CU | 52.8390215 | 0.52937644 | 0.22109257 | 2.39436559 | 0.01664914 |
| ACS1 | YAL054C | 276.911927 | -0.2536722 | 0.10594259 | -2.3944308 | 0.01664618 |
| SSU72 | YNL222W | 330.420922 | 0.26543099 | 0.11090402 | 2.39333977 | 0.01669577 |
| OAR1 | YKL055C | 132.714538 | -0.3643896 | 0.15226514 | -2.3931257 | 0.01670552 |
| ATG21 | YPL100W | 555.755182 | -0.2043256 | 0.08539922 | -2.3925938 | 0.01672975 |
| GIP2 | YER054C | 577.205811 | 0.22322479 | 0.09334151 | 2.39148477 | 0.01678038 |
| ORC5 | YNL261W | 518.540276 | 0.20068472 | 0.08392846 | 2.39114031 | 0.01679613 |
| PEX32 | YBR168W | 297.642838 | -0.2755203 | 0.1152498 | -2.3906357 | 0.01681923 |
| PRP4 | YPR178W | 401.139547 | 0.22429477 | 0.09383634 | 2.39027623 | 0.0168357 |
| Xu_2009_CUTs24 | Xu_2009_CU | 4.82878445 | 0.68391406 | 0.28663046 | 2.38604811 | 0.01703052 |
| GCR1 | YPL075W | 1271.23607 | 0.15511109 | 0.06502418 | 2.38543711 | 0.01705884 |
| GPI10 | YGL142C | 588.484841 | -0.1917827 | 0.0804328 | -2.3843846 | 0.01710771 |
| GIS1 | YDR096W | 1184.26028 | -0.1719169 | 0.07212485 | -2.3836019 | 0.01714414 |
| YBR284W | YBR284W | 195.873836 | 0.32187136 | 0.13509802 | 2.38250246 | 0.01719542 |
| MSH3 | YCR092C | 505.403668 | 0.20665746 | 0.08674541 | 2.38234468 | 0.01720279 |

| | | | | | | |
|---|---|---|---|---|---|---|
| YJL120W | YJL120W | 11.3261875 | 0.69963042 | 0.29387093 | 2.38074046 | 0.01727788 |
| BUL2 | YML111W | 1207.19474 | 0.1599942 | 0.06725233 | 2.37901356 | 0.01735904 |
| ATP8 | Q0080 | 51.6832418 | -0.5385234 | 0.22641825 | -2.3784452 | 0.01738582 |
| ABZ1 | YNR033W | 1337.187 | 0.15413243 | 0.06480637 | 2.37835303 | 0.01739017 |
| MPS2 | YGL075C | 293.984548 | 0.27079701 | 0.11391926 | 2.37709591 | 0.01744955 |
| RAX1 | YOR301W | 583.873275 | 0.20882081 | 0.08790316 | 2.37557802 | 0.01752149 |
| Xu_2009_SUTs35 | Xu_2009_SU | 143.545477 | 0.38174451 | 0.16071493 | 2.37528955 | 0.01753519 |
| Xu_2009_CUTs36 | Xu_2009_CU | 11.5215899 | 0.69354598 | 0.29202511 | 2.37495326 | 0.01755117 |
| Xu_2009_CUTs48 | Xu_2009_CU | 31.5265636 | 0.57318144 | 0.24143315 | 2.37407931 | 0.01759277 |
| BIT61 | YJL058C | 395.642036 | 0.24052284 | 0.1013477 | 2.37324431 | 0.0176326 |
| Xu_2009_CUTs76 | Xu_2009_CU | 35.7757109 | 0.56106413 | 0.23657015 | 2.37166066 | 0.01770835 |
| VAR1 | Q0140 | 7.74159517 | -0.7017065 | 0.29590697 | -2.3713754 | 0.01772202 |
| Xu_2009_SUTs14 | Xu_2009_SU | 10.246889 | 0.69695988 | 0.29397761 | 2.37079238 | 0.01775 |
| YOL019W | YOL019W | 1885.39211 | -0.1580747 | 0.06669495 | -2.3701146 | 0.01778257 |
| Xu_2009_SUTs75 | Xu_2009_SU | 725.053534 | 0.19368842 | 0.08173743 | 2.36964156 | 0.01780534 |
| YJL206C | YJL206C | 491.789274 | 0.22620455 | 0.09546701 | 2.36945256 | 0.01781444 |
| Xu_2009_SUTs51 | Xu_2009_SU | 40.8476692 | -0.55111 | 0.23264884 | -2.3688493 | 0.01784352 |
| Xu_2009_SUTs27 | Xu_2009_SU | 40.6131273 | 0.52883652 | 0.22330058 | 2.36827199 | 0.01787139 |
| ASK1 | YKL052C | 531.446225 | -0.2082574 | 0.08797338 | -2.3672779 | 0.01791947 |
| CDC73 | YLR418C | 364.102135 | -0.247045 | 0.10435971 | -2.3672448 | 0.01792108 |
| YVH1 | YIR026C | 694.336861 | 0.20089664 | 0.08489018 | 2.36654735 | 0.01795488 |
| LEA1 | YPL213W | 399.525651 | -0.2359019 | 0.09969951 | -2.3661289 | 0.01797519 |
| MUD1 | YBR119W | 240.149217 | -0.2850036 | 0.12046081 | -2.365945 | 0.01798411 |
| Xu_2009_SUTs70 | Xu_2009_SU | 66.3571028 | 0.4409381 | 0.18638465 | 2.3657426 | 0.01799395 |
| MPC3 | YGR243W | 156.828227 | -0.3269545 | 0.13822976 | -2.3652972 | 0.01801561 |
| PCC1 | YKR095W-A | 244.957198 | -0.2675811 | 0.11316038 | -2.3646182 | 0.01804866 |
| YOL164W-A | YOL164W-A | 4.60293307 | 0.67210408 | 0.2843948 | 2.36327842 | 0.01811406 |
| HXT11 | YOL156W | 16.2905701 | -0.6596227 | 0.27917026 | -2.3627971 | 0.0181376 |
| PEX8 | YGR077C | 1015.00679 | -0.1628927 | 0.06895922 | -2.3621605 | 0.01816878 |
| STN1 | YDR082W | 336.455815 | 0.25725588 | 0.1089196 | 2.36188782 | 0.01818214 |
| YIR036W-A | YIR036W-A | 5.94403967 | 0.68799638 | 0.2912812 | 2.36196631 | 0.01817829 |
| OPI10 | YOL032W | 195.689566 | -0.3375791 | 0.14293032 | -2.3618442 | 0.01818428 |
| MRX12 | YJR003C | 519.562525 | 0.21452692 | 0.09084658 | 2.36141975 | 0.01820511 |
| Xu_2009_CUTs28 | Xu_2009_CU | 15.1350419 | 0.67409518 | 0.28549775 | 2.36112258 | 0.01821971 |
| YLR365W | YLR365W | 18.8763756 | 0.6447392 | 0.27307526 | 2.36103115 | 0.0182242 |
| tQ(UUG)C | tQ(UUG)C | 7.51261443 | -0.6989417 | 0.29605829 | -2.3608248 | 0.01823434 |
| PHM6 | YDR281C | 68.0140123 | -0.4488082 | 0.19011437 | -2.3607273 | 0.01823914 |
| XRS2 | YDR369C | 420.197371 | 0.21889243 | 0.09273098 | 2.36051018 | 0.01824982 |
| MRX4 | YPL168W | 350.965125 | 0.23017015 | 0.09751602 | 2.36033166 | 0.0182586 |
| Xu_2009_SUTs61 | Xu_2009_SU | 77.7182564 | 0.42766098 | 0.18120582 | 2.36008416 | 0.01827079 |
| CCR4 | YAL021C | 2645.5936 | 0.12579243 | 0.05331873 | 2.35925403 | 0.01831172 |

| | | | | | | |
|---|---|---|---|---|---|---|
| FRE3 | YOR381W | 137.254469 | 0.3405075 | 0.14438526 | 2.35832582 | 0.01835757 |
| IKS1 | YJL057C | 453.335655 | -0.2437324 | 0.10341239 | -2.3568974 | 0.01842834 |
| ARR2 | YPR200C | 41.1223362 | -0.5123252 | 0.21742086 | -2.3563754 | 0.01845426 |
| PPZ1 | YML016C | 1478.59296 | 0.15826314 | 0.06717707 | 2.35591014 | 0.01847739 |
| Xu_2009_CUTs55 | Xu_2009_CU | 1.53720951 | -0.4921783 | 0.20896441 | -2.355321 | 0.01850671 |
| PHO81 | YGR233C | 919.035454 | 0.18365684 | 0.07798701 | 2.35496707 | 0.01852435 |
| MSS1 | YMR023C | 320.596942 | 0.23067038 | 0.09799249 | 2.35395985 | 0.01857462 |
| BCP1 | YDR361C | 768.279662 | -0.1880719 | 0.07990646 | -2.3536509 | 0.01859006 |
| YPL025C | YPL025C | 57.0712314 | 0.47138828 | 0.20063658 | 2.34946332 | 0.0188005 |
| RMT2 | YDR465C | 1675.86031 | 0.19798248 | 0.08427529 | 2.34923531 | 0.01881201 |
| Xu_2009_SUTs47 | Xu_2009_SU | 69.4727836 | -0.4475317 | 0.19050958 | -2.3491294 | 0.01881737 |
| DAT1 | YML113W | 859.794576 | 0.21000547 | 0.08941638 | 2.34862408 | 0.01884292 |
| FRE1 | YLR214W | 326.926936 | -0.2641856 | 0.112525 | -2.3477949 | 0.01888492 |
| Xu_2009_SUTs9 | Xu_2009_SU | 289.117888 | 0.26008075 | 0.11080626 | 2.34716663 | 0.01891679 |
| COS3 | YML132W | 111.1182 | -0.3570209 | 0.15211755 | -2.3470069 | 0.0189249 |
| BUD32 | YGR262C | 657.339001 | -0.1877916 | 0.08003987 | -2.3462256 | 0.01896462 |
| SWC4 | YGR002C | 538.976199 | -0.2035772 | 0.08680925 | -2.3451095 | 0.01902149 |
| RCH1 | YMR034C | 226.839622 | -0.2794108 | 0.1192783 | -2.3425114 | 0.01915445 |
| TCM62 | YBR044C | 666.843502 | -0.1955553 | 0.08359509 | -2.3393153 | 0.01931912 |
| REF2 | YDR195W | 664.566832 | -0.1819298 | 0.07779703 | -2.3385187 | 0.01936035 |
| SHC1 | YER096W | 96.0193225 | 0.38129021 | 0.16304436 | 2.33856726 | 0.01935784 |
| tE(UUC)G1/SOE1 | tE(UUC)G1 | 86.4948733 | -0.4030316 | 0.17236887 | -2.3381926 | 0.01937726 |
| YPS5 | YGL259W | 10.3384338 | 0.68666235 | 0.29370988 | 2.33789325 | 0.01939279 |
| SNF8 | YPL002C | 461.426871 | -0.2279443 | 0.09765765 | -2.3341158 | 0.01958966 |
| YOL024W | YOL024W | 16.8778133 | -0.6438774 | 0.27589996 | -2.333735 | 0.0196096 |
| UMP1 | YBR173C | 1369.72066 | -0.143344 | 0.06147454 | -2.3317614 | 0.01971325 |
| CAC2 | YML102W | 437.07061 | 0.21468838 | 0.09207739 | 2.33160801 | 0.01972132 |
| GLC8 | YMR311C | 1693.79479 | -0.1532849 | 0.06575057 | -2.3313093 | 0.01973706 |
| YPL277C | YPL277C | 471.887445 | -0.2158285 | 0.0925846 | -2.3311488 | 0.01974551 |
| Xu_2009_CUTs14 | Xu_2009_CU | 31.9779136 | 0.5643484 | 0.24210317 | 2.33102441 | 0.01975207 |
| DHR2 | YKL078W | 635.675115 | 0.23452296 | 0.1006261 | 2.33063748 | 0.01977248 |
| Xu_2009_CUTs77 | Xu_2009_CU | 24.1312402 | 0.61269185 | 0.26294534 | 2.3301111 | 0.01980028 |
| Xu_2009_CUTs26 | Xu_2009_CU | 26.0484464 | 0.60151983 | 0.25827713 | 2.32897058 | 0.01986062 |
| LSM2 | YBL026W | 656.052819 | -0.1871075 | 0.08034051 | -2.3289315 | 0.01986269 |
| Xu_2009_SUTs72 | Xu_2009_SU | 27.4084883 | 0.58188196 | 0.24997124 | 2.32779562 | 0.01992296 |
| YIL141W | YIL141W | 14.4985535 | 0.67290824 | 0.28914352 | 2.3272465 | 0.01995215 |
| CSM1 | YCR086W | 192.41031 | -0.2936171 | 0.12619707 | -2.3266554 | 0.01998361 |
| YKU70 | YMR284W | 688.863232 | -0.2066347 | 0.08882309 | -2.3263622 | 0.01999924 |
| Xu_2009_SUTs10 | Xu_2009_SU | 14.1740123 | 0.67045378 | 0.28823775 | 2.32604429 | 0.02001619 |
| ACF2 | YLR144C | 712.014236 | 0.17148368 | 0.07372503 | 2.32598996 | 0.02001909 |
| Xu_2009_CUTs2 | Xu_2009_CU | 40.6717883 | 0.5197352 | 0.22349897 | 2.32544792 | 0.02004802 |

| | | | | | | |
|---|---|---|---|---|---|---|
| MMS21 | YEL019C | 172.822784 | 0.31241933 | 0.13441448 | 2.32429807 | 0.02010952 |
| UBX6 | YJL048C | 878.621581 | -0.1806779 | 0.0777434 | -2.3240284 | 0.02012397 |
| KEL2 | YGR238C | 566.064915 | -0.223303 | 0.09611174 | -2.3233685 | 0.02015937 |
| YIL156W-A | YIL156W-A | 7.72176507 | 0.68537438 | 0.29501692 | 2.32316975 | 0.02017003 |
| BIK1 | YCL029C | 610.319645 | 0.19739476 | 0.08499626 | 2.32239337 | 0.02021176 |
| PDP3 | YLR455W | 521.626137 | -0.2074381 | 0.08932328 | -2.3223296 | 0.02021519 |
| Xu_2009_CUTs71 | Xu_2009_CU | 31.7347729 | 0.55351492 | 0.23840188 | 2.3217725 | 0.02024519 |
| YGR035C | YGR035C | 350.616537 | -0.2435635 | 0.10492218 | -2.3213729 | 0.02026673 |
| RTT101 | YJL047C | 506.955751 | 0.18989852 | 0.08184804 | 2.32013522 | 0.02033356 |
| Xu_2009_CUTs44 | Xu_2009_CU | 42.3844094 | -0.5055468 | 0.21791768 | -2.3198982 | 0.02034639 |
| Xu_2009_CUTs92 | Xu_2009_CU | 40.2297663 | 0.53183269 | 0.22933909 | 2.31897966 | 0.02039614 |
| ZNF1 | YFL052W | 107.557307 | 0.37209099 | 0.16050201 | 2.31829491 | 0.0204333 |
| Xu_2009_SUTs56 | Xu_2009_SU | 100.333158 | 0.38846595 | 0.16759173 | 2.31793032 | 0.02045311 |
| SNG1 | YGR197C | 468.911158 | 0.19957835 | 0.08610456 | 2.3178605 | 0.0204569 |
| PRP21 | YJL203W | 329.568882 | 0.24998905 | 0.10785789 | 2.31776332 | 0.02046219 |
| BUD23 | YCR047C | 657.098904 | 0.2096359 | 0.09050572 | 2.31627251 | 0.02054339 |
| YHL037C | YHL037C | 14.9597218 | 0.66696326 | 0.28794558 | 2.31628231 | 0.02054286 |
| MKK1 | YOR231W | 940.401194 | -0.1678704 | 0.07248957 | -2.3157876 | 0.02056987 |
| Xu_2009_CUTs39 | Xu_2009_CU | 13.8691588 | 0.66225319 | 0.28618537 | 2.31407076 | 0.02066384 |
| YMR153C-A | YMR153C-A | 46.0010473 | 0.49661546 | 0.21465402 | 2.31356236 | 0.02069174 |
| LEU9 | YOR108W | 1051.87596 | -0.1664131 | 0.07192865 | -2.3135854 | 0.02069047 |
| Xu_2009_SUTs66 | Xu_2009_SU | 28.1471269 | 0.57742763 | 0.2496293 | 2.31314047 | 0.02071492 |
| AQY2 | YLL052C | 54.6771438 | -0.4835639 | 0.20905611 | -2.3130818 | 0.02071814 |
| Xu_2009_CUTs21 | Xu_2009_CU | 12.7882872 | 0.67011341 | 0.28974583 | 2.31276289 | 0.02073568 |
| Xu_2009_SUTs61 | Xu_2009_SU | 29.1061977 | 0.59591076 | 0.25769285 | 2.31248464 | 0.02075099 |
| DAS2 | YDR020C | 238.564934 | 0.27501115 | 0.11895561 | 2.31188044 | 0.02078427 |
| STB1 | YNL309W | 608.596024 | 0.19427625 | 0.08406646 | 2.31098406 | 0.02083374 |
| Xu_2009_CUTs24 | Xu_2009_CU | 36.7976152 | 0.52133108 | 0.22567877 | 2.31005814 | 0.02088494 |
| KGD1 | YIL125W | 2948.2769 | -0.1215771 | 0.05263323 | -2.3098919 | 0.02089414 |
| Xu_2009_CUTs42 | Xu_2009_CU | 29.8655864 | 0.55971955 | 0.2423645 | 2.30941228 | 0.02092071 |
| SPC29 | YPL124W | 182.817828 | -0.3154751 | 0.13667927 | -2.3081415 | 0.02099127 |
| SNR62 | snR62 | 129.095733 | 0.34677722 | 0.15034953 | 2.30647368 | 0.02108418 |
| AVT2 | YEL064C | 247.574227 | -0.268187 | 0.11644458 | -2.3031299 | 0.02127153 |
| MAD1 | YGL086W | 642.449511 | 0.17829716 | 0.07742408 | 2.30286447 | 0.02128647 |
| YLR456W | YLR456W | 88.4920119 | -0.3964453 | 0.17221333 | -2.3020594 | 0.02133182 |
| IGD1 | YFR017C | 315.381399 | -0.2760997 | 0.12002948 | -2.300266 | 0.02143316 |
| YEL076C | YEL076C | 12.9610072 | 0.66708353 | 0.29002214 | 2.30011245 | 0.02144185 |
| GEM1 | YAL048C | 440.09776 | 0.20429745 | 0.08882872 | 2.2999032 | 0.0214537 |
| VPS70 | YJR126C | 1301.40553 | -0.1518693 | 0.06603687 | -2.2997655 | 0.02146151 |
| NUP1 | YOR098C | 3012.11144 | 0.14082907 | 0.06124384 | 2.2994815 | 0.02147761 |
| MDM10 | YAL010C | 622.266037 | 0.19149196 | 0.08332598 | 2.29810631 | 0.02155574 |

| | | | | | | |
|---|---|---|---|---|---|---|
| FAP7 | YDL166C | 374.463088 | -0.2355704 | 0.10250527 | -2.2981297 | 0.02155441 |
| Xu_2009_CUTs59 | Xu_2009_CU | 24.506123 | 0.59100987 | 0.25719077 | 2.29794358 | 0.021565 |
| Xu_2009_SUTs34 | Xu_2009_SU | 68.778609 | 0.4387347 | 0.1910177 | 2.29682754 | 0.02162861 |
| YDL071C | YDL071C | 16.9025135 | 0.63777998 | 0.27768222 | 2.29679807 | 0.02163029 |
| RSC30 | YHR056C | 773.228141 | -0.1708048 | 0.07436246 | -2.2969229 | 0.02162317 |
| MRM1 | YOR201C | 823.681809 | -0.1823722 | 0.07945093 | -2.2954064 | 0.02170984 |
| Xu_2009_CUTs45 | Xu_2009_CU | 35.9016794 | 0.54159869 | 0.2359628 | 2.29527151 | 0.02171757 |
| PAU17 | YLL025W | 35.4221951 | -0.5322418 | 0.23191332 | -2.2950031 | 0.02173295 |
| AMA1 | YGR225W | 122.689034 | 0.34749144 | 0.15148493 | 2.29390102 | 0.02179618 |
| Xu_2009_CUTs46 | Xu_2009_CU | 12.6073896 | 0.66274811 | 0.28901062 | 2.2931618 | 0.02183869 |
| SPO7 | YAL009W | 264.719801 | -0.2554548 | 0.11140556 | -2.2930164 | 0.02184706 |
| Xu_2009_CUTs63 | Xu_2009_CU | 41.8998604 | 0.51940562 | 0.2266035 | 2.29213413 | 0.0218979 |
| SLD2 | YKL108W | 447.673145 | -0.2171341 | 0.09478275 | -2.2908613 | 0.02197143 |
| ERP4 | YOR016C | 502.881433 | -0.2130696 | 0.09313989 | -2.2876296 | 0.0221591 |
| RRD1 | YIL153W | 596.370556 | -0.1853003 | 0.08101036 | -2.2873651 | 0.02217452 |
| Xu_2009_CUTs71 | Xu_2009_CU | 14.7767069 | 0.64305543 | 0.28115697 | 2.28717584 | 0.02218556 |
| SAT4 | YCR008W | 2306.6884 | -0.1332321 | 0.0582524 | -2.2871521 | 0.02218695 |
| Xu_2009_CUTs27 | Xu_2009_CU | 20.6468744 | 0.61184586 | 0.26754687 | 2.28687351 | 0.02220321 |
| FSH2 | YMR222C | 374.171731 | -0.2147357 | 0.09390955 | -2.2866226 | 0.02221786 |
| SHY1 | YGR112W | 362.661587 | -0.2394201 | 0.10474798 | -2.2856771 | 0.02227316 |
| AIM22 | YJL046W | 430.463031 | 0.20336707 | 0.0890123 | 2.28470762 | 0.02232998 |
| HIF1 | YLL022C | 611.912991 | -0.2000604 | 0.08766991 | -2.2819735 | 0.0224909 |
| CHL1 | YPL008W | 492.234262 | 0.20007742 | 0.08773838 | 2.28038642 | 0.02258478 |
| ROT2 | YBR229C | 1636.79258 | -0.1368537 | 0.06004891 | -2.2790373 | 0.02266485 |
| ROK1 | YGL171W | 633.541034 | 0.19946452 | 0.08754556 | 2.27840804 | 0.02270228 |
| TIP20 | YGL145W | 635.376414 | 0.1943897 | 0.08533024 | 2.27808702 | 0.02272139 |
| YLR345W | YLR345W | 971.86449 | -0.1905367 | 0.08371295 | -2.2760722 | 0.02284169 |
| SIR4 | YDR227W | 1016.67876 | 0.15629694 | 0.06869532 | 2.27521982 | 0.02289275 |
| RTS1 | YOR014W | 2402.53691 | -0.1227575 | 0.0539704 | -2.2745346 | 0.02293386 |
| ERP5 | YHR110W | 495.633855 | -0.2205538 | 0.09698807 | -2.2740306 | 0.02296415 |
| MIP6 | YHR015W | 154.823812 | -0.3306104 | 0.1454896 | -2.2723987 | 0.02306244 |
| SPC42 | YKL042W | 267.407194 | -0.2411862 | 0.10615077 | -2.2721094 | 0.02307991 |
| SAM3 | YPL274W | 213.84474 | 0.29695121 | 0.13072728 | 2.27153199 | 0.02311479 |
| Xu_2009_SUTs28 | Xu_2009_SU | 82.8688513 | 0.38706941 | 0.17053409 | 2.26974803 | 0.02322288 |
| HOG1 | YLR113W | 2128.07161 | 0.13455994 | 0.05928552 | 2.26969341 | 0.02322619 |
| TIM44 | YIL022W | 2911.12321 | -0.1392037 | 0.06137711 | -2.2680072 | 0.02332877 |
| Xu_2009_SUTs59 | Xu_2009_SU | 82.6005969 | 0.40262144 | 0.1775505 | 2.26764461 | 0.02335088 |
| PAH1 | YMR165C | 1428.23295 | 0.14624507 | 0.06451938 | 2.26668418 | 0.02340953 |
| SPS4 | YOR313C | 48.5350432 | -0.4873304 | 0.21515321 | -2.2650388 | 0.0235103 |
| Xu_2009_CUTs44 | Xu_2009_CU | 6.42884558 | 0.66813241 | 0.29501083 | 2.2647725 | 0.02352664 |
| AFI1 | YOR129C | 456.889406 | 0.20064797 | 0.08862413 | 2.26403321 | 0.02357207 |

| | | | | | | |
|---|---|---|---|---|---|---|
| tL(CAA)N | tL(CAA)N | 10.2738452 | -0.670233 | 0.29636112 | -2.2615417 | 0.02372573 |
| Xu_2009_CUTs52 | Xu_2009_CU | 6.61929263 | 0.66412072 | 0.29365374 | 2.2615776 | 0.02372351 |
| ERV15 | YBR210W | 300.654104 | -0.2476572 | 0.10956116 | -2.2604471 | 0.02379352 |
| Xu_2009_SUTs18 | Xu_2009_SU | 10.7304989 | 0.66575512 | 0.29464077 | 2.25954855 | 0.02384928 |
| Xu_2009_CUTs30 | Xu_2009_CU | 9.93101035 | 0.66374297 | 0.29402235 | 2.25745754 | 0.0239795 |
| PRP39 | YML046W | 673.520998 | -0.1782547 | 0.07899253 | -2.2566026 | 0.02403292 |
| RRT8 | YOL048C | 255.627335 | -0.2794681 | 0.12387344 | -2.2560775 | 0.02406578 |
| NUP60 | YAR002W | 938.767841 | -0.1602265 | 0.07103902 | -2.2554714 | 0.02410376 |
| SPR2 | YOR214C | 18.779735 | -0.6274104 | 0.27836753 | -2.253892 | 0.02420296 |
| tF(GAA)M | tF(GAA)M | 9.88535514 | -0.665508 | 0.29550783 | -2.2520823 | 0.02431707 |
| ZIM17 | YNL310C | 404.321781 | -0.2290248 | 0.10175825 | -2.2506749 | 0.02440613 |
| HOP1 | YIL072W | 50.6588056 | -0.477075 | 0.212046 | -2.2498656 | 0.02445748 |
| RMD9 | YGL107C | 834.82605 | -0.1682479 | 0.07479009 | -2.2496019 | 0.02447423 |
| NUP133 | YKR082W | 2249.42255 | 0.12729146 | 0.05658375 | 2.24961153 | 0.02447362 |
| Xu_2009_CUTs26 | Xu_2009_CU | 49.8884854 | 0.45836499 | 0.20380923 | 2.24899034 | 0.02451311 |
| NKP1 | YDR383C | 191.137943 | -0.2855183 | 0.12695706 | -2.2489362 | 0.02451656 |
| YOR032W-A | YOR032W-A | 35.9179857 | 0.53968911 | 0.23996364 | 2.24904533 | 0.02450961 |
| MKC7 | YDR144C | 953.858298 | 0.17331149 | 0.07707057 | 2.24873747 | 0.0245292 |
| ELP2 | YGR200C | 1483.685 | 0.14119342 | 0.06279702 | 2.24840974 | 0.02455007 |
| Xu_2009_SUTs15 | Xu_2009_SU | 26.4451491 | 0.56209916 | 0.25010607 | 2.24744314 | 0.02461172 |
| YGR161W-C | YGR161W-C | 205.355796 | -0.2740696 | 0.12195493 | -2.2473024 | 0.02462071 |
| GPR1 | YDL035C | 1262.24887 | 0.18041878 | 0.0802955 | 2.24693523 | 0.02464417 |
| FEX1 | YOR390W | 200.773387 | 0.26531592 | 0.11809512 | 2.24662899 | 0.02466375 |
| Xu_2009_SUTs81 | Xu_2009_SU | 67.5742608 | 0.41690465 | 0.1856769 | 2.24532324 | 0.02474739 |
| LEM3 | YNL323W | 555.584906 | -0.1879489 | 0.08372062 | -2.2449539 | 0.02477109 |
| SNR77 | snR77 | 29.1034248 | -0.5805207 | 0.2586559 | -2.2443745 | 0.02480832 |
| Xu_2009_CUTs41 | Xu_2009_CU | 59.082377 | 0.43134298 | 0.19226677 | 2.24346095 | 0.02486711 |
| SGM1 | YJR134C | 1014.39468 | 0.15146566 | 0.06751327 | 2.2434948 | 0.02486492 |
| YML045W | YML045W | 16.7478483 | 0.62515 | 0.2786777 | 2.2432724 | 0.02487925 |
| MEX67 | YPL169C | 1405.20328 | -0.1450088 | 0.06468423 | -2.241795 | 0.02497463 |
| Xu_2009_SUTs83 | Xu_2009_SU | 43.1637002 | 0.49977091 | 0.22297203 | 2.24140626 | 0.02499977 |
| tN(GUU)P | tN(GUU)P | 55.7039558 | 0.4530896 | 0.20217308 | 2.24109754 | 0.02501976 |
| Xu_2009_SUTs27 | Xu_2009_SU | 46.9436338 | 0.46676772 | 0.20838998 | 2.23987605 | 0.02509897 |
| Xu_2009_CUTs91 | Xu_2009_CU | 4.70150412 | 0.63766277 | 0.28498121 | 2.23756074 | 0.02524971 |
| SWI4 | YER111C | 1348.22649 | 0.16711134 | 0.07468973 | 2.23740724 | 0.02525974 |
| UBP8 | YMR223W | 536.039661 | 0.1879737 | 0.08404244 | 2.23665206 | 0.02530909 |
| SRP101 | YDR292C | 1314.25276 | 0.14464329 | 0.06467263 | 2.23654568 | 0.02531605 |
| Xu_2009_SUTs52 | Xu_2009_SU | 111.069953 | 0.33888161 | 0.15163209 | 2.23489369 | 0.02542433 |
| Xu_2009_CUTs43 | Xu_2009_CU | 74.455587 | 0.43311636 | 0.19389602 | 2.2337558 | 0.02549915 |
| Xu_2009_CUTs29 | Xu_2009_CU | 17.4574022 | 0.62664858 | 0.28055188 | 2.23362816 | 0.02550755 |
| UBX5 | YDR330W | 874.372489 | -0.1866182 | 0.08358886 | -2.2325723 | 0.02557716 |

| | | | | | | |
|---|---|---|---|---|---|---|
| PTA1 | YAL043C | 1447.72932 | 0.14166266 | 0.06351358 | 2.23043112 | 0.02571884 |
| Xu_2009_CUTs49 | Xu_2009_CU | 27.4788035 | 0.55068837 | 0.2469116 | 2.23030582 | 0.02572715 |
| YOR277C | YOR277C | 40.9558083 | 0.49651086 | 0.22270251 | 2.22948032 | 0.02578196 |
| CSN9 | YDR179C | 99.2952071 | -0.3845024 | 0.17247954 | -2.2292635 | 0.02579638 |
| Xu_2009_SUTs56 | Xu_2009_SU | 11.5262088 | 0.65426891 | 0.29355539 | 2.22877502 | 0.02582888 |
| YOL107W | YOL107W | 310.219362 | -0.2348859 | 0.10541849 | -2.2281278 | 0.025872 |
| Xu_2009_SUTs37 | Xu_2009_SU | 106.27719 | 0.37400428 | 0.16786304 | 2.22803228 | 0.02587836 |
| Xu_2009_CUTs25 | Xu_2009_CU | 3.77066408 | 0.60767164 | 0.27275133 | 2.22793281 | 0.025885 |
| UBP15 | YMR304W | 2447.44303 | 0.12729109 | 0.05717103 | 2.22649621 | 0.02598096 |
| Xu_2009_SUTs76 | Xu_2009_SU | 72.3239515 | 0.4021109 | 0.18071497 | 2.22511121 | 0.02607377 |
| YKL136W | YKL136W | 21.0887079 | -0.594064 | 0.26715213 | -2.2236916 | 0.0261692 |
| ALG1 | YBR110W | 699.239643 | -0.1871049 | 0.08416434 | -2.2230899 | 0.02620974 |
| GPC1 | YGR149W | 1085.82591 | -0.1842852 | 0.0829222 | -2.2223871 | 0.02625716 |
| Xu_2009_CUTs79 | Xu_2009_CU | 6.10812677 | 0.64777966 | 0.29169924 | 2.22071079 | 0.02637056 |
| Xu_2009_CUTs36 | Xu_2009_CU | 19.0446568 | 0.62381589 | 0.28097794 | 2.22015967 | 0.02640793 |
| HSP31 | YDR533C | 1530.8313 | -0.1629228 | 0.0734255 | -2.2188863 | 0.02649446 |
| RNH1 | YMR234W | 310.595592 | -0.2339272 | 0.10549902 | -2.2173399 | 0.02659987 |
| CCE1 | YKL011C | 380.373066 | -0.2063635 | 0.0930979 | -2.2166285 | 0.02664849 |
| YAL034C-B | YAL034C-B | 1.80372369 | 0.50270221 | 0.22682014 | 2.21630321 | 0.02667074 |
| Xu_2009_CUTs78 | Xu_2009_CU | 40.6965145 | 0.51547192 | 0.23268209 | 2.21534852 | 0.02673615 |
| ARP4 | YJL081C | 756.592788 | -0.1765994 | 0.07977298 | -2.2137751 | 0.02684425 |
| Xu_2009_CUTs26 | Xu_2009_CU | 14.2810925 | 0.63659882 | 0.28760521 | 2.21344674 | 0.02686686 |
| Xu_2009_CUTs92 | Xu_2009_CU | 19.1082029 | 0.5897755 | 0.26650887 | 2.21296759 | 0.02689988 |
| MDL2 | YPL270W | 961.42841 | -0.1474848 | 0.06667998 | -2.21183 | 0.02697841 |
| COG5 | YNL051W | 357.365346 | -0.2158559 | 0.09765269 | -2.2104449 | 0.0270743 |
| Xu_2009_CUTs55 | Xu_2009_CU | 2.57184803 | -0.5449483 | 0.24682003 | -2.2078771 | 0.02725284 |
| YML003W | YML003W | 222.467716 | 0.2647445 | 0.11992606 | 2.20756442 | 0.02727465 |
| XKS1 | YGR194C | 439.572865 | -0.2386213 | 0.10810116 | -2.2073891 | 0.02728689 |
| Xu_2009_CUTs40 | Xu_2009_CU | 98.3597327 | 0.35912907 | 0.16275879 | 2.20651114 | 0.02734823 |
| ARF3 | YOR094W | 187.109017 | -0.2775693 | 0.12587243 | -2.2051638 | 0.0274426 |
| AIM11 | YER093C-A | 160.074972 | -0.2985031 | 0.13538326 | -2.2048744 | 0.02746291 |
| HOP2 | YGL033W | 31.4215843 | 0.54450488 | 0.24703774 | 2.20413639 | 0.02751475 |
| GCV1 | YDR019C | 1378.04374 | 0.17605287 | 0.07996584 | 2.20160082 | 0.02769352 |
| MRX7 | YNL211C | 124.526206 | -0.3283308 | 0.14917576 | -2.200966 | 0.02773843 |
| YJL015C | YJL015C | 4.16815554 | 0.59990216 | 0.27300092 | 2.19743639 | 0.02798929 |
| TMA23 | YMR269W | 42.3829622 | 0.49496748 | 0.22541085 | 2.1958459 | 0.02810297 |
| LDS1 | YAL018C | 11.7893963 | -0.6402025 | 0.29178521 | -2.1940884 | 0.02822906 |
| CWH41 | YGL027C | 1432.19639 | -0.1380462 | 0.06291876 | -2.1940388 | 0.02823262 |
| NRG2 | YBR066C | 360.326 | -0.2217398 | 0.10108952 | -2.1934994 | 0.02827142 |
| YGR125W | YGR125W | 1444.14529 | 0.14858204 | 0.06774399 | 2.19328735 | 0.02828668 |
| MET7 | YOR241W | 747.8809 | 0.16269235 | 0.07420736 | 2.19240174 | 0.02835051 |

| | | | | | | |
|---|---|---|---|---|---|---|
| Xu_2009_CUTs72 | Xu_2009_CU | 32.4219936 | 0.52746119 | 0.24060099 | 2.19226528 | 0.02836036 |
| Xu_2009_SUTs71 | Xu_2009_SU | 73.7517623 | 0.39744314 | 0.18136344 | 2.19141818 | 0.02842154 |
| TOS2 | YGR221C | 335.127637 | -0.2195754 | 0.10019653 | -2.1914476 | 0.02841942 |
| Xu_2009_SUTs35 | Xu_2009_SU | 183.232184 | 0.27767509 | 0.12677993 | 2.19021335 | 0.02850877 |
| SCT1 | YBL011W | 2243.85351 | 0.12722302 | 0.05808152 | 2.19042142 | 0.02849369 |
| UBP13 | YBL067C | 1079.96753 | -0.1669311 | 0.07621063 | -2.1903917 | 0.02849584 |
| YBR209W | YBR209W | 12.8309511 | 0.62763312 | 0.28655603 | 2.19026315 | 0.02850516 |
| YGR038C-B | YGR038C-B | 262.024675 | 0.27200167 | 0.1241976 | 2.1900719 | 0.02851902 |
| Xu_2009_CUTs89 | Xu_2009_CU | 10.1187467 | 0.6488262 | 0.29635761 | 2.18933536 | 0.02857247 |
| THI74 | YDR438W | 178.8781 | 0.28218293 | 0.12895038 | 2.18830636 | 0.02864729 |
| MCO6 | YJL127C-B | 241.668932 | -0.2497884 | 0.11417586 | -2.1877511 | 0.02868773 |
| SIP5 | YMR140W | 661.854846 | -0.1948541 | 0.08913442 | -2.1860698 | 0.0288105 |
| FRM2 | YCL026C-A | 35.8365262 | -0.5152394 | 0.23570331 | -2.1859659 | 0.0288181 |
| NUT2 | YPR168W | 111.202894 | 0.34328461 | 0.15706374 | 2.18563886 | 0.02884204 |
| Xu_2009_CUTs67 | Xu_2009_CU | 16.3530285 | 0.61261058 | 0.28033324 | 2.18529415 | 0.02886729 |
| Xu_2009_CUTs88 | Xu_2009_CU | 43.7549061 | 0.47902775 | 0.21940243 | 2.18332924 | 0.02901157 |
| Xu_2009_SUTs61 | Xu_2009_SU | 110.628948 | 0.33324915 | 0.1527345 | 2.1818852 | 0.02911801 |
| JAC1 | YGL018C | 267.711974 | -0.2429129 | 0.11132903 | -2.1819364 | 0.02911423 |
| DDR2 | YOL052C-A | 898.299772 | 0.23021071 | 0.10553176 | 2.18143532 | 0.02915123 |
| Xu_2009_CUTs52 | Xu_2009_CU | 26.1865224 | 0.55498343 | 0.25444599 | 2.18114429 | 0.02917275 |
| Xu_2009_SUTs70 | Xu_2009_SU | 125.015948 | 0.34101199 | 0.15637257 | 2.1807661 | 0.02920072 |
| PAC11 | YDR488C | 371.978954 | 0.21940667 | 0.10067162 | 2.17942932 | 0.02929979 |
| YGL063C-A | YGL063C-A | 7.74523693 | 0.64419828 | 0.29569168 | 2.17861483 | 0.02936029 |
| GUD1 | YDL238C | 253.568728 | 0.27920808 | 0.12828556 | 2.17645756 | 0.02952106 |
| SKI3 | YPR189W | 1318.75587 | 0.13794719 | 0.06337957 | 2.1765245 | 0.02951606 |
| Xu_2009_SUTs65 | Xu_2009_SU | 248.117725 | 0.24616387 | 0.11310895 | 2.17634301 | 0.02952962 |
| YLR361C-A | YLR361C-A | 74.9401072 | -0.4048492 | 0.18603183 | -2.1762363 | 0.0295376 |
| MUK1 | YPL070W | 1268.99984 | 0.14987046 | 0.06889643 | 2.17530081 | 0.02960758 |
| SKI7 | YOR076C | 600.027069 | 0.17476043 | 0.08035457 | 2.1748661 | 0.02964015 |
| YPL071C | YPL071C | 283.81213 | -0.2316525 | 0.10651919 | -2.1747489 | 0.02964894 |
| YKL070W | YKL070W | 25.4295082 | -0.5611493 | 0.2580637 | -2.1744602 | 0.02967059 |
| Xu_2009_SUTs19 | Xu_2009_SU | 83.25135 | 0.37583051 | 0.17290991 | 2.17356251 | 0.02973801 |
| YOR318C | YOR318C | 69.8743795 | 0.41109223 | 0.18914732 | 2.17339707 | 0.02975045 |
| GCD14 | YJL125C | 837.366947 | -0.1674436 | 0.07709008 | -2.1720511 | 0.02985181 |
| Xu_2009_CUTs42 | Xu_2009_CU | 28.3211154 | 0.55831815 | 0.25720017 | 2.1707534 | 0.02994982 |
| Xu_2009_SUTs84 | Xu_2009_SU | 492.662064 | -0.1989427 | 0.09168166 | -2.1699287 | 0.03001225 |
| SIS2 | YKR072C | 2700.55758 | 0.14716734 | 0.06783934 | 2.169351 | 0.03005605 |
| DCN1 | YLR128W | 190.621957 | -0.2918972 | 0.13465255 | -2.1677809 | 0.03017536 |
| MCM4 | YPR019W | 1736.17392 | 0.13402082 | 0.06184253 | 2.1671303 | 0.03022492 |
| YJL077W-A | YJL077W-A | 85.0712599 | 0.36907098 | 0.17035296 | 2.16650756 | 0.03027242 |
| BUD19 | YJL188C | 2.48435418 | -0.5390481 | 0.24890429 | -2.1656844 | 0.03033531 |

| | | | | | | |
|---|---|---|---|---|---|---|
| Xu_2009_SUTs86 | Xu_2009_SU | 26.7039577 | 0.56178818 | 0.25948511 | 2.16501128 | 0.03038682 |
| YPL185W | YPL185W | 11.9296326 | 0.63041834 | 0.29122594 | 2.16470531 | 0.03041026 |
| YML039W | YML039W | 7.40097236 | 0.64045165 | 0.29593049 | 2.16419627 | 0.03044929 |
| tR(UCU)G3 | tR(UCU)G3 | 9.32973403 | -0.641292 | 0.29655481 | -2.1624739 | 0.03058167 |
| GBP2 | YCL011C | 2489.49818 | 0.1358391 | 0.06282228 | 2.16227595 | 0.03059691 |
| Xu_2009_CUTs65 | Xu_2009_CU | 133.974612 | 0.31946961 | 0.14788827 | 2.16020935 | 0.03075647 |
| YIL100W | YIL100W | 16.8844239 | 0.60313407 | 0.2791925 | 2.16028033 | 0.03075098 |
| SET2 | YJL168C | 1107.41844 | -0.1600944 | 0.07416383 | -2.1586583 | 0.03087668 |
| Xu_2009_SUTs11 | Xu_2009_SU | 42.3117714 | 0.46450302 | 0.21526627 | 2.15780677 | 0.03094286 |
| YIL152W | YIL152W | 220.332095 | -0.2650062 | 0.1228576 | -2.1570195 | 0.03100414 |
| Xu_2009_SUTs11 | Xu_2009_SU | 4.17982802 | -0.6063328 | 0.28130695 | -2.1554135 | 0.03112949 |
| SRP40 | YKR092C | 1811.39097 | -0.150024 | 0.06963614 | -2.1543982 | 0.03120896 |
| DOT6 | YER088C | 4421.26834 | 0.11953369 | 0.05548594 | 2.15430592 | 0.03121619 |
| Xu_2009_SUTs28 | Xu_2009_SU | 1.0004197 | 0.3897349 | 0.18093528 | 2.15400175 | 0.03124003 |
| Xu_2009_SUTs13 | Xu_2009_SU | 178.670566 | 0.27632251 | 0.12830955 | 2.15356147 | 0.03127458 |
| PET127 | YOR017W | 925.623058 | -0.1716838 | 0.07974852 | -2.1528151 | 0.03133321 |
| DOT1 | YDR440W | 544.183735 | 0.18473594 | 0.08585282 | 2.15177473 | 0.0314151 |
| GWT1 | YJL091C | 474.083239 | -0.1869152 | 0.08686061 | -2.1518985 | 0.03140535 |
| HAL1 | YPR005C | 362.781719 | -0.2138593 | 0.0993866 | -2.151792 | 0.03141374 |
| Xu_2009_SUTs83 | Xu_2009_SU | 76.7339554 | 0.40478126 | 0.18813843 | 2.15150759 | 0.03143616 |
| TCO89 | YPL180W | 823.043188 | 0.1576327 | 0.07326345 | 2.15158736 | 0.03142987 |
| YAL056C-A | YAL056C-A | 4.70724291 | 0.61222213 | 0.28459424 | 2.15121055 | 0.03145958 |
| VFA1 | YER128W | 174.166072 | -0.2810646 | 0.13075294 | -2.149585 | 0.03158805 |
| SDD4 | YPR022C | 1436.67724 | 0.14615061 | 0.06800306 | 2.14917703 | 0.03162037 |
| SPP381 | YBR152W | 527.025885 | 0.2139435 | 0.09955112 | 2.14908182 | 0.03162792 |
| LDS2 | YOL047C | 137.797985 | -0.3065035 | 0.14275054 | -2.1471269 | 0.03178318 |
| Xu_2009_CUTs28 | Xu_2009_CU | 7.59774827 | 0.63567768 | 0.29610435 | 2.1468029 | 0.03180898 |
| PLC1 | YPL268W | 1020.28641 | 0.17122794 | 0.07976297 | 2.14670962 | 0.03181641 |
| TRM5 | YHR070W | 2637.24526 | 0.17085093 | 0.07959306 | 2.14655565 | 0.03182867 |
| APA2 | YDR530C | 277.936396 | -0.2272691 | 0.10594731 | -2.1451147 | 0.03194368 |
| IKI3 | YLR384C | 2952.76133 | 0.12956325 | 0.060407 | 2.14483822 | 0.03196578 |
| FOB1 | YDR110W | 486.116656 | 0.20878109 | 0.09734768 | 2.14469508 | 0.03197723 |
| GMH1 | YKR030W | 320.889223 | -0.230604 | 0.10754258 | -2.1443044 | 0.0320085 |
| RMD5 | YDR255C | 347.00848 | 0.22202156 | 0.10360734 | 2.14291348 | 0.03212005 |
| YCR081C-A | YCR081C-A | 9.44262814 | 0.63205691 | 0.29516482 | 2.14136937 | 0.03224426 |
| Xu_2009_SUTs42 | Xu_2009_SU | 2.86013376 | 0.55878433 | 0.26096327 | 2.14123744 | 0.03225489 |
| Xu_2009_SUTs15 | Xu_2009_SU | 1.72682492 | 0.48429213 | 0.22629645 | 2.14007834 | 0.03234844 |
| CDC10 | YCR002C | 886.385523 | -0.1466978 | 0.06857876 | -2.1391139 | 0.03242645 |
| AIR2 | YDL175C | 358.982822 | 0.21025675 | 0.09830031 | 2.13892258 | 0.03244194 |
| ATG29 | YPL166W | 79.1893486 | -0.3727631 | 0.17429954 | -2.1386352 | 0.03246523 |
| SKM1 | YOL113W | 936.402302 | 0.17835608 | 0.08343126 | 2.13776092 | 0.03253615 |

| | | | | | | |
|---|---|---|---|---|---|---|
| INO4 | YOL108C | 252.727901 | -0.239383 | 0.11206523 | -2.1361041 | 0.03267093 |
| Xu_2009_SUTs53 | Xu_2009_SU | 124.950361 | 0.30812149 | 0.14430034 | 2.13527908 | 0.03273822 |
| NOB1 | YOR056C | 1195.50207 | 0.16145029 | 0.07565443 | 2.13404939 | 0.03283874 |
| SNL1 | YIL016W | 236.331117 | -0.2507638 | 0.11755107 | -2.1332324 | 0.03290567 |
| Xu_2009_CUTs17 | Xu_2009_CU | 29.6864848 | 0.52333159 | 0.24539353 | 2.13262183 | 0.03295576 |
| YCL002C | YCL002C | 313.209818 | -0.2288688 | 0.10731509 | -2.1326802 | 0.03295097 |
| YRF1-7 | YPL283C | 4.32449491 | 0.60304013 | 0.28283374 | 2.13213647 | 0.03299563 |
| NUP120 | YKL057C | 1460.42133 | -0.1464566 | 0.06872095 | -2.1311785 | 0.03307444 |
| GPI8 | YDR331W | 525.398949 | -0.1954486 | 0.09174783 | -2.1302808 | 0.03314844 |
| FAL1 | YDR021W | 452.914465 | 0.19851992 | 0.09320928 | 2.12983007 | 0.03318565 |
| Xu_2009_CUTs39 | Xu_2009_CU | 20.6966878 | 0.57599363 | 0.27052695 | 2.12915434 | 0.03324149 |
| Xu_2009_SUTs33 | Xu_2009_SU | 32.6661953 | -0.5033715 | 0.23645619 | -2.1288153 | 0.03326954 |
| CTF13 | YMR094W | 286.57089 | 0.22721438 | 0.10676611 | 2.1281508 | 0.03332458 |
| GPI18 | YBR004C | 341.64036 | -0.2426146 | 0.11402544 | -2.1277236 | 0.03336001 |
| SNR31 | snR31 | 887.805592 | -0.323033 | 0.15185106 | -2.1273013 | 0.03339506 |
| EFM2 | YBR271W | 408.096515 | -0.2114085 | 0.09938348 | -2.1271999 | 0.03340348 |
| tX(XXX)D | tX(XXX)D | 16.0838823 | -0.6019414 | 0.28321657 | -2.125375 | 0.03355533 |
| RMP1 | YLR145W | 201.373485 | 0.26773304 | 0.12609429 | 2.12327655 | 0.03373068 |
| ATG32 | YIL146C | 394.078262 | 0.21363609 | 0.10063363 | 2.12290938 | 0.03376145 |
| QRI7 | YDL104C | 112.872674 | 0.33945033 | 0.15991767 | 2.1226568 | 0.03378262 |
| Xu_2009_CUTs46 | Xu_2009_CU | 48.426642 | 0.43871555 | 0.20677541 | 2.12170077 | 0.03386287 |
| YIA6 | YIL006W | 311.039701 | -0.2180831 | 0.10281449 | -2.1211324 | 0.03391066 |
| Xu_2009_CUTs33 | Xu_2009_CU | 133.572164 | -0.3033589 | 0.14302876 | -2.1209642 | 0.03392481 |
| ARG82 | YDR173C | 326.962658 | 0.24184349 | 0.11404011 | 2.12068791 | 0.03394807 |
| UCC1 | YLR224W | 1188.30307 | -0.1457013 | 0.06873814 | -2.1196574 | 0.03403495 |
| ARA2 | YMR041C | 271.103405 | 0.26114868 | 0.12321429 | 2.11946745 | 0.03405098 |
| NAM8 | YHR086W | 700.984245 | -0.179847 | 0.08493703 | -2.1174158 | 0.03422458 |
| tT(AGU)I2 | tT(AGU)I2 | 5.8541418 | -0.6110847 | 0.28865338 | -2.1170192 | 0.03425822 |
| NAG1 | YGR031C-A | 17.5683966 | 0.58591851 | 0.27676296 | 2.11704096 | 0.03425637 |
| YMR310C | YMR310C | 902.908688 | 0.15940655 | 0.07533496 | 2.11597032 | 0.03434733 |
| Xu_2009_CUTs40 | Xu_2009_CU | 19.3706475 | 0.58158506 | 0.27491107 | 2.11553892 | 0.03438404 |
| BMT2 | YBR141C | 390.0257 | 0.23301001 | 0.11022915 | 2.11386914 | 0.03452645 |
| KCC4 | YCL024W | 504.337947 | 0.18994722 | 0.0898545 | 2.11394229 | 0.0345202 |
| Xu_2009_CUTs51 | Xu_2009_CU | 25.3063587 | 0.54171308 | 0.25628363 | 2.11372484 | 0.03453878 |
| YPL119C-A | YPL119C-A | 30.7988276 | -0.5235344 | 0.24779247 | -2.112794 | 0.03461841 |
| YER138C | YER138C | 16.142312 | 0.60616528 | 0.28704998 | 2.11170638 | 0.03471164 |
| ABM1 | YJR108W | 92.5683372 | 0.36136148 | 0.17120881 | 2.1106477 | 0.0348026 |
| YDR131C | YDR131C | 348.293875 | 0.21690752 | 0.10279233 | 2.11015272 | 0.0348452 |
| Xu_2009_SUTs62 | Xu_2009_SU | 42.6757258 | 0.46930468 | 0.22244932 | 2.10971509 | 0.0348829 |
| YPR150W | YPR150W | 58.9687916 | 0.43396276 | 0.20579384 | 2.10872573 | 0.03496826 |
| PUG1 | YER185W | 286.750339 | 0.27351389 | 0.12989191 | 2.10570379 | 0.03523009 |

| | | | | | | |
|---|---|---|---|---|---|---|
| YLR374C | YLR374C | 25.792336 | 0.5515818 | 0.26194314 | 2.10573099 | 0.03522773 |
| INP54 | YOL065C | 169.09061 | -0.2836298 | 0.13469668 | -2.1056923 | 0.03523109 |
| ATG23 | YLR431C | 380.510296 | 0.19666585 | 0.09341931 | 2.10519474 | 0.03527436 |
| Xu_2009_CUTs85 | Xu_2009_CU | 7.49748585 | 0.62046216 | 0.29493026 | 2.10375889 | 0.03539948 |
| Xu_2009_CUTs90 | Xu_2009_CU | 42.7396529 | 0.46960878 | 0.22349411 | 2.10121324 | 0.03562225 |
| GPD2 | YOL059W | 1694.58238 | -0.1309019 | 0.06230173 | -2.1010951 | 0.03563262 |
| SWD2 | YKL018W | 448.179715 | -0.18362 | 0.08740255 | -2.1008535 | 0.03565383 |
| Xu_2009_CUTs21 | Xu_2009_CU | 4.77701721 | 0.60609186 | 0.28853244 | 2.10060213 | 0.03567591 |
| UGA3 | YDL170W | 532.410869 | -0.1810169 | 0.08617615 | -2.1005452 | 0.03568091 |
| MUP1 | YGR055W | 1190.71674 | 0.14214586 | 0.06767188 | 2.10051578 | 0.03568349 |
| YNL097C-B | YNL097C-B | 15.2191812 | -0.612697 | 0.29187489 | -2.0991767 | 0.03580133 |
| ARP1 | YHR129C | 166.772172 | -0.2756905 | 0.13137808 | -2.0984516 | 0.03586527 |
| YNR061C | YNR061C | 374.936145 | -0.21371 | 0.10186271 | -2.09802 | 0.03590338 |
| YGR137W | YGR137W | 72.0368988 | -0.3750056 | 0.17882675 | -2.0970332 | 0.03599064 |
| BUB1 | YGR188C | 366.884252 | 0.20075833 | 0.09579424 | 2.09572436 | 0.03610665 |
| SEA4 | YBL104C | 825.548351 | 0.14648724 | 0.06995322 | 2.09407428 | 0.03625336 |
| Xu_2009_SUTs61 | Xu_2009_SU | 55.2107748 | -0.4288501 | 0.20480966 | -2.0938957 | 0.03626927 |
| YEL076C-A | YEL076C-A | 2.77495277 | 0.54254348 | 0.25921758 | 2.09300413 | 0.03634879 |
| PIH1 | YHR034C | 241.976563 | -0.2450243 | 0.11711272 | -2.0922095 | 0.03641978 |
| Xu_2009_SUTs59 | Xu_2009_SU | 12.3290586 | 0.61089274 | 0.29201944 | 2.09195912 | 0.03644217 |
| GRC3 | YLL035W | 953.437561 | 0.1745289 | 0.08343535 | 2.09178607 | 0.03645766 |
| SNR7-L | snR7-L | 6.47739059 | -0.6149022 | 0.29404964 | -2.0911511 | 0.03651452 |
| SAL1 | YNL083W | 223.809001 | 0.24539255 | 0.11734705 | 2.09116929 | 0.03651289 |
| IOC4 | YMR044W | 515.601044 | 0.20055317 | 0.09593184 | 2.09057969 | 0.03656576 |
| KEL3 | YPL263C | 1565.0151 | 0.13127887 | 0.06281795 | 2.08983064 | 0.03663302 |
| Xu_2009_CUTs67 | Xu_2009_CU | 15.5183761 | 0.59450476 | 0.28460408 | 2.08888348 | 0.03671821 |
| Xu_2009_SUTs14 | Xu_2009_SU | 44.1637892 | 0.4767395 | 0.2282253 | 2.08889856 | 0.03671686 |
| BRE4 | YDL231C | 1217.72359 | 0.14322468 | 0.06861573 | 2.08734472 | 0.03685699 |
| Xu_2009_CUTs27 | Xu_2009_CU | 17.6279084 | 0.57653387 | 0.2763886 | 2.08595388 | 0.0369828 |
| YER053C-A | YER053C-A | 75.9227344 | -0.4318311 | 0.20702262 | -2.0859127 | 0.03698653 |
| YPR203W | YPR203W | 2.36530201 | 0.52450664 | 0.25152702 | 2.08528944 | 0.03704304 |
| HOF1 | YMR032W | 653.178519 | 0.18017434 | 0.08641489 | 2.08499179 | 0.03707005 |
| DEG1 | YFL001W | 501.61506 | -0.1891557 | 0.0908708 | -2.0815899 | 0.03737994 |
| LSM12 | YHR121W | 490.6888 | -0.2069635 | 0.09944507 | -2.0811843 | 0.03741704 |
| Xu_2009_CUTs30 | Xu_2009_CU | 40.1667344 | 0.47091607 | 0.22639359 | 2.08007684 | 0.03751849 |
| RGD2 | YFL047W | 956.750551 | 0.17169845 | 0.08261264 | 2.07835564 | 0.03767661 |
| YGR064W | YGR064W | 39.331263 | 0.47283234 | 0.22750177 | 2.07836777 | 0.0376755 |
| GAT3 | YLR013W | 2.61230786 | 0.51819169 | 0.24934625 | 2.07820121 | 0.03769083 |
| JIP5 | YPR169W | 1160.32192 | 0.14839711 | 0.07143758 | 2.07729759 | 0.0377741 |
| FIN1 | YDR130C | 294.825353 | 0.23446246 | 0.11289891 | 2.07674699 | 0.03782492 |
| Xu_2009_CUTs64 | Xu_2009_CU | 91.1454524 | 0.34709657 | 0.16728289 | 2.07490776 | 0.03799508 |

| | | | | | | |
|---|---|---|---|---|---|---|
| LEE1 | YPL054W | 117.069568 | 0.31944225 | 0.15398836 | 2.07445709 | 0.03803688 |
| ROD1 | YOR018W | 704.758799 | 0.18785388 | 0.09059083 | 2.07365225 | 0.03811162 |
| RDS3 | YPR094W | 133.743361 | -0.3237696 | 0.15631068 | -2.0713208 | 0.03832882 |
| ALR1 | YOL130W | 1905.81775 | 0.14753266 | 0.07123557 | 2.07105317 | 0.03835383 |
| Xu_2009_SUTs35 | Xu_2009_SU | 30.2867598 | -0.5110308 | 0.24684544 | -2.070246 | 0.03842931 |
| Xu_2009_SUTs73 | Xu_2009_SU | 82.3643716 | -0.3617251 | 0.1747667 | -2.06976 | 0.03847483 |
| KSP1 | YHR082C | 3560.92269 | 0.11570628 | 0.05591261 | 2.06941306 | 0.03850734 |
| DCI1 | YOR180C | 185.395479 | 0.27552418 | 0.13328956 | 2.06710997 | 0.03872379 |
| Xu_2009_CUTs35 | Xu_2009_CU | 16.1514303 | 0.57744046 | 0.27945143 | 2.06633564 | 0.0387968 |
| IDS2 | YJL146W | 447.01978 | -0.185939 | 0.09005734 | -2.0646738 | 0.03895387 |
| Xu_2009_CUTs28 | Xu_2009_CU | 59.6582477 | 0.40746528 | 0.19750263 | 2.06308792 | 0.03910428 |
| SCM4 | YGR049W | 605.344802 | 0.16876014 | 0.0818099 | 2.06283274 | 0.03912852 |
| Xu_2009_CUTs90 | Xu_2009_CU | 53.3142485 | 0.41641587 | 0.20197934 | 2.06167555 | 0.03923864 |
| MDJ2 | YNL328C | 102.489949 | 0.334581 | 0.16228913 | 2.06163533 | 0.03924247 |
| PDR16 | YNL231C | 1937.97087 | 0.14798905 | 0.07180946 | 2.06085735 | 0.03931665 |
| BRX1 | YOL077C | 1993.59575 | -0.1477302 | 0.07169663 | -2.0604902 | 0.0393517 |
| GTB1 | YDR221W | 874.629604 | 0.1544583 | 0.07497507 | 2.06012881 | 0.03938623 |
| RSM28 | YDR494W | 868.404328 | -0.152726 | 0.07415502 | -2.0595507 | 0.03944151 |
| CCA1 | YER168C | 708.069596 | -0.1612437 | 0.07829782 | -2.0593639 | 0.03945939 |
| DCP2 | YNL118C | 1557.76565 | 0.13107114 | 0.06371578 | 2.05712204 | 0.03967449 |
| HOS3 | YPL116W | 817.591834 | 0.19344356 | 0.0940368 | 2.05710479 | 0.03967615 |
| CYC2 | YOR037W | 395.944187 | -0.2120454 | 0.10310989 | -2.0564987 | 0.03973447 |
| YAL059C-A | YAL059C-A | 10.9029473 | 0.60116887 | 0.29251005 | 2.05520754 | 0.03985896 |
| Xu_2009_SUTs28 | Xu_2009_SU | 632.720868 | 0.16759334 | 0.08156005 | 2.05484602 | 0.03989388 |
| GAR1 | YHR089C | 784.032668 | -0.1854793 | 0.09030307 | -2.0539645 | 0.03997913 |
| Xu_2009_CUTs57 | Xu_2009_CU | 36.4509041 | 0.46109256 | 0.22452846 | 2.05360403 | 0.04001403 |
| PMT5 | YDL093W | 542.794614 | -0.2078683 | 0.10122275 | -2.0535724 | 0.0400171 |
| Xu_2009_CUTs80 | Xu_2009_CU | 20.6757328 | 0.54132408 | 0.26372429 | 2.05261367 | 0.04011006 |
| YGR018C | YGR018C | 47.5450661 | 0.42761232 | 0.20836844 | 2.0521933 | 0.04015088 |
| YDR154C | YDR154C | 15.1953365 | -0.584999 | 0.28526278 | -2.0507371 | 0.04029256 |
| GUP2 | YPL189W | 82.3065307 | 0.34846958 | 0.17000847 | 2.04971895 | 0.04039186 |
| PDR11 | YIL013C | 100.44887 | -0.3302111 | 0.16114205 | -2.0491928 | 0.04044327 |
| EDE1 | YBL047C | 5842.14414 | -0.0953812 | 0.04661674 | -2.0460713 | 0.04074936 |
| PAU11 | YGL261C | 2.76040454 | 0.52702516 | 0.25762526 | 2.04570452 | 0.04078545 |
| BET4 | YJL031C | 400.002284 | -0.2011966 | 0.09843972 | -2.043856 | 0.04096777 |
| YMR160W | YMR160W | 464.335306 | -0.2165458 | 0.10597161 | -2.0434323 | 0.04100967 |
| RGA2 | YDR379W | 926.083656 | 0.14979527 | 0.07336101 | 2.04189209 | 0.04116224 |
| YKR033C | YKR033C | 5.30740563 | 0.59542125 | 0.29170903 | 2.04114781 | 0.04123614 |
| DOM34 | YNL001W | 321.408515 | -0.2111293 | 0.10347937 | -2.0403032 | 0.04132014 |
| MIX14 | YDR031W | 124.373544 | -0.3088068 | 0.15137062 | -2.040071 | 0.04134325 |
| Xu_2009_SUTs24 | Xu_2009_SU | 70.376707 | -0.3871146 | 0.1898373 | -2.0391912 | 0.04143095 |

| | | | | | | |
|---|---|---|---|---|---|---|
| Xu_2009_SUTs1 | Xu_2009_SU | 45.5197228 | 0.43822145 | 0.21499542 | 2.03828272 | 0.04152167 |
| YLR112W | YLR112W | 210.708169 | -0.2441013 | 0.11976291 | -2.0382048 | 0.04152945 |
| Xu_2009_CUTs52 | Xu_2009_CU | 32.1491611 | 0.48684504 | 0.2389268 | 2.03763262 | 0.04158669 |
| DPH2 | YKL191W | 1209.13034 | -0.1656998 | 0.08137483 | -2.0362538 | 0.04172488 |
| Xu_2009_SUTs42 | Xu_2009_SU | 16.0353638 | 0.56964885 | 0.27981571 | 2.03580009 | 0.04177043 |
| DAL4 | YIR028W | 42.5178286 | -0.4354467 | 0.21391998 | -2.0355588 | 0.04179468 |
| PWR1 | PWR1 | 8.57927348 | 0.60313411 | 0.29633779 | 2.0352926 | 0.04182144 |
| Xu_2009_CUTs26 | Xu_2009_CU | 7.2250675 | 0.59814535 | 0.29389873 | 2.03520904 | 0.04182984 |
| LTO1 | YNL260C | 391.19469 | -0.2015047 | 0.09901322 | -2.035129 | 0.04183789 |
| ULA1 | YPL003W | 306.16535 | -0.2108015 | 0.10356936 | -2.0353658 | 0.04181408 |
| ROQ1 | YJL144W | 291.279591 | 0.22954984 | 0.11290435 | 2.03313547 | 0.04203884 |
| YMR196W | YMR196W | 3507.99213 | 0.1247807 | 0.06139059 | 2.0325703 | 0.04209595 |
| tE(UUC)J | tE(UUC)J | 12.5834759 | -0.5923384 | 0.29148718 | -2.032125 | 0.042141 |
| Xu_2009_SUTs74 | Xu_2009_SU | 287.983933 | 0.23007283 | 0.11324236 | 2.03168528 | 0.04218553 |
| CDC55 | YGL190C | 1660.67229 | -0.1212525 | 0.0596831 | -2.0316047 | 0.04219369 |
| Xu_2009_CUTs62 | Xu_2009_CU | 19.7940322 | 0.54751044 | 0.26952608 | 2.03138207 | 0.04221625 |
| YMR085W | YMR085W | 94.2697943 | 0.34969555 | 0.17218453 | 2.03093482 | 0.04226161 |
| Xu_2009_SUTs41 | Xu_2009_SU | 108.920006 | 0.30790165 | 0.15163271 | 2.03057547 | 0.04229808 |
| UME1 | YPL139C | 329.549894 | -0.2083279 | 0.10262214 | -2.0300486 | 0.0423516 |
| FAR8 | YMR029C | 572.364242 | -0.1825709 | 0.09000396 | -2.0284759 | 0.0425117 |
| DRS2 | YAL026C | 1805.14245 | 0.12321128 | 0.06074757 | 2.02825044 | 0.04253469 |
| IZH2 | YOL002C | 1085.3901 | -0.1924462 | 0.09489643 | -2.0279608 | 0.04256424 |
| YDR261C-D | YDR261C-D | 392.653816 | 0.21494096 | 0.1060537 | 2.02671806 | 0.04269125 |
| MSS2 | YDL107W | 211.137478 | -0.2692947 | 0.13310681 | -2.0231476 | 0.04305793 |
| Xu_2009_CUTs72 | Xu_2009_CU | 49.8151076 | 0.45628423 | 0.22565165 | 2.02207351 | 0.04316876 |
| Xu_2009_CUTs80 | Xu_2009_CU | 23.5767882 | 0.52850056 | 0.26141695 | 2.02167667 | 0.04320977 |
| NUP42 | YDR192C | 295.854771 | -0.2154339 | 0.10657016 | -2.0215222 | 0.04322574 |
| Xu_2009_CUTs28 | Xu_2009_CU | 11.2174028 | 0.59049823 | 0.2922408 | 2.02058791 | 0.04332244 |
| YKR078W | YKR078W | 481.220706 | 0.1760679 | 0.08719635 | 2.01921169 | 0.04346522 |
| tG(GCC)G2 | tG(GCC)G2 | 10.3946204 | -0.5985514 | 0.29660949 | -2.0179779 | 0.04359356 |
| YPR099C | YPR099C | 31.8783197 | 0.48932301 | 0.24248702 | 2.01793486 | 0.04359805 |
| YOR364W | YOR364W | 39.0435226 | 0.47206849 | 0.23405092 | 2.01694782 | 0.04370096 |
| Xu_2009_CUTs50 | Xu_2009_CU | 21.8286792 | 0.5281992 | 0.26189356 | 2.01684683 | 0.0437115 |
| Xu_2009_SUTs71 | Xu_2009_SU | 98.9781691 | 0.3549434 | 0.17600897 | 2.01662106 | 0.04373507 |
| Xu_2009_CUTs21 | Xu_2009_CU | 7.66579862 | 0.59686194 | 0.29602647 | 2.01624518 | 0.04377434 |
| VPS21 | YOR089C | 1941.46089 | 0.15621229 | 0.07747642 | 2.01625593 | 0.04377322 |
| ADI1 | YMR009W | 339.3606 | 0.20092685 | 0.09966289 | 2.01606488 | 0.04379319 |
| Xu_2009_CUTs57 | Xu_2009_CU | 4.15013127 | 0.55979861 | 0.27784272 | 2.01480398 | 0.04392519 |
| TCB2 | YNL087W | 1983.64934 | -0.1191245 | 0.05912754 | -2.0147046 | 0.04393561 |
| TMN3 | YER113C | 717.824773 | -0.1563702 | 0.07770032 | -2.0124785 | 0.04416952 |
| NNR2 | YKL151C | 1134.05411 | -0.1758585 | 0.08742004 | -2.0116498 | 0.04425687 |

| | | | | | | |
|---|---|---|---|---|---|---|
| SPT14 | YPL175W | 297.224319 | 0.21150305 | 0.10520656 | 2.01035999 | 0.0443931 |
| Xu_2009_CUTs51 | Xu_2009_CU | 6.83398585 | 0.59246919 | 0.29483939 | 2.00946416 | 0.04448793 |
| Xu_2009_CUTs68 | Xu_2009_CU | 15.8486932 | 0.5571636 | 0.27739458 | 2.00855983 | 0.04458383 |
| SSL2 | YIL143C | 1352.30919 | 0.1255137 | 0.06248729 | 2.00862781 | 0.04457662 |
| Xu_2009_SUTs64 | Xu_2009_SU | 111.768865 | 0.30420268 | 0.15148595 | 2.00812476 | 0.04463004 |
| YOR248W | YOR248W | 2.91484038 | -0.5006374 | 0.24930992 | -2.0080925 | 0.04463347 |
| YLR283W | YLR283W | 225.687421 | -0.23344 | 0.11625877 | -2.0079347 | 0.04465023 |
| FRT2 | YAL028W | 214.353654 | -0.2459211 | 0.12250528 | -2.0074327 | 0.04470361 |
| tK(UUU)L | tK(UUU)L | 10.3210218 | -0.5946695 | 0.29649061 | -2.0056942 | 0.04488888 |
| Xu_2009_CUTs22 | Xu_2009_CU | 1.63544362 | 0.43610998 | 0.21751974 | 2.00492138 | 0.04497145 |
| Xu_2009_CUTs32 | Xu_2009_CU | 20.4028142 | -0.5467528 | 0.2727159 | -2.0048439 | 0.04497973 |
| Xu_2009_SUTs80 | Xu_2009_SU | 37.8729057 | 0.4494113 | 0.22419514 | 2.00455419 | 0.04501073 |
| YFL015C | YFL015C | 13.9317793 | 0.57118609 | 0.28494628 | 2.00453955 | 0.04501229 |
| SNR128 | snR128 | 633.080371 | -0.4451975 | 0.22231221 | -2.0025777 | 0.04522263 |
| Xu_2009_SUTs49 | Xu_2009_SU | 39.3382617 | 0.44128129 | 0.22034263 | 2.00270503 | 0.04520896 |
| ECI1 | YLR284C | 167.912359 | -0.287087 | 0.14336236 | -2.0025272 | 0.04522806 |
| YPL077C | YPL077C | 182.195677 | 0.25285676 | 0.12628446 | 2.00227937 | 0.04525469 |
| Xu_2009_CUTs76 | Xu_2009_CU | 18.6799114 | 0.55258172 | 0.27635306 | 1.99954987 | 0.04554889 |
| KTI12 | YKL110C | 680.444546 | 0.17569424 | 0.08791246 | 1.9985134 | 0.04566103 |
| Xu_2009_CUTs69 | Xu_2009_CU | 40.5793365 | 0.44045898 | 0.22045652 | 1.99794034 | 0.04572313 |
| YOR050C | YOR050C | 10.7017292 | 0.58710374 | 0.29403846 | 1.99669031 | 0.04585884 |
| Xu_2009_CUTs38 | Xu_2009_CU | 39.4143545 | 0.45055099 | 0.22578773 | 1.99546266 | 0.04599244 |
| ECM1 | YAL059W | 761.059511 | -0.1689371 | 0.08472376 | -1.9939751 | 0.04615477 |
| DBP8 | YHR169W | 1314.63904 | 0.15554721 | 0.07802294 | 1.99360863 | 0.04619484 |
| COY1 | YKL179C | 1189.44891 | 0.14536798 | 0.07300886 | 1.99110051 | 0.04646984 |
| Xu_2009_CUTs54 | Xu_2009_CU | 41.8370127 | -0.4442319 | 0.2231907 | -1.9903694 | 0.04655026 |
| REC104 | YHR157W | 100.061538 | 0.3286243 | 0.16510923 | 1.99034482 | 0.04655296 |
| SEC10 | YLR166C | 782.712 | -0.1573548 | 0.07906816 | -1.9901153 | 0.04657823 |
| YJL064W | YJL064W | 8.43057136 | 0.58906299 | 0.29608185 | 1.98952754 | 0.04664301 |
| SHU2 | YDR078C | 53.6116943 | 0.39900666 | 0.2007 | 1.98807503 | 0.04680339 |
| Xu_2009_SUTs53 | Xu_2009_SU | 231.531961 | 0.22683068 | 0.1141038 | 1.9879327 | 0.04681913 |
| SCO2 | YBR024W | 298.227017 | -0.228689 | 0.11513023 | -1.9863507 | 0.0469944 |
| BAP3 | YDR046C | 2074.31729 | 0.1314101 | 0.06617253 | 1.98587092 | 0.04704766 |
| CSI1 | YMR025W | 83.8895907 | 0.34596599 | 0.17427851 | 1.98513286 | 0.04712969 |
| Xu_2009_SUTs11 | Xu_2009_SU | 224.404108 | 0.25531005 | 0.12869571 | 1.98382712 | 0.04727512 |
| Xu_2009_CUTs77 | Xu_2009_CU | 4.0297252 | 0.5524193 | 0.27850392 | 1.98352433 | 0.04730889 |
| Xu_2009_CUTs92 | Xu_2009_CU | 13.1246135 | 0.57255116 | 0.28867177 | 1.98339851 | 0.04732293 |
| Xu_2009_SUTs37 | Xu_2009_SU | 84.0748985 | 0.35483364 | 0.1789444 | 1.98292673 | 0.04737561 |
| YDR018C | YDR018C | 108.278833 | -0.3044984 | 0.15356491 | -1.9828646 | 0.04738256 |
| Xu_2009_CUTs83 | Xu_2009_CU | 90.0147165 | 0.33720765 | 0.17015634 | 1.98175194 | 0.04750701 |
| Xu_2009_SUTs63 | Xu_2009_SU | 40.0789617 | 0.44063827 | 0.22251257 | 1.98028483 | 0.04767153 |

| | | | | | | |
|---|---|---|---|---|---|---|
| TMA16 | YOR252W | 429.602146 | 0.18849878 | 0.09520484 | 1.97992856 | 0.04771156 |
| OST3 | YOR085W | 1437.31792 | -0.14874 | 0.07513678 | -1.97959 | 0.04774962 |
| Xu_2009_CUTs74 | Xu_2009_CU | 2.15064439 | 0.47264828 | 0.2388157 | 1.97913401 | 0.04780092 |
| Xu_2009_CUTs81 | Xu_2009_CU | 17.2866041 | 0.54184065 | 0.27377412 | 1.97915216 | 0.04779888 |
| PET20 | YPL159C | 1274.19228 | 0.1405378 | 0.07102911 | 1.97859435 | 0.0478617 |
| YJR115W | YJR115W | 87.6592082 | -0.3321686 | 0.16799392 | -1.9772656 | 0.04801161 |
| KEX2 | YNL238W | 1277.29378 | 0.14750641 | 0.07459986 | 1.97730142 | 0.04800757 |
| Xu_2009_SUTs60 | Xu_2009_SU | 68.3294842 | 0.36365405 | 0.18398177 | 1.9765766 | 0.04808951 |
| tM(CAU)Q1 | tM(CAU)Q1 | 3.34472788 | -0.5181844 | 0.26236114 | -1.9750807 | 0.04825899 |
| OAF1 | YAL051W | 900.843137 | 0.14170232 | 0.07176564 | 1.97451498 | 0.04832322 |
| YAP5 | YIR018W | 336.607317 | -0.2036556 | 0.10319776 | -1.9734502 | 0.04844429 |
| STE4 | YOR212W | 1044.86487 | -0.1369909 | 0.06942113 | -1.9733308 | 0.04845789 |
| GOS1 | YHL031C | 390.814564 | -0.1947484 | 0.09872587 | -1.9726176 | 0.04853914 |
| CMG1 | YLR271W | 219.163621 | 0.24306518 | 0.12322564 | 1.97252121 | 0.04855014 |
| CEX1 | YOR112W | 725.547992 | -0.1597042 | 0.08097278 | -1.9723193 | 0.04857316 |
| AZF1 | YOR113W | 1283.9342 | 0.14328841 | 0.07265109 | 1.97228166 | 0.04857746 |
| KSS1 | YGR040W | 706.358878 | 0.15612891 | 0.07921218 | 1.97102165 | 0.0487214 |
| IXR1 | YKL032C | 1332.81403 | -0.1294935 | 0.06570002 | -1.9709814 | 0.04872601 |
| YHC1 | YLR298C | 324.058744 | -0.2075014 | 0.10528065 | -1.9709353 | 0.04873128 |
| YLR379W | YLR379W | 1.77614439 | 0.43992068 | 0.22320396 | 1.97093586 | 0.04873121 |
| GFA1 | YKL104C | 3942.07664 | 0.1153113 | 0.05852558 | 1.97027165 | 0.04880725 |
| Xu_2009_CUTs38 | Xu_2009_CU | 22.1878683 | 0.53464886 | 0.2714347 | 1.9697145 | 0.0488711 |
| ISC1 | YER019W | 754.940342 | -0.1543543 | 0.07836875 | -1.9695899 | 0.04888539 |
| GLO3 | YER122C | 2849.45289 | 0.11740932 | 0.05961892 | 1.9693299 | 0.04891522 |
| YNR064C | YNR064C | 22.1656648 | -0.5209235 | 0.26477089 | -1.9674501 | 0.04913134 |
| DXO1 | YDR370C | 550.984837 | 0.16537339 | 0.08406044 | 1.96731536 | 0.04914686 |
| DAD2 | YKR083C | 219.648797 | -0.2437503 | 0.12392196 | -1.9669661 | 0.04918712 |
| Xu_2009_CUTs28 | Xu_2009_CU | 15.3846946 | 0.55562421 | 0.28255241 | 1.96644657 | 0.04924705 |
| SRM1 | YGL097W | 1144.47 | -0.1253404 | 0.06377327 | -1.9654061 | 0.04936726 |
| tL(CAA)M | tL(CAA)M | 14.1928563 | -0.5663564 | 0.28823639 | -1.9649025 | 0.04942552 |
| Xu_2009_SUTs39 | Xu_2009_SU | 68.3450731 | 0.35457693 | 0.18049066 | 1.9645168 | 0.04947019 |
| ASF2 | YDL197C | 633.247455 | 0.17582075 | 0.08952008 | 1.96403709 | 0.04952579 |
| PGD1 | YGL025C | 460.004695 | 0.18223164 | 0.09278094 | 1.96410637 | 0.04951776 |
| AIM26 | YKL037W | 13.5330897 | -0.5660741 | 0.28833327 | -1.9632632 | 0.0496156 |
| Xu_2009_SUTs46 | Xu_2009_SU | 79.5760107 | 0.3324082 | 0.16943332 | 1.96188212 | 0.04977621 |
| Xu_2009_CUTs50 | Xu_2009_CU | 21.0803 | 0.51000798 | 0.26002411 | 1.96138726 | 0.04983386 |
| REX4 | YOL080C | 762.650085 | -0.1671759 | 0.08524577 | -1.9611046 | 0.04986683 |
| PFF1 | YBR074W | 1234.2939 | -0.1327649 | 0.06770644 | -1.9608905 | 0.0498918 |
| SDH2 | YLL041C | 739.041213 | -0.1765946 | 0.09006539 | -1.9607378 | 0.04990962 |
| COS8 | YHL048W | 570.30696 | 0.17554107 | 0.08958072 | 1.95958545 | 0.05004426 |
| Xu_2009_CUTs44 | Xu_2009_CU | 62.0614369 | 0.38116175 | 0.19462255 | 1.95846656 | 0.05017529 |

| | | | | | | |
|---|---|---|---|---|---|---|
| YBL012C | YBL012C | 3.14033179 | 0.51684177 | 0.26400884 | 1.95766843 | 0.05026893 |
| Xu_2009_CUTs13 | Xu_2009_CU | 23.7474058 | 0.51858275 | 0.26503008 | 1.95669391 | 0.05038347 |
| APC4 | YDR118W | 257.104978 | 0.22191877 | 0.11350349 | 1.95517139 | 0.05056284 |
| SME1 | YOR159C | 642.751273 | 0.19155599 | 0.09799314 | 1.95478975 | 0.05060789 |
| TFA2 | YKR062W | 790.262965 | -0.1439541 | 0.07365723 | -1.9543784 | 0.05065648 |
| tK(UUU)O | tK(UUU)O | 128.175613 | 0.29265624 | 0.14983557 | 1.95318271 | 0.05079795 |
| YPL229W | YPL229W | 411.725205 | 0.17941578 | 0.09188927 | 1.9525215 | 0.05087632 |
| Xu_2009_CUTs52 | Xu_2009_CU | 16.881297 | 0.53587097 | 0.27449037 | 1.95223962 | 0.05090976 |
| YPT53 | YNL093W | 78.1898917 | -0.3356894 | 0.1719564 | -1.9521774 | 0.05091715 |
| MIL1 | YFL034W | 1112.13677 | 0.13714393 | 0.07027188 | 1.95161882 | 0.05098348 |
| KAP104 | YBR017C | 2225.85008 | -0.1143685 | 0.05860736 | -1.9514351 | 0.05100531 |
| SLX8 | YER116C | 376.204116 | 0.19776274 | 0.10141928 | 1.94995204 | 0.05118184 |
| HTD2 | YHR067W | 376.419233 | -0.1810229 | 0.0928335 | -1.9499738 | 0.05117924 |
| SFB2 | YNL049C | 1027.0559 | -0.1580776 | 0.08106203 | -1.9500824 | 0.0511663 |
| UTP30 | YKR060W | 328.035926 | 0.21468198 | 0.11015905 | 1.94883663 | 0.05131494 |
| ZIP2 | YGL249W | 49.5882913 | -0.4035205 | 0.20709504 | -1.9484796 | 0.0513576 |
| Xu_2009_SUTs39 | Xu_2009_SU | 69.3599942 | 0.36193519 | 0.18578668 | 1.94812241 | 0.05140032 |
| SGF73 | YGL066W | 780.155407 | 0.14932461 | 0.07670309 | 1.94678744 | 0.05156022 |
| YNG2 | YHR090C | 361.997652 | -0.2093615 | 0.10761006 | -1.9455568 | 0.051708 |
| YHL044W | YHL044W | 159.445458 | 0.26576594 | 0.13661479 | 1.94536731 | 0.05173079 |
| OAZ1 | YPL052W | 303.459882 | -0.2119969 | 0.10912236 | -1.9427445 | 0.05204703 |
| CLP1 | YOR250C | 319.813726 | -0.1910913 | 0.09863979 | -1.9372638 | 0.05271311 |
| RIB1 | YBL033C | 1327.28773 | 0.13581386 | 0.07011829 | 1.93692477 | 0.05275454 |
| PAL1 | YDR348C | 1748.42745 | 0.12223109 | 0.06310683 | 1.93689153 | 0.0527586 |
| FAR11 | YNL127W | 669.475674 | 0.15169236 | 0.07833097 | 1.93655664 | 0.05279956 |
| Xu_2009_SUTs13 | Xu_2009_SU | 81.9928917 | 0.33361284 | 0.1723053 | 1.93617278 | 0.05284654 |
| YMR105W-A | YMR105W-A | 11.0613582 | 0.56878096 | 0.29394063 | 1.9350199 | 0.05298785 |
| Xu_2009_CUTs62 | Xu_2009_CU | 16.2505699 | 0.54562518 | 0.2822643 | 1.9330294 | 0.05323258 |
| YIH1 | YCR059C | 497.870648 | -0.1674111 | 0.08661327 | -1.9328575 | 0.05325375 |
| ATP23 | YNR020C | 318.767503 | 0.19653616 | 0.10175416 | 1.93148034 | 0.05342368 |
| Xu_2009_SUTs31 | Xu_2009_SU | 102.597368 | 0.3366199 | 0.17429393 | 1.93133465 | 0.05344168 |
| STE50 | YCL032W | 448.28333 | -0.1932275 | 0.10009165 | -1.9305061 | 0.05354416 |
| tR(ACG)D | tR(ACG)D | 12.0954021 | -0.5669228 | 0.29373036 | -1.9300789 | 0.05359706 |
| HEL2 | YDR266C | 1026.16076 | -0.1452482 | 0.07526998 | -1.9296967 | 0.05364443 |
| Xu_2009_SUTs81 | Xu_2009_SU | 2.47488534 | 0.48443617 | 0.25127322 | 1.92792601 | 0.05386433 |
| Xu_2009_CUTs64 | Xu_2009_CU | 36.9980271 | 0.4614529 | 0.2394948 | 1.92677625 | 0.05400752 |
| SPC3 | YLR066W | 349.825222 | -0.1886191 | 0.09789937 | -1.9266626 | 0.05402169 |
| PEX11 | YOL147C | 136.169586 | -0.2844764 | 0.14764957 | -1.9266998 | 0.05401705 |
| HMI1 | YOL095C | 196.565978 | -0.2523875 | 0.13113004 | -1.9247113 | 0.05426548 |
| TIR1 | YER011W | 31.4537088 | -0.4647546 | 0.24149185 | -1.9245148 | 0.05429009 |
| SMI1 | YGR229C | 2482.93452 | 0.11729526 | 0.0609573 | 1.92422003 | 0.05432701 |

| | | | | | | |
|---|---|---|---|---|---|---|
| YCR095W-A | YCR095W-A | 18.3174391 | 0.52275274 | 0.27170431 | 1.92397663 | 0.05435751 |
| LCD1 | YDR499W | 502.145338 | 0.18565734 | 0.09649563 | 1.92399736 | 0.05435491 |
| PFS2 | YNL317W | 659.189124 | -0.1593924 | 0.0828722 | -1.9233515 | 0.05443592 |
| YFR034W-A | YFR034W-A | 138.776673 | 0.26983309 | 0.14031031 | 1.92311664 | 0.0544654 |
| MST27 | YGL051W | 38.9500734 | -0.460071 | 0.23929395 | -1.9226185 | 0.05452798 |
| FOL3 | YMR113W | 321.618003 | -0.1958751 | 0.10190733 | -1.9220907 | 0.05459435 |
| IDH1 | YNL037C | 1026.4581 | -0.1671571 | 0.08695966 | -1.922237 | 0.05457595 |
| PEX25 | YPL112C | 478.161404 | -0.1745595 | 0.0908164 | -1.9221148 | 0.05459132 |
| ABZ2 | YMR289W | 619.879243 | -0.1507378 | 0.07842999 | -1.9219413 | 0.05461314 |
| STB6 | YKL072W | 541.771914 | 0.15808133 | 0.08228969 | 1.92103458 | 0.05472735 |
| SUV3 | YPL029W | 505.122154 | -0.1667211 | 0.08681272 | -1.9204687 | 0.05479872 |
| Xu_2009_CUTs91 | Xu_2009_CU | 49.198317 | 0.40353622 | 0.21019614 | 1.91980795 | 0.05488216 |
| Xu_2009_SUTs46 | Xu_2009_SU | 17.027263 | 0.54095375 | 0.28173978 | 1.92004745 | 0.05485191 |
| Xu_2009_SUTs81 | Xu_2009_SU | 43.9688934 | 0.41382414 | 0.2155524 | 1.9198308 | 0.05487928 |
| CAT8 | YMR280C | 316.78699 | 0.21150878 | 0.11016793 | 1.91987607 | 0.05487356 |
| AIM10 | YER087W | 407.67092 | -0.1763951 | 0.09188607 | -1.9197155 | 0.05489385 |
| YDL247W-A | YDL247W-A | 30.688129 | 0.49469809 | 0.25773995 | 1.91936905 | 0.05493765 |
| RML2 | YEL050C | 1102.51154 | -0.1397743 | 0.07282217 | -1.9193921 | 0.05493474 |
| Xu_2009_SUTs40 | Xu_2009_SU | 47.7615345 | 0.41283609 | 0.21520015 | 1.91838199 | 0.05506259 |
| RRN7 | YJL025W | 448.922589 | 0.18325321 | 0.09555428 | 1.91779172 | 0.05513743 |
| Xu_2009_SUTs67 | Xu_2009_SU | 19.1125638 | 0.51967958 | 0.27099127 | 1.91769861 | 0.05514924 |
| YDR098C-B | YDR098C-B | 131.972353 | 0.27017661 | 0.14099563 | 1.91620558 | 0.05533893 |
| PTC6 | YCR079W | 742.953525 | -0.1428024 | 0.07453623 | -1.9158797 | 0.05538041 |
| DET1 | YDR051C | 658.487057 | -0.1534041 | 0.08007152 | -1.9158388 | 0.05538561 |
| UFD1 | YGR048W | 869.310633 | -0.1401716 | 0.073159 | -1.9159852 | 0.05536698 |
| Xu_2009_CUTs81 | Xu_2009_CU | 20.9014455 | 0.51552306 | 0.26913434 | 1.91548598 | 0.05543056 |
| Xu_2009_CUTs32 | Xu_2009_CU | 18.2187392 | 0.51902592 | 0.27129223 | 1.91316175 | 0.05572735 |
| DIE2 | YGR227W | 450.00925 | -0.1858194 | 0.09718035 | -1.9121085 | 0.05586228 |
| Xu_2009_CUTs80 | Xu_2009_CU | 24.2833417 | 0.48180531 | 0.25211112 | 1.91108317 | 0.05599389 |
| VAM3 | YOR106W | 653.978707 | 0.1673039 | 0.08757056 | 1.91050387 | 0.05606837 |
| EMC4 | YGL231C | 518.324219 | -0.1938182 | 0.10159434 | -1.9077656 | 0.05642152 |
| PIF1 | YML061C | 707.657029 | -0.1418624 | 0.07445418 | -1.9053647 | 0.05673268 |
| Xu_2009_CUTs33 | Xu_2009_CU | 5.64713287 | 0.55162219 | 0.28953933 | 1.90517189 | 0.05675773 |
| SRX1 | YKL086W | 29.6889451 | 0.47479489 | 0.24927613 | 1.9046946 | 0.05681978 |
| YDR003W-A | YDR003W-A | 80.7266663 | -0.3255578 | 0.17098896 | -1.9039696 | 0.05691415 |
| SAF1 | YBR280C | 330.389262 | -0.1925345 | 0.10113841 | -1.9036733 | 0.05695275 |
| tP(UGG)F/SUF9 | tP(UGG)F | 3.8576586 | -0.5237524 | 0.27521239 | -1.9030843 | 0.05702954 |
| RRS1 | YOR294W | 1950.1411 | -0.1512819 | 0.07958638 | -1.9008515 | 0.05732146 |
| ALA1 | YOR335C | 4153.50934 | -0.120757 | 0.06356597 | -1.8997121 | 0.05747091 |
| YCR007C | YCR007C | 254.417876 | 0.26590565 | 0.13999514 | 1.89939201 | 0.05751295 |
| SRP68 | YPL243W | 1230.28433 | -0.1489814 | 0.07843297 | -1.8994738 | 0.05750221 |

| | | | | | | |
|---|---|---|---|---|---|---|
| Xu_2009_CUTs20 | Xu_2009_CU | 13.8449768 | 0.54613145 | 0.28766222 | 1.89851641 | 0.05762809 |
| Xu_2009_SUTs79 | Xu_2009_SU | 95.218893 | 0.3030712 | 0.15965232 | 1.89832008 | 0.05765393 |
| YDR094W | YDR094W | 4.13112485 | 0.52732882 | 0.27789919 | 1.89755436 | 0.05775481 |
| NMA111 | YNL123W | 2287.3708 | 0.13137482 | 0.06924204 | 1.89732733 | 0.05778475 |
| SMP3 | YOR149C | 404.343986 | 0.18399492 | 0.09700527 | 1.8967517 | 0.05786072 |
| TPA1 | YER049W | 1744.30184 | -0.1246745 | 0.06573823 | -1.8965293 | 0.05789009 |
| YIR020C-B | YIR020C-B | 7.77562973 | -0.5626571 | 0.29667391 | -1.8965507 | 0.05788726 |
| QDR3 | YBR043C | 772.018881 | 0.13834829 | 0.07297846 | 1.89574146 | 0.05799424 |
| FES1 | YBR101C | 949.026362 | -0.1532272 | 0.08084829 | -1.8952435 | 0.05806015 |
| FIS1 | YIL065C | 469.447085 | -0.1680892 | 0.08869181 | -1.8952058 | 0.05806515 |
| SNU56 | YDR240C | 524.444402 | -0.1672039 | 0.08828021 | -1.8940135 | 0.05822322 |
| RTT106 | YNL206C | 702.82707 | -0.157924 | 0.08339383 | -1.8937133 | 0.05826308 |
| Xu_2009_CUTs84 | Xu_2009_CU | 18.1010982 | 0.51914956 | 0.27432279 | 1.892477 | 0.05842745 |
| UBC6 | YER100W | 1312.37753 | -0.1252178 | 0.06617729 | -1.8921562 | 0.05847017 |
| IST1 | YNL265C | 379.403149 | -0.1813453 | 0.09594773 | -1.8900428 | 0.05875224 |
| PFA3 | YNL326C | 267.779592 | -0.2078136 | 0.10998299 | -1.8895072 | 0.0588239 |
| RSM27 | YGR215W | 552.76107 | 0.19351609 | 0.1024443 | 1.88898838 | 0.05889339 |
| SDH4 | YDR178W | 1032.71793 | -0.1414429 | 0.07489159 | -1.8886357 | 0.05894066 |
| SPO19 | YPL130W | 94.2598668 | -0.3209253 | 0.16992781 | -1.8885981 | 0.0589457 |
| VAB2 | YEL005C | 261.092771 | -0.2208676 | 0.11705421 | -1.8868832 | 0.05917604 |
| AI2 | Q0055 | 5561.16124 | 0.13041335 | 0.06917688 | 1.8852159 | 0.0594007 |
| Xu_2009_SUTs27 | Xu_2009_SU | 43.083979 | -0.4202513 | 0.2230845 | -1.8838214 | 0.05958914 |
| DAD4 | YDR320C-A | 245.630076 | -0.2256136 | 0.11987736 | -1.8820365 | 0.05983107 |
| Xu_2009_SUTs51 | Xu_2009_SU | 1.01226752 | 0.34912145 | 0.18573295 | 1.87969583 | 0.06014954 |
| PEX5 | YDR244W | 701.827777 | 0.14294732 | 0.07604795 | 1.87969981 | 0.060149 |
| SRB7 | YDR308C | 414.493105 | -0.1768824 | 0.09414372 | -1.8788553 | 0.06026425 |
| SNR54 | snR54 | 208.590625 | 0.24756732 | 0.13181739 | 1.87810823 | 0.06036636 |
| ADA2 | YDR448W | 686.39772 | -0.1457766 | 0.07761713 | -1.8781501 | 0.06036063 |
| Xu_2009_CUTs61 | Xu_2009_CU | 29.7791437 | 0.46577155 | 0.2480755 | 1.87753949 | 0.06044419 |
| LDB16 | YCL005W | 392.852572 | -0.1757265 | 0.09361312 | -1.8771572 | 0.06049655 |
| GNT1 | YOR320C | 787.377179 | -0.1387036 | 0.07393362 | -1.8760565 | 0.06064752 |
| Xu_2009_SUTs67 | Xu_2009_SU | 255.69652 | 0.22643663 | 0.12074768 | 1.87528768 | 0.06075316 |
| MCM10 | YIL150C | 459.141378 | 0.16408379 | 0.08753597 | 1.87447281 | 0.06086529 |
| SEN54 | YPL083C | 347.695574 | 0.18197981 | 0.09710873 | 1.87397986 | 0.0609332 |
| Xu_2009_CUTs23 | Xu_2009_CU | 12.730748 | 0.54032518 | 0.28837771 | 1.87367178 | 0.06097568 |
| ALG3 | YBL082C | 138.582212 | -0.2972081 | 0.15863545 | -1.8735287 | 0.06099541 |
| Xu_2009_SUTs79 | Xu_2009_SU | 7.01072809 | 0.55135663 | 0.29434602 | 1.87315809 | 0.06104656 |
| YAR075W | YAR075W | 0.98531035 | 0.33711138 | 0.17999847 | 1.87285692 | 0.06108815 |
| NOP12 | YOL041C | 1729.24019 | -0.1268016 | 0.06773749 | -1.8719556 | 0.06121274 |
| Xu_2009_SUTs48 | Xu_2009_SU | 156.760908 | 0.24482706 | 0.13085908 | 1.87092155 | 0.06135596 |
| IPL1 | YPL209C | 132.859702 | 0.265068 | 0.1419602 | 1.86719938 | 0.06187375 |

| | | | | | | |
|---|---|---|---|---|---|---|
| HXT12 | YIL170W | 5.59789296 | -0.5419541 | 0.29030801 | -1.8668243 | 0.06192612 |
| Xu_2009_SUTs74( | Xu_2009_SU | 13.0726769 | 0.54747319 | 0.2934857 | 1.8654169 | 0.06212299 |
| COS2 | YBR302C | 70.6493867 | 0.34687506 | 0.18612444 | 1.86367288 | 0.06236766 |
| Xu_2009_CUTs78 | Xu_2009_CU | 55.5784601 | 0.37256925 | 0.19994734 | 1.86333688 | 0.06241488 |
| Xu_2009_CUTs50 | Xu_2009_CU | 25.5828922 | 0.48630631 | 0.26111991 | 1.86238691 | 0.06254857 |
| RIB5 | YBR256C | 764.32891 | -0.1627419 | 0.08742909 | -1.8614158 | 0.06268548 |
| YNL235C | YNL235C | 6.632869 | 0.54777078 | 0.29448524 | 1.86009589 | 0.06287196 |
| TRM12 | YML005W | 312.191202 | -0.1990595 | 0.10703092 | -1.8598314 | 0.06290939 |
| UBX4 | YMR067C | 971.244516 | 0.13341175 | 0.07173287 | 1.85984117 | 0.062908 |
| SPP382 | YLR424W | 239.219894 | -0.2138796 | 0.11501382 | -1.8595995 | 0.06294221 |
| NAF1 | YNL124W | 1991.17245 | 0.25709834 | 0.13825466 | 1.85959981 | 0.06294217 |
| YDL172C | YDL172C | 21.9699946 | 0.48947142 | 0.26325752 | 1.85928749 | 0.0629864 |
| HXT10 | YFL011W | 31.9483935 | -0.4523465 | 0.24331403 | -1.8591056 | 0.06301217 |
| HSP33 | YOR391C | 23.0678713 | 0.48890184 | 0.26298655 | 1.85903744 | 0.06302183 |
| RMD8 | YFR048W | 846.824028 | -0.1360995 | 0.07322454 | -1.8586591 | 0.06307548 |
| Xu_2009_CUTs44 | Xu_2009_CU | 29.2012381 | 0.45053707 | 0.2425125 | 1.85778911 | 0.06319897 |
| DLD3 | YEL071W | 2477.17411 | -0.2455201 | 0.13216004 | -1.8577482 | 0.06320479 |
| MCM22 | YJR135C | 173.846034 | 0.24338666 | 0.13099318 | 1.85801026 | 0.06316756 |
| YNR068C | YNR068C | 97.6073151 | 0.29805259 | 0.1604359 | 1.85776741 | 0.06320205 |
| Xu_2009_CUTs31 | Xu_2009_CU | 20.4053302 | 0.50155354 | 0.27000595 | 1.85756475 | 0.06323085 |
| YGL052W | YGL052W | 38.1463 | -0.4382768 | 0.23601555 | -1.8569829 | 0.06331359 |
| tL(CAA)G1 | tL(CAA)G1 | 7.2973235 | -0.5501774 | 0.29637329 | -1.8563665 | 0.06340134 |
| ORM1 | YGR038W | 421.258913 | -0.1693522 | 0.09124676 | -1.8559802 | 0.06345638 |
| Xu_2009_CUTs57 | Xu_2009_CU | 9.37862859 | 0.54903272 | 0.29583676 | 1.85586378 | 0.06347298 |
| SAE3 | YHR079C-A | 5.49596172 | 0.54148202 | 0.29186037 | 1.8552776 | 0.0635566 |
| Xu_2009_CUTs17 | Xu_2009_CU | 29.6572377 | 0.46543603 | 0.25097957 | 1.85447779 | 0.06367083 |
| Xu_2009_SUTs76 | Xu_2009_SU | 62.9575272 | 0.35111995 | 0.18934576 | 1.85438511 | 0.06368408 |
| MEI4 | YER044C-A | 194.479886 | 0.22163705 | 0.11962569 | 1.85275457 | 0.06391754 |
| CDC36 | YDL165W | 430.748257 | -0.1670948 | 0.09022245 | -1.8520309 | 0.06402138 |
| Xu_2009_CUTs19 | Xu_2009_CU | 13.0557007 | 0.53301246 | 0.28786489 | 1.85160633 | 0.06408237 |
| APC9 | YLR102C | 115.969138 | 0.29105322 | 0.15728296 | 1.85050701 | 0.06424051 |
| YOL160W | YOL160W | 36.9888577 | 0.44025142 | 0.23791242 | 1.8504768 | 0.06424486 |
| CPR7 | YJR032W | 574.124362 | -0.1535485 | 0.08301423 | -1.849665 | 0.06436185 |
| MTL1 | YGR023W | 715.477317 | -0.1506007 | 0.08146489 | -1.8486573 | 0.06450731 |
| RNH202 | YDR279W | 328.148739 | -0.1810011 | 0.09803303 | -1.846328 | 0.06484459 |
| tL(UAG)J | tL(UAG)J | 0.72935938 | -0.2830448 | 0.15336247 | -1.8455938 | 0.06495121 |
| ERD1 | YDR414C | 133.991676 | -0.2616784 | 0.14184341 | -1.84484 | 0.06506082 |
| SKN7 | YHR206W | 2197.95652 | 0.12877957 | 0.06980404 | 1.84487286 | 0.06505604 |
| MPS3 | YJL019W | 606.473449 | 0.16248913 | 0.08808198 | 1.84474882 | 0.06507409 |
| SAS4 | YDR181C | 432.339767 | 0.16948576 | 0.09195755 | 1.84308685 | 0.06531633 |
| MNS1 | YJR131W | 710.494775 | 0.15943567 | 0.08653111 | 1.84252436 | 0.06539849 |

| | | | | | | |
|---|---|---|---|---|---|---|
| SEC72 | YLR292C | 1221.09735 | -0.1281014 | 0.0695369 | -1.8422072 | 0.06544485 |
| Xu_2009_CUTs85 | Xu_2009_CU | 14.1621536 | 0.53338261 | 0.28956518 | 1.84201224 | 0.06547336 |
| Xu_2009_CUTs67 | Xu_2009_CU | 25.4573128 | 0.48034801 | 0.2610182 | 1.84028548 | 0.06572634 |
| tE(UUC)C | tE(UUC)C | 6.88501039 | -0.5395665 | 0.29371651 | -1.8370316 | 0.06620524 |
| Xu_2009_CUTs10 | Xu_2009_CU | 104.020158 | 0.30888542 | 0.16824087 | 1.83597141 | 0.06636188 |
| DON1 | YDR273W | 147.443655 | 0.25877961 | 0.14095226 | 1.83593806 | 0.06636682 |
| Xu_2009_CUTs64 | Xu_2009_CU | 15.4722531 | 0.51522222 | 0.28084243 | 1.83455977 | 0.06657094 |
| GPB2 | YAL056W | 1048.326 | 0.12628682 | 0.0688591 | 1.83398879 | 0.06665566 |
| KAR3 | YPR141C | 461.195674 | -0.1718501 | 0.09378673 | -1.8323494 | 0.06689938 |
| JLP1 | YLL057C | 37.3646719 | -0.4172664 | 0.22775358 | -1.8320956 | 0.06693717 |
| SEI1 | YLR404W | 191.337841 | -0.2283915 | 0.12476543 | -1.8305675 | 0.06716512 |
| FHN1 | YGR131W | 47.6399166 | 0.39508502 | 0.21592626 | 1.82972191 | 0.06729153 |
| GIT1 | YCR098C | 35.9009506 | 0.42323046 | 0.23132217 | 1.82961473 | 0.06730757 |
| SPG1 | YGR236C | 34.0348967 | 0.44631024 | 0.24418312 | 1.82776857 | 0.06758429 |
| YOR142W-B | YOR142W-B | 116.86459 | 0.28332152 | 0.15501157 | 1.82774429 | 0.06758794 |
| tV(AAC)K1 | tV(AAC)K1 | 3.47165281 | -0.4804961 | 0.26294834 | -1.8273402 | 0.06764864 |
| MVP1 | YMR004W | 924.827619 | 0.14325954 | 0.07841303 | 1.82698636 | 0.06770182 |
| AAC3 | YBR085W | 28.402824 | -0.4901414 | 0.26838731 | -1.8262467 | 0.06781311 |
| YNL179C | YNL179C | 92.7696089 | 0.30366257 | 0.16629695 | 1.82602612 | 0.06784633 |
| tM(CAU)E/IMT4 | tM(CAU)E | 3.81960436 | -0.5009282 | 0.27436945 | -1.8257435 | 0.06788891 |
| GTO3 | YMR251W | 36.6463345 | -0.4288644 | 0.23509005 | -1.8242557 | 0.06811342 |
| Xu_2009_SUTs62 | Xu_2009_SU | 15.273074 | 0.51254279 | 0.28115368 | 1.82299872 | 0.06830359 |
| YCR025C | YCR025C | 36.2843284 | 0.41753906 | 0.22907103 | 1.82274931 | 0.06834137 |
| NEJ1 | YLR265C | 382.683867 | -0.1821062 | 0.09993726 | -1.8222052 | 0.06842386 |
| Xu_2009_CUTs83 | Xu_2009_CU | 8.69223449 | 0.53897739 | 0.29584831 | 1.82180316 | 0.06848486 |
| YDL144C | YDL144C | 840.554193 | -0.1519162 | 0.08340685 | -1.821388 | 0.0685479 |
| MUB1 | YMR100W | 1324.63446 | 0.11846207 | 0.06506596 | 1.82064602 | 0.06866068 |
| Xu_2009_SUTs44 | Xu_2009_SU | 36.2318677 | 0.42639463 | 0.23430298 | 1.819843 | 0.06878292 |
| YRF1-8 | YOR396W | 3.02529852 | 0.48161095 | 0.26465017 | 1.81980217 | 0.06878914 |
| PBS2 | YJL128C | 1884.68852 | 0.11276124 | 0.0619758 | 1.81943997 | 0.06884433 |
| VID22 | YLR373C | 1106.78786 | 0.11920751 | 0.06551606 | 1.81951591 | 0.06883276 |
| Xu_2009_SUTs27 | Xu_2009_SU | 95.3139917 | 0.31015167 | 0.17065491 | 1.81742012 | 0.06915281 |
| Xu_2009_SUTs77 | Xu_2009_SU | 38.8602225 | 0.40580188 | 0.22334211 | 1.81695197 | 0.06922447 |
| BRL1 | YHR036W | 890.245588 | 0.12615451 | 0.06944437 | 1.81662677 | 0.06927428 |
| PET8 | YNL003C | 433.6903 | -0.1648576 | 0.09074538 | -1.8167053 | 0.06926225 |
| DOG1 | YHR044C | 79.255352 | -0.3265769 | 0.17983442 | -1.8159869 | 0.06937239 |
| Xu_2009_CUTs46 | Xu_2009_CU | 18.8414015 | 0.51048997 | 0.28121207 | 1.81532028 | 0.0694747 |
| Xu_2009_CUTs41 | Xu_2009_CU | 11.1096592 | 0.52976232 | 0.29190845 | 1.81482349 | 0.06955104 |
| PDE2 | YOR360C | 1971.75064 | 0.13164556 | 0.07254134 | 1.81476593 | 0.06955989 |
| tG(GCC)P2 | tG(GCC)P2 | 10.5050073 | -0.5354894 | 0.2951123 | -1.8145277 | 0.06959653 |
| GRS2 | YPR081C | 904.893516 | 0.14671049 | 0.08087358 | 1.81407183 | 0.06966667 |

| | | | | | | |
|---|---|---|---|---|---|---|
| YGR176W | YGR176W | 16.1210804 | 0.51508523 | 0.28401828 | 1.81356367 | 0.06974493 |
| SNT309 | YPR101W | 349.896444 | 0.1934974 | 0.10676512 | 1.81236543 | 0.06992975 |
| AMS1 | YGL156W | 942.367154 | -0.140843 | 0.07779095 | -1.8105323 | 0.07021327 |
| MET16 | YPR167C | 334.174435 | 0.17723611 | 0.09789156 | 1.81053528 | 0.07021282 |
| LSB6 | YJL100W | 406.761807 | -0.1739282 | 0.09607754 | -1.8102894 | 0.07025092 |
| tK(CUU)I | tK(CUU)I | 2.9136992 | -0.4739525 | 0.26185538 | -1.809978 | 0.0702992 |
| YER186C | YER186C | 390.508839 | -0.1721144 | 0.09510531 | -1.809725 | 0.07033845 |
| SNU23 | YDL098C | 468.863957 | -0.1632129 | 0.09023252 | -1.808803 | 0.07048161 |
| Xu_2009_SUTs13 | Xu_2009_SU | 94.2580021 | -0.3026541 | 0.16739077 | -1.8080691 | 0.07059575 |
| YJL195C | YJL195C | 2.98839397 | 0.46563352 | 0.25757797 | 1.80773811 | 0.07064727 |
| CTR3 | YLR411W | 103.960381 | -0.2979827 | 0.16483449 | -1.807769 | 0.07064246 |
| Xu_2009_CUTs27 | Xu_2009_CU | 1.82077042 | -0.4159587 | 0.23023385 | -1.8066793 | 0.0708123 |
| INA17 | YPL099C | 257.992918 | -0.2012121 | 0.11137622 | -1.8065987 | 0.07082488 |
| YLR225C | YLR225C | 1593.09171 | -0.1076844 | 0.05960925 | -1.8065044 | 0.07083959 |
| Xu_2009_SUTs69 | Xu_2009_SU | 21.6032922 | 0.48577955 | 0.2689244 | 1.8063796 | 0.07085907 |
| TUB4 | YLR212C | 309.463387 | -0.1946669 | 0.10777079 | -1.8063052 | 0.07087068 |
| SHR5 | YOL110W | 161.733388 | -0.2360874 | 0.13075341 | -1.8055927 | 0.07098199 |
| Xu_2009_CUTs63 | Xu_2009_CU | 21.9567866 | 0.48154425 | 0.26679635 | 1.80491317 | 0.07108827 |
| RRG9 | YNL213C | 742.416759 | -0.1372749 | 0.07606954 | -1.8045972 | 0.07113773 |
| LCB2 | YDR062W | 1820.92287 | -0.1168911 | 0.06480293 | -1.8037928 | 0.0712638 |
| SKO1 | YNL167C | 1449.22303 | 0.11525234 | 0.06391981 | 1.80307711 | 0.07137611 |
| NSE4 | YDL105W | 202.733389 | -0.2140442 | 0.11896396 | -1.7992357 | 0.07198141 |
| RGR1 | YLR071C | 1225.42266 | 0.11984371 | 0.06663645 | 1.79847084 | 0.07210243 |
| HXT9 | YJL219W | 15.5447613 | -0.5148319 | 0.28629283 | -1.7982703 | 0.07213419 |
| Xu_2009_SUTs68 | Xu_2009_SU | 123.102289 | 0.26485649 | 0.14745714 | 1.7961592 | 0.0724692 |
| tD(GUC)J3 | tD(GUC)J3 | 5.35498771 | 0.5129747 | 0.28572862 | 1.79532137 | 0.07260251 |
| ZRG17 | YNR039C | 825.923944 | -0.130879 | 0.07292765 | -1.7946417 | 0.0727108 |
| YJL114W | YJL114W | 25.6062583 | 0.45766277 | 0.25524549 | 1.79302979 | 0.07296816 |
| YNL193W | YNL193W | 613.079074 | 0.14034926 | 0.07827589 | 1.79300744 | 0.07297173 |
| PET117 | YER058W | 104.106203 | -0.2830726 | 0.15801776 | -1.7913977 | 0.0732295 |
| RKI1 | YOR095C | 556.462092 | -0.1642711 | 0.09172456 | -1.7909176 | 0.07330652 |
| SNR81 | snR81 | 98.7983022 | -0.2993474 | 0.16751639 | -1.7869738 | 0.07394173 |
| BCK1 | YJL095W | 1204.63334 | 0.11891452 | 0.06655395 | 1.78673884 | 0.07397971 |
| MED1 | YPR070W | 550.300023 | 0.16768781 | 0.09389697 | 1.7858702 | 0.07412028 |
| LDB17 | YDL146W | 188.021435 | -0.23844 | 0.13353989 | -1.7855336 | 0.07417481 |
| YIP5 | YGL161C | 930.205196 | -0.1289086 | 0.07224217 | -1.7843952 | 0.07435947 |
| Xu_2009_CUTs20 | Xu_2009_CU | 4.50293426 | -0.5042637 | 0.28274068 | -1.7834847 | 0.07450743 |
| Xu_2009_SUTs38 | Xu_2009_SU | 39.3250812 | 0.4042538 | 0.22671446 | 1.78309666 | 0.07457057 |
| YCR099C | YCR099C | 36.5115275 | 0.41442689 | 0.23272637 | 1.78074744 | 0.07495372 |
| Xu_2009_SUTs52 | Xu_2009_SU | 23.4383278 | -0.4721455 | 0.26517671 | -1.780494 | 0.07499516 |
| HOM3 | YER052C | 1441.96953 | -0.1116768 | 0.06274197 | -1.7799384 | 0.07508604 |

| | | | | | | |
|---|---|---|---|---|---|---|
| ARP10 | YDR106W | 69.1047492 | 0.34484358 | 0.19375882 | 1.77975675 | 0.07511578 |
| Xu_2009_SUTs23 | Xu_2009_SU | 84.8490634 | 0.2999797 | 0.16856571 | 1.77960102 | 0.07514128 |
| YER152W-A | YER152W-A | 4.74513008 | 0.50289226 | 0.28261927 | 1.77939833 | 0.07517448 |
| Xu_2009_CUTs89 | Xu_2009_CU | 77.7839559 | 0.32540992 | 0.18297573 | 1.77843218 | 0.0753329 |
| YMR242W-A | YMR242W-A | 3.57785675 | -0.4764748 | 0.26800041 | -1.7778883 | 0.07542221 |
| GPI19 | YDR437W | 178.584186 | 0.22222966 | 0.12503588 | 1.77732709 | 0.07551444 |
| ISU1 | YPL135W | 1260.00846 | -0.1123863 | 0.06323865 | -1.7771773 | 0.07553907 |
| RNA14 | YMR061W | 1207.83987 | 0.11325152 | 0.06373294 | 1.77697004 | 0.07557317 |
| Xu_2009_CUTs20 | Xu_2009_CU | 72.0322733 | 0.32439979 | 0.18256947 | 1.77685668 | 0.07559182 |
| SWC3 | YAL011W | 553.465177 | 0.1503082 | 0.08462049 | 1.77626255 | 0.07568965 |
| Xu_2009_CUTs17 | Xu_2009_CU | 4.82768195 | 0.51378605 | 0.28929497 | 1.77599373 | 0.07573395 |
| YLR334C | YLR334C | 3.17383654 | 0.47453238 | 0.26726205 | 1.77553221 | 0.07581005 |
| YGL185C | YGL185C | 354.455091 | -0.1746018 | 0.09836028 | -1.7751255 | 0.07587717 |
| ANS1 | YHR126C | 29.1526787 | 0.45554125 | 0.25687405 | 1.77340315 | 0.07616193 |
| MON2 | YNL297C | 2182.3954 | 0.09848299 | 0.05553635 | 1.77330703 | 0.07617785 |
| VPS62 | YGR141W | 279.452809 | 0.20537314 | 0.11583272 | 1.77301486 | 0.07622624 |
| NQM1 | YGR043C | 701.7363 | 0.16239154 | 0.09165654 | 1.77173968 | 0.07643778 |
| YGK3 | YOL128C | 526.04944 | -0.1730546 | 0.09773806 | -1.7705962 | 0.07662788 |
| VTH2 | YJL222W | 18.1790535 | 0.48311991 | 0.27291513 | 1.77022031 | 0.07669045 |
| GRX7 | YBR014C | 812.13221 | -0.1469843 | 0.08306694 | -1.7694677 | 0.07681586 |
| YML082W | YML082W | 1018.68199 | 0.1265025 | 0.07152388 | 1.76867507 | 0.07694811 |
| SEN15 | YMR059W | 161.48895 | -0.2443073 | 0.13822114 | -1.7675106 | 0.07714275 |
| Xu_2009_CUTs29 | Xu_2009_CU | 1.55362517 | 0.38634257 | 0.21859798 | 1.7673657 | 0.077167 |
| MSI1 | YBR195C | 263.254812 | -0.2031161 | 0.11497013 | -1.7666856 | 0.07728089 |
| Xu_2009_SUTs44 | Xu_2009_SU | 43.9136422 | -0.3978215 | 0.22530615 | -1.7656931 | 0.07744736 |
| DDC1 | YPL194W | 398.974159 | 0.17730425 | 0.10049315 | 1.76434163 | 0.07767448 |
| SPT16 | YGL207W | 3213.91655 | 0.10638717 | 0.0603224 | 1.7636428 | 0.07779214 |
| NDT80 | YHR124W | 131.775263 | 0.25859058 | 0.14680953 | 1.76140178 | 0.07817042 |
| Xu_2009_CUTs65 | Xu_2009_CU | 18.3271747 | 0.49539515 | 0.28131474 | 1.76099961 | 0.07823847 |
| MRN1 | YPL184C | 983.910861 | -0.1222934 | 0.06946172 | -1.7605866 | 0.07830841 |
| Xu_2009_CUTs81 | Xu_2009_CU | 51.3228752 | 0.35989516 | 0.20451473 | 1.75975182 | 0.0784499 |
| SXM1 | YDR395W | 2637.96682 | -0.0989636 | 0.05625044 | -1.7593392 | 0.07851992 |
| YKR045C | YKR045C | 224.110036 | 0.2053539 | 0.11672529 | 1.75929224 | 0.07852788 |
| SLF1 | YDR515W | 616.997949 | -0.1409098 | 0.08013316 | -1.7584458 | 0.07867169 |
| PCL7 | YIL050W | 640.587133 | -0.1444548 | 0.08216194 | -1.7581717 | 0.0787183 |
| PTP2 | YOR208W | 1572.44506 | 0.1273592 | 0.0724391 | 1.75815548 | 0.07872106 |
| NOC4 | YPR144C | 952.066487 | 0.15532297 | 0.08837472 | 1.75754983 | 0.07882414 |
| SWD1 | YAR003W | 448.962907 | 0.15285454 | 0.08698354 | 1.75728115 | 0.0788699 |
| Xu_2009_SUTs75 | Xu_2009_SU | 131.733283 | 0.25795815 | 0.14680734 | 1.75712025 | 0.07889731 |
| YPR059C | YPR059C | 34.4956394 | -0.4101885 | 0.23351462 | -1.7565857 | 0.07898844 |
| Xu_2009_SUTs12 | Xu_2009_SU | 65.6493519 | -0.3282194 | 0.1869735 | -1.7554328 | 0.07918531 |

| | | | | | | |
|---|---|---|---|---|---|---|
| Xu_2009_SUTs42( | Xu_2009_SU | 78.9080689 | -0.3031494 | 0.17273783 | -1.7549679 | 0.0792648 |
| MNC1 | YBR056W-A | 500.186454 | 0.19970167 | 0.11380066 | 1.75483752 | 0.0792871 |
| TEX1 | YNL253W | 358.199807 | -0.177518 | 0.10115529 | -1.7549056 | 0.07927545 |
| NOP9 | YJL010C | 1197.61486 | 0.12670042 | 0.07221113 | 1.75458281 | 0.07933069 |
| KAE1 | YKR038C | 1128.7191 | 0.11947353 | 0.06811445 | 1.75401135 | 0.07942856 |
| PKR1 | YMR123W | 651.642994 | 0.15002645 | 0.08552884 | 1.75410361 | 0.07941275 |
| ATP19 | YOL077W-A | 1706.7279 | 0.16010222 | 0.09133901 | 1.75283517 | 0.0796303 |
| YOR293C-A | YOR293C-A | 3.30192119 | -0.4719559 | 0.26953708 | -1.750987 | 0.07994815 |
| OGG1 | YML060W | 322.280827 | -0.1794523 | 0.10249358 | -1.7508638 | 0.07996938 |
| Xu_2009_SUTs70 | Xu_2009_SU | 104.878061 | 0.28779381 | 0.1644117 | 1.75044604 | 0.08004138 |
| Xu_2009_CUTs62 | Xu_2009_CU | 10.1077653 | 0.51600423 | 0.29482409 | 1.75021055 | 0.08008199 |
| TRM2 | YKR056W | 1208.82648 | 0.14594784 | 0.08342778 | 1.74939135 | 0.08022339 |
| Xu_2009_SUTs26( | Xu_2009_SU | 38.2810156 | 0.38842387 | 0.22206379 | 1.74915445 | 0.08026432 |
| Xu_2009_CUTs28 | Xu_2009_CU | 4.7721622 | -0.4948616 | 0.28311819 | -1.7478975 | 0.08048177 |
| Xu_2009_CUTs57 | Xu_2009_CU | 10.0199405 | 0.51453482 | 0.29447354 | 1.74730406 | 0.08058461 |
| Xu_2009_SUTs78( | Xu_2009_SU | 28.7010904 | -0.4430994 | 0.25372517 | -1.7463753 | 0.08074575 |
| Xu_2009_SUTs54 | Xu_2009_SU | 30.2284183 | 0.42549283 | 0.24367502 | 1.74614875 | 0.08078511 |
| MED4 | YOR174W | 464.792817 | -0.1654535 | 0.0948155 | -1.7450049 | 0.08098401 |
| MET3 | YJR010W | 465.383209 | 0.16438935 | 0.09425806 | 1.74403493 | 0.081153 |
| MON1 | YGL124C | 470.466061 | 0.15916679 | 0.09134219 | 1.74253308 | 0.08141521 |
| STR3 | YGL184C | 63.7330315 | -0.3230965 | 0.18541121 | -1.7425939 | 0.08140457 |
| DPB2 | YPR175W | 588.083569 | 0.14877123 | 0.08538962 | 1.74226367 | 0.08146232 |
| YDR010C | YDR010C | 7.75285896 | 0.51561793 | 0.29600086 | 1.74194743 | 0.08151764 |
| GIN4 | YDR507C | 2041.30154 | 0.09879504 | 0.05676206 | 1.74051206 | 0.08176914 |
| YJL202C | YJL202C | 5.52328104 | 0.50559215 | 0.29062589 | 1.73966659 | 0.08191758 |
| GID7 | YCL039W | 957.388917 | 0.12167838 | 0.07000748 | 1.73807683 | 0.08219728 |
| ULI1 | YFR026C | 54.870842 | 0.37253185 | 0.21451181 | 1.7366496 | 0.08244904 |
| MFT1 | YML062C | 406.023696 | 0.17238015 | 0.09926683 | 1.73653337 | 0.08246957 |
| ATC1 | YDR184C | 1329.6775 | -0.1275204 | 0.07344576 | -1.7362524 | 0.08251922 |
| PHO91 | YNR013C | 1307.85084 | 0.10873029 | 0.06263435 | 1.7359531 | 0.08257213 |
| YFL002W-A | YFL002W-A | 6.40506623 | -0.5133497 | 0.29577366 | -1.7356165 | 0.08263166 |
| LYS20 | YDL182W | 290.415184 | -0.1927171 | 0.11106896 | -1.7351121 | 0.08272094 |
| GDI1 | YER136W | 2565.14473 | 0.11191812 | 0.06450541 | 1.7350191 | 0.08273742 |
| AGX1 | YFL030W | 187.465137 | 0.21116547 | 0.12172244 | 1.73481137 | 0.08277422 |
| SNM1 | YDR478W | 147.225272 | -0.2411069 | 0.13904187 | -1.7340597 | 0.0829075 |
| SUM1 | YDR310C | 1469.39381 | 0.10773214 | 0.06213676 | 1.73379083 | 0.0829552 |
| YGR130C | YGR130C | 2086.78448 | 0.11231136 | 0.06479874 | 1.73323363 | 0.08305415 |
| Xu_2009_CUTs58 | Xu_2009_CU | 13.0398182 | 0.50239584 | 0.28989518 | 1.73302581 | 0.08309108 |
| Xu_2009_CUTs26 | Xu_2009_CU | 7.56723913 | 0.51294566 | 0.29624908 | 1.73146754 | 0.08336841 |
| CYK3 | YDL117W | 1371.5598 | 0.10836561 | 0.06259978 | 1.7310863 | 0.08343637 |
| OCA6 | YDR067C | 254.916249 | -0.2028382 | 0.11718183 | -1.7309699 | 0.08345713 |

| | | | | | | |
|---|---|---|---|---|---|---|
| BET1 | YIL004C | 326.872452 | 0.19827582 | 0.11461775 | 1.72988754 | 0.08365037 |
| NTH2 | YBR001C | 1034.77724 | -0.1562575 | 0.09036469 | -1.729188 | 0.08377545 |
| YLR444C | YLR444C | 4.56063362 | 0.49422752 | 0.28581462 | 1.72918908 | 0.08377526 |
| HST1 | YOL068C | 816.77229 | 0.1303622 | 0.07542381 | 1.72839582 | 0.08391728 |
| HIS6 | YIL020C | 160.063564 | 0.23860835 | 0.13828628 | 1.72546652 | 0.08444343 |
| RDR1 | YOR380W | 316.587543 | 0.1711767 | 0.09920443 | 1.72549458 | 0.08443838 |
| YNL195C | YNL195C | 82.4726786 | -0.3016884 | 0.17494011 | -1.7245237 | 0.08461335 |
| HXT14 | YNL318C | 22.8865893 | 0.45189802 | 0.26209081 | 1.724204 | 0.08467102 |
| MUP3 | YHL036W | 161.212999 | -0.2264338 | 0.13135934 | -1.7237742 | 0.08474861 |
| YLR412C-A | YLR412C-A | 9.97780552 | 0.50867436 | 0.29531199 | 1.72249815 | 0.08497932 |
| Xu_2009_SUTs49 | Xu_2009_SU | 23.5437449 | 0.44520001 | 0.25848716 | 1.72232934 | 0.08500988 |
| STP1 | YDR463W | 1819.1904 | 0.10320278 | 0.05993816 | 1.72182108 | 0.08510194 |
| MDM12 | YOL009C | 293.697723 | 0.1999787 | 0.11615753 | 1.7216163 | 0.08513905 |
| Xu_2009_SUTs19 | Xu_2009_SU | 38.5113574 | 0.41055066 | 0.23894663 | 1.71816885 | 0.08576582 |
| YPR027C | YPR027C | 7.45998398 | -0.5078081 | 0.29561645 | -1.7177939 | 0.08583422 |
| LRS4 | YDR439W | 147.641581 | 0.25076623 | 0.14599323 | 1.71765649 | 0.08585929 |
| Xu_2009_SUTs52 | Xu_2009_SU | 22.5652937 | 0.45989169 | 0.26803937 | 1.71576171 | 0.08620566 |
| YJL169W | YJL169W | 11.166451 | 0.49990952 | 0.29138175 | 1.71565143 | 0.08622586 |
| YRF1-4 | YLR466W | 19.9186734 | 0.45724808 | 0.26654611 | 1.71545588 | 0.08626168 |
| HNT3 | YOR258W | 117.680055 | -0.2666891 | 0.15546045 | -1.7154786 | 0.08625751 |
| SEC20 | YDR498C | 389.976291 | -0.1640894 | 0.09568636 | -1.7148677 | 0.08636949 |
| YML101C-A | YML101C-A | 10.4540996 | 0.50112893 | 0.2923297 | 1.71425937 | 0.0864811 |
| SLA2 | YNL243W | 2079.07432 | -0.0972994 | 0.05676112 | -1.7141911 | 0.08649363 |
| YMR141W-A | YMR141W-A | 0.69369053 | 0.27325981 | 0.1594533 | 1.71372938 | 0.08657844 |
| tD(GUC)J1 | tD(GUC)J1 | 5.61028334 | -0.4936285 | 0.28816249 | -1.7130215 | 0.08670859 |
| YIR030W-A | YIR030W-A | 8.03439384 | 0.50643776 | 0.29572486 | 1.71253021 | 0.086799 |
| Xu_2009_CUTs55 | Xu_2009_CU | 18.2377284 | 0.47429087 | 0.27739407 | 1.70980898 | 0.0873012 |
| SAM1 | YLR180W | 3948.95129 | 0.12636488 | 0.07392874 | 1.70927954 | 0.08739918 |
| RRI1 | YDL216C | 223.086005 | 0.22986532 | 0.13455273 | 1.70836605 | 0.08756844 |
| SYN8 | YAL014C | 431.358179 | -0.1531225 | 0.08964622 | -1.7080751 | 0.08762242 |
| WHI5 | YOR083W | 109.440053 | 0.26590494 | 0.15566992 | 1.70813303 | 0.08761166 |
| tV(CAC)D | tV(CAC)D | 12.7848 | -0.5010713 | 0.29341411 | -1.7077274 | 0.08768694 |
| NCE101 | YJL205C | 299.030306 | -0.2212978 | 0.12958445 | -1.7077497 | 0.0876828 |
| YDR102C | YDR102C | 36.3357808 | 0.42538327 | 0.249146 | 1.70736544 | 0.08775415 |
| Xu_2009_SUTs81 | Xu_2009_SU | 27.6848191 | 0.4195381 | 0.2458283 | 1.70663056 | 0.08789074 |
| YLR152C | YLR152C | 207.975315 | -0.2159404 | 0.1266116 | -1.7055338 | 0.08809491 |
| Xu_2009_CUTs45 | Xu_2009_CU | 31.8241917 | 0.42336931 | 0.24825545 | 1.70537766 | 0.08812401 |
| Xu_2009_SUTs62 | Xu_2009_SU | 34.3000618 | 0.39986518 | 0.23449812 | 1.70519564 | 0.08815794 |
| LSM3 | YLR438C-A | 728.100973 | 0.16282058 | 0.09549369 | 1.70504019 | 0.08818693 |
| SEC6 | YIL068C | 636.34018 | -0.1339666 | 0.07862938 | -1.7037732 | 0.08842347 |
| NPR3 | YHL023C | 993.318955 | 0.11744931 | 0.0689615 | 1.70311407 | 0.08854672 |

| | | | | | | |
|---|---|---|---|---|---|---|
| PPT1 | YGR123C | 815.420115 | -0.1412925 | 0.08306761 | -1.7009342 | 0.08895535 |
| PCK1 | YKR097W | 120.325918 | 0.26858516 | 0.15790162 | 1.70096524 | 0.08894951 |
| Xu_2009_SUTs82 | Xu_2009_SU | 45.4564799 | 0.38521939 | 0.22649625 | 1.70077601 | 0.08898506 |
| Xu_2009_SUTs70 | Xu_2009_SU | 16.338564 | 0.46832741 | 0.2756232 | 1.69915815 | 0.08928939 |
| Xu_2009_CUTs36 | Xu_2009_CU | 12.4427464 | 0.49156945 | 0.28961158 | 1.69734044 | 0.08963232 |
| RHO1 | YPR165W | 2907.82686 | -0.0921473 | 0.0543392 | -1.6957797 | 0.08992761 |
| Xu_2009_SUTs65 | Xu_2009_SU | 231.33939 | 0.19417267 | 0.1145687 | 1.69481427 | 0.09011066 |
| SCH9 | YHR205W | 1815.82814 | 0.10475401 | 0.06182876 | 1.69426037 | 0.09021582 |
| YML057C-A | YML057C-A | 5.96144858 | 0.49769172 | 0.2937709 | 1.69414913 | 0.09023695 |
| AKR1 | YDR264C | 2109.26504 | -0.1084507 | 0.06402515 | -1.6938759 | 0.09028887 |
| SKG1 | YKR100C | 648.737575 | 0.14833652 | 0.0875734 | 1.69385368 | 0.09029309 |
| LRP1 | YHR081W | 437.247592 | 0.1886448 | 0.11138336 | 1.69365341 | 0.09033116 |
| Xu_2009_CUTs21 | Xu_2009_CU | 9.00252711 | 0.50139609 | 0.29610446 | 1.69330815 | 0.09039683 |
| YLR317W | YLR317W | 59.5542019 | 0.32165761 | 0.1900635 | 1.69236921 | 0.0905756 |
| YKL106C-A | YKL106C-A | 10.833133 | 0.49832663 | 0.29457011 | 1.69170804 | 0.09070165 |
| LCL1 | YPL056C | 107.504473 | 0.26525775 | 0.15683969 | 1.69126678 | 0.09078586 |
| SNR24 | snR24 | 51.7258901 | -0.3561733 | 0.21065605 | -1.6907811 | 0.09087862 |
| IPF1 | YDL176W | 448.664161 | 0.15747231 | 0.09316601 | 1.69023339 | 0.09098331 |
| ACL4 | YDR161W | 775.978909 | -0.1416388 | 0.08381585 | -1.6898805 | 0.09105083 |
| Xu_2009_CUTs24 | Xu_2009_CU | 8.75219294 | -0.5011857 | 0.29667058 | -1.6893676 | 0.09114901 |
| THI11 | YJR156C | 72.4603236 | -0.3092102 | 0.18315623 | -1.6882318 | 0.09136675 |
| Xu_2009_CUTs56 | Xu_2009_CU | 72.5378192 | 0.35273082 | 0.20895897 | 1.6880386 | 0.09140382 |
| YNR029C | YNR029C | 599.978181 | 0.1449551 | 0.08599709 | 1.6855815 | 0.09187643 |
| YIL163C | YIL163C | 18.2821886 | 0.46874837 | 0.27813715 | 1.68531375 | 0.09192805 |
| tN(GUU)K | tN(GUU)K | 8.91107504 | -0.4998618 | 0.29667821 | -1.6848619 | 0.09201521 |
| SAC3 | YDR159W | 1662.88181 | 0.10335868 | 0.06139988 | 1.68336937 | 0.09230361 |
| TPH3 | YJL016W | 1734.318 | 0.1104568 | 0.0656188 | 1.68331042 | 0.09231501 |
| POP2 | YNR052C | 1452.14618 | -0.1147252 | 0.06817546 | -1.682793 | 0.09241517 |
| YOR331C | YOR331C | 4.84631636 | 0.47975465 | 0.28509613 | 1.68278205 | 0.09241729 |
| EMC1 | YCL045C | 2485.17663 | -0.0943486 | 0.05609178 | -1.682039 | 0.09256128 |
| ZRT1 | YGL255W | 3136.5103 | -0.298483 | 0.17744518 | -1.6821141 | 0.09254671 |
| BSC4 | YNL269W | 9.57954228 | 0.49805846 | 0.29612526 | 1.68191819 | 0.0925847 |
| Xu_2009_CUTs59 | Xu_2009_CU | 23.4579715 | 0.42896969 | 0.25513613 | 1.68133653 | 0.09269757 |
| Xu_2009_SUTs20 | Xu_2009_SU | 24.897155 | 0.43885034 | 0.26132837 | 1.67930618 | 0.09309239 |
| PAU6 | YNR076W | 56.1763456 | 0.34468514 | 0.20533632 | 1.67863699 | 0.09322281 |
| Xu_2009_SUTs21 | Xu_2009_SU | 23.6671721 | 0.43394379 | 0.25861116 | 1.67797785 | 0.09335142 |
| ADY4 | YLR227C | 212.760046 | 0.20046438 | 0.11948878 | 1.67768368 | 0.09340887 |
| PET494 | YNR045W | 445.218525 | -0.1497708 | 0.08935402 | -1.6761508 | 0.09370866 |
| YOR365C | YOR365C | 9.87170052 | 0.49492492 | 0.29531139 | 1.67594252 | 0.09374945 |
| Xu_2009_CUTs39 | Xu_2009_CU | 26.3498524 | 0.4220659 | 0.25186599 | 1.67575586 | 0.09378602 |
| Xu_2009_CUTs43 | Xu_2009_CU | 126.333681 | 0.27865427 | 0.16635821 | 1.67502567 | 0.09392919 |

| | | | | | | |
|---|---|---|---|---|---|---|
| Xu_2009_CUTs90 | Xu_2009_CU | 30.9671954 | 0.3981863 | 0.23784575 | 1.67413667 | 0.09410374 |
| Xu_2009_SUTs14 | Xu_2009_SU | 144.670097 | 0.2423135 | 0.14480151 | 1.67341836 | 0.09424497 |
| PDX1 | YGR193C | 794.445105 | -0.1262293 | 0.07546086 | -1.6727783 | 0.09437094 |
| YCL048W-A | YCL048W-A | 48.2386812 | -0.3714691 | 0.22223677 | -1.6715016 | 0.09462265 |
| MTR4 | YJL050W | 2162.64502 | 0.11809705 | 0.07065168 | 1.67153916 | 0.09461523 |
| Xu_2009_CUTs18 | Xu_2009_CU | 215.035821 | -0.2034233 | 0.12172441 | -1.6711794 | 0.09468625 |
| Xu_2009_SUTs42 | Xu_2009_SU | 64.0307746 | 0.31172173 | 0.18658541 | 1.67066508 | 0.09478785 |
| Xu_2009_SUTs53 | Xu_2009_SU | 24.9634048 | 0.43357327 | 0.25955292 | 1.67046193 | 0.09482801 |
| HAA1 | YPR008W | 2287.0146 | 0.09789477 | 0.05860259 | 1.67048528 | 0.09482339 |
| DNF1 | YER166W | 2887.13496 | 0.08797251 | 0.05268087 | 1.66991377 | 0.09493643 |
| Xu_2009_SUTs49 | Xu_2009_SU | 45.107617 | 0.37170177 | 0.2227147 | 1.66895929 | 0.09512545 |
| YOR108C-A | YOR108C-A | 7.12888709 | 0.49380213 | 0.29604849 | 1.66797722 | 0.09532025 |
| YAL066W | YAL066W | 2.18133357 | 0.39646735 | 0.23786267 | 1.66679094 | 0.09555598 |
| Xu_2009_SUTs12 | Xu_2009_SU | 20.7055096 | 0.44323172 | 0.26601081 | 1.66621697 | 0.09567021 |
| Xu_2009_SUTs43 | Xu_2009_SU | 44.3983338 | 0.36117402 | 0.21677185 | 1.66614819 | 0.0956839 |
| ZAP1 | YJL056C | 1375.81399 | 0.12055041 | 0.07235212 | 1.6661627 | 0.09568101 |
| SPO12 | YHR152W | 904.172395 | 0.12007939 | 0.07209745 | 1.66551513 | 0.09581003 |
| Xu_2009_SUTs60 | Xu_2009_SU | 286.834798 | 0.17947075 | 0.10780256 | 1.66480981 | 0.09595071 |
| VNX1 | YNL321W | 1120.4581 | 0.11826317 | 0.07107483 | 1.66392474 | 0.09612747 |
| RMD1 | YDL001W | 552.145789 | 0.15776298 | 0.09485272 | 1.66324145 | 0.09626411 |
| Xu_2009_CUTs31 | Xu_2009_CU | 14.8645513 | 0.47317811 | 0.28452035 | 1.66307297 | 0.09629783 |
| Xu_2009_CUTs15 | Xu_2009_CU | 10.2360163 | 0.48901634 | 0.29416777 | 1.66237227 | 0.09643815 |
| Xu_2009_CUTs39 | Xu_2009_CU | 19.8457349 | 0.45233112 | 0.27219085 | 1.66181604 | 0.09654967 |
| TWF1 | YGR080W | 713.07857 | -0.144925 | 0.08724717 | -1.661085 | 0.09669638 |
| BRO1 | YPL084W | 1379.79588 | 0.11087921 | 0.0667765 | 1.66045262 | 0.09682343 |
| tG(CCC)O/SUF5 | tG(CCC)O | 2.14750056 | 0.39945066 | 0.24061797 | 1.6601032 | 0.09689369 |
| tQ(UUG)H | tQ(UUG)H | 7.47205846 | -0.4900243 | 0.29522597 | -1.659828 | 0.09694906 |
| PET191 | YJR034W | 172.374153 | -0.2385704 | 0.14375478 | -1.6595652 | 0.09700195 |
| SMD2 | YLR275W | 383.287108 | 0.15498234 | 0.09344341 | 1.65856904 | 0.09720267 |
| SNR13 | snR13 | 659.086527 | -0.3326111 | 0.20058783 | -1.658182 | 0.09728073 |
| TDA5 | YLR426W | 329.065179 | -0.165125 | 0.09967564 | -1.6566237 | 0.09759558 |
| YER187W | YER187W | 40.5167674 | 0.37995316 | 0.22947245 | 1.65576806 | 0.0977688 |
| YAL063C-A | YAL063C-A | 17.079161 | 0.45997958 | 0.27792692 | 1.65503789 | 0.09791681 |
| Xu_2009_SUTs47 | Xu_2009_SU | 9.21107311 | 0.48937778 | 0.29573937 | 1.65476031 | 0.09797313 |
| Xu_2009_CUTs13 | Xu_2009_CU | 19.5250877 | 0.44644953 | 0.26990916 | 1.65407328 | 0.09811263 |
| YCF1 | YDR135C | 3646.90258 | 0.08235642 | 0.04978863 | 1.65412088 | 0.09810295 |
| BNI5 | YNL166C | 692.373813 | -0.1310049 | 0.07923427 | -1.6533873 | 0.09825207 |
| Xu_2009_CUTs51 | Xu_2009_CU | 91.9185446 | -0.2839799 | 0.17191387 | -1.6518729 | 0.09856046 |
| Xu_2009_SUTs52 | Xu_2009_SU | 5.61389723 | -0.4794038 | 0.29022233 | -1.6518502 | 0.09856509 |
| TAF3 | YPL011C | 873.271677 | -0.1207194 | 0.07309134 | -1.6516233 | 0.09861137 |
| YOR082C | YOR082C | 4.26632711 | 0.45970792 | 0.27836401 | 1.65146319 | 0.09864403 |

| | | | | | | |
|---|---|---|---|---|---|---|
| NMA2 | YGR010W | 340.558442 | -0.1658509 | 0.10044188 | -1.6512131 | 0.09869506 |
| COX26 | YDR119W-A | 58.060285 | -0.334533 | 0.20271936 | -1.650227 | 0.09889651 |
| SPO1 | YNL012W | 44.5036894 | -0.3787452 | 0.22973824 | -1.6485944 | 0.09923077 |
| SNR45 | snR45 | 1854.55524 | -0.2444917 | 0.14836915 | -1.6478609 | 0.09938122 |
| Xu_2009_CUTs45 | Xu_2009_CU | 14.9113048 | 0.47022834 | 0.28538723 | 1.64768526 | 0.09941727 |
| Xu_2009_CUTs74 | Xu_2009_CU | 15.7161666 | 0.46176152 | 0.28013483 | 1.64835452 | 0.09927994 |
| Xu_2009_SUTs66 | Xu_2009_SU | 41.226257 | 0.3698697 | 0.22446152 | 1.64780896 | 0.09939188 |
| SHG1 | YBR258C | 219.673158 | 0.21605449 | 0.13110586 | 1.64793919 | 0.09936515 |
| MAG1 | YER142C | 267.464775 | -0.1777372 | 0.10786531 | -1.6477699 | 0.0993999 |
| NUP57 | YGR119C | 1518.41914 | 0.10242613 | 0.06215662 | 1.64787148 | 0.09937905 |
| FBP1 | YLR377C | 25.7830592 | -0.4371527 | 0.26527782 | -1.6479053 | 0.0993721 |
| YPL034W | YPL034W | 160.902609 | -0.247066 | 0.14988853 | -1.6483316 | 0.09928465 |
| TFC3 | YAL001C | 1021.57398 | 0.12827906 | 0.07786152 | 1.64752826 | 0.09944951 |
| ULP2 | YIL031W | 990.707313 | 0.11165665 | 0.06779299 | 1.64702366 | 0.09955318 |
| HPC2 | YBR215W | 657.195406 | 0.1312994 | 0.07972525 | 1.64689864 | 0.09957888 |
| TAF11 | YML015C | 729.161566 | -0.1245895 | 0.07565978 | -1.646707 | 0.09961829 |
| Xu_2009_SUTs65 | Xu_2009_SU | 132.509926 | -0.2359911 | 0.1433384 | -1.6463916 | 0.09968317 |
| SSH4 | YKL124W | 508.845565 | 0.14483978 | 0.08797565 | 1.64636213 | 0.09968923 |
| Xu_2009_CUTs60 | Xu_2009_CU | 23.5314872 | 0.41922944 | 0.25480141 | 1.6453184 | 0.09990417 |
| DCD1 | YHR144C | 329.578601 | -0.1698273 | 0.10333244 | -1.6435043 | 0.10027864 |
| ATG36 | YJL185C | 138.280917 | -0.2267413 | 0.13796543 | -1.6434647 | 0.10028681 |
| MIC60 | YKR016W | 989.003431 | -0.109763 | 0.06684025 | -1.642169 | 0.10055499 |
| Xu_2009_SUTs67 | Xu_2009_SU | 27.9543417 | 0.40725154 | 0.2480943 | 1.64151914 | 0.1006897 |
| SUL2 | YLR092W | 245.419359 | 0.1839843 | 0.11208776 | 1.64143074 | 0.10070803 |
| AFR1 | YDR085C | 816.252548 | 0.12658037 | 0.07713566 | 1.64100978 | 0.10079539 |
| TRM7 | YBR061C | 469.04438 | -0.1425943 | 0.08701198 | -1.6387889 | 0.10125722 |
| Xu_2009_CUTs75 | Xu_2009_CU | 0.91917759 | -0.2779765 | 0.16963583 | -1.6386661 | 0.10128281 |
| RKM3 | YBR030W | 534.015871 | -0.1509553 | 0.09224044 | -1.6365409 | 0.10172643 |
| CGR1 | YGL029W | 613.489894 | 0.15673964 | 0.09577874 | 1.63647629 | 0.10173994 |
| RPN2 | YIL075C | 6550.33128 | 0.07479786 | 0.04570291 | 1.63661051 | 0.10171188 |
| Xu_2009_CUTs67 | Xu_2009_CU | 16.3959183 | 0.46892932 | 0.2866557 | 1.63586255 | 0.10186836 |
| YFT2 | YDR319C | 260.880695 | 0.18477584 | 0.11304108 | 1.63459019 | 0.10213499 |
| Xu_2009_SUTs37 | Xu_2009_SU | 27.1869747 | 0.406446 | 0.24869908 | 1.63428832 | 0.10219832 |
| RNR3 | YIL066C | 109.657651 | -0.2584844 | 0.15817557 | -1.6341616 | 0.10222491 |
| QNS1 | YHR074W | 1314.64916 | -0.1039853 | 0.06364289 | -1.6338878 | 0.1022824 |
| YJL022W | YJL022W | 9.93434671 | 0.48411139 | 0.29638863 | 1.63336695 | 0.10239184 |
| Xu_2009_CUTs21 | Xu_2009_CU | 5.43517578 | 0.47637047 | 0.29172456 | 1.63294604 | 0.10248035 |
| YAT1 | YAR035W | 84.7704691 | -0.2877564 | 0.17624791 | -1.6326799 | 0.10253634 |
| MAD3 | YJL013C | 391.608418 | 0.1637255 | 0.10030001 | 1.63235772 | 0.10260415 |
| VIK1 | YPL253C | 220.581068 | -0.1883672 | 0.11549955 | -1.6308915 | 0.10291322 |
| YME1 | YPR024W | 2155.78342 | 0.09635061 | 0.05907779 | 1.63091091 | 0.10290912 |

| | | | | | | |
|---|---|---|---|---|---|---|
| Xu_2009_CUTs85 | Xu_2009_CU | 27.0276199 | 0.40389427 | 0.24773378 | 1.63035608 | 0.10302626 |
| PAF1 | YBR279W | 1865.67568 | 0.09948989 | 0.06103665 | 1.63000236 | 0.103101 |
| RPM2 | YML091C | 893.849188 | -0.1266924 | 0.07772289 | -1.6300528 | 0.10309034 |
| YGR045C | YGR045C | 13.422868 | 0.47345402 | 0.29051153 | 1.62972539 | 0.10315955 |
| Xu_2009_SUTs78 | Xu_2009_SU | 53.5985642 | -0.3521086 | 0.21613945 | -1.6290805 | 0.10329598 |
| Xu_2009_CUTs32 | Xu_2009_CU | 4.20821133 | -0.4524072 | 0.27778025 | -1.6286513 | 0.10338685 |
| Xu_2009_CUTs46 | Xu_2009_CU | 13.2984234 | 0.46556644 | 0.28585337 | 1.62868973 | 0.10337872 |
| CAB5 | YDR196C | 276.207325 | -0.1782807 | 0.10946891 | -1.6285967 | 0.10339842 |
| OMS1 | YDR316W | 409.12832 | 0.14881215 | 0.09147746 | 1.62676302 | 0.10378744 |
| Xu_2009_CUTs70 | Xu_2009_CU | 17.6574617 | 0.44482459 | 0.27367948 | 1.62534871 | 0.10408828 |
| IRC2 | YDR112W | 3.25249714 | 0.43488282 | 0.26761811 | 1.62501265 | 0.10415986 |
| YNL146W | YNL146W | 99.5800335 | -0.2535112 | 0.15601395 | -1.6249268 | 0.10417816 |
| Xu_2009_CUTs15 | Xu_2009_CU | 6.66186178 | 0.47923992 | 0.29495059 | 1.62481426 | 0.10420214 |
| DID4 | YKL002W | 593.582666 | -0.1462891 | 0.09014008 | -1.622909 | 0.10460887 |
| Xu_2009_CUTs60 | Xu_2009_CU | 47.4627149 | 0.34924229 | 0.21521545 | 1.62275662 | 0.10464144 |
| Xu_2009_SUTs16 | Xu_2009_SU | 47.4655052 | 0.33646547 | 0.20739292 | 1.62235752 | 0.10472682 |
| PRP6 | YBR055C | 980.234048 | 0.11664256 | 0.07190044 | 1.62227885 | 0.10474365 |
| WHI3 | YNL197C | 1738.27585 | 0.11616404 | 0.07160694 | 1.62224565 | 0.10475076 |
| PET18 | YCR020C | 77.3631706 | -0.2902307 | 0.17893786 | -1.6219634 | 0.10481118 |
| MAC1 | YMR021C | 482.382079 | -0.1377319 | 0.08493445 | -1.6216255 | 0.10488356 |
| VPH1 | YOR270C | 4209.61895 | -0.0912895 | 0.05631146 | -1.6211539 | 0.10498464 |
| SOA1 | YIL166C | 83.4958747 | -0.2713114 | 0.16740879 | -1.6206521 | 0.10509228 |
| FMP25 | YLR077W | 851.044664 | -0.1252425 | 0.07728029 | -1.620627 | 0.10509767 |
| tS(AGA)E | tS(AGA)E | 3.33937049 | 0.43591197 | 0.26904089 | 1.62024433 | 0.1051798 |
| MEP3 | YPR138C | 575.765724 | -0.1371062 | 0.08463829 | -1.6199077 | 0.10525211 |
| CTF8 | YHR191C | 394.044367 | 0.15738306 | 0.09720823 | 1.61903024 | 0.10544076 |
| Xu_2009_CUTs19 | Xu_2009_CU | 12.841418 | 0.47247146 | 0.2918681 | 1.61878417 | 0.10549371 |
| YSP2 | YDR326C | 2159.92965 | 0.08780446 | 0.05423936 | 1.6188329 | 0.10548322 |
| Xu_2009_CUTs23 | Xu_2009_CU | 2.30222431 | -0.395641 | 0.24453117 | -1.6179575 | 0.10567175 |
| YDL094C | YDL094C | 13.360939 | 0.46493992 | 0.28736458 | 1.61794442 | 0.10567458 |
| RKR1 | YMR247C | 1537.86822 | 0.10295929 | 0.06364269 | 1.61777089 | 0.10571198 |
| ERD2 | YBL040C | 610.218371 | -0.1347988 | 0.08333397 | -1.6175732 | 0.10575461 |
| Xu_2009_CUTs66 | Xu_2009_CU | 28.7112443 | 0.3989921 | 0.24671399 | 1.61722527 | 0.10582966 |
| Xu_2009_CUTs41 | Xu_2009_CU | 11.5356379 | 0.4738136 | 0.29321996 | 1.61589821 | 0.10611632 |
| SEC39 | YLR440C | 600.008075 | -0.1308791 | 0.08111488 | -1.6135033 | 0.10663521 |
| FBP26 | YJL155C | 760.170222 | -0.1382502 | 0.08573745 | -1.6124829 | 0.10685691 |
| YGR160W | YGR160W | 2.06752273 | 0.38506267 | 0.23884178 | 1.61220818 | 0.10691665 |
| IRC10 | YOL015W | 134.77064 | 0.23652047 | 0.14675104 | 1.61171245 | 0.10702453 |
| tD(GUC)O | tD(GUC)O | 2.41592737 | -0.3988575 | 0.24769133 | -1.6103007 | 0.10733222 |
| Xu_2009_SUTs13 | Xu_2009_SU | 76.279939 | 0.30928168 | 0.19207928 | 1.61017722 | 0.10735917 |
| Xu_2009_SUTs78 | Xu_2009_SU | 58.589689 | 0.31440871 | 0.19531209 | 1.609776 | 0.10744677 |

| | | | | | | |
|---|---|---|---|---|---|---|
| GTR2 | YGR163W | 604.135547 | 0.13092677 | 0.08136324 | 1.60916363 | 0.10758057 |
| Xu_2009_SUTs22 | Xu_2009_SU | 105.079947 | 0.2676964 | 0.16659808 | 1.60683969 | 0.10808954 |
| GIS3 | YLR094C | 1196.09764 | -0.105656 | 0.06576963 | -1.6064557 | 0.10817382 |
| YPT10 | YBR264C | 285.973639 | 0.17713388 | 0.11028656 | 1.60612394 | 0.10824668 |
| SNF5 | YBR289W | 988.187337 | 0.11270921 | 0.07019037 | 1.60576475 | 0.10832561 |
| RSE1 | YML049C | 1787.68292 | 0.09821671 | 0.06117723 | 1.60544554 | 0.10839579 |
| RIP1 | YEL024W | 790.140252 | -0.1584444 | 0.09872719 | -1.6048709 | 0.10852222 |
| YIL071W-A | YIL071W-A | 4.83048342 | -0.4634289 | 0.28877594 | -1.6048044 | 0.10853685 |
| SWI6 | YLR182W | 795.618037 | 0.12321674 | 0.07682417 | 1.60387995 | 0.10874052 |
| MDM31 | YHR194W | 514.671335 | 0.13547714 | 0.08450503 | 1.60318438 | 0.10889395 |
| FEX2 | YPL279C | 221.872715 | 0.19924761 | 0.12433489 | 1.60250767 | 0.10904339 |
| TAT2 | YOL020W | 1679.97382 | 0.10159297 | 0.0634129 | 1.60208679 | 0.10913642 |
| SEC1 | YDR164C | 728.358793 | 0.12662599 | 0.07908498 | 1.60113828 | 0.1093463 |
| Xu_2009_CUTs62 | Xu_2009_CU | 17.9508494 | 0.43591056 | 0.27251094 | 1.59960759 | 0.10968566 |
| YAR064W | YAR064W | 0.80203153 | 0.2708831 | 0.16943519 | 1.59874173 | 0.109878 |
| Xu_2009_SUTs69 | Xu_2009_SU | 19.7262234 | 0.42716687 | 0.26727726 | 1.59821625 | 0.10999486 |
| Xu_2009_SUTs19 | Xu_2009_SU | 27.3728555 | 0.39693704 | 0.24844176 | 1.59770664 | 0.11010828 |
| RIM11 | YMR139W | 369.152172 | -0.1607819 | 0.10063262 | -1.5977112 | 0.11010727 |
| YML018C | YML018C | 840.61904 | -0.1226747 | 0.07678825 | -1.5975707 | 0.11013856 |
| Xu_2009_SUTs53 | Xu_2009_SU | 29.6817783 | 0.40265613 | 0.25210278 | 1.59719033 | 0.11022329 |
| PAN2 | YGL094C | 1511.33772 | 0.10464226 | 0.06552444 | 1.59699587 | 0.11026663 |
| SNR50 | snR50 | 70.785858 | -0.3071177 | 0.1924553 | -1.5957873 | 0.1105363 |
| ICS3 | YJL077C | 1.83794471 | 0.36028616 | 0.2257962 | 1.59562544 | 0.11057244 |
| ATG3 | YNR007C | 269.462222 | -0.1684404 | 0.10557134 | -1.5955127 | 0.11059764 |
| GAL1 | YBR020W | 87.4541549 | 0.27365477 | 0.17165765 | 1.59418918 | 0.11089367 |
| ECM10 | YEL030W | 173.837287 | 0.21548225 | 0.13521756 | 1.59359659 | 0.11102642 |
| YDR194W-A | YDR194W-A | 2.20405273 | 0.38724222 | 0.24303442 | 1.59336374 | 0.11107862 |
| SHE9 | YDR393W | 549.510042 | 0.14044291 | 0.08814343 | 1.59334519 | 0.11108278 |
| NIP1 | YMR309C | 4736.33035 | -0.1182326 | 0.07424565 | -1.5924521 | 0.11128317 |
| FSH1 | YHR049W | 2752.29074 | -0.1135632 | 0.07135149 | -1.5916027 | 0.111474 |
| CLG1 | YGL215W | 4995.87677 | 0.08336358 | 0.05241632 | 1.5904128 | 0.11174179 |
| SRT1 | YMR101C | 49.5064531 | -0.3371101 | 0.21198101 | -1.5902845 | 0.1117707 |
| tP(UGG)M/SUF7 | tP(UGG)M | 23.173674 | -0.4239453 | 0.26661235 | -1.5901189 | 0.111808 |
| YEL028W | YEL028W | 109.17796 | 0.25828148 | 0.16243575 | 1.59005317 | 0.11182282 |
| POL3 | YDL102W | 835.07843 | 0.11924904 | 0.07502983 | 1.58935502 | 0.11198026 |
| RED1 | YLR263W | 301.746369 | 0.16191327 | 0.10187333 | 1.58935875 | 0.11197942 |
| PNS1 | YOR161C | 393.487783 | -0.1721562 | 0.10831212 | -1.5894458 | 0.11195979 |
| tG(GCC)E | tG(GCC)E | 2.82771776 | -0.4119723 | 0.25929671 | -1.5888065 | 0.1121041 |
| Xu_2009_SUTs62 | Xu_2009_SU | 73.0713796 | 0.30780414 | 0.19400075 | 1.58661317 | 0.11260028 |
| HEM25 | YDL119C | 388.340975 | -0.1515582 | 0.09552649 | -1.5865571 | 0.112613 |
| ECM25 | YJL201W | 656.384199 | -0.1212923 | 0.07647049 | -1.5861318 | 0.11270941 |

| | | | | | | |
|---|---|---|---|---|---|---|
| PHD1 | YKL043W | 1138.85155 | -0.1270072 | 0.08008461 | -1.5859129 | 0.11275906 |
| YMR158W-B | YMR158W-B | 1.58865154 | 0.3514171 | 0.22165201 | 1.58544515 | 0.11286523 |
| TRM44 | YPL030W | 672.422207 | -0.1306321 | 0.08239216 | -1.5854923 | 0.11285452 |
| POP8 | YBL018C | 218.11916 | 0.20181672 | 0.12730253 | 1.58533157 | 0.11289102 |
| IFA38 | YBR159W | 1154.25302 | -0.1069844 | 0.06753593 | -1.5841103 | 0.11316863 |
| MGR3 | YMR115W | 680.182839 | -0.1175575 | 0.0743004 | -1.582192 | 0.11360575 |
| KAP123 | YER110C | 3978.5768 | -0.0887044 | 0.05611483 | -1.5807661 | 0.11393153 |
| YGR068W-A | YGR068W-A | 0.82812676 | 0.27079313 | 0.17129518 | 1.5808567 | 0.11391081 |
| YNL095C | YNL095C | 327.286101 | 0.170082 | 0.10759946 | 1.58069563 | 0.11394765 |
| Xu_2009_CUTs27 | Xu_2009_CU | 6.12957814 | 0.4612315 | 0.29198073 | 1.57966415 | 0.1141838 |
| Xu_2009_SUTs11 | Xu_2009_SU | 72.8317928 | -0.2842608 | 0.17995962 | -1.5795808 | 0.1142029 |
| AAC1 | YMR056C | 176.874655 | 0.20708927 | 0.13118593 | 1.57859362 | 0.1144293 |
| FUN14 | YAL008W | 416.921727 | -0.1738721 | 0.11015898 | -1.5783744 | 0.11447963 |
| GAS2 | YLR343W | 257.743684 | 0.17498828 | 0.11089476 | 1.5779671 | 0.11457317 |
| YDL007C-A | YDL007C-A | 3.69870241 | 0.42804518 | 0.27150416 | 1.57656952 | 0.11489461 |
| YNL120C | YNL120C | 2.0550403 | 0.38339725 | 0.24329039 | 1.57588326 | 0.11505271 |
| YMR114C | YMR114C | 385.700635 | 0.16037138 | 0.10179837 | 1.57538249 | 0.11516818 |
| tE(UUC)E2 | tE(UUC)E2 | 9.701757 | -0.4672199 | 0.29668381 | -1.5748074 | 0.1153009 |
| AIM25 | YJR100C | 348.833305 | -0.1600698 | 0.10168056 | -1.5742418 | 0.11543156 |
| Xu_2009_SUTs66 | Xu_2009_SU | 35.2949674 | 0.36096314 | 0.22933129 | 1.5739812 | 0.1154918 |
| FMP48 | YGR052W | 103.322207 | -0.2670944 | 0.16969273 | -1.5739883 | 0.11549016 |
| Xu_2009_SUTs48 | Xu_2009_SU | 72.8825854 | 0.27887623 | 0.17728555 | 1.57303414 | 0.11571091 |
| ORC2 | YBR060C | 710.59662 | 0.1257375 | 0.07999116 | 1.57189259 | 0.11597546 |
| USB1 | YLR132C | 532.480716 | -0.1354909 | 0.08619685 | -1.5718776 | 0.11597894 |
| GIR2 | YDR152W | 1096.88404 | -0.1104996 | 0.07030297 | -1.5717623 | 0.11600569 |
| RME1 | YGR044C | 858.044384 | -0.1254804 | 0.07990038 | -1.5704609 | 0.11630791 |
| YMR001C-A | YMR001C-A | 10.7215034 | 0.46027641 | 0.29317245 | 1.56998522 | 0.11641855 |
| MLF3 | YNL074C | 1682.00743 | 0.10152954 | 0.06468291 | 1.56965015 | 0.11649653 |
| tV(AAC)G3 | tV(AAC)G3 | 13.4327892 | -0.449733 | 0.28659417 | -1.569233 | 0.11659367 |
| SIP2 | YGL208W | 427.489885 | -0.1608523 | 0.10249888 | -1.5693082 | 0.11657614 |
| NDC80 | YIL144W | 797.623806 | 0.11619838 | 0.07406191 | 1.56893568 | 0.11666293 |
| YCL021W-A | YCL021W-A | 288.757173 | 0.18697349 | 0.11919987 | 1.56857126 | 0.11674787 |
| YPR174C | YPR174C | 319.016784 | 0.16766869 | 0.106912 | 1.56828689 | 0.11681419 |
| Xu_2009_SUTs85 | Xu_2009_SU | 52.701592 | 0.32007438 | 0.20430194 | 1.56667318 | 0.1171911 |
| POL12 | YBL035C | 344.2877 | 0.14986955 | 0.09567098 | 1.56651006 | 0.11722926 |
| tN(GUU)N2 | tN(GUU)N2 | 1.13193693 | 0.31195798 | 0.19918721 | 1.56615468 | 0.11731241 |
| PBY1 | YBR094W | 826.731527 | 0.11163025 | 0.07128566 | 1.56595665 | 0.11735877 |
| YGL230C | YGL230C | 15.9441428 | -0.4446264 | 0.28402727 | -1.5654356 | 0.1174808 |
| PTP3 | YER075C | 1265.52192 | 0.11038327 | 0.07052729 | 1.56511426 | 0.11755612 |
| tE(UUC)G3 | tE(UUC)G3 | 3.4504507 | -0.4251475 | 0.2717533 | -1.5644613 | 0.11770929 |
| YGL088W | YGL088W | 7.53093571 | -0.463792 | 0.29644027 | -1.5645377 | 0.11769136 |

| | | | | | | |
|---|---|---|---|---|---|---|
| ACF4 | YJR083C | 282.83772 | -0.1630217 | 0.10420267 | -1.564468 | 0.11770769 |
| PUS1 | YPL212C | 1183.85723 | 0.12985495 | 0.08300717 | 1.56438238 | 0.1177278 |
| HDA2 | YDR295C | 625.437625 | -0.1250937 | 0.08010635 | -1.5615958 | 0.11838323 |
| tA(AGC)H | tA(AGC)H | 7.23496743 | -0.4612141 | 0.29578179 | -1.5593052 | 0.11892415 |
| PAU2 | YEL049W | 31.5309429 | 0.39139369 | 0.25137936 | 1.55698418 | 0.11947424 |
| NPA3 | YJR072C | 1221.52235 | -0.10766 | 0.06914727 | -1.5569671 | 0.11947829 |
| GSH1 | YJL101C | 1175.2511 | 0.10848098 | 0.06969327 | 1.55654885 | 0.11957764 |
| ATR1 | YML116W | 483.2303 | -0.1343338 | 0.08635699 | -1.5555642 | 0.11981176 |
| TFC8 | YPL007C | 260.816411 | -0.171581 | 0.11040699 | -1.5540768 | 0.12016609 |
| CHS7 | YHR142W | 534.626894 | -0.1273707 | 0.08199177 | -1.5534575 | 0.12031388 |
| GPI13 | YLL031C | 1499.47924 | 0.10780383 | 0.06939491 | 1.55348319 | 0.12030774 |
| Xu_2009_CUTs59 | Xu_2009_CU | 3.58337535 | 0.42037298 | 0.27064343 | 1.55323548 | 0.12036689 |
| MPC54 | YOR177C | 157.705461 | 0.21089829 | 0.1358398 | 1.55255156 | 0.12053031 |
| YCH1 | YGR203W | 201.318946 | -0.1881259 | 0.12118654 | -1.5523667 | 0.12057451 |
| CTK3 | YML112W | 535.508618 | 0.13485693 | 0.08689645 | 1.55192679 | 0.12067974 |
| MED8 | YBR193C | 308.401717 | -0.1587802 | 0.10235693 | -1.5512404 | 0.12084408 |
| Xu_2009_SUTs49 | Xu_2009_SU | 10.663775 | 0.45477366 | 0.29341785 | 1.54991819 | 0.12116115 |
| STP22 | YCL008C | 630.78445 | -0.1204811 | 0.07772645 | -1.5500652 | 0.12112587 |
| PML1 | YLR016C | 375.237376 | -0.152171 | 0.09818337 | -1.5498649 | 0.12117395 |
| IES4 | YOR189W | 415.903489 | 0.14269258 | 0.09206322 | 1.54994121 | 0.12115563 |
| Xu_2009_SUTs59 | Xu_2009_SU | 221.33374 | 0.17692168 | 0.11416105 | 1.54975521 | 0.12120028 |
| YGL177W | YGL177W | 137.108673 | 0.21164125 | 0.13663028 | 1.54900686 | 0.12138007 |
| SAS5 | YOR213C | 439.033983 | 0.15518562 | 0.10021809 | 1.54847916 | 0.12150698 |
| YDL199C | YDL199C | 194.827423 | -0.1967152 | 0.1271165 | -1.5475188 | 0.12173821 |
| Xu_2009_CUTs75 | Xu_2009_CU | 10.9890046 | 0.45350541 | 0.29323917 | 1.54653764 | 0.12197478 |
| GCN5 | YGR252W | 488.42822 | 0.13502516 | 0.08732654 | 1.54620981 | 0.1220539 |
| Xu_2009_SUTs56 | Xu_2009_SU | 186.236555 | 0.20556844 | 0.13299487 | 1.545687 | 0.12218018 |
| YDL177C | YDL177C | 292.646805 | 0.16516982 | 0.10689083 | 1.54521973 | 0.12229312 |
| SKG3 | YLR187W | 890.600606 | -0.1049095 | 0.06791016 | -1.5448274 | 0.12238802 |
| Xu_2009_CUTs34 | Xu_2009_CU | 141.153375 | 0.20906269 | 0.13539931 | 1.54404548 | 0.12257732 |
| BBP1 | YPL255W | 478.099857 | -0.1354759 | 0.08778167 | -1.5433275 | 0.12275133 |
| TMA108 | YIL137C | 1309.43419 | -0.1008124 | 0.06532673 | -1.5432024 | 0.12278168 |
| Xu_2009_CUTs35 | Xu_2009_CU | 54.3124726 | 0.32502625 | 0.21080334 | 1.54184577 | 0.12311108 |
| MSA1 | YOR066W | 1705.20775 | 0.10488865 | 0.06808094 | 1.54064627 | 0.1234029 |
| KRR1 | YCL059C | 805.288983 | -0.1281314 | 0.08317375 | -1.5405266 | 0.12343204 |
| NTC20 | YBR188C | 173.869355 | -0.1990482 | 0.12923497 | -1.5402038 | 0.12351069 |
| tP(UGG)A/TRN1 | tP(UGG)A | 2.11779973 | -0.3733593 | 0.24249813 | -1.5396377 | 0.12364869 |
| Xu_2009_SUTs45 | Xu_2009_SU | 191.349572 | 0.19792421 | 0.12859813 | 1.53909087 | 0.12378211 |
| YFL065C | YFL065C | 9.54025104 | 0.45269361 | 0.2945041 | 1.53713854 | 0.12425939 |
| ATG41 | YPL250C | 189.954511 | 0.202157 | 0.13151257 | 1.53716859 | 0.12425203 |
| AEP2 | YMR282C | 469.469995 | 0.14837551 | 0.09662794 | 1.53553429 | 0.12465264 |

| | | | | | | |
|---|---|---|---|---|---|---|
| YHL006W-A | YHL006W-A | 3.64163915 | 0.42179652 | 0.2747504 | 1.53519892 | 0.12473497 |
| Xu_2009_CUTs76 | Xu_2009_CU | 9.85316397 | 0.45163319 | 0.29430241 | 1.53458883 | 0.12488486 |
| PEX4 | YGR133W | 234.932416 | -0.1742663 | 0.11355042 | -1.5347039 | 0.12485657 |
| CAB2 | YIL083C | 560.93353 | -0.1426752 | 0.0929773 | -1.5345161 | 0.12490274 |
| YCR102C | YCR102C | 13.644469 | 0.43857388 | 0.285992 | 1.53351802 | 0.12514827 |
| GEP5 | YLR091W | 233.994283 | -0.1753689 | 0.11443386 | -1.5324916 | 0.12540116 |
| Xu_2009_CUTs33 | Xu_2009_CU | 20.0870515 | 0.42855888 | 0.27968475 | 1.53229259 | 0.12545025 |
| Xu_2009_CUTs33 | Xu_2009_CU | 29.973981 | 0.38007135 | 0.24813363 | 1.53172043 | 0.12559144 |
| YER067C-A | YER067C-A | 5.94030733 | 0.44936464 | 0.2934208 | 1.53146823 | 0.12565371 |
| Xu_2009_CUTs91 | Xu_2009_CU | 14.467098 | 0.43710113 | 0.28555635 | 1.53070007 | 0.12584354 |
| Xu_2009_SUTs25: | Xu_2009_SU | 18.9996571 | 0.42440267 | 0.27730086 | 1.5304773 | 0.12589863 |
| Xu_2009_CUTs57 | Xu_2009_CU | 50.4047111 | 0.31473159 | 0.2057365 | 1.52978004 | 0.12607118 |
| POR2 | YIL114C | 182.92669 | -0.1970784 | 0.12883172 | -1.5297354 | 0.12608224 |
| BUD21 | YOR078W | 1068.70722 | 0.12746557 | 0.08334407 | 1.52938981 | 0.12616784 |
| YPS3 | YLR121C | 3358.67447 | -0.0946298 | 0.06188016 | -1.5292433 | 0.12620413 |
| INP2 | YMR163C | 207.567833 | -0.2085065 | 0.13646458 | -1.5279168 | 0.1265332 |
| MCM5 | YLR274W | 983.599029 | 0.11401887 | 0.07465895 | 1.52719625 | 0.12671222 |
| Xu_2009_SUTs78! | Xu_2009_SU | 3.8902516 | 0.42496525 | 0.27834661 | 1.52674845 | 0.12682358 |
| Xu_2009_SUTs34! | Xu_2009_SU | 71.8678882 | 0.29414333 | 0.19272018 | 1.52627151 | 0.12694226 |
| Xu_2009_SUTs76! | Xu_2009_SU | 67.9515812 | 0.28871351 | 0.18917866 | 1.52614207 | 0.12697449 |
| Xu_2009_CUTs96 | Xu_2009_CU | 80.8230981 | 0.27631887 | 0.18110375 | 1.52574904 | 0.12707238 |
| Xu_2009_CUTs84 | Xu_2009_CU | 11.7986107 | 0.44240633 | 0.29000826 | 1.5254956 | 0.12713553 |
| RAD18 | YCR066W | 402.287101 | -0.1401944 | 0.09189998 | -1.5255104 | 0.12713184 |
| Xu_2009_CUTs77 | Xu_2009_CU | 6.10295578 | 0.4490624 | 0.29442409 | 1.52522302 | 0.12720348 |
| MRF1 | YGL143C | 353.970548 | -0.1480151 | 0.09710184 | -1.5243284 | 0.12742671 |
| CHA4 | YLR098C | 544.099343 | 0.12970527 | 0.08511426 | 1.52389591 | 0.12753471 |
| tH(GUG)G1 | tH(GUG)G1 | 10.0946282 | -0.4515852 | 0.29643019 | -1.5234117 | 0.12765574 |
| Xu_2009_SUTs63: | Xu_2009_SU | 1.13140961 | 0.27983287 | 0.18370176 | 1.52329991 | 0.12768369 |
| NGR1 | YBR212W | 1101.64281 | -0.1027042 | 0.06742866 | -1.523154 | 0.12772019 |
| LOC1 | YFR001W | 717.253275 | 0.13825197 | 0.09078886 | 1.52278556 | 0.12781236 |
| Xu_2009_CUTs10 | Xu_2009_CU | 8.65216865 | 0.45105839 | 0.29637722 | 1.52190642 | 0.12803253 |
| Xu_2009_CUTs24 | Xu_2009_CU | 2.09043351 | 0.36152566 | 0.23764571 | 1.52127999 | 0.12818959 |
| YDR510C-A | YDR510C-A | 4.08841581 | 0.4265982 | 0.28049198 | 1.52089266 | 0.12828678 |
| Xu_2009_SUTs39· | Xu_2009_SU | 39.0173106 | 0.34829504 | 0.22910453 | 1.52024513 | 0.12844938 |
| PAU20 | YOL161C | 2.21264004 | 0.36662283 | 0.24129519 | 1.51939551 | 0.12866297 |
| tS(GCU)L | tS(GCU)L | 4.53458562 | -0.4316684 | 0.28417675 | -1.5190138 | 0.12875902 |
| YBR144C | YBR144C | 7.88677191 | 0.4500249 | 0.29626188 | 1.51901048 | 0.12875986 |
| YKR075W-A | YKR075W-A | 15.3677416 | 0.42792985 | 0.28218712 | 1.5164755 | 0.12939916 |
| COX13 | YGL191W | 863.571558 | -0.1299937 | 0.08577786 | -1.5154693 | 0.12965361 |
| SPR1 | YOR190W | 98.7526322 | -0.2431541 | 0.16052842 | -1.5147107 | 0.12984568 |
| Xu_2009_CUTs29 | Xu_2009_CU | 10.9913984 | 0.44297034 | 0.29277664 | 1.51299756 | 0.13028028 |

| | | | | | | |
|---|---|---|---|---|---|---|
| GEX1 | YCL073C | 13.8988206 | 0.43450842 | 0.28731609 | 1.51230105 | 0.13045729 |
| GDB1 | YPR184W | 2142.1817 | 0.09445683 | 0.06251679 | 1.51090327 | 0.1308131 |
| DBP1 | YPL119C | 296.745659 | -0.1565313 | 0.10367197 | -1.5098715 | 0.13107623 |
| TGS1 | YPL157W | 595.598199 | 0.13456047 | 0.08911503 | 1.50996379 | 0.13105266 |
| Xu_2009_CUTs80 | Xu_2009_CU | 20.7681692 | 0.40007312 | 0.2650137 | 1.50963187 | 0.13113738 |
| DIG2 | YDR480W | 230.335976 | -0.1773057 | 0.11744469 | -1.5096953 | 0.13112119 |
| COB | Q0105 | 232.699841 | -0.4213405 | 0.2791632 | -1.5092982 | 0.1312226 |
| YBR206W | YBR206W | 53.5390308 | -0.312753 | 0.2072089 | -1.5093607 | 0.13120663 |
| YBL113W-A | YBL113W-A | 2.4025812 | 0.37464327 | 0.24842103 | 1.50809806 | 0.13152943 |
| Xu_2009_SUTs13 | Xu_2009_SU | 49.5163233 | 0.31009865 | 0.20563673 | 1.50799255 | 0.13155643 |
| SIZ1 | YDR409W | 534.591377 | -0.1323879 | 0.08781516 | -1.5075746 | 0.13166344 |
| YDR426C | YDR426C | 10.8928206 | 0.44392911 | 0.29453958 | 1.50719677 | 0.13176023 |
| RPC34 | YNR003C | 571.57507 | -0.1285399 | 0.08530385 | -1.5068477 | 0.13184969 |
| Xu_2009_SUTs55 | Xu_2009_SU | 16.905619 | -0.4180911 | 0.27765067 | -1.5058169 | 0.13211419 |
| Xu_2009_SUTs19 | Xu_2009_SU | 42.2848439 | 0.3337667 | 0.22175437 | 1.50511896 | 0.1322935 |
| YNR003W-A | YNR003W-A | 1.14617383 | -0.2877837 | 0.19159803 | -1.5020182 | 0.1330924 |
| YGL214W | YGL214W | 26.8554369 | -0.3704369 | 0.24671661 | -1.5014674 | 0.13323472 |
| YGR283C | YGR283C | 341.664249 | -0.1706048 | 0.11366197 | -1.5009841 | 0.13335966 |
| Xu_2009_CUTs91 | Xu_2009_CU | 19.7887303 | 0.4045839 | 0.26959566 | 1.5007063 | 0.13343154 |
| YNL162W-A | YNL162W-A | 118.114973 | 0.2311938 | 0.15407307 | 1.50054641 | 0.13347292 |
| YML084W | YML084W | 10.5982901 | 0.44166098 | 0.29436054 | 1.50040824 | 0.13350869 |
| Xu_2009_CUTs64 | Xu_2009_CU | 0.75109689 | -0.2278292 | 0.15185556 | -1.5003023 | 0.13353611 |
| YKL031W | YKL031W | 27.2807791 | -0.3881777 | 0.25878297 | -1.5000124 | 0.1336112 |
| HAP3 | YBL021C | 299.21896 | -0.1680423 | 0.11213511 | -1.4985699 | 0.13398524 |
| Xu_2009_SUTs47 | Xu_2009_SU | 808.218513 | 0.12096668 | 0.08073493 | 1.498319 | 0.13405039 |
| YOR379C | YOR379C | 74.0199457 | 0.26501035 | 0.17712036 | 1.49621619 | 0.13459733 |
| Xu_2009_SUTs16 | Xu_2009_SU | 44.9418294 | 0.32508985 | 0.21731279 | 1.49595362 | 0.13466574 |
| Xu_2009_SUTs24 | Xu_2009_SU | 48.1606991 | 0.32648799 | 0.21832591 | 1.49541567 | 0.13480599 |
| YLR222C-A | YLR222C-A | 2.66471552 | 0.37462388 | 0.2507307 | 1.4941285 | 0.13514204 |
| Xu_2009_CUTs55 | Xu_2009_CU | 5.72180897 | 0.43667942 | 0.29230742 | 1.49390468 | 0.13520054 |
| tF(GAA)G | tF(GAA)G | 12.9330496 | -0.4357525 | 0.29173724 | -1.493647 | 0.1352679 |
| Xu_2009_SUTs14 | Xu_2009_SU | 69.3697647 | 0.27182757 | 0.18207282 | 1.49296069 | 0.13544748 |
| MOT3 | YMR070W | 795.803105 | 0.10871999 | 0.07285459 | 1.4922874 | 0.13562382 |
| YER137C | YER137C | 98.7632285 | 0.24974034 | 0.16757122 | 1.49035348 | 0.13613132 |
| PNT1 | YOR266W | 193.466272 | -0.1850085 | 0.12424817 | -1.4890239 | 0.13648107 |
| YIL108W | YIL108W | 1516.04239 | 0.08891005 | 0.05971428 | 1.48892446 | 0.13650726 |
| VPS8 | YAL002W | 1015.08083 | 0.10193817 | 0.06852198 | 1.48767109 | 0.13683766 |
| PTP1 | YDL230W | 278.51491 | -0.1616908 | 0.10868891 | -1.4876473 | 0.13684392 |
| TPO4 | YOR273C | 1981.09133 | -0.1016135 | 0.06834154 | -1.486848 | 0.13705495 |
| HIP1 | YGR191W | 1950.75086 | 0.11430677 | 0.07691159 | 1.48620997 | 0.13722359 |
| GPG1 | YGL121C | 939.597757 | -0.1122807 | 0.07557033 | -1.4857774 | 0.13733801 |

| | | | | | | |
|---|---|---|---|---|---|---|
| Xu_2009_SUTs82: | Xu_2009_SU | 48.7640462 | 0.31814186 | 0.2141541 | 1.48557447 | 0.13739171 |
| NCA2 | YPR155C | 412.189384 | -0.1448464 | 0.09750416 | -1.4855408 | 0.13740062 |
| GMC1 | YDR506C | 336.722188 | 0.14441339 | 0.09726095 | 1.48480336 | 0.13759592 |
| YGR121W-A | YGR121W-A | 39.3141183 | 0.34244031 | 0.23069338 | 1.48439593 | 0.13770392 |
| YDR169C-A | YDR169C-A | 17.1654664 | 0.41796069 | 0.28165937 | 1.48392255 | 0.13782947 |
| Xu_2009_SUTs57: | Xu_2009_SU | 113.996922 | 0.22159555 | 0.14937628 | 1.48347217 | 0.13794901 |
| Xu_2009_CUTs68 | Xu_2009_CU | 13.8565623 | 0.43306398 | 0.29198511 | 1.48317148 | 0.13802886 |
| Xu_2009_SUTs51 | Xu_2009_SU | 36.8956417 | 0.34335661 | 0.231719 | 1.48178012 | 0.13839881 |
| COI1 | YDR381C-A | 312.261688 | 0.153502 | 0.1036677 | 1.48071188 | 0.13868337 |
| Xu_2009_CUTs60 | Xu_2009_CU | 6.90355969 | 0.4370285 | 0.29527165 | 1.4800896 | 0.13884934 |
| MATALPHA2 | YCR039C | 14.0455525 | -0.4272937 | 0.28866034 | -1.4802645 | 0.13880267 |
| YLL058W | YLL058W | 563.135759 | -0.1210425 | 0.08178043 | -1.4800912 | 0.13884892 |
| RRT12 | YCR045C | 23.3787142 | -0.3906145 | 0.26396465 | -1.4797984 | 0.13892706 |
| YIL177C | YIL177C | 1.17024196 | 0.28756488 | 0.1943497 | 1.47962608 | 0.13897306 |
| Xu_2009_CUTs78 | Xu_2009_CU | 1.495959 | 0.318175 | 0.21512667 | 1.47901238 | 0.13913701 |
| YKL183C-A | YKL183C-A | 2.44015576 | 0.3504966 | 0.23712037 | 1.4781379 | 0.13937087 |
| ALG5 | YPL227C | 540.136289 | -0.1295584 | 0.08764851 | -1.478159 | 0.13936523 |
| YLF2 | YHL014C | 215.090323 | -0.1842573 | 0.12473333 | -1.47721 | 0.13961936 |
| Xu_2009_SUTs54: | Xu_2009_SU | 3.01039375 | -0.3803786 | 0.25755867 | -1.4768619 | 0.13971265 |
| Xu_2009_CUTs65 | Xu_2009_CU | 11.2695697 | 0.43021596 | 0.29142392 | 1.47625482 | 0.1398755 |
| Xu_2009_CUTs67 | Xu_2009_CU | 34.7393756 | -0.364324 | 0.24683042 | -1.4760092 | 0.13994142 |
| tL(UAG)L1 | tL(UAG)L1 | 59.1114501 | 0.28955689 | 0.19661885 | 1.47268126 | 0.140837 |
| OSM1 | YJR051W | 819.208031 | 0.10792903 | 0.07328796 | 1.47267071 | 0.14083985 |
| UTP6 | YDR449C | 799.734504 | -0.1313052 | 0.08926231 | -1.4710044 | 0.14128993 |
| TFA1 | YKL028W | 1009.68859 | -0.1063943 | 0.0723391 | -1.470772 | 0.14135278 |
| tF(GAA)H1 | tF(GAA)H1 | 3.77659849 | -0.4059482 | 0.27605232 | -1.470548 | 0.1414134 |
| Xu_2009_CUTs30 | Xu_2009_CU | 29.9911195 | 0.38038426 | 0.25882372 | 1.46966539 | 0.1416524 |
| LOH1 | YJL038C | 52.895686 | -0.2961916 | 0.20163578 | -1.4689436 | 0.14184809 |
| VAC14 | YLR386W | 856.544842 | 0.10960659 | 0.07466632 | 1.46795231 | 0.14211718 |
| PRM3 | YPL192C | 13.3776662 | -0.4222948 | 0.2876684 | -1.4679917 | 0.14210647 |
| YAR030C | YAR030C | 14.8587158 | 0.41794404 | 0.28478894 | 1.46755713 | 0.14222456 |
| tE(UUC)Q | tE(UUC)Q | 3.0697946 | -0.3883041 | 0.26465456 | -1.4672112 | 0.14231861 |
| MID2 | YLR332W | 1079.19604 | -0.0975125 | 0.06646214 | -1.4671886 | 0.14232475 |
| HSK3 | YKL138C-A | 261.847201 | -0.1811823 | 0.12375085 | -1.4640896 | 0.14316947 |
| YER135C | YER135C | 4.57846972 | -0.4140281 | 0.28282966 | -1.4638778 | 0.14322734 |
| YOR121C | YOR121C | 62.2129529 | -0.2911025 | 0.19884471 | -1.463969 | 0.14320243 |
| Xu_2009_CUTs41 | Xu_2009_CU | 3.64334303 | -0.4011683 | 0.27418251 | -1.4631432 | 0.14342819 |
| YJL067W | YJL067W | 24.5118547 | 0.3663543 | 0.25053602 | 1.46228195 | 0.14366396 |
| Xu_2009_CUTs81 | Xu_2009_CU | 19.6999483 | 0.39352045 | 0.26917319 | 1.46196002 | 0.14375217 |
| YOR029W | YOR029W | 1.36961258 | 0.3025399 | 0.20701144 | 1.46146465 | 0.14388797 |
| JNM1 | YMR294W | 247.927273 | 0.16528188 | 0.11316641 | 1.46052065 | 0.14414704 |

| | | | | | | |
|---|---|---|---|---|---|---|
| AUA1 | YFL010W-A | 30.2943106 | 0.351685 | 0.24090974 | 1.45982058 | 0.14433939 |
| DIB1 | YPR082C | 232.71465 | -0.1711288 | 0.11727079 | -1.4592623 | 0.14449293 |
| YDL186W | YDL186W | 9.15152344 | -0.4312425 | 0.29556229 | -1.4590578 | 0.14454919 |
| YDR535C | YDR535C | 26.5305661 | 0.36145306 | 0.24812207 | 1.45675497 | 0.14518403 |
| Xu_2009_SUTs64 | Xu_2009_SU | 24.0052186 | 0.37625909 | 0.25837839 | 1.45623282 | 0.14532827 |
| YER137W-A | YER137W-A | 7.71787182 | 0.43132837 | 0.29622895 | 1.45606418 | 0.14537488 |
| SMT1 | YJL147C | 266.159657 | -0.1578876 | 0.10843108 | -1.4561105 | 0.14536206 |
| MED2 | YDL005C | 777.027921 | 0.1132095 | 0.07786234 | 1.45396982 | 0.14595467 |
| SEC27 | YGL137W | 4217.4162 | -0.0845088 | 0.0581423 | -1.4534825 | 0.14608984 |
| Xu_2009_SUTs78 | Xu_2009_SU | 50.3027968 | -0.3014121 | 0.20744877 | -1.4529472 | 0.14623842 |
| UBC12 | YLR306W | 121.246036 | 0.22747341 | 0.15668075 | 1.4518275 | 0.14654958 |
| Xu_2009_CUTs69 | Xu_2009_CU | 27.5020952 | 0.36360927 | 0.25082518 | 1.44965217 | 0.14715554 |
| Xu_2009_SUTs29 | Xu_2009_SU | 93.7413682 | 0.23401055 | 0.16144709 | 1.44945658 | 0.14721012 |
| YER023C-A | YER023C-A | 5.93409402 | 0.42577039 | 0.29388653 | 1.44875776 | 0.14740524 |
| YCL019W | YCL019W | 21.1768008 | 0.3796216 | 0.26215647 | 1.4480726 | 0.14759675 |
| APN2 | YBL019W | 295.303942 | -0.1526646 | 0.10549506 | -1.4471255 | 0.14786178 |
| tL(CAA)G3 | tL(CAA)G3 | 1.75602592 | -0.3152618 | 0.21791397 | -1.4467258 | 0.14797374 |
| YOL050C | YOL050C | 3.05549108 | 0.37748582 | 0.26100395 | 1.44628393 | 0.14809758 |
| UTP18 | YJL069C | 1086.92859 | 0.11370788 | 0.07870897 | 1.44466232 | 0.14855276 |
| NTG2 | YOL043C | 236.775368 | 0.16743978 | 0.1159054 | 1.44462447 | 0.14856339 |
| MTW1 | YAL034W-A | 170.209555 | -0.1894964 | 0.13119065 | -1.4444354 | 0.14861654 |
| Xu_2009_CUTs68 | Xu_2009_CU | 5.66304287 | 0.42128238 | 0.29176586 | 1.44390568 | 0.14876551 |
| TOF2 | YKR010C | 294.185164 | 0.15603095 | 0.10820726 | 1.44196384 | 0.14931257 |
| Xu_2009_CUTs14 | Xu_2009_CU | 24.268533 | 0.3752907 | 0.26057076 | 1.44026407 | 0.1497927 |
| YDL032W | YDL032W | 6.59732913 | 0.42330805 | 0.2939438 | 1.44009857 | 0.14983951 |
| YHL034W-A | YHL034W-A | 5.85154187 | 0.42052523 | 0.29201426 | 1.4400846 | 0.14984347 |
| YLR287C | YLR287C | 836.75964 | -0.1033511 | 0.07177484 | -1.4399351 | 0.14988576 |
| AMF1 | YOR378W | 28.236588 | -0.3619456 | 0.25153224 | -1.4389628 | 0.15016105 |
| YDR366C | YDR366C | 9.20226243 | 0.42604783 | 0.29613834 | 1.43867839 | 0.15024166 |
| SAE2 | YGL175C | 210.965337 | 0.18213157 | 0.1266068 | 1.4385607 | 0.15027503 |
| PEA2 | YER149C | 467.985095 | 0.12870263 | 0.08947595 | 1.43840482 | 0.15031923 |
| RRT13 | YER066W | 66.9708991 | -0.2733699 | 0.190092 | -1.4380929 | 0.15040771 |
| OTU1 | YFL044C | 392.162313 | -0.139289 | 0.09685401 | -1.4381333 | 0.15039622 |
| YDR061W | YDR061W | 636.776662 | -0.1222375 | 0.08506745 | -1.4369475 | 0.15073291 |
| YGR039W | YGR039W | 61.7457417 | 0.27648263 | 0.1924409 | 1.43671452 | 0.15079913 |
| MKS1 | YNL076W | 1004.73177 | 0.09741845 | 0.06780237 | 1.43680009 | 0.15077481 |
| tG(UCC)N/SUF6 | tG(UCC)N | 21.1823004 | -0.3874396 | 0.26972251 | -1.4364379 | 0.15087778 |
| MRX10 | YDR282C | 215.593187 | -0.1693846 | 0.1179242 | -1.4363852 | 0.15089277 |
| LSG1 | YGL099W | 1702.4714 | 0.09553553 | 0.06651324 | 1.43633846 | 0.15090606 |
| Xu_2009_SUTs33 | Xu_2009_SU | 11.3017119 | -0.4215313 | 0.29356086 | -1.4359247 | 0.15102378 |
| QCR9 | YGR183C | 584.079707 | -0.1220636 | 0.08504272 | -1.4353215 | 0.1511955 |

| | | | | | | |
|---|---|---|---|---|---|---|
| EXO70 | YJL085W | 462.483709 | 0.12803636 | 0.08920035 | 1.43537948 | 0.15117899 |
| EUC1 | YMR111C | 430.325727 | 0.12881893 | 0.08976609 | 1.43505108 | 0.15127254 |
| MND1 | YGL183C | 66.3246546 | 0.26274892 | 0.18322458 | 1.43402661 | 0.15156467 |
| FRE6 | YLL051C | 1051.5447 | -0.1014651 | 0.07076313 | -1.4338698 | 0.15160943 |
| IML3 | YBR107C | 140.859305 | 0.20522542 | 0.14321524 | 1.43298587 | 0.15186188 |
| YHL041W | YHL041W | 7.02769033 | 0.42246459 | 0.29500239 | 1.43207174 | 0.15212329 |
| HSP150 | YJL159W | 40038.7986 | 0.06225501 | 0.04348428 | 1.43166724 | 0.15223908 |
| Xu_2009_CUTs15 | Xu_2009_CU | 49.6497389 | 0.30066956 | 0.21003132 | 1.43154631 | 0.1522737 |
| STE14 | YDR410C | 244.95845 | -0.1653862 | 0.1155531 | -1.4312569 | 0.15235659 |
| TDA6 | YPR157W | 124.414071 | -0.2258282 | 0.15790834 | -1.4301219 | 0.15268204 |
| Xu_2009_CUTs32 | Xu_2009_CU | 50.1607571 | -0.2942359 | 0.20590929 | -1.4289589 | 0.15301605 |
| CDC45 | YLR103C | 502.74245 | 0.13706305 | 0.0959604 | 1.42832927 | 0.15319711 |
| Xu_2009_CUTs89 | Xu_2009_CU | 4.77923653 | -0.4048539 | 0.28350237 | -1.4280443 | 0.15327911 |
| YHR214C-E | YHR214C-E | 1.27911226 | 0.29004108 | 0.20314246 | 1.42777181 | 0.15335755 |
| SOL2 | YCR073W-A | 497.475946 | -0.1278668 | 0.08974457 | -1.4247856 | 0.15421919 |
| YPL062W | YPL062W | 11.4302598 | 0.41668439 | 0.29245665 | 1.42477315 | 0.15422279 |
| FAA3 | YIL009W | 542.721842 | -0.1264641 | 0.08879158 | -1.4242802 | 0.15436538 |
| Xu_2009_SUTs50 | Xu_2009_SU | 49.0927052 | 0.28824421 | 0.20256313 | 1.42298461 | 0.15474062 |
| GAL7 | YBR018C | 14.7904369 | -0.4092102 | 0.28761243 | -1.4227835 | 0.15479894 |
| Xu_2009_CUTs61 | Xu_2009_CU | 18.7290025 | 0.39386607 | 0.2769911 | 1.42194485 | 0.15504226 |
| RRB1 | YMR131C | 3059.94189 | -0.0991439 | 0.06972948 | -1.4218366 | 0.15507369 |
| YML054C-A | YML054C-A | 5.05017546 | 0.41164982 | 0.28955722 | 1.42165279 | 0.15512707 |
| YLR036C | YLR036C | 215.680436 | -0.1712176 | 0.12045423 | -1.4214332 | 0.15519087 |
| ATG17 | YLR423C | 135.250256 | -0.2122884 | 0.14937085 | -1.4212169 | 0.1552537 |
| Xu_2009_SUTs80 | Xu_2009_SU | 60.3307873 | -0.2756055 | 0.19416556 | -1.4194354 | 0.15577213 |
| NUG1 | YER006W | 1916.67485 | 0.1003954 | 0.07073875 | 1.4192419 | 0.15582851 |
| ALD2 | YMR170C | 389.487238 | -0.1454139 | 0.10259638 | -1.4173393 | 0.15638374 |
| tL(UAG)L2 | tL(UAG)L2 | 3.98461984 | -0.3914493 | 0.27622405 | -1.4171441 | 0.15644081 |
| Xu_2009_CUTs60 | Xu_2009_CU | 5.82628968 | 0.41495967 | 0.29305268 | 1.41599003 | 0.15677842 |
| APP1 | YNL094W | 1860.99238 | 0.09193711 | 0.06499925 | 1.41443329 | 0.15723472 |
| YAL037C-A | YAL037C-A | 3.57291753 | -0.3758935 | 0.26586605 | -1.4138454 | 0.1574073 |
| YIL024C | YIL024C | 228.578103 | 0.1722837 | 0.12195662 | 1.41266381 | 0.1577546 |
| EXO84 | YBR102C | 815.186814 | 0.10484694 | 0.07423695 | 1.41232821 | 0.15785335 |
| YIR023C-A | YIR023C-A | 2.76919614 | 0.35933314 | 0.25442925 | 1.41231067 | 0.15785851 |
| Xu_2009_SUTs47 | Xu_2009_SU | 16.5935388 | -0.3927884 | 0.27825035 | -1.4116367 | 0.15805697 |
| YLR230W | YLR230W | 33.6538688 | 0.36687949 | 0.25988471 | 1.41170095 | 0.15803803 |
| BSP1 | YPR171W | 713.966226 | -0.1081991 | 0.07665698 | -1.4114715 | 0.15810562 |
| EFR3 | YMR212C | 1361.23945 | -0.0895488 | 0.06345343 | -1.4112515 | 0.15817047 |
| APS2 | YJR058C | 365.928703 | 0.14135941 | 0.1002044 | 1.41071062 | 0.15832996 |
| YPK1 | YKL126W | 2060.74492 | -0.0883099 | 0.06260095 | -1.4106803 | 0.15833891 |
| Xu_2009_SUTs66 | Xu_2009_SU | 9.24557853 | 0.41814438 | 0.29643805 | 1.41056247 | 0.15837366 |

| | | | | | | |
|---|---|---|---|---|---|---|
| YDR327W | YDR327W | 7.86826545 | 0.41793719 | 0.29633592 | 1.4103494 | 0.15843654 |
| CEM1 | YER061C | 405.112143 | 0.13022052 | 0.09233291 | 1.41033704 | 0.15844019 |
| CUB1 | YPL260W | 1666.28728 | -0.0861104 | 0.06108396 | -1.4097051 | 0.15862679 |
| HSP32 | YPL280W | 21.0169585 | 0.37614401 | 0.26685217 | 1.40955951 | 0.15866979 |
| RPN9 | YDR427W | 1975.28278 | 0.08611201 | 0.0611547 | 1.40810112 | 0.15910113 |
| Xu_2009_CUTs20 | Xu_2009_CU | 0.73223885 | 0.22879306 | 0.16256025 | 1.40743548 | 0.1592983 |
| SCM3 | YDL139C | 166.725969 | 0.19012446 | 0.13509178 | 1.40737246 | 0.15931697 |
| YOL079W | YOL079W | 4.27800691 | 0.39959684 | 0.28388608 | 1.40759572 | 0.15925082 |
| ATG13 | YPR185W | 612.420038 | 0.13175384 | 0.09360991 | 1.40747745 | 0.15928586 |
| PET112 | YBL080C | 281.938554 | -0.1610951 | 0.11447682 | -1.4072291 | 0.15935946 |
| Xu_2009_CUTs11 | Xu_2009_CU | 5.63794457 | 0.40977076 | 0.29122123 | 1.40707723 | 0.15940449 |
| YER078W-A | YER078W-A | 12.8122637 | 0.4069008 | 0.28930657 | 1.40646927 | 0.15958482 |
| VHS2 | YIL135C | 927.329433 | -0.0998618 | 0.07099966 | -1.4065113 | 0.15957235 |
| HXT8 | YJL214W | 24.7716241 | -0.361972 | 0.2573359 | -1.4066128 | 0.15954223 |
| PBA1 | YLR199C | 526.278671 | -0.1176946 | 0.08370985 | -1.4059823 | 0.15972938 |
| Xu_2009_SUTs12 | Xu_2009_SU | 118.516805 | 0.20738667 | 0.14757714 | 1.40527638 | 0.15993911 |
| ITR2 | YOL103W | 1726.90953 | -0.0855415 | 0.06089704 | -1.4046899 | 0.16011352 |
| ACE2 | YLR131C | 824.3494 | 0.10701137 | 0.07620139 | 1.40432308 | 0.16022267 |
| Xu_2009_SUTs52 | Xu_2009_SU | 94.0444503 | 0.22846218 | 0.16271767 | 1.40404037 | 0.16030683 |
| YLR364C-A | YLR364C-A | 4.3292725 | -0.4000229 | 0.28491985 | -1.403984 | 0.16032363 |
| Xu_2009_SUTs11 | Xu_2009_SU | 7.37961494 | 0.41441051 | 0.29525 | 1.40359189 | 0.16044041 |
| CSR2 | YPR030W | 402.017774 | -0.1335014 | 0.095135 | -1.4032842 | 0.16053211 |
| YLR462W | YLR462W | 10.3450162 | 0.41469222 | 0.29569793 | 1.40241845 | 0.16079033 |
| HRD1 | YOL013C | 717.471022 | 0.10629063 | 0.07581492 | 1.40197511 | 0.16092268 |
| YNL043C | YNL043C | 12.7063681 | 0.40710165 | 0.29045885 | 1.40158116 | 0.16104036 |
| tK(UUU)G2 | tK(UUU)G2 | 27.3301099 | 0.34325449 | 0.24507959 | 1.40058376 | 0.16133858 |
| ALG14 | YBR070C | 198.644236 | -0.1799465 | 0.12848671 | -1.4005065 | 0.1613617 |
| PRM15 | YMR278W | 618.724671 | -0.1264934 | 0.09035328 | -1.3999869 | 0.16151725 |
| PRY1 | YJL079C | 2294.47581 | 0.08705936 | 0.06221296 | 1.39937667 | 0.16170006 |
| EST3 | YIL009C-A | 240.250012 | -0.1647091 | 0.11773275 | -1.3990081 | 0.16181055 |
| YJL026C-A | YJL026C-A | 5.72102967 | 0.40783948 | 0.29160537 | 1.3986007 | 0.16193276 |
| Xu_2009_CUTs47 | Xu_2009_CU | 4.6491777 | 0.39394864 | 0.28176143 | 1.39816382 | 0.16206388 |
| MDH2 | YOL126C | 1011.9186 | 0.1221382 | 0.08743693 | 1.39687198 | 0.16245207 |
| tG(GCC)J2/SUF23 | tG(GCC)J2 | 21.3686185 | 0.37224645 | 0.2665552 | 1.39650791 | 0.1625616 |
| Xu_2009_SUTs51 | Xu_2009_SU | 4.35859967 | 0.38696051 | 0.2771326 | 1.39630096 | 0.16262389 |
| NBA1 | YOL070C | 1242.27096 | 0.1091685 | 0.07821784 | 1.39569808 | 0.16280543 |
| Xu_2009_SUTs63 | Xu_2009_SU | 73.3540247 | 0.25851831 | 0.18529828 | 1.39514683 | 0.16297157 |
| ZDS2 | YML109W | 550.600667 | 0.12569428 | 0.0901407 | 1.39442311 | 0.16318987 |
| Xu_2009_SUTs28 | Xu_2009_SU | 52.1796038 | 0.28504613 | 0.20454168 | 1.39358458 | 0.16344309 |
| Xu_2009_SUTs72 | Xu_2009_SU | 38.4715575 | -0.3116302 | 0.22378103 | -1.3925676 | 0.16375058 |
| SWE1 | YJL187C | 951.057523 | 0.09591608 | 0.06887982 | 1.39251347 | 0.16376697 |

| | | | | | | |
|---|---|---|---|---|---|---|
| YOR186W | YOR186W | 20.8203195 | 0.36810394 | 0.26438315 | 1.39231243 | 0.16382781 |
| IMP2' | YIL154C | 945.823609 | 0.09933875 | 0.07135441 | 1.39218784 | 0.16386552 |
| MMT1 | YMR177W | 655.206596 | -0.1358288 | 0.09758698 | -1.391874 | 0.16396057 |
| JID1 | YPR061C | 159.135864 | 0.19524768 | 0.14030745 | 1.39157029 | 0.16405256 |
| Xu_2009_SUTs37 | Xu_2009_SU | 83.6148441 | -0.2603164 | 0.18715259 | -1.3909311 | 0.16424632 |
| KRE29 | YER038C | 226.871853 | 0.16259095 | 0.11690988 | 1.39073748 | 0.16430505 |
| EDC1 | YGL222C | 700.421972 | 0.12625279 | 0.0907925 | 1.39056399 | 0.16435768 |
| Xu_2009_SUTs38 | Xu_2009_SU | 22.2023552 | 0.36518737 | 0.26272342 | 1.39000692 | 0.16452678 |
| Xu_2009_CUTs47 | Xu_2009_CU | 4.05354951 | 0.39385077 | 0.2834072 | 1.38969925 | 0.16462022 |
| MGL2 | YMR210W | 224.236615 | -0.1681699 | 0.1210634 | -1.3891056 | 0.16480063 |
| APQ12 | YIL040W | 578.724584 | -0.1115366 | 0.08035216 | -1.3880968 | 0.16510757 |
| Xu_2009_SUTs56 | Xu_2009_SU | 10.9706892 | -0.4065667 | 0.29292071 | -1.3879753 | 0.16514457 |
| PEX29 | YDR479C | 458.979378 | 0.11844796 | 0.08538041 | 1.38729678 | 0.16535129 |
| HEM14 | YER014W | 779.365437 | -0.1153805 | 0.08324386 | -1.3860545 | 0.16573025 |
| CST26 | YBR042C | 365.791552 | -0.1436646 | 0.10368743 | -1.3855542 | 0.16588307 |
| YCR001W | YCR001W | 1.18741263 | 0.26826831 | 0.19366204 | 1.38523946 | 0.16597926 |
| YDL206W | YDL206W | 538.322306 | 0.11808151 | 0.0852531 | 1.38507 | 0.16603106 |
| Xu_2009_SUTs82 | Xu_2009_SU | 5.4078077 | 0.40103606 | 0.28964338 | 1.38458559 | 0.16617922 |
| THI21 | YPL258C | 265.551115 | 0.15224867 | 0.1100183 | 1.38384855 | 0.16640483 |
| YRB2 | YIL063C | 718.617845 | 0.12842833 | 0.09282862 | 1.38349932 | 0.16651182 |
| Xu_2009_CUTs38 | Xu_2009_CU | 21.2002346 | 0.37005978 | 0.26762689 | 1.38274511 | 0.16674304 |
| YOL103W-B | YOL103W-B | 127.354894 | 0.21854531 | 0.15807939 | 1.38250346 | 0.16681717 |
| MSC3 | YLR219W | 1373.01504 | 0.0951263 | 0.0688152 | 1.38234434 | 0.166866 |
| RRG7 | YOR305W | 379.36921 | 0.13327389 | 0.09645786 | 1.38167988 | 0.16707001 |
| YNG1 | YOR064C | 430.660698 | 0.12915623 | 0.09356921 | 1.38032833 | 0.16748557 |
| NDL1 | YLR254C | 408.533815 | 0.13609339 | 0.0986131 | 1.38007409 | 0.16756383 |
| PCL10 | YGL134W | 540.098132 | -0.1198026 | 0.08682529 | -1.3798128 | 0.1676443 |
| NEW1 | YPL226W | 4828.42924 | 0.08148436 | 0.05908 | 1.37922064 | 0.16782674 |
| tL(UAA)J/SUP51 | tL(UAA)J | 4.2211991 | -0.3861834 | 0.28040255 | -1.3772464 | 0.16843608 |
| NPC2 | YDL046W | 1685.84496 | 0.09706024 | 0.07047675 | 1.37719521 | 0.1684519 |
| Xu_2009_SUTs83 | Xu_2009_SU | 67.9210702 | 0.25796393 | 0.18733066 | 1.37705126 | 0.1684964 |
| Xu_2009_CUTs48 | Xu_2009_CU | 25.2092045 | 0.35309937 | 0.25654039 | 1.37638901 | 0.16870123 |
| HST2 | YPL015C | 1037.28878 | -0.0950242 | 0.06905308 | -1.3761044 | 0.16878932 |
| Xu_2009_SUTs15 | Xu_2009_SU | 113.489078 | 0.20633996 | 0.14997038 | 1.37587141 | 0.16886145 |
| IRC21 | YMR073C | 315.934928 | -0.1547376 | 0.11250239 | -1.3754164 | 0.16900239 |
| Xu_2009_SUTs77 | Xu_2009_SU | 103.442664 | -0.2143784 | 0.15597253 | -1.3744625 | 0.16929815 |
| RCI50 | YKL133C | 165.974715 | 0.18079344 | 0.13153133 | 1.37452757 | 0.16927796 |
| Xu_2009_SUTs46 | Xu_2009_SU | 12.1264885 | 0.39771609 | 0.28956645 | 1.37348813 | 0.16960065 |
| YNL194C | YNL194C | 68.2760434 | -0.2544331 | 0.18536039 | -1.37264 | 0.16986428 |
| SDD1 | YEL057C | 64.2352042 | -0.26027 | 0.18990701 | -1.3705126 | 0.17052693 |
| PPE1 | YHR075C | 365.533787 | 0.16675614 | 0.12170201 | 1.37020031 | 0.17062438 |

| | | | | | | |
|---|---|---|---|---|---|---|
| NDE2 | YDL085W | 102.356181 | -0.2381646 | 0.1739109 | -1.3694632 | 0.17085454 |
| PFK26 | YIL107C | 749.642504 | 0.12511344 | 0.0913592 | 1.36946738 | 0.17085323 |
| Xu_2009_CUTs64 | Xu_2009_CU | 24.0280929 | 0.35915002 | 0.26261589 | 1.36758677 | 0.17144146 |
| PRP18 | YGR006W | 191.635855 | 0.17800041 | 0.1302732 | 1.36636248 | 0.17182522 |
| ESBP6 | YNL125C | 974.386781 | 0.10776432 | 0.07890092 | 1.36581823 | 0.17199602 |
| SFH1 | YLR321C | 679.435993 | -0.1107356 | 0.08108367 | -1.3656953 | 0.17203463 |
| MUM2 | YBR057C | 940.339985 | -0.0940084 | 0.06888947 | -1.364627 | 0.17237031 |
| YJR079W | YJR079W | 23.309511 | 0.35797359 | 0.26235866 | 1.3644436 | 0.17242799 |
| PHO11 | YAR071W | 21.0733489 | 0.35568916 | 0.26077904 | 1.36394839 | 0.17258381 |
| SGF11 | YPL047W | 559.438914 | 0.12274702 | 0.09003028 | 1.36339713 | 0.17275739 |
| Xu_2009_CUTs47 | Xu_2009_CU | 5.45790113 | 0.39634119 | 0.29085085 | 1.36269567 | 0.17297844 |
| PRP31 | YGR091W | 346.531508 | 0.13165036 | 0.0966872 | 1.36161099 | 0.17332068 |
| Xu_2009_SUTs65 | Xu_2009_SU | 33.3796925 | 0.33429842 | 0.24556101 | 1.36136604 | 0.17339804 |
| Xu_2009_SUTs72 | Xu_2009_SU | 1.54010053 | -0.2937921 | 0.21581552 | -1.3613112 | 0.17341536 |
| Xu_2009_SUTs44 | Xu_2009_SU | 258.178338 | 0.16199952 | 0.11905455 | 1.36071672 | 0.17360323 |
| CSS3 | YOL159C | 167.35836 | -0.173631 | 0.12762264 | -1.3605033 | 0.17367072 |
| tE(UUC)L | tE(UUC)L | 6.14997803 | -0.3967316 | 0.29168689 | -1.3601283 | 0.17378932 |
| VTH1 | YIL173W | 12.6063453 | 0.39321489 | 0.28921309 | 1.3596027 | 0.17395568 |
| DPB4 | YDR121W | 417.073531 | -0.1246047 | 0.09177394 | -1.3577349 | 0.17454782 |
| GRX8 | YLR364W | 184.742809 | -0.1861806 | 0.13712037 | -1.3577896 | 0.17453047 |
| RPN13 | YLR421C | 696.06102 | -0.1115596 | 0.08220791 | -1.3570423 | 0.17476776 |
| DIN7 | YDR263C | 189.630249 | 0.17328724 | 0.1277476 | 1.35648137 | 0.17494606 |
| KTR6 | YPL053C | 823.950179 | -0.0975869 | 0.07195677 | -1.3561879 | 0.17503941 |
| CWC25 | YNL245C | 251.044318 | 0.17264203 | 0.12731471 | 1.35602574 | 0.17509098 |
| Xu_2009_CUTs48 | Xu_2009_CU | 55.6164655 | 0.27239562 | 0.20100242 | 1.35518574 | 0.17535839 |
| YDL057W | YDL057W | 226.149949 | 0.15544922 | 0.11480312 | 1.35405045 | 0.17572028 |
| Xu_2009_SUTs33 | Xu_2009_SU | 125.25306 | 0.19678301 | 0.14537532 | 1.35362051 | 0.17585748 |
| LSO1 | YJR005C-A | 7.45571835 | -0.4009375 | 0.29625224 | -1.3533653 | 0.17593897 |
| Xu_2009_SUTs59 | Xu_2009_SU | 36.9324796 | 0.3186025 | 0.23557014 | 1.35247404 | 0.17622372 |
| SPB4 | YFL002C | 991.039935 | 0.09067964 | 0.0670575 | 1.35226682 | 0.17628997 |
| CYT2 | YKL087C | 532.463073 | -0.1107177 | 0.08187372 | -1.3522989 | 0.1762797 |
| PDL32 | YMR087W | 328.280242 | 0.14817882 | 0.10957766 | 1.3522721 | 0.17628829 |
| YPR160C-A | YPR160C-A | 21.9352268 | 0.35645021 | 0.26357689 | 1.35235757 | 0.17626096 |
| PSP1 | YDR505C | 2551.48327 | 0.07826992 | 0.05790829 | 1.35161863 | 0.17649735 |
| MNT2 | YGL257C | 431.410306 | -0.1286122 | 0.09528641 | -1.3497434 | 0.17709832 |
| Xu_2009_SUTs75 | Xu_2009_SU | 8.89608761 | 0.39944385 | 0.29599518 | 1.34949442 | 0.17717821 |
| YBR277C | YBR277C | 11.0908735 | 0.39558556 | 0.29313373 | 1.34950544 | 0.17717467 |
| FMT1 | YBL013W | 158.28176 | 0.18616199 | 0.13796428 | 1.34934911 | 0.17722486 |
| YLR149C | YLR149C | 553.959641 | -0.1266799 | 0.09393149 | -1.3486417 | 0.17745209 |
| RNH70 | YGR276C | 692.88851 | 0.10318581 | 0.07657353 | 1.34753888 | 0.17780674 |
| CTP1 | YBR291C | 157.46951 | -0.1841291 | 0.13667284 | -1.3472251 | 0.17790776 |

| | | | | | | |
|---|---|---|---|---|---|---|
| Xu_2009_SUTs55( | Xu_2009_SU | 27.1758067 | 0.343441 | 0.25497023 | 1.34698468 | 0.17798517 |
| MFG1 | YDL233W | 439.832546 | -0.1245855 | 0.0925524 | -1.3461076 | 0.17826782 |
| CDC34 | YDR054C | 1827.80006 | -0.0774679 | 0.05756027 | -1.3458574 | 0.1783485 |
| BUD16 | YEL029C | 733.190571 | 0.10113456 | 0.07520701 | 1.34474915 | 0.17870626 |
| tV(AAC)J | tV(AAC)J | 3.89952358 | -0.3614642 | 0.26882502 | -1.3446076 | 0.17875199 |
| MAL32 | YBR299W | 22.2994074 | -0.3575856 | 0.26599751 | -1.3443194 | 0.17884512 |
| Xu_2009_CUTs35 | Xu_2009_CU | 11.7895624 | 0.39335875 | 0.29270129 | 1.34389139 | 0.17898351 |
| Xu_2009_CUTs59 | Xu_2009_CU | 34.8573861 | 0.32160192 | 0.23929177 | 1.343974 | 0.17895679 |
| AQY1 | YPR192W | 14.2263511 | -0.383449 | 0.28531301 | -1.3439589 | 0.17896167 |
| tS(AGA)M | tS(AGA)M | 5.57611072 | -0.3917875 | 0.29177084 | -1.3427917 | 0.17933944 |
| PEX28 | YHR150W | 369.603336 | -0.1325404 | 0.0987958 | -1.3415594 | 0.17973891 |
| CHK1 | YBR274W | 359.472788 | 0.13317362 | 0.09930002 | 1.34112378 | 0.17988027 |
| Xu_2009_SUTs26 | Xu_2009_SU | 22.9346049 | 0.34955591 | 0.26067872 | 1.34094529 | 0.17993821 |
| YCR038W-A | YCR038W-A | 10.9007602 | 0.39312098 | 0.29324474 | 1.34059005 | 0.18005359 |
| YHR050W-A | YHR050W-A | 73.4150684 | 0.25942258 | 0.19352877 | 1.34048589 | 0.18008743 |
| YDR109C | YDR109C | 565.911209 | 0.11248509 | 0.08394532 | 1.3399804 | 0.18025172 |
| Xu_2009_CUTs88 | Xu_2009_CU | 9.24215004 | 0.3961332 | 0.29569681 | 1.33966005 | 0.18035589 |
| EMA17 | YIL029C | 32.9621073 | 0.3099509 | 0.23144921 | 1.33917458 | 0.18051385 |
| VMR1 | YHL035C | 1263.09428 | -0.1118205 | 0.08351311 | -1.3389579 | 0.18058437 |
| PRS2 | YER099C | 831.53116 | 0.0949494 | 0.07094879 | 1.33828078 | 0.18080493 |
| CBP2 | YHL038C | 479.00413 | 0.11227988 | 0.08392188 | 1.33790945 | 0.18092596 |
| MRL1 | YPR079W | 630.605037 | -0.1159872 | 0.08674423 | -1.337117 | 0.18118444 |
| RRP15 | YPR143W | 956.745926 | 0.09962323 | 0.07451641 | 1.33693002 | 0.18124548 |
| tM(CAU)O1/IMT1 | tM(CAU)O1 | 1.50054591 | -0.2751927 | 0.20589613 | -1.3365606 | 0.18136612 |
| YAL019W-A | YAL019W-A | 6.75290974 | 0.39490091 | 0.29544555 | 1.3366284 | 0.18134396 |
| IRR1 | YIL026C | 822.875926 | -0.0957346 | 0.07169896 | -1.3352294 | 0.18180128 |
| CTO1 | YCR015C | 232.386354 | 0.16456589 | 0.12328295 | 1.33486327 | 0.1819211 |
| TCA17 | YEL048C | 132.256485 | -0.1967739 | 0.14740571 | -1.3349137 | 0.18190461 |
| TAF10 | YDR167W | 370.649159 | -0.1361818 | 0.1020935 | -1.3338934 | 0.1822388 |
| DLD2 | YDL178W | 939.152416 | 0.0889214 | 0.0666788 | 1.33357826 | 0.18234211 |
| RPB9 | YGL070C | 931.797491 | 0.10319321 | 0.07737745 | 1.33363413 | 0.18232379 |
| YKL063C | YKL063C | 705.92928 | 0.10300752 | 0.07726444 | 1.33318153 | 0.18247224 |
| Xu_2009_CUTs78 | Xu_2009_CU | 96.9314645 | 0.22341183 | 0.16775274 | 1.33179248 | 0.18292839 |
| NDE1 | YMR145C | 2377.16942 | 0.08195275 | 0.06156705 | 1.33111376 | 0.18315158 |
| HAT1 | YPL001W | 468.225757 | -0.1193003 | 0.08962065 | -1.3311697 | 0.18313317 |
| Xu_2009_SUTs38! | Xu_2009_SU | 62.0769894 | 0.26408685 | 0.19849154 | 1.33046902 | 0.18336379 |
| CDC9 | YDL164C | 713.381445 | -0.1016011 | 0.07638829 | -1.3300615 | 0.18349801 |
| YDR391C | YDR391C | 537.203698 | -0.1322611 | 0.09944141 | -1.3300409 | 0.18350479 |
| Xu_2009_CUTs62 | Xu_2009_CU | 26.2059249 | -0.3376711 | 0.25398979 | -1.3294673 | 0.18369384 |
| LPP1 | YDR503C | 191.814109 | -0.1698516 | 0.12776151 | -1.3294426 | 0.183702 |
| SEC2 | YNL272C | 760.893049 | 0.10602348 | 0.07976052 | 1.32927264 | 0.18375804 |

| | | | | | | |
|---|---|---|---|---|---|---|
| YPL142C | YPL142C | 0.9863861 | 0.24284722 | 0.1826925 | 1.32926763 | 0.18375969 |
| SUP35 | YDR172W | 4422.34892 | -0.0689555 | 0.05191338 | -1.3282809 | 0.18408531 |
| GPN3 | YLR243W | 461.021494 | -0.1263501 | 0.09519778 | -1.3272379 | 0.18443 |
| YHR131W-A | YHR131W-A | 2.77424722 | -0.3411501 | 0.25715452 | -1.3266345 | 0.18462961 |
| PEX22 | YAL055W | 152.499083 | -0.1883709 | 0.14208774 | -1.3257365 | 0.184927 |
| NPY1 | YGL067W | 561.590674 | -0.1103816 | 0.08326429 | -1.3256781 | 0.18494636 |
| YBR190W | YBR190W | 3.82880632 | 0.36439885 | 0.27500234 | 1.32507545 | 0.18514613 |
| Xu_2009_CUTs82 | Xu_2009_CU | 2.86913909 | 0.34431171 | 0.26019669 | 1.32327477 | 0.18574402 |
| CAK1 | YFL029C | 361.039422 | -0.1260943 | 0.0953254 | -1.322778 | 0.18590921 |
| YHR213W | YHR213W | 3.40046759 | 0.3582022 | 0.27100607 | 1.32174972 | 0.18625151 |
| LDO16 | YMR148W | 484.067203 | 0.11306029 | 0.08561006 | 1.32064259 | 0.18662057 |
| YNL150W | YNL150W | 12.5809803 | 0.37906462 | 0.28704314 | 1.32058415 | 0.18664006 |
| RSO55 | YLR281C | 0.70170754 | 0.20553968 | 0.15569544 | 1.32013938 | 0.18678849 |
| Xu_2009_SUTs71! | Xu_2009_SU | 125.59372 | -0.1966263 | 0.1489846 | -1.3197761 | 0.18690979 |
| FLC3 | YGL139W | 1123.16021 | -0.0921369 | 0.06985531 | -1.3189676 | 0.18717996 |
| Xu_2009_CUTs74 | Xu_2009_CU | 12.0590285 | 0.38539006 | 0.29225671 | 1.31866969 | 0.18727956 |
| YGL074C | YGL074C | 2.22955432 | 0.31849464 | 0.24154715 | 1.31856097 | 0.18731593 |
| YDR034C-A | YDR034C-A | 5.35126702 | -0.3828807 | 0.29041008 | -1.3184139 | 0.18736514 |
| SND2 | YLR065C | 293.734341 | -0.1465695 | 0.11120328 | -1.3180321 | 0.18749289 |
| NAM7 | YMR080C | 2213.076 | 0.08563261 | 0.06503013 | 1.31681441 | 0.18790084 |
| NDI1 | YML120C | 845.433311 | -0.106326 | 0.08078831 | -1.3161061 | 0.18813842 |
| Xu_2009_SUTs30! | Xu_2009_SU | 4.64686924 | 0.37657854 | 0.28617887 | 1.31588523 | 0.18821257 |
| HRQ1 | YDR291W | 993.554725 | 0.09407922 | 0.07150136 | 1.31576817 | 0.18825187 |
| YMR230W-A | YMR230W-A | 13.639353 | -0.3781765 | 0.28796463 | -1.313274 | 0.18909062 |
| SHB17 | YKR043C | 2210.05655 | -0.0867913 | 0.06617434 | -1.3115551 | 0.18967028 |
| CNL1 | YDR357C | 1002.29969 | 0.10126031 | 0.07726025 | 1.31063927 | 0.18997966 |
| SNR57 | snR57 | 423.428158 | 0.13590691 | 0.10374899 | 1.30995897 | 0.19020972 |
| YEL053W-A | YEL053W-A | 5.20223067 | -0.3783929 | 0.28886256 | -1.3099409 | 0.19021583 |
| EMP70 | YLR083C | 1919.14776 | -0.0867858 | 0.06624202 | -1.3101318 | 0.19015125 |
| Xu_2009_SUTs70 | Xu_2009_SU | 13.8780032 | -0.3733266 | 0.28509017 | -1.3095035 | 0.19036385 |
| ARL3 | YPL051W | 302.804978 | -0.1517729 | 0.11589302 | -1.3095952 | 0.19033283 |
| YOR325W | YOR325W | 1.73867055 | 0.28748301 | 0.21961325 | 1.3090422 | 0.19052005 |
| YDR157W | YDR157W | 16.1914363 | -0.3690271 | 0.28232078 | -1.3071197 | 0.19117205 |
| YGR174W-A | YGR174W-A | 34.7511628 | -0.3040461 | 0.23288549 | -1.3055607 | 0.19170199 |
| HRR25 | YPL204W | 2815.67858 | 0.09143279 | 0.07003402 | 1.30554822 | 0.19170624 |
| YLR217W | YLR217W | 3.75493384 | 0.36094683 | 0.27661605 | 1.30486583 | 0.19193854 |
| YML099W-A | YML099W-A | 4.01424199 | 0.36203382 | 0.2774812 | 1.30471479 | 0.19198998 |
| Xu_2009_SUTs71 | Xu_2009_SU | 76.2776509 | 0.23896741 | 0.18319172 | 1.30446616 | 0.19207469 |
| SVF1 | YDR346C | 2675.67747 | 0.07707906 | 0.05910584 | 1.30408529 | 0.1922045 |
| YDL196W | YDL196W | 7.60088462 | 0.38607323 | 0.29620893 | 1.30338146 | 0.19244456 |
| FET4 | YMR319C | 835.80118 | -0.1147145 | 0.08808301 | -1.3023448 | 0.19279855 |

| | | | | | | |
|---|---|---|---|---|---|---|
| SNO1 | YMR095C | 18.7080321 | 0.35905952 | 0.27587745 | 1.30151819 | 0.19308114 |
| Xu_2009_SUTs34 | Xu_2009_SU | 86.3571524 | 0.21378314 | 0.16426984 | 1.30141437 | 0.19311666 |
| EPS1 | YIL005W | 1076.47735 | -0.0889718 | 0.0683919 | -1.3009121 | 0.19328854 |
| RPC31 | YNL151C | 657.332713 | 0.11611736 | 0.08926255 | 1.30085192 | 0.19330915 |
| Xu_2009_CUTs56 | Xu_2009_CU | 22.7255704 | 0.33670928 | 0.25904002 | 1.299835 | 0.19365753 |
| DIM1 | YPL266W | 575.907769 | -0.1072331 | 0.0824918 | -1.2999243 | 0.19362692 |
| FDH1 | YOR388C | 15.7326858 | -0.3693294 | 0.28424437 | -1.2993376 | 0.19382811 |
| Xu_2009_SUTs73 | Xu_2009_SU | 55.2049289 | 0.260277 | 0.20057679 | 1.29764266 | 0.19441016 |
| SNF11 | YDR073W | 287.354784 | -0.1397717 | 0.10770486 | -1.2977289 | 0.19438051 |
| YOR316C-A | YOR316C-A | 12.5128012 | -0.3810439 | 0.29376592 | -1.2971004 | 0.19459664 |
| Xu_2009_SUTs3 | Xu_2009_SU | 49.8522471 | 0.30444603 | 0.23477755 | 1.29674252 | 0.19471979 |
| IDP3 | YNL009W | 116.24945 | 0.1926314 | 0.14854406 | 1.29679637 | 0.19470126 |
| AIM23 | YJL131C | 217.003617 | -0.1643627 | 0.12677424 | -1.2964994 | 0.19480348 |
| REB1 | YBR049C | 1165.25419 | -0.0869841 | 0.06718146 | -1.294764 | 0.19540165 |
| Xu_2009_SUTs48 | Xu_2009_SU | 31.0037291 | -0.3141073 | 0.24266893 | -1.2943862 | 0.19553207 |
| tK(CUU)E2 | tK(CUU)E2 | 11.7748878 | 0.37699584 | 0.29128583 | 1.29424706 | 0.19558009 |
| MBB1 | YJL199C | 60.9144692 | 0.24563743 | 0.18980262 | 1.29417303 | 0.19560566 |
| RFX1 | YLR176C | 434.2551 | 0.12505737 | 0.09668533 | 1.29344718 | 0.19585644 |
| tT(AGU)H | tT(AGU)H | 4.61615942 | -0.3684034 | 0.28495624 | -1.2928419 | 0.19606576 |
| Xu_2009_CUTs37 | Xu_2009_CU | 0.71721856 | -0.2069254 | 0.16003239 | -1.2930222 | 0.19600339 |
| CCL1 | YPR025C | 520.704883 | -0.1133431 | 0.08766535 | -1.2929061 | 0.19604353 |
| tQ(UUG)E1 | tQ(UUG)E1 | 3.63271026 | -0.3485005 | 0.26961286 | -1.2925958 | 0.19615088 |
| GYL1 | YMR192W | 550.10311 | 0.10853868 | 0.08399937 | 1.29213677 | 0.19630978 |
| ADH4 | YGL256W | 590.548542 | -0.1174503 | 0.09090626 | -1.2919938 | 0.19635928 |
| STB4 | YMR019W | 675.429884 | 0.10843555 | 0.08395393 | 1.29160775 | 0.19649302 |
| Xu_2009_CUTs43 | Xu_2009_CU | 26.5651586 | 0.32074616 | 0.2484435 | 1.29102254 | 0.19669586 |
| YDR431W | YDR431W | 2.57601293 | 0.32743228 | 0.25376634 | 1.29029042 | 0.19694984 |
| YLR366W | YLR366W | 36.01448 | 0.29907387 | 0.23183851 | 1.29000945 | 0.19704738 |
| PTH1 | YHR189W | 146.879356 | 0.18578798 | 0.14406545 | 1.28960815 | 0.19718674 |
| tT(AGU)B | tT(AGU)B | 3.17229657 | -0.3473652 | 0.26948585 | -1.2889924 | 0.19740073 |
| YKR047W | YKR047W | 21.5342545 | -0.3521348 | 0.27335928 | -1.2881757 | 0.1976848 |
| Xu_2009_SUTs73 | Xu_2009_SU | 167.804199 | 0.16165785 | 0.12565606 | 1.28651061 | 0.19826492 |
| SNO2 | YNL334C | 84.4769092 | 0.21503805 | 0.16736732 | 1.28482701 | 0.19885274 |
| YPR114W | YPR114W | 1747.61543 | -0.080054 | 0.06231432 | -1.2846801 | 0.19890409 |
| Xu_2009_CUTs48 | Xu_2009_CU | 54.5522761 | 0.25897591 | 0.20164932 | 1.28428851 | 0.19904103 |
| CST9 | YLR394W | 330.584331 | -0.1298817 | 0.10112684 | -1.2843448 | 0.19902135 |
| RFC3 | YNL290W | 946.920111 | -0.096849 | 0.07542354 | -1.2840692 | 0.19911775 |
| Xu_2009_SUTs58 | Xu_2009_SU | 44.5707829 | 0.27608476 | 0.21522794 | 1.28275521 | 0.19957785 |
| SET4 | YJL105W | 34.7451962 | 0.30272508 | 0.23614413 | 1.28195047 | 0.19986002 |
| FMO1 | YHR176W | 126.992004 | -0.1873411 | 0.14633095 | -1.280256 | 0.20045511 |
| YGR067C | YGR067C | 115.819454 | 0.19231739 | 0.1502311 | 1.28014371 | 0.2004946 |

| | | | | | | |
|---|---|---|---|---|---|---|
| GID10 | YGR066C | 28.7901477 | -0.3186863 | 0.24903597 | -1.2796798 | 0.20065776 |
| MTC6 | YHR151C | 828.732845 | 0.0984971 | 0.07697905 | 1.27953117 | 0.20071007 |
| YHR182W | YHR182W | 635.465351 | -0.1041438 | 0.08140643 | -1.2793071 | 0.20078893 |
| tT(AGU)N1 | tT(AGU)N1 | 7.08585543 | -0.3771449 | 0.29509167 | -1.2780601 | 0.20122822 |
| YBR224W | YBR224W | 6.33242959 | -0.375114 | 0.29357119 | -1.2777617 | 0.20133347 |
| RRP9 | YPR137W | 2355.28501 | 0.08260112 | 0.06466961 | 1.27727881 | 0.20150383 |
| Xu_2009_CUTs82 | Xu_2009_CU | 11.20829 | 0.3760413 | 0.2945111 | 1.27683234 | 0.20166145 |
| YMR304C-A | YMR304C-A | 38.7952653 | -0.292275 | 0.22889562 | -1.2768921 | 0.20164033 |
| PRY3 | YJL078C | 1369.93684 | -0.0914641 | 0.07169061 | -1.2758165 | 0.20202038 |
| Xu_2009_CUTs14 | Xu_2009_CU | 9.09512271 | -0.3777764 | 0.29615843 | -1.2755889 | 0.20210088 |
| Xu_2009_SUTs21 | Xu_2009_SU | 0.93730805 | 0.22821329 | 0.17899739 | 1.2749532 | 0.20232581 |
| YAL026C-A | YAL026C-A | 2.71725253 | -0.3210809 | 0.25193829 | -1.2744426 | 0.20250659 |
| YPR137C-B | YPR137C-B | 23.4264974 | 0.34831949 | 0.27347407 | 1.2736838 | 0.2027755 |
| SMC3 | YJL074C | 745.219657 | 0.10396678 | 0.08166377 | 1.27310775 | 0.20297981 |
| BOI1 | YBL085W | 1906.6733 | -0.0787451 | 0.06189066 | -1.2723262 | 0.20325724 |
| SPC24 | YMR117C | 207.471504 | 0.16736326 | 0.13156598 | 1.27208615 | 0.20334251 |
| VPS51 | YKR020W | 245.895684 | -0.1373285 | 0.10796727 | -1.2719453 | 0.20339256 |
| Xu_2009_SUTs72 | Xu_2009_SU | 2.61129833 | 0.32719208 | 0.25727646 | 1.27175286 | 0.20346094 |
| ATP22 | YDR350C | 728.396128 | -0.0957953 | 0.07535868 | -1.2711918 | 0.20366041 |
| Xu_2009_SUTs67 | Xu_2009_SU | 17.2190384 | 0.35109591 | 0.27626348 | 1.27087342 | 0.20377368 |
| YOL162W | YOL162W | 31.4876512 | 0.31011312 | 0.24420544 | 1.2698862 | 0.20412517 |
| tD(GUC)J2 | tD(GUC)J2 | 1.46698723 | 0.27182942 | 0.21425395 | 1.26872535 | 0.20453904 |
| ZIP1 | YDR285W | 287.809253 | 0.14285481 | 0.11259316 | 1.26876989 | 0.20452315 |
| Xu_2009_CUTs15 | Xu_2009_CU | 43.3630878 | 0.29335636 | 0.23132471 | 1.26815834 | 0.20474141 |
| PAU15 | YIR041W | 3.59790471 | 0.3421936 | 0.26987381 | 1.26797634 | 0.2048064 |
| JSN1 | YJR091C | 3630.59432 | 0.0731887 | 0.05773761 | 1.26760879 | 0.20493769 |
| Xu_2009_SUTs76 | Xu_2009_SU | 202.780217 | 0.15893511 | 0.12549214 | 1.26649463 | 0.20533605 |
| Xu_2009_CUTs11 | Xu_2009_CU | 7.34089522 | 0.37422853 | 0.29564403 | 1.26580782 | 0.20558189 |
| TAN1 | YGL232W | 393.256178 | -0.1186987 | 0.09377478 | -1.2657845 | 0.20559025 |
| Xu_2009_SUTs33 | Xu_2009_SU | 16.0782519 | 0.35988412 | 0.28455174 | 1.26474053 | 0.20596436 |
| PRP9 | YDL030W | 405.37491 | 0.13069347 | 0.10337283 | 1.26429223 | 0.20612516 |
| YJL218W | YJL218W | 37.037915 | -0.3141163 | 0.24865441 | -1.2632647 | 0.20649407 |
| HEM1 | YDR232W | 2291.13012 | -0.1005223 | 0.07961748 | -1.2625654 | 0.20674541 |
| CDC4 | YFL009W | 1147.68263 | -0.0825628 | 0.0654426 | -1.2616068 | 0.2070903 |
| HXT13 | YEL069C | 3.51126485 | 0.34005863 | 0.26959359 | 1.26137508 | 0.20717374 |
| CBT1 | YKL208W | 154.451198 | -0.1812993 | 0.14376829 | -1.261052 | 0.2072901 |
| PPT2 | YPL148C | 232.482345 | 0.15765783 | 0.12509547 | 1.26030006 | 0.20756114 |
| DPA10 | YLR307C-A | 65.283609 | 0.23604628 | 0.18732964 | 1.26005835 | 0.20764831 |
| KHA1 | YJL094C | 1222.35497 | -0.0885185 | 0.07034761 | -1.2583011 | 0.20828287 |
| YHR130C | YHR130C | 37.1285118 | -0.2911247 | 0.23148397 | -1.2576454 | 0.20852004 |
| Xu_2009_CUTs3 | Xu_2009_CU | 53.7657388 | 0.28170465 | 0.22401206 | 1.2575423 | 0.20855733 |

| | | | | | | |
|---|---|---|---|---|---|---|
| CLB4 | YLR210W | 266.787399 | -0.1394354 | 0.11092128 | -1.2570664 | 0.20872959 |
| UBP16 | YPL072W | 366.311667 | -0.1176505 | 0.0937158 | -1.2553967 | 0.2093348 |
| Xu_2009_CUTs25 | Xu_2009_CU | 5.57380246 | 0.36655379 | 0.29223736 | 1.25430161 | 0.2097324 |
| Xu_2009_SUTs72 | Xu_2009_SU | 11.7601021 | 0.36872487 | 0.2939447 | 1.25440218 | 0.20969586 |
| PHM7 | YOL084W | 582.666835 | -0.1308717 | 0.10433837 | -1.2543009 | 0.20973265 |
| Xu_2009_CUTs58 | Xu_2009_CU | 4.79501532 | 0.3575813 | 0.28522139 | 1.25369737 | 0.20995202 |
| TDP1 | YBR223C | 377.424682 | 0.12397344 | 0.09889023 | 1.25364704 | 0.20997032 |
| YTM1 | YOR272W | 1620.36514 | -0.0927538 | 0.07401679 | -1.2531459 | 0.21015261 |
| YER172C-A | YER172C-A | 1.51046974 | 0.25996447 | 0.20747651 | 1.25298267 | 0.21021201 |
| Xu_2009_CUTs87 | Xu_2009_CU | 32.5592572 | 0.29343267 | 0.2344323 | 1.25167341 | 0.21068889 |
| GYP8 | YFL027C | 274.806655 | -0.1402429 | 0.11213091 | -1.2507067 | 0.21104149 |
| SBE22 | YHR103W | 2109.49021 | 0.06855534 | 0.05483047 | 1.25031474 | 0.2111846 |
| YOR072W | YOR072W | 12.0800962 | 0.36367051 | 0.29089203 | 1.2501907 | 0.21122989 |
| ALK2 | YBL009W | 383.134739 | 0.13224484 | 0.10582126 | 1.24970012 | 0.21140911 |
| AST2 | YER101C | 242.365351 | -0.1423521 | 0.11393854 | -1.249376 | 0.21152759 |
| TPP1 | YMR156C | 178.076504 | -0.1637301 | 0.13112089 | -1.2486955 | 0.21177647 |
| Xu_2009_SUTs17 | Xu_2009_SU | 57.3802018 | 0.24893877 | 0.19939741 | 1.24845543 | 0.21186432 |
| ALT2 | YDR111C | 351.589819 | -0.128529 | 0.10294759 | -1.2484901 | 0.21185164 |
| Xu_2009_SUTs94 | Xu_2009_SU | 27.4804156 | 0.30693108 | 0.24588902 | 1.24825047 | 0.21193935 |
| RER2 | YBR002C | 762.587576 | -0.099935 | 0.08006483 | -1.2481763 | 0.2119665 |
| Xu_2009_SUTs66 | Xu_2009_SU | 261.623525 | 0.13837514 | 0.11092965 | 1.24741339 | 0.21224596 |
| Xu_2009_SUTs81 | Xu_2009_SU | 61.3711454 | -0.2542427 | 0.20382053 | -1.2473849 | 0.2122564 |
| YNL067W-B | YNL067W-B | 4.95582431 | 0.3589641 | 0.28783466 | 1.24711909 | 0.21235383 |
| NFI1 | YOR156C | 710.631432 | 0.1109431 | 0.08895839 | 1.24713484 | 0.21234806 |
| YPL251W | YPL251W | 2.31359579 | 0.30968646 | 0.24833537 | 1.24704937 | 0.2123794 |
| SKG6 | YHR149C | 1313.61867 | 0.09591574 | 0.07696527 | 1.24622103 | 0.21268326 |
| RRP6 | YOR001W | 946.858571 | -0.0907814 | 0.07287285 | -1.2457506 | 0.21285598 |
| tP(UGG)O3 | tP(UGG)O3 | 19.3822827 | -0.3474777 | 0.2789943 | -1.2454652 | 0.21296081 |
| PET111 | YMR257C | 589.99708 | -0.1080305 | 0.08675973 | -1.2451684 | 0.21306985 |
| FRE8 | YLR047C | 502.69625 | 0.10712664 | 0.08607739 | 1.24453871 | 0.21330136 |
| AI5_ALPHA | Q0070 | 13.3939792 | -0.3619029 | 0.29104058 | -1.2434792 | 0.21369131 |
| ATG31 | YDR022C | 135.466682 | -0.1802372 | 0.14496168 | -1.2433441 | 0.21374106 |
| ADF1 | YCL058W-A | 79.3581669 | 0.22403547 | 0.1802285 | 1.24306351 | 0.21384443 |
| IRC16 | YPR038W | 11.4356055 | 0.36549738 | 0.29402644 | 1.2430766 | 0.21383961 |
| SNR10 | snR10 | 34.5288905 | 0.30001286 | 0.24161986 | 1.24167301 | 0.21435723 |
| Xu_2009_SUTs24 | Xu_2009_SU | 2.62754589 | -0.3162164 | 0.25466936 | -1.2416744 | 0.2143567 |
| Xu_2009_SUTs79 | Xu_2009_SU | 26.4628339 | 0.33770456 | 0.27200625 | 1.24153234 | 0.21440916 |
| YUR1 | YJL139C | 1137.54903 | 0.08759932 | 0.07058147 | 1.24110928 | 0.21456538 |
| MTC2 | YKL098W | 381.172765 | 0.11844278 | 0.09542553 | 1.24120652 | 0.21452947 |
| GAB1 | YLR459W | 438.543313 | -0.1152527 | 0.09286299 | -1.2411053 | 0.21456686 |
| FMP33 | YJL161W | 236.525383 | 0.16181025 | 0.13042237 | 1.24066327 | 0.21473017 |

| | | | | | | |
|---|---|---|---|---|---|---|
| Xu_2009_CUTs85 | Xu_2009_CU | 14.0174267 | 0.3582208 | 0.28886516 | 1.24009693 | 0.21493954 |
| RRP36 | YOR287C | 400.385376 | 0.12966996 | 0.1046374 | 1.2392315 | 0.21525978 |
| YDR034W-B | YDR034W-B | 97.0923746 | 0.20389161 | 0.16456373 | 1.2389827 | 0.2153519 |
| Xu_2009_CUTs71 | Xu_2009_CU | 0.92228931 | -0.2065047 | 0.16682133 | -1.2378793 | 0.21576081 |
| PAU5 | YFL020C | 20.361466 | -0.3304141 | 0.26690743 | -1.2379351 | 0.21574012 |
| SSN8 | YNL025C | 322.756159 | -0.1293721 | 0.10450154 | -1.2379918 | 0.21571908 |
| Xu_2009_CUTs11 | Xu_2009_CU | 5.48054761 | 0.35946175 | 0.29045771 | 1.23757003 | 0.21587553 |
| RMA1 | YKL132C | 234.597165 | 0.14172856 | 0.1146741 | 1.23592485 | 0.2164865 |
| RRP17 | YDR412W | 1012.46711 | 0.10449571 | 0.0845862 | 1.2353753 | 0.21669086 |
| Xu_2009_SUTs50 | Xu_2009_SU | 20.956216 | -0.3264333 | 0.26440598 | -1.2345911 | 0.21698273 |
| YCK3 | YER123W | 926.146937 | 0.08703764 | 0.07051045 | 1.23439357 | 0.21705628 |
| YBR292C | YBR292C | 60.5013164 | 0.24050193 | 0.19486291 | 1.23421092 | 0.21712432 |
| VLD1 | YIR014W | 119.050209 | 0.19919521 | 0.16141542 | 1.23405314 | 0.2171831 |
| RBH2 | YJR030C | 336.350104 | -0.127542 | 0.10335119 | -1.2340641 | 0.21717901 |
| HAP2 | YGL237C | 325.27373 | -0.1227625 | 0.0995097 | -1.2336738 | 0.21732448 |
| YMR158C-A | YMR158C-A | 18.4962952 | -0.3395118 | 0.27521445 | -1.2336263 | 0.21734217 |
| GYP7 | YDL234C | 2179.99464 | 0.09327365 | 0.07561796 | 1.23348535 | 0.21739474 |
| YMR090W | YMR090W | 305.158732 | -0.1254748 | 0.10181192 | -1.2324174 | 0.2177932 |
| SMC5 | YOL034W | 748.495607 | 0.0963878 | 0.07822249 | 1.23222617 | 0.21786461 |
| CIR2 | YOR356W | 1127.96695 | 0.08044616 | 0.0652879 | 1.23217571 | 0.21788346 |
| IES6 | YEL044W | 423.987418 | -0.1190072 | 0.09661309 | -1.2317911 | 0.21802712 |
| CWC24 | YLR323C | 319.947987 | -0.1368701 | 0.11112657 | -1.2316595 | 0.21807629 |
| MRPL22 | YNL177C | 1081.10394 | -0.0858447 | 0.06972503 | -1.2311891 | 0.21825216 |
| Xu_2009_CUTs25 | Xu_2009_CU | 6.00904076 | 0.36185541 | 0.29397239 | 1.23091629 | 0.21835418 |
| SSP120 | YLR250W | 839.969639 | -0.1030506 | 0.08376063 | -1.2302986 | 0.21858532 |
| YEL074W | YEL074W | 18.6361829 | 0.33322252 | 0.27089249 | 1.2300914 | 0.21866288 |
| Xu_2009_CUTs7 | Xu_2009_CU | 8.61328225 | -0.3646964 | 0.29667392 | -1.2292838 | 0.21896543 |
| YBR116C | YBR116C | 0.83981909 | -0.2095631 | 0.17056771 | -1.2286212 | 0.21921387 |
| Xu_2009_SUTs26 | Xu_2009_SU | 25.6695081 | -0.3145447 | 0.25617573 | -1.2278476 | 0.2195042 |
| tS(AGA)B | tS(AGA)B | 5.20898519 | -0.3533554 | 0.28805479 | -1.2266953 | 0.21993715 |
| YER006C-A | YER006C-A | 9.02890004 | 0.36300171 | 0.29603089 | 1.22622918 | 0.22011244 |
| YPR130C | YPR130C | 13.8729085 | -0.3520912 | 0.28722555 | -1.2258351 | 0.22026074 |
| Xu_2009_SUTs12 | Xu_2009_SU | 20.0998577 | 0.32675152 | 0.26676495 | 1.22486675 | 0.22062543 |
| DRN1 | YGR093W | 541.029894 | -0.1124869 | 0.09184912 | -1.224692 | 0.22069128 |
| YPR003C | YPR003C | 526.852331 | 0.12027949 | 0.09822775 | 1.2244961 | 0.22076514 |
| SNX3 | YOR357C | 260.818146 | -0.157008 | 0.12829479 | -1.2238062 | 0.22102533 |
| YMR155W | YMR155W | 253.509746 | -0.1434792 | 0.11725511 | -1.2236497 | 0.22108441 |
| YPL257W | YPL257W | 125.625659 | 0.19221672 | 0.15728686 | 1.22207744 | 0.22167834 |
| ATG4 | YNL223W | 261.566681 | -0.1508808 | 0.12348251 | -1.2218798 | 0.22175308 |
| DGR2 | YKL121W | 592.283923 | 0.10749521 | 0.08807369 | 1.22051443 | 0.22226992 |
| FCY22 | YER060W-A | 66.7016275 | -0.2313217 | 0.18956415 | -1.2202817 | 0.22235811 |

| | | | | | | |
|---|---|---|---|---|---|---|
| YDR442W | YDR442W | 1.27895876 | 0.2478401 | 0.20316713 | 1.21988287 | 0.22250928 |
| Xu_2009_SUTs62 | Xu_2009_SU | 52.414649 | 0.26394602 | 0.21641599 | 1.21962347 | 0.22260764 |
| RTT102 | YGR275W | 545.398136 | 0.10289833 | 0.08437117 | 1.21959119 | 0.22261989 |
| YLR349W | YLR349W | 30.0342475 | 0.29895882 | 0.24514222 | 1.21953216 | 0.22264228 |
| YER076W-A | YER076W-A | 9.88966624 | 0.36007102 | 0.2953833 | 1.21899586 | 0.22284576 |
| Xu_2009_SUTs53 | Xu_2009_SU | 46.2965565 | 0.2612139 | 0.21434112 | 1.21868309 | 0.2229645 |
| Xu_2009_SUTs79 | Xu_2009_SU | 41.5115083 | 0.26518683 | 0.21766226 | 1.21834085 | 0.22309447 |
| MFA1 | YDR461W | 23.2283948 | 0.31903454 | 0.26184917 | 1.21839051 | 0.22307561 |
| YBL029W | YBL029W | 152.319892 | 0.16934149 | 0.13902927 | 1.21802764 | 0.22321347 |
| YPL150W | YPL150W | 830.356059 | 0.08641141 | 0.07094058 | 1.21808164 | 0.22319295 |
| MAD2 | YJL030W | 247.657851 | 0.14505422 | 0.11910133 | 1.21790594 | 0.22325972 |
| AFG3 | YER017C | 1750.6939 | 0.07160265 | 0.05882082 | 1.21730105 | 0.2234897 |
| YNR077C | YNR077C | 1.00635545 | 0.22935337 | 0.1885197 | 1.2166016 | 0.22375583 |
| Xu_2009_SUTs69 | Xu_2009_SU | 62.8162859 | 0.23486864 | 0.19316185 | 1.2159163 | 0.22401681 |
| RKM5 | YLR137W | 518.324219 | 0.10606147 | 0.08728642 | 1.21509705 | 0.22432908 |
| GPX2 | YBR244W | 351.457889 | -0.1334584 | 0.1098979 | -1.2143851 | 0.22460072 |
| IPI3 | YNL182C | 1465.24348 | 0.09401036 | 0.0774091 | 1.21446129 | 0.22457162 |
| PML39 | YML107C | 142.482482 | -0.172405 | 0.14210277 | -1.2132413 | 0.22503758 |
| YMR122C | YMR122C | 6.55215971 | 0.35664509 | 0.29421751 | 1.21218171 | 0.22544283 |
| ATG40 | YOR152C | 158.862794 | -0.1731732 | 0.14288077 | -1.212012 | 0.22550777 |
| AI1 | Q0050 | 9764.01927 | -0.0685075 | 0.05654583 | -1.2115394 | 0.22568874 |
| Xu_2009_SUTs32 | Xu_2009_SU | 4.74348461 | 0.34107741 | 0.28162361 | 1.21111084 | 0.22585293 |
| Xu_2009_SUTs22 | Xu_2009_SU | 42.4070904 | 0.28468715 | 0.23540267 | 1.20936247 | 0.22652362 |
| Xu_2009_SUTs49 | Xu_2009_SU | 29.4986465 | 0.30018284 | 0.24825394 | 1.20917653 | 0.22659503 |
| BDH2 | YAL061W | 605.304598 | -0.1256388 | 0.10391222 | -1.209086 | 0.22662982 |
| LCB5 | YLR260W | 570.267364 | -0.1059725 | 0.08765643 | -1.2089528 | 0.22668099 |
| Xu_2009_CUTs42 | Xu_2009_CU | 10.7381212 | 0.35424766 | 0.29309916 | 1.20862734 | 0.22680605 |
| Xu_2009_CUTs61 | Xu_2009_CU | 14.3719804 | 0.34831061 | 0.28834799 | 1.20795228 | 0.22706562 |
| YER087C-A | YER087C-A | 32.2803103 | -0.2889193 | 0.23921016 | -1.2078052 | 0.22712222 |
| YAL016C-A | YAL016C-A | 6.78650902 | 0.35641244 | 0.29529004 | 1.20699108 | 0.22743558 |
| SYG1 | YIL047C | 2509.09166 | 0.08508787 | 0.07051285 | 1.20670025 | 0.2275476 |
| SNF6 | YHL025W | 598.419599 | -0.1117544 | 0.09266495 | -1.2060053 | 0.22781544 |
| YFR054C | YFR054C | 22.3679752 | 0.31642579 | 0.26256816 | 1.20511869 | 0.22815748 |
| YRM1 | YOR172W | 779.904709 | 0.09720499 | 0.08068504 | 1.2047462 | 0.2283013 |
| RPA135 | YPR010C | 5486.67933 | 0.0915944 | 0.07604225 | 1.20451987 | 0.22838871 |
| SNR18 | snR18 | 64.4485663 | -0.2388744 | 0.19861579 | -1.2026956 | 0.22909412 |
| DYN3 | YMR299C | 356.210334 | 0.13107839 | 0.10898192 | 1.20275358 | 0.22907169 |
| ATX2 | YOR079C | 206.945356 | -0.1492066 | 0.12420351 | -1.2013073 | 0.22963203 |
| TCB1 | YOR086C | 3633.07206 | -0.0649502 | 0.05408716 | -1.2008434 | 0.22981194 |
| YFL031C-A | YFL031C-A | 1.93749558 | 0.28100651 | 0.23402766 | 1.20074057 | 0.22985185 |
| SNF12 | YNR023W | 557.899477 | 0.10522076 | 0.08775098 | 1.19908354 | 0.23049547 |

| | | | | | | |
|---|---|---|---|---|---|---|
| SSF1 | YHR066W | 1915.9623 | 0.09450976 | 0.07883106 | 1.19888998 | 0.23057073 |
| OPI11 | YPR044C | 1.75369872 | 0.27176376 | 0.22678105 | 1.19835304 | 0.23077961 |
| tT(AGU)I1 | tT(AGU)I1 | 4.061338 | -0.3363152 | 0.28079893 | -1.1977082 | 0.23103064 |
| HTL1 | YCR020W-B | 70.4288543 | -0.2293458 | 0.1915452 | -1.1973458 | 0.23117181 |
| MSL1 | YIR009W | 255.377692 | 0.1287922 | 0.10774002 | 1.195398 | 0.23193157 |
| NOT3 | YIL038C | 1081.10909 | 0.08415602 | 0.07044152 | 1.19469348 | 0.23220681 |
| BIL1 | YOR304C-A | 97.4580083 | 0.20248639 | 0.16962953 | 1.19369772 | 0.23259623 |
| YBR103C-A | YBR103C-A | 4.21026463 | 0.33772569 | 0.28320252 | 1.19252361 | 0.233056 |
| NUP84 | YDL116W | 1117.31002 | -0.0898461 | 0.07535326 | -1.1923326 | 0.23313086 |
| PFA5 | YDR459C | 283.04615 | -0.1269008 | 0.10647897 | -1.1917921 | 0.23334279 |
| YIL102C-A | YIL102C-A | 58.2622561 | -0.2409172 | 0.20222022 | -1.1913607 | 0.23351203 |
| PSR1 | YLL010C | 1465.84116 | -0.0855105 | 0.07185602 | -1.1900253 | 0.23403644 |
| YNL109W | YNL109W | 26.0530005 | 0.30915255 | 0.26027715 | 1.18778214 | 0.23491925 |
| CUR1 | YPR158W | 463.28473 | -0.1474351 | 0.12414712 | -1.1875839 | 0.2349974 |
| YPL068C | YPL068C | 611.304415 | -0.0986941 | 0.08320447 | -1.1861634 | 0.23555779 |
| Xu_2009_CUTs86 | Xu_2009_CU | 38.951536 | 0.26410044 | 0.22269503 | 1.18592879 | 0.23565042 |
| DUS1 | YML080W | 724.743041 | 0.11103933 | 0.09364166 | 1.18578986 | 0.2357053 |
| YPR202W | YPR202W | 25.732529 | 0.29750142 | 0.25104472 | 1.18505346 | 0.23599631 |
| tL(GAG)G | tL(GAG)G | 4.5543402 | 0.33748335 | 0.28481493 | 1.18492155 | 0.23604847 |
| YBR013C | YBR013C | 172.787487 | 0.15273313 | 0.12898609 | 1.18410541 | 0.23637134 |
| THI6 | YPL214C | 392.030649 | -0.1124528 | 0.09496114 | -1.1841979 | 0.23633472 |
| tY(GUA)D/SUP2 | tY(GUA)D | 9.98503013 | -0.3486313 | 0.29447047 | -1.1839262 | 0.23644228 |
| MEK1 | YOR351C | 8.21615273 | -0.3489468 | 0.29476263 | -1.1838229 | 0.23648318 |
| YEL030C-A | YEL030C-A | 3.67814389 | 0.32759538 | 0.27692718 | 1.18296579 | 0.2368227 |
| Xu_2009_SUTs72 | Xu_2009_SU | 177.312301 | -0.157134 | 0.132913 | -1.1822319 | 0.23711369 |
| PUL3 | YNR062C | 3.81984048 | -0.3240475 | 0.27431601 | -1.1812927 | 0.23748647 |
| Xu_2009_SUTs60 | Xu_2009_SU | 1.67659857 | -0.2597471 | 0.21999541 | -1.1806935 | 0.23772451 |
| YEL075W-A | YEL075W-A | 5.71091621 | 0.34239967 | 0.29004667 | 1.18049852 | 0.237802 |
| Xu_2009_CUTs70 | Xu_2009_CU | 4.74283123 | 0.33746479 | 0.28610132 | 1.17952895 | 0.23818762 |
| TIM22 | YDL217C | 334.591595 | -0.1268886 | 0.10758417 | -1.1794358 | 0.23822469 |
| Xu_2009_SUTs59 | Xu_2009_SU | 96.6828422 | -0.1970011 | 0.16745773 | -1.1764229 | 0.23942592 |
| FRA1 | YLL029W | 1679.91551 | -0.0719051 | 0.06114188 | -1.1760368 | 0.23958017 |
| YLR001C | YLR001C | 751.178274 | -0.0906915 | 0.07722895 | -1.1743202 | 0.24026678 |
| tA(UGC)O | tA(UGC)O | 0.81526209 | -0.1915299 | 0.1632005 | -1.1735863 | 0.24056079 |
| YER121W | YER121W | 126.215953 | 0.1801256 | 0.15346496 | 1.17372456 | 0.24050537 |
| FYV4 | YHR059W | 245.018403 | -0.1310695 | 0.11168818 | -1.1735303 | 0.2405832 |
| SDS3 | YIL084C | 295.028941 | 0.12307263 | 0.10485378 | 1.17375486 | 0.24049323 |
| SWI3 | YJL176C | 1656.89676 | 0.06912894 | 0.05889441 | 1.17377763 | 0.24048411 |
| Xu_2009_CUTs16 | Xu_2009_CU | 17.596407 | 0.31857381 | 0.27162118 | 1.17286074 | 0.24085166 |
| Xu_2009_SUTs66 | Xu_2009_SU | 29.2000381 | 0.28700339 | 0.2448134 | 1.17233531 | 0.24106246 |
| CLB1 | YGR108W | 2382.48611 | -0.067863 | 0.05788934 | -1.1722882 | 0.24108138 |

| | | | | | | |
|---|---|---|---|---|---|---|
| SNZ1 | YMR096W | 376.622495 | -0.1182625 | 0.10088941 | -1.1721992 | 0.24111709 |
| PGA1 | YNL158W | 330.960281 | 0.12006884 | 0.10244581 | 1.17202287 | 0.24118788 |
| POG1 | YIL122W | 848.727607 | -0.0846669 | 0.07224719 | -1.1719052 | 0.24123511 |
| YBR134W | YBR134W | 4.10324606 | 0.32583876 | 0.27812324 | 1.17156251 | 0.24137275 |
| Xu_2009_SUTs43 | Xu_2009_SU | 30.6097844 | 0.28335207 | 0.24195704 | 1.17108421 | 0.24156493 |
| PAD1 | YDR538W | 140.374481 | -0.1649464 | 0.14092478 | -1.1704573 | 0.241817 |
| NCW2 | YLR194C | 4786.95167 | -0.1475365 | 0.1260487 | -1.1704718 | 0.24181114 |
| YBP2 | YGL060W | 399.374662 | 0.11715215 | 0.10013146 | 1.16998341 | 0.24200765 |
| YMR082C | YMR082C | 1.13057215 | 0.22964651 | 0.19626406 | 1.17008947 | 0.24196496 |
| DPI34 | YOR114W | 157.087613 | 0.16770553 | 0.14334269 | 1.16996223 | 0.24201617 |
| TRM1 | YDR120C | 1400.5214 | 0.08762507 | 0.07490953 | 1.16974539 | 0.24210345 |
| SPO74 | YGL170C | 50.5846172 | -0.2365576 | 0.20238909 | -1.1688258 | 0.24247383 |
| SNR64 | snR64 | 290.809641 | -0.1415422 | 0.12111341 | -1.1686752 | 0.24253453 |
| YPR123C | YPR123C | 83.0476031 | 0.2012886 | 0.17235193 | 1.16789295 | 0.24284995 |
| CPR2 | YHR057C | 341.636722 | -0.1199943 | 0.10282869 | -1.166934 | 0.24323701 |
| tE(CUC)I | tE(CUC)I | 1.1817744 | -0.2328467 | 0.19957458 | -1.1667154 | 0.24332533 |
| Xu_2009_CUTs20 | Xu_2009_CU | 11.6814386 | 0.34156474 | 0.29284625 | 1.16636201 | 0.24346811 |
| Xu_2009_SUTs32 | Xu_2009_SU | 136.852318 | 0.16507713 | 0.14165285 | 1.16536398 | 0.24387169 |
| Xu_2009_SUTs64 | Xu_2009_SU | 86.7987623 | -0.208274 | 0.17872756 | -1.1653156 | 0.24389128 |
| YGL123C-A | YGL123C-A | 6.09113917 | 0.33700454 | 0.28917405 | 1.1654038 | 0.24385558 |
| Xu_2009_CUTs31 | Xu_2009_CU | 22.2977531 | 0.3076111 | 0.26409184 | 1.16478833 | 0.24410468 |
| ALB1 | YJL122W | 1166.64189 | -0.1040332 | 0.08937183 | -1.1640487 | 0.24440426 |
| MNN10 | YDR245W | 903.991032 | -0.0809137 | 0.06959455 | -1.1626449 | 0.24497359 |
| Xu_2009_CUTs59 | Xu_2009_CU | 40.9302488 | 0.26278019 | 0.22609237 | 1.16226916 | 0.24512615 |
| SRL4 | YPL033C | 40.9219611 | -0.2664678 | 0.22925974 | -1.1622965 | 0.24511507 |
| NIF3 | YGL221C | 1474.36318 | 0.08288447 | 0.07133851 | 1.16184742 | 0.24529745 |
| SVL3 | YPL032C | 2234.31267 | 0.06914161 | 0.0595325 | 1.16140948 | 0.24547542 |
| RRP8 | YDR083W | 1008.12705 | 0.10946896 | 0.09432321 | 1.16057286 | 0.24581565 |
| Xu_2009_CUTs55 | Xu_2009_CU | 4.94307073 | -0.3197127 | 0.27555569 | -1.1602471 | 0.24594823 |
| PGU1 | YJR153W | 12.9381435 | -0.3371093 | 0.2906689 | -1.1597709 | 0.24614211 |
| Xu_2009_CUTs17 | Xu_2009_CU | 13.5859957 | 0.33145015 | 0.28638161 | 1.15737234 | 0.24712027 |
| POC4 | YPL144W | 143.073139 | 0.15882858 | 0.13725629 | 1.15716791 | 0.24720377 |
| RSA3 | YLR221C | 1265.83526 | 0.08375117 | 0.07239445 | 1.15687282 | 0.24732433 |
| LAG2 | YOL025W | 424.765648 | 0.10151087 | 0.08776761 | 1.15658686 | 0.2474412 |
| Xu_2009_CUTs70 | Xu_2009_CU | 27.8348401 | 0.28730574 | 0.24863769 | 1.15551966 | 0.24787769 |
| YLR042C | YLR042C | 123.676335 | -0.1726233 | 0.14941984 | -1.1552903 | 0.24797155 |
| Xu_2009_SUTs8 | Xu_2009_SU | 3.60615193 | 0.31699319 | 0.27456798 | 1.15451625 | 0.24828858 |
| YRF1-3 | YGR296W | 3.45753012 | 0.3123412 | 0.27059384 | 1.15428053 | 0.24838518 |
| YJL136W-A | YJL136W-A | 1.63166586 | 0.2591323 | 0.22460565 | 1.1537212 | 0.24861449 |
| Xu_2009_CUTs38 | Xu_2009_CU | 2.94706827 | 0.3006235 | 0.26081775 | 1.15261904 | 0.24906678 |
| Xu_2009_SUTs75 | Xu_2009_SU | 180.357892 | 0.15574313 | 0.13530334 | 1.15106639 | 0.24970492 |

| | | | | | | |
|---|---|---|---|---|---|---|
| YDR433W | YDR433W | 18.3659307 | -0.3201417 | 0.27816362 | -1.1509115 | 0.24976864 |
| RGD1 | YBR260C | 890.340151 | -0.0858266 | 0.07467874 | -1.1492769 | 0.25044183 |
| GLN3 | YER040W | 913.745194 | 0.07869187 | 0.06849174 | 1.14892507 | 0.25058688 |
| YFH7 | YFR007W | 348.188278 | -0.1112749 | 0.09683724 | -1.1490924 | 0.25051788 |
| CIN4 | YMR138W | 107.492774 | 0.19031519 | 0.16564843 | 1.14891027 | 0.25059298 |
| tP(UGG)Q | tP(UGG)Q | 2.45306381 | -0.2882324 | 0.2509312 | -1.148651 | 0.25069993 |
| YND1 | YER005W | 866.357597 | -0.0856013 | 0.07456928 | -1.1479438 | 0.25099175 |
| tG(GCC)F1/SUF20 | tG(GCC)F1 | 3.24116264 | -0.2930675 | 0.25558267 | -1.1466643 | 0.25152038 |
| TPO5 | YKL174C | 840.962932 | 0.08837965 | 0.07707665 | 1.14664621 | 0.25152787 |
| WHI4 | YDL224C | 1379.08081 | 0.07368653 | 0.06428208 | 1.14629966 | 0.25167118 |
| IRE1 | YHR079C | 1224.59857 | 0.0765262 | 0.06676315 | 1.14623408 | 0.25169831 |
| RSC58 | YLR033W | 684.137519 | 0.0899571 | 0.07848434 | 1.14617896 | 0.25172111 |
| CGI121 | YML036W | 173.864579 | -0.1500725 | 0.13096863 | -1.1458658 | 0.25185068 |
| Xu_2009_SUTs67 | Xu_2009_SU | 9.71587377 | 0.33707503 | 0.2942361 | 1.14559373 | 0.25196329 |
| HSF1 | YGL073W | 1432.66886 | 0.07037911 | 0.06147721 | 1.14480003 | 0.252292 |
| Xu_2009_CUTs11 | Xu_2009_CU | 18.0613046 | 0.31232308 | 0.27338064 | 1.14244769 | 0.25326798 |
| CAF4 | YKR036C | 658.392704 | 0.09636154 | 0.08434658 | 1.14244751 | 0.25326805 |
| Xu_2009_CUTs86 | Xu_2009_CU | 7.82485457 | -0.3353752 | 0.29359924 | -1.1422889 | 0.25333395 |
| KIN1 | YDR122W | 1690.92423 | -0.0799842 | 0.07002975 | -1.1421466 | 0.25339309 |
| CDC37 | YDR168W | 1755.67209 | -0.0697513 | 0.06111758 | -1.1412641 | 0.25376002 |
| Xu_2009_SUTs73 | Xu_2009_SU | 185.435442 | 0.1450927 | 0.12751941 | 1.13780869 | 0.25520038 |
| tV(UAC)B | tV(UAC)B | 2.43169581 | -0.2728201 | 0.23995394 | -1.1369687 | 0.25555138 |
| YPL088W | YPL088W | 1473.11066 | 0.06942119 | 0.06113639 | 1.13551336 | 0.25616029 |
| YGR210C | YGR210C | 1288.88245 | 0.07363642 | 0.06488926 | 1.13480145 | 0.25645852 |
| RPB4 | YJL140W | 1120.69006 | -0.0815279 | 0.07184641 | -1.1347522 | 0.25647914 |
| MAK3 | YPR051W | 473.696896 | -0.1075122 | 0.09479288 | -1.1341798 | 0.25671912 |
| PDR18 | YNR070W | 70.0083131 | -0.2132314 | 0.18805719 | -1.1338649 | 0.25685124 |
| YDR278C | YDR278C | 0.83499829 | 0.19563376 | 0.1725865 | 1.1335403 | 0.25698743 |
| YPR084W | YPR084W | 560.101346 | -0.0938056 | 0.08284039 | -1.1323657 | 0.25748074 |
| Xu_2009_CUTs63 | Xu_2009_CU | 53.5654851 | 0.23322796 | 0.20598571 | 1.13225311 | 0.25752804 |
| FLO8 | YER109C | 786.069661 | 0.08382919 | 0.07409888 | 1.13131518 | 0.25792246 |
| TAH11 | YJR046W | 791.556725 | 0.08232115 | 0.07277246 | 1.13121295 | 0.25796547 |
| YPL039W | YPL039W | 367.4854 | 0.11014905 | 0.09743552 | 1.13048145 | 0.25827341 |
| BDS1 | YOL164W | 216.517242 | 0.13241102 | 0.1172875 | 1.12894401 | 0.25892145 |
| tV(AAC)G2 | tV(AAC)G2 | 2.77144895 | 0.29466644 | 0.26136299 | 1.12742219 | 0.25956402 |
| PHO4 | YFR034C | 304.451466 | -0.1194196 | 0.1059399 | -1.127239 | 0.25964146 |
| ATG10 | YLL042C | 36.7973806 | 0.26237843 | 0.23280422 | 1.12703468 | 0.25972782 |
| Xu_2009_SUTs2 | Xu_2009_SU | 26.6380765 | 0.28971989 | 0.2572711 | 1.12612683 | 0.26011184 |
| TRS120 | YDR407C | 1356.30072 | 0.07550231 | 0.0670454 | 1.12613702 | 0.26010753 |
| MSH2 | YOL090W | 915.15798 | -0.0895472 | 0.07950475 | -1.1263121 | 0.26003342 |
| Xu_2009_CUTs87 | Xu_2009_CU | 6.07518239 | 0.33034059 | 0.29347747 | 1.125608 | 0.26033148 |

| | | | | | | |
|---|---|---|---|---|---|---|
| ECM34 | YHL043W | 3.63838585 | 0.3080148 | 0.27366352 | 1.12552378 | 0.26036715 |
| SMD3 | YLR147C | 590.506961 | 0.11195979 | 0.09946485 | 1.12562169 | 0.26032568 |
| YML116W-A | YML116W-A | 43.0966822 | -0.2558548 | 0.22727696 | -1.12574 | 0.26027561 |
| PYK2 | YOR347C | 469.16583 | -0.1036289 | 0.09205081 | -1.1257792 | 0.26025898 |
| Xu_2009_CUTs57 | Xu_2009_CU | 10.3356133 | 0.33144473 | 0.29463568 | 1.12493071 | 0.2606184 |
| ENV11 | YGR071C | 470.040254 | 0.09777691 | 0.08692688 | 1.12481794 | 0.26066619 |
| SNR6 | snR6 | 4.8030726 | 0.3219932 | 0.28632795 | 1.12456082 | 0.26077519 |
| tL(UAA)B1 | tL(UAA)B1 | 1.76040259 | -0.2380676 | 0.21179107 | -1.1240682 | 0.26098411 |
| Xu_2009_CUTs80 | Xu_2009_CU | 24.7529037 | 0.28506471 | 0.25369585 | 1.12364752 | 0.26116259 |
| CKI1 | YLR133W | 1444.39418 | -0.0695106 | 0.06189866 | -1.1229746 | 0.26144827 |
| ECM8 | YBR076W | 72.8445476 | 0.20863529 | 0.18595823 | 1.12194704 | 0.26188496 |
| YBL036C | YBL036C | 1012.58885 | 0.08520943 | 0.07602744 | 1.12077205 | 0.26238491 |
| EMC5 | YIL027C | 316.091163 | -0.1146842 | 0.10241719 | -1.1197752 | 0.26280956 |
| Xu_2009_CUTs65 | Xu_2009_CU | 9.9877179 | 0.32855012 | 0.29351108 | 1.11937892 | 0.26297852 |
| YPL199C | YPL199C | 527.097875 | -0.0959435 | 0.08579743 | -1.1182562 | 0.26345758 |
| Xu_2009_CUTs18 | Xu_2009_CU | 15.0099281 | -0.3183919 | 0.28480193 | -1.1179414 | 0.263592 |
| Xu_2009_SUTs33 | Xu_2009_SU | 4.18134373 | 0.30960834 | 0.27723968 | 1.11675336 | 0.26409979 |
| Xu_2009_SUTs47 | Xu_2009_SU | 58.3872653 | -0.22844 | 0.20464269 | -1.1162871 | 0.26429928 |
| SSL1 | YLR005W | 745.660627 | -0.0887288 | 0.0796019 | -1.1146569 | 0.26499747 |
| FMP42 | YMR221C | 716.73116 | -0.0833182 | 0.07474211 | -1.1147421 | 0.26496095 |
| YOL099C | YOL099C | 6.41657832 | -0.329359 | 0.29547566 | -1.114674 | 0.26499016 |
| YCR047W-A | YCR047W-A | 6.50686071 | 0.32814583 | 0.29448614 | 1.11429974 | 0.26515062 |
| RPC10 | YHR143W-A | 901.8787 | -0.1106259 | 0.0993162 | -1.1138754 | 0.26533265 |
| YOR192C-C | YOR192C-C | 47.8676323 | 0.23087495 | 0.20728944 | 1.11378058 | 0.26537333 |
| YBL008W-A | YBL008W-A | 3.35648291 | -0.2866072 | 0.25745402 | -1.1132365 | 0.26560689 |
| CLD1 | YGR110W | 79.374075 | -0.1961629 | 0.17621654 | -1.1131922 | 0.2656259 |
| CYM1 | YDR430C | 1520.95833 | -0.0687808 | 0.06179531 | -1.1130421 | 0.26569036 |
| CKA1 | YIL035C | 992.810557 | -0.0783798 | 0.07042556 | -1.1129458 | 0.2657317 |
| ROT1 | YMR200W | 1940.38012 | -0.0668565 | 0.06011076 | -1.1122218 | 0.26604278 |
| PAU10 | YDR542W | 6.74972944 | 0.32779594 | 0.29501189 | 1.11112792 | 0.26651329 |
| Xu_2009_CUTs30 | Xu_2009_CU | 14.1393045 | 0.31745736 | 0.28574314 | 1.11098856 | 0.26657327 |
| TRP3 | YKL211C | 1029.60084 | -0.0821301 | 0.07393113 | -1.1108999 | 0.26661144 |
| YPL035C | YPL035C | 2.52608425 | 0.27990355 | 0.25236867 | 1.10910576 | 0.26738456 |
| RPB11 | YOL005C | 842.247987 | -0.0814604 | 0.07347543 | -1.1086754 | 0.26757025 |
| JEN1 | YKL217W | 152.247058 | 0.15710348 | 0.14171689 | 1.10857273 | 0.26761455 |
| YGL108C | YGL108C | 400.564523 | 0.11764937 | 0.10615076 | 1.10832344 | 0.26772216 |
| YAR062W | YAR062W | 1.75595669 | 0.25551162 | 0.23058074 | 1.10812215 | 0.26780907 |
| YLL054C | YLL054C | 493.390943 | 0.09288188 | 0.08385627 | 1.10763202 | 0.26802077 |
| YML083C | YML083C | 3.46361895 | -0.2918531 | 0.26348244 | -1.1076759 | 0.26800182 |
| RRT6 | YGL146C | 54.9863948 | -0.2314087 | 0.20904936 | -1.106957 | 0.26831251 |
| CUZ1 | YNL155W | 1423.2915 | 0.07106462 | 0.06421481 | 1.10667024 | 0.26843653 |

| | | | | | | |
|---|---|---|---|---|---|---|
| Xu_2009_SUTs71( | Xu_2009_SU | 44.6352608 | 0.23995372 | 0.21699233 | 1.1058166 | 0.26880591 |
| YIM2 | YMR151W | 39.7468937 | -0.2488999 | 0.22507859 | -1.1058356 | 0.26879767 |
| PXA1 | YPL147W | 210.533487 | -0.1378808 | 0.12466995 | -1.1059669 | 0.26874086 |
| Xu_2009_SUTs39 | Xu_2009_SU | 68.223648 | 0.20866407 | 0.18884751 | 1.10493423 | 0.26918809 |
| YGR073C | YGR073C | 25.049306 | -0.2931594 | 0.26531248 | -1.1049588 | 0.26917743 |
| Xu_2009_SUTs81 | Xu_2009_SU | 22.4880403 | 0.2918499 | 0.26436582 | 1.10396229 | 0.2696095 |
| GPI15 | YNL038W | 384.324005 | 0.11888963 | 0.10777406 | 1.10313775 | 0.26996735 |
| YIL165C | YIL165C | 51.4448799 | -0.229914 | 0.20857173 | -1.1023256 | 0.27032014 |
| YFL040W | YFL040W | 13.5980075 | -0.3161807 | 0.28702784 | -1.101568 | 0.27064952 |
| RIM2 | YBR192W | 540.196252 | -0.0934374 | 0.08483394 | -1.1014148 | 0.27071616 |
| Xu_2009_CUTs63 | Xu_2009_CU | 0.86619601 | 0.18473204 | 0.16778691 | 1.10099197 | 0.27090015 |
| TBS1 | YBR150C | 495.817053 | 0.09920576 | 0.09016771 | 1.10023592 | 0.27122934 |
| TBF1 | YPL128C | 1110.84109 | -0.0786916 | 0.07157943 | -1.0993604 | 0.27161088 |
| Xu_2009_CUTs12 | Xu_2009_CU | 23.8597855 | 0.27951384 | 0.25437529 | 1.09882467 | 0.27184455 |
| YBR071W | YBR071W | 1211.77636 | -0.0701084 | 0.06380731 | -1.0987522 | 0.27187619 |
| YOR161W-A | YOR161W-A | 4.06021473 | -0.2972875 | 0.27054881 | -1.0988312 | 0.27184168 |
| CIN5 | YOR028C | 176.665065 | 0.13893552 | 0.12653132 | 1.09803267 | 0.27219022 |
| tH(GUG)K | tH(GUG)K | 4.30246952 | 0.30710116 | 0.27977465 | 1.09767327 | 0.27234718 |
| ELP6 | YMR312W | 293.356073 | -0.1181523 | 0.10766722 | -1.0973845 | 0.27247337 |
| SPS100 | YHR139C | 22.0859362 | 0.28748928 | 0.26228206 | 1.0961073 | 0.27303182 |
| MAY24_ | YPR153W | 95.6422007 | 0.18177041 | 0.16594449 | 1.09536876 | 0.27335511 |
| YMR209C | YMR209C | 305.353832 | 0.10982359 | 0.10030213 | 1.09492778 | 0.27354828 |
| YGR291C | YGR291C | 4.6362675 | 0.31175937 | 0.28491192 | 1.09423072 | 0.2738538 |
| COG6 | YNL041C | 1295.90322 | 0.07866202 | 0.0718842 | 1.09428803 | 0.27382868 |
| AME1 | YBR211C | 392.73646 | 0.11004767 | 0.10062034 | 1.09369201 | 0.27409008 |
| YKR032W | YKR032W | 1.20850624 | 0.21798285 | 0.19943803 | 1.09298541 | 0.27440021 |
| Xu_2009_CUTs90 | Xu_2009_CU | 18.2622846 | 0.3010644 | 0.27557422 | 1.09249842 | 0.27461408 |
| ESL1 | YIL151C | 1536.33384 | 0.06572758 | 0.06017566 | 1.09226187 | 0.27471801 |
| Xu_2009_SUTs17 | Xu_2009_SU | 58.2285583 | 0.22172252 | 0.20304223 | 1.09200204 | 0.2748322 |
| ATF1 | YOR377W | 526.848847 | 0.091604 | 0.083965 | 1.09097842 | 0.27528238 |
| GAT2 | YMR136W | 668.047046 | 0.09188888 | 0.08424261 | 1.09076479 | 0.27537639 |
| Xu_2009_SUTs20 | Xu_2009_SU | 60.8786131 | 0.20553293 | 0.18852813 | 1.09019767 | 0.27562608 |
| YAL031W-A | YAL031W-A | 6.10728606 | 0.31998833 | 0.29363354 | 1.08975402 | 0.27582151 |
| Xu_2009_SUTs75 | Xu_2009_SU | 72.7759965 | 0.21316942 | 0.19563563 | 1.08962472 | 0.27587849 |
| Xu_2009_CUTs89 | Xu_2009_CU | 36.8245902 | 0.27178812 | 0.2494632 | 1.08949183 | 0.27593706 |
| STE7 | YDL159W | 558.084606 | -0.0895887 | 0.08228327 | -1.0887833 | 0.27624944 |
| YRF1-2 | YER190W | 20.8276016 | 0.29299848 | 0.26912696 | 1.08869985 | 0.27628627 |
| SLM6 | YBR266C | 2.54294505 | 0.2727155 | 0.2506745 | 1.08792678 | 0.27662743 |
| DML1 | YMR211W | 335.845298 | 0.10998471 | 0.10109469 | 1.08793751 | 0.2766227 |
| PRP45 | YAL032C | 537.66075 | -0.0925842 | 0.08511925 | -1.0876996 | 0.27672774 |
| tP(AGG)N/SUF10 | tP(AGG)N | 7.05319753 | 0.32183909 | 0.29607471 | 1.08701986 | 0.27702803 |

| | | | | | | |
|---|---|---|---|---|---|---|
| CNE1 | YAL058W | 439.464791 | -0.096813 | 0.08920153 | -1.0853295 | 0.27777575 |
| SPG3 | YDR504C | 169.683466 | -0.1464748 | 0.13493675 | -1.0855069 | 0.27769719 |
| YJL077W-B | YJL077W-B | 57.6340275 | 0.22208445 | 0.2046093 | 1.08540742 | 0.27774125 |
| SSO1 | YPL232W | 1706.52646 | -0.0705925 | 0.06504155 | -1.0853454 | 0.27776869 |
| YLR437C-A | YLR437C-A | 12.0371271 | 0.31474059 | 0.29009598 | 1.0849533 | 0.27794234 |
| Xu_2009_CUTs49 | Xu_2009_CU | 3.88992682 | 0.29827636 | 0.27502249 | 1.08455264 | 0.27811984 |
| GTT2 | YLL060C | 128.273368 | 0.1615425 | 0.1490097 | 1.08410728 | 0.27831724 |
| Xu_2009_SUTs13 | Xu_2009_SU | 17.1377427 | 0.30778309 | 0.28402507 | 1.08364761 | 0.27852107 |
| Xu_2009_SUTs96 | Xu_2009_SU | 9.95865525 | 0.32027478 | 0.2955885 | 1.0835157 | 0.27857959 |
| Xu_2009_SUTs17 | Xu_2009_SU | 103.41411 | -0.1689148 | 0.15595146 | -1.0831241 | 0.27875335 |
| Xu_2009_SUTs28 | Xu_2009_SU | 8.87825102 | -0.3212428 | 0.29661148 | -1.0830425 | 0.27878956 |
| KCH1 | YJR054W | 769.834952 | 0.08752361 | 0.08084636 | 1.08259185 | 0.27898963 |
| VPS34 | YLR240W | 714.370399 | 0.08444823 | 0.07809306 | 1.08137947 | 0.27952835 |
| Xu_2009_SUTs61 | Xu_2009_SU | 82.5357277 | 0.19231979 | 0.1779849 | 1.08053987 | 0.27990184 |
| Xu_2009_SUTs28 | Xu_2009_SU | 16.6601905 | 0.30033533 | 0.27827634 | 1.07927009 | 0.28046734 |
| FUR1 | YHR128W | 1773.5407 | -0.0649155 | 0.06014973 | -1.0792312 | 0.28048469 |
| VBA4 | YDR119W | 964.437669 | -0.0752645 | 0.06977627 | -1.0786548 | 0.28074165 |
| GTF1 | YGR102C | 653.774302 | -0.0965451 | 0.08956922 | -1.0778826 | 0.28108613 |
| THP1 | YOL072W | 461.532606 | -0.1031538 | 0.09572867 | -1.0775639 | 0.28122842 |
| Xu_2009_SUTs75 | Xu_2009_SU | 56.5569017 | -0.2231739 | 0.20719181 | -1.0771368 | 0.28141914 |
| GDH2 | YDL215C | 1165.36836 | 0.09408535 | 0.08738509 | 1.07667503 | 0.28162547 |
| SPC110 | YDR356W | 646.572275 | 0.0832165 | 0.07730461 | 1.07647524 | 0.28171477 |
| SDC1 | YDR469W | 371.503439 | 0.10809557 | 0.10041302 | 1.07650948 | 0.28169946 |
| COG1 | YGL223C | 619.564595 | -0.0960103 | 0.08921283 | -1.076194 | 0.28184049 |
| MSH6 | YDR097C | 1173.18463 | 0.07432085 | 0.0691573 | 1.07466385 | 0.28252526 |
| Xu_2009_SUTs49 | Xu_2009_SU | 23.2586874 | 0.29278314 | 0.2724852 | 1.07449186 | 0.28260229 |
| FCF2 | YLR051C | 1029.73458 | -0.0834923 | 0.07770758 | -1.0744419 | 0.28262466 |
| VMA21 | YGR105W | 373.338945 | -0.1088678 | 0.10143731 | -1.0732524 | 0.2831579 |
| ATG20 | YDL113C | 725.455644 | 0.0895109 | 0.08351417 | 1.07180496 | 0.28380764 |
| SMB1 | YER029C | 525.391342 | 0.09482329 | 0.08846841 | 1.07183224 | 0.28379539 |
| VHS1 | YDR247W | 299.311152 | -0.1216508 | 0.11353233 | -1.0715084 | 0.28394091 |
| GDH3 | YAL062W | 170.656881 | -0.1436912 | 0.13419504 | -1.0707637 | 0.28427571 |
| YDR406W-A | YDR406W-A | 1.02262495 | 0.18553298 | 0.17328172 | 1.07070141 | 0.28430371 |
| YNL276C | YNL276C | 63.2262244 | 0.20531409 | 0.1917782 | 1.070581 | 0.28435787 |
| Xu_2009_CUTs26 | Xu_2009_CU | 10.8225091 | 0.3116844 | 0.29123885 | 1.07020201 | 0.28452839 |
| DFG10 | YIL049W | 453.551106 | 0.0956293 | 0.08949647 | 1.06852591 | 0.28528335 |
| Xu_2009_CUTs46 | Xu_2009_CU | 21.3591882 | -0.286495 | 0.2681785 | -1.0682996 | 0.2853854 |
| SRF1 | YDL133W | 725.120846 | 0.08870875 | 0.0831183 | 1.06725895 | 0.28585492 |
| DLT1 | YMR126C | 175.454739 | -0.1404924 | 0.13163275 | -1.0673058 | 0.28583379 |
| EKI1 | YDR147W | 557.935579 | -0.0885552 | 0.08299299 | -1.0670208 | 0.28596245 |
| Xu_2009_SUTs37 | Xu_2009_SU | 70.0641615 | 0.19872348 | 0.18630104 | 1.0666794 | 0.28611663 |

| | | | | | | |
|---|---|---|---|---|---|---|
| DMA1 | YHR115C | 1146.67549 | 0.06753558 | 0.06330773 | 1.06678238 | 0.28607012 |
| NDJ1 | YOL104C | 199.43374 | 0.13768696 | 0.12907261 | 1.06674041 | 0.28608907 |
| Xu_2009_SUTs16( | Xu_2009_SU | 8.34065335 | 0.31551324 | 0.29606817 | 1.06567767 | 0.28656937 |
| YHR028W-A | YHR028W-A | 9.31127801 | 0.31472516 | 0.29554203 | 1.0649083 | 0.28691743 |
| ELA1 | YNL230C | 364.877445 | 0.10227217 | 0.09608537 | 1.06438857 | 0.28715271 |
| SWD3 | YBR175W | 415.477232 | -0.0977703 | 0.09186909 | -1.0642346 | 0.28722243 |
| YOR309C | YOR309C | 7.93114945 | -0.3151718 | 0.29626146 | -1.0638298 | 0.28740582 |
| GIP4 | YAL031C | 559.820638 | -0.0980141 | 0.09224672 | -1.0625209 | 0.28799927 |
| RIM4 | YHL024W | 251.993383 | 0.1260436 | 0.11864054 | 1.06239912 | 0.28805454 |
| BRE2 | YLR015W | 470.482454 | 0.1004532 | 0.0945851 | 1.06204036 | 0.28821736 |
| RRP1 | YDR087C | 895.167699 | 0.07818971 | 0.07375384 | 1.06014424 | 0.28907898 |
| YKR073C | YKR073C | 7.68407874 | -0.3145131 | 0.29664547 | -1.0602323 | 0.28903895 |
| YHL049C | YHL049C | 7.62949948 | 0.31399208 | 0.2964622 | 1.05913025 | 0.28954046 |
| BTN2 | YGR142W | 525.802218 | 0.20149989 | 0.19029402 | 1.05888714 | 0.28965118 |
| YDR274C | YDR274C | 3.35063053 | 0.28770961 | 0.27176437 | 1.05867304 | 0.28974871 |
| Xu_2009_SUTs34( | Xu_2009_SU | 12.1833142 | 0.30636541 | 0.28946024 | 1.0584024 | 0.28987203 |
| TAF5 | YBR198C | 1219.32562 | -0.0689494 | 0.06518386 | -1.0577687 | 0.2901609 |
| RAD59 | YDL059C | 143.994193 | 0.14544584 | 0.13761226 | 1.05692507 | 0.29054578 |
| DIA3 | YDL024C | 85.0832101 | 0.19518565 | 0.18470853 | 1.05672244 | 0.29063828 |
| Xu_2009_SUTs27! | Xu_2009_SU | 102.536495 | 0.1711504 | 0.16200802 | 1.05643163 | 0.29077106 |
| tM(CAU)O2/EMT2 | tM(CAU)O2 | 22.0361598 | 0.28392996 | 0.2689886 | 1.05554646 | 0.29117547 |
| YLR347W-A | YLR347W-A | 6.06928918 | 0.30797615 | 0.29182159 | 1.05535764 | 0.29126179 |
| Xu_2009_CUTs69 | Xu_2009_CU | 5.64774064 | 0.30919395 | 0.2931205 | 1.05483563 | 0.29150051 |
| tA(AGC)L | tA(AGC)L | 4.5857876 | -0.2956236 | 0.28057014 | -1.053653 | 0.2920418 |
| CNB1 | YKL190W | 982.663399 | -0.0777124 | 0.07382418 | -1.0526687 | 0.29249288 |
| UBP12 | YJL197W | 1232.19104 | 0.07107009 | 0.06756017 | 1.05195256 | 0.29282132 |
| TEN1 | YLR010C | 66.1981191 | -0.1948001 | 0.18516618 | -1.0520283 | 0.29278658 |
| YJL032W | YJL032W | 2.05330704 | -0.2415849 | 0.22981727 | -1.0512043 | 0.29316475 |
| TPK1 | YJL164C | 684.214964 | -0.1700353 | 0.16179517 | -1.0509296 | 0.29329093 |
| MER1 | YNL210W | 7.54632625 | -0.3111574 | 0.29648158 | -1.0494999 | 0.29394811 |
| PLP2 | YOR281C | 825.779659 | 0.07594964 | 0.07237788 | 1.04934884 | 0.2940176 |
| ANY1 | YMR010W | 996.031489 | -0.0846534 | 0.08068486 | -1.0491853 | 0.29409284 |
| IZH4 | YOL101C | 909.194013 | 0.0958464 | 0.09136489 | 1.04905072 | 0.29415477 |
| Xu_2009_SUTs49 | Xu_2009_SU | 20.7082343 | 0.28322721 | 0.27015257 | 1.04839725 | 0.29445562 |
| GAL3 | YDR009W | 90.1782711 | 0.17265925 | 0.16467766 | 1.04846797 | 0.29442305 |
| YMR052C-A | YMR052C-A | 3.0374572 | 0.28144875 | 0.26846092 | 1.04837884 | 0.2944641 |
| YMR141C | YMR141C | 7.7082548 | -0.3109092 | 0.29658164 | -1.048309 | 0.29449624 |
| Xu_2009_SUTs60( | Xu_2009_SU | 19.0131659 | -0.2841144 | 0.27119926 | -1.0476224 | 0.29481262 |
| SPC72 | YAL047C | 334.515019 | -0.1076109 | 0.10272599 | -1.047553 | 0.29484458 |
| NBP2 | YDR162C | 403.078877 | 0.10402868 | 0.09933165 | 1.04728633 | 0.29496754 |
| GIM5 | YML094W | 464.687196 | -0.0935228 | 0.089296 | -1.0473343 | 0.29494544 |

| | | | | | | |
|---|---|---|---|---|---|---|
| Xu_2009_SUTs19 | Xu_2009_SU | 118.280728 | 0.15245186 | 0.14564778 | 1.04671595 | 0.2952306 |
| YPR096C | YPR096C | 55.1093952 | 0.2118092 | 0.20245836 | 1.04618646 | 0.29547495 |
| APC2 | YLR127C | 460.207741 | 0.0917642 | 0.08774518 | 1.04580331 | 0.29565185 |
| SHO1 | YER118C | 804.934727 | -0.0806603 | 0.07715809 | -1.0453897 | 0.29584289 |
| Xu_2009_SUTs77 | Xu_2009_SU | 112.194725 | -0.163098 | 0.15605154 | -1.0451546 | 0.29595153 |
| Xu_2009_CUTs80 | Xu_2009_CU | 9.75075825 | 0.30821128 | 0.29507721 | 1.04451061 | 0.29624921 |
| MRPS17 | YMR188C | 761.201915 | -0.0847753 | 0.08117031 | -1.0444132 | 0.29629428 |
| Xu_2009_SUTs28 | Xu_2009_SU | 44.6524323 | 0.22852708 | 0.21905036 | 1.04326276 | 0.29682662 |
| Xu_2009_CUTs62 | Xu_2009_CU | 1.27547816 | 0.21127011 | 0.20256453 | 1.04297682 | 0.29695903 |
| Xu_2009_CUTs70 | Xu_2009_CU | 1.39266168 | 0.21713874 | 0.20831137 | 1.04237581 | 0.29723748 |
| Xu_2009_SUTs70 | Xu_2009_SU | 28.8648026 | -0.253529 | 0.2432293 | -1.0423456 | 0.2972515 |
| EFG1 | YGR271C-A | 485.702075 | -0.1066793 | 0.10231743 | -1.0426304 | 0.2971195 |
| SWC7 | YLR385C | 83.8680015 | 0.18660664 | 0.17902546 | 1.04234693 | 0.29725086 |
| YIL086C | YIL086C | 11.9138249 | -0.304616 | 0.29232162 | -1.0420577 | 0.29738491 |
| YOL083C-A | YOL083C-A | 4.07534802 | 0.28258389 | 0.27122617 | 1.04187543 | 0.29746944 |
| YBL086C | YBL086C | 410.280039 | 0.11263631 | 0.10815413 | 1.04144253 | 0.29767021 |
| CDC31 | YOR257W | 448.940773 | 0.10161433 | 0.0976093 | 1.04103128 | 0.29786103 |
| YNR042W | YNR042W | 11.8887546 | 0.30344209 | 0.29188577 | 1.03959194 | 0.29852952 |
| tS(AGA)D3 | tS(AGA)D3 | 3.66550211 | -0.2847095 | 0.27398732 | -1.039134 | 0.29874243 |
| SLP1 | YOR154W | 776.38648 | 0.07541517 | 0.07261803 | 1.03851841 | 0.29902877 |
| MDL1 | YLR188W | 1091.96118 | -0.0706579 | 0.0680746 | -1.0379483 | 0.29929413 |
| SEC24 | YIL109C | 3833.39811 | -0.0524249 | 0.05052151 | -1.0376744 | 0.29942165 |
| OLE1 | YGL055W | 16888.2767 | -0.1401073 | 0.13516085 | -1.0365968 | 0.2999238 |
| EFM3 | YJR129C | 128.216583 | -0.1475844 | 0.14242435 | -1.0362301 | 0.30009483 |
| MDM1 | YML104C | 680.277295 | 0.09135065 | 0.08817243 | 1.03604552 | 0.30018091 |
| VTA1 | YLR181C | 384.992721 | -0.1010337 | 0.09755014 | -1.0357101 | 0.30033741 |
| tV(UAC)D | tV(UAC)D | 154.51105 | 0.14440267 | 0.13945997 | 1.03544177 | 0.30046265 |
| YHR078W | YHR078W | 725.544641 | 0.08075939 | 0.07800305 | 1.03533628 | 0.30051189 |
| tD(GUC)N | tD(GUC)N | 27.892032 | 0.26130115 | 0.25246479 | 1.03500037 | 0.30066874 |
| FMP16 | YDR070C | 766.853595 | 0.08962821 | 0.08662637 | 1.03465281 | 0.30083108 |
| MSS11 | YMR164C | 572.792167 | 0.09591078 | 0.0927021 | 1.0346128 | 0.30084977 |
| YLR358C | YLR358C | 3.13431893 | 0.27755389 | 0.26872538 | 1.03285328 | 0.30167256 |
| RCL1 | YOL010W | 1205.77585 | -0.0730124 | 0.07068867 | -1.0328732 | 0.30166325 |
| APD1 | YBR151W | 1574.88445 | -0.0730938 | 0.0708138 | -1.0321964 | 0.3019801 |
| GLE1 | YDL207W | 1072.60121 | -0.0713447 | 0.06914979 | -1.0317408 | 0.30219355 |
| CDC8 | YJR057W | 450.290068 | -0.1001425 | 0.09724823 | -1.0297616 | 0.30312195 |
| THI73 | YLR004C | 55.7411326 | 0.20734532 | 0.20181071 | 1.02742477 | 0.30422049 |
| YJL215C | YJL215C | 39.0823196 | 0.23025378 | 0.22427911 | 1.02663946 | 0.30459027 |
| Xu_2009_SUTs12 | Xu_2009_SU | 86.1605646 | 0.17756638 | 0.1731437 | 1.02554341 | 0.30510685 |
| TRE2 | YOR256C | 1297.54401 | 0.0634436 | 0.06186064 | 1.02558912 | 0.3050853 |
| YLL044W | YLL044W | 1.33332537 | -0.2078372 | 0.20280874 | -1.0247943 | 0.30546026 |

| | | | | | | |
|---|---|---|---|---|---|---|
| DUN1 | YDL101C | 387.427213 | 0.10546903 | 0.10295721 | 1.02439671 | 0.30564794 |
| Xu_2009_CUTs47 | Xu_2009_CU | 2.95396321 | 0.26882366 | 0.26246289 | 1.02423493 | 0.30572433 |
| YHR045W | YHR045W | 384.160485 | -0.1025091 | 0.10014921 | -1.0235636 | 0.30604142 |
| Xu_2009_CUTs10 | Xu_2009_CU | 13.3970042 | 0.29283508 | 0.2862551 | 1.02298643 | 0.30631426 |
| RCN1 | YKL159C | 240.667086 | -0.1251886 | 0.12249937 | -1.0219529 | 0.3068032 |
| MIN4 | YML007C-A | 8.3613811 | 0.30258967 | 0.29610638 | 1.02189515 | 0.30683053 |
| Xu_2009_SUTs10 | Xu_2009_SU | 121.47205 | 0.15341588 | 0.15020219 | 1.02139575 | 0.30706698 |
| PPH22 | YDL188C | 1031.96097 | -0.0687437 | 0.06730309 | -1.0214051 | 0.30706255 |
| Xu_2009_SUTs37 | Xu_2009_SU | 17.2996691 | -0.2917659 | 0.28569442 | -1.0212516 | 0.30713525 |
| tG(GCC)P1 | tG(GCC)P1 | 50.0814202 | -0.2275501 | 0.2228537 | -1.021074 | 0.30721937 |
| FRE2 | YKL220C | 122.157015 | 0.15280621 | 0.14966857 | 1.02096392 | 0.30727153 |
| Xu_2009_SUTs45 | Xu_2009_SU | 10.7095219 | -0.2990237 | 0.29291688 | -1.0208481 | 0.30732639 |
| GPP2 | YER062C | 4502.48038 | -0.0682334 | 0.06691176 | -1.0197516 | 0.30784627 |
| RPL8A | YHL033C | 5284.87466 | -0.0629336 | 0.06173725 | -1.0193787 | 0.30802322 |
| MCM16 | YPR046W | 183.093234 | 0.12456515 | 0.12226803 | 1.01878756 | 0.30830383 |
| Xu_2009_CUTs51 | Xu_2009_CU | 24.9009317 | 0.25774655 | 0.25309323 | 1.01838581 | 0.30849464 |
| YLR296W | YLR296W | 5.0548808 | 0.29465124 | 0.28939403 | 1.01816624 | 0.30859896 |
| YOR011W-A | YOR011W-A | 3.24192249 | 0.27392906 | 0.26925212 | 1.01737012 | 0.30897739 |
| YGR127W | YGR127W | 429.771657 | -0.0925587 | 0.09104759 | -1.0165968 | 0.30934528 |
| Xu_2009_SUTs93 | Xu_2009_SU | 81.3154494 | 0.17626136 | 0.17343909 | 1.01627242 | 0.30949968 |
| SLZ1 | YNL196C | 16.3679998 | -0.2821501 | 0.27759745 | -1.0164003 | 0.30943879 |
| YPR159C-A | YPR159C-A | 2.66055735 | -0.2507876 | 0.24675774 | -1.0163312 | 0.30947172 |
| VPS35 | YJL154C | 1993.17235 | -0.0599378 | 0.05904917 | -1.0150482 | 0.31008283 |
| CTF18 | YMR078C | 508.892304 | -0.0939017 | 0.09252056 | -1.0149283 | 0.31014002 |
| DLS1 | YJL065C | 526.491261 | -0.0882768 | 0.08700036 | -1.0146722 | 0.31026212 |
| Xu_2009_CUTs46 | Xu_2009_CU | 2.15614349 | 0.24359242 | 0.24024707 | 1.01392461 | 0.31061873 |
| Xu_2009_CUTs26 | Xu_2009_CU | 15.4795803 | 0.28711397 | 0.28343174 | 1.01299157 | 0.31106419 |
| VHR2 | YER064C | 1461.14572 | 0.06619548 | 0.06537245 | 1.01258977 | 0.31125615 |
| VAN1 | YML115C | 1523.79414 | -0.0607187 | 0.06000189 | -1.0119465 | 0.31156364 |
| Xu_2009_CUTs30 | Xu_2009_CU | 16.8945794 | -0.2837588 | 0.28045327 | -1.0117863 | 0.31164023 |
| SRB4 | YER022W | 608.60116 | -0.0846926 | 0.08376144 | -1.0111163 | 0.31196079 |
| SLS1 | YLR139C | 529.738071 | -0.0974518 | 0.09644038 | -1.0104873 | 0.31226187 |
| Xu_2009_CUTs40 | Xu_2009_CU | 17.4561275 | 0.27749867 | 0.27480646 | 1.00979677 | 0.31259267 |
| YSF3 | YNL138W-A | 93.6912318 | -0.1848744 | 0.18334515 | -1.008341 | 0.31329081 |
| Xu_2009_SUTs64 | Xu_2009_SU | 30.9684573 | -0.2480647 | 0.24629834 | -1.0071717 | 0.31385226 |
| KAP122 | YGL016W | 1287.02751 | 0.06736314 | 0.06687232 | 1.00733967 | 0.31377157 |
| DSS1 | YMR287C | 787.262013 | -0.0773426 | 0.07678448 | -1.0072684 | 0.31380581 |
| YEL014C | YEL014C | 2.13903409 | 0.23992647 | 0.23832304 | 1.00672797 | 0.31406552 |
| SWA2 | YDR320C | 833.98295 | -0.0782609 | 0.07777042 | -1.0063063 | 0.31426826 |
| YPS6 | YIR039C | 178.258707 | 0.13396021 | 0.13313902 | 1.00616786 | 0.31433483 |
| Xu_2009_SUTs29 | Xu_2009_SU | 45.4910737 | 0.21020934 | 0.20904096 | 1.0055892 | 0.31461322 |

| | | | | | | |
|---|---|---|---|---|---|---|
| Xu_2009_SUTs55 | Xu_2009_SU | 68.4298414 | 0.19453422 | 0.19350837 | 1.0053013 | 0.31475179 |
| tY(GUA)F1/SUP11 | tY(GUA)F1 | 3.37259102 | 0.26959147 | 0.26827332 | 1.00491344 | 0.31493853 |
| Xu_2009_SUTs75 | Xu_2009_SU | 54.012419 | -0.205327 | 0.20442179 | -1.0044283 | 0.31517224 |
| YNL205C | YNL205C | 4.26984208 | -0.2803384 | 0.27913654 | -1.0043055 | 0.31523138 |
| Xu_2009_CUTs25 | Xu_2009_CU | 8.90593671 | 0.29748592 | 0.29637356 | 1.00375325 | 0.31549756 |
| KIN4 | YOR233W | 1136.64312 | 0.07163909 | 0.07137284 | 1.00373034 | 0.31550861 |
| YBR032W | YBR032W | 13.1458239 | -0.2884934 | 0.28761484 | -1.0030548 | 0.31583441 |
| tQ(UUG)L | tQ(UUG)L | 0.77158658 | -0.1530992 | 0.15277652 | -1.0021124 | 0.31628929 |
| BUD2 | YKL092C | 859.265732 | -0.0769235 | 0.07675932 | -1.0021392 | 0.31627637 |
| MAE1 | YKL029C | 1750.39085 | -0.0701922 | 0.0701348 | -1.0008189 | 0.31691439 |
| MTQ1 | YNL063W | 314.548601 | -0.1080117 | 0.10795694 | -1.0005069 | 0.31706528 |
| NOP15 | YNL110C | 1468.98089 | -0.0769603 | 0.07692699 | -1.0004331 | 0.31710096 |
| LEU5 | YHR002W | 187.448937 | -0.1375007 | 0.13760649 | -0.9992312 | 0.31768272 |
| Xu_2009_CUTs17 | Xu_2009_CU | 10.7708353 | -0.2959153 | 0.29638243 | -0.9984241 | 0.31807378 |
| Xu_2009_CUTs12 | Xu_2009_CU | 10.532854 | -0.2935381 | 0.29406339 | -0.9982138 | 0.3181757 |
| Xu_2009_SUTs46 | Xu_2009_SU | 104.744683 | 0.17088588 | 0.17127041 | 0.99775486 | 0.31839825 |
| MMS4 | YBR098W | 232.937348 | -0.1177684 | 0.11816324 | -0.9966588 | 0.31893016 |
| LUG1 | YLR352W | 543.07129 | -0.0864669 | 0.08677878 | -0.9964056 | 0.3190531 |
| ATP6 | Q0085 | 402.212841 | -0.2932908 | 0.29463183 | -0.9954484 | 0.31951822 |
| NNK1 | YKL171W | 577.088679 | 0.08310283 | 0.08348278 | 0.99544876 | 0.31951805 |
| Xu_2009_CUTs40 | Xu_2009_CU | 24.0554958 | 0.25900889 | 0.26028525 | 0.99509632 | 0.31968942 |
| MTG1 | YMR097C | 346.826874 | -0.1019317 | 0.10249298 | -0.9945233 | 0.31996817 |
| Xu_2009_CUTs70 | Xu_2009_CU | 13.2151197 | 0.28888447 | 0.29074527 | 0.9935999 | 0.32041769 |
| Xu_2009_CUTs73 | Xu_2009_CU | 8.11975446 | 0.29430733 | 0.29645055 | 0.9927704 | 0.32082186 |
| SCC4 | YER147C | 428.529545 | 0.0890635 | 0.08971678 | 0.99271847 | 0.32084717 |
| SPN1 | YPR133C | 1560.265 | 0.06330388 | 0.06376645 | 0.99274596 | 0.32083377 |
| Xu_2009_SUTs41 | Xu_2009_SU | 20.9411749 | -0.2630521 | 0.26512381 | -0.992186 | 0.32110679 |
| YLR053C | YLR053C | 17.6640878 | -0.2711701 | 0.27333865 | -0.9920665 | 0.3211651 |
| NOT5 | YPR072W | 1176.19439 | 0.06528263 | 0.06582169 | 0.99181036 | 0.32129004 |
| Xu_2009_SUTs75 | Xu_2009_SU | 9.04945706 | 0.29279725 | 0.29560876 | 0.99048908 | 0.32193513 |
| PAU23 | YLR037C | 53.8772181 | 0.19684114 | 0.19878204 | 0.99023602 | 0.32205877 |
| GND2 | YGR256W | 862.276765 | -0.0796836 | 0.08053055 | -0.989483 | 0.32242688 |
| PPZ2 | YDR436W | 394.543548 | -0.0936169 | 0.0946529 | -0.9890546 | 0.32263641 |
| Xu_2009_CUTs90 | Xu_2009_CU | 10.5840683 | -0.2890561 | 0.29294574 | -0.9867223 | 0.32377881 |
| YPR016W-A | YPR016W-A | 1.0540545 | -0.1913819 | 0.19393784 | -0.9868211 | 0.32373036 |
| YLL006W-A | YLL006W-A | 2.14793641 | -0.2242186 | 0.2273136 | -0.9863846 | 0.32394444 |
| YJR157W | YJR157W | 3.44987732 | 0.26146283 | 0.2652761 | 0.98562526 | 0.32431704 |
| Xu_2009_SUTs29 | Xu_2009_SU | 47.7684952 | 0.21208759 | 0.21521176 | 0.98548327 | 0.32438675 |
| CWC22 | YGR278W | 395.400431 | 0.10527532 | 0.10683601 | 0.98539171 | 0.3244317 |
| SNR61 | snR61 | 15.9678778 | -0.279083 | 0.28328397 | -0.9851705 | 0.32454035 |
| Xu_2009_SUTs24 | Xu_2009_SU | 56.8003689 | 0.20310285 | 0.20615462 | 0.98519666 | 0.32452748 |

| | | | | | | |
|---|---|---|---|---|---|---|
| PES4 | YFR023W | 161.206167 | 0.12852886 | 0.13048811 | 0.98498526 | 0.32463131 |
| YPR195C | YPR195C | 51.8396943 | 0.19804636 | 0.20113162 | 0.9846605 | 0.32479086 |
| Xu_2009_CUTs20 | Xu_2009_CU | 0.83432074 | -0.1673671 | 0.1699988 | -0.9845191 | 0.32486034 |
| CYC8 | YBR112C | 2646.83692 | -0.0535556 | 0.05441106 | -0.9842782 | 0.32497874 |
| YEL023C | YEL023C | 302.542964 | 0.10039978 | 0.10201869 | 0.98413122 | 0.325051 |
| YFL041W-A | YFL041W-A | 29.2413888 | -0.2430081 | 0.2469447 | -0.9840587 | 0.32508663 |
| MET2 | YNL277W | 96.0328065 | 0.15943021 | 0.16198912 | 0.98420325 | 0.32501559 |
| YIG1 | YPL201C | 129.938591 | -0.1476207 | 0.15002328 | -0.9839855 | 0.32512262 |
| APS1 | YLR170C | 273.921496 | -0.1052435 | 0.10697562 | -0.9838081 | 0.32520987 |
| YJR012C | YJR012C | 166.97533 | -0.129022 | 0.13124924 | -0.9830304 | 0.32559247 |
| Xu_2009_CUTs81 | Xu_2009_CU | 31.4360232 | 0.24208928 | 0.24638243 | 0.98257524 | 0.32581654 |
| Xu_2009_CUTs18 | Xu_2009_CU | 13.1362324 | 0.284567 | 0.2898172 | 0.98188444 | 0.32615678 |
| tA(AGC)J | tA(AGC)J | 1.40837538 | -0.1922954 | 0.19589049 | -0.9816472 | 0.32627368 |
| Xu_2009_SUTs30 | Xu_2009_SU | 51.7044988 | -0.2031545 | 0.20702305 | -0.9813132 | 0.32643831 |
| Xu_2009_CUTs83 | Xu_2009_CU | 5.32971625 | 0.28306037 | 0.28852099 | 0.98107378 | 0.32655636 |
| Xu_2009_CUTs93 | Xu_2009_CU | 12.9561396 | 0.28156906 | 0.28712254 | 0.98065817 | 0.32676134 |
| ATR2 | YMR279C | 7.47347753 | -0.2899855 | 0.29626389 | -0.978808 | 0.32767484 |
| PXL1 | YKR090W | 853.521061 | -0.0752249 | 0.07687586 | -0.978524 | 0.32781521 |
| YLR232W | YLR232W | 1.55976891 | 0.21273378 | 0.21742909 | 0.9784053 | 0.32787391 |
| Xu_2009_CUTs16 | Xu_2009_CU | 9.50135426 | 0.28962976 | 0.29609913 | 0.97815133 | 0.32799948 |
| YHR086W-A | YHR086W-A | 14.2081095 | 0.28207503 | 0.28856606 | 0.97750593 | 0.32831874 |
| YPR011C | YPR011C | 282.72026 | -0.1076654 | 0.11016159 | -0.9773411 | 0.32840031 |
| Xu_2009_SUTs80 | Xu_2009_SU | 16.602277 | -0.2751496 | 0.28225422 | -0.9748291 | 0.32964504 |
| ERR2 | YPL281C | 15.5865092 | 0.27646054 | 0.28365647 | 0.97463155 | 0.32974306 |
| Xu_2009_CUTs24 | Xu_2009_CU | 19.2918637 | 0.27185694 | 0.27919631 | 0.97371251 | 0.33019931 |
| YKL131W | YKL131W | 0.90568798 | -0.1707534 | 0.17549499 | -0.9729815 | 0.3305625 |
| Xu_2009_CUTs88 | Xu_2009_CU | 27.6205541 | 0.2474498 | 0.25437983 | 0.97275716 | 0.33067401 |
| BER1 | YLR412W | 581.070547 | 0.0779666 | 0.08018502 | 0.97233369 | 0.33088457 |
| YHR007C-A | YHR007C-A | 55.6320973 | 0.19855353 | 0.20423216 | 0.97219519 | 0.33095345 |
| BUD30 | YDL151C | 1.24858906 | 0.19604015 | 0.20183954 | 0.97126734 | 0.33141517 |
| Xu_2009_SUTs57 | Xu_2009_SU | 1.05744694 | -0.1857954 | 0.19140827 | -0.9706759 | 0.33170971 |
| YBR219C | YBR219C | 11.5922845 | -0.2816729 | 0.29014693 | -0.9707941 | 0.33165082 |
| ABD1 | YBR236C | 737.258419 | -0.0758412 | 0.078135 | -0.9706428 | 0.33172618 |
| Xu_2009_CUTs92 | Xu_2009_CU | 2.73632187 | 0.24825163 | 0.25593476 | 0.96998016 | 0.33205638 |
| MRT4 | YKL009W | 1907.7321 | 0.0760615 | 0.07844156 | 0.96965815 | 0.33221692 |
| Xu_2009_SUTs28 | Xu_2009_SU | 44.5644216 | -0.2172788 | 0.22414477 | -0.969368 | 0.33236162 |
| YNL171C | YNL171C | 3.73290272 | -0.2685574 | 0.27708243 | -0.969233 | 0.33242896 |
| LUC7 | YDL087C | 226.532207 | 0.11026295 | 0.11380993 | 0.96883421 | 0.33262792 |
| YJR038C | YJR038C | 4.20743895 | 0.27228949 | 0.28101926 | 0.96893535 | 0.33257745 |
| IRC23 | YOR044W | 569.449626 | -0.0850521 | 0.08790355 | -0.9675611 | 0.33326359 |
| Xu_2009_SUTs10 | Xu_2009_SU | 48.570817 | 0.20682416 | 0.21385244 | 0.96713492 | 0.33347659 |

| | | | | | | |
|---|---|---|---|---|---|---|
| SPE4 | YLR146C | 218.115382 | 0.12049421 | 0.12470428 | 0.96623963 | 0.33392429 |
| MTQ2 | YDR140W | 345.343754 | -0.0927536 | 0.09601571 | -0.9660252 | 0.3340316 |
| Xu_2009_CUTs58 | Xu_2009_CU | 17.2576178 | 0.27908045 | 0.28900948 | 0.96564462 | 0.33422204 |
| YML009W-B | YML009W-B | 4.02871986 | -0.2720273 | 0.2818081 | -0.9652925 | 0.33439831 |
| Xu_2009_CUTs61 | Xu_2009_CU | 8.48542082 | 0.28585204 | 0.29628576 | 0.96478493 | 0.33465254 |
| Xu_2009_CUTs22 | Xu_2009_CU | 1.7643619 | -0.2090518 | 0.21712336 | -0.9628249 | 0.33563538 |
| Xu_2009_SUTs48 | Xu_2009_SU | 67.6461989 | 0.18018386 | 0.18715209 | 0.96276703 | 0.33566445 |
| UBX3 | YDL091C | 428.777785 | -0.0936925 | 0.09739138 | -0.9620206 | 0.33603928 |
| YGL069C | YGL069C | 36.4161201 | 0.23929246 | 0.24884495 | 0.96161267 | 0.3362442 |
| MET28 | YIR017C | 107.711822 | -0.1483633 | 0.15442775 | -0.9607295 | 0.33668819 |
| MIH1 | YMR036C | 406.462378 | 0.10089602 | 0.10505597 | 0.96040257 | 0.33685264 |
| RRD2 | YPL152W | 403.255809 | 0.09925425 | 0.10347252 | 0.95923289 | 0.33744143 |
| tI(AAU)P2 | tI(AAU)P2 | 1.40073354 | -0.2046023 | 0.21348867 | -0.9583756 | 0.33787341 |
| Xu_2009_SUTs38 | Xu_2009_SU | 39.4401173 | 0.21983754 | 0.22938813 | 0.95836493 | 0.33787878 |
| YCL046W | YCL046W | 1.00311382 | -0.1692692 | 0.17658854 | -0.9585514 | 0.3377848 |
| YNL170W | YNL170W | 21.685795 | 0.25408136 | 0.2650901 | 0.9584717 | 0.33782496 |
| AOS1 | YPR180W | 346.79459 | -0.0982606 | 0.10253756 | -0.9582885 | 0.33791729 |
| KNH1 | YDL049C | 279.055568 | -0.1093722 | 0.11442214 | -0.9558655 | 0.33914018 |
| Xu_2009_CUTs82 | Xu_2009_CU | 2.95413173 | 0.25340232 | 0.26515206 | 0.95568677 | 0.33923051 |
| YNL034W | YNL034W | 7.76630767 | -0.2826122 | 0.29574888 | -0.9555815 | 0.3392837 |
| YAL037W | YAL037W | 129.078136 | 0.15429542 | 0.16154693 | 0.95511202 | 0.33952105 |
| Xu_2009_SUTs76 | Xu_2009_SU | 65.2750326 | 0.18632766 | 0.19528399 | 0.95413686 | 0.34001437 |
| YCR087W | YCR087W | 15.5175612 | -0.2671611 | 0.2801587 | -0.9536062 | 0.34028301 |
| YNL303W | YNL303W | 0.96469831 | 0.17577059 | 0.18439872 | 0.95320941 | 0.34048398 |
| SNR76 | snR76 | 940.743388 | 0.08631196 | 0.09057252 | 0.95295967 | 0.34061051 |
| YBR109W-A | YBR109W-A | 24.8749259 | -0.2533376 | 0.26585689 | -0.9529098 | 0.34063577 |
| Xu_2009_CUTs51 | Xu_2009_CU | 7.15178037 | 0.28165694 | 0.29579847 | 0.952192 | 0.34099962 |
| KEL1 | YHR158C | 1598.94946 | -0.0589575 | 0.06191403 | -0.9522483 | 0.34097108 |
| Xu_2009_SUTs13 | Xu_2009_SU | 123.171322 | -0.1448814 | 0.15227073 | -0.9514723 | 0.34136467 |
| SIP3 | YNL257C | 925.05756 | -0.0667855 | 0.07021777 | -0.95112 | 0.34154348 |
| YPR014C | YPR014C | 4.07588335 | 0.2657825 | 0.2799334 | 0.94944906 | 0.34239227 |
| YDR526C | YDR526C | 14.5812236 | 0.26676422 | 0.28120386 | 0.94865063 | 0.34279833 |
| KKQ8 | YKL168C | 600.098975 | -0.0902016 | 0.0951039 | -0.9484537 | 0.34289852 |
| Xu_2009_SUTs42 | Xu_2009_SU | 71.1844217 | 0.18479508 | 0.19499697 | 0.94768182 | 0.34329146 |
| UGX2 | YDL169C | 207.594842 | -0.1232948 | 0.13012858 | -0.9474845 | 0.34339195 |
| PWP1 | YLR196W | 2848.46444 | 0.0709575 | 0.07491952 | 0.94711637 | 0.34357948 |
| GLO4 | YOR040W | 214.565001 | -0.114446 | 0.12084572 | -0.947042 | 0.3436174 |
| YNL050C | YNL050C | 579.510069 | 0.07812133 | 0.08252728 | 0.94661218 | 0.34383643 |
| YOR329W-A | YOR329W-A | 3.42289716 | 0.25804443 | 0.27275831 | 0.94605525 | 0.34412041 |
| YPR136C | YPR136C | 1.13837821 | -0.1894876 | 0.20027106 | -0.9461556 | 0.34406922 |
| GZF3 | YJL110C | 980.235872 | 0.06464325 | 0.06838595 | 0.94527102 | 0.34452053 |

| | | | | | | |
|---|---|---|---|---|---|---|
| SPT23 | YKL020C | 1607.9261 | 0.05735243 | 0.0606743 | 0.94525082 | 0.34453084 |
| MSN5 | YDR335W | 1587.26032 | -0.060135 | 0.06363237 | -0.9450386 | 0.34463919 |
| MRE11 | YMR224C | 392.34364 | -0.0882114 | 0.09336199 | -0.9448317 | 0.34474481 |
| TDA7 | YNL176C | 965.861508 | 0.06609391 | 0.06998514 | 0.94439921 | 0.34496568 |
| YPI1 | YFR003C | 518.212119 | -0.0865026 | 0.09162751 | -0.9440684 | 0.34513468 |
| Xu_2009_SUTs22 | Xu_2009_SU | 17.9416006 | 0.25930332 | 0.27474016 | 0.94381296 | 0.34526523 |
| VID28 | YIL017C | 549.382791 | 0.08928375 | 0.09460238 | 0.94377908 | 0.34528255 |
| YLR149C-A | YLR149C-A | 2.33277907 | 0.23820912 | 0.25262102 | 0.94295052 | 0.34570621 |
| YSP3 | YOR003W | 204.54563 | -0.1136782 | 0.12061855 | -0.9424602 | 0.34595709 |
| Xu_2009_CUTs77 | Xu_2009_CU | 19.3262306 | 0.24985707 | 0.26544346 | 0.9412817 | 0.34656052 |
| FYV12 | YOR183W | 12.8766274 | 0.27102299 | 0.28849168 | 0.93944818 | 0.34750068 |
| YJL043W | YJL043W | 26.6362635 | -0.2448813 | 0.2609409 | -0.9384552 | 0.34801055 |
| tG(GCC)F2 | tG(GCC)F2 | 14.7545052 | -0.2730736 | 0.29132703 | -0.9373439 | 0.34858167 |
| Xu_2009_CUTs13 | Xu_2009_CU | 15.2039964 | 0.26202073 | 0.27970216 | 0.9367848 | 0.34886926 |
| YOL150C | YOL150C | 34.5485526 | -0.2200872 | 0.23505921 | -0.9363052 | 0.34911608 |
| OPI6 | YDL096C | 6.82206689 | 0.27692585 | 0.29586807 | 0.93597746 | 0.34928479 |
| BAR1 | YIL015W | 82.0046068 | 0.1649064 | 0.17619014 | 0.93595708 | 0.34929529 |
| PHO12 | YHR215W | 53.2076565 | 0.20038735 | 0.21414232 | 0.93576716 | 0.34939308 |
| Xu_2009_SUTs40 | Xu_2009_SU | 17.8120034 | 0.25714601 | 0.27485386 | 0.93557357 | 0.34949279 |
| OCA2 | YNL056W | 225.873539 | -0.1188527 | 0.12706474 | -0.9353713 | 0.34959698 |
| PEX13 | YLR191W | 389.501676 | -0.0918215 | 0.09822713 | -0.9347874 | 0.34989787 |
| CDC123 | YLR215C | 414.851577 | 0.08408637 | 0.09002795 | 0.93400301 | 0.35030234 |
| NUR1 | YDL089W | 276.906183 | -0.1046877 | 0.11211541 | -0.9337494 | 0.35043317 |
| Xu_2009_CUTs39 | Xu_2009_CU | 1.12722006 | 0.1851068 | 0.19870036 | 0.93158762 | 0.35154968 |
| YDL159W-A | YDL159W-A | 0.88406622 | -0.1555711 | 0.16703205 | -0.9313845 | 0.35165472 |
| YER147C-A | YER147C-A | 1.44133641 | -0.1977196 | 0.21226916 | -0.931457 | 0.35161723 |
| Xu_2009_SUTs31 | Xu_2009_SU | 111.134984 | 0.14575766 | 0.15654454 | 0.93109387 | 0.35180501 |
| YER079C-A | YER079C-A | 1.01468667 | 0.17765497 | 0.19085189 | 0.93085258 | 0.35192983 |
| SNR87 | snR87 | 121.32288 | 0.14395602 | 0.1547634 | 0.93016839 | 0.3522839 |
| Xu_2009_SUTs32 | Xu_2009_SU | 51.6662845 | 0.19462352 | 0.20931744 | 0.92980079 | 0.35247424 |
| tF(GAA)B | tF(GAA)B | 0.80396786 | 0.16181278 | 0.17417526 | 0.92902275 | 0.3528773 |
| Xu_2009_SUTs64 | Xu_2009_SU | 148.157483 | -0.1254969 | 0.13515609 | -0.9285329 | 0.3531312 |
| UTP13 | YLR222C | 1579.13242 | -0.0641897 | 0.0691702 | -0.9279968 | 0.35340921 |
| Xu_2009_SUTs43 | Xu_2009_SU | 18.201451 | -0.2541975 | 0.27414834 | -0.927226 | 0.35380921 |
| PTK1 | YKL198C | 305.032179 | -0.0977719 | 0.10544135 | -0.9272633 | 0.35378984 |
| Xu_2009_CUTs66 | Xu_2009_CU | 4.54061509 | 0.18276836 | 0.19720096 | 0.92681276 | 0.35402376 |
| TCD1 | YHR003C | 507.360334 | -0.0896947 | 0.09679055 | -0.9266884 | 0.35408833 |
| MPH3 | YJR160C | 20.0623771 | -0.2549385 | 0.27536855 | -0.9258083 | 0.35454559 |
| SSY1 | YDR160W | 533.926998 | -0.0774985 | 0.08375035 | -0.9253508 | 0.35478346 |
| UTP23 | YOR004W | 474.152408 | -0.0901419 | 0.09740825 | -0.9254033 | 0.35475614 |
| PXP2 | YJR111C | 618.622542 | -0.0930763 | 0.10059967 | -0.9252151 | 0.35485404 |

| | | | | | | |
|---|---|---|---|---|---|---|
| YOL085C | YOL085C | 4.65813194 | 0.26350247 | 0.28508175 | 0.92430492 | 0.35532758 |
| tV(AAC)H | tV(AAC)H | 2.6256906 | -0.2391951 | 0.25887557 | -0.9239771 | 0.35549825 |
| Xu_2009_SUTs30( | Xu_2009_SU | 35.9114462 | -0.2200756 | 0.23844251 | -0.9229713 | 0.35602218 |
| YIL134C-A | YIL134C-A | 11.0612733 | 0.27038538 | 0.29303059 | 0.92272064 | 0.3561528 |
| PCL6 | YER059W | 421.002201 | -0.0909585 | 0.09864226 | -0.9221045 | 0.35647405 |
| MPH2 | YDL247W | 8.69624038 | 0.27325474 | 0.29650584 | 0.92158299 | 0.35674614 |
| JEM1 | YJL073W | 630.575454 | -0.0719806 | 0.07817231 | -0.9207938 | 0.3571581 |
| YKL107W | YKL107W | 26.7053086 | -0.2318374 | 0.25176175 | -0.9208602 | 0.35712341 |
| YDR464C-A | YDR464C-A | 1.1529262 | -0.1769379 | 0.19254283 | -0.9189532 | 0.35812005 |
| YER145C-A | YER145C-A | 11.6417224 | 0.26906348 | 0.29331797 | 0.91730989 | 0.35898027 |
| Xu_2009_CUTs14 | Xu_2009_CU | 6.80526282 | 0.27079463 | 0.29529889 | 0.91701879 | 0.3591328 |
| RTT107 | YHR154W | 940.464252 | 0.06961893 | 0.07591523 | 0.91706139 | 0.35911047 |
| YGL188C-A | YGL188C-A | 144.164002 | -0.1319719 | 0.14397142 | -0.9166537 | 0.35932412 |
| Xu_2009_CUTs11 | Xu_2009_CU | 28.6550367 | -0.2280472 | 0.2488191 | -0.9165182 | 0.35939516 |
| SSP2 | YOR242C | 120.494833 | 0.1357041 | 0.14814658 | 0.91601236 | 0.35966041 |
| CDC14 | YFR028C | 868.81742 | -0.0691821 | 0.07553402 | -0.9159064 | 0.35971599 |
| ACC1 | YNR016C | 18298.4482 | -0.0483868 | 0.05284078 | -0.9157102 | 0.35981893 |
| ENP1 | YBR247C | 1249.27857 | 0.07586064 | 0.08289936 | 0.91509321 | 0.3601427 |
| Xu_2009_CUTs32 | Xu_2009_CU | 4.35374812 | -0.2619145 | 0.28629133 | -0.914853 | 0.3602688 |
| Xu_2009_CUTs36 | Xu_2009_CU | 14.78655 | 0.25893608 | 0.28304271 | 0.9148304 | 0.36028067 |
| SGO1 | YOR073W | 344.548258 | -0.0913699 | 0.09997602 | -0.9139181 | 0.36075989 |
| tY(GUA)J1/SUP7 | tY(GUA)J1 | 7.67075384 | -0.2706325 | 0.29649403 | -0.9127755 | 0.36136063 |
| YMR294W-A | YMR294W-A | 5.90842715 | 0.26704023 | 0.29270266 | 0.91232593 | 0.36159717 |
| CBP1 | YJL209W | 777.602188 | -0.067521 | 0.0740514 | -0.9118127 | 0.36186734 |
| LSM8 | YJR022W | 93.3970834 | -0.1548711 | 0.16982312 | -0.9119552 | 0.36179228 |
| ALD3 | YMR169C | 456.362437 | -0.094987 | 0.10417653 | -0.9117893 | 0.36187965 |
| YLR460C | YLR460C | 30.6562338 | 0.22826718 | 0.25040194 | 0.91160308 | 0.3619777 |
| Xu_2009_SUTs14( | Xu_2009_SU | 23.9408277 | -0.2391972 | 0.26247089 | -0.9113283 | 0.3621224 |
| TGL2 | YDR058C | 371.966754 | -0.0982636 | 0.10792309 | -0.9104968 | 0.36256056 |
| RMD6 | YEL072W | 190.644234 | -0.1155673 | 0.12698187 | -0.910109 | 0.36276503 |
| EMP46 | YLR080W | 256.474259 | -0.1151288 | 0.12665334 | -0.9090071 | 0.36334637 |
| PAU4 | YLR461W | 8.16664009 | 0.26933202 | 0.29641124 | 0.90864308 | 0.36353856 |
| tS(UGA)E/SUP17 | tS(UGA)E | 9.3050932 | 0.2688153 | 0.29605396 | 0.90799429 | 0.36388124 |
| YNL285W | YNL285W | 5.08559774 | 0.26222978 | 0.28885378 | 0.90782882 | 0.36396867 |
| Xu_2009_CUTs16 | Xu_2009_CU | 8.95398147 | 0.26840667 | 0.29577331 | 0.90747426 | 0.36415606 |
| CDC43 | YGL155W | 634.461873 | 0.07124257 | 0.0785299 | 0.90720313 | 0.36429939 |
| YGL072C | YGL072C | 141.103002 | -0.1303969 | 0.14383322 | -0.9065842 | 0.36462672 |
| PUS4 | YNL292W | 446.90992 | 0.08128594 | 0.08970339 | 0.90616354 | 0.3648493 |
| Xu_2009_SUTs50( | Xu_2009_SU | 192.115124 | 0.11274405 | 0.12443514 | 0.90604678 | 0.36491109 |
| FAA2 | YER015W | 206.968081 | -0.1108886 | 0.12242352 | -0.9057785 | 0.36505312 |
| tC(GCA)P1 | tC(GCA)P1 | 1.42112637 | -0.1896128 | 0.20958376 | -0.9047111 | 0.36561847 |

| | | | | | | |
|---|---|---|---|---|---|---|
| MET14 | YKL001C | 296.300591 | 0.09610303 | 0.10626195 | 0.90439735 | 0.36578474 |
| Q0255 | Q0255 | 18.9165028 | -0.2459939 | 0.27211151 | -0.9040187 | 0.36598547 |
| PRS3 | YHL011C | 1327.94164 | -0.0650036 | 0.07191033 | -0.9039538 | 0.36601987 |
| THP2 | YHR167W | 254.588388 | -0.1002827 | 0.11089891 | -0.9042713 | 0.36585154 |
| YOL035C | YOL035C | 3.1012821 | -0.2357712 | 0.26081474 | -0.9039796 | 0.36600621 |
| YLR154W-E | YLR154W-E | 4.42346717 | -0.2565137 | 0.28383873 | -0.9037306 | 0.36613828 |
| Xu_2009_SUTs66 | Xu_2009_SU | 53.0604989 | -0.1771634 | 0.19612043 | -0.90334 | 0.36634547 |
| Xu_2009_SUTs81 | Xu_2009_SU | 19.3665146 | -0.2418014 | 0.26801548 | -0.9021918 | 0.366955 |
| YJR140W-A | YJR140W-A | 1.81606723 | -0.2053258 | 0.22763895 | -0.90198 | 0.36706747 |
| FKS3 | YMR306W | 222.88982 | 0.11290835 | 0.1252057 | 0.90178285 | 0.36717223 |
| tS(UGA)Q2 | tS(UGA)Q2 | 4.06060147 | -0.2513195 | 0.27908728 | -0.9005049 | 0.36785161 |
| RIB2 | YOL066C | 442.422029 | 0.09092487 | 0.10104517 | 0.89984372 | 0.36820343 |
| COQ10 | YOL008W | 149.106379 | -0.1262597 | 0.14048196 | -0.8987607 | 0.36878014 |
| Xu_2009_CUTs87 | Xu_2009_CU | 0.80664241 | -0.1504289 | 0.16753557 | -0.897892 | 0.36924312 |
| RRM3 | YHR031C | 501.842483 | -0.0816643 | 0.09094294 | -0.8979733 | 0.36919981 |
| Xu_2009_CUTs22 | Xu_2009_CU | 19.4711456 | 0.24843157 | 0.27693765 | 0.8970668 | 0.36968328 |
| ENA2 | YDR039C | 7.32411232 | 0.26570101 | 0.29624421 | 0.89689856 | 0.36977305 |
| YER034W | YER034W | 468.821944 | -0.0840932 | 0.09376522 | -0.8968486 | 0.36979968 |
| PMC1 | YGL006W | 1778.13098 | 0.05463255 | 0.061014 | 0.89541001 | 0.37056795 |
| Xu_2009_SUTs50 | Xu_2009_SU | 43.4959691 | 0.20944319 | 0.2339906 | 0.89509232 | 0.37073774 |
| YBL096C | YBL096C | 41.9223071 | -0.1951866 | 0.21836073 | -0.8938724 | 0.37139017 |
| SPE1 | YKL184W | 834.094112 | -0.0687527 | 0.07691348 | -0.8938963 | 0.37137739 |
| YEL032C-A | YEL032C-A | 0.93523404 | 0.1505981 | 0.16855086 | 0.89348762 | 0.3715961 |
| RFT1 | YBL020W | 854.770174 | -0.0669774 | 0.07498472 | -0.8932136 | 0.37174279 |
| YPL197C | YPL197C | 5.53427827 | 0.25680402 | 0.28751344 | 0.89318962 | 0.37175564 |
| HFI1 | YPL254W | 622.267671 | 0.07075656 | 0.07924499 | 0.89288363 | 0.3719195 |
| Xu_2009_CUTs86 | Xu_2009_CU | 3.88542257 | 0.24491398 | 0.2743287 | 0.89277563 | 0.37197734 |
| YPR092W | YPR092W | 4.8172946 | 0.25355671 | 0.28419729 | 0.89218554 | 0.37229349 |
| MTF1 | YMR228W | 239.086999 | -0.1052328 | 0.11799907 | -0.8918106 | 0.37249445 |
| MSL5 | YLR116W | 1248.73826 | -0.0567082 | 0.06360057 | -0.8916311 | 0.37259069 |
| YIL066W-A | YIL066W-A | 0.8141993 | 0.15067355 | 0.16920449 | 0.89048201 | 0.37320712 |
| SEY1 | YOR165W | 1208.80687 | 0.05817199 | 0.06533083 | 0.89042166 | 0.37323951 |
| UTP7 | YER082C | 1960.31612 | 0.06233752 | 0.07004161 | 0.890007 | 0.37346213 |
| SNR58 | snR58 | 23.2531243 | 0.22993503 | 0.25853149 | 0.88938888 | 0.37379412 |
| Xu_2009_CUTs13 | Xu_2009_CU | 3.35690887 | 0.2408938 | 0.27085029 | 0.88939834 | 0.37378904 |
| DAL3 | YIR032C | 50.1259098 | -0.1857632 | 0.20883663 | -0.8895144 | 0.3737267 |
| OPI9 | YLR338W | 1.71666264 | -0.1991976 | 0.2239312 | -0.8895483 | 0.3737085 |
| VPS60 | YDR486C | 879.881633 | 0.06252955 | 0.07040085 | 0.88819307 | 0.37443691 |
| RRG8 | YPR116W | 193.32798 | 0.1124548 | 0.12666759 | 0.88779456 | 0.37465127 |
| Xu_2009_CUTs20 | Xu_2009_CU | 7.77365351 | 0.2628222 | 0.29637707 | 0.88678315 | 0.37519566 |
| Xu_2009_CUTs47 | Xu_2009_CU | 1.42510339 | 0.18786589 | 0.2118589 | 0.88675004 | 0.37521349 |

| | | | | | | |
|---|---|---|---|---|---|---|
| RAP1 | YNL216W | 1221.93985 | 0.05819511 | 0.06577883 | 0.88470878 | 0.37631372 |
| YPR064W | YPR064W | 7.65314688 | 0.25963602 | 0.29457577 | 0.8813896 | 0.37810698 |
| Xu_2009_CUTs55 | Xu_2009_CU | 4.69150784 | 0.25290465 | 0.28715954 | 0.88071128 | 0.37847411 |
| Xu_2009_CUTs50 | Xu_2009_CU | 10.1952031 | 0.26003008 | 0.29550077 | 0.87996413 | 0.37887874 |
| YOL166C | YOL166C | 0.79473845 | 0.14149969 | 0.16089432 | 0.87945731 | 0.37915337 |
| MMM1 | YLL006W | 235.300527 | -0.1000545 | 0.11384606 | -0.8788581 | 0.3794782 |
| VPS64 | YDR200C | 841.237092 | 0.06425241 | 0.07312466 | 0.87866948 | 0.37958051 |
| Xu_2009_CUTs45 | Xu_2009_CU | 8.75009818 | 0.26024249 | 0.29645471 | 0.87784907 | 0.38002563 |
| RFU1 | YLR073C | 540.819769 | 0.0826214 | 0.09422886 | 0.87681637 | 0.38058639 |
| RDS2 | YPL133C | 447.343925 | 0.0802959 | 0.0916733 | 0.87589194 | 0.38108878 |
| YHR022C-A | YHR022C-A | 4.14491045 | 0.2438734 | 0.27900005 | 0.87409804 | 0.38206486 |
| Xu_2009_SUTs62 | Xu_2009_SU | 127.966659 | -0.1310422 | 0.1499719 | -0.8737784 | 0.38223892 |
| BUD25 | YER014C-A | 4.15177039 | -0.2473408 | 0.28305907 | -0.8738134 | 0.38221987 |
| Xu_2009_SUTs20 | Xu_2009_SU | 151.485548 | 0.12547792 | 0.14367875 | 0.87332271 | 0.38248721 |
| MCO32 | YGR053C | 66.5111619 | -0.1668175 | 0.19102893 | -0.873258 | 0.38252246 |
| MID1 | YNL291C | 795.073018 | -0.0693747 | 0.07943263 | -0.8733775 | 0.38245736 |
| YGL176C | YGL176C | 470.877522 | -0.0883062 | 0.10115719 | -0.8729603 | 0.38268472 |
| VPS36 | YLR417W | 506.210126 | 0.08274017 | 0.0947946 | 0.87283629 | 0.38275232 |
| YML037C | YML037C | 265.809281 | 0.10013883 | 0.11479261 | 0.87234562 | 0.38301986 |
| Xu_2009_CUTs64 | Xu_2009_CU | 6.47615836 | 0.25698543 | 0.29465038 | 0.87217072 | 0.38311525 |
| Xu_2009_SUTs18 | Xu_2009_SU | 18.9786199 | 0.24049134 | 0.27594164 | 0.87152971 | 0.38346499 |
| APN1 | YKL114C | 641.779034 | -0.067587 | 0.07757037 | -0.8712988 | 0.383591 |
| tR(UCU)K | tR(UCU)K | 21.1531802 | -0.2341848 | 0.26896876 | -0.8706767 | 0.38393068 |
| YML094C-A | YML094C-A | 16.9868728 | 0.24216288 | 0.27829886 | 0.87015405 | 0.38421622 |
| Xu_2009_SUTs36 | Xu_2009_SU | 67.8739136 | 0.16013588 | 0.18417697 | 0.86946741 | 0.38459153 |
| Xu_2009_SUTs74 | Xu_2009_SU | 192.61023 | 0.12030239 | 0.13845699 | 0.86887912 | 0.38491325 |
| TEL2 | YGR099W | 304.257117 | 0.09012957 | 0.10376359 | 0.86860503 | 0.3850632 |
| OSW7 | YFR039C | 540.478446 | -0.0755695 | 0.08701881 | -0.868427 | 0.3851606 |
| Xu_2009_CUTs22 | Xu_2009_CU | 5.19965793 | -0.2511062 | 0.28930763 | -0.8679558 | 0.38541855 |
| API2 | YDR525W | 8.064823 | 0.25721164 | 0.2963862 | 0.86782596 | 0.38548962 |
| tY(GUA)O/SUP3 | tY(GUA)O | 0.98431536 | -0.1474512 | 0.1700494 | -0.8671079 | 0.38588288 |
| ILT1 | YDR090C | 247.940992 | -0.0977327 | 0.11281462 | -0.8663125 | 0.38631883 |
| Xu_2009_CUTs84 | Xu_2009_CU | 19.6777782 | 0.2336639 | 0.2700366 | 0.86530458 | 0.38687164 |
| YOR387C | YOR387C | 2.67170045 | 0.21938134 | 0.25360273 | 0.86505908 | 0.38700636 |
| YMR193C-A | YMR193C-A | 1.87295692 | 0.20423823 | 0.23616121 | 0.86482545 | 0.3871346 |
| RAD24 | YER173W | 411.259606 | -0.0783423 | 0.09066149 | -0.864119 | 0.38752252 |
| Xu_2009_CUTs70 | Xu_2009_CU | 6.61862828 | 0.25490838 | 0.29503756 | 0.8639862 | 0.38759547 |
| YDL114W | YDL114W | 32.129854 | -0.2057788 | 0.23821954 | -0.8638199 | 0.38768684 |
| CDC27 | YBL084C | 501.017461 | 0.07755407 | 0.08980879 | 0.86354655 | 0.38783704 |
| POM152 | YMR129W | 1944.03101 | 0.04897382 | 0.05671418 | 0.86351991 | 0.38785168 |
| POL31 | YJR006W | 528.516909 | 0.07998151 | 0.09264412 | 0.86331985 | 0.38796163 |

| | | | | | | |
|---|---|---|---|---|---|---|
| YOR008C-A | YOR008C-A | 13.2226178 | 0.24798744 | 0.28746126 | 0.86268124 | 0.38831275 |
| Xu_2009_SUTs51! | Xu_2009_SU | 9.33069097 | 0.2548977 | 0.29557469 | 0.86238002 | 0.38847843 |
| PGS1 | YCL004W | 338.389438 | -0.0835601 | 0.0969119 | -0.8622278 | 0.38856218 |
| ROG1 | YGL144C | 739.0448 | -0.0705933 | 0.08187431 | -0.862216 | 0.38856867 |
| MOH1 | YBL049W | 95.0775164 | 0.15126296 | 0.17545889 | 0.86209914 | 0.38863297 |
| APC11 | YDL008W | 271.059451 | 0.09453252 | 0.10968514 | 0.86185346 | 0.38876816 |
| HRA1 | HRA1 | 3.27377541 | -0.2352736 | 0.27315198 | -0.8613287 | 0.38905705 |
| YDR182W-A | YDR182W-A | 22.3814474 | 0.22868883 | 0.26561077 | 0.86099231 | 0.38924228 |
| APM4 | YOL062C | 592.498524 | -0.0684193 | 0.07949097 | -0.8607173 | 0.38939377 |
| Xu_2009_SUTs46 | Xu_2009_SU | 13.5637011 | -0.2469437 | 0.2872728 | -0.859614 | 0.39000186 |
| PHO90 | YJL198W | 1301.91846 | 0.07009306 | 0.08159906 | 0.85899345 | 0.39034413 |
| PRP42 | YDR235W | 649.714567 | -0.0739318 | 0.08607972 | -0.8588763 | 0.39040874 |
| YJL150W | YJL150W | 12.4799504 | 0.25053751 | 0.29175841 | 0.85871564 | 0.39049742 |
| tR(UCU)G1 | tR(UCU)G1 | 3.70454151 | -0.2363345 | 0.27535917 | -0.8582773 | 0.39073935 |
| DPB3 | YBR278W | 185.297861 | -0.1073498 | 0.12512011 | -0.8579744 | 0.39090662 |
| RPC37 | YKR025W | 388.59688 | 0.09079992 | 0.10584325 | 0.85787166 | 0.39096333 |
| Xu_2009_SUTs48 | Xu_2009_SU | 2.13324341 | -0.2019118 | 0.23547058 | -0.8574821 | 0.39117852 |
| PET309 | YLR067C | 854.404624 | 0.06546497 | 0.07641258 | 0.85673034 | 0.39159393 |
| YIL068W-A | YIL068W-A | 3.66020764 | 0.23779734 | 0.27773374 | 0.85620616 | 0.39188376 |
| Xu_2009_CUTs34 | Xu_2009_CU | 6.35812352 | 0.25211177 | 0.29452664 | 0.8559897 | 0.39200348 |
| SRL2 | YLR082C | 207.757735 | -0.1010578 | 0.11810689 | -0.8556467 | 0.39219326 |
| Xu_2009_SUTs30 | Xu_2009_SU | 17.5375722 | 0.23689771 | 0.27713002 | 0.85482514 | 0.39264796 |
| SUT2 | YPR009W | 613.221419 | 0.07043916 | 0.08241848 | 0.85465255 | 0.39274353 |
| GIM3 | YNL153C | 833.867342 | 0.06908465 | 0.08084728 | 0.85450811 | 0.39282352 |
| PIR5 | YJL160C | 136.529568 | -0.1225631 | 0.14348725 | -0.8541745 | 0.39300833 |
| YOR385W | YOR385W | 2592.23131 | 0.11092062 | 0.12989701 | 0.853912 | 0.39315375 |
| tM(CAU)D/EMT1 | tM(CAU)D | 15.6391717 | 0.24159458 | 0.28301735 | 0.85363877 | 0.39330517 |
| Xu_2009_SUTs36 | Xu_2009_SU | 22.05482 | 0.22408686 | 0.2625667 | 0.85344738 | 0.39341126 |
| Xu_2009_SUTs38 | Xu_2009_SU | 13.4805776 | 0.24267481 | 0.28457452 | 0.85276366 | 0.39379038 |
| YKL177W | YKL177W | 3.54610866 | 0.23195214 | 0.27200016 | 0.85276472 | 0.39378979 |
| Xu_2009_CUTs71 | Xu_2009_CU | 11.8774997 | 0.24600319 | 0.28936429 | 0.85015048 | 0.39524143 |
| YPR002C-A | YPR002C-A | 8.2395138 | -0.2510793 | 0.29541681 | -0.8499154 | 0.39537215 |
| AIM33 | YML087C | 61.5186169 | 0.17025255 | 0.20036551 | 0.84970987 | 0.39548641 |
| QCR2 | YPR191W | 2704.58084 | -0.055628 | 0.06546788 | -0.8496997 | 0.39549208 |
| Xu_2009_CUTs50 | Xu_2009_CU | 2.73893739 | 0.2213051 | 0.26066761 | 0.84899348 | 0.39588492 |
| tE(UUC)P | tE(UUC)P | 4.59993802 | -0.2418773 | 0.28503078 | -0.8486007 | 0.39610352 |
| MNT4 | YNR059W | 459.675304 | 0.07732733 | 0.09115113 | 0.84834199 | 0.39624754 |
| YHR145C | YHR145C | 9.13145507 | 0.25076853 | 0.29570253 | 0.84804323 | 0.39641389 |
| CCT3 | YJL014W | 3376.30558 | 0.04953614 | 0.05842384 | 0.84787534 | 0.3965074 |
| EFM1 | YHL039W | 693.874636 | -0.0737846 | 0.08713762 | -0.8467592 | 0.39712933 |
| SRB6 | YBR253W | 259.816671 | -0.0933615 | 0.11036285 | -0.8459505 | 0.39758038 |

| | | | | | | |
|---|---|---|---|---|---|---|
| RFM1 | YOR279C | 345.045995 | 0.08577542 | 0.10149039 | 0.84515808 | 0.39802258 |
| CDC48 | YDL126C | 9325.09538 | -0.041929 | 0.04962696 | -0.8448825 | 0.39817644 |
| Xu_2009_SUTs77: | Xu_2009_SU | 27.6447361 | 0.21644799 | 0.25646352 | 0.84397182 | 0.39868515 |
| Xu_2009_CUTs46 | Xu_2009_CU | 20.7167921 | 0.22205699 | 0.26315713 | 0.84381902 | 0.39877055 |
| Xu_2009_CUTs11 | Xu_2009_CU | 9.17569342 | 0.24913164 | 0.29536883 | 0.84345948 | 0.39897152 |
| DCR2 | YLR361C | 446.045886 | 0.07561575 | 0.08965959 | 0.84336482 | 0.39902444 |
| ELG1 | YOR144C | 654.28199 | 0.06655469 | 0.07906254 | 0.84179806 | 0.399901 |
| FMP30 | YPL103C | 257.11126 | -0.0982073 | 0.11666856 | -0.841763 | 0.39992064 |
| YKR012C | YKR012C | 13.6755252 | 0.24194168 | 0.28762826 | 0.84116102 | 0.40025773 |
| MIC12 | YBR262C | 610.076406 | 0.07906131 | 0.09402263 | 0.84087535 | 0.40041777 |
| Xu_2009_SUTs24: | Xu_2009_SU | 16.3883769 | 0.2336685 | 0.27816428 | 0.84003778 | 0.40088721 |
| CTF4 | YPR135W | 680.936326 | 0.06593912 | 0.07850705 | 0.83991335 | 0.40095697 |
| SPO16 | YHR153C | 16.049716 | -0.2347295 | 0.27993246 | -0.838522 | 0.40173759 |
| Xu_2009_SUTs23 | Xu_2009_SU | 16.5654683 | 0.2367558 | 0.28252368 | 0.83800336 | 0.40202882 |
| MRX6 | YNL295W | 694.910506 | 0.06729432 | 0.08030665 | 0.83796696 | 0.40204926 |
| MSN1 | YOL116W | 377.014993 | -0.0788233 | 0.09412287 | -0.8374512 | 0.40233899 |
| Xu_2009_SUTs88 | Xu_2009_SU | 10.6668247 | -0.2449107 | 0.2926949 | -0.8367438 | 0.40273657 |
| Xu_2009_CUTs23 | Xu_2009_CU | 6.21202669 | 0.23993395 | 0.28718677 | 0.8354631 | 0.40345702 |
| YDL011C | YDL011C | 1.52645855 | -0.1811657 | 0.21703757 | -0.8347203 | 0.4038752 |
| PAN6 | YIL145C | 991.877974 | 0.05975203 | 0.07157257 | 0.83484539 | 0.40380476 |
| NIT1 | YIL164C | 91.8351924 | -0.1390701 | 0.16659824 | -0.8347634 | 0.40385096 |
| SNR9 | snR9 | 937.161237 | -0.08393 | 0.10061543 | -0.8341667 | 0.40418703 |
| YJL118W | YJL118W | 279.143111 | -0.0868724 | 0.1041346 | -0.8342317 | 0.40415045 |
| TEC1 | YBR083W | 908.12362 | -0.0666097 | 0.07987782 | -0.8338953 | 0.40433998 |
| PEP3 | YLR148W | 705.70414 | 0.06778896 | 0.08135102 | 0.8332896 | 0.40468142 |
| MAN2 | YNR073C | 37.1510719 | -0.1944 | 0.23327875 | -0.8333379 | 0.40465417 |
| MAM1 | YER106W | 38.0321641 | -0.1993324 | 0.23965884 | -0.8317339 | 0.40555913 |
| GYP6 | YJL044C | 737.950687 | 0.06223493 | 0.07482741 | 0.83171303 | 0.40557094 |
| YHL008C | YHL008C | 1512.0402 | 0.05598977 | 0.06733612 | 0.83149677 | 0.40569305 |
| tL(CAA)G2/SUP54 | tL(CAA)G2 | 32.756081 | 0.201721 | 0.2428624 | 0.83059792 | 0.40620081 |
| Xu_2009_CUTs23 | Xu_2009_CU | 106.373077 | 0.12957562 | 0.15597058 | 0.83076962 | 0.40610379 |
| PEX12 | YMR026C | 486.563718 | 0.07107357 | 0.08556911 | 0.83059844 | 0.40620051 |
| Xu_2009_CUTs45 | Xu_2009_CU | 12.5327927 | 0.2382503 | 0.28704747 | 0.83000312 | 0.40653702 |
| Xu_2009_SUTs83: | Xu_2009_SU | 64.0796291 | -0.15785 | 0.19020714 | -0.8298845 | 0.40660407 |
| KIP1 | YBL063W | 566.650884 | 0.0684309 | 0.08241817 | 0.83028892 | 0.40637545 |
| YCR050C | YCR050C | 4.92752107 | 0.23708933 | 0.28567786 | 0.82991847 | 0.40658488 |
| SEN2 | YLR105C | 562.835818 | -0.0681561 | 0.08211402 | -0.8300174 | 0.40652895 |
| tG(GCC)D2 | tG(GCC)D2 | 167.462634 | 0.11209164 | 0.13516946 | 0.82926744 | 0.40695309 |
| Xu_2009_SUTs80 | Xu_2009_SU | 29.0994457 | 0.1998899 | 0.24117202 | 0.82882708 | 0.40720226 |
| RTN2 | YDL204W | 1301.07403 | -0.0658609 | 0.07955014 | -0.8279172 | 0.40771741 |
| LOS1 | YKL205W | 1426.24892 | 0.06045114 | 0.07301909 | 0.8278813 | 0.40773773 |

| | | | | | | |
|---|---|---|---|---|---|---|
| YGL081W | YGL081W | 170.933024 | 0.11134099 | 0.134518 | 0.82770328 | 0.40783856 |
| Xu_2009_CUTs68 | Xu_2009_CU | 29.2380664 | 0.20144576 | 0.24341474 | 0.82758244 | 0.40790702 |
| VAM10 | YOR068C | 1.53374141 | 0.16612749 | 0.2009831 | 0.82657442 | 0.40847832 |
| Xu_2009_CUTs60 | Xu_2009_CU | 2.1510207 | 0.19284621 | 0.23335306 | 0.8264139 | 0.40856934 |
| HRD3 | YLR207W | 1048.9346 | -0.0597038 | 0.07228614 | -0.8259369 | 0.40883991 |
| YIL059C | YIL059C | 101.941032 | -0.1405049 | 0.17038286 | -0.824642 | 0.40957487 |
| YJL052C-A | YJL052C-A | 1.02357746 | 0.1584958 | 0.19222582 | 0.82452919 | 0.40963893 |
| UTR1 | YJR049C | 619.821763 | -0.0693213 | 0.08408214 | -0.8244477 | 0.40968522 |
| YNL203C | YNL203C | 163.005245 | 0.10934744 | 0.13268119 | 0.82413672 | 0.40986188 |
| EDC2 | YER035W | 965.474197 | -0.0621332 | 0.07547417 | -0.8232386 | 0.41037232 |
| SPT4 | YGR063C | 329.170465 | -0.0832328 | 0.10112062 | -0.8231044 | 0.41044861 |
| MHP1 | YJL042W | 2867.4304 | -0.0453279 | 0.05504795 | -0.8234257 | 0.41026594 |
| YNL277W-A | YNL277W-A | 32.5064612 | 0.19972373 | 0.24264353 | 0.82311582 | 0.41044214 |
| YOL159C-A | YOL159C-A | 139.598053 | -0.1314694 | 0.159676 | -0.8233511 | 0.41030839 |
| VPS5 | YOR069W | 832.240771 | -0.0586235 | 0.07121196 | -0.8232254 | 0.41037983 |
| NSR1 | YGR159C | 5217.94993 | 0.11365944 | 0.13815746 | 0.82268046 | 0.41068973 |
| UBP2 | YOR124C | 2234.58858 | 0.05356065 | 0.0651322 | 0.82233755 | 0.41088481 |
| Xu_2009_SUTs41 | Xu_2009_SU | 15.2751932 | -0.2301426 | 0.27997835 | -0.8220014 | 0.41107609 |
| tH(GUG)H | tH(GUG)H | 0.91187586 | -0.1428808 | 0.17404665 | -0.8209338 | 0.41168396 |
| FUN12 | YAL035W | 5876.64228 | -0.0459593 | 0.05599982 | -0.8207047 | 0.41181448 |
| Xu_2009_CUTs16 | Xu_2009_CU | 9.11426053 | 0.24310948 | 0.29648389 | 0.81997535 | 0.41223016 |
| YOR345C | YOR345C | 4.4149998 | 0.23269758 | 0.2838168 | 0.81988659 | 0.41228076 |
| YBR089W | YBR089W | 2.74173366 | -0.2113299 | 0.25786727 | -0.8195297 | 0.41248426 |
| GSM1 | YJL103C | 380.344934 | 0.08391521 | 0.10241691 | 0.81934916 | 0.41258723 |
| Xu_2009_SUTs45 | Xu_2009_SU | 92.2702597 | -0.1374077 | 0.16794294 | -0.818181 | 0.41325382 |
| CIT2 | YCR005C | 3219.79363 | -0.1214316 | 0.14857919 | -0.8172855 | 0.4137653 |
| POT1 | YIL160C | 10.0296501 | -0.24045 | 0.2943159 | -0.8169792 | 0.41394031 |
| Xu_2009_CUTs32 | Xu_2009_CU | 3.63298736 | -0.2176587 | 0.26729564 | -0.8142995 | 0.4154734 |
| CAD1 | YDR423C | 631.570922 | 0.066404 | 0.08159015 | 0.81387269 | 0.41571789 |
| YBR141W-A | YBR141W-A | 38.1529234 | 0.18080624 | 0.22225319 | 0.81351472 | 0.41592301 |
| SRB5 | YGR104C | 359.040722 | -0.0781416 | 0.09615858 | -0.8126324 | 0.41642886 |
| ERO1 | YML130C | 2847.26968 | 0.04526973 | 0.0557289 | 0.81232048 | 0.41660776 |
| Xu_2009_CUTs15 | Xu_2009_CU | 15.3908801 | 0.23220714 | 0.28609118 | 0.8116543 | 0.41699002 |
| Xu_2009_SUTs28 | Xu_2009_SU | 4.21009216 | 0.22660608 | 0.27940864 | 0.81102032 | 0.417354 |
| VPH2 | YKL119C | 111.500861 | -0.1248736 | 0.15405263 | -0.8105907 | 0.41760078 |
| YHR213W-B | YHR213W-B | 1.18762242 | 0.15961203 | 0.19695459 | 0.81040014 | 0.41771024 |
| Xu_2009_CUTs17 | Xu_2009_CU | 17.0142922 | 0.22788916 | 0.28158312 | 0.80931401 | 0.41833455 |
| MET32 | YDR253C | 40.5803824 | 0.1893167 | 0.23414439 | 0.80854685 | 0.41877585 |
| PIC2 | YER053C | 1090.30072 | -0.070472 | 0.0871701 | -0.8084429 | 0.41883564 |
| RAD53 | YPL153C | 425.792746 | -0.0765903 | 0.09474832 | -0.8083552 | 0.41888616 |
| Xu_2009_CUTs31 | Xu_2009_CU | 5.32371411 | 0.23522376 | 0.29128228 | 0.80754572 | 0.41935215 |

| | | | | | | |
|---|---|---|---|---|---|---|
| YHR140W | YHR140W | 104.043272 | -0.1327779 | 0.16445514 | -0.8073809 | 0.41944708 |
| HPM1 | YIL110W | 536.429595 | -0.0701535 | 0.08688803 | -0.8074014 | 0.41943527 |
| YLR124W | YLR124W | 40.6873022 | -0.1830572 | 0.22671588 | -0.80743 | 0.41941882 |
| RVB2 | YPL235W | 1217.50664 | -0.0561821 | 0.06958519 | -0.807386 | 0.41944415 |
| YBL055C | YBL055C | 533.850151 | 0.07560025 | 0.09372952 | 0.80657888 | 0.41990915 |
| YMR194C-A | YMR194C-A | 17.1848918 | -0.2288812 | 0.283781 | -0.8065416 | 0.41993064 |
| ASG1 | YIL130W | 1684.84309 | 0.04763252 | 0.05915247 | 0.80524979 | 0.42067555 |
| IAH1 | YOR126C | 166.996205 | -0.1073809 | 0.13358595 | -0.8038339 | 0.42149293 |
| STE11 | YLR362W | 648.557316 | 0.06141854 | 0.07646343 | 0.80324073 | 0.42183561 |
| FLC2 | YAL053W | 3841.76057 | 0.04342644 | 0.05407797 | 0.80303379 | 0.42195521 |
| PKP1 | YIL042C | 431.11296 | -0.0859518 | 0.10706999 | -0.8027631 | 0.42211167 |
| YGL138C | YGL138C | 4.13742269 | 0.22617138 | 0.28180886 | 0.80257015 | 0.42222323 |
| YBR242W | YBR242W | 437.39661 | -0.0772074 | 0.09621928 | -0.8024108 | 0.42231537 |
| RQC1 | YDR333C | 962.452542 | 0.05804327 | 0.07236743 | 0.80206348 | 0.42251624 |
| YOR161C-C | YOR161C-C | 13.7687873 | 0.23004476 | 0.286923 | 0.80176479 | 0.42268903 |
| Xu_2009_CUTs80 | Xu_2009_CU | 51.2937184 | 0.1618282 | 0.2020703 | 0.80085103 | 0.42321789 |
| Xu_2009_SUTs69 | Xu_2009_SU | 35.2469042 | 0.18436348 | 0.2302839 | 0.80059213 | 0.42336781 |
| GIS4 | YML006C | 1684.09986 | -0.0461118 | 0.05760668 | -0.8004596 | 0.42344454 |
| PIP2 | YOR363C | 778.56735 | 0.05756544 | 0.07194881 | 0.80008887 | 0.42365931 |
| YDL241W | YDL241W | 29.0231428 | 0.20933551 | 0.2616993 | 0.79990858 | 0.42376377 |
| YJL220W | YJL220W | 0.9474253 | 0.13938853 | 0.17431523 | 0.7996348 | 0.42392242 |
| YLL032C | YLL032C | 509.208661 | 0.06509383 | 0.08142465 | 0.79943645 | 0.42403738 |
| UBX2 | YML013W | 815.305179 | -0.0587382 | 0.07346808 | -0.7995064 | 0.42399683 |
| YDL068W | YDL068W | 9.37849089 | 0.23667085 | 0.29616046 | 0.79913049 | 0.42421475 |
| YOR055W | YOR055W | 1.38037329 | -0.1534475 | 0.19219106 | -0.7984111 | 0.42463199 |
| Xu_2009_CUTs40 | Xu_2009_CU | 8.08354409 | 0.23664977 | 0.29645781 | 0.79825783 | 0.42472088 |
| STL1 | YDR536W | 8.83079276 | 0.23583176 | 0.2954651 | 0.79817131 | 0.42477109 |
| tE(UUC)M | tE(UUC)M | 3.95602656 | -0.2219282 | 0.27832669 | -0.797366 | 0.42523851 |
| VPS65 | YLR322W | 2.35954558 | -0.1854548 | 0.23257882 | -0.7973848 | 0.42522757 |
| PTC1 | YDL006W | 398.644564 | 0.07582812 | 0.09513788 | 0.79703393 | 0.42543133 |
| USA1 | YML029W | 907.486882 | -0.0592115 | 0.07428213 | -0.7971169 | 0.42538316 |
| FUN26 | YAL022C | 760.794848 | 0.06566692 | 0.08241137 | 0.79681871 | 0.42555633 |
| FUS1 | YCL027W | 182.122444 | 0.10271101 | 0.12890683 | 0.79678481 | 0.42557602 |
| tK(CUU)F | tK(CUU)F | 1.61267351 | 0.17088414 | 0.2146999 | 0.79592092 | 0.426078 |
| YAR070C | YAR070C | 1.04599662 | 0.14803489 | 0.18600457 | 0.79586696 | 0.42610936 |
| EDS1 | YBR033W | 173.919618 | -0.103507 | 0.13006668 | -0.7957999 | 0.42614835 |
| YAP1801 | YHR161C | 1514.65665 | 0.05308949 | 0.06670415 | 0.79589496 | 0.42609309 |
| Xu_2009_SUTs47 | Xu_2009_SU | 72.6763829 | 0.14236734 | 0.17895329 | 0.79555588 | 0.42629022 |
| RPT2 | YDL007W | 2170.05228 | 0.04953308 | 0.06226356 | 0.79553875 | 0.42630018 |
| SMY1 | YKL079W | 805.197707 | 0.06134757 | 0.07714695 | 0.79520407 | 0.42649481 |
| YER188W | YER188W | 28.4884266 | 0.20433589 | 0.25708966 | 0.794804 | 0.42672752 |

| | | | | | | |
|---|---|---|---|---|---|---|
| CCM1 | YGR150C | 926.197668 | 0.05513409 | 0.06938055 | 0.794662 | 0.42681014 |
| YMR187C | YMR187C | 260.846517 | -0.0882935 | 0.11126117 | -0.7935694 | 0.42744613 |
| MLS1 | YNL117W | 20.3518124 | -0.2135214 | 0.26902815 | -0.7936768 | 0.42738364 |
| YLR339C | YLR339C | 2.2692087 | -0.1830134 | 0.23071288 | -0.793252 | 0.42763103 |
| YOR342C | YOR342C | 2103.12641 | 0.04759078 | 0.06000216 | 0.79315111 | 0.42768979 |
| YDL041W | YDL041W | 5.19705526 | 0.22785382 | 0.28744941 | 0.79267451 | 0.42796749 |
| YDR354C-A | YDR354C-A | 1.99322624 | -0.183854 | 0.23227988 | -0.7915192 | 0.42864111 |
| Xu_2009_CUTs71 | Xu_2009_CU | 24.5908533 | 0.19931731 | 0.25205941 | 0.79075526 | 0.42908682 |
| RAS1 | YOR101W | 865.067895 | -0.0582843 | 0.07372219 | -0.7905934 | 0.42918128 |
| Xu_2009_SUTs71 | Xu_2009_SU | 1.94306165 | 0.18827406 | 0.23843688 | 0.78961802 | 0.42975088 |
| SQT1 | YIR012W | 1682.38239 | -0.0527999 | 0.06690817 | -0.7891402 | 0.43003004 |
| Xu_2009_SUTs65 | Xu_2009_SU | 77.6434529 | 0.14078772 | 0.17843452 | 0.78901614 | 0.43010257 |
| Xu_2009_CUTs54 | Xu_2009_CU | 9.80941244 | 0.23342496 | 0.29595066 | 0.78872932 | 0.43027023 |
| RRP43 | YCR035C | 957.903016 | -0.0588472 | 0.07460139 | -0.7888224 | 0.43021582 |
| TFB4 | YPR056W | 433.632039 | -0.0729199 | 0.09255187 | -0.7878819 | 0.43076582 |
| YLR285C-A | YLR285C-A | 5.38049897 | -0.2242777 | 0.28476306 | -0.7875941 | 0.43093415 |
| Xu_2009_SUTs35 | Xu_2009_SU | 30.3134745 | 0.18655778 | 0.23728976 | 0.78620241 | 0.43174892 |
| VPS27 | YNR006W | 1189.05506 | 0.05265533 | 0.06701843 | 0.78568435 | 0.43205243 |
| IAI11 | YBL059W | 223.956265 | 0.09209353 | 0.11742675 | 0.78426363 | 0.43288544 |
| YDL026W | YDL026W | 3.31352333 | 0.21188392 | 0.27015067 | 0.78431758 | 0.43285379 |
| SLD3 | YGL113W | 804.752052 | 0.05608032 | 0.07149881 | 0.7843532 | 0.43283289 |
| Xu_2009_SUTs31 | Xu_2009_SU | 35.0100678 | 0.18478256 | 0.23585535 | 0.78345715 | 0.43335871 |
| HSH155 | YMR288W | 585.394599 | 0.06486373 | 0.08292061 | 0.78223896 | 0.43407415 |
| tK(UUU)G1 | tK(UUU)G1 | 12.9550601 | 0.22600058 | 0.28896318 | 0.78210856 | 0.43415078 |
| YIR016W | YIR016W | 273.252829 | -0.0862884 | 0.11039785 | -0.7816134 | 0.4344418 |
| Xu_2009_CUTs14 | Xu_2009_CU | 12.0251321 | 0.22853365 | 0.29245643 | 0.78142801 | 0.4345508 |
| Xu_2009_SUTs69 | Xu_2009_SU | 74.0184726 | 0.1483736 | 0.18991031 | 0.78128252 | 0.43463635 |
| COG3 | YER157W | 843.833756 | 0.06200367 | 0.07947671 | 0.7801489 | 0.43530324 |
| Xu_2009_CUTs36 | Xu_2009_CU | 11.9348055 | 0.22792669 | 0.29228024 | 0.77982244 | 0.4354954 |
| CDA2 | YLR308W | 2.74538239 | 0.20096488 | 0.25769479 | 0.7798562 | 0.43547552 |
| NGG1 | YDR176W | 920.76035 | 0.05656973 | 0.07255278 | 0.77970451 | 0.43556483 |
| TRM82 | YDR165W | 879.060243 | -0.0588645 | 0.07559593 | -0.7786724 | 0.43617273 |
| Xu_2009_SUTs45 | Xu_2009_SU | 7.32436038 | -0.2308155 | 0.29663998 | -0.7780999 | 0.4365101 |
| YAR047C | YAR047C | 9.20804513 | 0.23035152 | 0.29609753 | 0.77795826 | 0.43659362 |
| MSG5 | YNL053W | 636.639858 | -0.0676091 | 0.0869367 | -0.7776823 | 0.43675632 |
| Xu_2009_CUTs66 | Xu_2009_CU | 6.27216594 | 0.22809722 | 0.29350499 | 0.77714939 | 0.43707064 |
| SCY1 | YGL083W | 433.093543 | -0.0734613 | 0.09457613 | -0.776742 | 0.43731098 |
| CUE4 | YML101C | 124.548325 | -0.1255389 | 0.16162693 | -0.7767199 | 0.43732402 |
| YDL114W-A | YDL114W-A | 6.55955413 | 0.22983334 | 0.29598393 | 0.77650615 | 0.43745019 |
| FRE4 | YNR060W | 112.182586 | 0.11780473 | 0.15190662 | 0.77550756 | 0.4380398 |
| Xu_2009_SUTs45 | Xu_2009_SU | 86.3774128 | -0.1341914 | 0.17307344 | -0.7753437 | 0.4381366 |

| | | | | | | |
|---|---|---|---|---|---|---|
| Xu_2009_SUTs79: | Xu_2009_SU | 6.10080793 | 0.22722162 | 0.29346372 | 0.77427499 | 0.43876819 |
| NAP1 | YKR048C | 3906.62307 | 0.0398705 | 0.05154063 | 0.7735742 | 0.43918263 |
| Xu_2009_SUTs73 | Xu_2009_SU | 14.705507 | -0.2215227 | 0.28677819 | -0.7724531 | 0.43984614 |
| YJL135W | YJL135W | 10.6370259 | 0.22644026 | 0.29323014 | 0.77222711 | 0.43997993 |
| NNT1 | YLR285W | 718.051731 | -0.0615324 | 0.07979732 | -0.7711082 | 0.44064282 |
| MIG3 | YER028C | 188.946896 | 0.10054498 | 0.13059265 | 0.76991301 | 0.4413515 |
| WSC3 | YOL105C | 428.556245 | 0.08057912 | 0.10472493 | 0.76943588 | 0.4416346 |
| YKL075C | YKL075C | 291.225608 | -0.0836921 | 0.10880743 | -0.7691762 | 0.4417887 |
| YLR464W | YLR464W | 1.95853437 | 0.17243398 | 0.22430317 | 0.76875409 | 0.44203931 |
| Xu_2009_SUTs60 | Xu_2009_SU | 15.7459926 | -0.2155724 | 0.28058484 | -0.7682966 | 0.44231099 |
| PHO92 | YDR374C | 15.7925762 | 0.21611145 | 0.28147029 | 0.76779487 | 0.44260907 |
| Xu_2009_SUTs82 | Xu_2009_SU | 61.4472604 | 0.14492512 | 0.18879115 | 0.76764786 | 0.44269642 |
| DSN1 | YIR010W | 929.71293 | -0.0549025 | 0.07163077 | -0.7664646 | 0.4433999 |
| YGR079W | YGR079W | 2893.34402 | 0.05464809 | 0.07136312 | 0.76577496 | 0.44381022 |
| YER088C-A | YER088C-A | 35.6712704 | -0.1759458 | 0.23025072 | -0.7641486 | 0.44477867 |
| tY(GUA)J2/SUP4 | tY(GUA)J2 | 2.86881303 | -0.1916519 | 0.2510804 | -0.7633087 | 0.4452793 |
| YGR139W | YGR139W | 1.69848098 | 0.17343868 | 0.22732181 | 0.7629654 | 0.44548403 |
| RPB2 | YOR151C | 4075.1523 | -0.0394911 | 0.05176753 | -0.7628543 | 0.44555032 |
| YJR084W | YJR084W | 331.936187 | 0.07751967 | 0.1016494 | 0.76261809 | 0.44569119 |
| PLB3 | YOL011W | 3114.15547 | 0.04239227 | 0.05559593 | 0.76250661 | 0.4457577 |
| YOR376W | YOR376W | 1.09579181 | 0.1494729 | 0.19601986 | 0.7625396 | 0.44573802 |
| Xu_2009_SUTs77 | Xu_2009_SU | 15.3149861 | 0.21950268 | 0.28791549 | 0.7623858 | 0.44582978 |
| Xu_2009_SUTs37 | Xu_2009_SU | 34.2689784 | 0.1736047 | 0.22784545 | 0.76194061 | 0.44609545 |
| YBL100W-C | YBL100W-C | 2.20501063 | -0.1848092 | 0.24255912 | -0.7619141 | 0.44611127 |
| MIC19 | YFR011C | 433.923275 | 0.07285289 | 0.09562246 | 0.76188049 | 0.44613134 |
| YGR016W | YGR016W | 94.7386363 | -0.1231349 | 0.16161709 | -0.7618925 | 0.44612414 |
| tF(GAA)P2 | tF(GAA)P2 | 2.54958019 | 0.19437029 | 0.25515938 | 0.76176032 | 0.44620307 |
| AHA1 | YDR214W | 4973.8559 | -0.0492901 | 0.06473908 | -0.7613657 | 0.44643867 |
| YMR030W-A | YMR030W-A | 3.34860551 | -0.1985435 | 0.2611097 | -0.7603836 | 0.44702534 |
| tV(AAC)G1 | tV(AAC)G1 | 3.62145965 | 0.20517538 | 0.26991141 | 0.76015823 | 0.44716001 |
| INH1 | YDL181W | 644.669942 | -0.0659203 | 0.08676571 | -0.7597506 | 0.4474037 |
| Xu_2009_SUTs34 | Xu_2009_SU | 44.5824131 | -0.1628233 | 0.2145742 | -0.7588204 | 0.44796003 |
| SDT1 | YGL224C | 457.734261 | 0.06923549 | 0.09121789 | 0.75901217 | 0.44784527 |
| MNL1 | YHR204W | 866.994566 | 0.05504217 | 0.07253506 | 0.75883541 | 0.44795102 |
| LRO1 | YNR008W | 1037.48536 | 0.0541409 | 0.0713199 | 0.7591275 | 0.44777629 |
| UAF30 | YOR295W | 425.836767 | -0.0700868 | 0.0923106 | -0.7592501 | 0.44770297 |
| ELP3 | YPL086C | 761.526993 | -0.0563515 | 0.07430638 | -0.7583663 | 0.44823171 |
| URM1 | YIL008W | 406.395882 | -0.0704004 | 0.09302367 | -0.7568004 | 0.44916943 |
| FPR4 | YLR449W | 3830.66027 | 0.04790408 | 0.06334205 | 0.75627604 | 0.44948371 |
| YOR203W | YOR203W | 45.7731023 | -0.1614697 | 0.21360659 | -0.7559212 | 0.44969647 |
| ALG13 | YGL047W | 293.191633 | 0.07704361 | 0.10205082 | 0.7549534 | 0.45027694 |

| | | | | | | |
|---|---|---|---|---|---|---|
| CRC1 | YOR100C | 78.3008743 | -0.1341351 | 0.17769515 | -0.7548605 | 0.4503327 |
| MUS81 | YDR386W | 256.627765 | -0.0897028 | 0.11890921 | -0.7543808 | 0.45062057 |
| STE13 | YOR219C | 1437.07082 | 0.0488026 | 0.06470721 | 0.75420656 | 0.4507252 |
| Xu_2009_SUTs59 | Xu_2009_SU | 43.1365912 | 0.17002394 | 0.22557067 | 0.75375022 | 0.45099922 |
| Xu_2009_SUTs64 | Xu_2009_SU | 29.7391062 | -0.1803964 | 0.23945004 | -0.7533782 | 0.45122269 |
| Xu_2009_CUTs59 | Xu_2009_CU | 5.67907016 | 0.21940915 | 0.29152621 | 0.75262237 | 0.45167687 |
| HST4 | YDR191W | 309.877567 | -0.0758106 | 0.10074948 | -0.7524664 | 0.45177062 |
| YDL062W | YDL062W | 1.15513756 | 0.15132828 | 0.2013378 | 0.75161389 | 0.45228329 |
| YEA4 | YEL004W | 175.790154 | -0.1066805 | 0.14195063 | -0.7515322 | 0.45233242 |
| Xu_2009_SUTs45 | Xu_2009_SU | 5.69455161 | 0.22082488 | 0.29395577 | 0.75121806 | 0.45252143 |
| IRC14 | YOR135C | 1.49981715 | -0.1591762 | 0.21229503 | -0.7497876 | 0.45338264 |
| RTT105 | YER104W | 115.413009 | 0.11529642 | 0.15407224 | 0.74832698 | 0.45426296 |
| tK(CUU)K | tK(CUU)K | 10.9210561 | -0.2214686 | 0.29621858 | -0.7476526 | 0.45466975 |
| YGL235W | YGL235W | 11.2865506 | -0.2200335 | 0.2943347 | -0.7475621 | 0.45472432 |
| Xu_2009_SUTs20 | Xu_2009_SU | 21.2937834 | -0.1986962 | 0.266072 | -0.7467762 | 0.45519866 |
| GCD6 | YDR211W | 1810.29808 | 0.04356193 | 0.05834157 | 0.7466705 | 0.45526248 |
| ENA1 | YDR040C | 382.965977 | -0.0836488 | 0.11211555 | -0.7460948 | 0.45561016 |
| MTR10 | YOR160W | 723.87142 | 0.05673686 | 0.07610245 | 0.74553265 | 0.45594978 |
| RPC53 | YDL150W | 1346.04473 | 0.05339466 | 0.07171539 | 0.74453561 | 0.45655251 |
| ACK1 | YDL203C | 629.50318 | -0.0593145 | 0.07968007 | -0.7444083 | 0.45662948 |
| SNN1 | YNL086W | 175.911129 | 0.0934603 | 0.1255612 | 0.74434064 | 0.45667042 |
| DBP6 | YNR038W | 1464.93919 | 0.0590944 | 0.07935988 | 0.74463819 | 0.45649048 |
| SPS1 | YDR523C | 228.995326 | 0.10727273 | 0.14419428 | 0.74394582 | 0.45690925 |
| YKR005C | YKR005C | 152.357678 | 0.10367574 | 0.13939025 | 0.74378045 | 0.45700931 |
| YKL165C-A | YKL165C-A | 28.1142905 | -0.1882307 | 0.25311058 | -0.7436699 | 0.45707623 |
| ECM27 | YJR106W | 421.419762 | -0.0731939 | 0.09844929 | -0.7434675 | 0.45719869 |
| ECM21 | YBL101C | 1373.9692 | 0.0501628 | 0.06751702 | 0.7429652 | 0.45750275 |
| RGI2 | YIL057C | 13.1513265 | -0.2154912 | 0.29018705 | -0.7425943 | 0.45772737 |
| YIR020W-A | YIR020W-A | 3.14289415 | 0.19872187 | 0.26759134 | 0.74263191 | 0.45770457 |
| MIP1 | YOR330C | 1438.80161 | 0.04765241 | 0.0641712 | 0.74258257 | 0.45773444 |
| SPP2 | YOR148C | 241.746905 | 0.08416049 | 0.11341589 | 0.74205203 | 0.45805581 |
| PHO80 | YOL001W | 387.135848 | 0.07187833 | 0.09688137 | 0.74192102 | 0.45813519 |
| Xu_2009_CUTs49 | Xu_2009_CU | 6.10579339 | 0.21663296 | 0.29208581 | 0.74167574 | 0.45828382 |
| ACO2 | YJL200C | 1121.11698 | -0.0491817 | 0.0663057 | -0.7417409 | 0.45824435 |
| RAI1 | YGL246C | 671.448911 | -0.0665279 | 0.08991045 | -0.7399353 | 0.45933927 |
| Xu_2009_SUTs70 | Xu_2009_SU | 19.1592176 | 0.2010131 | 0.27179851 | 0.73956659 | 0.45956302 |
| tV(AAC)M3 | tV(AAC)M3 | 11.9239121 | 0.21667996 | 0.29330849 | 0.73874427 | 0.4600623 |
| Xu_2009_SUTs35 | Xu_2009_SU | 19.8884313 | -0.1977993 | 0.26771952 | -0.7388305 | 0.46000995 |
| RUD3 | YOR216C | 926.539929 | 0.0534827 | 0.07240293 | 0.73868138 | 0.4601005 |
| GAL80 | YML051W | 1475.92785 | -0.0470671 | 0.0637949 | -0.7377877 | 0.46064346 |
| UTP9 | YHR196W | 1829.62598 | 0.04777764 | 0.06483359 | 0.73692726 | 0.46116659 |

| | | | | | | |
|---|---|---|---|---|---|---|
| YMR244W | YMR244W | 93.9572363 | 0.1262796 | 0.17136258 | 0.73691466 | 0.46117425 |
| OCT1_ | YKL134C | 613.89587 | 0.05923505 | 0.08048452 | 0.73598065 | 0.46174248 |
| Xu_2009_SUTs20! | Xu_2009_SU' | 165.749006 | 0.0960239 | 0.13055262 | 0.73551872 | 0.46202365 |
| YJR128W | YJR128W | 6.52875073 | -0.2169398 | 0.29496118 | -0.7354859 | 0.46204363 |
| SSP1 | YHR184W | 73.7241019 | -0.1379105 | 0.18784899 | -0.7341562 | 0.46285357 |
| YIL021C-A | YIL021C-A | 1.2659688 | 0.14752963 | 0.20124816 | 0.73307318 | 0.4635138 |
| PUF2 | YPR042C | 743.052011 | -0.0532484 | 0.07263169 | -0.7331289 | 0.46347984 |
| RPM1 | RPM1 | 11.1756239 | -0.2170189 | 0.296325 | -0.7323677 | 0.46394414 |
| YHL005C | YHL005C | 4.90217912 | -0.21154 | 0.28891051 | -0.7321991 | 0.46404702 |
| YLR235C | YLR235C | 4.19130975 | 0.20214635 | 0.27606969 | 0.73222943 | 0.46402855 |
| Xu_2009_CUTs25 | Xu_2009_CU | 10.7191728 | 0.21397477 | 0.29271459 | 0.73100139 | 0.4647783 |
| Xu_2009_SUTs41! | Xu_2009_SU' | 30.4291043 | -0.1767703 | 0.24185592 | -0.7308908 | 0.46484583 |
| HSH49 | YOR319W | 237.79742 | 0.08527606 | 0.11668621 | 0.73081529 | 0.46489199 |
| ATG1 | YGL180W | 741.739206 | 0.0574291 | 0.07863541 | 0.73032105 | 0.46519396 |
| ISC10 | YER180C | 381.184651 | -0.0681739 | 0.09340027 | -0.7299116 | 0.4654442 |
| PFK27 | YOL136C | 2825.05184 | -0.0472866 | 0.06478132 | -0.7299417 | 0.46542584 |
| RDN37-2 | RDN37-2 | 2.45904166 | -0.1781291 | 0.24434424 | -0.7290086 | 0.46599638 |
| YGL114W | YGL114W | 683.11806 | -0.0598807 | 0.08220717 | -0.7284125 | 0.46636109 |
| Xu_2009_CUTs89 | Xu_2009_CU | 25.9749682 | 0.18748045 | 0.25755795 | 0.7279156 | 0.46666525 |
| tL(UAA)D | tL(UAA)D | 2.77197902 | 0.18530298 | 0.25468724 | 0.72757072 | 0.46687641 |
| MTC3 | YGL226W | 225.500419 | 0.0947583 | 0.13043159 | 0.72649807 | 0.46753349 |
| Xu_2009_SUTs57( | Xu_2009_SU' | 9.55373503 | -0.2143695 | 0.29531274 | -0.7259068 | 0.46789591 |
| RRT7 | YLL030C | 1.85709068 | -0.1699714 | 0.23411987 | -0.7260016 | 0.4678378 |
| SNP1 | YIL061C | 341.766164 | 0.07367599 | 0.10151767 | 0.72574548 | 0.46799482 |
| Xu_2009_SUTs31! | Xu_2009_SU' | 293.387766 | 0.07964024 | 0.10986281 | 0.72490624 | 0.46850955 |
| PTK2 | YJR059W | 1808.77622 | -0.047737 | 0.06584334 | -0.7250088 | 0.4684466 |
| EFM4 | YIL064W | 705.702747 | 0.05676939 | 0.07844295 | 0.72370296 | 0.46924811 |
| YPR147C | YPR147C | 518.010403 | -0.0661273 | 0.0913642 | -0.723777 | 0.46920268 |
| BUD13 | YGL174W | 162.379866 | -0.0963517 | 0.13321584 | -0.7232751 | 0.46951086 |
| MOB1 | YIL106W | 891.708959 | 0.05756194 | 0.0795793 | 0.723328 | 0.4694784 |
| RTR2 | YDR066C | 84.5960768 | -0.1275387 | 0.17637988 | -0.7230907 | 0.46962413 |
| YJL182C | YJL182C | 2.26800147 | -0.1767392 | 0.24461576 | -0.7225178 | 0.4699762 |
| PAP2 | YOL115W | 603.01184 | -0.0595086 | 0.0824358 | -0.7218782 | 0.47036936 |
| tY(GUA)Q | tY(GUA)Q | 2.37516341 | -0.174847 | 0.24234383 | -0.7214832 | 0.47061224 |
| SNR68 | snR68 | 401.000649 | 0.07756553 | 0.1076794 | 0.72033768 | 0.47131711 |
| YKL069W | YKL069W | 340.963665 | -0.0840211 | 0.11685907 | -0.7189952 | 0.47214386 |
| SPO23 | YBR250W | 86.5652234 | 0.12086758 | 0.16814159 | 0.71884405 | 0.47223701 |
| YSH1 | YLR277C | 627.211155 | -0.0621178 | 0.08648421 | -0.7182556 | 0.47259969 |
| YAR053W | YAR053W | 4.75836395 | 0.20261508 | 0.28231853 | 0.71768253 | 0.47295305 |
| RPF2 | YKR081C | 1336.53854 | 0.059331 | 0.08284093 | 0.71620396 | 0.47386542 |
| Xu_2009_SUTs70! | Xu_2009_SU' | 1.47198601 | -0.1492231 | 0.20855061 | -0.7155246 | 0.47428496 |

| | | | | | | |
|---|---|---|---|---|---|---|
| RHO2 | YNL090W | 628.28846 | 0.06560863 | 0.09170699 | 0.71541584 | 0.47435213 |
| Xu_2009_CUTs57 | Xu_2009_CU | 80.8651988 | 0.12318444 | 0.17226775 | 0.71507544 | 0.47456243 |
| YER148W-A | YER148W-A | 4.62360914 | -0.2048394 | 0.28662199 | -0.7146676 | 0.47481449 |
| YLR123C | YLR123C | 16.8391086 | 0.20062718 | 0.28099275 | 0.71399414 | 0.47523081 |
| YHR138C | YHR138C | 1225.66595 | 0.05588241 | 0.07830404 | 0.71365929 | 0.47543789 |
| YCR045W-A | YCR045W-A | 99.0251832 | 0.11442347 | 0.16046481 | 0.71307516 | 0.47579925 |
| Xu_2009_SUTs17 | Xu_2009_SU | 40.1901422 | 0.16435852 | 0.23055321 | 0.71288758 | 0.47591532 |
| MTF2 | YDL044C | 300.08108 | 0.07573191 | 0.10623834 | 0.71284913 | 0.47593912 |
| VPS30 | YPL120W | 223.692938 | 0.08622438 | 0.12103734 | 0.7123784 | 0.47623049 |
| YIL105W-A | YIL105W-A | 2.13193044 | 0.17109864 | 0.2405581 | 0.71125702 | 0.47692498 |
| HRP1 | YOL123W | 3198.90411 | -0.0392429 | 0.05524736 | -0.7103133 | 0.47750986 |
| TRI1 | YMR233W | 165.756541 | -0.096409 | 0.13599759 | -0.7089024 | 0.47838503 |
| tE(UUC)E3 | tE(UUC)E3 | 1.86600278 | -0.1604642 | 0.22647244 | -0.7085373 | 0.47861168 |
| SIN4 | YNL236W | 1195.10459 | -0.046961 | 0.06639607 | -0.7072854 | 0.47938916 |
| PBP2 | YBR233W | 583.586207 | -0.062131 | 0.08788333 | -0.7069709 | 0.47958459 |
| Xu_2009_SUTs41 | Xu_2009_SU | 118.629438 | -0.1038284 | 0.14691498 | -0.7067245 | 0.4797377 |
| Xu_2009_SUTs15 | Xu_2009_SU | 106.100835 | 0.12374741 | 0.17523938 | 0.7061621 | 0.48008734 |
| SMM1 | YNR015W | 547.956432 | 0.06364529 | 0.09018202 | 0.70574257 | 0.48034824 |
| Xu_2009_CUTs89 | Xu_2009_CU | 4.99987 | 0.20372597 | 0.28884228 | 0.70531907 | 0.4806117 |
| NCR1 | YPL006W | 1253.33448 | -0.0492688 | 0.06986095 | -0.7052411 | 0.48066019 |
| FIG1 | YBR040W | 37.9064571 | -0.1631638 | 0.231576 | -0.7045801 | 0.48107159 |
| YHR218W | YHR218W | 5.13832141 | 0.20513459 | 0.29122991 | 0.70437337 | 0.48120029 |
| MAK10 | YEL053C | 899.801406 | 0.05694452 | 0.08087222 | 0.70412956 | 0.48135209 |
| Xu_2009_SUTs78 | Xu_2009_SU | 33.992525 | -0.1658729 | 0.23584037 | -0.7033268 | 0.4818521 |
| YAE1 | YJR067C | 123.564354 | 0.10401058 | 0.14809891 | 0.70230485 | 0.48248907 |
| Xu_2009_CUTs79 | Xu_2009_CU | 1.50624713 | -0.1451593 | 0.20677484 | -0.7020161 | 0.4826691 |
| YHL026C | YHL026C | 477.952668 | -0.0623071 | 0.08878212 | -0.7017976 | 0.48280539 |
| Xu_2009_CUTs85 | Xu_2009_CU | 8.30327367 | 0.20775221 | 0.29651112 | 0.70065573 | 0.48351789 |
| Xu_2009_CUTs91 | Xu_2009_CU | 6.8029604 | 0.20714594 | 0.29573931 | 0.70043424 | 0.48365616 |
| YBL094C | YBL094C | 99.3352301 | 0.11321503 | 0.16172293 | 0.7000555 | 0.48389264 |
| Xu_2009_SUTs61 | Xu_2009_SU | 10.4325285 | -0.2062161 | 0.29482768 | -0.6994462 | 0.48427322 |
| tQ(UUG)D2 | tQ(UUG)D2 | 1.71881954 | -0.1612392 | 0.23061558 | -0.6991688 | 0.48444653 |
| OPI8 | YKR035C | 30.5674436 | 0.16836697 | 0.24080085 | 0.69919589 | 0.48442962 |
| CWC23 | YGL128C | 384.809868 | 0.06727126 | 0.09624861 | 0.69893223 | 0.48459438 |
| Xu_2009_CUTs73 | Xu_2009_CU | 5.73575602 | -0.2022642 | 0.28975717 | -0.6980473 | 0.48514761 |
| RVS161 | YCR009C | 2264.77887 | -0.0437852 | 0.062765 | -0.697606 | 0.48542365 |
| CAF130 | YGR134W | 621.560377 | -0.0642949 | 0.09222365 | -0.6971633 | 0.48570063 |
| tL(UAA)L | tL(UAA)L | 3.52348158 | -0.1905651 | 0.27394825 | -0.6956246 | 0.48666396 |
| YDR415C | YDR415C | 457.803207 | -0.0625769 | 0.08996906 | -0.6955377 | 0.48671837 |
| Xu_2009_SUTs27 | Xu_2009_SU | 175.398019 | 0.09201847 | 0.13232666 | 0.69538875 | 0.48681171 |
| Xu_2009_CUTs73 | Xu_2009_CU | 2.8933196 | 0.17716532 | 0.25496602 | 0.69485853 | 0.48714396 |

| | | | | | | |
|---|---|---|---|---|---|---|
| YLR286W-A | YLR286W-A | 13.9055624 | 0.19766855 | 0.28457164 | 0.69461788 | 0.4872948 |
| DTR1 | YBR180W | 5.14583238 | -0.1986832 | 0.28618858 | -0.6942387 | 0.48753254 |
| POP4 | YBR257W | 401.520027 | 0.06806457 | 0.09807013 | 0.69403974 | 0.48765729 |
| NOP56 | YLR197W | 5726.52447 | -0.0443755 | 0.06396807 | -0.6937134 | 0.48786195 |
| Xu_2009_SUTs62! | Xu_2009_SU | 1.96609554 | -0.1618867 | 0.23352063 | -0.6932437 | 0.48815665 |
| Xu_2009_CUTs57 | Xu_2009_CU | 9.75184928 | -0.2046428 | 0.2953124 | -0.6929707 | 0.48832796 |
| Xu_2009_SUTs81 | Xu_2009_SU | 10.481335 | 0.20530418 | 0.29627346 | 0.69295503 | 0.48833777 |
| Xu_2009_SUTs44 | Xu_2009_SU | 5.7589542 | 0.20242909 | 0.29226195 | 0.69262896 | 0.48854243 |
| SOV1 | YMR066W | 604.375573 | -0.0584176 | 0.08439425 | -0.6921989 | 0.48881242 |
| RFC4 | YOL094C | 231.809867 | 0.07918532 | 0.11438847 | 0.69224913 | 0.48878089 |
| PXP1 | YEL020C | 470.496063 | -0.0618478 | 0.089435 | -0.6915393 | 0.48922666 |
| ISW1 | YBR245C | 1405.13609 | 0.04567387 | 0.06610289 | 0.69095117 | 0.48959622 |
| YKL102C | YKL102C | 7.8301173 | 0.20449557 | 0.29646628 | 0.68977682 | 0.49033455 |
| YLR434C | YLR434C | 3.38389009 | 0.187885 | 0.27244852 | 0.68961653 | 0.49043537 |
| DAD3 | YBR233W-A | 233.08962 | -0.0796396 | 0.11585239 | -0.6874228 | 0.49181631 |
| Xu_2009_CUTs19 | Xu_2009_CU | 0.84502763 | 0.11565295 | 0.16842834 | 0.6866597 | 0.4922972 |
| Xu_2009_SUTs51 | Xu_2009_SU | 72.7511356 | -0.123565 | 0.17993061 | -0.686737 | 0.49224847 |
| Xu_2009_CUTs64 | Xu_2009_CU | 34.0983376 | 0.16386276 | 0.23873081 | 0.68639132 | 0.49246638 |
| YOL019W-A | YOL019W-A | 1.70667744 | -0.1530816 | 0.22324348 | -0.6857159 | 0.4928923 |
| YPL162C | YPL162C | 280.566236 | -0.0760865 | 0.11095281 | -0.6857556 | 0.49286724 |
| DOS2 | YDR068W | 611.548567 | 0.06288453 | 0.09174861 | 0.68540033 | 0.49309134 |
| BBC1 | YJL020C | 3677.37735 | -0.0434266 | 0.06336829 | -0.6853049 | 0.49315156 |
| tS(GCU)O | tS(GCU)O | 16.2041562 | -0.1939201 | 0.28324502 | -0.6846374 | 0.49357277 |
| Xu_2009_SUTs39 | Xu_2009_SU | 12.2073136 | -0.1998281 | 0.29225901 | -0.6837363 | 0.49414168 |
| FZO1 | YBR179C | 715.710233 | 0.05213646 | 0.07628613 | 0.68343301 | 0.49433327 |
| IGO1 | YNL157W | 1071.56379 | -0.0445609 | 0.06522329 | -0.6832058 | 0.4944768 |
| RTA1 | YGR213C | 39.6083716 | 0.15608307 | 0.22856026 | 0.6828968 | 0.49467205 |
| VPS29 | YHR012W | 728.403703 | 0.05293374 | 0.07759532 | 0.68217692 | 0.49512709 |
| ARG2 | YJL071W | 373.616982 | -0.0691319 | 0.10142133 | -0.681631 | 0.49547228 |
| ERF2 | YLR246W | 481.548803 | -0.0628797 | 0.09227408 | -0.6814444 | 0.49559032 |
| RPN5 | YDL147W | 3947.12004 | 0.03514686 | 0.05161949 | 0.68088338 | 0.49594528 |
| tD(GUC)I2 | tD(GUC)I2 | 13.4107952 | 0.19732567 | 0.28998796 | 0.68046158 | 0.49621224 |
| Xu_2009_CUTs67 | Xu_2009_CU | 27.9943512 | 0.17663952 | 0.25964775 | 0.68030447 | 0.4963117 |
| UIP5 | YKR044W | 572.362621 | -0.0658052 | 0.09675317 | -0.6801348 | 0.49641913 |
| YNL067W-A | YNL067W-A | 1.06903534 | -0.1303952 | 0.19187498 | -0.6795842 | 0.49676777 |
| BUD28 | YLR062C | 2.79355058 | 0.17072227 | 0.2513175 | 0.67930913 | 0.49694202 |
| YER084W | YER084W | 34.142709 | -0.1608765 | 0.23687127 | -0.6791726 | 0.49702849 |
| MEC3 | YLR288C | 378.632388 | 0.0660124 | 0.09720892 | 0.67907762 | 0.49708868 |
| PAU7 | YAR020C | 39.830478 | -0.1492056 | 0.22001996 | -0.6781459 | 0.49767917 |
| COX9 | YDL067C | 893.293721 | -0.0582181 | 0.0858452 | -0.6781748 | 0.49766084 |
| YOL036W | YOL036W | 2195.15713 | 0.03841163 | 0.05664085 | 0.67816137 | 0.49766938 |

| | | | | | | |
|---|---|---|---|---|---|---|
| Xu_2009_CUTs12 | Xu_2009_CU | 5.17606439 | -0.1966881 | 0.2903794 | -0.6773488 | 0.4981847 |
| Xu_2009_CUTs33 | Xu_2009_CU | 16.3212893 | 0.18617399 | 0.27481707 | 0.67744696 | 0.49812241 |
| FLR1 | YBR008C | 256.838051 | -0.0735342 | 0.10878649 | -0.6759495 | 0.49907274 |
| Xu_2009_SUTs71 | Xu_2009_SU | 95.1755385 | -0.1107153 | 0.16391925 | -0.6754256 | 0.4994054 |
| YER158W-A | YER158W-A | 18.3863535 | 0.18569847 | 0.27516128 | 0.67487136 | 0.4997575 |
| YGL007C-A | YGL007C-A | 3.71923529 | -0.1822414 | 0.27024933 | -0.6743454 | 0.50009173 |
| ADH6 | YMR318C | 2961.94806 | -0.040899 | 0.06067353 | -0.6740833 | 0.50025837 |
| RPN14 | YGL004C | 222.488909 | -0.0820844 | 0.12181387 | -0.6738511 | 0.50040596 |
| Xu_2009_CUTs12 | Xu_2009_CU | 9.83634689 | 0.19891901 | 0.29550593 | 0.67314726 | 0.50085361 |
| Xu_2009_CUTs28 | Xu_2009_CU | 37.5649394 | 0.15323655 | 0.2277022 | 0.67296913 | 0.50096693 |
| SLX1 | YBR228W | 249.921032 | -0.076988 | 0.1144432 | -0.6727182 | 0.5011266 |
| LIN1 | YHR156C | 154.980571 | 0.09068296 | 0.13501118 | 0.67167002 | 0.50179379 |
| HAP5 | YOR358W | 322.205546 | -0.069248 | 0.10309747 | -0.6716749 | 0.50179068 |
| PRK1 | YIL095W | 623.852264 | 0.05210175 | 0.07760576 | 0.67136452 | 0.50198834 |
| FPK1 | YNR047W | 1941.6386 | 0.03835355 | 0.05714195 | 0.6711977 | 0.50209459 |
| Xu_2009_CUTs81 | Xu_2009_CU | 16.0468893 | 0.18916976 | 0.28228177 | 0.67014515 | 0.50276526 |
| Xu_2009_SUTs39 | Xu_2009_SU | 58.2721343 | 0.1325732 | 0.19807209 | 0.6693179 | 0.50329271 |
| Xu_2009_CUTs92 | Xu_2009_CU | 0.84044401 | 0.11605068 | 0.17358351 | 0.66855819 | 0.50377735 |
| Xu_2009_SUTs27 | Xu_2009_SU | 0.69917978 | 0.10756996 | 0.16088088 | 0.66863111 | 0.50373082 |
| Xu_2009_CUTs56 | Xu_2009_CU | 10.4266594 | -0.1978981 | 0.29614291 | -0.668252 | 0.50397277 |
| tF(GAA)Q | tF(GAA)Q | 1.03616762 | -0.1204236 | 0.18043989 | -0.6673889 | 0.50452374 |
| Xu_2009_SUTs78 | Xu_2009_SU | 1.23061856 | -0.1306923 | 0.19588386 | -0.6671928 | 0.50464899 |
| GAL10 | YBR019C | 37.1080725 | 0.15406489 | 0.23079294 | 0.66754595 | 0.50442347 |
| CRP1 | YHR146W | 1082.20772 | -0.050154 | 0.07518073 | -0.6671124 | 0.50470033 |
| YIR021W-A | YIR021W-A | 1.13269762 | -0.1311294 | 0.19653006 | -0.667223 | 0.50462971 |
| YLR122C | YLR122C | 21.3225562 | 0.18176753 | 0.27236223 | 0.66737422 | 0.50453313 |
| tN(GUU)G | tN(GUU)G | 14.9667092 | 0.18889218 | 0.2836078 | 0.66603309 | 0.50538995 |
| YMR119W-A | YMR119W-A | 5.57484014 | 0.1947483 | 0.29270358 | 0.66534308 | 0.50583108 |
| NTR2 | YKR022C | 264.139152 | 0.07834296 | 0.11780187 | 0.66504009 | 0.50602485 |
| YPT52 | YKR014C | 1447.18385 | 0.0542412 | 0.08162366 | 0.66452788 | 0.50635251 |
| DBP3 | YGL078C | 2679.31873 | 0.04196429 | 0.0631804 | 0.66419799 | 0.5065636 |
| Xu_2009_SUTs50 | Xu_2009_SU | 0.92511789 | -0.1187572 | 0.17883336 | -0.6640662 | 0.50664796 |
| YCR018C-A | YCR018C-A | 29.4053563 | -0.1610328 | 0.24271255 | -0.6634713 | 0.50702874 |
| NAR1 | YNL240C | 525.638282 | 0.06384682 | 0.09652785 | 0.66143423 | 0.50833388 |
| PRP11 | YDL043C | 242.043819 | -0.0820007 | 0.12426176 | -0.6599032 | 0.50931593 |
| tF(GAA)H2 | tF(GAA)H2 | 8.86594394 | -0.1956395 | 0.29655519 | -0.6597069 | 0.50944196 |
| Xu_2009_SUTs19 | Xu_2009_SU | 18.7942234 | 0.17778132 | 0.26959724 | 0.659433 | 0.50961776 |
| YDR187C | YDR187C | 4.71856194 | 0.18750315 | 0.28436426 | 0.65937663 | 0.50965395 |
| YLR282C | YLR282C | 1.1077254 | 0.12052421 | 0.18308405 | 0.65829992 | 0.51034543 |
| PEX35 | YGR168C | 191.509717 | -0.0845037 | 0.12851319 | -0.6575491 | 0.51082789 |
| CSC1 | YLR241W | 1552.50167 | 0.04063154 | 0.06180114 | 0.65745608 | 0.5108877 |

| | | | | | | |
|---|---|---|---|---|---|---|
| tI(AAU)E2 | tI(AAU)E2 | 11.7389124 | 0.19107094 | 0.29078416 | 0.65708852 | 0.511124 |
| CEG1 | YGL130W | 697.046926 | 0.05149609 | 0.07834543 | 0.65729546 | 0.51099095 |
| AIR1 | YIL079C | 750.2855 | -0.0518846 | 0.07896629 | -0.6570474 | 0.51115042 |
| YLR326W | YLR326W | 415.059615 | 0.06487805 | 0.09874828 | 0.65700438 | 0.5111781 |
| COS1 | YNL336W | 242.6302 | 0.07217055 | 0.10990389 | 0.6566697 | 0.51139332 |
| DBP9 | YLR276C | 2196.22681 | 0.04755201 | 0.07252267 | 0.65568475 | 0.51202699 |
| YGL159W | YGL159W | 235.866435 | 0.07313741 | 0.11158 | 0.65547062 | 0.51216479 |
| FAR7 | YFR008W | 114.006032 | -0.1034807 | 0.15797248 | -0.655055 | 0.51243236 |
| Xu_2009_SUTs58 | Xu_2009_SU | 24.1026146 | -0.1698577 | 0.25941357 | -0.6547758 | 0.51261209 |
| MED6 | YHR058C | 412.200404 | -0.0649603 | 0.09922962 | -0.6546466 | 0.51269533 |
| YOR218C | YOR218C | 9.00059391 | -0.19309 | 0.29507054 | -0.6543858 | 0.51286328 |
| YPR078C | YPR078C | 55.6934577 | -0.1310577 | 0.20024897 | -0.6544736 | 0.51280674 |
| ADY2 | YCR010C | 22.116026 | -0.1697498 | 0.25948124 | -0.654189 | 0.51299007 |
| YCL041C | YCL041C | 5.05313209 | 0.18982328 | 0.29045065 | 0.65354745 | 0.51340341 |
| YCR102W-A | YCR102W-A | 5.27267447 | 0.19056209 | 0.29203065 | 0.65254139 | 0.51405198 |
| CHL4 | YDR254W | 183.399862 | -0.0795231 | 0.12186542 | -0.6525486 | 0.51404732 |
| SAC7 | YDR389W | 2051.6057 | -0.0403192 | 0.06180595 | -0.6523511 | 0.51417469 |
| YDR048C | YDR048C | 4.84571051 | 0.18567388 | 0.28468436 | 0.65220962 | 0.51426596 |
| YGR237C | YGR237C | 727.444609 | 0.05471968 | 0.08408394 | 0.65077446 | 0.51519209 |
| SMA1 | YPL027W | 17.8542164 | 0.18203224 | 0.27968577 | 0.65084554 | 0.5151462 |
| YBR174C | YBR174C | 2.29371503 | 0.16206765 | 0.24920397 | 0.65034137 | 0.51547174 |
| YHR070C-A | YHR070C-A | 0.93508073 | -0.1161181 | 0.17915915 | -0.6481285 | 0.51690186 |
| Xu_2009_SUTs21 | Xu_2009_SU | 31.5897696 | -0.1596963 | 0.24651183 | -0.6478243 | 0.51709863 |
| SFC1 | YJR095W | 113.898649 | 0.1009175 | 0.15590145 | 0.64731599 | 0.51742745 |
| YBL073W | YBL073W | 0.85754191 | -0.1059464 | 0.16384531 | -0.6466248 | 0.51787478 |
| SWF1 | YDR126W | 249.393508 | -0.0731381 | 0.11323467 | -0.6458982 | 0.51834526 |
| YKL111C | YKL111C | 0.92351311 | 0.114274 | 0.17694495 | 0.64581672 | 0.51839806 |
| Xu_2009_SUTs19 | Xu_2009_SU | 7.58905508 | 0.19138921 | 0.29645264 | 0.64559795 | 0.51853977 |
| CRT10 | YOL063C | 763.66461 | -0.0507065 | 0.07861674 | -0.644984 | 0.51893754 |
| tE(UUC)B | tE(UUC)B | 1.14377518 | -0.1234199 | 0.19164596 | -0.6439992 | 0.51957593 |
| Xu_2009_CUTs66 | Xu_2009_CU | 27.9324588 | -0.1566377 | 0.24341447 | -0.6435019 | 0.51989849 |
| DAM1 | YGR113W | 444.19913 | -0.0586354 | 0.09110575 | -0.6435973 | 0.51983659 |
| tR(CCG)L/TRR4 | tR(CCG)L | 18.5370024 | -0.1796131 | 0.27921921 | -0.6432691 | 0.52004948 |
| Xu_2009_CUTs44 | Xu_2009_CU | 1.77669541 | -0.1441561 | 0.22433139 | -0.6426035 | 0.52048143 |
| Xu_2009_CUTs16 | Xu_2009_CU | 7.88805815 | -0.1895169 | 0.29518485 | -0.6420279 | 0.52085505 |
| YER107W-A | YER107W-A | 0.80207957 | 0.11106247 | 0.173102 | 0.64160132 | 0.52113207 |
| AVT4 | YNL101W | 1489.23505 | -0.0458343 | 0.07145823 | -0.6414137 | 0.52125392 |
| YFL015W-A | YFL015W-A | 1.83523719 | -0.1495595 | 0.23323303 | -0.641245 | 0.52136351 |
| Xu_2009_CUTs72 | Xu_2009_CU | 2.54595167 | 0.15505975 | 0.24198747 | 0.64077595 | 0.52166826 |
| Xu_2009_SUTs32 | Xu_2009_SU | 63.7951793 | -0.1248952 | 0.19506335 | -0.6402799 | 0.52199061 |
| PRM6 | YML047C | 5.21732418 | 0.18507591 | 0.28955179 | 0.63918069 | 0.52270539 |

| | | | | | | |
|---|---|---|---|---|---|---|
| SED4 | YCR067C | 1481.1786 | 0.03825148 | 0.05985576 | 0.63906101 | 0.52278324 |
| Xu_2009_SUTs57 | Xu_2009_SU | 144.84106 | 0.09187685 | 0.14391877 | 0.63839381 | 0.52321736 |
| RAD17 | YOR368W | 234.56865 | -0.0713782 | 0.11187292 | -0.6380293 | 0.52345458 |
| DPC7 | YKL065W-A | 16.7228057 | -0.1798608 | 0.28218017 | -0.637397 | 0.52386629 |
| SRP21 | YKL122C | 1617.73034 | 0.04698943 | 0.07379896 | 0.63672217 | 0.52430583 |
| YMR084W | YMR084W | 45.6250071 | 0.14146378 | 0.22213818 | 0.63682783 | 0.52423699 |
| PCT1 | YGR202C | 1254.21704 | 0.04584964 | 0.07208561 | 0.63604423 | 0.52474759 |
| YKL083W | YKL083W | 0.82789459 | -0.1050883 | 0.16519758 | -0.6361372 | 0.52468702 |
| Xu_2009_CUTs83 | Xu_2009_CU | 3.58728546 | 0.1739036 | 0.27354834 | 0.63573261 | 0.52495072 |
| YNL115C | YNL115C | 951.963183 | 0.05768441 | 0.09084199 | 0.63499726 | 0.5254302 |
| YDR230W | YDR230W | 3.76704702 | -0.1750262 | 0.27576824 | -0.634686 | 0.52563323 |
| WAR1 | YML076C | 1134.7989 | 0.04574967 | 0.07223567 | 0.63333901 | 0.52651228 |
| Xu_2009_SUTs54 | Xu_2009_SU | 48.9215161 | -0.1332028 | 0.21037241 | -0.6331761 | 0.52661865 |
| tL(CAA)A/SUP56 | tL(CAA)A | 7.27903003 | -0.1868267 | 0.2953269 | -0.6326099 | 0.52698842 |
| YAL047W-A | YAL047W-A | 4.33821692 | -0.1811881 | 0.28654799 | -0.6323133 | 0.52718216 |
| Xu_2009_SUTs54 | Xu_2009_SU | 44.3629228 | 0.13274434 | 0.2103599 | 0.63103441 | 0.52801803 |
| BRE1 | YDL074C | 1371.55121 | 0.03993446 | 0.06339451 | 0.62993566 | 0.52873668 |
| ART5 | YGR068C | 319.749199 | 0.06528317 | 0.10362316 | 0.63000552 | 0.52869097 |
| CIA2 | YHR122W | 276.426415 | -0.0668528 | 0.10614212 | -0.6298419 | 0.52879804 |
| RAV1 | YJR033C | 1010.13286 | -0.0448782 | 0.07122903 | -0.6300545 | 0.52865894 |
| SPS2 | YDR522C | 8.20246692 | -0.1867971 | 0.29663406 | -0.6297223 | 0.52887628 |
| YJL027C | YJL027C | 4.46572187 | -0.178128 | 0.28344887 | -0.6284306 | 0.52972187 |
| TDA11 | YHR159W | 325.578481 | 0.06595724 | 0.10513242 | 0.62737305 | 0.53041473 |
| STE18 | YJR086W | 121.851856 | -0.0985844 | 0.15717901 | -0.6272112 | 0.53052083 |
| YPL276W | YPL276W | 2.58148754 | -0.1599174 | 0.25500261 | -0.6271208 | 0.53058004 |
| Xu_2009_CUTs23 | Xu_2009_CU | 6.70001171 | 0.18501116 | 0.29516388 | 0.62680828 | 0.53078491 |
| tN(GUU)N1 | tN(GUU)N1 | 2.77398424 | -0.1654203 | 0.26405112 | -0.6264706 | 0.53100631 |
| YPR197C | YPR197C | 6.8238157 | -0.1822638 | 0.29116043 | -0.6259911 | 0.53132079 |
| SWP82 | YFL049W | 717.45022 | 0.06025636 | 0.09635862 | 0.62533443 | 0.53175159 |
| Xu_2009_CUTs15 | Xu_2009_CU | 4.10981644 | 0.17515931 | 0.28044015 | 0.62458713 | 0.53224207 |
| PDC5 | YLR134W | 381.911746 | -0.0640864 | 0.10282033 | -0.6232855 | 0.5330969 |
| PAU19 | YMR325W | 4.23685799 | 0.1745082 | 0.28016901 | 0.6228676 | 0.53337153 |
| Xu_2009_SUTs14 | Xu_2009_SU | 30.627651 | -0.1507669 | 0.24214754 | -0.6226242 | 0.53353152 |
| tI(AAU)G | tI(AAU)G | 46.468222 | -0.135169 | 0.2174457 | -0.6216216 | 0.53419069 |
| Xu_2009_CUTs49 | Xu_2009_CU | 19.0598121 | 0.16687976 | 0.2684149 | 0.62172316 | 0.53412392 |
| Xu_2009_SUTs51 | Xu_2009_SU | 6.01170458 | -0.1783751 | 0.28695823 | -0.6216065 | 0.53420062 |
| Xu_2009_CUTs52 | Xu_2009_CU | 14.9146942 | 0.17674695 | 0.2844132 | 0.62144424 | 0.53430737 |
| SWM1 | YDR260C | 442.790059 | -0.0578072 | 0.09311339 | -0.620826 | 0.53471414 |
| tA(UGC)A/TGA1 | tA(UGC)A | 3.52310211 | 0.16410778 | 0.26449221 | 0.62046355 | 0.53495265 |
| COS111 | YBR203W | 671.431976 | 0.05107863 | 0.08236862 | 0.62012239 | 0.53517721 |
| Xu_2009_CUTs44 | Xu_2009_CU | 22.3135948 | 0.16295203 | 0.26300979 | 0.61956639 | 0.5355433 |

| | | | | | | |
|---|---|---|---|---|---|---|
| AKL1 | YBR059C | 2411.61822 | 0.04275272 | 0.06903335 | 0.61930528 | 0.53571527 |
| YJL152W | YJL152W | 12.8852885 | 0.181145 | 0.29269514 | 0.61888628 | 0.53599128 |
| BI4 | Q0120 | 37.0516078 | -0.1817427 | 0.29375467 | -0.6186888 | 0.53612136 |
| YRF1-6 | YNL339C | 1.21017087 | 0.12274196 | 0.1984573 | 0.61848044 | 0.53625869 |
| YDL159C-B | YDL159C-B | 1.8743116 | 0.14163635 | 0.22946717 | 0.61724015 | 0.53707634 |
| Xu_2009_SUTs46! | Xu_2009_SU | 22.4136549 | 0.16231884 | 0.26333037 | 0.6164076 | 0.53762554 |
| GRX3 | YDR098C | 930.880268 | -0.050671 | 0.08221526 | -0.616321 | 0.53768269 |
| PLM2 | YDR501W | 144.854038 | -0.0868274 | 0.14095675 | -0.6159863 | 0.53790357 |
| YLR030W | YLR030W | 159.787185 | 0.08298961 | 0.13481877 | 0.61556418 | 0.5381822 |
| GCD2 | YGR083C | 2213.5615 | 0.03701321 | 0.06022269 | 0.61460575 | 0.53881511 |
| FAR3 | YMR052W | 160.15808 | 0.08358294 | 0.13601341 | 0.61451986 | 0.53887185 |
| YNL140C | YNL140C | 12.4871932 | 0.17867834 | 0.29069894 | 0.61465082 | 0.53878534 |
| HAL9 | YOL089C | 687.821494 | 0.04718609 | 0.07680509 | 0.61436153 | 0.53897645 |
| Xu_2009_CUTs74 | Xu_2009_CU | 4.56269904 | -0.1751809 | 0.28572224 | -0.6131159 | 0.53979968 |
| YCL022C | YCL022C | 3.39061697 | 0.16762205 | 0.27360819 | 0.61263534 | 0.54011748 |
| KAR4 | YCL055W | 315.432188 | 0.06129426 | 0.10006694 | 0.6125326 | 0.54018544 |
| VPS4 | YPR173C | 1121.92298 | 0.04305884 | 0.07035787 | 0.61199741 | 0.54053947 |
| tG(GCC)J1 | tG(GCC)J1 | 0.71601084 | -0.0998237 | 0.16338297 | -0.6109798 | 0.54121292 |
| SPO77 | YLR341W | 42.3983173 | 0.13382632 | 0.21902198 | 0.61101777 | 0.54118782 |
| tR(ACG)K | tR(ACG)K | 3.91270402 | 0.16840647 | 0.27575056 | 0.6107203 | 0.54138477 |
| GDS1 | YOR355W | 1220.68822 | -0.0402311 | 0.06589046 | -0.6105755 | 0.54148064 |
| tF(GAA)D | tF(GAA)D | 6.55241455 | 0.1796856 | 0.29465952 | 0.60980756 | 0.5419893 |
| Xu_2009_SUTs54: | Xu_2009_SU | 62.2568165 | -0.1153317 | 0.18934804 | -0.6090987 | 0.54245899 |
| MRS3 | YJL133W | 694.818194 | -0.0495585 | 0.08141496 | -0.6087147 | 0.54271356 |
| Xu_2009_SUTs98 | Xu_2009_SU | 66.9351877 | 0.1132768 | 0.18617976 | 0.608427 | 0.54290431 |
| CET1 | YPL228W | 586.722837 | -0.0498821 | 0.08203967 | -0.6080238 | 0.54317171 |
| YBR241C | YBR241C | 511.619565 | -0.0532998 | 0.08768781 | -0.6078358 | 0.5432964 |
| YLR264C-A | YLR264C-A | 2.97052434 | 0.15797681 | 0.26012741 | 0.6073055 | 0.54364819 |
| tW(CCA)G2 | tW(CCA)G2 | 3.14225487 | -0.1581628 | 0.26099554 | -0.605998 | 0.54451608 |
| MME1 | YMR166C | 359.191922 | -0.0580989 | 0.0959376 | -0.6055905 | 0.54478674 |
| Xu_2009_SUTs72 | Xu_2009_SU | 42.0559056 | 0.14153144 | 0.23384544 | 0.60523496 | 0.54502289 |
| FAD1 | YDL045C | 528.313583 | 0.04985719 | 0.08236525 | 0.6053182 | 0.54496759 |
| YER066C-A | YER066C-A | 14.2441858 | 0.17242692 | 0.28493552 | 0.60514365 | 0.54508355 |
| RAD55 | YDR076W | 145.80926 | -0.0895931 | 0.14819901 | -0.6045456 | 0.54548094 |
| tR(ACG)J | tR(ACG)J | 23.2073678 | -0.1554732 | 0.25728104 | -0.6042934 | 0.54564856 |
| YAL004W | YAL004W | 1.34120365 | 0.12033499 | 0.19944593 | 0.6033464 | 0.54627827 |
| tI(AAU)I1 | tI(AAU)I1 | 0.88396618 | -0.1073062 | 0.17789867 | -0.6031871 | 0.54638421 |
| tN(GUU)O2 | tN(GUU)O2 | 8.98384096 | 0.17834307 | 0.2958165 | 0.60288413 | 0.54658578 |
| NUC1 | YJL208C | 864.999865 | 0.04646886 | 0.07710746 | 0.60265062 | 0.54674114 |
| PEX27 | YOR193W | 224.101572 | 0.07786234 | 0.12920453 | 0.60262855 | 0.54675582 |
| YLR446W | YLR446W | 106.411104 | -0.099754 | 0.16556728 | -0.6024983 | 0.54684249 |

| | | | | | | |
|---|---|---|---|---|---|---|
| UTP11 | YKL099C | 850.283781 | -0.049775 | 0.08271682 | -0.6017523 | 0.547339 |
| SED1 | YDR077W | 106343.713 | 0.03307699 | 0.05498533 | 0.60156017 | 0.54746695 |
| YAL042C-A | YAL042C-A | 3.13517753 | -0.1623047 | 0.26988083 | -0.6013939 | 0.54757769 |
| YJR023C | YJR023C | 1.6593138 | 0.1274159 | 0.21205101 | 0.60087383 | 0.54792403 |
| PDH1 | YPR002W | 189.156683 | 0.08802797 | 0.1465456 | 0.60068654 | 0.54804878 |
| GYP1 | YOR070C | 907.075049 | 0.04494257 | 0.07490665 | 0.59998104 | 0.54851887 |
| Xu_2009_CUTs25 | Xu_2009_CU | 138.575997 | -0.0862619 | 0.14384956 | -0.5996676 | 0.54872777 |
| YNR025C | YNR025C | 12.6264528 | 0.17376041 | 0.28998543 | 0.59920392 | 0.54903691 |
| LIH1 | YJR107W | 368.740335 | -0.0572914 | 0.09569523 | -0.5986865 | 0.54938193 |
| YHL017W | YHL017W | 841.525278 | -0.0534244 | 0.08937327 | -0.5977675 | 0.54999511 |
| JIP3 | YLR331C | 10.630226 | -0.176455 | 0.2951573 | -0.5978336 | 0.54995096 |
| KAR5 | YMR065W | 179.391076 | 0.07758828 | 0.12992323 | 0.59718563 | 0.55038345 |
| AAD6 | YFL056C | 156.863615 | -0.0845803 | 0.14167736 | -0.5969923 | 0.55051249 |
| NSP1 | YJL041W | 2142.09708 | -0.0352878 | 0.05912425 | -0.5968416 | 0.55061315 |
| YPL182C | YPL182C | 0.9915391 | 0.11070828 | 0.18579072 | 0.59587625 | 0.5512579 |
| tG(UCC)G/SUF4 | tG(UCC)G | 5.72806632 | -0.1732955 | 0.290914 | -0.5956931 | 0.55138027 |
| Xu_2009_SUTs78 | Xu_2009_SU | 149.708711 | 0.08092111 | 0.13595202 | 0.59521816 | 0.55169765 |
| YDR413C | YDR413C | 5.05043419 | 0.16810351 | 0.28242215 | 0.59522072 | 0.55169594 |
| YER085C | YER085C | 30.4063648 | -0.1466688 | 0.24649531 | -0.5950167 | 0.5518323 |
| YGL218W | YGL218W | 2.22707657 | -0.1419426 | 0.23855752 | -0.5950037 | 0.55184098 |
| MDM32 | YOR147W | 888.306662 | 0.04790153 | 0.08047902 | 0.59520522 | 0.5517063 |
| COS9 | YKL219W | 290.925046 | 0.06489704 | 0.10953863 | 0.59245801 | 0.55354393 |
| MDM38 | YOL027C | 2153.70466 | 0.03437269 | 0.05818459 | 0.5907524 | 0.55468633 |
| YSW1 | YBR148W | 107.6725 | -0.0911109 | 0.15434194 | -0.5903184 | 0.55497722 |
| ATG9 | YDL149W | 429.566658 | 0.0546393 | 0.09255923 | 0.59031714 | 0.55497805 |
| Xu_2009_SUTs17 | Xu_2009_SU | 71.1864058 | 0.1055664 | 0.17896931 | 0.58985758 | 0.55528613 |
| YCL001W-A | YCL001W-A | 32.0248852 | -0.1435699 | 0.24381396 | -0.5888502 | 0.55596179 |
| YPL264C | YPL264C | 90.0986073 | -0.1075581 | 0.1827529 | -0.5885439 | 0.55616726 |
| TRS31 | YDR472W | 512.062333 | -0.0513901 | 0.08734269 | -0.5883736 | 0.55628158 |
| MUD2 | YKL074C | 929.185327 | 0.04377392 | 0.07449333 | 0.58762198 | 0.55678605 |
| YMR103C | YMR103C | 113.17912 | 0.09125725 | 0.15530774 | 0.58758983 | 0.55680764 |
| YKL162C-A | YKL162C-A | 24.1595209 | 0.15489 | 0.26389364 | 0.58694102 | 0.55724332 |
| YOL106W | YOL106W | 13.2557871 | -0.1675457 | 0.28558035 | -0.5866851 | 0.5574152 |
| tV(AAC)M1 | tV(AAC)M1 | 8.19226929 | 0.17369196 | 0.29634035 | 0.5861232 | 0.55779273 |
| tI(AAU)I2 | tI(AAU)I2 | 1.53195937 | -0.1176345 | 0.2008231 | -0.5857616 | 0.55803576 |
| Xu_2009_CUTs55 | Xu_2009_CU | 11.4644059 | -0.1726167 | 0.29476682 | -0.5856041 | 0.55814156 |
| Xu_2009_SUTs73 | Xu_2009_SU | 56.1364315 | -0.1141889 | 0.19496509 | -0.5856889 | 0.55808459 |
| DMA2 | YNL116W | 423.851839 | 0.05406196 | 0.09235857 | 0.58534856 | 0.55831337 |
| YOR034C-A | YOR034C-A | 22.1750607 | -0.1541884 | 0.26344484 | -0.5852779 | 0.55836088 |
| POB3 | YML069W | 1236.02878 | 0.04110008 | 0.07034268 | 0.58428364 | 0.5590295 |
| HUR1 | YGL168W | 4.38379823 | -0.1661077 | 0.28440326 | -0.5840569 | 0.55918204 |

| | | | | | | |
|---|---|---|---|---|---|---|
| Xu_2009_SUTs79 | Xu_2009_SU | 3.80702558 | -0.161611 | 0.27718989 | -0.5830335 | 0.55987074 |
| IRC3 | YDR332W | 565.953307 | -0.0491595 | 0.08439945 | -0.582462 | 0.56025553 |
| RCF2 | YNR018W | 1259.23347 | -0.0466417 | 0.08014378 | -0.5819758 | 0.56058299 |
| tK(CUU)D1 | tK(CUU)D1 | 4.59703014 | -0.165537 | 0.28465809 | -0.5815291 | 0.56088393 |
| YKR040C | YKR040C | 3.06932069 | 0.15631001 | 0.26898992 | 0.58109987 | 0.56117315 |
| Xu_2009_CUTs1 | Xu_2009_CU | 5.30443335 | 0.16735754 | 0.28817868 | 0.58074228 | 0.56141417 |
| Xu_2009_SUTs78 | Xu_2009_SU | 8.88061008 | -0.1716259 | 0.29618739 | -0.5794504 | 0.56228528 |
| YFR012W-A | YFR012W-A | 34.0629492 | -0.1397459 | 0.24116 | -0.5794738 | 0.56226953 |
| NKP2 | YLR315W | 51.5136491 | 0.11863555 | 0.20481756 | 0.5792255 | 0.56243703 |
| SGF29 | YCL010C | 409.07556 | 0.05456563 | 0.0942571 | 0.57890209 | 0.56265524 |
| NOG1 | YPL093W | 4366.72575 | 0.04562943 | 0.07884022 | 0.57875825 | 0.56275231 |
| Xu_2009_CUTs20 | Xu_2009_CU | 2.96042343 | -0.1491496 | 0.25791692 | -0.5782855 | 0.56307135 |
| VPS3 | YDR495C | 591.223427 | -0.0467133 | 0.08075471 | -0.5784595 | 0.56295394 |
| ATG16 | YMR159C | 93.3109177 | 0.09859441 | 0.17049116 | 0.57829631 | 0.56306408 |
| EAP1 | YKL204W | 1172.32955 | -0.0390732 | 0.06763125 | -0.5777395 | 0.56344 |
| HIM1 | YDR317W | 123.358089 | -0.0894883 | 0.1549384 | -0.5775734 | 0.5635522 |
| CTM1 | YHR109W | 184.260598 | 0.07231538 | 0.12532158 | 0.57703853 | 0.56391343 |
| YLR236C | YLR236C | 2.25453777 | -0.1439016 | 0.24943612 | -0.5769076 | 0.56400188 |
| tG(GCC)O2/SUF17 | tG(GCC)O2 | 1.17088385 | 0.10318257 | 0.17921739 | 0.57573974 | 0.56479111 |
| TFB1 | YDR311W | 650.346523 | 0.0435366 | 0.07570493 | 0.57508277 | 0.56523532 |
| DIT1 | YDR403W | 131.975511 | -0.08621 | 0.14976184 | -0.5756473 | 0.5648536 |
| YGL217C | YGL217C | 3.91432386 | 0.15564249 | 0.27034682 | 0.57571414 | 0.56480842 |
| TTI2 | YJR136C | 388.157609 | -0.0533773 | 0.09274234 | -0.5755439 | 0.56492351 |
| PRI2 | YKL045W | 498.400846 | 0.05254134 | 0.09136186 | 0.5750905 | 0.56523009 |
| YRF1-5 | YLR467W | 4.28243671 | 0.1631107 | 0.28359075 | 0.57516227 | 0.56518156 |
| YMR172C-A | YMR172C-A | 2.26365618 | -0.1429475 | 0.2485124 | -0.5752126 | 0.56514752 |
| LAM6 | YLR072W | 838.126352 | -0.0406222 | 0.0708004 | -0.5737567 | 0.56613245 |
| Xu_2009_SUTs68 | Xu_2009_SU | 133.803465 | -0.0914365 | 0.15941883 | -0.5735615 | 0.56626456 |
| MAL12 | YGR292W | 21.7011582 | -0.153165 | 0.26706695 | -0.5735077 | 0.56630096 |
| STD1 | YOR047C | 1200.8003 | 0.04200634 | 0.0733469 | 0.57270783 | 0.56684254 |
| TPK3 | YKL166C | 659.038969 | -0.0468524 | 0.08183486 | -0.5725241 | 0.56696695 |
| KTR7 | YIL085C | 879.974677 | -0.0404681 | 0.07074761 | -0.5720062 | 0.56731775 |
| PHO2 | YDL106C | 458.817919 | 0.05089663 | 0.08901394 | 0.57178267 | 0.56746922 |
| HER2 | YMR293C | 162.546384 | -0.0782674 | 0.13688795 | -0.5717627 | 0.56748272 |
| Xu_2009_SUTs70 | Xu_2009_SU | 30.9836515 | 0.13956153 | 0.24440239 | 0.57103177 | 0.56797811 |
| YJR037W | YJR037W | 0.83798996 | -0.0941314 | 0.16492445 | -0.5707548 | 0.56816584 |
| UBA2 | YDR390C | 931.095954 | 0.03972958 | 0.06965574 | 0.57037052 | 0.56842642 |
| Xu_2009_SUTs62 | Xu_2009_SU | 37.6976607 | 0.13294165 | 0.23315234 | 0.57019223 | 0.56854733 |
| tP(UGG)L | tP(UGG)L | 3.45683576 | 0.14797593 | 0.26044825 | 0.56815866 | 0.56992724 |
| YJR029W | YJR029W | 101.119974 | 0.09072011 | 0.15974697 | 0.56789881 | 0.57010368 |
| PRP38 | YGR075C | 76.1852253 | -0.1079437 | 0.19026732 | -0.5673268 | 0.57049219 |

| | | | | | | |
|---|---|---|---|---|---|---|
| YOL155W-A | YOL155W-A | 0.79747345 | -0.0969018 | 0.17091033 | -0.5669744 | 0.57073154 |
| TYC1 | YBR296C-A | 1.33772145 | -0.1145531 | 0.20260029 | -0.5654145 | 0.57179187 |
| NUP82 | YJL061W | 919.636022 | 0.03922829 | 0.06939085 | 0.56532359 | 0.57185368 |
| YGL039W | YGL039W | 223.793048 | 0.06439803 | 0.11398827 | 0.56495317 | 0.57210562 |
| FAR10 | YLR238W | 494.733262 | 0.05065428 | 0.08981857 | 0.56396222 | 0.57277984 |
| EXP1 | YDL121C | 606.605242 | 0.04560704 | 0.08088672 | 0.5638384 | 0.57286412 |
| YDL163W | YDL163W | 2.09385015 | -0.1357944 | 0.24088939 | -0.5637209 | 0.57294409 |
| Xu_2009_SUTs47( | Xu_2009_SU | 6.70413615 | 0.16622846 | 0.29525639 | 0.56299697 | 0.57343695 |
| SPT6 | YGR116W | 3300.83271 | 0.0306898 | 0.05451622 | 0.56294803 | 0.57347027 |
| MSN4 | YKL062W | 1125.75991 | 0.03934321 | 0.06987481 | 0.56305282 | 0.57339892 |
| DDI2 | YFL061W | 3.26300782 | 0.15229513 | 0.27077385 | 0.56244399 | 0.57381355 |
| ISA1 | YLL027W | 660.830613 | -0.0449357 | 0.08004237 | -0.5613986 | 0.57452581 |
| SNR65 | snR65 | 18.2369651 | 0.16195847 | 0.28907478 | 0.56026496 | 0.57529872 |
| tL(UAA)N | tL(UAA)N | 2.43099711 | 0.13818009 | 0.24659138 | 0.5603606 | 0.5752335 |
| YLR428C | YLR428C | 2.56510409 | 0.14250999 | 0.25449787 | 0.55996538 | 0.57550305 |
| YIL020C-A | YIL020C-A | 2.35413455 | -0.1277825 | 0.22836289 | -0.5595591 | 0.57578018 |
| HOS2 | YGL194C | 599.940804 | 0.04392212 | 0.07861477 | 0.55870064 | 0.57636604 |
| Xu_2009_SUTs15 | Xu_2009_SU | 334.645571 | -0.0582106 | 0.10424008 | -0.5584281 | 0.57655212 |
| Xu_2009_CUTs23 | Xu_2009_CU | 4.4952846 | -0.1560404 | 0.27955848 | -0.5581674 | 0.57673009 |
| Xu_2009_SUTs31 | Xu_2009_SU | 8.02959926 | 0.16538157 | 0.29627742 | 0.55819837 | 0.57670894 |
| Xu_2009_CUTs52 | Xu_2009_CU | 5.06220903 | 0.16128032 | 0.2893731 | 0.55734387 | 0.57729251 |
| MIN3 | YMR182W-A | 24.7450839 | 0.14403792 | 0.25858693 | 0.55701933 | 0.57751422 |
| YNL035C | YNL035C | 609.720362 | -0.0437878 | 0.07879345 | -0.5557294 | 0.57839585 |
| YLR012C | YLR012C | 4.36532305 | -0.1567317 | 0.28221549 | -0.5553618 | 0.57864725 |
| YBP1 | YBR216C | 498.252253 | 0.05015243 | 0.09047948 | 0.55429621 | 0.57937615 |
| YPT35 | YHR105W | 175.593316 | -0.0730543 | 0.13178521 | -0.5543436 | 0.57934372 |
| Xu_2009_CUTs42 | Xu_2009_CU | 22.3705411 | 0.14525747 | 0.2623043 | 0.55377466 | 0.57973307 |
| FAR1 | YJL157C | 1923.60433 | 0.03600216 | 0.06516652 | 0.5524641 | 0.58063043 |
| MMO1 | YKL044W | 70.152719 | 0.10138451 | 0.18358918 | 0.55223578 | 0.58078683 |
| ANP1 | YEL036C | 2364.03377 | -0.0392209 | 0.07104354 | -0.552069 | 0.5809011 |
| YOR019W | YOR019W | 245.038885 | 0.06658022 | 0.12071552 | 0.55154645 | 0.58125913 |
| IMP3 | YHR148W | 462.597354 | 0.05241606 | 0.09513972 | 0.55093775 | 0.58167635 |
| Xu_2009_CUTs30 | Xu_2009_CU | 23.7242392 | -0.1486576 | 0.27018565 | -0.5502055 | 0.5821784 |
| PHB1 | YGR132C | 2451.25052 | 0.03811934 | 0.06940197 | 0.54925439 | 0.58283088 |
| Xu_2009_CUTs31 | Xu_2009_CU | 10.0293703 | 0.16109349 | 0.29368168 | 0.54853094 | 0.58332739 |
| YMR317W | YMR317W | 77.4535353 | 0.09631623 | 0.1757645 | 0.54798453 | 0.58370253 |
| Xu_2009_CUTs65 | Xu_2009_CU | 21.8191305 | -0.1425861 | 0.26047575 | -0.5474062 | 0.58409967 |
| RBS1 | YDL189W | 855.46869 | 0.04159372 | 0.07608049 | 0.54670682 | 0.58458016 |
| PAC1 | YOR269W | 564.018462 | -0.0458741 | 0.08405387 | -0.5457702 | 0.5852239 |
| NOP2 | YNL061W | 3457.33269 | 0.04268374 | 0.0782331 | 0.54559686 | 0.58534308 |
| BSD2 | YBR290W | 804.305799 | 0.03875859 | 0.07106882 | 0.54536708 | 0.58550107 |

| | | | | | | |
|---|---|---|---|---|---|---|
| RPB3 | YIL021W | 1106.77443 | -0.042585 | 0.07811367 | -0.5451668 | 0.58563879 |
| Xu_2009_CUTs78 | Xu_2009_CU | 10.7602487 | 0.1603196 | 0.29413967 | 0.54504582 | 0.585722 |
| YDL023C | YDL023C | 3.92455942 | -0.1487783 | 0.27330969 | -0.5443581 | 0.58619509 |
| Xu_2009_SUTs83 | Xu_2009_SU | 42.7191715 | 0.11890379 | 0.21861682 | 0.54389131 | 0.58651627 |
| SIP4 | YJL089W | 82.6479026 | -0.0967769 | 0.17821052 | -0.5430484 | 0.58709647 |
| YMR046W-A | YMR046W-A | 10.0172393 | -0.1603191 | 0.29520509 | -0.543077 | 0.58707681 |
| NOC2 | YOR206W | 2753.68774 | 0.03862053 | 0.07112329 | 0.54300817 | 0.58712418 |
| RAD10 | YML095C | 137.137404 | 0.07815515 | 0.14407932 | 0.54244529 | 0.58751179 |
| Xu_2009_SUTs37 | Xu_2009_SU | 38.2519373 | 0.1208243 | 0.22281424 | 0.54226472 | 0.58763615 |
| YGR204C-A | YGR204C-A | 2.41072568 | 0.13673625 | 0.25225539 | 0.54205483 | 0.58778074 |
| VAM6 | YDL077C | 943.401418 | -0.0380887 | 0.07034263 | -0.5414734 | 0.58818131 |
| KIN82 | YCR091W | 317.957411 | -0.0599485 | 0.11076066 | -0.5412433 | 0.58833987 |
| MLC2 | YPR188C | 338.176933 | -0.0611561 | 0.11300797 | -0.5411665 | 0.58839283 |
| YBL062W | YBL062W | 1.68197931 | -0.1228726 | 0.22746184 | -0.5401901 | 0.58906596 |
| BOR1 | YNL275W | 397.071093 | 0.05027894 | 0.09305835 | 0.5402948 | 0.58899375 |
| MPP6 | YNR024W | 279.518708 | 0.06084901 | 0.11270756 | 0.53988403 | 0.58927701 |
| YCR061W | YCR061W | 526.493376 | -0.0525333 | 0.09738133 | -0.5394597 | 0.58956972 |
| PRR2 | YDL214C | 150.934075 | 0.07988594 | 0.14830693 | 0.53865282 | 0.59012644 |
| ECM11 | YDR446W | 25.582675 | 0.13457498 | 0.25049155 | 0.53724357 | 0.59109938 |
| BRP1 | YGL007W | 6.08277886 | -0.1568191 | 0.29210354 | -0.5368613 | 0.59136344 |
| YOR394C-A | YOR394C-A | 5.32868991 | -0.156733 | 0.29226925 | -0.5362624 | 0.59177723 |
| Xu_2009_SUTs14 | Xu_2009_SU | 5.32207823 | 0.15571341 | 0.29062765 | 0.53578319 | 0.5921084 |
| YBR184W | YBR184W | 64.0311855 | -0.1059106 | 0.19769038 | -0.5357399 | 0.5921383 |
| XDJ1 | YLR090W | 1372.43785 | -0.0352965 | 0.06587282 | -0.5358285 | 0.59207705 |
| HIS5 | YIL116W | 868.008869 | 0.03895095 | 0.0727928 | 0.53509347 | 0.59258522 |
| Xu_2009_SUTs12 | Xu_2009_SU | 150.429772 | 0.0706271 | 0.13205998 | 0.53481079 | 0.5927807 |
| YJL163C | YJL163C | 203.273465 | -0.0761037 | 0.14236056 | -0.5345839 | 0.59293762 |
| PRM2 | YIL037C | 42.141996 | -0.1172368 | 0.21973047 | -0.5335483 | 0.59365406 |
| ZPS1 | YOL154W | 88.2685513 | 0.0902192 | 0.16909096 | 0.53355428 | 0.59364995 |
| YBR196C-A | YBR196C-A | 14.3176172 | -0.1522518 | 0.2856149 | -0.5330667 | 0.59398739 |
| YOR302W | YOR302W | 417.368604 | -0.0558848 | 0.10481645 | -0.5331679 | 0.59391734 |
| YBR300C | YBR300C | 0.83812672 | 0.08745155 | 0.16445436 | 0.53176786 | 0.59488679 |
| SNR38 | snR38 | 54.2851568 | -0.1124834 | 0.21176326 | -0.5311753 | 0.59529728 |
| Xu_2009_SUTs76 | Xu_2009_SU | 15.9675028 | 0.14907314 | 0.28058438 | 0.53129523 | 0.59521421 |
| YBR197C | YBR197C | 317.037024 | 0.05867647 | 0.11043102 | 0.53134043 | 0.5951829 |
| YGL034C | YGL034C | 4.6394409 | 0.15004969 | 0.28248764 | 0.53117257 | 0.5952992 |
| IRC9 | YJL142C | 2.51939965 | -0.1340269 | 0.25239754 | -0.5310151 | 0.59540832 |
| ERG29 | YMR134W | 270.781749 | 0.06041512 | 0.11401609 | 0.52988237 | 0.59619349 |
| Xu_2009_CUTs85 | Xu_2009_CU | 1.12972532 | 0.09756081 | 0.18456102 | 0.52861002 | 0.59707601 |
| Xu_2009_CUTs62 | Xu_2009_CU | 22.6266667 | -0.1426432 | 0.26991483 | -0.5284748 | 0.59716982 |
| TMA17 | YDL110C | 990.565981 | -0.0457067 | 0.08656926 | -0.5279781 | 0.59751457 |

| | | | | | | |
|---|---|---|---|---|---|---|
| YDR053W | YDR053W | 2.23385856 | -0.1309915 | 0.24809143 | -0.5279968 | 0.59750158 |
| YDR193W | YDR193W | 1.36041782 | 0.10352712 | 0.19605575 | 0.52804943 | 0.59746503 |
| REC107 | YJR021C | 297.680932 | 0.05455088 | 0.1032917 | 0.5281245 | 0.59741293 |
| YGL006W-A | YGL006W-A | 56.2506528 | 0.10902234 | 0.20671961 | 0.52739233 | 0.59792117 |
| YNL226W | YNL226W | 2.83327035 | 0.13699882 | 0.2599552 | 0.52700936 | 0.59818709 |
| Xu_2009_SUTs10 | Xu_2009_SU | 51.9990439 | 0.10917271 | 0.20727161 | 0.52671328 | 0.59839271 |
| YKL036C | YKL036C | 7.03342026 | -0.155345 | 0.29502574 | -0.5265471 | 0.5985081 |
| BUG1 | YDL099W | 660.981782 | -0.0412203 | 0.07831844 | -0.5263165 | 0.59866829 |
| YGR122C-A | YGR122C-A | 42.5167974 | -0.1129525 | 0.2151359 | -0.5250284 | 0.59956345 |
| SLM1 | YIL105C | 785.310538 | -0.0390294 | 0.07436502 | -0.524835 | 0.59969787 |
| YDL118W | YDL118W | 3.55355944 | -0.139273 | 0.26542532 | -0.5247164 | 0.59978036 |
| YML002W | YML002W | 370.909038 | 0.05039039 | 0.09613447 | 0.5241657 | 0.6001633 |
| YAL064W | YAL064W | 4.98394702 | 0.14842548 | 0.28324476 | 0.52401845 | 0.60026571 |
| Xu_2009_SUTs74 | Xu_2009_SU | 11.9279875 | 0.15201178 | 0.29024518 | 0.52373574 | 0.60046235 |
| Xu_2009_CUTs25 | Xu_2009_CU | 22.077621 | 0.13738742 | 0.26249786 | 0.52338492 | 0.60070641 |
| RXT2 | YBR095C | 694.27626 | -0.0402036 | 0.07684688 | -0.5231653 | 0.6008592 |
| GID8 | YMR135C | 347.274348 | -0.0503155 | 0.09624858 | -0.5227657 | 0.60113734 |
| YPQ2 | YDR352W | 397.127437 | 0.05088487 | 0.09737177 | 0.52258343 | 0.60126418 |
| PSK1 | YAL017W | 1971.91181 | 0.03307956 | 0.06332321 | 0.52239237 | 0.60139717 |
| Xu_2009_SUTs29 | Xu_2009_SU | 59.870909 | 0.10113085 | 0.19374027 | 0.52199187 | 0.60167599 |
| tG(CCC)D/SUF3 | tG(CCC)D | 3.79938301 | -0.1456098 | 0.27951945 | -0.5209291 | 0.60241617 |
| DYN2 | YDR424C | 291.62944 | 0.06046715 | 0.11607447 | 0.52093409 | 0.60241269 |
| tS(UGA)I/SUP16 | tS(UGA)I | 2.60994466 | -0.1312036 | 0.25214836 | -0.520343 | 0.60282454 |
| Xu_2009_SUTs44 | Xu_2009_SU | 18.6639171 | 0.14843343 | 0.28555586 | 0.51980524 | 0.60319933 |
| tK(CUU)P | tK(CUU)P | 3.07759737 | -0.1354891 | 0.26088835 | -0.5193377 | 0.6035253 |
| YLR140W | YLR140W | 1.20335045 | 0.09802708 | 0.1887597 | 0.51932209 | 0.60353615 |
| NPL6 | YMR091C | 891.869408 | 0.04338485 | 0.08355926 | 0.51921058 | 0.6036139 |
| YLR169W | YLR169W | 2.5848004 | -0.1311771 | 0.2528676 | -0.5187581 | 0.60392947 |
| MAL13 | YGR288W | 263.548003 | 0.05614372 | 0.1083 | 0.51840922 | 0.60417278 |
| SOR1 | YJR159W | 5.90552156 | -0.1510271 | 0.29167914 | -0.517785 | 0.60460826 |
| YKL169C | YKL169C | 12.1191798 | 0.15049982 | 0.29089423 | 0.51736957 | 0.6048982 |
| Xu_2009_SUTs84 | Xu_2009_SU | 34.6617221 | 0.11935554 | 0.23124544 | 0.51614224 | 0.60575507 |
| TFC6 | YDR362C | 502.378046 | 0.04598871 | 0.0891039 | 0.5161246 | 0.60576739 |
| YHR032W-A | YHR032W-A | 49.3887875 | 0.10817197 | 0.2096684 | 0.51591931 | 0.60591077 |
| AAD10 | YJR155W | 313.235716 | -0.054209 | 0.10508598 | -0.5158539 | 0.60595648 |
| Xu_2009_SUTs77 | Xu_2009_SU | 4.61519032 | 0.14564943 | 0.28270703 | 0.51519563 | 0.60641632 |
| CBS1 | YDL069C | 139.617435 | -0.0742285 | 0.1441719 | -0.514861 | 0.60665018 |
| Xu_2009_SUTs15 | Xu_2009_SU | 22.0578927 | 0.13485064 | 0.26216685 | 0.51436955 | 0.60699364 |
| GTA1 | YEL043W | 1350.1492 | -0.0313393 | 0.06092138 | -0.5144228 | 0.60695639 |
| COX1 | Q0045 | 1286.17039 | -0.0804712 | 0.15668237 | -0.5135948 | 0.60753535 |
| Xu_2009_CUTs80 | Xu_2009_CU | 14.1119405 | -0.1454293 | 0.2830509 | -0.5137921 | 0.60739732 |

| | | | | | | |
|---|---|---|---|---|---|---|
| FYV1 | YDR024W | 11.6212642 | -0.1500578 | 0.29203511 | -0.5138348 | 0.60736751 |
| STR2 | YJR130C | 498.965733 | -0.0440467 | 0.08576179 | -0.5135937 | 0.60753606 |
| YOR333C | YOR333C | 1.30231142 | 0.10392592 | 0.20220494 | 0.51396332 | 0.60727763 |
| CIT3 | YPR001W | 106.951793 | -0.0825245 | 0.16058213 | -0.5139081 | 0.60731622 |
| YHL002C-A | YHL002C-A | 5.18089954 | -0.1472819 | 0.28684885 | -0.5134479 | 0.60763806 |
| Xu_2009_CUTs61 | Xu_2009_CU | 21.8862322 | -0.1344161 | 0.26195751 | -0.5131217 | 0.6078662 |
| NTH1 | YDR001C | 4015.52927 | 0.02965918 | 0.057812 | 0.51302815 | 0.60793163 |
| Xu_2009_SUTs41 | Xu_2009_SU | 7.90436147 | 0.15192383 | 0.29644516 | 0.51248543 | 0.60831131 |
| RPC11 | YDR045C | 197.110689 | 0.06835916 | 0.13344696 | 0.51225717 | 0.60847104 |
| Xu_2009_CUTs38 | Xu_2009_CU | 35.7441087 | -0.1191931 | 0.23279861 | -0.5120009 | 0.60865037 |
| YJL028W | YJL028W | 6.42956951 | 0.15046704 | 0.29433996 | 0.51120152 | 0.60920996 |
| CNN1 | YFR046C | 309.870735 | 0.05502289 | 0.10769174 | 0.51092953 | 0.6094004 |
| YOR300W | YOR300W | 27.0635197 | 0.13066373 | 0.25608137 | 0.51024303 | 0.60988121 |
| SPO73 | YER046W | 193.736932 | 0.06656287 | 0.13050257 | 0.51005029 | 0.61001623 |
| RSC8 | YFR037C | 1404.18449 | 0.03310321 | 0.06507003 | 0.50873199 | 0.6109401 |
| YOR376W-A | YOR376W-A | 174.111282 | 0.06321836 | 0.1243968 | 0.50819929 | 0.61131359 |
| tH(GUG)M | tH(GUG)M | 12.7585479 | -0.1472267 | 0.28995559 | -0.5077562 | 0.61162435 |
| tP(UGG)H/SUF8 | tP(UGG)H | 5.33304273 | -0.1478822 | 0.2912205 | -0.5078016 | 0.61159252 |
| YLR407W | YLR407W | 917.734787 | 0.03748902 | 0.07383193 | 0.50776165 | 0.61162051 |
| TAH1 | YCR060W | 312.094527 | 0.05362081 | 0.10572473 | 0.50717377 | 0.6120329 |
| Xu_2009_SUTs58 | Xu_2009_SU | 182.307717 | 0.06677455 | 0.13180685 | 0.5066091 | 0.61242912 |
| YDR203W | YDR203W | 7.65037444 | -0.1501458 | 0.29643289 | -0.5065085 | 0.61249971 |
| OTU2 | YHL013C | 728.001242 | 0.04385029 | 0.08668219 | 0.50587426 | 0.61294493 |
| PMS1 | YNL082W | 381.53297 | 0.05266835 | 0.10409202 | 0.50597879 | 0.61287154 |
| IMO32 | YGR031W | 540.486645 | 0.04261985 | 0.08432103 | 0.5054475 | 0.61324457 |
| SPT3 | YDR392W | 484.884434 | -0.0450913 | 0.08928938 | -0.5050024 | 0.61355718 |
| NTG1 | YAL015C | 546.675727 | 0.04382148 | 0.08680677 | 0.50481644 | 0.61368777 |
| Xu_2009_CUTs55 | Xu_2009_CU | 26.9955542 | 0.12837086 | 0.25442143 | 0.50455993 | 0.61386796 |
| YKL162C | YKL162C | 194.544256 | 0.0616164 | 0.12224638 | 0.5040345 | 0.61423713 |
| tA(UGC)L | tA(UGC)L | 3.21393978 | -0.1332384 | 0.26464486 | -0.5034611 | 0.6146401 |
| Xu_2009_SUTs19 | Xu_2009_SU | 44.9328559 | 0.10654881 | 0.21228684 | 0.50190964 | 0.61573109 |
| PHO89 | YBR296C | 18.8215336 | 0.13842315 | 0.27583318 | 0.50183648 | 0.61578255 |
| ANT1 | YPR128C | 276.98758 | -0.0550505 | 0.10965953 | -0.5020125 | 0.61565874 |
| YLL017W | YLL017W | 21.1160097 | -0.1384746 | 0.27613726 | -0.5014702 | 0.61604024 |
| DDL1 | YOR022C | 400.981148 | 0.04754994 | 0.09482074 | 0.5014719 | 0.61603905 |
| YLR416C | YLR416C | 7.91924576 | 0.1479048 | 0.29551734 | 0.50049449 | 0.61672693 |
| YDL180W | YDL180W | 666.126047 | -0.0405674 | 0.08111718 | -0.5001088 | 0.61699847 |
| Xu_2009_CUTs95 | Xu_2009_CU | 10.7543215 | 0.14709974 | 0.29461932 | 0.49928748 | 0.61757687 |
| YPL238C | YPL238C | 12.3349269 | -0.1450207 | 0.29049554 | -0.4992183 | 0.61762562 |
| RGS2 | YOR107W | 1208.46342 | -0.0406224 | 0.08145245 | -0.4987254 | 0.61797285 |
| Xu_2009_CUTs37 | Xu_2009_CU | 5.68027757 | 0.14377229 | 0.28878669 | 0.49784946 | 0.61859015 |

| | | | | | | |
|---|---|---|---|---|---|---|
| GPI1 | YGR216C | 405.723183 | -0.0456259 | 0.09163251 | -0.4979227 | 0.61853852 |
| CIC1 | YHR052W | 1430.89844 | 0.04132493 | 0.08316085 | 0.49692767 | 0.61924006 |
| Xu_2009_SUTs23 | Xu_2009_SU | 15.2028766 | -0.1419779 | 0.28582763 | -0.4967256 | 0.61938258 |
| tA(AGC)M2 | tA(AGC)M2 | 1.12940597 | -0.0872925 | 0.17586058 | -0.4963733 | 0.61963107 |
| ANR2 | YKL047W | 626.060423 | -0.0411011 | 0.08281761 | -0.4962851 | 0.61969328 |
| ASG7 | YJL170C | 25.6055205 | -0.1282382 | 0.25884867 | -0.4954177 | 0.62030529 |
| SPC25 | YER018C | 117.000911 | -0.0793398 | 0.16020614 | -0.495236 | 0.62043354 |
| LAA2 | YBL010C | 303.282016 | -0.0559144 | 0.11296128 | -0.4949876 | 0.62060888 |
| YMR175W-A | YMR175W-A | 18.162651 | 0.13844389 | 0.27970205 | 0.49496915 | 0.6206219 |
| YMR315W-A | YMR315W-A | 40.3179108 | -0.1104676 | 0.22333465 | -0.494628 | 0.62086274 |
| SPS18 | YNL204C | 167.40788 | 0.06941821 | 0.14040631 | 0.49440946 | 0.62101704 |
| tM(CAU)J2/EMT3 | tM(CAU)J2 | 3.20367085 | 0.13062126 | 0.26442791 | 0.49397685 | 0.62132253 |
| TRS33 | YOR115C | 507.822721 | -0.0451083 | 0.0913475 | -0.4938099 | 0.62144042 |
| TFB2 | YPL122C | 445.831485 | 0.04864505 | 0.09852261 | 0.49374505 | 0.62148625 |
| Xu_2009_CUTs60 | Xu_2009_CU | 14.6820814 | 0.14025892 | 0.2843023 | 0.49334429 | 0.62176934 |
| FUR4 | YBR021W | 272.659334 | 0.05723728 | 0.11599258 | 0.49345636 | 0.62169017 |
| UFO1 | YML088W | 1131.69039 | 0.03645611 | 0.07386759 | 0.49353319 | 0.62163589 |
| UBC5 | YDR059C | 443.833238 | 0.0492209 | 0.0998402 | 0.49299679 | 0.62201485 |
| YVC1 | YOR087W | 516.835623 | -0.0434434 | 0.08822138 | -0.492436 | 0.62241113 |
| YPR146C | YPR146C | 15.3774574 | 0.14183145 | 0.288062 | 0.49236431 | 0.62246182 |
| Xu_2009_SUTs71 | Xu_2009_SU | 17.2761729 | 0.14125466 | 0.2871068 | 0.49199343 | 0.62272399 |
| VBA5 | YKR105C | 4.03240851 | -0.1384252 | 0.28167341 | -0.4914387 | 0.62311618 |
| MSB1 | YOR188W | 1151.33186 | 0.03195501 | 0.06503914 | 0.49131974 | 0.62320032 |
| APS3 | YJL024C | 543.544386 | -0.0436064 | 0.08899382 | -0.4899936 | 0.6241384 |
| YNL198C | YNL198C | 6.6854358 | 0.14414856 | 0.29455577 | 0.48937613 | 0.62457543 |
| YDR118W-A | YDR118W-A | 5.96904805 | 0.14341379 | 0.29315116 | 0.48921449 | 0.62468985 |
| PAU13 | YHL046C | 3.73481534 | -0.1345672 | 0.27518645 | -0.4890038 | 0.62483898 |
| Xu_2009_CUTs61 | Xu_2009_CU | 44.5001019 | 0.10588847 | 0.21676723 | 0.48848929 | 0.62520331 |
| YDR095C | YDR095C | 10.535223 | 0.14258719 | 0.29199026 | 0.48832859 | 0.62531711 |
| PAC10 | YGR078C | 1091.78958 | 0.03640496 | 0.07458652 | 0.4880903 | 0.62548588 |
| YDR524C-A | YDR524C-A | 4.34815071 | -0.1358267 | 0.27860835 | -0.4875184 | 0.625891 |
| Xu_2009_SUTs58 | Xu_2009_SU | 34.4225987 | 0.1159967 | 0.23804238 | 0.48729431 | 0.62604978 |
| HUL4 | YJR036C | 193.44298 | 0.06386623 | 0.13108104 | 0.48722706 | 0.62609743 |
| MST28 | YAR033W | 108.607444 | -0.0743186 | 0.15269059 | -0.486727 | 0.62645182 |
| ENA5 | YDR038C | 272.820045 | -0.0544276 | 0.11180722 | -0.486799 | 0.62640077 |
| YPR050C | YPR050C | 15.1061353 | 0.13874614 | 0.28568835 | 0.48565557 | 0.62721139 |
| QCR7 | YDR529C | 3248.47088 | 0.02880297 | 0.05952134 | 0.48391002 | 0.62844973 |
| YLR202C | YLR202C | 23.4842204 | -0.1268132 | 0.26203502 | -0.483955 | 0.62841778 |
| Xu_2009_SUTs29 | Xu_2009_SU | 58.7482555 | 0.1001852 | 0.20750995 | 0.48279708 | 0.62923983 |
| STE2 | YFL026W | 101.581747 | 0.08198072 | 0.17023556 | 0.48157224 | 0.63010985 |
| OSW1 | YOR255W | 10.392433 | 0.14249585 | 0.29609826 | 0.48124514 | 0.63034228 |

| | | | | | | |
|---|---|---|---|---|---|---|
| Xu_2009_CUTs67 | Xu_2009_CU | 9.9430355 | 0.14184881 | 0.29617027 | 0.47894343 | 0.63197887 |
| DPI35 | YMR130W | 343.394673 | -0.0475496 | 0.09928513 | -0.4789198 | 0.63199566 |
| Xu_2009_CUTs20 | Xu_2009_CU | 4.34173281 | 0.13293076 | 0.27766155 | 0.47875107 | 0.63211573 |
| FIG4 | YNL325C | 313.998453 | 0.05036571 | 0.10521786 | 0.47868026 | 0.63216611 |
| Xu_2009_SUTs46 | Xu_2009_SU | 20.615159 | 0.12948154 | 0.27088136 | 0.47800091 | 0.63264955 |
| ESP1 | YGR098C | 1269.19669 | -0.0324678 | 0.06794612 | -0.4778465 | 0.63275947 |
| AIM39 | YOL053W | 481.266997 | -0.0502604 | 0.10555779 | -0.4761408 | 0.63397405 |
| Xu_2009_CUTs86 | Xu_2009_CU | 20.2858429 | 0.13050012 | 0.27491999 | 0.474684 | 0.63501222 |
| PAU18 | YLL064C | 2.19914062 | 0.11524696 | 0.242926 | 0.47441181 | 0.63520627 |
| YNL057W | YNL057W | 18.0668829 | -0.1323195 | 0.27887349 | -0.4744786 | 0.63515867 |
| RUF23 | RUF23 | 7.69543584 | -0.140063 | 0.29556056 | -0.4738893 | 0.63557888 |
| YMR272W-B | YMR272W-B | 3.26951708 | 0.12279894 | 0.25935475 | 0.47347868 | 0.63587171 |
| CDC53 | YDL132W | 2289.13949 | -0.0288604 | 0.06105886 | -0.472665 | 0.63645222 |
| YER068C-A | YER068C-A | 2.29438656 | 0.07317024 | 0.15476408 | 0.47278566 | 0.6363661 |
| RAD51 | YER095W | 794.750261 | -0.0339428 | 0.07179894 | -0.4727475 | 0.63639332 |
| FDH2 | YPL275W | 3.69856784 | 0.13197305 | 0.27931539 | 0.47248758 | 0.63657881 |
| DMC1 | YER179W | 61.1569853 | 0.08888843 | 0.18861406 | 0.4712715 | 0.63744687 |
| SEC8 | YPR055W | 719.605085 | -0.0395595 | 0.08397283 | -0.4710991 | 0.63756996 |
| Xu_2009_SUTs45 | Xu_2009_SU | 19.296746 | 0.12775782 | 0.27130026 | 0.47090933 | 0.63770548 |
| YNL320W | YNL320W | 179.127412 | 0.06371037 | 0.13544792 | 0.47036801 | 0.63809212 |
| HUB1 | YNR032C-A | 106.217981 | 0.07436395 | 0.15824595 | 0.46992643 | 0.63840758 |
| Xu_2009_CUTs75 | Xu_2009_CU | 11.2006508 | 0.1373187 | 0.29267542 | 0.46918426 | 0.63893793 |
| IRC19 | YLL033W | 80.1435586 | -0.0836983 | 0.17839815 | -0.4691655 | 0.63895132 |
| Xu_2009_SUTs58 | Xu_2009_SU | 21.8549852 | 0.12551332 | 0.26792529 | 0.46846387 | 0.6394529 |
| Xu_2009_SUTs30 | Xu_2009_SU | 56.0118966 | 0.09108827 | 0.19460036 | 0.46807864 | 0.63972835 |
| ATH1 | YPR026W | 771.397005 | -0.0472012 | 0.1008548 | -0.468011 | 0.63977673 |
| Xu_2009_SUTs42 | Xu_2009_SU | 63.9083684 | 0.08861745 | 0.18954776 | 0.46752045 | 0.64012756 |
| YBR063C | YBR063C | 361.907519 | 0.04345803 | 0.09305063 | 0.46703642 | 0.64047382 |
| YOL131W | YOL131W | 8.92338358 | -0.1384595 | 0.29660841 | -0.466809 | 0.64063651 |
| Xu_2009_CUTs26 | Xu_2009_CU | 20.5862003 | -0.1230392 | 0.26391871 | -0.466201 | 0.64107167 |
| YHR049C-A | YHR049C-A | 7.25856457 | 0.13786251 | 0.29580707 | 0.46605549 | 0.64117578 |
| ABP140 | YOR239W | 1640.42617 | 0.02902251 | 0.06233488 | 0.46559025 | 0.64150882 |
| YNR001W-A | YNR001W-A | 1.46439902 | 0.09919149 | 0.21329956 | 0.46503374 | 0.6419073 |
| CDC50 | YCR094W | 604.294236 | -0.0414507 | 0.08922525 | -0.464563 | 0.64224446 |
| YDR015C | YDR015C | 0.88265337 | 0.0835637 | 0.17990869 | 0.46447842 | 0.64230502 |
| YJR146W | YJR146W | 14.4118751 | 0.13115422 | 0.28240653 | 0.4644164 | 0.64234944 |
| VPS69 | YPR087W | 9.45733088 | -0.1376837 | 0.29656022 | -0.4642688 | 0.64245516 |
| Xu_2009_SUTs23 | Xu_2009_SU | 83.7240389 | 0.08581827 | 0.18490857 | 0.4641119 | 0.64256757 |
| Xu_2009_SUTs29 | Xu_2009_SU | 2.08846986 | 0.11397958 | 0.24621402 | 0.46292886 | 0.64341536 |
| RRT2 | YBR246W | 639.589213 | -0.0364913 | 0.07899129 | -0.4619666 | 0.64410528 |
| YFL013W-A | YFL013W-A | 7.12791269 | 0.13640193 | 0.29520885 | 0.4620523 | 0.64404382 |

| | | | | | | |
|---|---|---|---|---|---|---|
| HMS2 | YJR147W | 290.283463 | 0.0492656 | 0.10660844 | 0.46211723 | 0.64399726 |
| TAD3 | YLR316C | 194.899076 | -0.057985 | 0.12543622 | -0.4622667 | 0.64389007 |
| Xu_2009_CUTs55 | Xu_2009_CU | 22.1697122 | -0.1243499 | 0.26946094 | -0.4614764 | 0.64445688 |
| MGT1 | YDL200C | 176.600803 | -0.0622931 | 0.13527819 | -0.4604812 | 0.64517083 |
| VTS1 | YOR359W | 1692.91857 | 0.03062894 | 0.06657665 | 0.46005523 | 0.64547658 |
| YMR321C | YMR321C | 9.96803236 | -0.1353308 | 0.29472084 | -0.459183 | 0.64610277 |
| RSB1 | YOR049C | 60.5623772 | 0.09002009 | 0.19608872 | 0.45907836 | 0.6461779 |
| tQ(UUG)B | tQ(UUG)B | 6.38288488 | -0.1353883 | 0.29508132 | -0.4588169 | 0.64636563 |
| YSY6 | YBR162W-A | 1159.00666 | -0.0316493 | 0.06900332 | -0.4586641 | 0.64647539 |
| HIT1 | YJR055W | 116.805576 | -0.0780061 | 0.17023267 | -0.4582325 | 0.64678541 |
| BUB3 | YOR026W | 363.73594 | -0.0428977 | 0.09364187 | -0.4581043 | 0.64687754 |
| YDR114C | YDR114C | 2.63739239 | -0.1182675 | 0.25833774 | -0.4578018 | 0.64709482 |
| PAU24 | YBR301W | 5.68114784 | -0.1323024 | 0.2895018 | -0.4570004 | 0.64767077 |
| ATG14 | YBR128C | 112.685736 | -0.0754533 | 0.16542363 | -0.4561217 | 0.64830245 |
| LOT6 | YLR011W | 160.431017 | 0.06169034 | 0.13522475 | 0.45620596 | 0.64824188 |
| Xu_2009_SUTs35 | Xu_2009_SU | 59.1403673 | -0.0896669 | 0.19694174 | -0.4552965 | 0.64889595 |
| ICS2 | YBR157C | 250.185226 | 0.05143681 | 0.11292537 | 0.45549383 | 0.648754 |
| YGR240C-A | YGR240C-A | 1.60670575 | 0.10002939 | 0.21966472 | 0.45537305 | 0.64884088 |
| INP1 | YMR204C | 141.746417 | -0.061934 | 0.13600672 | -0.4553746 | 0.64883974 |
| SPS22 | YCL048W | 18.821224 | -0.1241162 | 0.27279817 | -0.4549745 | 0.64912758 |
| ATP14 | YLR295C | 786.497761 | -0.0342956 | 0.07540819 | -0.4547993 | 0.64925363 |
| PZF1 | YPR186C | 511.976363 | 0.03953563 | 0.08698909 | 0.45448952 | 0.64947654 |
| Xu_2009_SUTs34 | Xu_2009_SU | 26.3150738 | 0.11308273 | 0.24950339 | 0.45323123 | 0.65038224 |
| GLY1 | YEL046C | 4165.62369 | -0.0308212 | 0.06819427 | -0.4519614 | 0.65129676 |
| HMG2 | YLR450W | 1420.13647 | -0.0306844 | 0.06800383 | -0.451215 | 0.65183459 |
| ARN2 | YHL047C | 489.019197 | 0.04007422 | 0.08890881 | 0.45073393 | 0.65218132 |
| MEO1 | YBR126W-A | 1.73994374 | -0.0989864 | 0.22037176 | -0.4491791 | 0.65330243 |
| YDR249C | YDR249C | 213.35555 | -0.0614121 | 0.13668909 | -0.4492834 | 0.65322725 |
| YER046W-A | YER046W-A | 8.91190371 | 0.13279843 | 0.29561503 | 0.44922759 | 0.65326748 |
| HHY1 | YEL059W | 21.0179515 | 0.12012692 | 0.2675632 | 0.44896652 | 0.65345581 |
| RCF3 | YBR255C-A | 121.574937 | 0.06500438 | 0.14529491 | 0.44739616 | 0.65458905 |
| AIM21 | YIR003W | 777.249943 | 0.03442188 | 0.07693714 | 0.44740262 | 0.65458439 |
| YDR417C | YDR417C | 4.48533482 | 0.12648113 | 0.2827845 | 0.44727038 | 0.65467985 |
| Xu_2009_CUTs83 | Xu_2009_CU | 2.78933763 | 0.11770076 | 0.2633088 | 0.44700654 | 0.65487033 |
| YLR294C | YLR294C | 648.273303 | -0.0372871 | 0.08354023 | -0.4463371 | 0.65535374 |
| FCP1 | YMR277W | 1394.47659 | 0.03037114 | 0.06816739 | 0.44553775 | 0.65593119 |
| TAP42 | YMR028W | 438.18515 | 0.03925503 | 0.08817495 | 0.44519486 | 0.65617894 |
| Xu_2009_CUTs86 | Xu_2009_CU | 15.9357608 | -0.1288599 | 0.28961146 | -0.4449407 | 0.65636262 |
| RAM1 | YDL090C | 197.015844 | -0.0541827 | 0.12176932 | -0.444962 | 0.65634721 |
| Xu_2009_SUTs29 | Xu_2009_SU | 33.6328906 | 0.10928061 | 0.24592074 | 0.44437328 | 0.65677273 |
| YPR015C | YPR015C | 88.0413818 | 0.07428231 | 0.16730259 | 0.44399975 | 0.65704276 |

| | | | | | | |
|---|---|---|---|---|---|---|
| Xu_2009_CUTs10 | Xu_2009_CU | 14.7652197 | -0.129433 | 0.29254671 | -0.4424352 | 0.65817432 |
| YGR273C | YGR273C | 10.1668839 | -0.1302432 | 0.29441334 | -0.442382 | 0.65821278 |
| SUA7 | YPR086W | 1397.80889 | 0.02744667 | 0.06203417 | 0.44244431 | 0.65816772 |
| ATM1 | YMR301C | 889.608917 | -0.0328924 | 0.07440137 | -0.4420934 | 0.65842165 |
| SKP2 | YNL311C | 953.253118 | 0.03228311 | 0.07312988 | 0.44144896 | 0.65888801 |
| SDP1 | YIL113W | 120.040661 | 0.07528401 | 0.17091159 | 0.44048508 | 0.65958581 |
| NUP53 | YMR153W | 469.823835 | 0.03760594 | 0.0855559 | 0.43954814 | 0.66026441 |
| tW(CCA)J | tW(CCA)J | 2.03980695 | -0.1001307 | 0.22817806 | -0.4388271 | 0.66078683 |
| EXG2 | YDR261C | 1371.6624 | 0.02928645 | 0.06675145 | 0.43873877 | 0.66085083 |
| DCV1 | YFR012W | 43.9350782 | -0.0949486 | 0.21639734 | -0.4387699 | 0.66082827 |
| Xu_2009_CUTs39 | Xu_2009_CU | 13.8375222 | 0.1263675 | 0.2888458 | 0.43749122 | 0.66175514 |
| YNL013C | YNL013C | 1.23321888 | -0.088569 | 0.202431 | -0.4375269 | 0.66172928 |
| YGR228W | YGR228W | 1.25978481 | -0.0900645 | 0.206281 | -0.4366106 | 0.66239378 |
| PEX14 | YGL153W | 432.368314 | -0.0418377 | 0.09588062 | -0.4363524 | 0.66258107 |
| YOR199W | YOR199W | 3.02538457 | -0.1153167 | 0.2644698 | -0.4360298 | 0.66281508 |
| YGL165C | YGL165C | 2.0843665 | 0.1017012 | 0.23334284 | 0.43584455 | 0.66294952 |
| MAL11 | YGR289C | 97.2615911 | -0.0718924 | 0.165331 | -0.4348393 | 0.66367908 |
| YKR041W | YKR041W | 78.6694858 | 0.0757599 | 0.1744626 | 0.43424722 | 0.66410893 |
| STB5 | YHR178W | 336.148035 | -0.0475919 | 0.10966706 | -0.4339671 | 0.66431234 |
| GUF1 | YLR289W | 300.455082 | 0.04456113 | 0.1028347 | 0.43332779 | 0.66477665 |
| ABF1 | YKL112W | 1142.15765 | -0.0340954 | 0.07873646 | -0.4330316 | 0.66499182 |
| Xu_2009_SUTs10! | Xu_2009_SU | 345.902271 | 0.04305435 | 0.09946947 | 0.43283984 | 0.66513113 |
| TRR2 | YHR106W | 392.087988 | -0.0413606 | 0.09562804 | -0.4325149 | 0.66536723 |
| Xu_2009_SUTs25! | Xu_2009_SU | 6.3314375 | 0.12721209 | 0.2942171 | 0.43237492 | 0.66546894 |
| BDF2 | YDL070W | 3594.44676 | 0.02535735 | 0.05870634 | 0.43193542 | 0.66578835 |
| VMA22 | YHR060W | 309.473297 | 0.04615767 | 0.10693163 | 0.43165586 | 0.66599155 |
| YNL042W-B | YNL042W-B | 12.5287374 | 0.12563708 | 0.29102095 | 0.43171145 | 0.66595114 |
| Xu_2009_SUTs64 | Xu_2009_SU | 8.94027263 | -0.1277452 | 0.29642609 | -0.4309511 | 0.66650391 |
| YMR075C-A | YMR075C-A | 3.3170532 | -0.1163156 | 0.27057979 | -0.4298756 | 0.66728614 |
| Xu_2009_CUTs89 | Xu_2009_CU | 1.14203974 | 0.08695274 | 0.20243612 | 0.42953176 | 0.66753628 |
| Xu_2009_CUTs89 | Xu_2009_CU | 1.04810662 | 0.0811494 | 0.18914634 | 0.42902972 | 0.6679016 |
| YEH2 | YLR020C | 964.935294 | 0.02972781 | 0.06936124 | 0.42859397 | 0.66821874 |
| tD(GUC)I1 | tD(GUC)I1 | 6.03698448 | -0.12492 | 0.29172545 | -0.4282107 | 0.66849771 |
| MUM3 | YOR298W | 109.069802 | 0.06742055 | 0.15745221 | 0.42819687 | 0.66850779 |
| APQ13 | YJL075C | 1.04033995 | -0.0794751 | 0.1857577 | -0.4278427 | 0.66876568 |
| tA(UGC)E | tA(UGC)E | 9.0607015 | -0.1267559 | 0.296507 | -0.427497 | 0.66901736 |
| YDR199W | YDR199W | 3.10215679 | -0.1099826 | 0.25767854 | -0.4268208 | 0.66950983 |
| QDR1 | YIL120W | 259.207077 | 0.04766565 | 0.11186538 | 0.42609831 | 0.67003621 |
| YGR259C | YGR259C | 4.27700836 | 0.11935597 | 0.28067823 | 0.42524129 | 0.67066078 |
| KTI11 | YBL071W-A | 421.950342 | 0.0396074 | 0.09322418 | 0.42486184 | 0.6709374 |
| YBR099C | YBR099C | 16.5640982 | -0.1166418 | 0.27471377 | -0.4245939 | 0.67113272 |

| | | | | | | |
|---|---|---|---|---|---|---|
| SLO1 | YER180C-A | 31.857353 | -0.1016878 | 0.23977326 | -0.4240997 | 0.67149313 |
| PTI1 | YGR156W | 421.160812 | 0.03903858 | 0.09228193 | 0.423036 | 0.672269 |
| DAL81 | YIR023W | 1743.08165 | 0.0250859 | 0.05932853 | 0.42283027 | 0.6724191 |
| BUD26 | YDR241W | 89.3605018 | 0.06960234 | 0.16474376 | 0.42248847 | 0.67266851 |
| YPR012W | YPR012W | 8.05237745 | 0.12525937 | 0.29652419 | 0.42242548 | 0.67271448 |
| CTT1 | YGR088W | 742.51536 | 0.04145963 | 0.09827076 | 0.42189184 | 0.67310396 |
| RUF21 | RUF21 | 1.14257108 | 0.083972 | 0.19919785 | 0.42155071 | 0.67335299 |
| Xu_2009_CUTs57 | Xu_2009_CU | 2.0114629 | -0.0970794 | 0.23040059 | -0.4213503 | 0.67349929 |
| Xu_2009_SUTs58 | Xu_2009_SU | 5.87127877 | 0.12170911 | 0.28889762 | 0.42128802 | 0.67354477 |
| ASH1 | YKL185W | 2663.93811 | 0.02658092 | 0.06312673 | 0.42107227 | 0.67370231 |
| Xu_2009_SUTs44 | Xu_2009_SU | 2.42019753 | 0.10531756 | 0.2502914 | 0.42077977 | 0.67391591 |
| YFL019C | YFL019C | 20.8363628 | 0.11423094 | 0.27140204 | 0.42089198 | 0.67383396 |
| Xu_2009_CUTs15 | Xu_2009_CU | 7.33282266 | -0.1240361 | 0.29514231 | -0.4202586 | 0.67429655 |
| Xu_2009_SUTs46 | Xu_2009_SU | 190.085185 | 0.05027684 | 0.11980548 | 0.41965389 | 0.67473832 |
| Xu_2009_CUTs52 | Xu_2009_CU | 15.7866021 | -0.1168935 | 0.27909352 | -0.4188327 | 0.67533841 |
| YIL089W | YIL089W | 72.8978475 | 0.075186 | 0.17950334 | 0.41885569 | 0.6753216 |
| YBR012C | YBR012C | 82.5448023 | -0.0748257 | 0.17900558 | -0.4180079 | 0.67594134 |
| YAK1 | YJL141C | 1677.45845 | -0.0296348 | 0.07094668 | -0.4177054 | 0.67616251 |
| YPR077C | YPR077C | 0.98999233 | 0.07722266 | 0.18497979 | 0.41746537 | 0.67633805 |
| Xu_2009_SUTs48 | Xu_2009_SU | 58.6544158 | 0.08394587 | 0.20124711 | 0.41712833 | 0.67658454 |
| BFA1 | YJR053W | 286.968453 | 0.04777034 | 0.11449514 | 0.41722592 | 0.67651317 |
| Xu_2009_CUTs13 | Xu_2009_CU | 37.7648331 | 0.09616675 | 0.23080389 | 0.41666 | 0.67692712 |
| YJR071W | YJR071W | 1.9804957 | 0.09580515 | 0.22992515 | 0.41667971 | 0.6769127 |
| DAL80 | YKR034W | 34.7201663 | 0.09593952 | 0.23015173 | 0.41685336 | 0.67678567 |
| YCK1 | YHR135C | 1693.51792 | 0.02602203 | 0.06250639 | 0.41630996 | 0.6771832 |
| Xu_2009_CUTs54 | Xu_2009_CU | 38.2613299 | 0.09443731 | 0.22702893 | 0.41597036 | 0.67743169 |
| MSC7 | YHR039C | 1413.77607 | -0.0320277 | 0.07709046 | -0.4154557 | 0.67780831 |
| Xu_2009_SUTs60 | Xu_2009_SU | 63.7724216 | -0.0778337 | 0.1874782 | -0.4151615 | 0.67802369 |
| CUP2 | YGL166W | 543.413467 | -0.0405386 | 0.09771041 | -0.4148849 | 0.67822613 |
| RRN9 | YMR270C | 582.123462 | -0.0349956 | 0.08432461 | -0.4150108 | 0.678134 |
| NAT5 | YOR253W | 465.033853 | -0.0406322 | 0.09791064 | -0.4149923 | 0.67814751 |
| MSC2 | YDR205W | 612.35561 | 0.03282224 | 0.07913576 | 0.41475864 | 0.6783186 |
| tD(GUC)B | tD(GUC)B | 1.77561396 | 0.08969885 | 0.21664023 | 0.41404519 | 0.67884101 |
| YHL030W-A | YHL030W-A | 5.22648032 | 0.11882473 | 0.28725073 | 0.41366205 | 0.67912162 |
| HEL1 | YKR017C | 415.554404 | -0.0378551 | 0.09155877 | -0.4134517 | 0.67927571 |
| YML089C | YML089C | 7.46087265 | -0.1224745 | 0.29655433 | -0.4129917 | 0.6796127 |
| YCR013C | YCR013C | 11.2397883 | 0.12201993 | 0.29594363 | 0.41230802 | 0.68011367 |
| CTK1 | YKL139W | 551.289725 | 0.03668696 | 0.08899696 | 0.41222712 | 0.68017296 |
| Xu_2009_CUTs88 | Xu_2009_CU | 10.4969932 | 0.12014495 | 0.29198391 | 0.41147799 | 0.68072207 |
| GEP3 | YOR205C | 957.824957 | -0.0281666 | 0.068466 | -0.4113952 | 0.68078277 |
| Xu_2009_SUTs12 | Xu_2009_SU | 198.364155 | 0.04990368 | 0.12141868 | 0.41100496 | 0.6810689 |

| | | | | | | |
|---|---|---|---|---|---|---|
| YAR023C | YAR023C | 49.5605765 | -0.0843988 | 0.20533304 | -0.4110338 | 0.68104778 |
| Xu_2009_SUTs38 | Xu_2009_SU | 2.61443376 | -0.1044218 | 0.25423641 | -0.4107271 | 0.68127262 |
| HXT15 | YDL245C | 1.77360399 | 0.09177208 | 0.2236831 | 0.41027723 | 0.68160259 |
| RSC1 | YGR056W | 1158.2209 | 0.02628096 | 0.06411249 | 0.40991948 | 0.68186501 |
| ASM4 | YDL088C | 894.690838 | -0.0323669 | 0.07906846 | -0.4093532 | 0.68228049 |
| YDR467C | YDR467C | 21.1030059 | -0.113237 | 0.27661291 | -0.4093699 | 0.68226826 |
| MRH4 | YGL064C | 553.899879 | -0.0327641 | 0.08021976 | -0.4084295 | 0.68295835 |
| YHR202W | YHR202W | 269.385502 | 0.04570423 | 0.11203539 | 0.40794461 | 0.68331434 |
| ERG8 | YMR220W | 849.078908 | 0.02951382 | 0.07238861 | 0.40771357 | 0.68348397 |
| Xu_2009_CUTs88 | Xu_2009_CU | 23.3363896 | 0.10729779 | 0.26392916 | 0.4065401 | 0.6843458 |
| YDR401W | YDR401W | 34.4735977 | -0.0982642 | 0.24164355 | -0.4066494 | 0.68426553 |
| RUF22 | RUF22 | 1.21822591 | -0.0846434 | 0.20831932 | -0.4063157 | 0.68451066 |
| EAF5 | YEL018W | 253.469242 | -0.0458227 | 0.11282031 | -0.4061563 | 0.68462777 |
| GFD1 | YMR255W | 853.416417 | -0.0282915 | 0.06970006 | -0.4059036 | 0.68481346 |
| UFE1 | YOR075W | 303.090666 | -0.0450453 | 0.11098255 | -0.4058773 | 0.68483275 |
| YPL152W-A | YPL152W-A | 4.16637028 | -0.1138222 | 0.2804124 | -0.4059102 | 0.68480862 |
| YDR509W | YDR509W | 0.85212813 | 0.06674649 | 0.16476699 | 0.40509622 | 0.68540679 |
| YDR250C | YDR250C | 12.0490147 | 0.11646206 | 0.28785371 | 0.40458767 | 0.68578062 |
| YOR105W | YOR105W | 6.46429286 | 0.11852212 | 0.29292804 | 0.40461174 | 0.68576293 |
| UBP9 | YER098W | 572.926013 | 0.03312753 | 0.08198746 | 0.40405609 | 0.68617147 |
| YLR111W | YLR111W | 4.96426129 | -0.1140589 | 0.28271421 | -0.4034423 | 0.68662286 |
| BDF1 | YLR399C | 2743.98598 | 0.02655253 | 0.06580888 | 0.40347944 | 0.68659556 |
| Xu_2009_SUTs41 | Xu_2009_SU | 21.2261983 | -0.1052275 | 0.26118793 | -0.4028803 | 0.68703628 |
| SUA5 | YGL169W | 541.1461 | -0.0345324 | 0.08577783 | -0.40258 | 0.68725721 |
| YOL013W-A | YOL013W-A | 2.47975887 | -0.0999608 | 0.24843902 | -0.4023555 | 0.68742243 |
| YOR296W | YOR296W | 617.551924 | -0.0340631 | 0.08463373 | -0.4024764 | 0.68733346 |
| ECM12 | YHR021W-A | 27.0175894 | 0.10571503 | 0.26283488 | 0.40221082 | 0.68752888 |
| YNL033W | YNL033W | 9.56064073 | 0.11842185 | 0.29479647 | 0.40170715 | 0.68789956 |
| HAS1 | YMR290C | 2349.83521 | -0.0309284 | 0.07705448 | -0.4013834 | 0.68813787 |
| YNL028W | YNL028W | 5.98684546 | 0.11706799 | 0.29178033 | 0.40121962 | 0.68825844 |
| YER088W-B | YER088W-B | 20.1568332 | 0.11282871 | 0.2815792 | 0.40069975 | 0.6886412 |
| ECM30 | YLR436C | 1525.12527 | -0.0258507 | 0.06454371 | -0.4005147 | 0.68877747 |
| Xu_2009_CUTs69 | Xu_2009_CU | 6.92944986 | 0.11858408 | 0.29618654 | 0.4003696 | 0.68888431 |
| MET1 | YKR069W | 299.109597 | -0.0413803 | 0.10338319 | -0.4002617 | 0.68896375 |
| Xu_2009_SUTs49 | Xu_2009_SU | 82.9296616 | 0.07092657 | 0.17733764 | 0.39995215 | 0.68919176 |
| DUS4 | YLR405W | 374.768843 | -0.0372618 | 0.093266 | -0.3995222 | 0.68950844 |
| RIB7 | YBR153W | 351.754792 | -0.043189 | 0.10826875 | -0.3989054 | 0.68996288 |
| RCM1 | YNL022C | 1005.53585 | 0.02777023 | 0.06963301 | 0.39880843 | 0.69003437 |
| Xu_2009_SUTs10 | Xu_2009_SU | 34.3205038 | -0.092253 | 0.23147346 | -0.398547 | 0.69022704 |
| tT(AGU)N2 | tT(AGU)N2 | 1.5907951 | -0.0820336 | 0.20692314 | -0.3964448 | 0.69177693 |
| Xu_2009_CUTs37 | Xu_2009_CU | 1.68331007 | 0.08912489 | 0.22489083 | 0.39630291 | 0.69188158 |

| | | | | | | |
|---|---|---|---|---|---|---|
| YGR219W | YGR219W | 1.49720266 | -0.0847315 | 0.21393987 | -0.3960528 | 0.69206606 |
| SPP1 | YPL138C | 504.724469 | 0.03584157 | 0.09052715 | 0.39592072 | 0.69216352 |
| STS1 | YIR011C | 1420.36541 | -0.0246695 | 0.06243148 | -0.395145 | 0.69273587 |
| YGL193C | YGL193C | 266.28371 | -0.0433082 | 0.109834 | -0.3943063 | 0.69335492 |
| THI7 | YLR237W | 1629.77961 | -0.0278208 | 0.0706967 | -0.3935239 | 0.69393257 |
| Xu_2009_CUTs37 | Xu_2009_CU | 5.79343705 | -0.115206 | 0.29334088 | -0.3927375 | 0.69451337 |
| AAD16 | YFL057C | 99.8905658 | 0.06413821 | 0.16331921 | 0.39271684 | 0.69452863 |
| BIO3 | YNR058W | 383.39618 | -0.041162 | 0.1048654 | -0.3925224 | 0.69467228 |
| Xu_2009_SUTs56 | Xu_2009_SU | 41.0468035 | 0.08554696 | 0.2181862 | 0.39208235 | 0.69499737 |
| PAU12 | YGR294W | 62.7955875 | -0.0766807 | 0.19557323 | -0.3920818 | 0.69499781 |
| tT(AGU)O2/TRT2 | tT(AGU)O2 | 1.95450089 | -0.0896187 | 0.22900617 | -0.3913374 | 0.69554783 |
| CSS2 | YFR020W | 222.102341 | -0.0483247 | 0.12359681 | -0.3909866 | 0.69580712 |
| ECM9 | YKR004C | 181.080429 | -0.0510023 | 0.13055921 | -0.3906451 | 0.69605959 |
| Xu_2009_CUTs63 | Xu_2009_CU | 4.51473242 | 0.10828675 | 0.27729854 | 0.39050603 | 0.6961624 |
| UBS1 | YBR165W | 423.354825 | -0.0377382 | 0.0968917 | -0.3894882 | 0.69691505 |
| YPL108W | YPL108W | 151.941719 | 0.05635073 | 0.14474807 | 0.38930213 | 0.69705266 |
| PIG2 | YIL045W | 227.555212 | -0.048309 | 0.12425627 | -0.3887856 | 0.69743477 |
| VPS53 | YJL029C | 1088.99166 | -0.0275195 | 0.07086317 | -0.3883464 | 0.6977597 |
| Xu_2009_CUTs66 | Xu_2009_CU | 5.49593796 | 0.1121344 | 0.2891262 | 0.38783895 | 0.69813522 |
| YIL102C | YIL102C | 1.09761587 | -0.0702114 | 0.18125841 | -0.3873555 | 0.69849305 |
| UIP4 | YPL186C | 300.228217 | -0.042159 | 0.1091429 | -0.3862735 | 0.69929412 |
| Xu_2009_SUTs23 | Xu_2009_SU | 6.21239779 | 0.11284399 | 0.29237433 | 0.38595724 | 0.69952834 |
| FUS3 | YBL016W | 899.441875 | 0.03067363 | 0.07953751 | 0.3856498 | 0.69975604 |
| YNL266W | YNL266W | 5.89718881 | -0.1132077 | 0.29348656 | -0.385734 | 0.69969367 |
| TMC1 | YOR052C | 2568.74998 | -0.0218249 | 0.05652869 | -0.3860858 | 0.69943312 |
| YPR126C | YPR126C | 1.08498489 | -0.0749028 | 0.19423842 | -0.3856232 | 0.69977577 |
| TLG2 | YOL018C | 488.38622 | 0.03291187 | 0.0854224 | 0.38528386 | 0.70002712 |
| ERR1 | YOR393W | 2.70475176 | -0.0981855 | 0.25476193 | -0.3854008 | 0.69994047 |
| Xu_2009_CUTs16 | Xu_2009_CU | 58.5677168 | -0.0762203 | 0.19825515 | -0.3844558 | 0.70064063 |
| REV7 | YIL139C | 179.414665 | -0.0480109 | 0.12492367 | -0.3843219 | 0.70073986 |
| Xu_2009_CUTs19 | Xu_2009_CU | 7.87076676 | 0.11300689 | 0.29629139 | 0.38140455 | 0.70290308 |
| YDR340W | YDR340W | 15.9338855 | -0.1060446 | 0.27811639 | -0.3812959 | 0.7029837 |
| Xu_2009_CUTs8 | Xu_2009_CU | 16.0584809 | 0.10640162 | 0.27925531 | 0.38101915 | 0.70318904 |
| YIL142C-A | YIL142C-A | 2.14403347 | 0.09399478 | 0.24680796 | 0.38084178 | 0.70332065 |
| PTC4 | YBR125C | 864.976737 | 0.02861559 | 0.07529907 | 0.38002576 | 0.70392629 |
| SNR72 | snR72 | 41.2102754 | 0.08192196 | 0.21590289 | 0.3794389 | 0.70436197 |
| Xu_2009_SUTs23 | Xu_2009_SU | 82.0932756 | 0.06653589 | 0.17555651 | 0.37899987 | 0.70468796 |
| Xu_2009_SUTs36 | Xu_2009_SU | 13.4481241 | 0.10835541 | 0.28611029 | 0.37871901 | 0.70489654 |
| SNZ2 | YNL333W | 191.508182 | -0.048553 | 0.12813827 | -0.3789114 | 0.70475368 |
| VPS16 | YPL045W | 666.942074 | 0.03307897 | 0.08732723 | 0.37879325 | 0.7048414 |
| SUL1 | YBR294W | 9.6157392 | -0.1117722 | 0.29525976 | -0.3785554 | 0.70501803 |

| | | | | | | |
|---|---|---|---|---|---|---|
| YEL010W | YEL010W | 3.09764011 | 0.10146399 | 0.26814989 | 0.37838536 | 0.70514434 |
| YPL073C | YPL073C | 8.41649022 | 0.11216732 | 0.29650142 | 0.37830282 | 0.70520565 |
| SNR53 | snR53 | 2.46490116 | 0.09521016 | 0.25204531 | 0.37775019 | 0.70561618 |
| Xu_2009_CUTs72 | Xu_2009_CU | 1.01170886 | -0.0735658 | 0.19499926 | -0.3772622 | 0.70597879 |
| RTT109 | YLL002W | 175.725614 | 0.04966446 | 0.13169676 | 0.37711221 | 0.70609022 |
| POL1 | YNL102W | 1102.03507 | 0.02523019 | 0.06705567 | 0.37625737 | 0.70672557 |
| Xu_2009_SUTs41 | Xu_2009_SU | 4.14254169 | 0.10253058 | 0.27263708 | 0.37606982 | 0.70686499 |
| FIR1 | YER032W | 655.087843 | -0.0312629 | 0.08324185 | -0.3755665 | 0.70723919 |
| Xu_2009_CUTs75 | Xu_2009_CU | 3.92261328 | 0.10439985 | 0.27831117 | 0.37511917 | 0.70757184 |
| Xu_2009_SUTs20 | Xu_2009_SU | 12.2076221 | -0.1090042 | 0.29080683 | -0.3748338 | 0.70778409 |
| REC114 | YMR133W | 7.94731142 | 0.11102301 | 0.29616123 | 0.37487355 | 0.70775451 |
| Xu_2009_SUTs82 | Xu_2009_SU | 5.77637574 | 0.10986948 | 0.29346543 | 0.37438645 | 0.70811682 |
| YJL070C | YJL070C | 516.789752 | -0.0355578 | 0.09502953 | -0.3741761 | 0.70827329 |
| ARR3 | YPR201W | 70.7639722 | 0.06835101 | 0.18269614 | 0.37412398 | 0.70831208 |
| AIM3 | YBR108W | 940.506488 | -0.028582 | 0.07657237 | -0.3732677 | 0.70894925 |
| Xu_2009_CUTs80 | Xu_2009_CU | 2.05337926 | -0.0874279 | 0.23442238 | -0.3729504 | 0.70918539 |
| Xu_2009_SUTs18 | Xu_2009_SU | 16.7298362 | -0.1049122 | 0.28142501 | -0.3727893 | 0.70930523 |
| SIN3 | YOL004W | 4355.5756 | 0.01852097 | 0.04975641 | 0.37223281 | 0.70971952 |
| Xu_2009_CUTs31 | Xu_2009_CU | 9.25082617 | 0.11004847 | 0.29623458 | 0.37149096 | 0.71027189 |
| NOP13 | YNL175C | 1098.89914 | -0.0292976 | 0.07885085 | -0.3715578 | 0.71022214 |
| Xu_2009_CUTs52 | Xu_2009_CU | 33.7799286 | 0.09151704 | 0.24676456 | 0.37086783 | 0.71073597 |
| Xu_2009_SUTs42 | Xu_2009_SU | 42.5448875 | 0.087847 | 0.23676039 | 0.37103758 | 0.71060954 |
| YFL063W | YFL063W | 0.78899112 | 0.0611874 | 0.16495234 | 0.37093988 | 0.71068231 |
| RTG2 | YGL252C | 847.653602 | 0.02824872 | 0.07616316 | 0.37089735 | 0.71071399 |
| Xu_2009_SUTs68 | Xu_2009_SU | 11.0036213 | -0.1087229 | 0.29351743 | -0.3704137 | 0.7110743 |
| YOL014W | YOL014W | 30.2172792 | -0.0883782 | 0.23851486 | -0.3705353 | 0.71098369 |
| YGL102C | YGL102C | 3.0863546 | 0.09577368 | 0.25898832 | 0.36979921 | 0.7115321 |
| YJR142W | YJR142W | 498.741228 | -0.0358287 | 0.09708692 | -0.3690373 | 0.71209995 |
| SWM2 | YNR004W | 100.823716 | -0.0614661 | 0.16653828 | -0.3690806 | 0.71206761 |
| Xu_2009_SUTs60 | Xu_2009_SU | 133.971081 | -0.0530576 | 0.14382754 | -0.3688974 | 0.7122042 |
| Xu_2009_CUTs10 | Xu_2009_CU | 4.9772347 | 0.10543385 | 0.28717744 | 0.36713834 | 0.71351583 |
| TMA22 | YJR014W | 1824.71612 | 0.02429364 | 0.06618969 | 0.36703052 | 0.71359626 |
| YBR196C-B | YBR196C-B | 16.8271558 | 0.10489666 | 0.28593337 | 0.36685702 | 0.71372567 |
| YFL032W | YFL032W | 4.06517568 | 0.10326174 | 0.28187722 | 0.36633589 | 0.71411445 |
| YLR342W-A | YLR342W-A | 77.9868256 | 0.06440264 | 0.17576633 | 0.36641058 | 0.71405873 |
| YDL162C | YDL162C | 8.38804834 | 0.1081908 | 0.29594733 | 0.36557451 | 0.7146826 |
| Xu_2009_SUTs57 | Xu_2009_SU | 12.1929125 | -0.1063076 | 0.29102691 | -0.3652845 | 0.71489901 |
| AIM14 | YGL160W | 740.444383 | 0.0326108 | 0.0893845 | 0.36483726 | 0.71523289 |
| tA(AGC)G | tA(AGC)G | 0.7524086 | 0.05706234 | 0.15646743 | 0.3646915 | 0.71534171 |
| tN(GUU)F | tN(GUU)F | 2.1870659 | -0.08913 | 0.24471701 | -0.3642165 | 0.71569633 |
| Xu_2009_CUTs12 | Xu_2009_CU | 32.4776757 | -0.0871951 | 0.23925707 | -0.364441 | 0.71552876 |

| | | | | | | |
|---|---|---|---|---|---|---|
| FMP45 | YDL222C | 161.706983 | 0.04733568 | 0.13000148 | 0.36411648 | 0.71577103 |
| YGR190C | YGR190C | 2.86712468 | 0.09135285 | 0.25083196 | 0.3641994 | 0.71570912 |
| MIN10 | YFR032C-B | 73.858118 | 0.0639612 | 0.17581537 | 0.36379754 | 0.7160092 |
| YOR024W | YOR024W | 2.42318888 | -0.0915246 | 0.25176925 | -0.3635257 | 0.71621219 |
| YGL015C | YGL015C | 13.2583931 | -0.1045163 | 0.28801084 | -0.3628902 | 0.71668691 |
| tG(GCC)B | tG(GCC)B | 1.08121348 | -0.0689203 | 0.19028881 | -0.362188 | 0.71721151 |
| Xu_2009_SUTs73( | Xu_2009_SU' | 76.575137 | -0.0662744 | 0.1830614 | -0.3620338 | 0.71732678 |
| CKA2 | YOR061W | 1745.21299 | -0.0218222 | 0.06026437 | -0.3621078 | 0.71727145 |
| Xu_2009_CUTs46 | Xu_2009_CU | 4.81698569 | -0.1022941 | 0.28299139 | -0.3614741 | 0.71774506 |
| BUR2 | YLR226W | 649.25445 | -0.0319808 | 0.08853722 | -0.3612127 | 0.71794045 |
| DSF1 | YEL070W | 35.1771598 | 0.08244729 | 0.22842515 | 0.36093787 | 0.71814589 |
| tQ(CUG)M/CDC65 | tQ(CUG)M | 5.00688104 | -0.1028059 | 0.28505923 | -0.3606476 | 0.71836292 |
| ADP1 | YCR011C | 1691.38931 | 0.02462003 | 0.06838278 | 0.36003261 | 0.71882275 |
| YLR415C | YLR415C | 34.5756409 | 0.08624567 | 0.23965121 | 0.35987998 | 0.71893689 |
| tQ(UUG)E2 | tQ(UUG)E2 | 2.40456361 | -0.0837391 | 0.23351709 | -0.3585993 | 0.71989487 |
| YLR031W | YLR031W | 62.9107619 | -0.0678996 | 0.18996312 | -0.3574356 | 0.72076569 |
| BIM1 | YER016W | 471.243861 | 0.03339218 | 0.09369591 | 0.35638886 | 0.72154937 |
| SNZ3 | YFL059W | 199.652163 | 0.04343955 | 0.12210767 | 0.35574794 | 0.72202934 |
| YOR282W | YOR282W | 2.19104716 | -0.0831006 | 0.23354922 | -0.3558162 | 0.72197825 |
| YRF1-1 | YDR545W | 2.33409189 | 0.08970017 | 0.25229197 | 0.3555411 | 0.72218426 |
| MAG2 | YLR427W | 595.882405 | 0.03126618 | 0.08803853 | 0.35514199 | 0.72248322 |
| Xu_2009_CUTs36 | Xu_2009_CU | 13.0319537 | -0.1028838 | 0.28989025 | -0.3549062 | 0.72265989 |
| Xu_2009_SUTs56 | Xu_2009_SU' | 2.2945647 | 0.08657845 | 0.24432534 | 0.35435721 | 0.7230712 |
| ULP1 | YPL020C | 759.260024 | 0.02572177 | 0.07267186 | 0.35394403 | 0.72338083 |
| UPS3 | YDR185C | 123.342461 | 0.0508891 | 0.14412321 | 0.35309441 | 0.72401766 |
| YDR521W | YDR521W | 1.23470405 | 0.07254292 | 0.206055 | 0.35205611 | 0.72479619 |
| UBA3 | YPR066W | 126.837459 | -0.0513084 | 0.14574332 | -0.3520461 | 0.72480372 |
| YDR246W-A | YDR246W-A | 30.2871226 | -0.086989 | 0.2472353 | -0.3518469 | 0.72495311 |
| PSP2 | YML017W | 1897.54548 | -0.0211623 | 0.06042844 | -0.3502042 | 0.72618546 |
| YLL020C | YLL020C | 4.63113354 | 0.09977923 | 0.2855589 | 0.34941733 | 0.72677603 |
| Xu_2009_SUTs49 | Xu_2009_SU' | 17.6301009 | 0.09479495 | 0.27156634 | 0.34906739 | 0.72703872 |
| tD(GUC)M | tD(GUC)M | 5.11497142 | -0.1003343 | 0.28781089 | -0.3486117 | 0.72738081 |
| CDC26 | YFR036W | 497.731111 | -0.0337402 | 0.09682074 | -0.3484809 | 0.72747904 |
| Xu_2009_CUTs81 | Xu_2009_CU | 8.58185446 | 0.10222455 | 0.29632131 | 0.34497873 | 0.73011036 |
| OCA4 | YCR095C | 349.486613 | -0.0357718 | 0.10372744 | -0.3448634 | 0.73019709 |
| MF(ALPHA)2 | YGL089C | 56.7123052 | -0.0684688 | 0.198634 | -0.3446983 | 0.73032119 |
| MHF2 | YDL160C-A | 108.269882 | -0.0541122 | 0.15715062 | -0.3443335 | 0.73059551 |
| Xu_2009_SUTs36 | Xu_2009_SU' | 12.5964149 | 0.10025495 | 0.29162066 | 0.34378549 | 0.73100761 |
| Xu_2009_CUTs51 | Xu_2009_CU | 2.35166259 | -0.0842071 | 0.24640634 | -0.3417408 | 0.732546 |
| YHL042W | YHL042W | 26.2027793 | -0.0870596 | 0.25496402 | -0.3414586 | 0.7327584 |
| ITT1 | YML068W | 230.451923 | 0.04319361 | 0.12645495 | 0.34157314 | 0.73267215 |

| | | | | | | |
|---|---|---|---|---|---|---|
| Xu_2009_SUTs33( | Xu_2009_SU | 53.1957134 | 0.06815962 | 0.19996776 | 0.34085307 | 0.7332142 |
| YCR049C | YCR049C | 0.71428019 | -0.0555305 | 0.16311684 | -0.3404341 | 0.7335296 |
| Xu_2009_CUTs75 | Xu_2009_CU | 49.7904787 | -0.0761817 | 0.22403655 | -0.3400413 | 0.73382539 |
| RRP3 | YHR065C | 1251.59564 | -0.0260772 | 0.07688743 | -0.3391614 | 0.73448815 |
| YKE2 | YLR200W | 840.913592 | 0.02544678 | 0.07500479 | 0.33926871 | 0.73440731 |
| tQ(UUG)D3 | tQ(UUG)D3 | 1.56053467 | -0.0726223 | 0.21462698 | -0.3383651 | 0.73508805 |
| YER090C-A | YER090C-A | 1.12821658 | -0.0659362 | 0.19578117 | -0.3367854 | 0.73627871 |
| Xu_2009_SUTs43! | Xu_2009_SU | 2.73114774 | 0.08547025 | 0.25397151 | 0.3365348 | 0.73646761 |
| BET2 | YPR176C | 427.157632 | 0.0301585 | 0.08959525 | 0.33660821 | 0.73641226 |
| HUA2 | YOR284W | 138.906212 | 0.04852234 | 0.14425694 | 0.33636055 | 0.736599 |
| YKE4 | YIL023C | 586.151533 | 0.03104436 | 0.09236769 | 0.33609541 | 0.73679892 |
| KAR1 | YNL188W | 476.654696 | -0.0302849 | 0.09009442 | -0.3361467 | 0.73676025 |
| YCT1 | YLL055W | 189.456225 | -0.0412742 | 0.1235059 | -0.3341877 | 0.73823793 |
| Xu_2009_CUTs75 | Xu_2009_CU | 13.3600199 | 0.09808851 | 0.29379543 | 0.33386669 | 0.73848016 |
| Xu_2009_SUTs76 | Xu_2009_SU | 24.1913045 | -0.0879827 | 0.26415281 | -0.333075 | 0.7390777 |
| YPL205C | YPL205C | 4.00431239 | 0.09236 | 0.27755986 | 0.33275706 | 0.73931767 |
| SLM2 | YNL047C | 710.707061 | -0.0283298 | 0.08519806 | -0.3325174 | 0.73949861 |
| IME1 | YJR094C | 167.483944 | 0.04521145 | 0.13626804 | 0.33178324 | 0.74005294 |
| HXT5 | YHR096C | 210.436292 | 0.04101266 | 0.12377687 | 0.33134352 | 0.74038502 |
| NDC1 | YML031W | 1433.76867 | -0.021481 | 0.0647933 | -0.3315316 | 0.74024301 |
| PPG1 | YNR032W | 489.893986 | -0.0283786 | 0.08565995 | -0.3312943 | 0.74042223 |
| tF(GAA)F | tF(GAA)F | 3.93824458 | 0.09170486 | 0.2775649 | 0.33039069 | 0.74110477 |
| YFL021C-A | YFL021C-A | 15.9619743 | 0.09268302 | 0.28059168 | 0.33031277 | 0.74116364 |
| POP3 | YNL282W | 189.134376 | 0.04133394 | 0.12556972 | 0.32917123 | 0.74202627 |
| Xu_2009_SUTs66! | Xu_2009_SU | 23.3708389 | 0.08559383 | 0.26040268 | 0.32869797 | 0.74238399 |
| WTM2 | YOR229W | 617.441631 | -0.0252972 | 0.07701083 | -0.3284882 | 0.74254255 |
| RTR1 | YER139C | 270.450018 | -0.0385249 | 0.11742847 | -0.3280712 | 0.7428578 |
| tE(UUC)G2 | tE(UUC)G2 | 2.36515582 | -0.0775197 | 0.23676291 | -0.3274147 | 0.74335424 |
| NPR2 | YEL062W | 545.981842 | -0.0279999 | 0.08575179 | -0.3265231 | 0.74402859 |
| GGA1 | YDR358W | 588.64392 | -0.0342416 | 0.10497741 | -0.3261809 | 0.74428748 |
| Xu_2009_SUTs74! | Xu_2009_SU | 11.0874764 | 0.09558417 | 0.29317215 | 0.32603426 | 0.74439843 |
| KRE28 | YDR532C | 338.194267 | 0.03099632 | 0.09540352 | 0.324897 | 0.74525903 |
| tF(GAA)N | tF(GAA)N | 10.6245972 | -0.095359 | 0.29409595 | -0.3242445 | 0.74575295 |
| PPS1 | YBR276C | 933.365708 | -0.0223203 | 0.06891674 | -0.3238728 | 0.74603431 |
| PUT3 | YKL015W | 1231.70694 | 0.0253059 | 0.07818639 | 0.32366122 | 0.74619453 |
| ECM18 | YDR125C | 164.133172 | 0.04625883 | 0.14315345 | 0.32314158 | 0.74658802 |
| NTO1 | YPR031W | 386.121844 | 0.03371679 | 0.10430812 | 0.32324229 | 0.74651176 |
| IME4 | YGL192W | 2.94119031 | -0.0809872 | 0.25078228 | -0.3229384 | 0.74674192 |
| YOL029C | YOL029C | 269.096256 | 0.03532557 | 0.10962891 | 0.32222857 | 0.74727954 |
| GSH2 | YOL049W | 1381.85197 | 0.01999915 | 0.0621326 | 0.32187853 | 0.74754472 |
| Xu_2009_SUTs18! | Xu_2009_SU | 46.1784358 | 0.06680569 | 0.2085853 | 0.32027997 | 0.7487561 |

| | | | | | | |
|---|---|---|---|---|---|---|
| EMP65 | YER140W | 351.323607 | 0.03094978 | 0.09659718 | 0.32040046 | 0.74866477 |
| MRX11 | YPL041C | 125.008059 | -0.0461001 | 0.14400295 | -0.3201332 | 0.74886733 |
| tT(UGU)G2 | tT(UGU)G2 | 43.3167726 | -0.0732539 | 0.22945699 | -0.3192488 | 0.74953788 |
| Xu_2009_SUTs25 | Xu_2009_SU | 10.3632335 | -0.0937165 | 0.29351477 | -0.3192904 | 0.74950629 |
| POP7 | YBR167C | 170.788869 | -0.0428497 | 0.13407742 | -0.319589 | 0.7492799 |
| SEG1 | YMR086W | 1314.52464 | 0.019623 | 0.06144954 | 0.3193352 | 0.74947234 |
| Xu_2009_CUTs69 | Xu_2009_CU | 8.17139353 | -0.0945502 | 0.29659274 | -0.3187881 | 0.74988723 |
| AAD4 | YDL243C | 111.099784 | 0.04885224 | 0.15338787 | 0.3184883 | 0.75011457 |
| Xu_2009_SUTs50 | Xu_2009_SU | 60.3237263 | -0.0609575 | 0.19175228 | -0.3178971 | 0.75056299 |
| Xu_2009_CUTs53 | Xu_2009_CU | 5.24942556 | 0.09256679 | 0.29139804 | 0.31766442 | 0.75073951 |
| Xu_2009_CUTs50 | Xu_2009_CU | 11.6520373 | 0.09236191 | 0.29158581 | 0.31675721 | 0.75142784 |
| Xu_2009_CUTs86 | Xu_2009_CU | 13.7418983 | -0.0907265 | 0.28671639 | -0.3164328 | 0.75167403 |
| YDR220C | YDR220C | 27.3602197 | -0.0800594 | 0.25302347 | -0.316411 | 0.75169057 |
| YIL058W | YIL058W | 10.3544749 | 0.09329887 | 0.29567451 | 0.31554588 | 0.75234723 |
| Xu_2009_SUTs50 | Xu_2009_SU | 15.2098314 | 0.08876206 | 0.28176621 | 0.31502025 | 0.75274628 |
| Xu_2009_SUTs54 | Xu_2009_SU | 16.2945971 | 0.09061425 | 0.28746353 | 0.31521999 | 0.75259463 |
| SEM1 | YDR363W-A | 1775.49462 | -0.0279839 | 0.08887379 | -0.3148726 | 0.75285836 |
| COG7 | YGL005C | 392.703532 | 0.03601469 | 0.11439211 | 0.31483548 | 0.75288657 |
| PUT1 | YLR142W | 397.903231 | -0.0334164 | 0.10616909 | -0.3147471 | 0.7529537 |
| Xu_2009_CUTs54 | Xu_2009_CU | 4.60121337 | 0.09012374 | 0.28659041 | 0.3144688 | 0.75316501 |
| DIF1 | YLR437C | 415.347804 | -0.0309202 | 0.09846085 | -0.3140359 | 0.7534938 |
| VEL1 | YGL258W | 7.8095454 | -0.0927113 | 0.29626726 | -0.3129312 | 0.7543329 |
| MPS1 | YDL028C | 679.169958 | -0.0236225 | 0.07556595 | -0.3126072 | 0.75457911 |
| DAN1 | YJR150C | 3.14373745 | -0.0832365 | 0.26629761 | -0.3125696 | 0.75460768 |
| YPL136W | YPL136W | 1.93560153 | -0.0751936 | 0.24085361 | -0.3121964 | 0.75489127 |
| Xu_2009_SUTs92 | Xu_2009_SU | 100.117464 | -0.0495423 | 0.15903943 | -0.3115092 | 0.75541352 |
| IMP4 | YNL075W | 1001.03841 | 0.02201209 | 0.07065186 | 0.3115571 | 0.75537714 |
| APL1 | YJR005W | 744.736412 | -0.0229049 | 0.07361834 | -0.3111304 | 0.75570151 |
| YJL009W | YJL009W | 3.92373919 | -0.0867014 | 0.27909741 | -0.3106494 | 0.75606719 |
| MSS4 | YDR208W | 2033.53953 | -0.0195484 | 0.06295565 | -0.3105102 | 0.75617303 |
| FUN19 | YAL034C | 570.93292 | -0.0260505 | 0.08394299 | -0.3103359 | 0.75630552 |
| SDL1 | YIL167W | 27.9147931 | 0.07723877 | 0.2493019 | 0.30982021 | 0.75669768 |
| PET122 | YER153C | 131.264284 | -0.0482927 | 0.1567354 | -0.3081162 | 0.75799392 |
| UBP11 | YKR098C | 346.463383 | 0.03040977 | 0.09870798 | 0.30807816 | 0.75802286 |
| YBR113W | YBR113W | 4.61804266 | 0.08563284 | 0.27850314 | 0.30747531 | 0.75848161 |
| Xu_2009_SUTs19 | Xu_2009_SU | 19.4971892 | -0.0856816 | 0.27878879 | -0.307335 | 0.75858839 |
| PRP43 | YGL120C | 2526.15169 | -0.0238493 | 0.07767196 | -0.3070521 | 0.75880374 |
| CWC27 | YPL064C | 545.784973 | -0.0264675 | 0.08637331 | -0.3064318 | 0.75927592 |
| MMP1 | YLL061W | 84.0658914 | 0.05195184 | 0.16963353 | 0.30625925 | 0.75940726 |
| SDC25 | YLL016W | 341.879687 | -0.0320119 | 0.10481799 | -0.3054046 | 0.76005799 |
| YGL042C | YGL042C | 4.6527852 | -0.086713 | 0.28537938 | -0.3038516 | 0.76124094 |

| | | | | | | |
|---|---|---|---|---|---|---|
| AEP1 | YMR064W | 642.237322 | -0.0230768 | 0.07597754 | -0.3037319 | 0.76133214 |
| BEM3 | YPL115C | 1053.20599 | 0.02323839 | 0.07654068 | 0.30360832 | 0.76142631 |
| NBP1 | YLR457C | 182.037667 | 0.04106406 | 0.13541814 | 0.30323896 | 0.76170776 |
| GAL2 | YLR081W | 8.36895587 | 0.08955726 | 0.29628868 | 0.30226351 | 0.76245119 |
| URE2 | YNL229C | 993.354338 | -0.0201312 | 0.06660932 | -0.3022277 | 0.76247848 |
| RSA1 | YPL193W | 686.303342 | 0.02565121 | 0.08482891 | 0.30238758 | 0.76235662 |
| tW(CCA)K | tW(CCA)K | 7.61960301 | -0.0894684 | 0.29642043 | -0.3018294 | 0.76278215 |
| SSK1 | YLR006C | 813.602234 | 0.02197239 | 0.07278975 | 0.30186106 | 0.76275798 |
| SPO22 | YIL073C | 195.650217 | 0.03726984 | 0.12354454 | 0.30167125 | 0.76290269 |
| Xu_2009_SUTs39 | Xu_2009_SU | 4.00002014 | 0.08441007 | 0.28065861 | 0.30075711 | 0.76359972 |
| LST7 | YGR057C | 343.161694 | -0.0334943 | 0.11133743 | -0.300836 | 0.76353959 |
| CBF5 | YLR175W | 5345.91265 | 0.01948792 | 0.06489517 | 0.30029853 | 0.76394946 |
| Xu_2009_CUTs71 | Xu_2009_CU | 26.830332 | 0.07483777 | 0.24937929 | 0.30009618 | 0.76410379 |
| PRP24 | YMR268C | 387.658486 | 0.02808557 | 0.0936345 | 0.29994899 | 0.76421606 |
| Xu_2009_CUTs72 | Xu_2009_CU | 1.12494897 | 0.05764702 | 0.19341605 | 0.29804675 | 0.76566748 |
| TCD2 | YKL027W | 898.985505 | 0.02313555 | 0.07769937 | 0.29775729 | 0.76588842 |
| Xu_2009_CUTs35 | Xu_2009_CU | 28.6961173 | 0.07370103 | 0.24802626 | 0.2971501 | 0.76635192 |
| Xu_2009_CUTs37 | Xu_2009_CU | 10.7310896 | -0.0873115 | 0.29374446 | -0.2972364 | 0.76628607 |
| YCL023C | YCL023C | 2.55366194 | 0.07375425 | 0.24834641 | 0.29698134 | 0.76648075 |
| YNL165W | YNL165W | 267.692567 | 0.03481669 | 0.11784333 | 0.29544893 | 0.76765096 |
| YPL044C | YPL044C | 1.57626662 | 0.06488733 | 0.21986211 | 0.29512737 | 0.76789658 |
| Xu_2009_CUTs78 | Xu_2009_CU | 17.9892166 | 0.08049684 | 0.27311184 | 0.29473948 | 0.7681929 |
| SLM5 | YCR024C | 403.20181 | 0.02919307 | 0.09901899 | 0.29482291 | 0.76812916 |
| Xu_2009_CUTs44 | Xu_2009_CU | 32.3263629 | -0.0698349 | 0.23713212 | -0.2944977 | 0.76837762 |
| Xu_2009_SUTs50 | Xu_2009_SU | 83.2517813 | -0.0490646 | 0.16760099 | -0.2927462 | 0.76971616 |
| Xu_2009_SUTs24 | Xu_2009_SU | 18.0101126 | 0.08036618 | 0.27475336 | 0.29250298 | 0.76990208 |
| tI(AAU)N1 | tI(AAU)N1 | 1.22792319 | 0.05833211 | 0.2000264 | 0.29162207 | 0.7705756 |
| ESC8 | YOL017W | 237.136223 | 0.03291071 | 0.11301338 | 0.29121077 | 0.77089012 |
| YOL134C | YOL134C | 2.15145611 | -0.0668957 | 0.22967182 | -0.2912664 | 0.77084759 |
| Xu_2009_SUTs12 | Xu_2009_SU | 30.0254157 | 0.07118662 | 0.2450252 | 0.29052776 | 0.77141252 |
| TAX4 | YJL083W | 664.762201 | -0.022849 | 0.07862186 | -0.2906188 | 0.7713429 |
| ETT1 | YOR051C | 1189.77407 | -0.0194456 | 0.06717377 | -0.2894823 | 0.77221236 |
| YHR125W | YHR125W | 0.75367007 | -0.0483345 | 0.16793593 | -0.2878151 | 0.77348825 |
| PAM1 | YDR251W | 1464.45216 | -0.0186463 | 0.06505224 | -0.2866363 | 0.7743908 |
| Xu_2009_CUTs63 | Xu_2009_CU | 3.5491237 | 0.07859399 | 0.27444711 | 0.28637209 | 0.77459315 |
| YOR161W-B | YOR161W-B | 2.20247725 | 0.06847664 | 0.23921347 | 0.28625746 | 0.77468094 |
| Xu_2009_SUTs12 | Xu_2009_SU | 4.76626536 | -0.0827517 | 0.28953835 | -0.2858055 | 0.7750271 |
| YIL115W-A | YIL115W-A | 2.15439568 | 0.06777857 | 0.23718851 | 0.28575824 | 0.77506329 |
| AAD14 | YNL331C | 236.875641 | 0.03194639 | 0.11182775 | 0.28567499 | 0.77512706 |
| FRE7 | YOL152W | 72.1794044 | 0.05125058 | 0.17938164 | 0.28570696 | 0.77510257 |
| ENV7 | YPL236C | 196.088671 | 0.03428541 | 0.12051869 | 0.28448209 | 0.77604096 |

| | | | | | | |
|---|---|---|---|---|---|---|
| YJR114W | YJR114W | 12.8690436 | -0.0809143 | 0.28670079 | -0.2822255 | 0.77777062 |
| Xu_2009_CUTs76 | Xu_2009_CU | 6.9587154 | 0.08171364 | 0.29061809 | 0.28117191 | 0.77857855 |
| STB3 | YDR169C | 1331.36392 | 0.0180642 | 0.06420564 | 0.28134912 | 0.77844264 |
| ATG38 | YLR211C | 185.087516 | -0.0350148 | 0.12451018 | -0.28122 | 0.77854166 |
| RIT1 | YMR283C | 290.494005 | -0.0306 | 0.10884817 | -0.2811253 | 0.77861433 |
| YGL041C | YGL041C | 13.0938656 | -0.0808209 | 0.28807214 | -0.2805579 | 0.7790495 |
| IPK1 | YDR315C | 169.200255 | 0.03689922 | 0.13166915 | 0.28024197 | 0.77929187 |
| ICL1 | YER065C | 336.709333 | 0.02718699 | 0.09708618 | 0.28002945 | 0.77945491 |
| YHR063W-A | YHR063W-A | 1.09526322 | 0.05357269 | 0.19268053 | 0.27803891 | 0.78098249 |
| YDR320W-B | YDR320W-B | 11.4564786 | -0.0814158 | 0.29335206 | -0.2775361 | 0.78136846 |
| PSY1 | YKL076C | 8.57040293 | -0.082329 | 0.29657987 | -0.2775947 | 0.78132349 |
| YOR072W-B | YOR072W-B | 0.80691835 | -0.0471242 | 0.17007685 | -0.2770757 | 0.781722 |
| MRPL51 | YPR100W | 436.442203 | -0.0277161 | 0.10014964 | -0.2767468 | 0.78197456 |
| Xu_2009_SUTs29 | Xu_2009_SU | 78.3197362 | 0.04855639 | 0.17560643 | 0.27650689 | 0.78215877 |
| Xu_2009_SUTs40 | Xu_2009_SU | 21.0283585 | -0.0736089 | 0.26657006 | -0.2761334 | 0.78244557 |
| ATP20 | YPR020W | 254.87013 | -0.0323065 | 0.11726573 | -0.2754978 | 0.78293379 |
| ASI3 | YNL008C | 425.938047 | -0.0247986 | 0.09009712 | -0.2752435 | 0.78312917 |
| CUP9 | YPL177C | 701.411024 | -0.0212814 | 0.07751441 | -0.2745474 | 0.78366395 |
| PMT6 | YGR199W | 456.398387 | -0.0235235 | 0.08588198 | -0.2739054 | 0.78415731 |
| Xu_2009_SUTs54 | Xu_2009_SU | 119.72603 | 0.04130606 | 0.15103558 | 0.27348561 | 0.78447995 |
| GMC2 | YLR445W | 10.3930324 | 0.0809497 | 0.29590923 | 0.2735626 | 0.78442078 |
| NCS2 | YNL119W | 880.908213 | 0.02031262 | 0.07424255 | 0.27359811 | 0.78439348 |
| Xu_2009_CUTs10 | Xu_2009_CU | 4.82211234 | 0.07749831 | 0.28427373 | 0.27261862 | 0.78514639 |
| PKP2 | YGL059W | 365.347389 | -0.0288316 | 0.10583507 | -0.2724198 | 0.78529925 |
| Xu_2009_SUTs43 | Xu_2009_SU | 44.0883895 | -0.058733 | 0.21605497 | -0.2718428 | 0.78574293 |
| HYM1 | YKL189W | 328.82175 | -0.0280288 | 0.10352962 | -0.2707322 | 0.78659702 |
| DNM1 | YLL001W | 1032.66816 | 0.02011398 | 0.07445541 | 0.27014802 | 0.78704638 |
| YSR3 | YKR053C | 195.691199 | -0.0342498 | 0.12688832 | -0.2699206 | 0.78722135 |
| tT(CGU)K | tT(CGU)K | 1.39562743 | 0.05564945 | 0.2063488 | 0.26968634 | 0.78740157 |
| YDR537C | YDR537C | 9.05221487 | -0.0793501 | 0.29554001 | -0.2684919 | 0.78832074 |
| YLR311C | YLR311C | 5.76769123 | 0.07836514 | 0.292005 | 0.26836916 | 0.78841518 |
| UTR5 | YEL035C | 49.0523455 | 0.05696762 | 0.21239158 | 0.26821978 | 0.78853015 |
| tK(CUU)G3 | tK(CUU)G3 | 1.0497075 | -0.0479231 | 0.17885585 | -0.2679423 | 0.78874369 |
| Xu_2009_CUTs53 | Xu_2009_CU | 1.48158353 | -0.0579287 | 0.2161875 | -0.2679557 | 0.78873343 |
| Xu_2009_SUTs50 | Xu_2009_SU | 29.8035975 | 0.06623619 | 0.24761372 | 0.26749806 | 0.78908571 |
| Xu_2009_CUTs69 | Xu_2009_CU | 5.56780625 | -0.0781504 | 0.29244346 | -0.2672325 | 0.78929015 |
| Xu_2009_SUTs16 | Xu_2009_SU | 0.96555562 | -0.0239526 | 0.08967362 | -0.2671088 | 0.78938537 |
| YDR396W | YDR396W | 20.4726306 | -0.0712014 | 0.26633078 | -0.2673419 | 0.7892059 |
| YLR076C | YLR076C | 8.16564381 | 0.0790556 | 0.29647466 | 0.26665214 | 0.789737 |
| Xu_2009_CUTs14 | Xu_2009_CU | 3.25705176 | -0.0706174 | 0.26601247 | -0.2654663 | 0.79065024 |
| HCS1 | YKL017C | 542.017526 | -0.0237332 | 0.08936227 | -0.2655838 | 0.79055977 |

| | | | | | | |
|---|---|---|---|---|---|---|
| Xu_2009_CUTs60 | Xu_2009_CU | 14.0006141 | -0.0757927 | 0.28587819 | -0.2651225 | 0.7909151 |
| YAR029W | YAR029W | 109.248243 | 0.04204235 | 0.15869799 | 0.26492051 | 0.7910707 |
| YDR371C-A | YDR371C-A | 1.46616227 | -0.0546 | 0.20607152 | -0.2649565 | 0.79104299 |
| TSR4 | YOL022C | 1216.71707 | 0.01934694 | 0.0729839 | 0.26508504 | 0.79094395 |
| SSQ1 | YLR369W | 745.38642 | -0.0209731 | 0.07924317 | -0.2646681 | 0.79126517 |
| YKL100W-A | YKL100W-A | 1.09303539 | -0.0499938 | 0.18910031 | -0.2643773 | 0.79148923 |
| APC5 | YOR249C | 347.559076 | -0.0266125 | 0.1006147 | -0.2644995 | 0.79139506 |
| tS(AGA)A | tS(AGA)A | 7.37060153 | 0.07785442 | 0.29549995 | 0.26346677 | 0.79219083 |
| PNG1 | YPL096W | 283.351099 | 0.03015541 | 0.11457596 | 0.26319138 | 0.79240308 |
| DDI3 | YNL335W | 5.61635451 | 0.07616565 | 0.29235919 | 0.2605208 | 0.79446207 |
| SUE1 | YPR151C | 74.077405 | 0.05122625 | 0.19679375 | 0.26030427 | 0.79462908 |
| Xu_2009_CUTs75 | Xu_2009_CU | 109.711345 | 0.04010917 | 0.15436233 | 0.25983781 | 0.79498888 |
| FRT1 | YOR324C | 849.697153 | 0.02108194 | 0.08112712 | 0.25986307 | 0.7949694 |
| Xu_2009_SUTs74 | Xu_2009_SU | 28.5498216 | 0.06404307 | 0.24685295 | 0.25943813 | 0.79529721 |
| SEN34 | YAR008W | 276.907656 | 0.02748975 | 0.10622424 | 0.25878978 | 0.79579745 |
| YHR193C-A | YHR193C-A | 10.976619 | 0.07605649 | 0.29391117 | 0.25877375 | 0.79580981 |
| HOL1 | YNR055C | 1026.47126 | 0.01862548 | 0.07202231 | 0.25860715 | 0.79593837 |
| tT(UGU)G1 | tT(UGU)G1 | 6.01026361 | -0.0755108 | 0.29296092 | -0.2577505 | 0.79659948 |
| Xu_2009_CUTs62 | Xu_2009_CU | 2.26276679 | -0.0621889 | 0.24143932 | -0.2575757 | 0.79673435 |
| AZR1 | YGR224W | 8.26970832 | 0.07646158 | 0.29652563 | 0.25785824 | 0.79651631 |
| RPA43 | YOR340C | 1016.06252 | 0.01954915 | 0.07586934 | 0.25766864 | 0.79666265 |
| tD(GUC)L2 | tD(GUC)L2 | 2.01689625 | 0.05997953 | 0.23383945 | 0.25649875 | 0.79756574 |
| Xu_2009_CUTs87 | Xu_2009_CU | 13.731624 | -0.0746057 | 0.29179328 | -0.2556799 | 0.79819801 |
| YMR031W-A | YMR031W-A | 1.38812054 | 0.05437518 | 0.21285349 | 0.25545823 | 0.7983692 |
| YDL050C | YDL050C | 6.78829371 | 0.07495265 | 0.29413224 | 0.25482635 | 0.79885722 |
| GRX6 | YDL010W | 578.242258 | 0.02091249 | 0.08221945 | 0.2543497 | 0.7992254 |
| MCM21 | YDR318W | 164.702771 | -0.0353962 | 0.13923034 | -0.2542277 | 0.79931966 |
| RCY1 | YJL204C | 483.794561 | 0.02287565 | 0.09002493 | 0.25410348 | 0.79941561 |
| Xu_2009_SUTs21 | Xu_2009_SU | 15.8059859 | 0.07142049 | 0.28290393 | 0.25245491 | 0.80068946 |
| YFR010W-A | YFR010W-A | 1.48412751 | -0.0518176 | 0.20531379 | -0.2523825 | 0.8007454 |
| ICE2 | YIL090W | 910.26567 | -0.0180371 | 0.07135602 | -0.2527756 | 0.80044165 |
| INA22 | YIR024C | 245.311626 | 0.02888404 | 0.11442881 | 0.25241927 | 0.80071701 |
| COA6 | YMR244C-A | 750.358911 | 0.01968156 | 0.07798966 | 0.25236111 | 0.80076196 |
| Xu_2009_SUTs35 | Xu_2009_SU | 16.253581 | 0.07141666 | 0.28359606 | 0.2518253 | 0.8011761 |
| PRD1 | YCL057W | 2077.8455 | 0.01587422 | 0.0630355 | 0.25182979 | 0.80117263 |
| Xu_2009_CUTs66 | Xu_2009_CU | 7.84135889 | -0.0746355 | 0.29653332 | -0.2516935 | 0.80127799 |
| YBL006W-A | YBL006W-A | 12.1124345 | -0.0741377 | 0.29495093 | -0.251356 | 0.80153886 |
| Xu_2009_SUTs23 | Xu_2009_SU | 12.6553555 | 0.07230811 | 0.28806839 | 0.25101021 | 0.80180621 |
| YJL086C | YJL086C | 1.61595162 | 0.05624525 | 0.2245788 | 0.25044774 | 0.80224112 |
| YKL118W | YKL118W | 11.3289729 | -0.0732663 | 0.29398155 | -0.2492208 | 0.80318995 |
| YNL324W | YNL324W | 2.603814 | 0.06278226 | 0.25329154 | 0.2478656 | 0.8042384 |

| | | | | | | |
|---|---|---|---|---|---|---|
| Xu_2009_CUTs35 | Xu_2009_CU | 10.3630431 | -0.0729703 | 0.29485301 | -0.2474802 | 0.80453663 |
| PDS5 | YMR076C | 903.548721 | -0.0191017 | 0.07718818 | -0.247469 | 0.80454529 |
| Xu_2009_CUTs68 | Xu_2009_CU | 4.68170771 | 0.07129963 | 0.28851335 | 0.24712767 | 0.80480942 |
| YKL115C | YKL115C | 26.2694728 | -0.0631166 | 0.25550795 | -0.2470239 | 0.80488972 |
| YEL050W-A | YEL050W-A | 3.30777886 | 0.06487493 | 0.26319991 | 0.24648538 | 0.80530652 |
| SPT21 | YMR179W | 736.647395 | 0.0204768 | 0.08326304 | 0.24592901 | 0.80573718 |
| AHT1 | YHR093W | 0.7513913 | 0.04146533 | 0.16907478 | 0.24524843 | 0.80626408 |
| YOR314W-A | YOR314W-A | 2.39978778 | -0.0623465 | 0.25410936 | -0.245353 | 0.80618311 |
| Xu_2009_SUTs58 | Xu_2009_SU | 9.23679604 | -0.0722606 | 0.29482481 | -0.2450968 | 0.80638152 |
| tL(UAA)B2 | tL(UAA)B2 | 46.8899977 | 0.05141195 | 0.21019413 | 0.24459271 | 0.80677181 |
| HEM3 | YDL205C | 2422.63251 | 0.01399915 | 0.0574623 | 0.24362315 | 0.80752269 |
| RHO5 | YNL180C | 1674.9718 | -0.0204432 | 0.08437455 | -0.2422909 | 0.80855475 |
| VTC3 | YPL019C | 808.626438 | 0.01913791 | 0.07896555 | 0.24235772 | 0.80850299 |
| PHA2 | YNL316C | 285.66245 | -0.0264332 | 0.10952905 | -0.2413349 | 0.80929558 |
| YPR142C | YPR142C | 1.77788828 | 0.05458698 | 0.22634629 | 0.2411658 | 0.80942662 |
| SRL3 | YKR091W | 591.163362 | -0.0231928 | 0.09656995 | -0.2401662 | 0.81020141 |
| BI2 | Q0110 | 46.8204763 | 0.05240835 | 0.21844733 | 0.239913 | 0.8103977 |
| NPP2 | YEL016C | 599.031619 | -0.019463 | 0.08119587 | -0.2397042 | 0.81055955 |
| ECM13 | YBL043W | 149.636278 | 0.03303334 | 0.13854631 | 0.23842813 | 0.81154905 |
| Xu_2009_CUTs82 | Xu_2009_CU | 11.6112525 | -0.0698355 | 0.29331188 | -0.238093 | 0.81180898 |
| SPG4 | YMR107W | 35.3616224 | -0.0555777 | 0.23362973 | -0.2378879 | 0.81196801 |
| Xu_2009_CUTs84 | Xu_2009_CU | 24.8784586 | -0.0617583 | 0.26067971 | -0.2369127 | 0.81272448 |
| Xu_2009_SUTs11 | Xu_2009_SU | 55.4289079 | 0.04617968 | 0.19489645 | 0.23694469 | 0.8126997 |
| AFG2 | YLR397C | 814.550731 | -0.0182838 | 0.0772593 | -0.2366548 | 0.81292459 |
| FIP1 | YJR093C | 552.784301 | -0.0204418 | 0.08679891 | -0.2355078 | 0.81381463 |
| Xu_2009_CUTs76 | Xu_2009_CU | 11.2138139 | 0.06843199 | 0.29203902 | 0.23432481 | 0.81473282 |
| YGL041C-B | YGL041C-B | 7.68057454 | -0.0690303 | 0.2964584 | -0.23285 | 0.8158779 |
| PUS2 | YGL063W | 287.057094 | 0.02536015 | 0.10881036 | 0.23306745 | 0.81570903 |
| YHL045W | YHL045W | 0.94943727 | -0.0402064 | 0.17272061 | -0.2327826 | 0.81593019 |
| YKL147C | YKL147C | 9.43574044 | 0.06905156 | 0.29639408 | 0.23297215 | 0.81578303 |
| Xu_2009_CUTs30 | Xu_2009_CU | 18.8116894 | -0.0642819 | 0.2769338 | -0.2321201 | 0.81644476 |
| Xu_2009_CUTs87 | Xu_2009_CU | 9.88476664 | -0.0684908 | 0.29596285 | -0.2314167 | 0.81699107 |
| TFC4 | YGR047C | 896.573999 | -0.0157415 | 0.06799067 | -0.2315243 | 0.81690754 |
| Xu_2009_CUTs45 | Xu_2009_CU | 21.2929537 | 0.06063802 | 0.26227854 | 0.23119705 | 0.81716172 |
| tM(CAU)P/IMT2 | tM(CAU)P | 2.68727562 | -0.0582799 | 0.25410171 | -0.2293566 | 0.81859173 |
| YDR239C | YDR239C | 464.449769 | 0.02060474 | 0.08984284 | 0.22934201 | 0.81860311 |
| SED5 | YLR026C | 996.123774 | 0.01525961 | 0.06661922 | 0.22905712 | 0.81882452 |
| CAT2 | YML042W | 244.765762 | 0.02580239 | 0.11264807 | 0.22905312 | 0.81882763 |
| OAF3 | YKR064W | 646.90785 | -0.0175625 | 0.07677599 | -0.2287496 | 0.81906354 |
| Xu_2009_CUTs63 | Xu_2009_CU | 12.0869954 | 0.06745423 | 0.29522811 | 0.22848172 | 0.81927176 |
| YPL135C-A | YPL135C-A | 0.70024326 | -0.0350285 | 0.15329178 | -0.2285084 | 0.81925106 |

| | | | | | | |
|---|---|---|---|---|---|---|
| RSA4 | YCR072C | 1524.66022 | 0.01592238 | 0.06977362 | 0.22820051 | 0.81949037 |
| YKL153W | YKL153W | 5.80694165 | -0.0667135 | 0.29273252 | -0.2278993 | 0.81972454 |
| RNP1 | YLL046C | 82.1680437 | -0.0411272 | 0.18059638 | -0.2277301 | 0.81985605 |
| Xu_2009_SUTs18 | Xu_2009_SU | 60.514782 | 0.04393376 | 0.19344661 | 0.2271105 | 0.82033782 |
| Xu_2009_CUTs14 | Xu_2009_CU | 46.3593297 | 0.04996486 | 0.2205743 | 0.22652169 | 0.82079569 |
| CDC6 | YJL194W | 758.705748 | 0.01861442 | 0.08219778 | 0.22645885 | 0.82084456 |
| PDC6 | YGR087C | 95.1622676 | -0.0368797 | 0.16333886 | -0.2257864 | 0.82136755 |
| Xu_2009_CUTs72 | Xu_2009_CU | 0.96031068 | 0.04193978 | 0.18667957 | 0.22466185 | 0.82224235 |
| TSL1 | YML100W | 8773.35968 | 0.0184007 | 0.0820671 | 0.22421532 | 0.82258977 |
| Xu_2009_SUTs65 | Xu_2009_SU | 24.6955589 | -0.0575567 | 0.25752122 | -0.2235028 | 0.82314421 |
| ENT4 | YLL038C | 397.617964 | -0.02128 | 0.09535089 | -0.2231755 | 0.82339894 |
| RNA170 | RNA170 | 23.1434821 | 0.05777326 | 0.26076614 | 0.22155198 | 0.82466266 |
| YNL146C-A | YNL146C-A | 11.3522749 | 0.0648517 | 0.2928267 | 0.22146786 | 0.82472816 |
| tD(GUC)G2 | tD(GUC)G2 | 1.9489246 | -0.0515819 | 0.23364727 | -0.2207683 | 0.82527286 |
| SPO75 | YLL005C | 24.3798312 | 0.05891902 | 0.26722507 | 0.22048463 | 0.82549374 |
| Xu_2009_CUTs87 | Xu_2009_CU | 9.33449932 | 0.06511124 | 0.29587795 | 0.22006116 | 0.82582353 |
| EXG1 | YLR300W | 10590.7948 | 0.01028863 | 0.0468955 | 0.21939487 | 0.82634247 |
| YMR206W | YMR206W | 27.0612274 | 0.05775161 | 0.26332287 | 0.21931864 | 0.82640184 |
| YHR180W | YHR180W | 84.7614602 | 0.03652185 | 0.16705873 | 0.21861681 | 0.82694856 |
| SLG1 | YOR008C | 1599.33067 | 0.01296976 | 0.05929524 | 0.21873191 | 0.82685889 |
| Xu_2009_CUTs18 | Xu_2009_CU | 41.2220861 | 0.04790209 | 0.219714 | 0.21802022 | 0.82741336 |
| Xu_2009_CUTs11 | Xu_2009_CU | 12.6337828 | -0.0622628 | 0.28641426 | -0.2173873 | 0.8279065 |
| ERG7 | YHR072W | 1540.82374 | -0.0131044 | 0.06038643 | -0.2170088 | 0.82820145 |
| FYV10 | YIL097W | 292.105921 | -0.0249908 | 0.11618763 | -0.2150899 | 0.82969721 |
| GCD1 | YOR260W | 3009.9022 | 0.01139426 | 0.05297202 | 0.21509964 | 0.82968963 |
| YER138W-A | YER138W-A | 203.676795 | 0.02952886 | 0.13818868 | 0.21368509 | 0.83079264 |
| GPA2 | YER020W | 1373.0346 | 0.0139472 | 0.06546881 | 0.21303583 | 0.83129902 |
| YPL261C | YPL261C | 1.50638371 | -0.0431555 | 0.20329017 | -0.212285 | 0.8318847 |
| SNR33 | snR33 | 1443.68444 | 0.01830025 | 0.08645377 | 0.21167666 | 0.8323593 |
| Xu_2009_SUTs82 | Xu_2009_SU | 23.6685209 | -0.0552601 | 0.26175224 | -0.211116 | 0.83279677 |
| PAN1 | YIR006C | 3006.40615 | 0.01057429 | 0.05032656 | 0.21011352 | 0.83357907 |
| YBL071C-B | YBL071C-B | 9.5015804 | 0.06201699 | 0.29552619 | 0.20985279 | 0.83378257 |
| Xu_2009_SUTs52 | Xu_2009_SU | 10.9082606 | -0.0611303 | 0.29222715 | -0.2091875 | 0.83430187 |
| Xu_2009_SUTs20 | Xu_2009_SU | 158.62397 | -0.0276425 | 0.13314066 | -0.2076189 | 0.83552651 |
| YIL028W | YIL028W | 16.5683731 | -0.0591068 | 0.28530418 | -0.2071712 | 0.83587613 |
| YPL038W-A | YPL038W-A | 2.07476396 | -0.0487645 | 0.2358721 | -0.2067412 | 0.836212 |
| Xu_2009_CUTs83 | Xu_2009_CU | 14.6336993 | 0.05826958 | 0.28216088 | 0.20651189 | 0.83639108 |
| EIS1 | YMR031C | 1665.42568 | 0.01585443 | 0.07679345 | 0.20645547 | 0.83643514 |
| ARX1 | YDR101C | 2130.79043 | 0.01505641 | 0.07329203 | 0.20543041 | 0.83723587 |
| PCD1 | YLR151C | 124.190878 | 0.03308843 | 0.1614278 | 0.20497355 | 0.83759279 |
| ARO7 | YPR060C | 965.049747 | 0.01407582 | 0.0686665 | 0.20498811 | 0.83758141 |

| | | | | | | |
|---|---|---|---|---|---|---|
| SIF2 | YBR103W | 638.952953 | -0.0193423 | 0.09472486 | -0.2041946 | 0.8382014 |
| Xu_2009_CUTs34 | Xu_2009_CU | 7.81450867 | 0.06047563 | 0.29647804 | 0.20398014 | 0.83836902 |
| Xu_2009_CUTs14 | Xu_2009_CU | 7.62977918 | 0.0599937 | 0.29520202 | 0.2032293 | 0.83895582 |
| YDL129W | YDL129W | 215.247717 | 0.02626182 | 0.12932203 | 0.20307306 | 0.83907793 |
| tY(GUA)F2/SUP6 | tY(GUA)F2 | 1.29499012 | -0.0427154 | 0.21087896 | -0.2025587 | 0.83947998 |
| CHS2 | YBR038W | 1250.00107 | 0.01357909 | 0.0671085 | 0.20234528 | 0.8396468 |
| YLR101C | YLR101C | 5.74687646 | -0.0589049 | 0.29094954 | -0.2024576 | 0.83955901 |
| YBR178W | YBR178W | 1.3535494 | -0.0418036 | 0.20731724 | -0.2016406 | 0.84019774 |
| ARE2 | YNR019W | 1544.87864 | -0.0140097 | 0.06968417 | -0.201046 | 0.84066261 |
| Xu_2009_SUTs21 | Xu_2009_SU | 12.8995822 | 0.05732083 | 0.28692118 | 0.19977901 | 0.84165342 |
| Xu_2009_SUTs53 | Xu_2009_SU | 23.2626853 | 0.05213469 | 0.26102481 | 0.19973078 | 0.84169114 |
| Xu_2009_CUTs71 | Xu_2009_CU | 18.735206 | 0.05357638 | 0.26937297 | 0.19889294 | 0.8423465 |
| Xu_2009_SUTs74 | Xu_2009_SU | 41.6638817 | 0.04476995 | 0.22503029 | 0.19895079 | 0.84230124 |
| CIS1 | YLR346C | 41.7120478 | -0.0450489 | 0.22668478 | -0.1987294 | 0.84247444 |
| SAN1 | YDR143C | 1170.95923 | 0.01492511 | 0.07544411 | 0.19783008 | 0.84317801 |
| NBP35 | YGL091C | 522.994537 | 0.01637935 | 0.08369084 | 0.19571257 | 0.84483515 |
| tI(AAU)N2 | tI(AAU)N2 | 1.72638412 | -0.0449524 | 0.23000344 | -0.1954421 | 0.84504687 |
| Xu_2009_SUTs26 | Xu_2009_SU | 59.9908039 | 0.03730757 | 0.19213375 | 0.19417501 | 0.84603885 |
| YGR164W | YGR164W | 1.05850659 | 0.03639557 | 0.18793655 | 0.19365883 | 0.84644303 |
| Xu_2009_CUTs82 | Xu_2009_CU | 2.7223958 | -0.0490753 | 0.25593883 | -0.1917462 | 0.84794101 |
| YOR139C | YOR139C | 4.95755237 | 0.05542721 | 0.28990847 | 0.19118867 | 0.84837778 |
| YIL060W | YIL060W | 10.7682699 | 0.05623411 | 0.29434599 | 0.19104766 | 0.84848825 |
| HMG1 | YML075C | 1390.23428 | 0.01201406 | 0.06295401 | 0.19083871 | 0.84865196 |
| MTC1 | YJL123C | 1403.0688 | 0.01426231 | 0.07509887 | 0.18991383 | 0.84937665 |
| YPR074W-A | YPR074W-A | 3.55372189 | 0.05195491 | 0.27360745 | 0.18988852 | 0.84939649 |
| Xu_2009_SUTs38 | Xu_2009_SU | 68.3997276 | -0.0365567 | 0.19372218 | -0.1887067 | 0.85032269 |
| YDL228C | YDL228C | 0.77792067 | -0.0312084 | 0.16603372 | -0.1879641 | 0.85090482 |
| Xu_2009_CUTs15 | Xu_2009_CU | 25.2176495 | -0.0476918 | 0.25456437 | -0.1873468 | 0.85138876 |
| Xu_2009_CUTs72 | Xu_2009_CU | 15.9972378 | -0.0534685 | 0.2851135 | -0.1875341 | 0.85124188 |
| YDR215C | YDR215C | 4.0651195 | -0.0498754 | 0.2660335 | -0.1874779 | 0.85128598 |
| PNO1 | YOR145C | 2036.57343 | 0.01427482 | 0.07632024 | 0.18703844 | 0.8516305 |
| Xu_2009_CUTs78 | Xu_2009_CU | 63.7698502 | -0.0343765 | 0.1839995 | -0.1868292 | 0.85179456 |
| HSP42 | YDR171W | 3043.89921 | 0.0275621 | 0.14836362 | 0.18577401 | 0.85262198 |
| CSM3 | YMR048W | 245.786564 | 0.02393644 | 0.12913177 | 0.18536441 | 0.85294322 |
| Xu_2009_CUTs37 | Xu_2009_CU | 2.14417289 | 0.04416038 | 0.2399054 | 0.18407414 | 0.85395529 |
| tR(ACG)L | tR(ACG)L | 9.06585639 | -0.0540144 | 0.2964748 | -0.1821887 | 0.85543466 |
| tW(CCA)G1 | tW(CCA)G1 | 42.3537218 | -0.0409617 | 0.22524757 | -0.1818519 | 0.85569895 |
| Xu_2009_SUTs38 | Xu_2009_SU | 33.2148112 | -0.0428178 | 0.23433299 | -0.1827221 | 0.85501604 |
| Xu_2009_SUTs50 | Xu_2009_SU | 138.914844 | 0.02558018 | 0.14037229 | 0.18223094 | 0.85540149 |
| YBR191W-A | YBR191W-A | 1.89389819 | -0.0417815 | 0.22879337 | -0.1826166 | 0.85509885 |
| GCD11 | YER025W | 4605.40065 | -0.0115909 | 0.06349798 | -0.1825399 | 0.85515903 |

| | | | | | | |
|---|---|---|---|---|---|---|
| CRM1 | YGR218W | 4220.78835 | -0.0097437 | 0.05351878 | -0.1820621 | 0.85553398 |
| PLN1 | YKR046C | 1185.06557 | 0.012544 | 0.06898817 | 0.18182823 | 0.85571753 |
| BUD20 | YLR074C | 1245.11236 | 0.01534331 | 0.08420534 | 0.18221302 | 0.85541556 |
| YLR163W-A | YLR163W-A | 0.76131152 | -0.0288678 | 0.15864975 | -0.1819596 | 0.85561446 |
| YNL018C | YNL018C | 15.0494986 | -0.0514185 | 0.28361654 | -0.1812958 | 0.85613537 |
| Xu_2009_CUTs65 | Xu_2009_CU | 3.62992943 | 0.05012956 | 0.2772506 | 0.18080958 | 0.85651705 |
| TMA46 | YOR091W | 1723.10271 | -0.0124951 | 0.06920364 | -0.1805553 | 0.85671668 |
| HRB1 | YNL004W | 1689.24109 | 0.01190711 | 0.06606998 | 0.18021962 | 0.85698016 |
| CIN8 | YEL061C | 644.458297 | 0.01621743 | 0.09021328 | 0.17976761 | 0.85733501 |
| YGL258W-A | YGL258W-A | 4.98003442 | -0.0516918 | 0.28816959 | -0.1793799 | 0.85763945 |
| PUS6 | YGR169C | 380.393911 | 0.01961494 | 0.10988506 | 0.1785042 | 0.85832703 |
| Xu_2009_CUTs77 | Xu_2009_CU | 154.605168 | -0.0244222 | 0.13713306 | -0.1780913 | 0.8586513 |
| VTC5 | YDR089W | 607.857019 | -0.0201817 | 0.11330309 | -0.1781216 | 0.85862745 |
| Xu_2009_CUTs53 | Xu_2009_CU | 6.35664452 | -0.0520683 | 0.29341825 | -0.1774542 | 0.85915164 |
| YEL034C-A | YEL034C-A | 92.9923161 | 0.02874026 | 0.16180546 | 0.17762231 | 0.8590196 |
| MSN2 | YMR037C | 1568.93494 | 0.01287701 | 0.07251849 | 0.17756868 | 0.85906173 |
| YDL211C | YDL211C | 318.905335 | 0.01893203 | 0.10697116 | 0.17698254 | 0.85952211 |
| MET18 | YIL128W | 983.043536 | -0.0121508 | 0.06865048 | -0.1769951 | 0.85951224 |
| PEF1 | YGR058W | 404.524085 | 0.01711001 | 0.09689867 | 0.17657634 | 0.85984118 |
| PDR8 | YLR266C | 353.172241 | -0.0172174 | 0.09760835 | -0.1763932 | 0.85998505 |
| LRG1 | YDL240W | 737.407635 | 0.01368105 | 0.07765844 | 0.17616955 | 0.86016074 |
| VPS33 | YLR396C | 537.405606 | -0.0145489 | 0.08281287 | -0.1756846 | 0.86054177 |
| YML122C | YML122C | 17.3800833 | 0.04865799 | 0.2770025 | 0.17565904 | 0.86056182 |
| YNL019C | YNL019C | 1.95681683 | -0.040107 | 0.22918525 | -0.1749983 | 0.86108094 |
| SMA2 | YML066C | 9.738568 | 0.05158273 | 0.29517189 | 0.17475489 | 0.86127223 |
| MIX23 | YBL107C | 257.668403 | 0.02137046 | 0.12310538 | 0.17359481 | 0.86218391 |
| RRT15 | YLR162W-A | 2.00605201 | 0.03964636 | 0.2284026 | 0.17358103 | 0.86219475 |
| YGL024W | YGL024W | 7.44139024 | -0.0511716 | 0.29540239 | -0.1732267 | 0.86247322 |
| Xu_2009_CUTs56 | Xu_2009_CU | 9.54095536 | 0.05093583 | 0.29502947 | 0.17264658 | 0.86292923 |
| YNL319W | YNL319W | 10.7226749 | -0.0507563 | 0.29489637 | -0.1721157 | 0.86334657 |
| SDS23 | YGL056C | 1688.04013 | -0.0111636 | 0.06496194 | -0.171849 | 0.86355624 |
| HSV2 | YGR223C | 498.283176 | -0.0149369 | 0.0869449 | -0.1717969 | 0.86359718 |
| Xu_2009_SUTs30 | Xu_2009_SU | 4.82977851 | 0.04928417 | 0.28785146 | 0.17121389 | 0.86405559 |
| Xu_2009_CUTs35 | Xu_2009_CU | 25.8655737 | -0.0421163 | 0.24704168 | -0.1704824 | 0.86463076 |
| PIN4 | YBL051C | 2110.25698 | 0.00993674 | 0.05844952 | 0.17000558 | 0.86500575 |
| GLG2 | YJL137C | 546.816987 | 0.01660366 | 0.09786168 | 0.16966457 | 0.86527394 |
| CRS5 | YOR031W | 106.763824 | 0.02812523 | 0.1659847 | 0.16944471 | 0.86544686 |
| Xu_2009_SUTs53 | Xu_2009_SU | 2.8353837 | 0.04351453 | 0.25821972 | 0.16851747 | 0.8661762 |
| Xu_2009_CUTs32 | Xu_2009_CU | 41.2083377 | 0.03933341 | 0.23393874 | 0.16813551 | 0.86647667 |
| SYC1 | YOR179C | 193.323144 | -0.0211887 | 0.12640402 | -0.1676265 | 0.86687713 |
| MRM2 | YGL136C | 428.313163 | 0.01541874 | 0.09231229 | 0.16702803 | 0.86734799 |

| | | | | | | |
|---|---|---|---|---|---|---|
| Xu_2009_CUTs65 | Xu_2009_CU | 1.60172614 | -0.035686 | 0.21487954 | -0.1660747 | 0.8680982 |
| MSR1 | YHR091C | 708.150229 | -0.0132884 | 0.08047732 | -0.1651194 | 0.86885002 |
| SAP30 | YMR263W | 395.358417 | 0.01552395 | 0.09436385 | 0.16451167 | 0.86932836 |
| HPT1 | YDR399W | 1636.01407 | 0.0109295 | 0.06649604 | 0.16436311 | 0.86944531 |
| PBR1 | YNL181W | 601.815184 | -0.0142815 | 0.08695453 | -0.1642407 | 0.86954169 |
| INP53 | YOR109W | 1637.52895 | -0.0102714 | 0.06269645 | -0.1638282 | 0.86986642 |
| YDR183C-A | YDR183C-A | 16.6255709 | -0.0457931 | 0.28022048 | -0.163418 | 0.87018934 |
| APL3 | YBL037W | 699.358549 | 0.01271742 | 0.07791907 | 0.1632132 | 0.87035057 |
| AGA2 | YGL032C | 28.0045154 | 0.0397807 | 0.24402182 | 0.1630211 | 0.87050182 |
| YOR200W | YOR200W | 4.37852417 | 0.04619687 | 0.28353962 | 0.16292915 | 0.87057422 |
| YPL191C | YPL191C | 115.143589 | 0.02432795 | 0.14944547 | 0.16278814 | 0.87068525 |
| YAL045C | YAL045C | 4.31592459 | -0.0456844 | 0.28095606 | -0.1626032 | 0.87083085 |
| YBL109W | YBL109W | 7.91986392 | 0.04807679 | 0.29658437 | 0.16210155 | 0.87122589 |
| YFH1 | YDL120W | 623.548678 | -0.0127301 | 0.07855176 | -0.1620598 | 0.87125876 |
| CST6 | YIL036W | 1309.87993 | -0.0105246 | 0.0651649 | -0.1615075 | 0.87169372 |
| YHR210C | YHR210C | 103.183963 | -0.0258143 | 0.16029515 | -0.161042 | 0.87206032 |
| VPS74 | YDR372C | 1818.30878 | 0.0100987 | 0.06286374 | 0.16064422 | 0.87237362 |
| DUO1 | YGL061C | 184.056303 | -0.0209464 | 0.13030345 | -0.1607507 | 0.87228973 |
| YDR491C | YDR491C | 5.52087093 | 0.04686208 | 0.29230849 | 0.16031719 | 0.87263122 |
| MDM36 | YPR083W | 530.542629 | -0.0137371 | 0.08572512 | -0.1602464 | 0.872687 |
| Xu_2009_SUTs80 | Xu_2009_SU | 13.5905777 | -0.0463582 | 0.28970494 | -0.1600187 | 0.87286633 |
| Xu_2009_SUTs19 | Xu_2009_SU | 15.7356546 | -0.0452894 | 0.28409067 | -0.159419 | 0.87333878 |
| Xu_2009_SUTs10 | Xu_2009_SU | 1.32060169 | 0.03186536 | 0.20016835 | 0.15919278 | 0.873517 |
| MGA1 | YGR249W | 236.81859 | 0.01807885 | 0.11369912 | 0.15900604 | 0.87366412 |
| tS(UGA)P | tS(UGA)P | 49.5938936 | -0.0347834 | 0.21907613 | -0.1587729 | 0.87384777 |
| Xu_2009_SUTs54 | Xu_2009_SU | 40.4250571 | 0.03495328 | 0.22021931 | 0.15872034 | 0.87388921 |
| Xu_2009_SUTs73 | Xu_2009_SU | 46.6479827 | -0.0332523 | 0.21027675 | -0.1581358 | 0.8743498 |
| CDC40 | YDR364C | 590.045865 | 0.01323409 | 0.08381464 | 0.15789713 | 0.87453786 |
| CHM7 | YJL049W | 329.865507 | -0.0165283 | 0.10468316 | -0.1578889 | 0.87454437 |
| RTC4 | YNL254C | 328.387717 | 0.01547593 | 0.09815432 | 0.1576694 | 0.87471731 |
| YOR381W-A | YOR381W-A | 55.5777918 | 0.0331484 | 0.21022237 | 0.15768255 | 0.87470695 |
| Xu_2009_SUTs55 | Xu_2009_SU | 41.0987012 | -0.0355674 | 0.2259072 | -0.1574426 | 0.87489604 |
| tA(AGC)P | tA(AGC)P | 6.61213215 | 0.04572359 | 0.29175852 | 0.15671725 | 0.87546769 |
| Xu_2009_CUTs54 | Xu_2009_CU | 3.3763077 | -0.0412988 | 0.26342602 | -0.1567756 | 0.87542169 |
| JLP2 | YMR132C | 153.527009 | -0.0222126 | 0.1420425 | -0.15638 | 0.87573353 |
| YML100W-A | YML100W-A | 67.6937831 | -0.0288197 | 0.18637766 | -0.1546307 | 0.87711246 |
| Xu_2009_CUTs86 | Xu_2009_CU | 24.0837726 | 0.04037725 | 0.26238168 | 0.15388747 | 0.87769847 |
| YEL009C-A | YEL009C-A | 12.5722006 | -0.0441771 | 0.28769992 | -0.1535526 | 0.87796251 |
| DAL7 | YIR031C | 501.765817 | 0.01410889 | 0.09184643 | 0.15361396 | 0.87791413 |
| YPL102C | YPL102C | 7.63296166 | 0.04551594 | 0.29651372 | 0.15350365 | 0.87800111 |
| SMK1 | YPR054W | 29.2062295 | 0.03707533 | 0.24092406 | 0.15388804 | 0.87769801 |

| | | | | | | |
|---|---|---|---|---|---|---|
| SNR59 | snR59 | 28.1649466 | -0.0380658 | 0.24871238 | -0.1530515 | 0.87835766 |
| YPR170C | YPR170C | 23.0109542 | 0.04020768 | 0.26359241 | 0.15253733 | 0.87876315 |
| tD(GUC)L1 | tD(GUC)L1 | 1.98242904 | 0.03619379 | 0.23888781 | 0.15150956 | 0.87957377 |
| FYV6 | YNL133C | 316.997487 | -0.0162303 | 0.10753474 | -0.1509311 | 0.8800301 |
| Xu_2009_CUTs53 | Xu_2009_CU | 11.1529165 | -0.0441397 | 0.29425966 | -0.1500024 | 0.88076273 |
| UBP6 | YFR010W | 1369.42109 | -0.0110769 | 0.07408016 | -0.1495255 | 0.88113898 |
| tD(GUC)D | tD(GUC)D | 1.47776456 | -0.031262 | 0.20962736 | -0.1491313 | 0.88145002 |
| Xu_2009_SUTs95 | Xu_2009_SU | 102.295014 | 0.02310359 | 0.15574467 | 0.14834271 | 0.88207231 |
| IES5 | YER092W | 423.930219 | -0.0157739 | 0.10624895 | -0.1484613 | 0.88197872 |
| YLR198C | YLR198C | 41.9994705 | 0.03242704 | 0.21982143 | 0.14751537 | 0.88272525 |
| VPS9 | YML097C | 866.044606 | 0.01102722 | 0.07489322 | 0.14723925 | 0.88294318 |
| Xu_2009_SUTs47 | Xu_2009_SU | 3.10091257 | -0.0341577 | 0.23434524 | -0.1457581 | 0.88411237 |
| PAL2 | YHR097C | 856.895241 | -0.0102698 | 0.07054956 | -0.1455685 | 0.88426204 |
| YEL018C-A | YEL018C-A | 13.0945557 | -0.0418318 | 0.2876438 | -0.1454292 | 0.88437201 |
| RPA49 | YNL248C | 1827.3218 | 0.01128551 | 0.07769637 | 0.14525148 | 0.88451232 |
| Xu_2009_SUTs22 | Xu_2009_SU | 0.76528399 | -0.0233793 | 0.16129476 | -0.1449477 | 0.88475214 |
| Xu_2009_SUTs56 | Xu_2009_SU | 102.49759 | -0.0225916 | 0.15579809 | -0.1450057 | 0.88470635 |
| YCR006C | YCR006C | 96.3070575 | 0.02358795 | 0.1637956 | 0.14400843 | 0.88549381 |
| IRC25 | YLR021W | 345.81221 | -0.0139898 | 0.09756993 | -0.1433822 | 0.8859883 |
| YAL065C | YAL065C | 3.74396549 | -0.0392381 | 0.27395722 | -0.1432272 | 0.88611072 |
| TEM1 | YML064C | 404.384687 | 0.01367563 | 0.09591461 | 0.14258126 | 0.8866209 |
| IKI1 | YHR187W | 584.577524 | -0.0119847 | 0.08417788 | -0.1423731 | 0.88678531 |
| CMS1 | YLR003C | 654.254097 | 0.01227024 | 0.08625425 | 0.14225663 | 0.8868773 |
| PEX7 | YDR142C | 287.359328 | -0.0153694 | 0.10869639 | -0.1413975 | 0.8875559 |
| YGR161W-B | YGR161W-B | 3.03953287 | 0.03810298 | 0.26928229 | 0.14149829 | 0.88747631 |
| YBR064W | YBR064W | 0.94776326 | 0.02382826 | 0.17065906 | 0.13962496 | 0.88895632 |
| AMD2 | YDR242W | 209.541131 | 0.0174096 | 0.12465726 | 0.13965975 | 0.88892883 |
| YJR151W-A | YJR151W-A | 1.73594245 | -0.0312575 | 0.22400128 | -0.1395417 | 0.88902211 |
| RSM22 | YKL155C | 999.985562 | -0.0099387 | 0.07109627 | -0.1397927 | 0.88882376 |
| Xu_2009_CUTs14 | Xu_2009_CU | 7.27137982 | 0.04120177 | 0.2963437 | 0.13903374 | 0.88942349 |
| GIP1 | YBR045C | 159.328706 | -0.0180934 | 0.13050391 | -0.1386427 | 0.88973246 |
| Xu_2009_SUTs20 | Xu_2009_SU | 44.6420945 | -0.0299563 | 0.21739096 | -0.137799 | 0.89039929 |
| KXD1 | YGL079W | 258.625166 | 0.01598351 | 0.11623344 | 0.13751212 | 0.89062601 |
| EAF6 | YJR082C | 642.863528 | 0.01237024 | 0.09076637 | 0.13628664 | 0.89159469 |
| SDD3 | YOL098C | 2257.27717 | 0.00728005 | 0.05345139 | 0.13619946 | 0.8916636 |
| YAR019W-A | YAR019W-A | 16.2368169 | 0.03722162 | 0.27464008 | 0.13552874 | 0.89219384 |
| Xu_2009_CUTs79 | Xu_2009_CU | 3.43122438 | -0.0354962 | 0.26501132 | -0.133942 | 0.89344842 |
| Xu_2009_CUTs42 | Xu_2009_CU | 21.2775123 | -0.0356222 | 0.26664088 | -0.1335961 | 0.89372196 |
| Xu_2009_CUTs33 | Xu_2009_CU | 6.55501307 | -0.039297 | 0.29487294 | -0.1332675 | 0.8939818 |
| SUT1 | YGL162W | 122.807421 | -0.0202468 | 0.15266503 | -0.1326225 | 0.89449195 |
| Xu_2009_CUTs28 | Xu_2009_CU | 24.3924514 | 0.03464993 | 0.26222443 | 0.13213845 | 0.89487478 |

| | | | | | | |
|---|---|---|---|---|---|---|
| MSB2 | YGR014W | 4182.8529 | -0.0076876 | 0.05821799 | -0.1320481 | 0.89494627 |
| YJR087W | YJR087W | 62.8797644 | -0.0252762 | 0.19112298 | -0.1322508 | 0.89478588 |
| YIR018C-A | YIR018C-A | 14.0993194 | -0.0374465 | 0.28474394 | -0.1315095 | 0.89537226 |
| YIL032C | YIL032C | 2.15900802 | 0.03138271 | 0.2401583 | 0.13067509 | 0.89603234 |
| YDL242W | YDL242W | 5.39686872 | -0.0379135 | 0.29188962 | -0.1298897 | 0.89665369 |
| PHO84 | YML123C | 147.334835 | 0.03839022 | 0.29586872 | 0.12975422 | 0.89676088 |
| MTE1 | YGR042W | 380.028506 | 0.01268551 | 0.09788052 | 0.129602 | 0.89688132 |
| YHR180W-A | YHR180W-A | 0.7962379 | -0.0113971 | 0.0887961 | -0.128351 | 0.89787121 |
| YHR095W | YHR095W | 5.59142832 | 0.03736587 | 0.29156041 | 0.12815823 | 0.89802376 |
| AEP3 | YPL005W | 384.460604 | -0.0119977 | 0.09375483 | -0.1279684 | 0.89817396 |
| Xu_2009_CUTs53 | Xu_2009_CU | 20.3531407 | -0.0350836 | 0.27457938 | -0.1277721 | 0.89832935 |
| SHQ1 | YIL104C | 555.390344 | 0.01245775 | 0.09784053 | 0.12732711 | 0.8986815 |
| Xu_2009_SUTs41 | Xu_2009_SU | 28.4198658 | 0.03364381 | 0.26904949 | 0.12504691 | 0.90048641 |
| Xu_2009_SUTs47 | Xu_2009_SU | 2.01462362 | -0.0291453 | 0.23433791 | -0.1243732 | 0.90101982 |
| THI12 | YNL332W | 48.186871 | -0.0261065 | 0.21000264 | -0.1243151 | 0.90106575 |
| BIO5 | YNR056C | 50.8466937 | -0.0257158 | 0.20796913 | -0.1236519 | 0.90159092 |
| YPR177C | YPR177C | 5.44346316 | 0.03590737 | 0.29028407 | 0.12369736 | 0.9015549 |
| SNO3 | YFL060C | 68.5058275 | 0.0229843 | 0.18626818 | 0.12339359 | 0.90179543 |
| YDL022C-A | YDL022C-A | 4.78321203 | -0.0344927 | 0.28334707 | -0.121733 | 0.90311049 |
| HED1 | YDR014W-A | 73.3473023 | -0.0219405 | 0.18021612 | -0.1217455 | 0.90310054 |
| VIP1 | YLR410W | 2657.95042 | 0.00694533 | 0.05697172 | 0.12190846 | 0.90297152 |
| MEI5 | YPL121C | 4.94164 | 0.0350877 | 0.28780929 | 0.12191302 | 0.9029679 |
| TUS1 | YLR425W | 796.723968 | 0.00863554 | 0.07112921 | 0.12140637 | 0.90336917 |
| tR(ACG)O | tR(ACG)O | 1.26800704 | 0.02403594 | 0.1991759 | 0.12067695 | 0.90394691 |
| Xu_2009_CUTs79 | Xu_2009_CU | 1.20220165 | -0.0241975 | 0.19987492 | -0.1210631 | 0.90364105 |
| SRO9 | YCL037C | 4317.54826 | -0.0079865 | 0.06618898 | -0.1206618 | 0.90395888 |
| YGR022C | YGR022C | 1.96515212 | 0.02866848 | 0.23665047 | 0.12114273 | 0.90357799 |
| AVT1 | YJR001W | 1215.87036 | 0.00897536 | 0.0743003 | 0.12079845 | 0.90385067 |
| Xu_2009_SUTs45 | Xu_2009_SU | 56.6097058 | -0.0239999 | 0.20088454 | -0.1194714 | 0.90490193 |
| IME2 | YJL106W | 154.808029 | -0.0157743 | 0.13294063 | -0.1186568 | 0.90554724 |
| YBL070C | YBL070C | 2.41010399 | 0.02795374 | 0.23776117 | 0.11757066 | 0.90640786 |
| DBF20 | YPR111W | 455.970404 | 0.01171231 | 0.09951761 | 0.11769081 | 0.90631265 |
| KRE6 | YPR159W | 4906.45516 | 0.0061084 | 0.05308563 | 0.11506686 | 0.90839213 |
| NVJ3 | YDR179W-A | 193.73297 | -0.0146398 | 0.12763413 | -0.1147011 | 0.90868206 |
| Xu_2009_CUTs25 | Xu_2009_CU | 7.91060868 | 0.03367624 | 0.29652126 | 0.11357107 | 0.90957782 |
| PLB1 | YMR008C | 1483.42024 | 0.00758848 | 0.06674093 | 0.11370047 | 0.90947524 |
| YLR146W-A | YLR146W-A | 116.397077 | -0.0174263 | 0.1540458 | -0.1131241 | 0.90993218 |
| YBL065W | YBL065W | 3.12252094 | 0.02978816 | 0.26454842 | 0.11260003 | 0.91034766 |
| Xu_2009_CUTs34 | Xu_2009_CU | 19.5357566 | -0.0298991 | 0.26642782 | -0.1122221 | 0.91064733 |
| PIM1 | YBL022C | 2903.76275 | -0.0060377 | 0.05405254 | -0.1116997 | 0.91106154 |
| YHR214W | YHR214W | 4.6303052 | 0.03158112 | 0.2830522 | 0.1115735 | 0.91116158 |

| | | | | | | |
|---|---|---|---|---|---|---|
| TED1 | YIL039W | 1115.73274 | 0.00721553 | 0.06464794 | 0.11161263 | 0.91113056 |
| Xu_2009_SUTs80 | Xu_2009_SU | 33.7244763 | 0.02566894 | 0.23130845 | 0.11097278 | 0.91163793 |
| tK(CUU)J | tK(CUU)J | 0.87776363 | -0.0192064 | 0.17390445 | -0.1104423 | 0.91205864 |
| SET5 | YHR207C | 652.735456 | -0.0097358 | 0.08820062 | -0.1103824 | 0.91210609 |
| SPO21 | YOL091W | 197.900374 | -0.0135776 | 0.12321173 | -0.1101972 | 0.91225297 |
| Xu_2009_SUTs82 | Xu_2009_SU | 5.39763206 | 0.03204955 | 0.29159631 | 0.10991069 | 0.9124802 |
| tW(CCA)M | tW(CCA)M | 7.37018745 | -0.0322887 | 0.2953755 | -0.109314 | 0.91295348 |
| NSE5 | YML023C | 444.702238 | -0.0094985 | 0.08743855 | -0.1086302 | 0.91349582 |
| Xu_2009_SUTs83 | Xu_2009_SU | 14.8540003 | 0.03072784 | 0.28394833 | 0.10821631 | 0.91382411 |
| SWT1 | YOR166C | 807.485603 | 0.00793144 | 0.07320068 | 0.10835196 | 0.91371651 |
| Xu_2009_CUTs63 | Xu_2009_CU | 21.1910608 | 0.02846543 | 0.26495993 | 0.10743294 | 0.91444552 |
| Xu_2009_CUTs74 | Xu_2009_CU | 10.3003043 | 0.03146303 | 0.29545955 | 0.10648844 | 0.91519482 |
| BEM1 | YBR200W | 1098.31431 | -0.0070912 | 0.06658499 | -0.1064988 | 0.91518662 |
| MCO14 | YHL018W | 66.5008855 | -0.0200296 | 0.18836818 | -0.1063322 | 0.91531876 |
| FMP32 | YFL046W | 553.383869 | 0.00971312 | 0.09231512 | 0.10521698 | 0.91620364 |
| MPE1 | YKL059C | 656.208435 | -0.0078961 | 0.07505047 | -0.1052109 | 0.91620844 |
| Xu_2009_CUTs88 | Xu_2009_CU | 14.9918148 | -0.0294922 | 0.28071071 | -0.1050625 | 0.91632618 |
| YAR010C | YAR010C | 7.80364556 | 0.03079572 | 0.2956761 | 0.10415357 | 0.91704748 |
| TFG2 | YGR005C | 778.503277 | -0.0076634 | 0.0741252 | -0.1033843 | 0.917658 |
| PTC5 | YOR090C | 967.312403 | 0.00750165 | 0.07278832 | 0.10306118 | 0.91791442 |
| Xu_2009_CUTs56 | Xu_2009_CU | 1.36441445 | 0.02066441 | 0.2010711 | 0.10277166 | 0.9181442 |
| tA(UGC)G | tA(UGC)G | 2.74443145 | -0.025633 | 0.25022498 | -0.1024399 | 0.91840751 |
| PAU3 | YCR104W | 4.01948018 | 0.02896012 | 0.28260508 | 0.10247559 | 0.91837919 |
| YKL096C-B | YKL096C-B | 2.04068177 | -0.0241914 | 0.23729138 | -0.1019479 | 0.91879801 |
| YGR025W | YGR025W | 6.57978356 | 0.03011468 | 0.29605113 | 0.1017212 | 0.91897797 |
| YOL046C | YOL046C | 1.18356104 | 0.02041933 | 0.20094087 | 0.1016186 | 0.91905941 |
| Xu_2009_SUTs46 | Xu_2009_SU | 52.3199437 | -0.0204327 | 0.20308423 | -0.1006121 | 0.91985839 |
| YCK2 | YNL154C | 1956.65161 | 0.00646372 | 0.06421979 | 0.10064992 | 0.91982836 |
| Xu_2009_SUTs77 | Xu_2009_SU | 11.7661227 | 0.02966314 | 0.29540756 | 0.10041428 | 0.92001543 |
| ATG8 | YBL078C | 369.824594 | -0.0100473 | 0.10070159 | -0.0997731 | 0.92052448 |
| YDL016C | YDL016C | 68.4552669 | 0.01926193 | 0.19311341 | 0.09974411 | 0.92054748 |
| YHR137C-A | YHR137C-A | 3.59766504 | 0.02746912 | 0.27519255 | 0.09981782 | 0.92048896 |
| BZZ1 | YHR114W | 904.867029 | -0.0069639 | 0.07034066 | -0.0990027 | 0.92113609 |
| YOR062C | YOR062C | 492.706483 | 0.0107655 | 0.10870009 | 0.09903858 | 0.92110764 |
| YOR314W | YOR314W | 10.5822 | -0.0289047 | 0.2942275 | -0.0982394 | 0.92174223 |
| LCB1 | YMR296C | 2540.04973 | -0.0055773 | 0.05690628 | -0.098008 | 0.92192598 |
| Xu_2009_CUTs82 | Xu_2009_CU | 57.0203243 | -0.0191796 | 0.19647335 | -0.0976192 | 0.92223465 |
| PMR1 | YGL167C | 3828.12947 | -0.00524 | 0.05377883 | -0.0974355 | 0.92238058 |
| NCE103 | YNL036W | 2086.63473 | 0.00649209 | 0.06655334 | 0.0975472 | 0.92229185 |
| Xu_2009_CUTs29 | Xu_2009_CU | 14.1451154 | -0.0278136 | 0.28897547 | -0.096249 | 0.92332284 |
| YCR023C | YCR023C | 245.804461 | -0.0107648 | 0.11246461 | -0.0957172 | 0.92374519 |

| | | | | | | |
|---|---|---|---|---|---|---|
| YLR269C | YLR269C | 0.92083484 | -0.0172395 | 0.18113748 | -0.0951738 | 0.92417678 |
| YDL009C | YDL009C | 27.0955648 | -0.023899 | 0.25401049 | -0.0940867 | 0.92504026 |
| THI5 | YFL058W | 33.7755713 | 0.02271603 | 0.24111618 | 0.09421196 | 0.92494079 |
| YLL056C | YLL056C | 87.7753308 | -0.0160819 | 0.17030122 | -0.0944321 | 0.92476593 |
| TIF5 | YPR041W | 2224.75258 | -0.0070154 | 0.07446309 | -0.0942133 | 0.92493974 |
| RAV2 | YDR202C | 468.065316 | 0.00900565 | 0.09616839 | 0.09364455 | 0.92539152 |
| Xu_2009_SUTs76 | Xu_2009_SU | 17.8090132 | -0.0256301 | 0.27611495 | -0.0928242 | 0.92604324 |
| GEP4 | YHR100C | 137.414349 | -0.0133392 | 0.14380725 | -0.0927572 | 0.92609647 |
| SEC59 | YMR013C | 235.839394 | -0.0108233 | 0.11668467 | -0.0927565 | 0.926097 |
| tN(GUU)O1 | tN(GUU)O1 | 2.49365429 | 0.02312749 | 0.25140791 | 0.09199191 | 0.92670447 |
| STP2 | YHR006W | 1052.21243 | -0.0063771 | 0.0697886 | -0.0913772 | 0.9271929 |
| MTR2 | YKL186C | 1014.69957 | -0.0065426 | 0.07206567 | -0.0907863 | 0.92766243 |
| SHU1 | YHL006C | 77.1121056 | -0.0159908 | 0.17667256 | -0.0905112 | 0.92788103 |
| Xu_2009_CUTs88 | Xu_2009_CU | 3.38813326 | -0.0241185 | 0.26921992 | -0.0895867 | 0.92861567 |
| DSF2 | YBR007C | 801.638884 | 0.00694824 | 0.07755236 | 0.08959415 | 0.92860974 |
| SPO13 | YHR014W | 26.7417977 | -0.0226083 | 0.25327886 | -0.0892626 | 0.92887325 |
| ECM38 | YLR299W | 852.657136 | 0.00649612 | 0.07382129 | 0.0879979 | 0.92987835 |
| Xu_2009_SUTs80 | Xu_2009_SU | 63.0039189 | -0.0168469 | 0.19280977 | -0.0873758 | 0.93037283 |
| RRT1 | YBL048W | 14.5070577 | -0.0247406 | 0.28351129 | -0.0872648 | 0.93046101 |
| KTR5 | YNL029C | 628.722724 | -0.0077056 | 0.08863093 | -0.0869405 | 0.9307188 |
| EGH1 | YIR007W | 647.196061 | -0.0072282 | 0.08327672 | -0.0867969 | 0.93083294 |
| tV(AAC)E1 | tV(AAC)E1 | 0.89164419 | -0.0153295 | 0.17886872 | -0.0857027 | 0.93170275 |
| Xu_2009_CUTs70 | Xu_2009_CU | 49.7844095 | -0.0180974 | 0.21174107 | -0.0854696 | 0.93188805 |
| BIT2 | YBR270C | 268.593409 | -0.0095659 | 0.11221013 | -0.0852503 | 0.9320624 |
| OPI7 | YDR360W | 1.04338565 | -0.0162809 | 0.19124801 | -0.08513 | 0.93215802 |
| Xu_2009_CUTs19 | Xu_2009_CU | 32.7899484 | 0.0199222 | 0.23632083 | 0.08430148 | 0.93281673 |
| Xu_2009_CUTs28 | Xu_2009_CU | 1.87006797 | 0.01943678 | 0.23134322 | 0.08401706 | 0.93304287 |
| Xu_2009_CUTs19 | Xu_2009_CU | 1.68995044 | -0.0182446 | 0.21808699 | -0.0836573 | 0.9333289 |
| MOT2 | YER068W | 1326.26517 | -0.0054436 | 0.06544863 | -0.0831742 | 0.93371303 |
| Xu_2009_CUTs87 | Xu_2009_CU | 11.8343887 | -0.0245531 | 0.29591222 | -0.0829744 | 0.93387187 |
| RRN10 | YBL025W | 240.52325 | 0.00980118 | 0.11878303 | 0.08251329 | 0.93423855 |
| RTT103 | YDR289C | 358.366094 | 0.0081615 | 0.09894542 | 0.08248483 | 0.93426118 |
| AI3 | Q0060 | 47.7757061 | 0.01828661 | 0.22221122 | 0.08229383 | 0.93441306 |
| Xu_2009_CUTs48 | Xu_2009_CU | 19.3172528 | 0.02211103 | 0.2706452 | 0.08169747 | 0.93488729 |
| Xu_2009_SUTs57 | Xu_2009_SU | 12.0790437 | -0.0238374 | 0.29230532 | -0.0815495 | 0.93500495 |
| Xu_2009_SUTs26 | Xu_2009_SU | 37.8413898 | -0.0190801 | 0.23585324 | -0.0808983 | 0.93552287 |
| DUR3 | YHL016C | 370.363341 | -0.0077832 | 0.09736293 | -0.0799401 | 0.93628487 |
| POL32 | YJR043C | 464.449626 | -0.0080313 | 0.10041621 | -0.0799803 | 0.93625292 |
| Xu_2009_CUTs70 | Xu_2009_CU | 2.38711852 | 0.01960404 | 0.24745503 | 0.07922262 | 0.93685555 |
| HFM1 | YGL251C | 207.294494 | -0.0098641 | 0.12443039 | -0.0792744 | 0.93681437 |
| DAS1 | YJL149W | 674.989627 | 0.0069565 | 0.08923377 | 0.07795812 | 0.93786137 |

| | | | | | | |
|---|---|---|---|---|---|---|
| Xu_2009_SUTs44 | Xu_2009_SU | 103.575417 | 0.01263589 | 0.16304041 | 0.0775016 | 0.93822452 |
| Xu_2009_SUTs77 | Xu_2009_SU | 41.5176462 | 0.01684423 | 0.21812075 | 0.07722433 | 0.93844509 |
| OYE3 | YPL171C | 50.1884605 | 0.01682343 | 0.21792656 | 0.07719769 | 0.93846628 |
| tS(GCU)F | tS(GCU)F | 8.93259018 | -0.0227119 | 0.29652342 | -0.0765938 | 0.9389467 |
| URK1 | YNR012W | 507.201125 | -0.0066437 | 0.08662459 | -0.0766955 | 0.93886575 |
| YOR146W | YOR146W | 1.18852804 | -0.0146935 | 0.19290737 | -0.0761688 | 0.9392848 |
| tE(UUC)I | tE(UUC)I | 1.50366913 | -0.0159715 | 0.21233796 | -0.0752172 | 0.94004191 |
| Xu_2009_SUTs84 | Xu_2009_SU | 48.0899648 | -0.0153362 | 0.20327644 | -0.0754448 | 0.93986081 |
| YNL011C | YNL011C | 249.102068 | 0.00870232 | 0.11557403 | 0.07529654 | 0.93997878 |
| INN1 | YNL152W | 571.702731 | -0.0062551 | 0.0831957 | -0.0751854 | 0.94006718 |
| YPL080C | YPL080C | 4.43135064 | -0.0210219 | 0.28099113 | -0.0748132 | 0.9403633 |
| RLP24 | YLR009W | 2202.86323 | -0.0052853 | 0.07096671 | -0.0744765 | 0.94063127 |
| HMT1 | YBR034C | 2017.11447 | -0.00688 | 0.09269968 | -0.0742183 | 0.94083665 |
| PEX15 | YOL044W | 145.26678 | -0.0112919 | 0.1527005 | -0.0739478 | 0.94105189 |
| HVG1 | YER039C | 56.7011928 | -0.0148039 | 0.20149933 | -0.0734685 | 0.94143331 |
| STV1 | YMR054W | 1793.52572 | -0.0041786 | 0.05797012 | -0.0720823 | 0.94253642 |
| tS(CGA)C/SUP61 | tS(CGA)C | 11.5541907 | -0.0209038 | 0.29238195 | -0.071495 | 0.94300383 |
| YBL053W | YBL053W | 2.87683023 | -0.0187125 | 0.26092746 | -0.0717153 | 0.94282851 |
| PAC2 | YER007W | 97.0427031 | -0.0118727 | 0.16537725 | -0.0717914 | 0.94276789 |
| SNX4 | YJL036W | 613.096263 | 0.00571999 | 0.07985198 | 0.07163247 | 0.9428944 |
| YME2 | YMR302C | 1020.64334 | 0.00494616 | 0.06958324 | 0.07108264 | 0.94333198 |
| YAL037C-B | YAL037C-B | 5.18978357 | -0.0197307 | 0.28443591 | -0.0693679 | 0.94469682 |
| CNM67 | YNL225C | 435.701437 | 0.00645561 | 0.09645447 | 0.06692906 | 0.94663818 |
| YOR268C | YOR268C | 4.55399442 | 0.01887075 | 0.28398313 | 0.06645024 | 0.94701937 |
| SOR2 | YDL246C | 8.97911807 | -0.019445 | 0.29647409 | -0.0655876 | 0.94770615 |
| Xu_2009_SUTs77 | Xu_2009_SU | 0.86357311 | 0.01147484 | 0.17557696 | 0.06535506 | 0.9478913 |
| Xu_2009_SUTs67 | Xu_2009_SU | 15.5175016 | 0.01849588 | 0.28588413 | 0.06469712 | 0.94841515 |
| SSE2 | YBR169C | 1773.14793 | 0.0049891 | 0.0773388 | 0.0645097 | 0.94856438 |
| YLR171W | YLR171W | 9.33375309 | 0.01904296 | 0.29519199 | 0.06451042 | 0.94856381 |
| YML009C-A | YML009C-A | 5.61704316 | 0.01881922 | 0.29290409 | 0.06425045 | 0.9487708 |
| CSL4 | YNL232W | 1039.44587 | 0.00490257 | 0.07691181 | 0.06374273 | 0.94917508 |
| tR(UCU)E | tR(UCU)E | 1.00541839 | 0.01139374 | 0.18115777 | 0.06289403 | 0.94985089 |
| CTF3 | YLR381W | 346.384204 | -0.0062987 | 0.10010711 | -0.0629197 | 0.94983041 |
| PRP19 | YLL036C | 855.784542 | -0.0047646 | 0.07645456 | -0.0623192 | 0.95030867 |
| Xu_2009_CUTs60 | Xu_2009_CU | 21.1228851 | -0.0174778 | 0.2829374 | -0.0617726 | 0.95074393 |
| Xu_2009_SUTs27 | Xu_2009_SU | 8.16207273 | 0.01785025 | 0.29652845 | 0.06019743 | 0.95199839 |
| Xu_2009_CUTs70 | Xu_2009_CU | 40.3214697 | -0.0136016 | 0.22741036 | -0.0598107 | 0.9523064 |
| YPL257W-B | YPL257W-B | 9.34857106 | 0.01767652 | 0.29495358 | 0.05992985 | 0.9522115 |
| KRE5 | YOR336W | 1347.53353 | -0.0036467 | 0.06258566 | -0.0582667 | 0.95353621 |
| UBC13 | YDR092W | 691.633097 | -0.0043054 | 0.07496266 | -0.0574339 | 0.95419957 |
| VHS3 | YOR054C | 713.464689 | -0.0054023 | 0.09414532 | -0.057383 | 0.9542401 |

| | | | | | | |
|---|---|---|---|---|---|---|
| MSA2 | YKR077W | 259.192054 | 0.00618665 | 0.10855656 | 0.05699012 | 0.95455307 |
| ECM23 | YPL021W | 2.41843659 | -0.0139284 | 0.24455805 | -0.0569532 | 0.9545825 |
| YHL048C-A | YHL048C-A | 1.63883373 | -0.0128324 | 0.22627677 | -0.0567111 | 0.95477532 |
| Xu_2009_SUTs59! | Xu_2009_SU | 12.7624177 | -0.0161613 | 0.28937129 | -0.0558498 | 0.95546144 |
| Xu_2009_CUTs41 | Xu_2009_CU | 16.4760582 | -0.0154458 | 0.27983132 | -0.0551968 | 0.95598167 |
| Xu_2009_CUTs80 | Xu_2009_CU | 8.22558481 | -0.0161902 | 0.29624237 | -0.0546517 | 0.95641592 |
| YGR242W | YGR242W | 1.81291155 | 0.0119667 | 0.21736128 | 0.05505444 | 0.95609509 |
| RIM20 | YOR275C | 1009.298 | -0.0037809 | 0.06912174 | -0.0546988 | 0.95637841 |
| APL4 | YPR029C | 1690.84952 | -0.0036634 | 0.06681288 | -0.054831 | 0.95627311 |
| RSC2 | YLR357W | 1699.53953 | -0.0037195 | 0.06875305 | -0.0541001 | 0.95685543 |
| NPL4 | YBR170C | 1221.89856 | -0.0036276 | 0.06724524 | -0.0539463 | 0.95697798 |
| YER181C | YER181C | 13.2792135 | 0.01529125 | 0.28857122 | 0.05298952 | 0.95774025 |
| TCB3 | YML072C | 6374.7834 | 0.00258081 | 0.04895693 | 0.05271588 | 0.95795829 |
| Xu_2009_CUTs12 | Xu_2009_CU | 2.36438461 | 0.01280111 | 0.24481467 | 0.05228896 | 0.95829845 |
| YER119C-A | YER119C-A | 4.72429004 | 0.01470961 | 0.28430849 | 0.05173822 | 0.95873729 |
| MLH3 | YPL164C | 360.356283 | 0.00478723 | 0.09437791 | 0.05072403 | 0.95954543 |
| UBX7 | YBR273C | 771.990388 | 0.00432754 | 0.08566805 | 0.05051525 | 0.9597118 |
| TFC1 | YBR123C | 524.572885 | -0.004663 | 0.09295719 | -0.0501631 | 0.95999244 |
| Xu_2009_CUTs47 | Xu_2009_CU | 22.6210317 | -0.0133282 | 0.26811932 | -0.0497098 | 0.96035363 |
| YHR139C-A | YHR139C-A | 4.81467638 | 0.01438846 | 0.28945474 | 0.04970883 | 0.96035442 |
| SNR60 | snR60 | 32.2112593 | 0.01132972 | 0.23873934 | 0.04745646 | 0.96214943 |
| tI(AAU)E1 | tI(AAU)E1 | 1.10437114 | 0.00899919 | 0.18640018 | 0.04827887 | 0.961494 |
| Xu_2009_CUTs10 | Xu_2009_CU | 42.2042815 | 0.01034437 | 0.21821643 | 0.04740417 | 0.96219111 |
| Xu_2009_CUTs62 | Xu_2009_CU | 3.2042153 | -0.0126599 | 0.26277662 | -0.0481773 | 0.96157495 |
| Xu_2009_SUTs78! | Xu_2009_SU | 0.9154876 | -0.0082797 | 0.17392254 | -0.0476057 | 0.96203048 |
| PAU8 | YAL068C | 3.11933624 | -0.0129416 | 0.26830727 | -0.0482343 | 0.96152948 |
| MIN7 | YBR201C-A | 160.677005 | -0.006269 | 0.13043216 | -0.0480629 | 0.9616661 |
| VAC8 | YEL013W | 1408.26499 | -0.0033417 | 0.07025314 | -0.047567 | 0.96206137 |
| ECO1 | YFR027W | 120.75916 | -0.007397 | 0.15558134 | -0.047544 | 0.96207967 |
| tA(AGC)D | tA(AGC)D | 3.35941527 | 0.01251284 | 0.27135258 | 0.04611284 | 0.96322031 |
| YDL152W | YDL152W | 4.58915598 | -0.013074 | 0.28395001 | -0.0460433 | 0.96327575 |
| YOR012W | YOR012W | 52.4665914 | 0.00940289 | 0.20423091 | 0.04604049 | 0.96327798 |
| Xu_2009_SUTs30 | Xu_2009_SU | 25.6955073 | -0.0113372 | 0.25429923 | -0.044582 | 0.9644405 |
| Xu_2009_CUTs94 | Xu_2009_CU | 16.8807365 | -0.0121352 | 0.27757537 | -0.0437187 | 0.96512863 |
| Xu_2009_SUTs37 | Xu_2009_SU | 15.7577977 | 0.01203789 | 0.27607359 | 0.0436039 | 0.96522014 |
| VPS15 | YBR097W | 694.115706 | -0.0036889 | 0.08394052 | -0.0439466 | 0.96494698 |
| AUS1 | YOR011W | 385.126558 | -0.0045929 | 0.10516058 | -0.0436756 | 0.96516302 |
| TRM11 | YOL124C | 1530.83254 | -0.0037042 | 0.08528339 | -0.0434337 | 0.96535581 |
| YEL067C | YEL067C | 83.3655967 | 0.00767577 | 0.17831072 | 0.04304715 | 0.96566395 |
| NIC96 | YFR002W | 1422.19343 | -0.003197 | 0.07421315 | -0.0430788 | 0.96563869 |
| COA2 | YPL189C-A | 179.260106 | -0.0060868 | 0.14323368 | -0.0424957 | 0.9661035 |

| | | | | | | |
|---|---|---|---|---|---|---|
| tR(UCU)M2 | tR(UCU)M2 | 1.84709863 | 0.00982236 | 0.23348438 | 0.0420686 | 0.96644401 |
| Xu_2009_CUTs73 | Xu_2009_CU | 2.29148406 | -0.0097594 | 0.23234757 | -0.0420035 | 0.96649595 |
| Xu_2009_SUTs57 | Xu_2009_SU | 46.5491794 | 0.00876269 | 0.2103093 | 0.04166574 | 0.96676516 |
| YFR036W-A | YFR036W-A | 2.71961627 | -0.010349 | 0.26130107 | -0.0396058 | 0.96840741 |
| YNR071C | YNR071C | 42.4640344 | 0.00890958 | 0.22576861 | 0.03946332 | 0.96852099 |
| Xu_2009_SUTs32 | Xu_2009_SU | 21.4595293 | 0.01024173 | 0.2624734 | 0.03902008 | 0.96887438 |
| SPT20 | YOL148C | 793.760219 | -0.0029834 | 0.07629804 | -0.0391013 | 0.9688096 |
| TFC7 | YOR110W | 613.276604 | -0.0031209 | 0.08005101 | -0.038986 | 0.96890158 |
| Xu_2009_CUTs38 | Xu_2009_CU | 10.15634 | -0.0114005 | 0.29627565 | -0.0384795 | 0.96930537 |
| YEH1 | YLL012W | 511.125207 | 0.00331187 | 0.08665525 | 0.03821887 | 0.96951317 |
| Xu_2009_SUTs12 | Xu_2009_SU | 15.4206674 | 0.0106557 | 0.28125723 | 0.03788594 | 0.96977862 |
| BDP1 | YNL039W | 756.898381 | 0.00277984 | 0.07350308 | 0.03781939 | 0.96983168 |
| YCR085W | YCR085W | 16.2516963 | -0.0105846 | 0.28376864 | -0.0373001 | 0.9702457 |
| YGL117W | YGL117W | 224.301204 | 0.00424871 | 0.11410398 | 0.03723545 | 0.97029728 |
| Xu_2009_CUTs41 | Xu_2009_CU | 11.6522852 | -0.0106982 | 0.29293999 | -0.0365202 | 0.97086754 |
| Xu_2009_SUTs83 | Xu_2009_SU | 119.902562 | 0.00600127 | 0.16209427 | 0.03702335 | 0.97046639 |
| NCL1 | YBL024W | 2805.23806 | 0.00211881 | 0.05791853 | 0.03658257 | 0.97081784 |
| YBR124W | YBR124W | 2.68039401 | 0.00941758 | 0.25772892 | 0.03654066 | 0.97085126 |
| CFD1 | YIL003W | 312.299231 | -0.0036958 | 0.10122194 | -0.0365119 | 0.97087416 |
| Xu_2009_SUTs10 | Xu_2009_SU | 2.77230924 | 0.00885435 | 0.24543086 | 0.03607676 | 0.97122115 |
| ATP10 | YLR393W | 322.876331 | 0.00367656 | 0.10169128 | 0.03615417 | 0.97115943 |
| YTA12 | YMR089C | 2570.96478 | -0.0021399 | 0.0593462 | -0.0360587 | 0.97123559 |
| KRE33 | YNL132W | 2130.4316 | 0.00300776 | 0.08370452 | 0.03593311 | 0.97133569 |
| Xu_2009_SUTs48 | Xu_2009_SU | 37.4445119 | 0.00794675 | 0.2263421 | 0.03510948 | 0.97199244 |
| OSW2 | YLR054C | 155.377844 | 0.0050544 | 0.14428618 | 0.03503039 | 0.97205551 |
| DSC3 | YOR223W | 219.494627 | -0.0048058 | 0.13703159 | -0.0350709 | 0.97202324 |
| RUF5-2 | RUF5-2 | 3.52468548 | 0.00948279 | 0.27313207 | 0.03471869 | 0.97230406 |
| CDC7 | YDL017W | 358.759321 | -0.0032692 | 0.09474104 | -0.0345071 | 0.97247281 |
| YFR057W | YFR057W | 1.18787355 | 0.00680768 | 0.19949405 | 0.03412473 | 0.97277769 |
| Xu_2009_SUTs29 | Xu_2009_SU | 74.3890183 | -0.006096 | 0.1805838 | -0.033757 | 0.97307094 |
| YML047W-A | YML047W-A | 5.37777036 | -0.0096336 | 0.29040521 | -0.0331731 | 0.97353654 |
| Xu_2009_CUTs84 | Xu_2009_CU | 41.5737826 | -0.0077029 | 0.24117193 | -0.0319395 | 0.9745203 |
| YMR007W | YMR007W | 11.2629383 | 0.009385 | 0.29420776 | 0.03189923 | 0.97455242 |
| Xu_2009_CUTs29 | Xu_2009_CU | 5.41483975 | -0.0091984 | 0.28988676 | -0.0317309 | 0.97468667 |
| YNL017C | YNL017C | 4.04747404 | -0.0088835 | 0.28138397 | -0.0315708 | 0.97481435 |
| LOT5 | YKL183W | 294.915178 | -0.0031924 | 0.10277546 | -0.0310614 | 0.97522056 |
| HEH2 | YDR458C | 881.769761 | -0.0021226 | 0.06978067 | -0.0304189 | 0.97573299 |
| YPR071W | YPR071W | 301.564788 | -0.0032957 | 0.10885458 | -0.0302762 | 0.97584678 |
| DAL82 | YNL314W | 278.839596 | -0.0033848 | 0.12072975 | -0.0280365 | 0.97763301 |
| YDR541C | YDR541C | 291.632619 | 0.00293331 | 0.10777971 | 0.02721577 | 0.97828764 |
| SAS2 | YMR127C | 298.156158 | -0.0028137 | 0.10319655 | -0.0272653 | 0.97824814 |

| | | | | | | |
|---|---|---|---|---|---|---|
| MGM1 | YOR211C | 790.165055 | 0.00200396 | 0.07317529 | 0.02738572 | 0.97815209 |
| Xu_2009_CUTs37 | Xu_2009_CU | 17.2221226 | -0.0074147 | 0.27560829 | -0.0269031 | 0.97853702 |
| PET130 | YJL023C | 208.516654 | 0.00346608 | 0.12824039 | 0.02702796 | 0.97843744 |
| Xu_2009_SUTs16 | Xu_2009_SU | 3.29978712 | -0.0071104 | 0.26763417 | -0.0265677 | 0.97880454 |
| BMT6 | YLR063W | 321.029545 | -0.0026881 | 0.1022275 | -0.0262953 | 0.97902182 |
| YER039C-A | YER039C-A | 15.4637826 | 0.00712654 | 0.28326459 | 0.02515861 | 0.97992845 |
| YOR102W | YOR102W | 35.7792037 | 0.00608515 | 0.24207974 | 0.02513696 | 0.97994572 |
| YPR076W | YPR076W | 8.06821797 | 0.007404 | 0.2965236 | 0.02496933 | 0.98007943 |
| PLP1 | YDR183W | 202.475774 | -0.0029267 | 0.11981503 | -0.0244267 | 0.98051228 |
| YKL097C | YKL097C | 2.97620529 | 0.00657323 | 0.26837782 | 0.02449245 | 0.98045981 |
| Xu_2009_SUTs73 | Xu_2009_SU | 2.8243477 | -0.0061454 | 0.263657 | -0.0233084 | 0.98140426 |
| tG(GCC)O1 | tG(GCC)O1 | 2.1293343 | -0.0055352 | 0.23946718 | -0.0231147 | 0.9815588 |
| YGL132W | YGL132W | 2.05946018 | 0.00541343 | 0.2352139 | 0.02301493 | 0.98163837 |
| tE(UUC)E1 | tE(UUC)E1 | 10.4850128 | 0.00663479 | 0.29519594 | 0.02247588 | 0.98206835 |
| LYS4 | YDR234W | 1269.21912 | 0.00147878 | 0.06669064 | 0.02217373 | 0.98230937 |
| tG(UCC)O/SUF1 | tG(UCC)O | 0.83741744 | -0.0036151 | 0.16682852 | -0.0216693 | 0.98271171 |
| MTR3 | YGR158C | 398.708015 | 0.00187047 | 0.09221718 | 0.02028333 | 0.98381735 |
| Xu_2009_CUTs72 | Xu_2009_CU | 73.640869 | 0.00356191 | 0.1805258 | 0.01973073 | 0.98425818 |
| YBL044W | YBL044W | 3.56157582 | 0.00521607 | 0.27151516 | 0.01921096 | 0.98467281 |
| Xu_2009_CUTs19 | Xu_2009_CU | 62.0973635 | -0.00361 | 0.19511537 | -0.0185019 | 0.98523848 |
| THI13 | YDL244W | 42.3941394 | -0.0040428 | 0.22032528 | -0.0183491 | 0.98536033 |
| Xu_2009_CUTs49 | Xu_2009_CU | 10.5812086 | -0.0051216 | 0.29443614 | -0.0173948 | 0.98612168 |
| TRS85 | YDR108W | 517.838261 | 0.00151372 | 0.08880069 | 0.01704632 | 0.98639966 |
| Xu_2009_CUTs60 | Xu_2009_CU | 22.5747964 | -0.0043901 | 0.26252956 | -0.0167225 | 0.98665803 |
| GPI2 | YPL076W | 184.814668 | 0.00200739 | 0.12675773 | 0.01583645 | 0.98736487 |
| Xu_2009_SUTs58 | Xu_2009_SU | 45.4654293 | 0.00339547 | 0.21866665 | 0.01552805 | 0.98761091 |
| TYR1 | YBR166C | 606.159587 | -0.0012696 | 0.08320781 | -0.0152576 | 0.98782665 |
| YLR041W | YLR041W | 21.7598008 | -0.0040291 | 0.26447331 | -0.0152344 | 0.9878452 |
| TIR3 | YIL011W | 44.7972736 | -0.0032057 | 0.21564647 | -0.0148654 | 0.98813954 |
| YMR057C | YMR057C | 6.00166587 | -0.0042304 | 0.29398773 | -0.0143897 | 0.98851908 |
| HPF1 | YOL155C | 3728.62808 | -0.0007459 | 0.05994596 | -0.0124422 | 0.9900728 |
| YCR022C | YCR022C | 8.63167732 | 0.00352084 | 0.29633896 | 0.01188113 | 0.99052045 |
| DGR1 | YNL130C-A | 3.65733824 | 0.00325647 | 0.27740362 | 0.01173909 | 0.99063378 |
| YML034C-A | YML034C-A | 3.1676925 | 0.00284863 | 0.26518868 | 0.01074191 | 0.99142936 |
| Xu_2009_CUTs45 | Xu_2009_CU | 0.86722578 | -0.0016635 | 0.17578679 | -0.009463 | 0.99244977 |
| Xu_2009_SUTs25 | Xu_2009_SU | 17.3209938 | 0.00260351 | 0.27571387 | 0.00944279 | 0.99246585 |
| GNP1 | YDR508C | 2816.78486 | -0.0006632 | 0.07044436 | -0.009414 | 0.99248882 |
| tK(CUU)G1 | tK(CUU)G1 | 2.20726838 | -0.002182 | 0.24254192 | -0.0089963 | 0.99282205 |
| SFK1 | YKL051W | 627.717929 | -0.000672 | 0.08089673 | -0.0083073 | 0.9933718 |
| Xu_2009_CUTs67 | Xu_2009_CU | 2.55993307 | 0.00202058 | 0.25576815 | 0.00790004 | 0.99369675 |
| Xu_2009_SUTs55 | Xu_2009_SU | 13.7187175 | -0.0022328 | 0.28601477 | -0.0078065 | 0.99377134 |

| | | | | | | |
|---|---|---|---|---|---|---|
| Xu_2009_CUTs27 | Xu_2009_CU | 15.9532869 | 0.00205304 | 0.27487932 | 0.00746886 | 0.99404076 |
| YLR406C-A | YLR406C-A | 10.8062556 | 0.00209497 | 0.29513108 | 0.00709843 | 0.99433632 |
| MAK32 | YCR019W | 324.756762 | -0.0007586 | 0.11015474 | -0.0068866 | 0.99450534 |
| VID30 | YGL227W | 647.81834 | -0.0007041 | 0.10371192 | -0.0067886 | 0.99458355 |
| Xu_2009_CUTs74 | Xu_2009_CU | 13.8906266 | -0.0016987 | 0.28600225 | -0.0059396 | 0.99526095 |
| SOF1 | YLL011W | 1246.97714 | 0.00042786 | 0.0709307 | 0.00603209 | 0.99518712 |
| SMP1 | YBR182C | 827.380891 | -0.0004626 | 0.08392686 | -0.0055114 | 0.9956026 |
| DUF1 | YOL087C | 1468.45313 | -0.0003819 | 0.06834567 | -0.0055871 | 0.99554214 |
| VBA2 | YBR293W | 330.936278 | 0.00056927 | 0.10816379 | 0.00526307 | 0.99580069 |
| NUP85 | YJR042W | 1135.47874 | 0.00030475 | 0.06947926 | 0.00438621 | 0.99650032 |
| Xu_2009_SUTs55 | Xu_2009_SU | 69.4122662 | -0.0006908 | 0.18894057 | -0.0036564 | 0.99708261 |
| Xu_2009_CUTs85 | Xu_2009_CU | 8.04974906 | 0.0009926 | 0.29634399 | 0.00334948 | 0.9973275 |
| SPR3 | YGR059W | 73.7805964 | 0.00053142 | 0.18252848 | 0.00291142 | 0.99767702 |
| Xu_2009_CUTs41 | Xu_2009_CU | 22.7095715 | 0.00067079 | 0.26374575 | 0.00254332 | 0.99797073 |
| tQ(UUG)D1 | tQ(UUG)D1 | 1.10264459 | 0.00025727 | 0.19159708 | 0.00134279 | 0.99892861 |
| VID27 | YNL212W | 905.467633 | -9.07E-05 | 0.06879143 | -0.0013181 | 0.99894828 |
| tP(UGG)N2 | tP(UGG)N2 | 3.95191217 | 0.00022798 | 0.27646694 | 0.00082463 | 0.99934204 |
| YEF1 | YEL041W | 189.104681 | -0.000131 | 0.12454202 | -0.0010523 | 0.99916042 |
| YPR053C | YPR053C | 94.3616791 | -0.0001362 | 0.16124723 | -0.0008445 | 0.99932617 |
| Xu_2009_SUTs67 | Xu_2009_SU | 18.9357696 | 0.00016162 | 0.27031876 | 0.00059788 | 0.99952296 |
| YOR389W | YOR389W | 796.821676 | 2.52E-05 | 0.07207995 | 0.00034901 | 0.99972153 |
| Q0010 | Q0010 | 0.35678379 | 0.01468767 | 0.1143886 | 0.12840153 | 0.89783122 |
| Q0017 | Q0017 | 0 | NA | NA | NA | NA |
| Q0032 | Q0032 | 0.274372 | -0.0842871 | 0.10660478 | -0.7906499 | 0.4291483 |
| Q0092 | Q0092 | 0 | NA | NA | NA | NA |
| Q0142 | Q0142 | 0 | NA | NA | NA | NA |
| Q0143 | Q0143 | 0 | NA | NA | NA | NA |
| Q0144 | Q0144 | 0 | NA | NA | NA | NA |
| Q0297 | Q0297 | 0 | NA | NA | NA | NA |
| RDN18-1 | RDN18-1 | 0 | NA | NA | NA | NA |
| RDN18-2 | RDN18-2 | 0 | NA | NA | NA | NA |
| RDN25-1 | RDN25-1 | 0 | NA | NA | NA | NA |
| RDN25-2 | RDN25-2 | 0 | NA | NA | NA | NA |
| RDN37-1 | RDN37-1 | 0.33047709 | 0 | 0.11278862 | 0 | 1 |
| RDN5-1 | RDN5-1 | 0.15192853 | 0.02739462 | 0.08450585 | 0.32417424 | 0.74580613 |
| RDN5-2 | RDN5-2 | 0 | NA | NA | NA | NA |
| RDN5-3 | RDN5-3 | 0 | NA | NA | NA | NA |
| RDN5-4 | RDN5-4 | 0 | NA | NA | NA | NA |
| RDN5-5 | RDN5-5 | 0 | NA | NA | NA | NA |
| RDN5-6 | RDN5-6 | 0.12935759 | -0.03236 | 0.08450585 | -0.3829325 | 0.70176978 |
| RDN58-1 | RDN58-1 | 0 | NA | NA | NA | NA |

| | | | | | | |
|---|---|---|---|---|---|---|
| RDN58-2 | RDN58-2 | 0 | NA | NA | NA | NA |
| RUF20 | RUF20 | 0 | NA | NA | NA | NA |
| RUF5-1 | RUF5-1 | 0 | NA | NA | NA | NA |
| SNR7-S | snR7-S | 0 | NA | NA | NA | NA |
| SNR73 | snR73 | 0.20532185 | -0.0164717 | 0.09689938 | -0.1699872 | 0.86502017 |
| tA(AGC)F | tA(AGC)F | 0 | NA | NA | NA | NA |
| tA(AGC)K1 | tA(AGC)K1 | 0.42679838 | -0.1080644 | 0.120126 | -0.8995925 | 0.36833716 |
| tA(AGC)K2 | tA(AGC)K2 | 0 | NA | NA | NA | NA |
| tA(AGC)M1 | tA(AGC)M1 | 0 | NA | NA | NA | NA |
| tA(UGC)Q | tA(UGC)Q | 0.53473626 | -0.0311231 | 0.13525534 | -0.2301065 | 0.81800898 |
| tC(GCA)B | tC(GCA)B | 0.12222156 | 0 | 0.08450585 | 0 | 1 |
| tC(GCA)G | tC(GCA)G | 0.07596426 | 0.0074764 | 0.08450585 | 0.08847198 | 0.92950156 |
| tC(GCA)P2 | tC(GCA)P2 | 0.23228177 | -0.0649031 | 0.09391765 | -0.6910642 | 0.48952517 |
| tC(GCA)Q | tC(GCA)Q | 0.23202636 | 0 | 0.10088369 | 0 | 1 |
| tD(GUC)J4 | tD(GUC)J4 | 0.50866259 | 0.1046443 | 0.13570402 | 0.77112157 | 0.44063488 |
| tD(GUC)K | tD(GUC)K | 0.53553161 | 0.06706544 | 0.13594727 | 0.49331953 | 0.62178683 |
| tD(GUC)Q | tD(GUC)Q | 0 | NA | NA | NA | NA |
| tE(CUC)D | tE(CUC)D | 0 | NA | NA | NA | NA |
| tG(GCC)C/SUF16 | tG(GCC)C | 0.39025748 | -0.0719166 | 0.12433962 | -0.5783881 | 0.56300215 |
| tG(GCC)D1 | tG(GCC)D1 | 0.62448636 | -0.0075273 | 0.14850454 | -0.0506872 | 0.95957478 |
| tG(GCC)M | tG(GCC)M | 0.54418116 | 0.01354464 | 0.14215078 | 0.09528365 | 0.92408953 |
| tG(UCC)Q | tG(UCC)Q | 0 | NA | NA | NA | NA |
| tH(GUG)Q | tH(GUG)Q | 0.11614088 | -0.03236 | 0.08450585 | -0.3829325 | 0.70176978 |
| tI(AAU)B | tI(AAU)B | 0 | NA | NA | NA | NA |
| tI(AAU)D | tI(AAU)D | 0.28002867 | 0.07734829 | 0.10684798 | 0.72390973 | 0.46912116 |
| tI(AAU)L1 | tI(AAU)L1 | 0.52866291 | 0.02083904 | 0.14241424 | 0.14632691 | 0.88366332 |
| tI(AAU)L2 | tI(AAU)L2 | 0.61024664 | -0.2033485 | 0.14858023 | -1.3686107 | 0.17112101 |
| tI(GAU)Q | tI(GAU)Q | 0.274372 | -0.0842871 | 0.10660478 | -0.7906499 | 0.4291483 |
| tK(CUU)C | tK(CUU)C | 0 | NA | NA | NA | NA |
| tK(CUU)D2 | tK(CUU)D2 | 0 | NA | NA | NA | NA |
| tK(CUU)E1 | tK(CUU)E1 | 0.11614088 | -0.03236 | 0.08450585 | -0.3829325 | 0.70176978 |
| tK(CUU)G2 | tK(CUU)G2 | 0.23202636 | 0 | 0.10088369 | 0 | 1 |
| tK(UUU)Q | tK(UUU)Q | 0.44262895 | 0.01349203 | 0.12125665 | 0.11126839 | 0.91140352 |
| tL(UAA)K | tL(UAA)K | 0.60799609 | -0.093587 | 0.14521158 | -0.6444871 | 0.51925963 |
| tL(UAA)Q | tL(UAA)Q | 0 | NA | NA | NA | NA |
| tM(CAU)C | tM(CAU)C | 0 | NA | NA | NA | NA |
| tM(CAU)J1/EMT5 | tM(CAU)J1 | 0.41559819 | 0.02082891 | 0.13240125 | 0.15731657 | 0.87499536 |
| tM(CAU)Q2 | tM(CAU)Q2 | 0 | NA | NA | NA | NA |
| tN(GUU)C | tN(GUU)C | 0.43273868 | 0.02166069 | 0.13424878 | 0.16134738 | 0.87181981 |
| tN(GUU)L | tN(GUU)L | 0.67327014 | -0.0508888 | 0.16091583 | -0.3162448 | 0.75181673 |
| tN(GUU)Q | tN(GUU)Q | 0.26803592 | -0.0515605 | 0.10599835 | -0.4864278 | 0.62666386 |

| | | | | | | |
|---|---|---|---|---|---|---|
| tP(UGG)N1 | tP(UGG)N1 | 0.15823112 | -0.03236 | 0.08450585 | -0.3829325 | 0.70176978 |
| tP(UGG)O1 | tP(UGG)O1 | 0 | NA | NA | NA | NA |
| tP(UGG)O2/SUF1 | tP(UGG)O2 | 0.31469355 | 0.0135753 | 0.1100811 | 0.12332088 | 0.90185301 |
| tQ(UUG)Q | tQ(UUG)Q | 0.37485605 | -0.1419663 | 0.1173058 | -1.2102241 | 0.2261929 |
| tR(ACG)Q2 | tR(ACG)Q2 | 0 | NA | NA | NA | NA |
| tR(CCU)J/HSX1 | tR(CCU)J | 0.33628183 | -0.0663286 | 0.11965012 | -0.5543551 | 0.57933588 |
| tR(UCU)B | tR(UCU)B | 0 | NA | NA | NA | NA |
| tR(UCU)D | tR(UCU)D | 0 | NA | NA | NA | NA |
| tR(UCU)J1 | tR(UCU)J1 | 0 | NA | NA | NA | NA |
| tR(UCU)J2 | tR(UCU)J2 | 0 | NA | NA | NA | NA |
| tR(UCU)Q1 | tR(UCU)Q1 | 0 | NA | NA | NA | NA |
| tS(AGA)D1 | tS(AGA)D1 | 0.41869361 | 0.06625379 | 0.12652595 | 0.52363793 | 0.6005304 |
| tS(AGA)H | tS(AGA)H | 0 | NA | NA | NA | NA |
| tS(AGA)L | tS(AGA)L | 0 | NA | NA | NA | NA |
| tS(GCU)Q1/SUP1 | tS(GCU)Q1 | 0.4887082 | -0.0766723 | 0.13233286 | -0.5793898 | 0.56232616 |
| tT(AGU)J | tT(AGU)J | 0 | NA | NA | NA | NA |
| tT(AGU)O1 | tT(AGU)O1 | 0 | NA | NA | NA | NA |
| tT(UGU)H | tT(UGU)H | 0 | NA | NA | NA | NA |
| tT(UGU)P | tT(UGU)P | 0 | NA | NA | NA | NA |
| tT(UGU)Q1 | tT(UGU)Q1 | 0 | NA | NA | NA | NA |
| tT(XXX)Q2 | tT(XXX)Q2 | 0.2196096 | 0 | 0.09176909 | 0 | 1 |
| tV(UAC)Q | tV(UAC)Q | 0 | NA | NA | NA | NA |
| tW(UCA)Q | tW(UCA)Q | 0.11614088 | -0.03236 | 0.08450585 | -0.3829325 | 0.70176978 |
| tY(GUA)M2/SUP8 | tY(GUA)M2 | 0.60688242 | -0.194831 | 0.14694419 | -1.3258841 | 0.18487808 |
| Xu_2009_CUTs10 | Xu_2009_CU | 0.1752406 | 0.03229832 | 0.0915842 | 0.35266261 | 0.72434139 |
| Xu_2009_CUTs18 | Xu_2009_CU | 0.1752406 | 0.03229832 | 0.0915842 | 0.35266261 | 0.72434139 |
| Xu_2009_CUTs24 | Xu_2009_CU | 0.68608196 | 0.03020595 | 0.1531996 | 0.19716731 | 0.84369661 |
| Xu_2009_CUTs29 | Xu_2009_CU | 0.15823112 | -0.03236 | 0.08450585 | -0.3829325 | 0.70176978 |
| Xu_2009_CUTs44 | Xu_2009_CU | 0.09927633 | 0.0074764 | 0.08450585 | 0.08847198 | 0.92950156 |
| Xu_2009_CUTs48 | Xu_2009_CU | 0 | NA | NA | NA | NA |
| Xu_2009_CUTs54 | Xu_2009_CU | 0.58127476 | 0.10594475 | 0.15299052 | 0.69249227 | 0.48862824 |
| Xu_2009_CUTs54 | Xu_2009_CU | 0 | NA | NA | NA | NA |
| Xu_2009_CUTs61 | Xu_2009_CU | 0.63507527 | 0.01442768 | 0.15471534 | 0.09325305 | 0.92570253 |
| Xu_2009_CUTs64 | Xu_2009_CU | 0.33822894 | 0.07291837 | 0.11291998 | 0.64575262 | 0.51843958 |
| Xu_2009_CUTs69 | Xu_2009_CU | 0.55919549 | 0.14795296 | 0.14472852 | 1.02227922 | 0.30664877 |
| Xu_2009_CUTs75 | Xu_2009_CU | 0.11614088 | -0.03236 | 0.08450585 | -0.3829325 | 0.70176978 |
| Xu_2009_CUTs80 | Xu_2009_CU | 0.09927633 | 0.0074764 | 0.08450585 | 0.08847198 | 0.92950156 |
| Xu_2009_SUTs17 | Xu_2009_SU | 0 | NA | NA | NA | NA |
| Xu_2009_SUTs23 | Xu_2009_SU | 0.09037617 | 0.0074764 | 0.08450585 | 0.08847198 | 0.92950156 |
| Xu_2009_SUTs24 | Xu_2009_SU | 0.63978432 | 0.11042605 | 0.15009951 | 0.73568564 | 0.46192203 |
| Xu_2009_SUTs30 | Xu_2009_SU | 0.60797927 | 0.13167659 | 0.15731873 | 0.83700516 | 0.40258967 |

| | | | | | | |
|---|---|---|---|---|---|---|
| Xu_2009_SUTs39 | Xu_2009_SU | 0.34869859 | -0.0674752 | 0.12062313 | -0.5593883 | 0.57589673 |
| Xu_2009_SUTs51 | Xu_2009_SU | 0.11033614 | 0 | 0.08450585 | 0 | 1 |
| Xu_2009_SUTs61 | Xu_2009_SU | 0.41832677 | 0.01651154 | 0.13301247 | 0.12413531 | 0.90120814 |
| Xu_2009_SUTs73 | Xu_2009_SU | 0.26806941 | 0.010341 | 0.10553485 | 0.09798663 | 0.92194291 |
| Xu_2009_SUTs77 | Xu_2009_SU | 0 | NA | NA | NA | NA |
| Xu_2009_SUTs80 | Xu_2009_SU | 0.1098048 | 0 | 0.08450585 | 0 | 1 |
| Xu_2009_SUTs84 | Xu_2009_SU | 0.22863392 | -0.0177707 | 0.10053583 | -0.1767595 | 0.8596973 |
| YAL016C-B | YAL016C-B | 0.56450856 | -0.0896265 | 0.14453486 | -0.620103 | 0.53518995 |
| YAL067W-A | YAL067W-A | 0 | NA | NA | NA | NA |
| YAL068W-A | YAL068W-A | 0 | NA | NA | NA | NA |
| YAL069W | YAL069W | 0 | NA | NA | NA | NA |
| YAR035C-A | YAR035C-A | 0.11614088 | -0.03236 | 0.08450585 | -0.3829325 | 0.70176978 |
| YAR060C | YAR060C | 0.1098048 | 0 | 0.08450585 | 0 | 1 |
| YAR061W | YAR061W | 0 | NA | NA | NA | NA |
| YAR066W | YAR066W | 0 | NA | NA | NA | NA |
| YAR068W | YAR068W | 0.34183117 | 0 | 0.11380921 | 0 | 1 |
| YAR069C | YAR069C | 0 | NA | NA | NA | NA |
| YBL005W-A | YBL005W-A | 0 | NA | NA | NA | NA |
| YBL039C-A | YBL039C-A | 0 | NA | NA | NA | NA |
| YBL068W-A | YBL068W-A | 0 | NA | NA | NA | NA |
| YBL077W | YBL077W | 0.1752406 | 0.03229832 | 0.0915842 | 0.35266261 | 0.72434139 |
| YBL083C | YBL083C | 0 | NA | NA | NA | NA |
| YBL100C | YBL100C | 0.33587464 | -0.0660913 | 0.11989564 | -0.55124 | 0.58146916 |
| YBL100W-A | YBL100W-A | 0 | NA | NA | NA | NA |
| YBL107W-A | YBL107W-A | 0 | NA | NA | NA | NA |
| PAU9 | YBL108C-A | 0.38432173 | 0.10027193 | 0.12445923 | 0.80566083 | 0.42043844 |
| YBL108W | YBL108W | 0 | NA | NA | NA | NA |
| YBR012W-A | YBR012W-A | 0 | NA | NA | NA | NA |
| YBR027C | YBR027C | 0.40267424 | -0.0651127 | 0.11860745 | -0.5489763 | 0.5830217 |
| LYS2 | YBR115C | 0.09927633 | 0.0074764 | 0.08450585 | 0.08847198 | 0.92950156 |
| YBR121C-A | YBR121C-A | 0 | NA | NA | NA | NA |
| YBR126W-B | YBR126W-B | 0.48817686 | -0.076638 | 0.13230662 | -0.5792457 | 0.56242341 |
| YBR131C-A | YBR131C-A | 0.56506952 | -0.1207806 | 0.12913269 | -0.935322 | 0.34962238 |
| YBR200W-A | YBR200W-A | 0.64816527 | 0.05247914 | 0.15052184 | 0.34864799 | 0.7273536 |
| YBR223W-A | YBR223W-A | 0.20908114 | 0.0100266 | 0.09750913 | 0.10282727 | 0.91810006 |
| YBR226C | YBR226C | 0.59680803 | 0.00238165 | 0.14843193 | 0.01604542 | 0.98719816 |
| YCL020W | YCL020W | 0 | NA | NA | NA | NA |
| YCL042W | YCL042W | 0.43409563 | 0.06482206 | 0.12733936 | 0.50904968 | 0.6107174 |
| YCL065W | YCL065W | 0 | NA | NA | NA | NA |
| HMLALPHA1 | YCL066W | 0.12935759 | -0.03236 | 0.08450585 | -0.3829325 | 0.70176978 |
| HMLALPHA2 | YCL067C | 0.20651705 | -0.0165399 | 0.09709427 | -0.170349 | 0.86473566 |

| | | | | | | |
|---|---|---|---|---|---|---|
| YCL068C | YCL068C | 0.11614088 | -0.03236 | 0.08450585 | -0.3829325 | 0.70176978 |
| VBA3 | YCL069W | 0.20651705 | -0.0165399 | 0.09709427 | -0.170349 | 0.86473566 |
| YCL074W | YCL074W | 0.26226467 | 0.0411228 | 0.10478352 | 0.39245483 | 0.69472218 |
| YCL075W | YCL075W | 0 | NA | NA | NA | NA |
| YCL076W | YCL076W | 0 | NA | NA | NA | NA |
| MATALPHA1 | YCR040W | 0 | NA | NA | NA | NA |
| HMRA2 | YCR096C | 0 | NA | NA | NA | NA |
| HMRA1 | YCR097W | 0 | NA | NA | NA | NA |
| YCR097W-A | YCR097W-A | 0 | NA | NA | NA | NA |
| YCR108C | YCR108C | 0.09037617 | 0.0074764 | 0.08450585 | 0.08847198 | 0.92950156 |
| YDL034W | YDL034W | 0.11033614 | 0 | 0.08450585 | 0 | 1 |
| YDL185C-A | YDL185C-A | 0.07596426 | 0.0074764 | 0.08450585 | 0.08847198 | 0.92950156 |
| YDL240C-A | YDL240C-A | 0.29945731 | 0.05280657 | 0.11550884 | 0.4571648 | 0.64755259 |
| YDR029W | YDR029W | 0.2423047 | 0.06813122 | 0.10238535 | 0.66543908 | 0.50576969 |
| YDR034C-C | YDR034C-C | 0 | NA | NA | NA | NA |
| YDR098C-A | YDR098C-A | 0 | NA | NA | NA | NA |
| YDR149C | YDR149C | 0.61716075 | -0.054773 | 0.15035426 | -0.364293 | 0.71563921 |
| YDR209C | YDR209C | 0.42718659 | 0.05847832 | 0.12040439 | 0.48568266 | 0.62719218 |
| YDR210C-C | YDR210C-C | 0 | NA | NA | NA | NA |
| YDR210W-A | YDR210W-A | 0 | NA | NA | NA | NA |
| YDR261C-C | YDR261C-C | 0 | NA | NA | NA | NA |
| YDR261W-A | YDR261W-A | 0 | NA | NA | NA | NA |
| YDR290W | YDR290W | 0.07596426 | 0.0074764 | 0.08450585 | 0.08847198 | 0.92950156 |
| YDR316W-A | YDR316W-A | 0 | NA | NA | NA | NA |
| YDR355C | YDR355C | 0 | NA | NA | NA | NA |
| YDR365W-A | YDR365W-A | 0 | NA | NA | NA | NA |
| YDR445C | YDR445C | 0.1098048 | 0 | 0.08450585 | 0 | 1 |
| YDR455C | YDR455C | 0.52430603 | 0.06400932 | 0.13445084 | 0.47607974 | 0.63401757 |
| YDR524W-C | YDR524W-C | 0.61855061 | 0.16348858 | 0.14603742 | 1.11949783 | 0.26292782 |
| YDR543C | YDR543C | 0.20908114 | 0.0100266 | 0.09750913 | 0.10282727 | 0.91810006 |
| YDR545C-A | YDR545C-A | 0 | NA | NA | NA | NA |
| URA3 | YEL021W | 0.12222156 | 0 | 0.08450585 | 0 | 1 |
| YER084W-A | YER084W-A | 0.38056245 | 0.05106827 | 0.1243231 | 0.41077057 | 0.68124078 |
| YER091C-A | YER091C-A | 0.34158103 | 0.1294187 | 0.12185073 | 1.06210856 | 0.2881864 |
| YER133W-A | YER133W-A | 0.288562 | 0.05071908 | 0.11478063 | 0.44187839 | 0.65857721 |
| YER137C-A | YER137C-A | 0 | NA | NA | NA | NA |
| YER159C-A | YER159C-A | 0 | NA | NA | NA | NA |
| YER165C-A | YER165C-A | 0 | NA | NA | NA | NA |
| YER190C-A | YER190C-A | 0 | NA | NA | NA | NA |
| YER190C-B | YER190C-B | 0 | NA | NA | NA | NA |
| YFL002W-B | YFL002W-B | 0 | NA | NA | NA | NA |

| | | | | | | |
|---|---|---|---|---|---|---|
| YFL012W-A | YFL012W-A | 0.12935759 | -0.03236 | 0.08450585 | -0.3829325 | 0.70176978 |
| YFL067W | YFL067W | 0.39423323 | 0.14098304 | 0.11973584 | 1.17745064 | 0.23901569 |
| YFL068W | YFL068W | 0 | NA | NA | NA | NA |
| YFR009W-A | YFR009W-A | 0 | NA | NA | NA | NA |
| YFR052C-A | YFR052C-A | 0.41078356 | 0.0571292 | 0.12581663 | 0.45406713 | 0.64978051 |
| YFR056C | YFR056C | 0.2850454 | 0.05016066 | 0.11417129 | 0.43934566 | 0.66041109 |
| YGL014C-A | YGL014C-A | 0.20651705 | -0.0165399 | 0.09709427 | -0.170349 | 0.86473566 |
| YGL149W | YGL149W | 0.39497435 | 0.06170728 | 0.13164129 | 0.46875319 | 0.63924606 |
| YGL239C | YGL239C | 0.09037617 | 0.0074764 | 0.08450585 | 0.08847198 | 0.92950156 |
| YGL260W | YGL260W | 0.22789279 | 0.05479485 | 0.09257103 | 0.5919222 | 0.55390269 |
| YGR027W-A | YGR027W-A | 0 | NA | NA | NA | NA |
| YGR035W-A | YGR035W-A | 0.31187407 | 0.0540975 | 0.11686369 | 0.46291111 | 0.64342809 |
| YGR038C-A | YGR038C-A | 0 | NA | NA | NA | NA |
| YGR051C | YGR051C | 0.43238121 | 0.00905067 | 0.12820359 | 0.07059609 | 0.94371923 |
| YGR109W-A | YGR109W-A | 0 | NA | NA | NA | NA |
| YGR114C | YGR114C | 0.21259773 | 0.01014594 | 0.09807106 | 0.10345503 | 0.91760184 |
| YGR115C | YGR115C | 0.12222156 | 0 | 0.08450585 | 0 | 1 |
| YGR151C | YGR151C | 0 | NA | NA | NA | NA |
| YGR161C-C | YGR161C-C | 0 | NA | NA | NA | NA |
| YGR161W-A | YGR161W-A | 0 | NA | NA | NA | NA |
| YGR226C | YGR226C | 0 | NA | NA | NA | NA |
| YGR265W | YGR265W | 0.39497435 | 0.06170728 | 0.13164129 | 0.46875319 | 0.63924606 |
| YGR270C-A | YGR270C-A | 0.2325577 | 0 | 0.10095377 | 0 | 1 |
| YGR290W | YGR290W | 0.26561677 | 0.08385168 | 0.11233388 | 0.74645052 | 0.45539531 |
| YGR293C | YGR293C | 0.62467932 | 0.06758518 | 0.14880875 | 0.45417479 | 0.64970303 |
| YGR296C-A | YGR296C-A | 0 | NA | NA | NA | NA |
| YGR296C-B | YGR296C-B | 0 | NA | NA | NA | NA |
| YHL009W-A | YHL009W-A | 0 | NA | NA | NA | NA |
| YHL015W-A | YHL015W-A | 0.25750745 | -0.0192917 | 0.10461325 | -0.1844094 | 0.85369228 |
| YHL046W-A | YHL046W-A | 0.23836245 | -0.0475798 | 0.10195837 | -0.4666589 | 0.64074394 |
| YHR032C-A | YHR032C-A | 0.49587899 | 0.05858947 | 0.13974011 | 0.4192745 | 0.67501553 |
| YHR052W-A | YHR052W-A | 0 | NA | NA | NA | NA |
| CUP1-1 | YHR053C | 0.34236251 | 0 | 0.12064729 | 0 | 1 |
| YHR054W-A | YHR054W-A | 0 | NA | NA | NA | NA |
| CUP1-2 | YHR055C | 0.58124895 | -0.1752297 | 0.14013829 | -1.2504053 | 0.21115153 |
| YHR073C-B | YHR073C-B | 0.19855267 | 0.02977116 | 0.0880143 | 0.33825362 | 0.73517207 |
| YHR073W-A | YHR073W-A | 0 | NA | NA | NA | NA |
| YHR165W-A | YHR165W-A | 0.4493196 | 0.01746053 | 0.1290274 | 0.13532421 | 0.89235555 |
| YHR180C-B | YHR180C-B | 0.20961248 | 0.01004469 | 0.09759455 | 0.10292266 | 0.91802435 |
| YHR212C | YHR212C | 0 | NA | NA | NA | NA |
| YHR212W-A | YHR212W-A | 0 | NA | NA | NA | NA |

| | | | | | | |
|---|---|---|---|---|---|---|
| YHR213W-A | YHR213W-A | 0.12935759 | -0.03236 | 0.08450585 | -0.3829325 | 0.70176978 |
| YHR214C-C | YHR214C-C | 0 | NA | NA | NA | NA |
| YHR214C-D | YHR214C-D | 0 | NA | NA | NA | NA |
| YHR218W-A | YHR218W-A | 0.16634043 | 0.03117685 | 0.09001923 | 0.34633541 | 0.72909066 |
| YHR219C-A | YHR219C-A | 0 | NA | NA | NA | NA |
| YIL025C | YIL025C | 0.57171911 | 0.09801715 | 0.13909947 | 0.70465509 | 0.48102491 |
| YIL029W-A | YIL029W-A | 0.21541722 | -0.0170424 | 0.09851585 | -0.1729919 | 0.86265776 |
| YIL030W-A | YIL030W-A | 0.3924265 | -0.0639558 | 0.11733872 | -0.5450524 | 0.58571745 |
| YIL047C-A | YIL047C-A | 0 | NA | NA | NA | NA |
| YIL054W | YIL054W | 0.20018097 | 0.00972027 | 0.09604992 | 0.10120015 | 0.91939158 |
| YIL082W | YIL082W | 0 | NA | NA | NA | NA |
| YIL082W-A | YIL082W-A | 0 | NA | NA | NA | NA |
| YIL171W | YIL171W | 0.58342658 | 0.01628083 | 0.13851782 | 0.11753601 | 0.90643531 |
| YIL171W-A | YIL171W-A | 0 | NA | NA | NA | NA |
| IMA3 | YIL172C | 0 | NA | NA | NA | NA |
| YIL174W | YIL174W | 0 | NA | NA | NA | NA |
| YIL175W | YIL175W | 0 | NA | NA | NA | NA |
| PAU14 | YIL176C | 0 | NA | NA | NA | NA |
| YIL177W-A | YIL177W-A | 0 | NA | NA | NA | NA |
| YIR020C | YIR020C | 0 | NA | NA | NA | NA |
| YIR040C | YIR040C | 0 | NA | NA | NA | NA |
| YJL007C | YJL007C | 0.60933623 | 0.16227837 | 0.14840632 | 1.09347345 | 0.27418598 |
| YJL020W-A | YJL020W-A | 0.5676671 | 0.00287313 | 0.13749069 | 0.02089692 | 0.98332789 |
| YJL127W-A | YJL127W-A | 0.11614088 | -0.03236 | 0.08450585 | -0.3829325 | 0.70176978 |
| YJL175W | YJL175W | 0.09927633 | 0.0074764 | 0.08450585 | 0.08847198 | 0.92950156 |
| YJL197C-A | YJL197C-A | 0.07596426 | 0.0074764 | 0.08450585 | 0.08847198 | 0.92950156 |
| IMA4 | YJL221C | 0.45821976 | 0.02171308 | 0.13544137 | 0.16031353 | 0.8726341 |
| YJL222W-A | YJL222W-A | 0 | NA | NA | NA | NA |
| YJL222W-B | YJL222W-B | 0 | NA | NA | NA | NA |
| PAU1 | YJL223C | 0.09037617 | 0.0074764 | 0.08450585 | 0.08847198 | 0.92950156 |
| YJL225W-A | YJL225W-A | 0 | NA | NA | NA | NA |
| YJR026W | YJR026W | 0 | NA | NA | NA | NA |
| YJR028W | YJR028W | 0 | NA | NA | NA | NA |
| HXT16 | YJR158W | 0.30824603 | -0.0562224 | 0.11026301 | -0.5098935 | 0.61012605 |
| YJR162C | YJR162C | 0.15823112 | -0.03236 | 0.08450585 | -0.3829325 | 0.70176978 |
| YKL123W | YKL123W | 0 | NA | NA | NA | NA |
| YKL145W-A | YKL145W-A | 0 | NA | NA | NA | NA |
| YKL156C-A | YKL156C-A | 0.34909134 | -0.0596624 | 0.11418776 | -0.5224938 | 0.60132655 |
| YKL223W | YKL223W | 0 | NA | NA | NA | NA |
| PAU16 | YKL224C | 0.2196096 | 0 | 0.09176909 | 0 | 1 |
| YKL225W | YKL225W | 0.09037617 | 0.0074764 | 0.08450585 | 0.08847198 | 0.92950156 |

| | | | | | | |
|---|---|---|---|---|---|---|
| YLL019W-A | YLL019W-A | 0.09037617 | 0.0074764 | 0.08450585 | 0.08847198 | 0.92950156 |
| YLL065W | YLL065W | 0.12222156 | 0 | 0.08450585 | 0 | 1 |
| YLL066W-A | YLL066W-A | 0.54524384 | 0.0135297 | 0.14220606 | 0.09514153 | 0.92420241 |
| YLL067W-A | YLL067W-A | 0 | NA | NA | NA | NA |
| YLR154C-G | YLR154C-G | 0 | NA | NA | NA | NA |
| YLR154C-H | YLR154C-H | 0.07596426 | 0.0074764 | 0.08450585 | 0.08847198 | 0.92950156 |
| ASP3-1 | YLR155C | 0 | NA | NA | NA | NA |
| YLR156C-A | YLR156C-A | 0 | NA | NA | NA | NA |
| YLR156W | YLR156W | 0 | NA | NA | NA | NA |
| ASP3-2 | YLR157C | 0 | NA | NA | NA | NA |
| YLR157C-A | YLR157C-A | 0 | NA | NA | NA | NA |
| YLR157C-C | YLR157C-C | 0 | NA | NA | NA | NA |
| YLR157W-D | YLR157W-D | 0 | NA | NA | NA | NA |
| YLR157W-E | YLR157W-E | 0 | NA | NA | NA | NA |
| ASP3-3 | YLR158C | 0 | NA | NA | NA | NA |
| YLR159C-A | YLR159C-A | 0 | NA | NA | NA | NA |
| YLR159W | YLR159W | 0 | NA | NA | NA | NA |
| ASP3-4 | YLR160C | 0 | NA | NA | NA | NA |
| YLR161W | YLR161W | 0 | NA | NA | NA | NA |
| YLR162W | YLR162W | 0.64722949 | -0.0511575 | 0.151759 | -0.3370972 | 0.73604365 |
| YLR184W | YLR184W | 0.288562 | 0.05071908 | 0.11478063 | 0.44187839 | 0.65857721 |
| YLR227W-A | YLR227W-A | 0 | NA | NA | NA | NA |
| YLR280C | YLR280C | 0.09927633 | 0.0074764 | 0.08450585 | 0.08847198 | 0.92950156 |
| YLR399W-A | YLR399W-A | 0.09927633 | 0.0074764 | 0.08450585 | 0.08847198 | 0.92950156 |
| YLR410W-A | YLR410W-A | 0 | NA | NA | NA | NA |
| YLR458W | YLR458W | 0.6008029 | 0.19736179 | 0.14308255 | 1.37935613 | 0.16778498 |
| YLR466C-A | YLR466C-A | 0 | NA | NA | NA | NA |
| YLR467C-A | YLR467C-A | 0 | NA | NA | NA | NA |
| YML031C-A | YML031C-A | 0.26561677 | 0.08385168 | 0.11233388 | 0.74645052 | 0.45539531 |
| YML040W | YML040W | 0 | NA | NA | NA | NA |
| YML045W-A | YML045W-A | 0 | NA | NA | NA | NA |
| YML133W-A | YML133W-A | 0 | NA | NA | NA | NA |
| YML133W-B | YML133W-B | 0 | NA | NA | NA | NA |
| YMR013C-A | YMR013C-A | 0 | NA | NA | NA | NA |
| YMR013W-A | YMR013W-A | 0 | NA | NA | NA | NA |
| YMR046C | YMR046C | 0 | NA | NA | NA | NA |
| YMR051C | YMR051C | 0 | NA | NA | NA | NA |
| YMR135W-A | YMR135W-A | 0.35599294 | 0.13532851 | 0.12320284 | 1.09842036 | 0.27202098 |
| YMR247W-A | YMR247W-A | 0.504655 | 0.0630618 | 0.13327572 | 0.47316798 | 0.63609334 |
| YMR272W-A | YMR272W-A | 0.66018169 | -0.0819898 | 0.14266116 | -0.5747174 | 0.56548247 |
| YMR290W-A | YMR290W-A | 0.58772503 | 0.10251457 | 0.13993889 | 0.7325667 | 0.46382274 |

| | | | | | | |
|---|---|---|---|---|---|---|
| YMR307C-A | YMR307C-A | 0.6117632 | 0.00657856 | 0.14854323 | 0.04428714 | 0.96467552 |
| YMR324C | YMR324C | 0 | NA | NA | NA | NA |
| YMR326C | YMR326C | 0 | NA | NA | NA | NA |
| YNL054W-A | YNL054W-A | 0 | NA | NA | NA | NA |
| YNL103W-A | YNL103W-A | 0.11614088 | -0.03236 | 0.08450585 | -0.3829325 | 0.70176978 |
| RRT16 | YNL105W | 0.622743 | 0.16743792 | 0.14404128 | 1.16243009 | 0.24506081 |
| YNL144W-A | YNL144W-A | 0.49981215 | 0.10509856 | 0.14132141 | 0.74368463 | 0.45706729 |
| YNL184C | YNL184C | 0.5090866 | -0.0830078 | 0.13835603 | -0.5999579 | 0.54853428 |
| YNL284C-A | YNL284C-A | 0 | NA | NA | NA | NA |
| YNL296W | YNL296W | 0.54642564 | -0.0228602 | 0.13639363 | -0.1676046 | 0.86689435 |
| YNL337W | YNL337W | 0.63292902 | 0.11340147 | 0.15577084 | 0.72800191 | 0.46661242 |
| YNL339W-A | YNL339W-A | 0 | NA | NA | NA | NA |
| YNL339W-B | YNL339W-B | 0 | NA | NA | NA | NA |
| YNR005C | YNR005C | 0.39065781 | 0.05744444 | 0.12501311 | 0.45950735 | 0.64586988 |
| YNR075C-A | YNR075C-A | 0 | NA | NA | NA | NA |
| YOL013W-B | YOL013W-B | 0.46931434 | 0.14037926 | 0.13217472 | 1.06207347 | 0.28820233 |
| YOL037C | YOL037C | 0.29663655 | 0.01216779 | 0.10857617 | 0.11206689 | 0.91077037 |
| YOL103W-A | YOL103W-A | 0 | NA | NA | NA | NA |
| YOL166W-A | YOL166W-A | 0 | NA | NA | NA | NA |
| YOR073W-A | YOR073W-A | 0.667867 | 0.25096552 | 0.15620278 | 1.60666494 | 0.10812789 |
| YOR142W-A | YOR142W-A | 0 | NA | NA | NA | NA |
| YOR169C | YOR169C | 0 | NA | NA | NA | NA |
| YOR170W | YOR170W | 0 | NA | NA | NA | NA |
| YOR192C-A | YOR192C-A | 0 | NA | NA | NA | NA |
| YOR231C-A | YOR231C-A | 0.68590451 | -0.138977 | 0.15373363 | -0.9040119 | 0.36598909 |
| YOR263C | YOR263C | 0.66466297 | -0.1026196 | 0.1526335 | -0.6723269 | 0.50137564 |
| YOR335W-A | YOR335W-A | 0 | NA | NA | NA | NA |
| YOR343W-A | YOR343W-A | 0 | NA | NA | NA | NA |
| PAU21 | YOR394W | 0.55241462 | 0.12837877 | 0.1403865 | 0.91446665 | 0.36047169 |
| YOR396C-A | YOR396C-A | 0 | NA | NA | NA | NA |
| YPL257W-A | YPL257W-A | 0 | NA | NA | NA | NA |
| PAU22 | YPL282C | 0.3924265 | -0.0639558 | 0.11733872 | -0.5450524 | 0.58571745 |
| YPL283W-A | YPL283W-A | 0 | NA | NA | NA | NA |
| YPL283W-B | YPL283W-B | 0 | NA | NA | NA | NA |
| YPR039W | YPR039W | 0.52433194 | 0.00709937 | 0.14164676 | 0.05012027 | 0.96002654 |
| YPR108W-A | YPR108W-A | 0.55696476 | 0.06955784 | 0.1432962 | 0.48541304 | 0.62738338 |
| YPR137C-A | YPR137C-A | 0 | NA | NA | NA | NA |
| YPR158C-C | YPR158C-C | 0 | NA | NA | NA | NA |
| YPR158W-A | YPR158W-A | 0 | NA | NA | NA | NA |
| YPR160W-A | YPR160W-A | 0 | NA | NA | NA | NA |
| YPR169W-A | YPR169W-A | 0 | NA | NA | NA | NA |

| | | | | | | |
|---|---|---|---|---|---|---|
| YPR170W-A | YPR170W-A | 0 | NA | NA | NA | NA |
| YPR204C-A | YPR204C-A | 0 | NA | NA | NA | NA |

| p-value adjusted |
|---|
| 0 |
| 2.92E-190 |
| 4.69E-152 |
| 6.85E-141 |
| 4.29E-140 |
| 7.24E-124 |
| 1.31E-119 |
| 1.34E-112 |
| 1.78E-105 |
| 6.53E-105 |
| 6.17E-101 |
| 5.74E-100 |
| 1.75E-99 |
| 3.45E-99 |
| 5.15E-94 |
| 9.47E-94 |
| 5.42E-91 |
| 5.70E-89 |
| 4.89E-88 |
| 1.39E-84 |
| 1.48E-84 |
| 1.02E-82 |
| 6.40E-82 |
| 4.90E-81 |
| 1.41E-79 |
| 2.10E-78 |
| 6.34E-78 |
| 9.37E-78 |
| 1.33E-77 |
| 2.54E-76 |
| 1.25E-75 |
| 8.35E-75 |
| 1.25E-74 |
| 1.48E-74 |
| 2.74E-73 |
| 1.24E-70 |
| 8.71E-70 |
| 1.24E-69 |
| 1.92E-69 |
| 1.27E-68 |

| |
|---|
| 1.47E-68 |
| 1.46E-66 |
| 1.95E-66 |
| 5.05E-65 |
| 8.94E-65 |
| 1.59E-64 |
| 1.24E-63 |
| 1.66E-62 |
| 3.56E-62 |
| 1.12E-61 |
| 2.22E-61 |
| 3.38E-61 |
| 7.96E-61 |
| 1.78E-60 |
| 4.11E-60 |
| 1.51E-59 |
| 2.72E-59 |
| 2.81E-59 |
| 6.71E-59 |
| 1.94E-58 |
| 2.90E-58 |
| 2.90E-58 |
| 3.08E-58 |
| 7.27E-58 |
| 1.31E-57 |
| 1.43E-57 |
| 1.73E-57 |
| 5.20E-57 |
| 1.56E-56 |
| 8.89E-56 |
| 2.37E-55 |
| 2.66E-55 |
| 9.95E-55 |
| 5.98E-54 |
| 7.85E-54 |
| 8.20E-54 |
| 1.72E-53 |
| 3.13E-53 |
| 5.20E-53 |
| 1.43E-52 |
| 1.27E-51 |

9.04E-51
1.01E-50
3.52E-50
4.33E-50
8.39E-50
9.59E-50
1.07E-49
1.52E-49
1.65E-49
5.79E-49
6.94E-49
1.43E-48
3.00E-48
4.23E-48
4.61E-48
4.69E-48
6.00E-48
8.51E-48
1.29E-47
2.03E-47
2.60E-47
6.31E-47
1.34E-46
1.54E-46
2.99E-46
3.07E-46
3.36E-46
4.35E-46
4.91E-46
9.85E-46
1.20E-45
1.93E-45
2.53E-45
2.63E-45
5.62E-45
1.16E-44
3.09E-44
3.45E-44
3.78E-44
4.15E-44
4.34E-44

| |
|---|
| 1.17E-43 |
| 1.21E-43 |
| 1.39E-43 |
| 2.22E-43 |
| 3.19E-43 |
| 3.74E-43 |
| 4.55E-43 |
| 6.17E-43 |
| 8.37E-43 |
| 1.20E-42 |
| 1.47E-42 |
| 2.51E-42 |
| 4.29E-42 |
| 4.31E-42 |
| 8.16E-42 |
| 2.01E-41 |
| 3.51E-41 |
| 4.75E-41 |
| 6.80E-41 |
| 6.93E-41 |
| 1.00E-40 |
| 1.23E-40 |
| 1.30E-40 |
| 1.81E-40 |
| 1.83E-40 |
| 1.85E-40 |
| 2.46E-40 |
| 3.11E-40 |
| 3.52E-40 |
| 4.34E-40 |
| 8.07E-40 |
| 1.43E-39 |
| 2.46E-39 |
| 3.37E-39 |
| 7.71E-39 |
| 8.96E-39 |
| 1.13E-38 |
| 1.35E-38 |
| 1.59E-38 |
| 4.12E-38 |
| 4.22E-38 |

4.24E-38
4.52E-38
5.76E-38
5.78E-38
6.19E-38
7.00E-38
7.86E-38
8.86E-38
8.90E-38
1.01E-37
1.07E-37
1.17E-37
1.73E-37
1.74E-37
2.04E-37
2.31E-37
2.70E-37
2.97E-37
3.19E-37
3.21E-37
6.56E-37
8.82E-37
1.06E-36
1.24E-36
1.38E-36
1.86E-36
2.59E-36
6.18E-36
7.83E-36
9.81E-36
9.81E-36
1.07E-35
1.41E-35
1.47E-35
2.19E-35
3.24E-35
4.68E-35
5.64E-35
6.93E-35
6.95E-35
8.44E-35

9.79E-35
1.02E-34
1.05E-34
1.87E-34
1.89E-34
1.91E-34
2.24E-34
2.29E-34
2.33E-34
2.95E-34
3.18E-34
3.36E-34
3.77E-34
4.29E-34
4.55E-34
7.79E-34
8.95E-34
1.11E-33
1.12E-33
1.36E-33
1.45E-33
1.50E-33
1.63E-33
1.70E-33
2.23E-33
2.48E-33
2.83E-33
4.89E-33
5.17E-33
9.60E-33
1.72E-32
1.75E-32
1.92E-32
2.29E-32
2.35E-32
3.18E-32
3.64E-32
4.47E-32
5.19E-32
5.37E-32
5.53E-32

6.00E-32
7.85E-32
9.87E-32
9.89E-32
1.21E-31
1.26E-31
1.33E-31
1.53E-31
1.99E-31
3.42E-31
5.05E-31
5.05E-31
6.48E-31
7.25E-31
8.89E-31
1.06E-30
1.07E-30
1.15E-30
1.28E-30
1.28E-30
1.63E-30
2.17E-30
2.84E-30
3.01E-30
3.43E-30
3.93E-30
3.94E-30
5.11E-30
5.21E-30
5.68E-30
6.31E-30
7.37E-30
1.36E-29
1.55E-29
1.60E-29
1.65E-29
2.39E-29
3.43E-29
3.74E-29
4.56E-29
5.59E-29

5.84E-29
6.05E-29
6.19E-29
7.25E-29
9.93E-29
1.13E-28
1.19E-28
3.22E-28
3.97E-28
4.59E-28
5.29E-28
5.35E-28
5.58E-28
8.36E-28
8.36E-28
9.31E-28
9.33E-28
1.09E-27
1.23E-27
1.33E-27
1.44E-27
1.56E-27
1.56E-27
1.59E-27
1.89E-27
1.92E-27
2.05E-27
2.19E-27
2.46E-27
2.46E-27
3.29E-27
3.55E-27
3.60E-27
4.46E-27
4.49E-27
4.76E-27
4.76E-27
5.07E-27
5.94E-27
6.35E-27
7.46E-27

7.55E-27
7.93E-27
8.66E-27
8.66E-27
9.87E-27
1.00E-26
1.19E-26
1.23E-26
1.39E-26
1.79E-26
1.81E-26
2.06E-26
2.58E-26
2.65E-26
3.44E-26
3.47E-26
3.69E-26
3.79E-26
3.95E-26
5.75E-26
6.86E-26
7.72E-26
7.78E-26
1.02E-25
1.13E-25
1.13E-25
1.18E-25
1.41E-25
1.41E-25
1.52E-25
1.65E-25
1.67E-25
1.67E-25
1.80E-25
2.33E-25
2.36E-25
2.63E-25
3.70E-25
3.75E-25
3.95E-25
4.19E-25

4.24E-25
4.55E-25
5.30E-25
5.67E-25
5.77E-25
9.49E-25
1.03E-24
1.03E-24
1.20E-24
1.24E-24
1.26E-24
1.40E-24
1.48E-24
1.51E-24
1.52E-24
1.53E-24
1.67E-24
1.76E-24
1.78E-24
1.98E-24
2.04E-24
2.04E-24
2.10E-24
2.11E-24
2.22E-24
2.39E-24
2.52E-24
2.54E-24
2.71E-24
2.77E-24
3.81E-24
3.94E-24
3.97E-24
5.78E-24
6.07E-24
6.27E-24
6.95E-24
6.99E-24
7.92E-24
9.09E-24
9.90E-24

| |
|---|
| 1.13E-23 |
| 1.17E-23 |
| 1.26E-23 |
| 1.53E-23 |
| 1.56E-23 |
| 1.64E-23 |
| 2.05E-23 |
| 2.11E-23 |
| 2.34E-23 |
| 2.40E-23 |
| 2.40E-23 |
| 2.42E-23 |
| 2.67E-23 |
| 2.88E-23 |
| 2.97E-23 |
| 3.17E-23 |
| 3.28E-23 |
| 3.82E-23 |
| 3.91E-23 |
| 4.19E-23 |
| 4.26E-23 |
| 4.33E-23 |
| 4.34E-23 |
| 4.57E-23 |
| 4.66E-23 |
| 4.75E-23 |
| 5.60E-23 |
| 5.93E-23 |
| 6.68E-23 |
| 7.55E-23 |
| 7.61E-23 |
| 7.80E-23 |
| 8.98E-23 |
| 9.02E-23 |
| 1.05E-22 |
| 1.08E-22 |
| 1.17E-22 |
| 1.32E-22 |
| 1.39E-22 |
| 1.72E-22 |
| 1.79E-22 |

| |
|---|
| 1.87E-22 |
| 1.90E-22 |
| 2.09E-22 |
| 2.59E-22 |
| 2.88E-22 |
| 3.10E-22 |
| 3.27E-22 |
| 3.34E-22 |
| 3.61E-22 |
| 3.65E-22 |
| 4.18E-22 |
| 4.21E-22 |
| 4.26E-22 |
| 5.04E-22 |
| 9.37E-22 |
| 1.22E-21 |
| 1.32E-21 |
| 1.32E-21 |
| 1.66E-21 |
| 1.97E-21 |
| 2.21E-21 |
| 2.27E-21 |
| 3.88E-21 |
| 5.36E-21 |
| 5.36E-21 |
| 5.36E-21 |
| 6.99E-21 |
| 7.38E-21 |
| 8.70E-21 |
| 8.81E-21 |
| 9.15E-21 |
| 9.35E-21 |
| 9.55E-21 |
| 9.90E-21 |
| 1.02E-20 |
| 1.04E-20 |
| 1.12E-20 |
| 1.22E-20 |
| 1.22E-20 |
| 1.34E-20 |
| 1.36E-20 |

| |
|---|
| 1.53E-20 |
| 1.57E-20 |
| 1.64E-20 |
| 1.68E-20 |
| 1.76E-20 |
| 1.89E-20 |
| 2.23E-20 |
| 2.51E-20 |
| 2.51E-20 |
| 2.85E-20 |
| 2.88E-20 |
| 2.88E-20 |
| 3.28E-20 |
| 3.51E-20 |
| 3.92E-20 |
| 4.05E-20 |
| 4.76E-20 |
| 5.10E-20 |
| 5.17E-20 |
| 5.21E-20 |
| 5.29E-20 |
| 5.59E-20 |
| 6.06E-20 |
| 6.06E-20 |
| 6.42E-20 |
| 8.31E-20 |
| 8.32E-20 |
| 8.77E-20 |
| 1.06E-19 |
| 1.16E-19 |
| 1.17E-19 |
| 1.17E-19 |
| 1.20E-19 |
| 1.37E-19 |
| 1.46E-19 |
| 1.85E-19 |
| 1.96E-19 |
| 2.19E-19 |
| 2.20E-19 |
| 2.47E-19 |
| 2.53E-19 |

|  |
| --- |
| 2.59E-19 |
| 2.59E-19 |
| 2.60E-19 |
| 2.86E-19 |
| 3.02E-19 |
| 3.04E-19 |
| 3.23E-19 |
| 3.31E-19 |
| 3.77E-19 |
| 4.62E-19 |
| 4.94E-19 |
| 5.98E-19 |
| 6.27E-19 |
| 6.40E-19 |
| 6.69E-19 |
| 6.87E-19 |
| 7.39E-19 |
| 8.15E-19 |
| 1.04E-18 |
| 1.14E-18 |
| 1.18E-18 |
| 1.23E-18 |
| 1.31E-18 |
| 1.49E-18 |
| 1.73E-18 |
| 1.73E-18 |
| 1.91E-18 |
| 1.92E-18 |
| 2.34E-18 |
| 2.52E-18 |
| 2.55E-18 |
| 2.82E-18 |
| 3.29E-18 |
| 4.11E-18 |
| 4.25E-18 |
| 4.42E-18 |
| 4.91E-18 |
| 5.18E-18 |
| 5.27E-18 |
| 5.40E-18 |
| 5.52E-18 |

| |
|---|
| 5.62E-18 |
| 5.98E-18 |
| 6.03E-18 |
| 6.12E-18 |
| 7.86E-18 |
| 8.52E-18 |
| 8.83E-18 |
| 8.93E-18 |
| 9.37E-18 |
| 1.02E-17 |
| 1.07E-17 |
| 1.09E-17 |
| 1.13E-17 |
| 1.15E-17 |
| 1.22E-17 |
| 1.34E-17 |
| 1.34E-17 |
| 1.48E-17 |
| 1.63E-17 |
| 1.84E-17 |
| 1.96E-17 |
| 2.19E-17 |
| 2.41E-17 |
| 2.43E-17 |
| 2.66E-17 |
| 3.20E-17 |
| 3.29E-17 |
| 3.36E-17 |
| 3.37E-17 |
| 3.50E-17 |
| 3.64E-17 |
| 3.67E-17 |
| 3.70E-17 |
| 3.81E-17 |
| 4.03E-17 |
| 4.23E-17 |
| 5.35E-17 |
| 5.98E-17 |
| 6.02E-17 |
| 8.80E-17 |
| 9.31E-17 |

9.71E-17
9.95E-17
1.02E-16
1.06E-16
1.11E-16
1.17E-16
1.24E-16
1.31E-16
1.32E-16
1.41E-16
1.42E-16
1.46E-16
1.52E-16
1.83E-16
1.87E-16
1.97E-16
2.01E-16
2.06E-16
2.06E-16
2.08E-16
2.08E-16
2.13E-16
2.15E-16
2.78E-16
2.79E-16
2.80E-16
2.81E-16
2.82E-16
3.63E-16
3.71E-16
3.88E-16
4.10E-16
4.57E-16
4.71E-16
4.77E-16
4.88E-16
4.92E-16
5.43E-16
5.55E-16
5.62E-16
6.77E-16

7.06E-16
7.12E-16
7.78E-16
8.13E-16
8.20E-16
8.94E-16
9.56E-16
9.56E-16
9.60E-16
1.02E-15
1.03E-15
1.15E-15
1.18E-15
1.19E-15
1.23E-15
1.29E-15
1.30E-15
1.33E-15
1.40E-15
1.47E-15
1.48E-15
1.64E-15
1.72E-15
1.73E-15
1.91E-15
1.94E-15
2.00E-15
2.06E-15
2.16E-15
2.47E-15
2.60E-15
2.74E-15
2.76E-15
2.81E-15
2.84E-15
2.90E-15
2.96E-15
3.05E-15
3.07E-15
3.20E-15
3.28E-15

3.43E-15
3.60E-15
3.74E-15
3.86E-15
3.99E-15
4.03E-15
4.49E-15
4.96E-15
4.99E-15
5.00E-15
5.02E-15
5.07E-15
5.10E-15
5.64E-15
5.78E-15
5.98E-15
6.12E-15
6.27E-15
6.40E-15
6.42E-15
6.62E-15
6.81E-15
6.84E-15
6.88E-15
7.31E-15
7.40E-15
7.78E-15
8.08E-15
9.01E-15
9.08E-15
1.06E-14
1.09E-14
1.22E-14
1.31E-14
1.32E-14
1.34E-14
1.47E-14
1.49E-14
1.51E-14
1.61E-14
1.70E-14

| |
|---|
| 1.76E-14 |
| 1.84E-14 |
| 1.94E-14 |
| 2.04E-14 |
| 2.15E-14 |
| 2.16E-14 |
| 2.24E-14 |
| 2.31E-14 |
| 2.45E-14 |
| 2.52E-14 |
| 2.60E-14 |
| 2.62E-14 |
| 3.10E-14 |
| 3.12E-14 |
| 3.14E-14 |
| 3.16E-14 |
| 3.22E-14 |
| 3.34E-14 |
| 3.40E-14 |
| 3.43E-14 |
| 3.68E-14 |
| 3.68E-14 |
| 4.25E-14 |
| 4.86E-14 |
| 5.59E-14 |
| 6.57E-14 |
| 6.65E-14 |
| 6.65E-14 |
| 6.90E-14 |
| 6.96E-14 |
| 7.00E-14 |
| 7.03E-14 |
| 7.13E-14 |
| 7.16E-14 |
| 7.29E-14 |
| 7.50E-14 |
| 7.65E-14 |
| 7.65E-14 |
| 7.77E-14 |
| 8.09E-14 |
| 8.56E-14 |

8.59E-14
8.64E-14
8.79E-14
8.97E-14
9.24E-14
9.41E-14
1.03E-13
1.05E-13
1.07E-13
1.20E-13
1.21E-13
1.28E-13
1.28E-13
1.32E-13
1.39E-13
1.39E-13
1.40E-13
1.41E-13
1.58E-13
1.62E-13
1.64E-13
1.71E-13
1.74E-13
1.83E-13
1.83E-13
1.90E-13
1.92E-13
1.94E-13
2.04E-13
2.11E-13
2.12E-13
2.12E-13
2.15E-13
2.22E-13
2.28E-13
2.30E-13
2.54E-13
2.55E-13
2.56E-13
2.57E-13
2.62E-13

| |
|---|
| 2.74E-13 |
| 2.83E-13 |
| 2.83E-13 |
| 2.84E-13 |
| 2.94E-13 |
| 3.07E-13 |
| 3.08E-13 |
| 3.41E-13 |
| 3.52E-13 |
| 3.61E-13 |
| 3.65E-13 |
| 3.73E-13 |
| 3.73E-13 |
| 3.82E-13 |
| 4.13E-13 |
| 4.20E-13 |
| 4.36E-13 |
| 4.39E-13 |
| 5.24E-13 |
| 5.34E-13 |
| 5.51E-13 |
| 5.55E-13 |
| 5.82E-13 |
| 5.88E-13 |
| 6.08E-13 |
| 6.18E-13 |
| 6.34E-13 |
| 6.42E-13 |
| 6.49E-13 |
| 6.53E-13 |
| 6.60E-13 |
| 7.23E-13 |
| 7.30E-13 |
| 7.30E-13 |
| 7.45E-13 |
| 8.43E-13 |
| 8.82E-13 |
| 9.89E-13 |
| 9.90E-13 |
| 1.02E-12 |
| 1.09E-12 |

1.11E-12
1.12E-12
1.13E-12
1.18E-12
1.21E-12
1.23E-12
1.25E-12
1.25E-12
1.25E-12
1.30E-12
1.32E-12
1.35E-12
1.39E-12
1.41E-12
1.41E-12
1.56E-12
1.59E-12
1.67E-12
1.70E-12
1.71E-12
1.71E-12
1.75E-12
1.75E-12
1.77E-12
2.02E-12
2.04E-12
2.04E-12
2.04E-12
2.04E-12
2.15E-12
2.28E-12
2.30E-12
2.30E-12
2.44E-12
2.62E-12
2.70E-12
2.81E-12
2.85E-12
3.19E-12
3.30E-12
3.37E-12

3.48E-12
3.50E-12
3.50E-12
3.52E-12
3.52E-12
3.55E-12
3.58E-12
3.69E-12
3.69E-12
3.71E-12
4.13E-12
4.31E-12
4.36E-12
4.47E-12
4.53E-12
4.88E-12
4.99E-12
5.09E-12
5.13E-12
5.13E-12
5.18E-12
5.35E-12
5.39E-12
5.44E-12
5.46E-12
5.46E-12
5.69E-12
5.71E-12
6.01E-12
6.09E-12
6.60E-12
6.80E-12
6.88E-12
7.31E-12
7.74E-12
8.05E-12
8.24E-12
8.29E-12
8.40E-12
8.68E-12
8.92E-12

| |
|---|
| 9.04E-12 |
| 9.41E-12 |
| 9.61E-12 |
| 1.02E-11 |
| 1.03E-11 |
| 1.04E-11 |
| 1.09E-11 |
| 1.09E-11 |
| 1.09E-11 |
| 1.09E-11 |
| 1.11E-11 |
| 1.12E-11 |
| 1.13E-11 |
| 1.13E-11 |
| 1.19E-11 |
| 1.21E-11 |
| 1.24E-11 |
| 1.28E-11 |
| 1.30E-11 |
| 1.38E-11 |
| 1.41E-11 |
| 1.41E-11 |
| 1.41E-11 |
| 1.43E-11 |
| 1.44E-11 |
| 1.46E-11 |
| 1.47E-11 |
| 1.48E-11 |
| 1.51E-11 |
| 1.53E-11 |
| 1.56E-11 |
| 1.75E-11 |
| 1.92E-11 |
| 1.93E-11 |
| 1.96E-11 |
| 1.97E-11 |
| 2.14E-11 |
| 2.14E-11 |
| 2.21E-11 |
| 2.23E-11 |
| 2.23E-11 |

| |
|---|
| 2.50E-11 |
| 2.64E-11 |
| 2.66E-11 |
| 2.89E-11 |
| 2.89E-11 |
| 3.01E-11 |
| 3.15E-11 |
| 3.46E-11 |
| 3.60E-11 |
| 3.73E-11 |
| 3.81E-11 |
| 3.83E-11 |
| 3.86E-11 |
| 3.97E-11 |
| 4.03E-11 |
| 4.18E-11 |
| 4.25E-11 |
| 4.27E-11 |
| 4.29E-11 |
| 4.34E-11 |
| 4.48E-11 |
| 4.53E-11 |
| 4.64E-11 |
| 4.85E-11 |
| 4.90E-11 |
| 5.00E-11 |
| 5.17E-11 |
| 5.19E-11 |
| 5.41E-11 |
| 5.46E-11 |
| 5.47E-11 |
| 5.68E-11 |
| 5.79E-11 |
| 5.93E-11 |
| 5.96E-11 |
| 6.26E-11 |
| 6.46E-11 |
| 6.71E-11 |
| 6.82E-11 |
| 6.82E-11 |
| 6.83E-11 |

6.84E-11
7.30E-11
7.48E-11
7.54E-11
8.52E-11
8.52E-11
8.91E-11
8.96E-11
9.13E-11
1.00E-10
1.03E-10
1.04E-10
1.04E-10
1.04E-10
1.06E-10
1.08E-10
1.13E-10
1.14E-10
1.20E-10
1.20E-10
1.21E-10
1.29E-10
1.29E-10
1.33E-10
1.36E-10
1.38E-10
1.41E-10
1.45E-10
1.51E-10
1.55E-10
1.55E-10
1.55E-10
1.65E-10
1.66E-10
1.67E-10
1.72E-10
1.75E-10
1.83E-10
1.83E-10
1.85E-10
1.88E-10

| |
|---|
| 1.93E-10 |
| 2.01E-10 |
| 2.02E-10 |
| 2.25E-10 |
| 2.29E-10 |
| 2.29E-10 |
| 2.32E-10 |
| 2.33E-10 |
| 2.40E-10 |
| 2.40E-10 |
| 2.40E-10 |
| 2.40E-10 |
| 2.41E-10 |
| 2.45E-10 |
| 2.55E-10 |
| 2.56E-10 |
| 2.56E-10 |
| 2.57E-10 |
| 2.60E-10 |
| 2.73E-10 |
| 2.84E-10 |
| 3.38E-10 |
| 3.61E-10 |
| 3.70E-10 |
| 3.90E-10 |
| 3.92E-10 |
| 4.02E-10 |
| 4.07E-10 |
| 4.09E-10 |
| 4.31E-10 |
| 4.61E-10 |
| 4.62E-10 |
| 4.74E-10 |
| 4.78E-10 |
| 4.83E-10 |
| 4.84E-10 |
| 4.86E-10 |
| 4.99E-10 |
| 5.38E-10 |
| 5.41E-10 |
| 5.46E-10 |

| |
|---|
| 5.52E-10 |
| 6.07E-10 |
| 6.17E-10 |
| 6.19E-10 |
| 6.25E-10 |
| 6.33E-10 |
| 6.40E-10 |
| 6.64E-10 |
| 6.90E-10 |
| 6.91E-10 |
| 7.06E-10 |
| 7.46E-10 |
| 7.51E-10 |
| 7.55E-10 |
| 7.60E-10 |
| 7.98E-10 |
| 8.30E-10 |
| 8.37E-10 |
| 8.62E-10 |
| 8.64E-10 |
| 8.82E-10 |
| 8.90E-10 |
| 9.01E-10 |
| 9.10E-10 |
| 9.15E-10 |
| 9.32E-10 |
| 9.41E-10 |
| 9.42E-10 |
| 9.54E-10 |
| 1.01E-09 |
| 1.01E-09 |
| 1.03E-09 |
| 1.03E-09 |
| 1.05E-09 |
| 1.07E-09 |
| 1.08E-09 |
| 1.09E-09 |
| 1.10E-09 |
| 1.12E-09 |
| 1.14E-09 |
| 1.29E-09 |

1.30E-09
1.30E-09
1.31E-09
1.36E-09
1.39E-09
1.39E-09
1.39E-09
1.41E-09
1.41E-09
1.41E-09
1.44E-09
1.49E-09
1.51E-09
1.53E-09
1.55E-09
1.55E-09
1.55E-09
1.58E-09
1.61E-09
1.66E-09
1.68E-09
1.70E-09
1.72E-09
1.77E-09
1.77E-09
1.84E-09
1.90E-09
1.92E-09
1.94E-09
1.96E-09
1.98E-09
2.03E-09
2.06E-09
2.08E-09
2.09E-09
2.20E-09
2.20E-09
2.21E-09
2.22E-09
2.23E-09
2.27E-09

| |
|---|
| 2.28E-09 |
| 2.36E-09 |
| 2.37E-09 |
| 2.37E-09 |
| 2.38E-09 |
| 2.43E-09 |
| 2.50E-09 |
| 2.53E-09 |
| 2.55E-09 |
| 2.58E-09 |
| 2.59E-09 |
| 2.60E-09 |
| 2.61E-09 |
| 2.62E-09 |
| 2.86E-09 |
| 2.93E-09 |
| 3.05E-09 |
| 3.05E-09 |
| 3.14E-09 |
| 3.18E-09 |
| 3.21E-09 |
| 3.25E-09 |
| 3.40E-09 |
| 3.41E-09 |
| 3.42E-09 |
| 3.46E-09 |
| 3.52E-09 |
| 3.76E-09 |
| 3.78E-09 |
| 3.82E-09 |
| 3.96E-09 |
| 4.04E-09 |
| 4.17E-09 |
| 4.27E-09 |
| 4.29E-09 |
| 4.35E-09 |
| 4.47E-09 |
| 4.54E-09 |
| 4.69E-09 |
| 4.71E-09 |
| 4.81E-09 |

4.81E-09
4.99E-09
5.04E-09
5.17E-09
5.21E-09
5.37E-09
5.46E-09
5.49E-09
5.75E-09
5.76E-09
5.84E-09
5.90E-09
5.97E-09
6.03E-09
6.15E-09
6.20E-09
6.31E-09
6.33E-09
6.41E-09
6.42E-09
6.42E-09
6.46E-09
6.51E-09
6.59E-09
6.60E-09
6.60E-09
6.81E-09
6.86E-09
7.26E-09
7.39E-09
7.67E-09
7.74E-09
7.86E-09
8.05E-09
8.06E-09
8.85E-09
8.89E-09
8.90E-09
9.05E-09
9.15E-09
9.32E-09

9.33E-09
9.41E-09
9.65E-09
9.65E-09
9.86E-09
9.94E-09
1.01E-08
1.02E-08
1.05E-08
1.06E-08
1.06E-08
1.07E-08
1.09E-08
1.10E-08
1.10E-08
1.14E-08
1.15E-08
1.18E-08
1.23E-08
1.24E-08
1.26E-08
1.28E-08
1.28E-08
1.32E-08
1.36E-08
1.37E-08
1.38E-08
1.43E-08
1.45E-08
1.47E-08
1.53E-08
1.53E-08
1.62E-08
1.66E-08
1.68E-08
1.68E-08
1.71E-08
1.72E-08
1.74E-08
1.75E-08
1.76E-08

1.76E-08
1.81E-08
1.95E-08
1.96E-08
1.97E-08
2.01E-08
2.02E-08
2.15E-08
2.30E-08
2.36E-08
2.37E-08
2.38E-08
2.38E-08
2.39E-08
2.40E-08
2.43E-08
2.43E-08
2.49E-08
2.49E-08
2.53E-08
2.53E-08
2.54E-08
2.57E-08
2.63E-08
2.66E-08
2.76E-08
2.78E-08
2.78E-08
2.80E-08
2.80E-08
2.82E-08
2.94E-08
2.99E-08
3.01E-08
3.07E-08
3.17E-08
3.18E-08
3.25E-08
3.33E-08
3.40E-08
3.42E-08

|  |
|---|
| 3.47E-08 |
| 3.58E-08 |
| 3.63E-08 |
| 3.87E-08 |
| 3.97E-08 |
| 4.01E-08 |
| 4.05E-08 |
| 4.15E-08 |
| 4.20E-08 |
| 4.20E-08 |
| 4.24E-08 |
| 4.24E-08 |
| 4.27E-08 |
| 4.30E-08 |
| 4.31E-08 |
| 4.34E-08 |
| 4.35E-08 |
| 4.37E-08 |
| 4.47E-08 |
| 4.50E-08 |
| 4.57E-08 |
| 4.78E-08 |
| 4.84E-08 |
| 5.02E-08 |
| 5.04E-08 |
| 5.05E-08 |
| 5.29E-08 |
| 5.32E-08 |
| 5.37E-08 |
| 5.39E-08 |
| 5.49E-08 |
| 5.56E-08 |
| 5.58E-08 |
| 5.64E-08 |
| 5.74E-08 |
| 5.77E-08 |
| 5.78E-08 |
| 5.83E-08 |
| 5.90E-08 |
| 5.93E-08 |
| 6.02E-08 |

6.02E-08
6.09E-08
6.17E-08
6.17E-08
6.30E-08
6.40E-08
6.55E-08
6.60E-08
6.71E-08
6.84E-08
6.88E-08
6.90E-08
7.05E-08
7.13E-08
7.26E-08
7.41E-08
7.42E-08
7.61E-08
7.70E-08
7.93E-08
7.98E-08
7.98E-08
7.99E-08
8.01E-08
8.17E-08
8.23E-08
8.40E-08
8.42E-08
8.47E-08
8.47E-08
8.48E-08
8.79E-08
8.97E-08
9.54E-08
9.61E-08
9.63E-08
9.84E-08
9.86E-08
1.01E-07
1.02E-07
1.04E-07

| |
|---|
| 1.04E-07 |
| 1.06E-07 |
| 1.06E-07 |
| 1.07E-07 |
| 1.10E-07 |
| 1.11E-07 |
| 1.11E-07 |
| 1.11E-07 |
| 1.12E-07 |
| 1.15E-07 |
| 1.16E-07 |
| 1.16E-07 |
| 1.18E-07 |
| 1.26E-07 |
| 1.29E-07 |
| 1.29E-07 |
| 1.36E-07 |
| 1.37E-07 |
| 1.38E-07 |
| 1.39E-07 |
| 1.40E-07 |
| 1.40E-07 |
| 1.41E-07 |
| 1.42E-07 |
| 1.42E-07 |
| 1.44E-07 |
| 1.46E-07 |
| 1.46E-07 |
| 1.48E-07 |
| 1.59E-07 |
| 1.62E-07 |
| 1.63E-07 |
| 1.64E-07 |
| 1.64E-07 |
| 1.71E-07 |
| 1.73E-07 |
| 1.73E-07 |
| 1.73E-07 |
| 1.75E-07 |
| 1.75E-07 |
| 1.76E-07 |

1.79E-07
1.79E-07
1.80E-07
1.83E-07
1.93E-07
2.00E-07
2.05E-07
2.05E-07
2.07E-07
2.10E-07
2.10E-07
2.18E-07
2.23E-07
2.23E-07
2.23E-07
2.24E-07
2.26E-07
2.29E-07
2.33E-07
2.33E-07
2.37E-07
2.42E-07
2.42E-07
2.45E-07
2.46E-07
2.48E-07
2.50E-07
2.51E-07
2.54E-07
2.54E-07
2.56E-07
2.56E-07
2.59E-07
2.65E-07
2.77E-07
2.81E-07
2.81E-07
2.85E-07
2.90E-07
2.92E-07
2.95E-07

| |
|---|
| 2.97E-07 |
| 2.99E-07 |
| 3.04E-07 |
| 3.10E-07 |
| 3.12E-07 |
| 3.19E-07 |
| 3.20E-07 |
| 3.29E-07 |
| 3.31E-07 |
| 3.36E-07 |
| 3.39E-07 |
| 3.42E-07 |
| 3.42E-07 |
| 3.48E-07 |
| 3.48E-07 |
| 3.48E-07 |
| 3.50E-07 |
| 3.50E-07 |
| 3.52E-07 |
| 3.60E-07 |
| 3.68E-07 |
| 3.68E-07 |
| 3.69E-07 |
| 3.73E-07 |
| 3.74E-07 |
| 3.75E-07 |
| 3.82E-07 |
| 3.82E-07 |
| 3.84E-07 |
| 3.87E-07 |
| 3.89E-07 |
| 3.92E-07 |
| 3.93E-07 |
| 3.93E-07 |
| 3.96E-07 |
| 3.96E-07 |
| 3.96E-07 |
| 3.97E-07 |
| 3.97E-07 |
| 4.01E-07 |
| 4.01E-07 |

4.04E-07
4.04E-07
4.04E-07
4.06E-07
4.15E-07
4.26E-07
4.27E-07
4.28E-07
4.38E-07
4.42E-07
4.46E-07
4.47E-07
4.50E-07
4.66E-07
4.72E-07
4.72E-07
4.72E-07
4.75E-07
4.81E-07
4.84E-07
5.04E-07
5.07E-07
5.09E-07
5.13E-07
5.17E-07
5.17E-07
5.17E-07
5.18E-07
5.23E-07
5.47E-07
5.49E-07
5.51E-07
5.54E-07
5.55E-07
5.56E-07
5.60E-07
5.60E-07
5.69E-07
5.82E-07
5.83E-07
5.91E-07

| |
|---|
| 5.94E-07 |
| 6.02E-07 |
| 6.09E-07 |
| 6.13E-07 |
| 6.14E-07 |
| 6.16E-07 |
| 6.20E-07 |
| 6.39E-07 |
| 6.55E-07 |
| 6.56E-07 |
| 6.67E-07 |
| 6.89E-07 |
| 6.90E-07 |
| 6.91E-07 |
| 6.94E-07 |
| 6.96E-07 |
| 7.03E-07 |
| 7.04E-07 |
| 7.05E-07 |
| 7.09E-07 |
| 7.13E-07 |
| 7.24E-07 |
| 7.26E-07 |
| 7.48E-07 |
| 7.48E-07 |
| 7.48E-07 |
| 7.67E-07 |
| 7.69E-07 |
| 7.70E-07 |
| 7.72E-07 |
| 7.87E-07 |
| 8.10E-07 |
| 8.37E-07 |
| 8.46E-07 |
| 8.63E-07 |
| 8.70E-07 |
| 8.78E-07 |
| 8.81E-07 |
| 8.82E-07 |
| 9.02E-07 |
| 9.08E-07 |

| |
|---|
| 9.12E-07 |
| 9.24E-07 |
| 9.24E-07 |
| 9.34E-07 |
| 9.47E-07 |
| 9.50E-07 |
| 9.62E-07 |
| 9.91E-07 |
| 9.93E-07 |
| 9.99E-07 |
| 1.00E-06 |
| 1.00E-06 |
| 1.02E-06 |
| 1.06E-06 |
| 1.06E-06 |
| 1.06E-06 |
| 1.06E-06 |
| 1.06E-06 |
| 1.07E-06 |
| 1.08E-06 |
| 1.09E-06 |
| 1.09E-06 |
| 1.11E-06 |
| 1.12E-06 |
| 1.13E-06 |
| 1.13E-06 |
| 1.15E-06 |
| 1.15E-06 |
| 1.18E-06 |
| 1.22E-06 |
| 1.23E-06 |
| 1.23E-06 |
| 1.24E-06 |
| 1.26E-06 |
| 1.29E-06 |
| 1.30E-06 |
| 1.30E-06 |
| 1.31E-06 |
| 1.32E-06 |
| 1.34E-06 |
| 1.36E-06 |

| |
|---|
| 1.37E-06 |
| 1.38E-06 |
| 1.43E-06 |
| 1.44E-06 |
| 1.44E-06 |
| 1.45E-06 |
| 1.50E-06 |
| 1.50E-06 |
| 1.55E-06 |
| 1.56E-06 |
| 1.56E-06 |
| 1.57E-06 |
| 1.58E-06 |
| 1.60E-06 |
| 1.63E-06 |
| 1.63E-06 |
| 1.63E-06 |
| 1.64E-06 |
| 1.66E-06 |
| 1.66E-06 |
| 1.68E-06 |
| 1.71E-06 |
| 1.72E-06 |
| 1.75E-06 |
| 1.75E-06 |
| 1.76E-06 |
| 1.77E-06 |
| 1.78E-06 |
| 1.82E-06 |
| 1.82E-06 |
| 1.83E-06 |
| 1.83E-06 |
| 1.84E-06 |
| 1.84E-06 |
| 1.88E-06 |
| 1.91E-06 |
| 1.93E-06 |
| 1.97E-06 |
| 1.97E-06 |
| 2.00E-06 |
| 2.00E-06 |

2.02E-06
2.04E-06
2.04E-06
2.06E-06
2.06E-06
2.06E-06
2.09E-06
2.14E-06
2.16E-06
2.19E-06
2.19E-06
2.27E-06
2.34E-06
2.37E-06
2.39E-06
2.41E-06
2.42E-06
2.42E-06
2.42E-06
2.44E-06
2.45E-06
2.48E-06
2.50E-06
2.53E-06
2.58E-06
2.59E-06
2.60E-06
2.60E-06
2.64E-06
2.67E-06
2.67E-06
2.69E-06
2.70E-06
2.71E-06
2.74E-06
2.85E-06
2.85E-06
2.87E-06
2.87E-06
2.88E-06
2.90E-06

| |
|---|
| 2.97E-06 |
| 3.06E-06 |
| 3.08E-06 |
| 3.18E-06 |
| 3.20E-06 |
| 3.22E-06 |
| 3.23E-06 |
| 3.25E-06 |
| 3.31E-06 |
| 3.32E-06 |
| 3.32E-06 |
| 3.34E-06 |
| 3.40E-06 |
| 3.43E-06 |
| 3.51E-06 |
| 3.53E-06 |
| 3.56E-06 |
| 3.61E-06 |
| 3.61E-06 |
| 3.61E-06 |
| 3.64E-06 |
| 3.66E-06 |
| 3.67E-06 |
| 3.72E-06 |
| 3.72E-06 |
| 3.78E-06 |
| 3.79E-06 |
| 3.79E-06 |
| 3.81E-06 |
| 3.83E-06 |
| 3.83E-06 |
| 3.94E-06 |
| 3.94E-06 |
| 4.03E-06 |
| 4.06E-06 |
| 4.12E-06 |
| 4.12E-06 |
| 4.17E-06 |
| 4.18E-06 |
| 4.19E-06 |
| 4.21E-06 |

| |
|---|
| 4.23E-06 |
| 4.25E-06 |
| 4.28E-06 |
| 4.38E-06 |
| 4.39E-06 |
| 4.40E-06 |
| 4.41E-06 |
| 4.47E-06 |
| 4.51E-06 |
| 4.51E-06 |
| 4.56E-06 |
| 4.69E-06 |
| 4.76E-06 |
| 4.84E-06 |
| 4.94E-06 |
| 4.96E-06 |
| 4.97E-06 |
| 5.11E-06 |
| 5.18E-06 |
| 5.33E-06 |
| 5.46E-06 |
| 5.47E-06 |
| 5.49E-06 |
| 5.49E-06 |
| 5.53E-06 |
| 5.79E-06 |
| 5.86E-06 |
| 5.86E-06 |
| 5.88E-06 |
| 5.88E-06 |
| 5.90E-06 |
| 5.95E-06 |
| 5.95E-06 |
| 5.95E-06 |
| 5.97E-06 |
| 5.98E-06 |
| 5.98E-06 |
| 6.08E-06 |
| 6.19E-06 |
| 6.26E-06 |
| 6.29E-06 |

| |
|---|
| 6.30E-06 |
| 6.31E-06 |
| 6.32E-06 |
| 6.32E-06 |
| 6.45E-06 |
| 6.51E-06 |
| 6.57E-06 |
| 6.94E-06 |
| 7.02E-06 |
| 7.03E-06 |
| 7.09E-06 |
| 7.17E-06 |
| 7.34E-06 |
| 7.39E-06 |
| 7.40E-06 |
| 7.45E-06 |
| 7.47E-06 |
| 7.47E-06 |
| 7.47E-06 |
| 7.57E-06 |
| 7.58E-06 |
| 7.64E-06 |
| 7.65E-06 |
| 7.73E-06 |
| 7.80E-06 |
| 7.80E-06 |
| 7.84E-06 |
| 7.84E-06 |
| 7.92E-06 |
| 8.09E-06 |
| 8.15E-06 |
| 8.25E-06 |
| 8.29E-06 |
| 8.31E-06 |
| 8.40E-06 |
| 8.45E-06 |
| 8.55E-06 |
| 8.61E-06 |
| 8.67E-06 |
| 8.67E-06 |
| 8.74E-06 |

8.77E-06
8.79E-06
8.84E-06
8.86E-06
8.88E-06
8.88E-06
8.90E-06
9.09E-06
9.09E-06
9.09E-06
9.25E-06
9.30E-06
9.41E-06
9.55E-06
9.77E-06
9.88E-06
9.89E-06
1.00E-05
1.01E-05
1.03E-05
1.05E-05
1.05E-05
1.05E-05
1.05E-05
1.06E-05
1.08E-05
1.09E-05
1.10E-05
1.10E-05
1.14E-05
1.14E-05
1.15E-05
1.17E-05
1.18E-05
1.18E-05
1.18E-05
1.18E-05
1.18E-05
1.19E-05
1.19E-05
1.19E-05

| |
|---|
| 1.20E-05 |
| 1.20E-05 |
| 1.20E-05 |
| 1.21E-05 |
| 1.24E-05 |
| 1.24E-05 |
| 1.24E-05 |
| 1.26E-05 |
| 1.27E-05 |
| 1.27E-05 |
| 1.27E-05 |
| 1.28E-05 |
| 1.30E-05 |
| 1.31E-05 |
| 1.32E-05 |
| 1.33E-05 |
| 1.33E-05 |
| 1.34E-05 |
| 1.34E-05 |
| 1.34E-05 |
| 1.35E-05 |
| 1.35E-05 |
| 1.38E-05 |
| 1.40E-05 |
| 1.40E-05 |
| 1.40E-05 |
| 1.42E-05 |
| 1.43E-05 |
| 1.44E-05 |
| 1.45E-05 |
| 1.45E-05 |
| 1.45E-05 |
| 1.46E-05 |
| 1.48E-05 |
| 1.48E-05 |
| 1.49E-05 |
| 1.49E-05 |
| 1.50E-05 |
| 1.50E-05 |
| 1.50E-05 |
| 1.54E-05 |

| |
|---|
| 1.54E-05 |
| 1.54E-05 |
| 1.57E-05 |
| 1.58E-05 |
| 1.62E-05 |
| 1.63E-05 |
| 1.64E-05 |
| 1.64E-05 |
| 1.65E-05 |
| 1.68E-05 |
| 1.70E-05 |
| 1.70E-05 |
| 1.72E-05 |
| 1.72E-05 |
| 1.74E-05 |
| 1.74E-05 |
| 1.75E-05 |
| 1.76E-05 |
| 1.79E-05 |
| 1.79E-05 |
| 1.79E-05 |
| 1.79E-05 |
| 1.79E-05 |
| 1.84E-05 |
| 1.85E-05 |
| 1.88E-05 |
| 1.88E-05 |
| 1.88E-05 |
| 1.89E-05 |
| 1.90E-05 |
| 1.90E-05 |
| 1.91E-05 |
| 1.96E-05 |
| 1.96E-05 |
| 1.98E-05 |
| 1.99E-05 |
| 2.02E-05 |
| 2.03E-05 |
| 2.04E-05 |
| 2.05E-05 |
| 2.05E-05 |

| |
|---|
| 2.10E-05 |
| 2.12E-05 |
| 2.16E-05 |
| 2.19E-05 |
| 2.21E-05 |
| 2.21E-05 |
| 2.21E-05 |
| 2.27E-05 |
| 2.27E-05 |
| 2.28E-05 |
| 2.31E-05 |
| 2.32E-05 |
| 2.33E-05 |
| 2.34E-05 |
| 2.37E-05 |
| 2.39E-05 |
| 2.40E-05 |
| 2.41E-05 |
| 2.42E-05 |
| 2.42E-05 |
| 2.43E-05 |
| 2.43E-05 |
| 2.46E-05 |
| 2.52E-05 |
| 2.52E-05 |
| 2.52E-05 |
| 2.54E-05 |
| 2.57E-05 |
| 2.57E-05 |
| 2.57E-05 |
| 2.58E-05 |
| 2.60E-05 |
| 2.66E-05 |
| 2.66E-05 |
| 2.69E-05 |
| 2.69E-05 |
| 2.72E-05 |
| 2.73E-05 |
| 2.73E-05 |
| 2.86E-05 |
| 2.88E-05 |

| |
|---|
| 2.88E-05 |
| 2.90E-05 |
| 2.91E-05 |
| 2.92E-05 |
| 2.93E-05 |
| 2.95E-05 |
| 2.97E-05 |
| 2.97E-05 |
| 2.98E-05 |
| 3.00E-05 |
| 3.03E-05 |
| 3.06E-05 |
| 3.09E-05 |
| 3.09E-05 |
| 3.13E-05 |
| 3.14E-05 |
| 3.15E-05 |
| 3.24E-05 |
| 3.25E-05 |
| 3.34E-05 |
| 3.37E-05 |
| 3.40E-05 |
| 3.41E-05 |
| 3.45E-05 |
| 3.47E-05 |
| 3.47E-05 |
| 3.47E-05 |
| 3.48E-05 |
| 3.49E-05 |
| 3.52E-05 |
| 3.52E-05 |
| 3.57E-05 |
| 3.58E-05 |
| 3.73E-05 |
| 3.75E-05 |
| 3.76E-05 |
| 3.76E-05 |
| 3.77E-05 |
| 3.77E-05 |
| 3.81E-05 |
| 3.84E-05 |

| |
|---|
| 3.88E-05 |
| 3.89E-05 |
| 3.99E-05 |
| 3.99E-05 |
| 3.99E-05 |
| 4.00E-05 |
| 4.00E-05 |
| 4.01E-05 |
| 4.08E-05 |
| 4.08E-05 |
| 4.13E-05 |
| 4.13E-05 |
| 4.14E-05 |
| 4.17E-05 |
| 4.17E-05 |
| 4.18E-05 |
| 4.19E-05 |
| 4.19E-05 |
| 4.24E-05 |
| 4.25E-05 |
| 4.26E-05 |
| 4.26E-05 |
| 4.27E-05 |
| 4.27E-05 |
| 4.28E-05 |
| 4.31E-05 |
| 4.33E-05 |
| 4.34E-05 |
| 4.34E-05 |
| 4.39E-05 |
| 4.39E-05 |
| 4.40E-05 |
| 4.45E-05 |
| 4.50E-05 |
| 4.51E-05 |
| 4.51E-05 |
| 4.51E-05 |
| 4.51E-05 |
| 4.55E-05 |
| 4.56E-05 |
| 4.57E-05 |

| |
|---|
| 4.65E-05 |
| 4.66E-05 |
| 4.66E-05 |
| 4.67E-05 |
| 4.68E-05 |
| 4.74E-05 |
| 4.76E-05 |
| 4.76E-05 |
| 4.79E-05 |
| 4.82E-05 |
| 4.82E-05 |
| 4.86E-05 |
| 4.86E-05 |
| 4.88E-05 |
| 4.91E-05 |
| 4.94E-05 |
| 4.94E-05 |
| 4.99E-05 |
| 5.01E-05 |
| 5.02E-05 |
| 5.04E-05 |
| 5.05E-05 |
| 5.07E-05 |
| 5.07E-05 |
| 5.07E-05 |
| 5.08E-05 |
| 5.15E-05 |
| 5.16E-05 |
| 5.25E-05 |
| 5.26E-05 |
| 5.35E-05 |
| 5.36E-05 |
| 5.36E-05 |
| 5.40E-05 |
| 5.44E-05 |
| 5.47E-05 |
| 5.50E-05 |
| 5.50E-05 |
| 5.50E-05 |
| 5.56E-05 |
| 5.65E-05 |

| |
|---|
| 5.68E-05 |
| 5.77E-05 |
| 5.80E-05 |
| 5.80E-05 |
| 5.85E-05 |
| 5.93E-05 |
| 5.93E-05 |
| 5.94E-05 |
| 5.97E-05 |
| 5.99E-05 |
| 6.12E-05 |
| 6.13E-05 |
| 6.22E-05 |
| 6.22E-05 |
| 6.23E-05 |
| 6.26E-05 |
| 6.31E-05 |
| 6.31E-05 |
| 6.31E-05 |
| 6.33E-05 |
| 6.34E-05 |
| 6.34E-05 |
| 6.45E-05 |
| 6.46E-05 |
| 6.47E-05 |
| 6.49E-05 |
| 6.54E-05 |
| 6.56E-05 |
| 6.56E-05 |
| 6.56E-05 |
| 6.58E-05 |
| 6.58E-05 |
| 6.58E-05 |
| 6.63E-05 |
| 6.66E-05 |
| 6.69E-05 |
| 6.69E-05 |
| 6.69E-05 |
| 6.71E-05 |
| 6.75E-05 |
| 6.78E-05 |

6.79E-05
6.79E-05
6.82E-05
6.91E-05
6.99E-05
7.01E-05
7.07E-05
7.07E-05
7.09E-05
7.12E-05
7.18E-05
7.40E-05
7.40E-05
7.51E-05
7.61E-05
7.68E-05
7.70E-05
7.79E-05
7.83E-05
7.83E-05
7.84E-05
7.85E-05
7.87E-05
7.88E-05
7.90E-05
7.93E-05
8.02E-05
8.02E-05
8.12E-05
8.22E-05
8.23E-05
8.24E-05
8.28E-05
8.29E-05
8.39E-05
8.46E-05
8.47E-05
8.67E-05
8.74E-05
8.76E-05
8.77E-05

| |
|---|
| 8.81E-05 |
| 8.84E-05 |
| 8.84E-05 |
| 8.86E-05 |
| 8.93E-05 |
| 9.04E-05 |
| 9.29E-05 |
| 9.29E-05 |
| 9.31E-05 |
| 9.31E-05 |
| 9.60E-05 |
| 9.63E-05 |
| 9.66E-05 |
| 9.67E-05 |
| 9.68E-05 |
| 9.71E-05 |
| 9.73E-05 |
| 9.86E-05 |
| 9.91E-05 |
| 9.93E-05 |
| 9.98E-05 |
| 0.00010152 |
| 0.00010188 |
| 0.00010188 |
| 0.00010227 |
| 0.00010281 |
| 0.00010608 |
| 0.00010674 |
| 0.00010711 |
| 0.00010796 |
| 0.00010885 |
| 0.00010967 |
| 0.00011019 |
| 0.0001107 |
| 0.00011118 |
| 0.00011143 |
| 0.00011176 |
| 0.00011176 |
| 0.00011176 |
| 0.00011381 |
| 0.00011431 |

| |
|---|
| 0.00011434 |
| 0.00011495 |
| 0.00011502 |
| 0.00011515 |
| 0.00011558 |
| 0.00011642 |
| 0.00011642 |
| 0.00011642 |
| 0.00011642 |
| 0.00011663 |
| 0.00011671 |
| 0.00011692 |
| 0.00011692 |
| 0.00011766 |
| 0.00011779 |
| 0.0001183 |
| 0.00011837 |
| 0.00011985 |
| 0.00011987 |
| 0.00012068 |
| 0.00012125 |
| 0.00012158 |
| 0.00012392 |
| 0.00012549 |
| 0.00012713 |
| 0.00012856 |
| 0.00012889 |
| 0.00012949 |
| 0.00012954 |
| 0.00013037 |
| 0.00013082 |
| 0.00013112 |
| 0.00013304 |
| 0.00013444 |
| 0.00013444 |
| 0.00013485 |
| 0.00013569 |
| 0.00013815 |
| 0.0001409 |
| 0.00014102 |
| 0.00014196 |

| |
|---|
| 0.00014253 |
| 0.00014344 |
| 0.00014448 |
| 0.0001452 |
| 0.00014584 |
| 0.00014715 |
| 0.00014725 |
| 0.00014912 |
| 0.00014987 |
| 0.00015008 |
| 0.00015008 |
| 0.00015112 |
| 0.00015242 |
| 0.00015287 |
| 0.00015318 |
| 0.00015411 |
| 0.00015602 |
| 0.00015636 |
| 0.00016137 |
| 0.00016156 |
| 0.00016209 |
| 0.0001631 |
| 0.0001631 |
| 0.00016485 |
| 0.00016598 |
| 0.00016662 |
| 0.00016713 |
| 0.00016879 |
| 0.0001705 |
| 0.00017118 |
| 0.00017556 |
| 0.00017639 |
| 0.00017659 |
| 0.00017759 |
| 0.00017843 |
| 0.00017847 |
| 0.00017875 |
| 0.00018033 |
| 0.00018145 |
| 0.00018316 |
| 0.00018381 |

| |
|---|
| 0.00018402 |
| 0.00018454 |
| 0.00019087 |
| 0.00019354 |
| 0.00019422 |
| 0.0001947 |
| 0.00019747 |
| 0.00019802 |
| 0.0001981 |
| 0.00019889 |
| 0.00019921 |
| 0.00020057 |
| 0.00020057 |
| 0.00020057 |
| 0.0002017 |
| 0.00020374 |
| 0.00020479 |
| 0.00020595 |
| 0.00020805 |
| 0.00020865 |
| 0.00021242 |
| 0.00021242 |
| 0.00021411 |
| 0.00021514 |
| 0.0002192 |
| 0.00022267 |
| 0.00022284 |
| 0.00022515 |
| 0.0002257 |
| 0.00022785 |
| 0.00022857 |
| 0.00022862 |
| 0.00022977 |
| 0.00023086 |
| 0.00023255 |
| 0.0002338 |
| 0.00023445 |
| 0.00023491 |
| 0.00023524 |
| 0.00023581 |
| 0.00023638 |

| |
|---|
| 0.00023716 |
| 0.00023716 |
| 0.00024037 |
| 0.0002406 |
| 0.00024066 |
| 0.0002426 |
| 0.0002426 |
| 0.00024384 |
| 0.00024467 |
| 0.00024597 |
| 0.00024597 |
| 0.00024637 |
| 0.00024869 |
| 0.00024905 |
| 0.00025242 |
| 0.00025352 |
| 0.00025442 |
| 0.00025688 |
| 0.00025821 |
| 0.00025955 |
| 0.00026382 |
| 0.00026703 |
| 0.0002706 |
| 0.00027129 |
| 0.00027435 |
| 0.00027757 |
| 0.00027805 |
| 0.00028085 |
| 0.00028288 |
| 0.00028373 |
| 0.00028421 |
| 0.00028537 |
| 0.00028715 |
| 0.00028755 |
| 0.00028808 |
| 0.00028968 |
| 0.00028998 |
| 0.00029024 |
| 0.00029093 |
| 0.00029093 |
| 0.00029343 |

| |
|---|
| 0.00029623 |
| 0.00029859 |
| 0.00029895 |
| 0.00030278 |
| 0.00030435 |
| 0.00030647 |
| 0.00030647 |
| 0.000307 |
| 0.0003077 |
| 0.00031068 |
| 0.00031079 |
| 0.0003145 |
| 0.0003145 |
| 0.00031536 |
| 0.00031755 |
| 0.00031762 |
| 0.00031841 |
| 0.000322 |
| 0.00032286 |
| 0.000323 |
| 0.00032425 |
| 0.00032425 |
| 0.00032502 |
| 0.00032551 |
| 0.00032637 |
| 0.00032648 |
| 0.00032678 |
| 0.00032696 |
| 0.00033282 |
| 0.00033331 |
| 0.00033484 |
| 0.00033784 |
| 0.00033823 |
| 0.00033966 |
| 0.00034353 |
| 0.00034426 |
| 0.00034429 |
| 0.00034536 |
| 0.00034575 |
| 0.00034654 |
| 0.0003475 |

| |
|---|
| 0.00034812 |
| 0.00034825 |
| 0.00035128 |
| 0.0003537 |
| 0.0003554 |
| 0.00035942 |
| 0.00035987 |
| 0.00036139 |
| 0.00036194 |
| 0.00036409 |
| 0.00036443 |
| 0.00036495 |
| 0.00036574 |
| 0.00036611 |
| 0.00037235 |
| 0.00037522 |
| 0.0003774 |
| 0.00037815 |
| 0.00037916 |
| 0.00038044 |
| 0.00038617 |
| 0.00038888 |
| 0.00038889 |
| 0.00039106 |
| 0.00039333 |
| 0.00039534 |
| 0.00039622 |
| 0.00039646 |
| 0.0004004 |
| 0.00040303 |
| 0.0004035 |
| 0.00040382 |
| 0.00040449 |
| 0.00040714 |
| 0.00040959 |
| 0.00040959 |
| 0.00041024 |
| 0.00041144 |
| 0.00041482 |
| 0.00041698 |
| 0.00041821 |

| |
|---|
| 0.00042164 |
| 0.00042164 |
| 0.00042319 |
| 0.00042377 |
| 0.00042377 |
| 0.00042757 |
| 0.00042817 |
| 0.00043274 |
| 0.00043308 |
| 0.00043413 |
| 0.00043434 |
| 0.0004346 |
| 0.00043463 |
| 0.00043711 |
| 0.00043729 |
| 0.00043794 |
| 0.00043888 |
| 0.00043904 |
| 0.00044274 |
| 0.00044404 |
| 0.0004446 |
| 0.00044544 |
| 0.00044702 |
| 0.00044905 |
| 0.00045024 |
| 0.00045106 |
| 0.00045441 |
| 0.00045873 |
| 0.00045873 |
| 0.00045982 |
| 0.00046014 |
| 0.00046157 |
| 0.00046792 |
| 0.00046894 |
| 0.00047219 |
| 0.0004781 |
| 0.0004801 |
| 0.00048039 |
| 0.00048041 |
| 0.0004805 |
| 0.00048073 |

| |
|---|
| 0.00048078 |
| 0.0004809 |
| 0.0004816 |
| 0.00048218 |
| 0.00048696 |
| 0.00048796 |
| 0.00048827 |
| 0.00049068 |
| 0.00049072 |
| 0.00049689 |
| 0.00049843 |
| 0.00049926 |
| 0.00050188 |
| 0.0005032 |
| 0.00050324 |
| 0.00050605 |
| 0.00050829 |
| 0.00050866 |
| 0.00050928 |
| 0.00051464 |
| 0.00051464 |
| 0.0005172 |
| 0.00052157 |
| 0.00052173 |
| 0.00052318 |
| 0.00052485 |
| 0.00052644 |
| 0.00052929 |
| 0.00053044 |
| 0.00053442 |
| 0.0005389 |
| 0.00053921 |
| 0.00054709 |
| 0.0005586 |
| 0.00055915 |
| 0.00055973 |
| 0.00056081 |
| 0.0005653 |
| 0.0005653 |
| 0.00056561 |
| 0.00056717 |

| |
|---|
| 0.00056731 |
| 0.00057073 |
| 0.00057295 |
| 0.00057568 |
| 0.00057807 |
| 0.0005792 |
| 0.00057925 |
| 0.00058396 |
| 0.00058396 |
| 0.00058785 |
| 0.00059357 |
| 0.00059377 |
| 0.00059553 |
| 0.00059749 |
| 0.00060471 |
| 0.00061473 |
| 0.00061884 |
| 0.00061935 |
| 0.00061971 |
| 0.00061971 |
| 0.00062671 |
| 0.00062901 |
| 0.00063238 |
| 0.0006331 |
| 0.00063419 |
| 0.00063672 |
| 0.00063881 |
| 0.00064082 |
| 0.00065226 |
| 0.00065533 |
| 0.00065533 |
| 0.00065752 |
| 0.00065844 |
| 0.00066084 |
| 0.00066267 |
| 0.00066375 |
| 0.00066375 |
| 0.00066453 |
| 0.00067381 |
| 0.00067899 |
| 0.00067916 |

| |
|---|
| 0.00068245 |
| 0.00068564 |
| 0.00068733 |
| 0.00068833 |
| 0.00069112 |
| 0.00069285 |
| 0.00069534 |
| 0.00069727 |
| 0.00069914 |
| 0.00069968 |
| 0.00070215 |
| 0.00070215 |
| 0.00070373 |
| 0.00070672 |
| 0.00070733 |
| 0.00070912 |
| 0.00070916 |
| 0.00070935 |
| 0.00071175 |
| 0.00071526 |
| 0.00071703 |
| 0.00071976 |
| 0.00072378 |
| 0.00072573 |
| 0.00073023 |
| 0.00074592 |
| 0.00074975 |
| 0.0007504 |
| 0.00075401 |
| 0.0007641 |
| 0.00076438 |
| 0.00076694 |
| 0.00076701 |
| 0.00077021 |
| 0.00077021 |
| 0.00077522 |
| 0.00079165 |
| 0.00079552 |
| 0.00079791 |
| 0.00079923 |
| 0.00080874 |

| |
|---|
| 0.00081195 |
| 0.00082127 |
| 0.00082265 |
| 0.00082265 |
| 0.00082377 |
| 0.00082542 |
| 0.00082705 |
| 0.00082795 |
| 0.00082896 |
| 0.00082984 |
| 0.00083549 |
| 0.00083549 |
| 0.00083639 |
| 0.00083639 |
| 0.00083639 |
| 0.00083996 |
| 0.00084642 |
| 0.00084792 |
| 0.00084792 |
| 0.00084909 |
| 0.00085364 |
| 0.00085527 |
| 0.00085585 |
| 0.00085737 |
| 0.00085849 |
| 0.00086343 |
| 0.0008668 |
| 0.00086698 |
| 0.0008684 |
| 0.00087052 |
| 0.00087052 |
| 0.00087464 |
| 0.00087576 |
| 0.00087756 |
| 0.00087789 |
| 0.00087811 |
| 0.00087867 |
| 0.00088391 |
| 0.0008868 |
| 0.00088781 |
| 0.00088962 |

0.00089129
0.00089203
0.0008971
0.00090748
0.00090783
0.00090873
0.0009092
0.00092317
0.00092317
0.0009351
0.00094646
0.00094704
0.00095134
0.00095306
0.00095961
0.00096157
0.00096598
0.00096746
0.00096808
0.00096998
0.00097475
0.00097584
0.00097656
0.0009785
0.00098151
0.0009952
0.00099868
0.00100423
0.00101633
0.00101739
0.00101746
0.00102667
0.00103989
0.00103989
0.00104034
0.00104128
0.00104798
0.00104798
0.00105228
0.00105462
0.00105853

| |
|---|
| 0.0010609 |
| 0.00106199 |
| 0.00107026 |
| 0.00107026 |
| 0.00107544 |
| 0.00107672 |
| 0.0010852 |
| 0.00108757 |
| 0.00108814 |
| 0.00108994 |
| 0.00109371 |
| 0.00109555 |
| 0.0010973 |
| 0.00112105 |
| 0.00112481 |
| 0.00113016 |
| 0.00113818 |
| 0.00114231 |
| 0.00114487 |
| 0.00114912 |
| 0.00115097 |
| 0.00115097 |
| 0.00115128 |
| 0.00115321 |
| 0.00115943 |
| 0.00116009 |
| 0.00116127 |
| 0.00116127 |
| 0.00117447 |
| 0.00117447 |
| 0.00117447 |
| 0.0011762 |
| 0.00117911 |
| 0.00118086 |
| 0.0011839 |
| 0.00118639 |
| 0.0011976 |
| 0.00120517 |
| 0.00121386 |
| 0.00121849 |
| 0.00123114 |

| |
|---|
| 0.00123118 |
| 0.00124255 |
| 0.00124902 |
| 0.00126247 |
| 0.00126247 |
| 0.00126509 |
| 0.00128012 |
| 0.00128289 |
| 0.00128742 |
| 0.00129201 |
| 0.00129272 |
| 0.00129431 |
| 0.00131961 |
| 0.00132222 |
| 0.00132377 |
| 0.00132636 |
| 0.00133365 |
| 0.00133377 |
| 0.00134109 |
| 0.00134577 |
| 0.00134577 |
| 0.00135208 |
| 0.00135756 |
| 0.00136924 |
| 0.00137822 |
| 0.00138208 |
| 0.00138257 |
| 0.00139026 |
| 0.00139026 |
| 0.00139137 |
| 0.00139385 |
| 0.0013952 |
| 0.0014036 |
| 0.00140689 |
| 0.00140712 |
| 0.00140741 |
| 0.00140818 |
| 0.00140893 |
| 0.00141012 |
| 0.00141151 |
| 0.0014197 |

| |
|---|
| 0.00142268 |
| 0.00142696 |
| 0.00143037 |
| 0.001441 |
| 0.00144182 |
| 0.001459 |
| 0.00145953 |
| 0.00146705 |
| 0.00147928 |
| 0.00148066 |
| 0.00148354 |
| 0.00148747 |
| 0.00148872 |
| 0.00149004 |
| 0.00149313 |
| 0.00149313 |
| 0.00149542 |
| 0.0015024 |
| 0.00150973 |
| 0.00151078 |
| 0.0015129 |
| 0.00151296 |
| 0.00151338 |
| 0.00151586 |
| 0.00151809 |
| 0.00152961 |
| 0.00153012 |
| 0.00153012 |
| 0.00156007 |
| 0.00157754 |
| 0.00158607 |
| 0.00159046 |
| 0.00159346 |
| 0.00159346 |
| 0.00159498 |
| 0.00160062 |
| 0.00160183 |
| 0.00160639 |
| 0.00163083 |
| 0.00163248 |
| 0.00163868 |

| |
|---|
| 0.00163997 |
| 0.00164202 |
| 0.00164735 |
| 0.00164735 |
| 0.00165185 |
| 0.00166533 |
| 0.00168261 |
| 0.00168261 |
| 0.00170225 |
| 0.00170249 |
| 0.00171644 |
| 0.00171644 |
| 0.00171889 |
| 0.0017193 |
| 0.00172575 |
| 0.00172891 |
| 0.00172891 |
| 0.0017371 |
| 0.0017371 |
| 0.0017371 |
| 0.00174065 |
| 0.00174654 |
| 0.00176643 |
| 0.00176738 |
| 0.00177224 |
| 0.00177326 |
| 0.00177326 |
| 0.00178283 |
| 0.00178994 |
| 0.00179087 |
| 0.00179177 |
| 0.00179454 |
| 0.00180065 |
| 0.00180088 |
| 0.00181334 |
| 0.00181415 |
| 0.00183821 |
| 0.00183942 |
| 0.00184332 |
| 0.00185508 |
| 0.00185788 |

| |
|---|
| 0.00186063 |
| 0.0018632 |
| 0.00188012 |
| 0.00190047 |
| 0.0019076 |
| 0.00191145 |
| 0.00191629 |
| 0.00192932 |
| 0.00192952 |
| 0.00192952 |
| 0.00193532 |
| 0.00193695 |
| 0.00195374 |
| 0.00195966 |
| 0.00197149 |
| 0.00198169 |
| 0.00199031 |
| 0.00199632 |
| 0.00200135 |
| 0.00200136 |
| 0.00200298 |
| 0.00200938 |
| 0.00201139 |
| 0.00201139 |
| 0.00201359 |
| 0.00201434 |
| 0.00202111 |
| 0.00202316 |
| 0.00203175 |
| 0.00203916 |
| 0.00203916 |
| 0.00203916 |
| 0.00204749 |
| 0.00207163 |
| 0.00207163 |
| 0.00207389 |
| 0.00207389 |
| 0.00207389 |
| 0.00207921 |
| 0.00207921 |
| 0.00208171 |

| |
|---|
| 0.00209232 |
| 0.00212891 |
| 0.00214394 |
| 0.00214518 |
| 0.00215317 |
| 0.00216452 |
| 0.00220109 |
| 0.00220371 |
| 0.00221657 |
| 0.00221682 |
| 0.00221728 |
| 0.00222077 |
| 0.00222525 |
| 0.00222528 |
| 0.00222646 |
| 0.00224619 |
| 0.00224623 |
| 0.00225227 |
| 0.00225664 |
| 0.00225977 |
| 0.00228164 |
| 0.00230665 |
| 0.00232728 |
| 0.00233246 |
| 0.00233246 |
| 0.00234361 |
| 0.00236187 |
| 0.00236637 |
| 0.0023711 |
| 0.0023711 |
| 0.00238274 |
| 0.00238446 |
| 0.00238446 |
| 0.00238524 |
| 0.00241408 |
| 0.00242055 |
| 0.0024235 |
| 0.00242864 |
| 0.00242864 |
| 0.00243308 |
| 0.00243382 |

| |
|---|
| 0.00244039 |
| 0.00244615 |
| 0.00244615 |
| 0.00244888 |
| 0.00245523 |
| 0.00245963 |
| 0.00246119 |
| 0.002465 |
| 0.00248429 |
| 0.00248565 |
| 0.00248676 |
| 0.00248908 |
| 0.00249588 |
| 0.00249661 |
| 0.00250808 |
| 0.00251968 |
| 0.00252988 |
| 0.00253001 |
| 0.00253001 |
| 0.0025605 |
| 0.00256756 |
| 0.00257073 |
| 0.00257094 |
| 0.00257301 |
| 0.00257466 |
| 0.00257873 |
| 0.00259054 |
| 0.00259285 |
| 0.00259533 |
| 0.00259533 |
| 0.0026067 |
| 0.00265232 |
| 0.0026537 |
| 0.00266191 |
| 0.00266377 |
| 0.00266642 |
| 0.0026834 |
| 0.00270011 |
| 0.00270568 |
| 0.00271168 |
| 0.00271648 |

| |
|---|
| 0.00274154 |
| 0.00275683 |
| 0.00275683 |
| 0.00277431 |
| 0.00277835 |
| 0.00278181 |
| 0.00278294 |
| 0.00281808 |
| 0.00281964 |
| 0.00281964 |
| 0.00282539 |
| 0.00282702 |
| 0.00283933 |
| 0.00284216 |
| 0.00285369 |
| 0.00285369 |
| 0.00285369 |
| 0.00286973 |
| 0.00286996 |
| 0.00287489 |
| 0.00287489 |
| 0.00288196 |
| 0.00289294 |
| 0.00289294 |
| 0.00289669 |
| 0.00291295 |
| 0.00291685 |
| 0.00291734 |
| 0.00292635 |
| 0.0029308 |
| 0.00294051 |
| 0.00294051 |
| 0.00294589 |
| 0.00297993 |
| 0.00299008 |
| 0.00300532 |
| 0.00304793 |
| 0.00305105 |
| 0.00309266 |
| 0.003094 |
| 0.00311663 |

| |
|---|
| 0.00314741 |
| 0.00315409 |
| 0.00316036 |
| 0.00316829 |
| 0.00317001 |
| 0.00317958 |
| 0.00319089 |
| 0.00320716 |
| 0.00321099 |
| 0.00322073 |
| 0.00324834 |
| 0.00327162 |
| 0.00329505 |
| 0.00331108 |
| 0.00331901 |
| 0.00332184 |
| 0.00332504 |
| 0.00333197 |
| 0.00335476 |
| 0.00336177 |
| 0.00337044 |
| 0.00337178 |
| 0.00338166 |
| 0.00339777 |
| 0.00340812 |
| 0.0034183 |
| 0.00343409 |
| 0.00344397 |
| 0.00348034 |
| 0.00348034 |
| 0.00350424 |
| 0.00351419 |
| 0.00351526 |
| 0.0035267 |
| 0.00353095 |
| 0.00353427 |
| 0.00354282 |
| 0.00355399 |
| 0.00356034 |
| 0.0035839 |
| 0.00359897 |

| |
|---|
| 0.00365161 |
| 0.0036551 |
| 0.00365996 |
| 0.00368249 |
| 0.00368896 |
| 0.0036898 |
| 0.00369014 |
| 0.00369667 |
| 0.00371535 |
| 0.00372682 |
| 0.00372797 |
| 0.00373693 |
| 0.00374083 |
| 0.00379722 |
| 0.0038124 |
| 0.00383139 |
| 0.00383858 |
| 0.00384956 |
| 0.00385474 |
| 0.00386599 |
| 0.00386848 |
| 0.00389002 |
| 0.00390031 |
| 0.00390255 |
| 0.00390724 |
| 0.00392409 |
| 0.00394959 |
| 0.00395258 |
| 0.00396096 |
| 0.00396732 |
| 0.00397544 |
| 0.00398481 |
| 0.00400872 |
| 0.00401111 |
| 0.00402074 |
| 0.00402559 |
| 0.00403583 |
| 0.00406825 |
| 0.00408345 |
| 0.00410372 |
| 0.00410372 |

| |
|---|
| 0.00410635 |
| 0.00411747 |
| 0.00413227 |
| 0.00417464 |
| 0.00418483 |
| 0.00419184 |
| 0.00419768 |
| 0.00419768 |
| 0.00421224 |
| 0.00422171 |
| 0.00422803 |
| 0.00424427 |
| 0.00424678 |
| 0.00425966 |
| 0.00429724 |
| 0.00429724 |
| 0.0042984 |
| 0.00432429 |
| 0.00433014 |
| 0.00433014 |
| 0.00433755 |
| 0.0043423 |
| 0.00437398 |
| 0.00437584 |
| 0.00441526 |
| 0.00441872 |
| 0.00443878 |
| 0.00446705 |
| 0.00446705 |
| 0.00447096 |
| 0.00447778 |
| 0.00449659 |
| 0.00451628 |
| 0.00451628 |
| 0.00452517 |
| 0.00452963 |
| 0.00453617 |
| 0.00453837 |
| 0.00453837 |
| 0.00453893 |
| 0.00455713 |

| |
|---|
| 0.00456027 |
| 0.00461201 |
| 0.00465351 |
| 0.00465535 |
| 0.00465634 |
| 0.00465634 |
| 0.00468358 |
| 0.00469434 |
| 0.00469794 |
| 0.00471233 |
| 0.00471575 |
| 0.0047176 |
| 0.00472702 |
| 0.00472702 |
| 0.00474017 |
| 0.00474017 |
| 0.00474168 |
| 0.00475643 |
| 0.00478508 |
| 0.0047956 |
| 0.00480218 |
| 0.00481467 |
| 0.004836 |
| 0.00484698 |
| 0.00486729 |
| 0.00487743 |
| 0.00488442 |
| 0.00499231 |
| 0.00500447 |
| 0.00500927 |
| 0.00501037 |
| 0.00501907 |
| 0.00502594 |
| 0.00507842 |
| 0.00508026 |
| 0.00508515 |
| 0.00510371 |
| 0.00511549 |
| 0.00514712 |
| 0.00516208 |
| 0.0051818 |

| |
|---|
| 0.00519577 |
| 0.00521241 |
| 0.00521856 |
| 0.00523058 |
| 0.00523418 |
| 0.00526849 |
| 0.00528534 |
| 0.00529261 |
| 0.00529587 |
| 0.00532004 |
| 0.00532905 |
| 0.00533024 |
| 0.00536608 |
| 0.0053777 |
| 0.00540431 |
| 0.00542214 |
| 0.00544715 |
| 0.00545413 |
| 0.00546335 |
| 0.00548751 |
| 0.00553754 |
| 0.00554336 |
| 0.00554714 |
| 0.00555273 |
| 0.00556717 |
| 0.00557493 |
| 0.00558909 |
| 0.00561161 |
| 0.00562122 |
| 0.00562833 |
| 0.00563676 |
| 0.00563848 |
| 0.00568327 |
| 0.00569082 |
| 0.00576447 |
| 0.00576586 |
| 0.00577115 |
| 0.00577704 |
| 0.00577835 |
| 0.00578076 |
| 0.00580741 |

| |
|---|
| 0.00580741 |
| 0.00585059 |
| 0.00588173 |
| 0.00588307 |
| 0.00588746 |
| 0.00589286 |
| 0.00589767 |
| 0.00590299 |
| 0.00592288 |
| 0.00593093 |
| 0.00593093 |
| 0.00594586 |
| 0.00595744 |
| 0.00597907 |
| 0.00598152 |
| 0.0061019 |
| 0.00616294 |
| 0.00618034 |
| 0.00619198 |
| 0.00619358 |
| 0.00619512 |
| 0.00622024 |
| 0.00624466 |
| 0.00624893 |
| 0.00628547 |
| 0.00635225 |
| 0.00635432 |
| 0.00636303 |
| 0.00641159 |
| 0.00645142 |
| 0.00651444 |
| 0.00651791 |
| 0.00652265 |
| 0.00657142 |
| 0.00657532 |
| 0.00663799 |
| 0.00664524 |
| 0.00670757 |
| 0.00671191 |
| 0.00671191 |
| 0.00673738 |

| |
|---|
| 0.0067888 |
| 0.00678983 |
| 0.00679166 |
| 0.00679394 |
| 0.00679394 |
| 0.00682334 |
| 0.00684583 |
| 0.00686056 |
| 0.00686157 |
| 0.00686157 |
| 0.00686343 |
| 0.00687006 |
| 0.00687036 |
| 0.0068753 |
| 0.00687726 |
| 0.00690261 |
| 0.00690261 |
| 0.00693989 |
| 0.00700633 |
| 0.0070182 |
| 0.00704906 |
| 0.00706381 |
| 0.00711843 |
| 0.00722682 |
| 0.00723257 |
| 0.00727597 |
| 0.00727915 |
| 0.0072987 |
| 0.00732085 |
| 0.00735836 |
| 0.00739365 |
| 0.00743454 |
| 0.00743993 |
| 0.00743993 |
| 0.0074581 |
| 0.00747501 |
| 0.00749101 |
| 0.00749642 |
| 0.00749788 |
| 0.00749788 |
| 0.00750078 |

| |
|---|
| 0.00750078 |
| 0.00750407 |
| 0.00751272 |
| 0.00751384 |
| 0.00753727 |
| 0.00753783 |
| 0.00753783 |
| 0.00755514 |
| 0.00755795 |
| 0.0075958 |
| 0.00760767 |
| 0.00761135 |
| 0.00762243 |
| 0.00762962 |
| 0.00762962 |
| 0.00764 |
| 0.00765158 |
| 0.0076606 |
| 0.00773861 |
| 0.00774883 |
| 0.00776257 |
| 0.0077706 |
| 0.00778447 |
| 0.00778447 |
| 0.00779675 |
| 0.00779675 |
| 0.00783335 |
| 0.00785568 |
| 0.00785819 |
| 0.00786321 |
| 0.00786337 |
| 0.00789398 |
| 0.00789844 |
| 0.00799254 |
| 0.00800157 |
| 0.00802357 |
| 0.0080509 |
| 0.0080509 |
| 0.00808959 |
| 0.00809797 |
| 0.00811083 |

| |
|---|
| 0.00812715 |
| 0.00812876 |
| 0.0081359 |
| 0.00814865 |
| 0.00820285 |
| 0.00821437 |
| 0.00821437 |
| 0.00830265 |
| 0.00830835 |
| 0.00831472 |
| 0.00832386 |
| 0.00833876 |
| 0.00835787 |
| 0.0083681 |
| 0.00846243 |
| 0.00846489 |
| 0.00849017 |
| 0.00849142 |
| 0.00849497 |
| 0.00852602 |
| 0.00856857 |
| 0.00856918 |
| 0.00856918 |
| 0.00857266 |
| 0.00857586 |
| 0.00858675 |
| 0.00860442 |
| 0.0086361 |
| 0.008637 |
| 0.00863729 |
| 0.00865999 |
| 0.00866314 |
| 0.00869199 |
| 0.00872891 |
| 0.008749 |
| 0.008749 |
| 0.00875169 |
| 0.00877639 |
| 0.0087823 |
| 0.0087823 |
| 0.0087823 |

| |
|---|
| 0.00881776 |
| 0.0088731 |
| 0.00887934 |
| 0.00894171 |
| 0.00896528 |
| 0.00896528 |
| 0.0089739 |
| 0.00898383 |
| 0.00906159 |
| 0.0090618 |
| 0.00907059 |
| 0.00911132 |
| 0.00911493 |
| 0.00912915 |
| 0.00915989 |
| 0.00917549 |
| 0.00918103 |
| 0.00922888 |
| 0.00926001 |
| 0.00932296 |
| 0.0093397 |
| 0.0093397 |
| 0.00934633 |
| 0.00934753 |
| 0.00938893 |
| 0.00941859 |
| 0.00947974 |
| 0.00947974 |
| 0.00952057 |
| 0.00953186 |
| 0.0095364 |
| 0.0095364 |
| 0.00953908 |
| 0.00953973 |
| 0.00954022 |
| 0.00955053 |
| 0.00958683 |
| 0.00958683 |
| 0.00963548 |
| 0.00964551 |
| 0.00968723 |

| |
|---|
| 0.00978195 |
| 0.00979154 |
| 0.00985278 |
| 0.0098818 |
| 0.0098818 |
| 0.00991191 |
| 0.00992816 |
| 0.00994739 |
| 0.01000855 |
| 0.01005145 |
| 0.01008014 |
| 0.01016931 |
| 0.01019505 |
| 0.01023524 |
| 0.01026649 |
| 0.01028156 |
| 0.01028227 |
| 0.0103514 |
| 0.0103514 |
| 0.01039912 |
| 0.01043137 |
| 0.01049317 |
| 0.01054189 |
| 0.01058149 |
| 0.01062783 |
| 0.01063988 |
| 0.0106421 |
| 0.01065125 |
| 0.01066602 |
| 0.0108775 |
| 0.01089198 |
| 0.01089999 |
| 0.0109197 |
| 0.01102967 |
| 0.01103749 |
| 0.01108241 |
| 0.01108241 |
| 0.01110857 |
| 0.01111174 |
| 0.01115736 |
| 0.01123534 |

| |
|---|
| 0.01123777 |
| 0.01124089 |
| 0.01135097 |
| 0.01135898 |
| 0.01137473 |
| 0.011411 |
| 0.011411 |
| 0.01143491 |
| 0.01147559 |
| 0.0114922 |
| 0.0114922 |
| 0.0115387 |
| 0.01155792 |
| 0.01157565 |
| 0.01158382 |
| 0.01163192 |
| 0.0116636 |
| 0.01167649 |
| 0.01170282 |
| 0.0117849 |
| 0.01178647 |
| 0.01179743 |
| 0.01179743 |
| 0.01180157 |
| 0.01187088 |
| 0.01188794 |
| 0.01188988 |
| 0.01190464 |
| 0.01207053 |
| 0.01210906 |
| 0.0121434 |
| 0.01241237 |
| 0.01244177 |
| 0.01245154 |
| 0.01245169 |
| 0.01246278 |
| 0.01246278 |
| 0.01246278 |
| 0.01250379 |
| 0.01252345 |
| 0.01252982 |

| |
|---|
| 0.01255639 |
| 0.0125627 |
| 0.0126052 |
| 0.01263064 |
| 0.01263201 |
| 0.01265431 |
| 0.01265778 |
| 0.01268678 |
| 0.0126989 |
| 0.01271558 |
| 0.01273253 |
| 0.01278195 |
| 0.01278697 |
| 0.01280235 |
| 0.01284173 |
| 0.01290256 |
| 0.01290638 |
| 0.01292464 |
| 0.01301692 |
| 0.01305783 |
| 0.01311069 |
| 0.01315847 |
| 0.01320609 |
| 0.0132816 |
| 0.01332183 |
| 0.01333441 |
| 0.01336173 |
| 0.01336867 |
| 0.01339783 |
| 0.01345657 |
| 0.01346219 |
| 0.01355406 |
| 0.01355406 |
| 0.01355406 |
| 0.01356013 |
| 0.01356013 |
| 0.01358794 |
| 0.01360114 |
| 0.01361433 |
| 0.01363752 |
| 0.01371337 |

| |
|---|
| 0.01373815 |
| 0.01379075 |
| 0.01386418 |
| 0.01386418 |
| 0.01392099 |
| 0.01396027 |
| 0.01397752 |
| 0.01403741 |
| 0.01410956 |
| 0.01412269 |
| 0.01414044 |
| 0.01416442 |
| 0.01425057 |
| 0.0142924 |
| 0.01436655 |
| 0.01437047 |
| 0.01443068 |
| 0.01443342 |
| 0.01445239 |
| 0.01446695 |
| 0.01448032 |
| 0.01450145 |
| 0.01453364 |
| 0.01458951 |
| 0.01461483 |
| 0.01463848 |
| 0.01466113 |
| 0.01472417 |
| 0.01472609 |
| 0.01477545 |
| 0.01478721 |
| 0.01487563 |
| 0.01497361 |
| 0.01497885 |
| 0.01509127 |
| 0.01509444 |
| 0.01511576 |
| 0.01512542 |
| 0.01516028 |
| 0.01516165 |
| 0.01519421 |

| |
|---|
| 0.01522274 |
| 0.01533013 |
| 0.01539048 |
| 0.01539298 |
| 0.01544164 |
| 0.01548409 |
| 0.01553764 |
| 0.01558373 |
| 0.01558373 |
| 0.01558541 |
| 0.01558541 |
| 0.01558541 |
| 0.01559029 |
| 0.01559029 |
| 0.01562802 |
| 0.01569181 |
| 0.01570271 |
| 0.01570651 |
| 0.01582386 |
| 0.01583113 |
| 0.0158421 |
| 0.01599192 |
| 0.01603779 |
| 0.01606057 |
| 0.01608678 |
| 0.01612433 |
| 0.01612433 |
| 0.01623188 |
| 0.01624125 |
| 0.01630121 |
| 0.01635224 |
| 0.01636609 |
| 0.01636609 |
| 0.01637166 |
| 0.01639616 |
| 0.01639616 |
| 0.01640115 |
| 0.01651301 |
| 0.016524 |
| 0.01660642 |
| 0.01662693 |

| |
|---|
| 0.0166707 |
| 0.01673696 |
| 0.01673813 |
| 0.01674715 |
| 0.01678387 |
| 0.01678788 |
| 0.01679011 |
| 0.01679027 |
| 0.01681852 |
| 0.01685321 |
| 0.01689357 |
| 0.01690699 |
| 0.01695305 |
| 0.01696436 |
| 0.01696963 |
| 0.01712447 |
| 0.01713348 |
| 0.01722096 |
| 0.01722096 |
| 0.0173833 |
| 0.0173833 |
| 0.01738479 |
| 0.01747623 |
| 0.01755283 |
| 0.01757405 |
| 0.01764662 |
| 0.01764662 |
| 0.01771697 |
| 0.01772188 |
| 0.01780999 |
| 0.01780999 |
| 0.01783607 |
| 0.01786351 |
| 0.01787299 |
| 0.01789227 |
| 0.01789227 |
| 0.01789227 |
| 0.01794598 |
| 0.0179974 |
| 0.01815889 |
| 0.0183555 |

| |
|---|
| 0.01837331 |
| 0.0184909 |
| 0.0184909 |
| 0.018518 |
| 0.01853976 |
| 0.01859223 |
| 0.01859789 |
| 0.01870463 |
| 0.01875121 |
| 0.01875353 |
| 0.01877765 |
| 0.01881511 |
| 0.01890005 |
| 0.01890691 |
| 0.01896146 |
| 0.01901892 |
| 0.01905729 |
| 0.01905729 |
| 0.01912359 |
| 0.01913012 |
| 0.0191354 |
| 0.01917956 |
| 0.01921332 |
| 0.0193483 |
| 0.01944964 |
| 0.01947974 |
| 0.01949112 |
| 0.01951683 |
| 0.01961292 |
| 0.01969171 |
| 0.01970405 |
| 0.01971398 |
| 0.01976178 |
| 0.01979767 |
| 0.01994728 |
| 0.01995495 |
| 0.01995495 |
| 0.01996585 |
| 0.02000758 |
| 0.02019649 |
| 0.02026635 |

| |
|---|
| 0.02033072 |
| 0.02044142 |
| 0.02044778 |
| 0.02046138 |
| 0.02056779 |
| 0.02064353 |
| 0.02065252 |
| 0.02066038 |
| 0.02072374 |
| 0.02075192 |
| 0.02079838 |
| 0.0209829 |
| 0.02100802 |
| 0.02101916 |
| 0.02101916 |
| 0.02106179 |
| 0.02108879 |
| 0.02109863 |
| 0.02116632 |
| 0.02127009 |
| 0.02130137 |
| 0.02130797 |
| 0.02139941 |
| 0.02141299 |
| 0.02144845 |
| 0.02145498 |
| 0.02146866 |
| 0.02164687 |
| 0.02165738 |
| 0.02166275 |
| 0.02166275 |
| 0.02171186 |
| 0.02172201 |
| 0.02173752 |
| 0.02181075 |
| 0.02182326 |
| 0.02190312 |
| 0.02194584 |
| 0.02200829 |
| 0.0220647 |
| 0.02215935 |

| |
|---|
| 0.02223501 |
| 0.02225593 |
| 0.02232656 |
| 0.0224388 |
| 0.02244213 |
| 0.02244306 |
| 0.02255335 |
| 0.02258423 |
| 0.02258423 |
| 0.02271044 |
| 0.02277916 |
| 0.02280529 |
| 0.02286292 |
| 0.0229377 |
| 0.0229377 |
| 0.0229653 |
| 0.0229653 |
| 0.02309696 |
| 0.02313618 |
| 0.02316499 |
| 0.02317757 |
| 0.02319277 |
| 0.0232582 |
| 0.0232582 |
| 0.02326079 |
| 0.02328813 |
| 0.02328946 |
| 0.02329777 |
| 0.02331977 |
| 0.02334783 |
| 0.02335754 |
| 0.02337848 |
| 0.02340882 |
| 0.02342284 |
| 0.02359909 |
| 0.02361967 |
| 0.02361967 |
| 0.02372978 |
| 0.02383192 |
| 0.0239374 |
| 0.02394668 |

| |
|---|
| 0.0241418 |
| 0.0241418 |
| 0.02417967 |
| 0.02420918 |
| 0.02420918 |
| 0.0242345 |
| 0.02429576 |
| 0.02437106 |
| 0.02437106 |
| 0.02448846 |
| 0.02455031 |
| 0.02457397 |
| 0.02469106 |
| 0.02471405 |
| 0.02476797 |
| 0.0248409 |
| 0.02485273 |
| 0.02487705 |
| 0.02489544 |
| 0.02497916 |
| 0.02498158 |
| 0.02504273 |
| 0.02526947 |
| 0.02527354 |
| 0.02529914 |
| 0.02529914 |
| 0.02539546 |
| 0.02542603 |
| 0.0255867 |
| 0.02561911 |
| 0.02564551 |
| 0.02565759 |
| 0.02574748 |
| 0.02578065 |
| 0.02582722 |
| 0.02584601 |
| 0.02603315 |
| 0.02607892 |
| 0.02614313 |
| 0.02622517 |
| 0.02639833 |

| |
|---|
| 0.02643237 |
| 0.02652007 |
| 0.02655987 |
| 0.02655987 |
| 0.02657052 |
| 0.02660432 |
| 0.02668312 |
| 0.02671057 |
| 0.02682374 |
| 0.02689594 |
| 0.02697475 |
| 0.0270585 |
| 0.02709062 |
| 0.02720271 |
| 0.02720836 |
| 0.02722147 |
| 0.0272373 |
| 0.02724533 |
| 0.0272802 |
| 0.02733861 |
| 0.02738381 |
| 0.0274456 |
| 0.0274456 |
| 0.02745609 |
| 0.02752151 |
| 0.02752167 |
| 0.02752642 |
| 0.02755947 |
| 0.02755947 |
| 0.02756774 |
| 0.02768596 |
| 0.02772547 |
| 0.02779537 |
| 0.02799705 |
| 0.0280011 |
| 0.02812675 |
| 0.02819727 |
| 0.02830576 |
| 0.02832504 |
| 0.02836256 |
| 0.02838317 |

0.02839681
0.02846174
0.02864489
0.02865492
0.02870593
0.0287148
0.02874043
0.02874339
0.02877912
0.02880436
0.02880832
0.02881117
0.02891939
0.02891939
0.02897877
0.0290333
0.0290333
0.0291096
0.02914656
0.02930758
0.0293077
0.02943378
0.02953965
0.0295567
0.029586
0.0295937
0.02959811
0.02960503
0.02962635
0.02974428
0.02977519
0.02978304
0.02985056
0.02985056
0.02987747
0.02995239
0.03004386
0.03004386
0.03009285
0.0300935
0.03015019

| |
|---|
| 0.03024252 |
| 0.03026301 |
| 0.03030851 |
| 0.03036483 |
| 0.03039344 |
| 0.03042559 |
| 0.03044421 |
| 0.03066093 |
| 0.03067829 |
| 0.03068831 |
| 0.0306887 |
| 0.03075679 |
| 0.03087947 |
| 0.03097414 |
| 0.03113708 |
| 0.03126389 |
| 0.03126389 |
| 0.03126389 |
| 0.03126389 |
| 0.03128055 |
| 0.03134097 |
| 0.03134097 |
| 0.03134097 |
| 0.031461 |
| 0.03153642 |
| 0.03159527 |
| 0.0317254 |
| 0.031842 |
| 0.03187827 |
| 0.03187827 |
| 0.03198947 |
| 0.03219151 |
| 0.03224672 |
| 0.03224672 |
| 0.03237502 |
| 0.03237662 |
| 0.03246271 |
| 0.03252199 |
| 0.03257182 |
| 0.03267119 |
| 0.03270614 |

| |
|---|
| 0.03274228 |
| 0.03279195 |
| 0.03286928 |
| 0.0329279 |
| 0.03309667 |
| 0.03312181 |
| 0.03312521 |
| 0.03317825 |
| 0.033188 |
| 0.0332461 |
| 0.03326802 |
| 0.03361405 |
| 0.03377142 |
| 0.03382025 |
| 0.03398206 |
| 0.03401424 |
| 0.03406017 |
| 0.03414328 |
| 0.03431412 |
| 0.03436348 |
| 0.0344896 |
| 0.03451345 |
| 0.03454882 |
| 0.0346026 |
| 0.03464004 |
| 0.03478594 |
| 0.03483037 |
| 0.03483037 |
| 0.03491937 |
| 0.03493121 |
| 0.03497332 |
| 0.03507058 |
| 0.03509493 |
| 0.03513461 |
| 0.03516042 |
| 0.0355586 |
| 0.03560902 |
| 0.03570233 |
| 0.0357696 |
| 0.03586784 |
| 0.03587446 |

| |
|---|
| 0.03602227 |
| 0.03618265 |
| 0.03622965 |
| 0.03622987 |
| 0.03634473 |
| 0.03648567 |
| 0.03650531 |
| 0.03652969 |
| 0.03660735 |
| 0.03668129 |
| 0.03682991 |
| 0.03684939 |
| 0.03689859 |
| 0.03695731 |
| 0.03699563 |
| 0.03700555 |
| 0.03705696 |
| 0.03710582 |
| 0.03719091 |
| 0.03719091 |
| 0.03725201 |
| 0.0372851 |
| 0.03729457 |
| 0.03730591 |
| 0.03734176 |
| 0.03740121 |
| 0.03752762 |
| 0.03756728 |
| 0.03762275 |
| 0.03762751 |
| 0.03762751 |
| 0.03762751 |
| 0.03766149 |
| 0.03768257 |
| 0.03768274 |
| 0.0376946 |
| 0.03769539 |
| 0.03770835 |
| 0.03771739 |
| 0.03773344 |
| 0.03780883 |

| |
|---|
| 0.03789436 |
| 0.03803125 |
| 0.03807555 |
| 0.03811407 |
| 0.03816534 |
| 0.0381925 |
| 0.03828691 |
| 0.03830951 |
| 0.03873383 |
| 0.03874822 |
| 0.0387499 |
| 0.03879318 |
| 0.03887028 |
| 0.03892652 |
| 0.03893384 |
| 0.03900616 |
| 0.03911372 |
| 0.03937764 |
| 0.03970663 |
| 0.03977224 |
| 0.03977224 |
| 0.03979741 |
| 0.03981973 |
| 0.04021432 |
| 0.04024559 |
| 0.04044859 |
| 0.04045545 |
| 0.04047802 |
| 0.04048564 |
| 0.04048938 |
| 0.0405215 |
| 0.04056874 |
| 0.04067712 |
| 0.04067712 |
| 0.04079077 |
| 0.04084075 |
| 0.04089536 |
| 0.04091754 |
| 0.04093855 |
| 0.04093855 |
| 0.04098792 |

| |
|---|
| 0.04110383 |
| 0.04112352 |
| 0.04118601 |
| 0.04119796 |
| 0.04127047 |
| 0.04127047 |
| 0.04132184 |
| 0.04135593 |
| 0.04148241 |
| 0.04149866 |
| 0.04159022 |
| 0.04165606 |
| 0.04168429 |
| 0.04168429 |
| 0.04168512 |
| 0.04183062 |
| 0.04183062 |
| 0.04187455 |
| 0.04205584 |
| 0.04209257 |
| 0.04209257 |
| 0.04212624 |
| 0.04212624 |
| 0.04215186 |
| 0.04217296 |
| 0.04223056 |
| 0.042321 |
| 0.04241493 |
| 0.04242354 |
| 0.04246741 |
| 0.04260051 |
| 0.04277891 |
| 0.04314881 |
| 0.04316886 |
| 0.04325057 |
| 0.04344572 |
| 0.04345304 |
| 0.04346676 |
| 0.04347227 |
| 0.04349458 |
| 0.04363212 |

| |
|---|
| 0.04363212 |
| 0.04364054 |
| 0.0437416 |
| 0.0437416 |
| 0.0437416 |
| 0.04389208 |
| 0.04389732 |
| 0.04391802 |
| 0.0440354 |
| 0.04411085 |
| 0.04411733 |
| 0.04420956 |
| 0.04434754 |
| 0.04471577 |
| 0.04473632 |
| 0.04474027 |
| 0.04474027 |
| 0.0447625 |
| 0.04478149 |
| 0.04488235 |
| 0.04498625 |
| 0.04529977 |
| 0.04547814 |
| 0.04562862 |
| 0.04569321 |
| 0.04572092 |
| 0.04595217 |
| 0.04604405 |
| 0.04611589 |
| 0.04616593 |
| 0.04635264 |
| 0.04637683 |
| 0.04643602 |
| 0.04663789 |
| 0.04663789 |
| 0.04683286 |
| 0.04686623 |
| 0.04697292 |
| 0.04716405 |
| 0.04718577 |
| 0.04726579 |

0.04755159
0.04755159
0.04767627
0.04777682
0.04802642
0.04812214
0.04817665
0.04824138
0.04842858
0.04864552
0.04881227
0.04890351
0.04891412
0.04891412
0.04896437
0.04896437
0.04896437
0.04897819
0.04900842
0.04912001
0.04912648
0.04916181
0.04918939
0.04934469
0.04938043
0.04944311
0.04953717
0.04953717
0.04954983
0.04972819
0.04976666
0.04979485
0.04994088
0.05022913
0.05023737
0.05032382
0.05032595
0.05052945
0.05066636
0.05067128
0.05079777

| |
|---|
| 0.05106727 |
| 0.05107191 |
| 0.05116885 |
| 0.05118557 |
| 0.05123816 |
| 0.05131179 |
| 0.05131251 |
| 0.05131376 |
| 0.05149206 |
| 0.05166403 |
| 0.0518411 |
| 0.05190936 |
| 0.05199124 |
| 0.05220369 |
| 0.05226557 |
| 0.05242469 |
| 0.05262108 |
| 0.05270506 |
| 0.05273688 |
| 0.05285398 |
| 0.05305541 |
| 0.05308782 |
| 0.05314078 |
| 0.05328361 |
| 0.05346064 |
| 0.05380076 |
| 0.05383138 |
| 0.0538431 |
| 0.05395169 |
| 0.05412537 |
| 0.05415293 |
| 0.05424265 |
| 0.05458247 |
| 0.05465839 |
| 0.05514001 |
| 0.0553512 |
| 0.05558094 |
| 0.05558094 |
| 0.05564451 |
| 0.05566173 |
| 0.05577449 |

| |
|---|
| 0.05578103 |
| 0.05587566 |
| 0.05587566 |
| 0.05599562 |
| 0.05599562 |
| 0.05599562 |
| 0.05599562 |
| 0.0560029 |
| 0.05609497 |
| 0.05622895 |
| 0.0562954 |
| 0.05652333 |
| 0.05652527 |
| 0.05655924 |
| 0.05659577 |
| 0.05686561 |
| 0.05704807 |
| 0.05704807 |
| 0.05710008 |
| 0.05712912 |
| 0.05717081 |
| 0.05735164 |
| 0.05745691 |
| 0.05774509 |
| 0.05774509 |
| 0.05774862 |
| 0.057751 |
| 0.0578746 |
| 0.05792502 |
| 0.05792894 |
| 0.057958 |
| 0.05807642 |
| 0.05808744 |
| 0.05827204 |
| 0.05845001 |
| 0.05855848 |
| 0.05863055 |
| 0.05884987 |
| 0.05893308 |
| 0.05901224 |
| 0.05912135 |

| |
|---|
| 0.05920824 |
| 0.05924041 |
| 0.05930292 |
| 0.05954717 |
| 0.05956329 |
| 0.05984664 |
| 0.05984664 |
| 0.06006688 |
| 0.06018192 |
| 0.0602874 |
| 0.06051738 |
| 0.06065807 |
| 0.06065833 |
| 0.06069087 |
| 0.06074417 |
| 0.06084423 |
| 0.0609617 |
| 0.0609617 |
| 0.0609617 |
| 0.06097487 |
| 0.06097487 |
| 0.06100647 |
| 0.06124171 |
| 0.06129046 |
| 0.06129119 |
| 0.06157811 |
| 0.06161412 |
| 0.06161455 |
| 0.06162435 |
| 0.06183301 |
| 0.06186179 |
| 0.06186994 |
| 0.06191643 |
| 0.06211813 |
| 0.06234426 |
| 0.06235071 |
| 0.06251739 |
| 0.06265399 |
| 0.06266976 |
| 0.06270058 |
| 0.06282336 |

| |
|---|
| 0.06306936 |
| 0.06318499 |
| 0.06336468 |
| 0.06347949 |
| 0.06354745 |
| 0.06354745 |
| 0.06360998 |
| 0.06374753 |
| 0.06387576 |
| 0.06393304 |
| 0.0640262 |
| 0.0640658 |
| 0.06415733 |
| 0.06421107 |
| 0.06426406 |
| 0.06426579 |
| 0.06454343 |
| 0.06486612 |
| 0.06491067 |
| 0.06493678 |
| 0.0650764 |
| 0.0651536 |
| 0.06516613 |
| 0.06519617 |
| 0.06534832 |
| 0.06536443 |
| 0.06568291 |
| 0.06571797 |
| 0.06571797 |
| 0.06587413 |
| 0.06592974 |
| 0.06617311 |
| 0.06617311 |
| 0.0661819 |
| 0.06631962 |
| 0.06648333 |
| 0.06664262 |
| 0.06670925 |
| 0.06676647 |
| 0.06691487 |
| 0.06737256 |

| |
|---|
| 0.06737256 |
| 0.06737256 |
| 0.06744022 |
| 0.06766431 |
| 0.0680749 |
| 0.06807949 |
| 0.06810479 |
| 0.06811579 |
| 0.06811579 |
| 0.06811579 |
| 0.06832547 |
| 0.06843221 |
| 0.06848964 |
| 0.06864078 |
| 0.06884667 |
| 0.06911101 |
| 0.06912591 |
| 0.06926202 |
| 0.06938183 |
| 0.06940902 |
| 0.06942305 |
| 0.06950036 |
| 0.06950036 |
| 0.06958239 |
| 0.06969486 |
| 0.06982587 |
| 0.06982587 |
| 0.07007418 |
| 0.0702892 |
| 0.0702892 |
| 0.07038094 |
| 0.07041661 |
| 0.0709895 |
| 0.07104417 |
| 0.07122097 |
| 0.07148939 |
| 0.07148939 |
| 0.07150049 |
| 0.07164256 |
| 0.07172302 |
| 0.07202971 |

0.07209296
0.0722186
0.07261409
0.07264536
0.07277221
0.07284227
0.07288768
0.07328116
0.07340306
0.07368394
0.07395207
0.07398155
0.07416419
0.07416419
0.07428796
0.07433776
0.07438654
0.0744745
0.07449181
0.07486793
0.07486793
0.07496143
0.0751797
0.07522896
0.07537308
0.07541141
0.07541141
0.07556994
0.07563017
0.07588032
0.07605059
0.0761306
0.0766899
0.07674093
0.07706701
0.07712885
0.07739877
0.07752067
0.07766151
0.07768787
0.07783556

| |
|---|
| 0.07798633 |
| 0.07798633 |
| 0.07807665 |
| 0.0783189 |
| 0.07838719 |
| 0.07841548 |
| 0.0784277 |
| 0.0784277 |
| 0.0784277 |
| 0.0784277 |
| 0.07878711 |
| 0.07887686 |
| 0.07894396 |
| 0.07900803 |
| 0.07900803 |
| 0.07903297 |
| 0.07910055 |
| 0.07915148 |
| 0.0792343 |
| 0.07951642 |
| 0.07954202 |
| 0.07957987 |
| 0.07979988 |
| 0.08046768 |
| 0.08065717 |
| 0.08071614 |
| 0.08072834 |
| 0.08089127 |
| 0.08114014 |
| 0.08135256 |
| 0.08135256 |
| 0.0815268 |
| 0.08152867 |
| 0.08155484 |
| 0.08159247 |
| 0.08159247 |
| 0.08160981 |
| 0.08183796 |
| 0.08183952 |
| 0.0822573 |
| 0.08240202 |

| |
|---|
| 0.08263768 |
| 0.08279618 |
| 0.08293855 |
| 0.08293855 |
| 0.08299475 |
| 0.08299475 |
| 0.08300787 |
| 0.08308903 |
| 0.08341525 |
| 0.08354775 |
| 0.08354775 |
| 0.08357192 |
| 0.08357192 |
| 0.08391783 |
| 0.08391783 |
| 0.08391783 |
| 0.08394903 |
| 0.08447644 |
| 0.08466605 |
| 0.08476281 |
| 0.08499595 |
| 0.0852251 |
| 0.08550737 |
| 0.08556306 |
| 0.08605377 |
| 0.08617036 |
| 0.08617036 |
| 0.08619847 |
| 0.08629966 |
| 0.08657767 |
| 0.08658805 |
| 0.08689338 |
| 0.08697305 |
| 0.08710586 |
| 0.08735575 |
| 0.08739927 |
| 0.08740632 |
| 0.08747864 |
| 0.08747864 |
| 0.08768947 |
| 0.08797415 |

| |
|---|
| 0.08802901 |
| 0.08808022 |
| 0.08813681 |
| 0.08813681 |
| 0.08822984 |
| 0.08846805 |
| 0.08846805 |
| 0.08859248 |
| 0.08888555 |
| 0.08898467 |
| 0.08918841 |
| 0.08919424 |
| 0.08932456 |
| 0.08932557 |
| 0.08933738 |
| 0.08933738 |
| 0.0895432 |
| 0.0895432 |
| 0.0895432 |
| 0.0895432 |
| 0.08966352 |
| 0.08976153 |
| 0.0897685 |
| 0.08980397 |
| 0.09018139 |
| 0.09019051 |
| 0.09024501 |
| 0.09033557 |
| 0.09053665 |
| 0.09062405 |
| 0.09068651 |
| 0.09074951 |
| 0.09074951 |
| 0.09089458 |
| 0.09116928 |
| 0.09125533 |
| 0.09129613 |
| 0.09132229 |
| 0.09133536 |
| 0.09156215 |
| 0.09178224 |

| |
|---|
| 0.09193385 |
| 0.09212361 |
| 0.09243182 |
| 0.09249439 |
| 0.09256341 |
| 0.09280207 |
| 0.09292538 |
| 0.09296023 |
| 0.09296023 |
| 0.09306145 |
| 0.09308142 |
| 0.09334379 |
| 0.09334379 |
| 0.09334379 |
| 0.09356657 |
| 0.09362439 |
| 0.09368229 |
| 0.0939537 |
| 0.0942029 |
| 0.09422433 |
| 0.09478015 |
| 0.09597266 |
| 0.09601459 |
| 0.09601459 |
| 0.09606868 |
| 0.0961337 |
| 0.09637025 |
| 0.09679474 |
| 0.09681265 |
| 0.09710091 |
| 0.09711299 |
| 0.09727853 |
| 0.09735395 |
| 0.09741929 |
| 0.09779785 |
| 0.0980211 |
| 0.0980211 |
| 0.0980211 |
| 0.09844256 |
| 0.0984663 |
| 0.09851236 |

| |
|---|
| 0.09852587 |
| 0.09852587 |
| 0.09864707 |
| 0.09867958 |
| 0.09877203 |
| 0.09882942 |
| 0.09882942 |
| 0.09882942 |
| 0.09884251 |
| 0.09902823 |
| 0.0991364 |
| 0.09920337 |
| 0.09920337 |
| 0.09920337 |
| 0.09920337 |
| 0.09920351 |
| 0.0992407 |
| 0.0992407 |
| 0.09944539 |
| 0.09955951 |
| 0.09955981 |
| 0.09988117 |
| 0.09990214 |
| 0.09990214 |
| 0.09990214 |
| 0.09996211 |
| 0.10047614 |
| 0.10069818 |
| 0.10091415 |
| 0.10102707 |
| 0.10164197 |
| 0.10218098 |
| 0.10220457 |
| 0.10229475 |
| 0.10244307 |
| 0.10249096 |
| 0.10260756 |
| 0.10311108 |
| 0.10335815 |
| 0.10339027 |
| 0.10339027 |

| |
|---|
| 0.10357546 |
| 0.10360012 |
| 0.10375959 |
| 0.10379156 |
| 0.10390618 |
| 0.10391526 |
| 0.10391526 |
| 0.10408036 |
| 0.10416388 |
| 0.10416388 |
| 0.10442553 |
| 0.1044751 |
| 0.10474788 |
| 0.10480248 |
| 0.10528599 |
| 0.1053923 |
| 0.10549469 |
| 0.10554416 |
| 0.10554416 |
| 0.1059344 |
| 0.10631431 |
| 0.10662926 |
| 0.10703976 |
| 0.1075645 |
| 0.1075645 |
| 0.10774708 |
| 0.10788452 |
| 0.10788452 |
| 0.10800103 |
| 0.10807201 |
| 0.10831906 |
| 0.10848506 |
| 0.10866259 |
| 0.10876112 |
| 0.10881422 |
| 0.10882672 |
| 0.10889524 |
| 0.10894669 |
| 0.10914612 |
| 0.10937867 |
| 0.11027874 |

| |
|---|
| 0.11034907 |
| 0.11067681 |
| 0.11108955 |
| 0.11115051 |
| 0.11136538 |
| 0.11158591 |
| 0.11189455 |
| 0.11191456 |
| 0.11191456 |
| 0.11192636 |
| 0.11192636 |
| 0.11198164 |
| 0.11199805 |
| 0.11199805 |
| 0.11207009 |
| 0.11220655 |
| 0.11220655 |
| 0.11220655 |
| 0.11220655 |
| 0.1122295 |
| 0.11235304 |
| 0.11248539 |
| 0.11255968 |
| 0.11256577 |
| 0.11269068 |
| 0.11286982 |
| 0.1128699 |
| 0.11326019 |
| 0.11342068 |
| 0.11350521 |
| 0.11374587 |
| 0.11374587 |
| 0.1139294 |
| 0.11416325 |
| 0.1147364 |
| 0.11490128 |
| 0.11504721 |
| 0.11504721 |
| 0.11504721 |
| 0.11545161 |
| 0.11557293 |

| |
|---|
| 0.11563095 |
| 0.11565743 |
| 0.11608031 |
| 0.11690196 |
| 0.11713887 |
| 0.11713887 |
| 0.1174749 |
| 0.11760012 |
| 0.11800575 |
| 0.11804806 |
| 0.11842563 |
| 0.11862405 |
| 0.11862786 |
| 0.11907289 |
| 0.11907289 |
| 0.11915527 |
| 0.11922438 |
| 0.11939577 |
| 0.11942965 |
| 0.11948001 |
| 0.11985046 |
| 0.12016033 |
| 0.12020206 |
| 0.12032239 |
| 0.12040489 |
| 0.12049094 |
| 0.12066437 |
| 0.12084043 |
| 0.12084043 |
| 0.12088771 |
| 0.12088771 |
| 0.12140444 |
| 0.12150529 |
| 0.12154281 |
| 0.12154281 |
| 0.12168996 |
| 0.12184444 |
| 0.12194379 |
| 0.12194379 |
| 0.121983 |
| 0.12208091 |

| |
|---|
| 0.122193 |
| 0.1224917 |
| 0.12293795 |
| 0.12293795 |
| 0.12297868 |
| 0.123038 |
| 0.12308149 |
| 0.12330675 |
| 0.12348116 |
| 0.12352073 |
| 0.12352073 |
| 0.12378063 |
| 0.12378063 |
| 0.12378103 |
| 0.12378476 |
| 0.12378476 |
| 0.12395384 |
| 0.12411408 |
| 0.12417507 |
| 0.12436973 |
| 0.1245403 |
| 0.12557083 |
| 0.12575628 |
| 0.12578602 |
| 0.12634443 |
| 0.12655103 |
| 0.12671395 |
| 0.12711685 |
| 0.12711685 |
| 0.12753989 |
| 0.12764802 |
| 0.12872788 |
| 0.12876777 |
| 0.12898618 |
| 0.1290548 |
| 0.12934976 |
| 0.12958077 |
| 0.1296642 |
| 0.13030392 |
| 0.13034945 |
| 0.13048088 |

| |
|---|
| 0.13050603 |
| 0.13052381 |
| 0.13055495 |
| 0.1308035 |
| 0.13093197 |
| 0.13106547 |
| 0.1310816 |
| 0.13111416 |
| 0.13111991 |
| 0.13126296 |
| 0.13131315 |
| 0.13141844 |
| 0.13150811 |
| 0.13197489 |
| 0.13197572 |
| 0.13203281 |
| 0.1323724 |
| 0.13267472 |
| 0.13275616 |
| 0.13294634 |
| 0.13314828 |
| 0.13345806 |
| 0.13347299 |
| 0.13364294 |
| 0.13390372 |
| 0.13426925 |
| 0.13444545 |
| 0.13507192 |
| 0.13516217 |
| 0.13525566 |
| 0.13547267 |
| 0.13555258 |
| 0.13555258 |
| 0.13577339 |
| 0.13580376 |
| 0.13580376 |
| 0.13595414 |
| 0.13600563 |
| 0.13602546 |
| 0.13615512 |
| 0.13646695 |

| |
|---|
| 0.13655978 |
| 0.13655978 |
| 0.13655978 |
| 0.13660733 |
| 0.13672077 |
| 0.13672077 |
| 0.13704044 |
| 0.13755974 |
| 0.13756857 |
| 0.13766473 |
| 0.13770687 |
| 0.13792228 |
| 0.1379649 |
| 0.13831085 |
| 0.13845975 |
| 0.13870874 |
| 0.13874846 |
| 0.13906214 |
| 0.13932433 |
| 0.13971837 |
| 0.13971837 |
| 0.13977115 |
| 0.13983801 |
| 0.1402413 |
| 0.1404677 |
| 0.14091904 |
| 0.14132231 |
| 0.14132917 |
| 0.14138591 |
| 0.14144821 |
| 0.14152183 |
| 0.1416462 |
| 0.1416462 |
| 0.14168083 |
| 0.14188054 |
| 0.14193376 |
| 0.14207462 |
| 0.14210936 |
| 0.14255514 |
| 0.14264283 |
| 0.14264978 |

| |
|---|
| 0.14295149 |
| 0.14310802 |
| 0.14310802 |
| 0.14332167 |
| 0.14416266 |
| 0.14416266 |
| 0.1444239 |
| 0.14449349 |
| 0.14459703 |
| 0.14496173 |
| 0.14498492 |
| 0.14511298 |
| 0.14514731 |
| 0.14618669 |
| 0.14627411 |
| 0.14628766 |
| 0.14684855 |
| 0.14685369 |
| 0.14685617 |
| 0.14685617 |
| 0.14701044 |
| 0.14716313 |
| 0.14716313 |
| 0.14727811 |
| 0.14747015 |
| 0.14759456 |
| 0.14841899 |
| 0.14855601 |
| 0.14881412 |
| 0.14884667 |
| 0.14884667 |
| 0.14889709 |
| 0.14889709 |
| 0.14898162 |
| 0.14918389 |
| 0.14950075 |
| 0.14952045 |
| 0.14954834 |
| 0.14956783 |
| 0.14993927 |
| 0.15011848 |

| |
|---|
| 0.15075146 |
| 0.15075146 |
| 0.1507719 |
| 0.15125757 |
| 0.15180841 |
| 0.15227836 |
| 0.15255811 |
| 0.1527059 |
| 0.15271143 |
| 0.15274596 |
| 0.15274596 |
| 0.15278013 |
| 0.15286095 |
| 0.15313296 |
| 0.15331575 |
| 0.15342775 |
| 0.15355415 |
| 0.15370066 |
| 0.15378432 |
| 0.15391974 |
| 0.15425695 |
| 0.15428907 |
| 0.15505622 |
| 0.15511271 |
| 0.15522914 |
| 0.15567346 |
| 0.15567346 |
| 0.15578446 |
| 0.15578446 |
| 0.15596566 |
| 0.15596566 |
| 0.15597439 |
| 0.15613375 |
| 0.15676787 |
| 0.15695659 |
| 0.15714218 |
| 0.15720791 |
| 0.15768143 |
| 0.15771902 |
| 0.1577495 |
| 0.15795923 |

0.15822163
0.15842792
0.15860849
0.15896903
0.15896903
0.15904462
0.159184
0.15918888
0.15918888
0.1593396
0.15962551
0.159921
0.16028504
0.16040518
0.16040518
0.16040518
0.16058512
0.16078938
0.16105402
0.16125135
0.16127622
0.1614796
0.16163465
0.16184857
0.1620295
0.16211534
0.16217624
0.16223296
0.16253685
0.16263557
0.16313005
0.16338762
0.16360299
0.1636651
0.16383409
0.16383409
0.1640349
0.16449324
0.16449324
0.16453835
0.16456073

| |
|---|
| 0.16461373 |
| 0.16491756 |
| 0.16544267 |
| 0.16546317 |
| 0.16546317 |
| 0.16546317 |
| 0.16546317 |
| 0.16546317 |
| 0.16546317 |
| 0.16546317 |
| 0.16546317 |
| 0.16546317 |
| 0.16548461 |
| 0.16562488 |
| 0.16563539 |
| 0.16566871 |
| 0.16572219 |
| 0.16572219 |
| 0.16604722 |
| 0.1666184 |
| 0.1666184 |
| 0.16703149 |
| 0.16722072 |
| 0.16722072 |
| 0.16733326 |
| 0.16806733 |
| 0.16807716 |
| 0.16873749 |
| 0.16873749 |
| 0.16873749 |
| 0.16891769 |
| 0.16932697 |
| 0.16939912 |
| 0.16941034 |
| 0.16947276 |
| 0.16962122 |
| 0.16973493 |
| 0.16979478 |
| 0.16987416 |
| 0.17031989 |
| 0.17031989 |

0.17047396
0.17053161
0.17053161
0.17059544
0.17078802
0.17085824
0.17085824
0.17085824
0.17146792
0.17193171
0.17201369
0.17201369
0.17202007
0.17265816
0.17267858
0.17275892
0.17275892
0.17275892
0.17282523
0.17291122
0.17304449
0.17316401
0.17316401
0.17326594
0.17335164
0.17362889
0.17364919
0.17364919
0.17387994
0.17387994
0.17390801
0.17394467
0.17403462
0.17447246
0.17529189
0.17562254
0.17568694
0.17583042
0.17630204
0.17631243
0.17642239

| |
|---|
| 0.17660816 |
| 0.17740964 |
| 0.17751389 |
| 0.17759935 |
| 0.17769474 |
| 0.17777575 |
| 0.17793881 |
| 0.17793881 |
| 0.17823853 |
| 0.17845581 |
| 0.17866646 |
| 0.17878461 |
| 0.17909411 |
| 0.17961553 |
| 0.17989603 |
| 0.18005286 |
| 0.18016952 |
| 0.18016952 |
| 0.18018458 |
| 0.18028868 |
| 0.18032507 |
| 0.18073149 |
| 0.18075602 |
| 0.18076264 |
| 0.18121183 |
| 0.18139408 |
| 0.18141681 |
| 0.18141681 |
| 0.18170936 |
| 0.1819862 |
| 0.18238854 |
| 0.1824009 |
| 0.18241631 |
| 0.18241631 |
| 0.18256863 |
| 0.18256863 |
| 0.18256863 |
| 0.18273566 |
| 0.18349521 |
| 0.18349521 |
| 0.18361732 |

| |
|---|
| 0.1836632 |
| 0.18376609 |
| 0.18376609 |
| 0.18377308 |
| 0.18418992 |
| 0.18486616 |
| 0.18531668 |
| 0.18531668 |
| 0.18531668 |
| 0.18566117 |
| 0.18566117 |
| 0.18599385 |
| 0.18604028 |
| 0.1861569 |
| 0.1866437 |
| 0.18686501 |
| 0.18701703 |
| 0.18719698 |
| 0.18737353 |
| 0.18740013 |
| 0.18740013 |
| 0.18772004 |
| 0.18808347 |
| 0.18808347 |
| 0.18809117 |
| 0.18854543 |
| 0.188689 |
| 0.18877958 |
| 0.18886539 |
| 0.18886539 |
| 0.18894178 |
| 0.18904353 |
| 0.1891151 |
| 0.18968937 |
| 0.1897152 |
| 0.18981383 |
| 0.1898529 |
| 0.19001436 |
| 0.19010021 |
| 0.19023387 |
| 0.19023387 |

| |
|---|
| 0.19023387 |
| 0.19023387 |
| 0.19125682 |
| 0.19209442 |
| 0.19291659 |
| 0.19291659 |
| 0.19304053 |
| 0.19338197 |
| 0.19391726 |
| 0.19408248 |
| 0.19408248 |
| 0.19413135 |
| 0.19435825 |
| 0.19439285 |
| 0.19452582 |
| 0.194754 |
| 0.19513846 |
| 0.19513846 |
| 0.19513846 |
| 0.19513846 |
| 0.1951441 |
| 0.19539677 |
| 0.19556423 |
| 0.19589951 |
| 0.19624325 |
| 0.1963336 |
| 0.19649975 |
| 0.1966444 |
| 0.19675997 |
| 0.19702725 |
| 0.19726986 |
| 0.19728154 |
| 0.19777363 |
| 0.19820518 |
| 0.19821474 |
| 0.1983038 |
| 0.19848808 |
| 0.19866495 |
| 0.19935609 |
| 0.19935609 |
| 0.19994948 |

| |
|---|
| 0.20004401 |
| 0.20020038 |
| 0.20020038 |
| 0.20020038 |
| 0.20055632 |
| 0.20092392 |
| 0.2009649 |
| 0.20115339 |
| 0.20121542 |
| 0.20148165 |
| 0.20153211 |
| 0.20175046 |
| 0.20175046 |
| 0.20184965 |
| 0.20186994 |
| 0.20235843 |
| 0.20260682 |
| 0.20274695 |
| 0.20289874 |
| 0.20291231 |
| 0.20303078 |
| 0.20305578 |
| 0.20305578 |
| 0.20312635 |
| 0.20344481 |
| 0.20357922 |
| 0.20373437 |
| 0.20374093 |
| 0.20376113 |
| 0.20387014 |
| 0.20418322 |
| 0.20439556 |
| 0.20451238 |
| 0.20473341 |
| 0.20503562 |
| 0.20511354 |
| 0.20511354 |
| 0.20609354 |
| 0.20646032 |
| 0.20672768 |
| 0.20738097 |

| |
|---|
| 0.20762408 |
| 0.20815159 |
| 0.20849267 |
| 0.20849267 |
| 0.20851235 |
| 0.20851235 |
| 0.20857025 |
| 0.20857025 |
| 0.20901907 |
| 0.20902312 |
| 0.20915426 |
| 0.20926913 |
| 0.20937232 |
| 0.20975337 |
| 0.20999903 |
| 0.21122797 |
| 0.21141459 |
| 0.21157357 |
| 0.21164832 |
| 0.21167468 |
| 0.21169212 |
| 0.21169634 |
| 0.2117761 |
| 0.2123296 |
| 0.21239346 |
| 0.21322053 |
| 0.21328938 |
| 0.21347196 |
| 0.21396446 |
| 0.21401744 |
| 0.21408443 |
| 0.21432896 |
| 0.21456828 |
| 0.21533133 |
| 0.21584462 |
| 0.21584611 |
| 0.21629842 |
| 0.21629842 |
| 0.21659192 |
| 0.21681835 |
| 0.21695903 |

| |
|---|
| 0.21697773 |
| 0.21697773 |
| 0.21724602 |
| 0.21737637 |
| 0.2175344 |
| 0.21768287 |
| 0.21776867 |
| 0.21831206 |
| 0.21872056 |
| 0.21886117 |
| 0.21886117 |
| 0.21886117 |
| 0.2189433 |
| 0.21897544 |
| 0.21919335 |
| 0.21948088 |
| 0.21948088 |
| 0.21983169 |
| 0.21993807 |
| 0.22015389 |
| 0.22021709 |
| 0.22154932 |
| 0.22154932 |
| 0.22221644 |
| 0.22227439 |
| 0.2223288 |
| 0.22266361 |
| 0.22293023 |
| 0.22327102 |
| 0.22327102 |
| 0.22339866 |
| 0.22347391 |
| 0.22347391 |
| 0.22475898 |
| 0.22476727 |
| 0.22476727 |
| 0.22504114 |
| 0.22536971 |
| 0.2254667 |
| 0.2256383 |
| 0.22600309 |

| |
|---|
| 0.22626318 |
| 0.22646234 |
| 0.22650899 |
| 0.22746208 |
| 0.22763592 |
| 0.22763592 |
| 0.22763592 |
| 0.22850193 |
| 0.22867166 |
| 0.22886232 |
| 0.2293073 |
| 0.23021332 |
| 0.23025656 |
| 0.23051958 |
| 0.23077684 |
| 0.23114895 |
| 0.23128168 |
| 0.23143293 |
| 0.232076 |
| 0.232076 |
| 0.2321166 |
| 0.23230683 |
| 0.23311852 |
| 0.23381927 |
| 0.23381927 |
| 0.23381927 |
| 0.23384257 |
| 0.23422932 |
| 0.2343123 |
| 0.23432159 |
| 0.23434775 |
| 0.2344002 |
| 0.2344002 |
| 0.23486419 |
| 0.23488173 |
| 0.23488173 |
| 0.23491986 |
| 0.23491986 |
| 0.23491986 |
| 0.23506031 |
| 0.23524193 |

| |
|---|
| 0.23524193 |
| 0.23531896 |
| 0.23573049 |
| 0.23575722 |
| 0.23610682 |
| 0.23647025 |
| 0.23660721 |
| 0.236618 |
| 0.23670377 |
| 0.23716629 |
| 0.23764193 |
| 0.2378799 |
| 0.237964 |
| 0.23804255 |
| 0.23929867 |
| 0.23929867 |
| 0.23947645 |
| 0.24001502 |
| 0.24006192 |
| 0.24039566 |
| 0.24040078 |
| 0.24043993 |
| 0.2404952 |
| 0.24054896 |
| 0.24130847 |
| 0.24135207 |
| 0.24216815 |
| 0.24221265 |
| 0.24269141 |
| 0.24335368 |
| 0.24357668 |
| 0.24406992 |
| 0.2441423 |
| 0.2441423 |
| 0.24436083 |
| 0.24436083 |
| 0.24439184 |
| 0.24444788 |
| 0.24461973 |
| 0.24461973 |
| 0.24462921 |

| |
|---|
| 0.24464356 |
| 0.24464356 |
| 0.24488745 |
| 0.2449096 |
| 0.24553104 |
| 0.24568672 |
| 0.24568672 |
| 0.24568672 |
| 0.24568672 |
| 0.24570791 |
| 0.24573301 |
| 0.24587799 |
| 0.24587799 |
| 0.24587799 |
| 0.24605636 |
| 0.24633504 |
| 0.24655925 |
| 0.24668288 |
| 0.24674944 |
| 0.24674944 |
| 0.24688472 |
| 0.24698136 |
| 0.24733412 |
| 0.24749317 |
| 0.2476296 |
| 0.24803449 |
| 0.24803449 |
| 0.24822896 |
| 0.24846524 |
| 0.24859033 |
| 0.24873337 |
| 0.24889005 |
| 0.24944141 |
| 0.24956475 |
| 0.24961555 |
| 0.24984934 |
| 0.2500594 |
| 0.25034943 |
| 0.2506929 |
| 0.25109956 |
| 0.25109956 |

| |
|---|
| 0.2511478 |
| 0.25116057 |
| 0.25126119 |
| 0.2513571 |
| 0.25160888 |
| 0.25165374 |
| 0.25168926 |
| 0.25190308 |
| 0.25200102 |
| 0.25223202 |
| 0.25265655 |
| 0.25266794 |
| 0.25293894 |
| 0.25347328 |
| 0.25366162 |
| 0.25376331 |
| 0.25379713 |
| 0.25397818 |
| 0.25427753 |
| 0.25439553 |
| 0.25470327 |
| 0.25477098 |
| 0.25480002 |
| 0.25506599 |
| 0.25565477 |
| 0.25572856 |
| 0.25580569 |
| 0.25603836 |
| 0.25690042 |
| 0.25690042 |
| 0.25692244 |
| 0.25718889 |
| 0.25727731 |
| 0.25734136 |
| 0.25751024 |
| 0.25786896 |
| 0.25786896 |
| 0.25828369 |
| 0.25863909 |
| 0.25960182 |
| 0.25970392 |

| |
|---|
| 0.25996166 |
| 0.25996166 |
| 0.26080826 |
| 0.26134555 |
| 0.2615588 |
| 0.26157097 |
| 0.26203475 |
| 0.26207583 |
| 0.26226603 |
| 0.26248314 |
| 0.2627723 |
| 0.26324542 |
| 0.26329565 |
| 0.26329565 |
| 0.26353408 |
| 0.26358973 |
| 0.2637229 |
| 0.2639285 |
| 0.26473292 |
| 0.26473292 |
| 0.26501948 |
| 0.2652428 |
| 0.26533727 |
| 0.26536838 |
| 0.26572656 |
| 0.26622775 |
| 0.2663884 |
| 0.26646462 |
| 0.2667599 |
| 0.2667599 |
| 0.2667599 |
| 0.2667599 |
| 0.2667599 |
| 0.26702643 |
| 0.26788823 |
| 0.26791426 |
| 0.26791426 |
| 0.2679374 |
| 0.26823351 |
| 0.26872207 |
| 0.26882721 |

| |
|---|
| 0.26889664 |
| 0.26927607 |
| 0.26935035 |
| 0.26984297 |
| 0.26986435 |
| 0.26995728 |
| 0.27002313 |
| 0.27002313 |
| 0.27002313 |
| 0.27051235 |
| 0.27106707 |
| 0.27123241 |
| 0.27127193 |
| 0.27139801 |
| 0.27140115 |
| 0.27160086 |
| 0.27170993 |
| 0.27189997 |
| 0.27195828 |
| 0.27224247 |
| 0.27237673 |
| 0.27271784 |
| 0.27276169 |
| 0.27284716 |
| 0.27284716 |
| 0.27345369 |
| 0.27353765 |
| 0.27353765 |
| 0.27396716 |
| 0.27402608 |
| 0.27402608 |
| 0.27417344 |
| 0.27481052 |
| 0.27504914 |
| 0.27504914 |
| 0.27531945 |
| 0.27543441 |
| 0.27543441 |
| 0.27562341 |
| 0.27562341 |
| 0.27562341 |

| |
|---|
| 0.27562341 |
| 0.27606338 |
| 0.27653177 |
| 0.27678252 |
| 0.27716014 |
| 0.27716014 |
| 0.27741088 |
| 0.27825795 |
| 0.27845662 |
| 0.27892044 |
| 0.27940442 |
| 0.27940442 |
| 0.27957765 |
| 0.27971024 |
| 0.28006552 |
| 0.28016551 |
| 0.28017088 |
| 0.28019545 |
| 0.28033746 |
| 0.28089829 |
| 0.28120427 |
| 0.28126591 |
| 0.28127547 |
| 0.28247931 |
| 0.28329574 |
| 0.28370828 |
| 0.28391216 |
| 0.28391216 |
| 0.28391216 |
| 0.28403389 |
| 0.28403389 |
| 0.28421735 |
| 0.28514024 |
| 0.28583724 |
| 0.28583724 |
| 0.2861337 |
| 0.28616049 |
| 0.28623683 |
| 0.28638035 |
| 0.28668806 |
| 0.28716535 |

| |
|---|
| 0.28753617 |
| 0.28753897 |
| 0.28772533 |
| 0.28772533 |
| 0.28814352 |
| 0.28814352 |
| 0.28834714 |
| 0.28911239 |
| 0.28911239 |
| 0.28933937 |
| 0.2894218 |
| 0.2894218 |
| 0.28949586 |
| 0.29033431 |
| 0.2904776 |
| 0.29048596 |
| 0.29048596 |
| 0.29080786 |
| 0.29096703 |
| 0.29096703 |
| 0.29096703 |
| 0.29104283 |
| 0.29122805 |
| 0.29125094 |
| 0.29139874 |
| 0.29164896 |
| 0.29197489 |
| 0.29206884 |
| 0.29222474 |
| 0.29249115 |
| 0.2928613 |
| 0.29366982 |
| 0.29448947 |
| 0.29451449 |
| 0.29461519 |
| 0.29461519 |
| 0.29467773 |
| 0.29530751 |
| 0.29567384 |
| 0.2965029 |
| 0.29651 |

| |
|---|
| 0.29669997 |
| 0.29672599 |
| 0.29679124 |
| 0.29738915 |
| 0.29749325 |
| 0.29769353 |
| 0.29782344 |
| 0.29782344 |
| 0.29830198 |
| 0.29836932 |
| 0.2986498 |
| 0.29886504 |
| 0.29921025 |
| 0.29946003 |
| 0.29981759 |
| 0.29989161 |
| 0.29991367 |
| 0.29996277 |
| 0.30020507 |
| 0.30032025 |
| 0.30078641 |
| 0.30129238 |
| 0.30129238 |
| 0.30153852 |
| 0.30158228 |
| 0.30172363 |
| 0.30225806 |
| 0.30252805 |
| 0.30252805 |
| 0.30302639 |
| 0.30321078 |
| 0.30370119 |
| 0.30401855 |
| 0.30447333 |
| 0.30454362 |
| 0.30466228 |
| 0.30500818 |
| 0.30508384 |
| 0.30596356 |
| 0.30625933 |
| 0.30626148 |

| |
|---|
| 0.30646178 |
| 0.30729759 |
| 0.30772309 |
| 0.30772309 |
| 0.30772309 |
| 0.30796608 |
| 0.30796608 |
| 0.30818057 |
| 0.3082148 |
| 0.30886103 |
| 0.30932488 |
| 0.30948156 |
| 0.30949488 |
| 0.30970439 |
| 0.30982485 |
| 0.31013624 |
| 0.31015862 |
| 0.31015862 |
| 0.31020196 |
| 0.31020196 |
| 0.31051989 |
| 0.31051989 |
| 0.31054038 |
| 0.31054038 |
| 0.31054038 |
| 0.31093151 |
| 0.3111308 |
| 0.31123081 |
| 0.31133694 |
| 0.31162195 |
| 0.3121383 |
| 0.31215763 |
| 0.31220191 |
| 0.31220191 |
| 0.31284371 |
| 0.31284371 |
| 0.31286608 |
| 0.31293594 |
| 0.31293594 |
| 0.31293594 |
| 0.31312069 |

| |
|---|
| 0.31337254 |
| 0.31378591 |
| 0.31386668 |
| 0.31430188 |
| 0.31430188 |
| 0.31430188 |
| 0.31441541 |
| 0.31525155 |
| 0.3154954 |
| 0.31586655 |
| 0.31591982 |
| 0.31594301 |
| 0.31594301 |
| 0.31594301 |
| 0.31606684 |
| 0.31606684 |
| 0.31608951 |
| 0.31661502 |
| 0.31663852 |
| 0.31663852 |
| 0.31679344 |
| 0.31681104 |
| 0.31701265 |
| 0.31710695 |
| 0.31738871 |
| 0.3174474 |
| 0.31783266 |
| 0.31813925 |
| 0.31850652 |
| 0.31908057 |
| 0.3192807 |
| 0.3194416 |
| 0.31991623 |
| 0.31995743 |
| 0.32001024 |
| 0.32033308 |
| 0.32036439 |
| 0.32117059 |
| 0.32122443 |
| 0.32191855 |
| 0.32199173 |

| |
|---|
| 0.32215605 |
| 0.32218489 |
| 0.32218489 |
| 0.32218489 |
| 0.32242478 |
| 0.32254197 |
| 0.3226208 |
| 0.3226208 |
| 0.3226837 |
| 0.3226837 |
| 0.32269599 |
| 0.32297378 |
| 0.32330373 |
| 0.3236261 |
| 0.32402247 |
| 0.32430523 |
| 0.32430523 |
| 0.32488114 |
| 0.32541123 |
| 0.32545001 |
| 0.3256562 |
| 0.32583811 |
| 0.32675057 |
| 0.32679347 |
| 0.32679347 |
| 0.32681212 |
| 0.32693728 |
| 0.32725627 |
| 0.32728266 |
| 0.32767897 |
| 0.32778512 |
| 0.32811564 |
| 0.32855293 |
| 0.32870465 |
| 0.32877512 |
| 0.32967955 |
| 0.32967955 |
| 0.33039802 |
| 0.33060122 |
| 0.330603 |
| 0.33147295 |

| |
|---|
| 0.33152541 |
| 0.33176994 |
| 0.33207498 |
| 0.33222203 |
| 0.33325787 |
| 0.33359728 |
| 0.33410059 |
| 0.33470475 |
| 0.33475603 |
| 0.33500405 |
| 0.33519073 |
| 0.33588708 |
| 0.3370975 |
| 0.33715304 |
| 0.33790032 |
| 0.33797648 |
| 0.33799847 |
| 0.33835902 |
| 0.33837704 |
| 0.33872627 |
| 0.33872627 |
| 0.33877113 |
| 0.33877295 |
| 0.33920249 |
| 0.33956238 |
| 0.34003925 |
| 0.34032309 |
| 0.340377 |
| 0.34086787 |
| 0.34086787 |
| 0.34252933 |
| 0.34269265 |
| 0.34361727 |
| 0.34378221 |
| 0.34378221 |
| 0.34378221 |
| 0.34378221 |
| 0.34378221 |
| 0.3441083 |
| 0.34431489 |
| 0.34431489 |

| |
|---|
| 0.34431489 |
| 0.34435844 |
| 0.34436835 |
| 0.3445073 |
| 0.34472404 |
| 0.34496857 |
| 0.34496857 |
| 0.34507994 |
| 0.34507994 |
| 0.34507994 |
| 0.34514681 |
| 0.3456172 |
| 0.34564607 |
| 0.3460379 |
| 0.34653166 |
| 0.34659972 |
| 0.34674532 |
| 0.34717447 |
| 0.34717447 |
| 0.34717447 |
| 0.34742038 |
| 0.34778885 |
| 0.34854099 |
| 0.34864199 |
| 0.34864199 |
| 0.34882758 |
| 0.3490226 |
| 0.34944822 |
| 0.34957857 |
| 0.34979597 |
| 0.35112767 |
| 0.35118793 |
| 0.35130082 |
| 0.35140843 |
| 0.35196984 |
| 0.35204465 |
| 0.3524362 |
| 0.35251477 |
| 0.35278165 |
| 0.35336479 |
| 0.35421137 |

| |
|---|
| 0.35424297 |
| 0.35513881 |
| 0.35517639 |
| 0.35517639 |
| 0.35517639 |
| 0.35526907 |
| 0.35562365 |
| 0.35626516 |
| 0.35626516 |
| 0.35636172 |
| 0.35636172 |
| 0.35636172 |
| 0.3564861 |
| 0.35658646 |
| 0.35699256 |
| 0.35825506 |
| 0.35825506 |
| 0.35828898 |
| 0.35831334 |
| 0.35877285 |
| 0.3607496 |
| 0.36118605 |
| 0.3619868 |
| 0.36231759 |
| 0.36231759 |
| 0.36259667 |
| 0.36272335 |
| 0.36285572 |
| 0.36349221 |
| 0.36349895 |
| 0.36399557 |
| 0.36399618 |
| 0.36437054 |
| 0.36522452 |
| 0.36607049 |
| 0.3661193 |
| 0.36618067 |
| 0.36654072 |
| 0.36654072 |
| 0.36654072 |
| 0.36659832 |

| |
|---|
| 0.36659832 |
| 0.36659832 |
| 0.36659832 |
| 0.36659832 |
| 0.36689164 |
| 0.3668985 |
| 0.36699149 |
| 0.36722505 |
| 0.3674157 |
| 0.36775709 |
| 0.36831073 |
| 0.36895315 |
| 0.36948949 |
| 0.36966624 |
| 0.37027876 |
| 0.37040678 |
| 0.37105934 |
| 0.37127859 |
| 0.37207595 |
| 0.37207595 |
| 0.37207595 |
| 0.37222984 |
| 0.37242017 |
| 0.37242017 |
| 0.37265226 |
| 0.37265226 |
| 0.37267835 |
| 0.37267835 |
| 0.37305343 |
| 0.3736519 |
| 0.37366693 |
| 0.37366693 |
| 0.37468905 |
| 0.37488781 |
| 0.37488842 |
| 0.37497772 |
| 0.37503799 |
| 0.37521152 |
| 0.37521152 |
| 0.37555842 |
| 0.37567051 |

| |
|---|
| 0.37600277 |
| 0.37600277 |
| 0.37600277 |
| 0.37641416 |
| 0.37641416 |
| 0.37694177 |
| 0.37738036 |
| 0.37781173 |
| 0.37821025 |
| 0.37824154 |
| 0.37843675 |
| 0.37883471 |
| 0.37930563 |
| 0.37949014 |
| 0.37949014 |
| 0.37949014 |
| 0.37986646 |
| 0.38002346 |
| 0.38013748 |
| 0.38085444 |
| 0.38124319 |
| 0.38145035 |
| 0.38175184 |
| 0.38175184 |
| 0.38201891 |
| 0.38238881 |
| 0.38262448 |
| 0.38270691 |
| 0.3828036 |
| 0.38336816 |
| 0.38343662 |
| 0.38372178 |
| 0.38393133 |
| 0.38394812 |
| 0.38396711 |
| 0.38432791 |
| 0.38432791 |
| 0.38467729 |
| 0.38467729 |
| 0.38475418 |
| 0.38510906 |

| |
|---|
| 0.38589747 |
| 0.38589747 |
| 0.38589747 |
| 0.38589747 |
| 0.38606616 |
| 0.38624994 |
| 0.38646129 |
| 0.38668152 |
| 0.38669995 |
| 0.38686577 |
| 0.38686577 |
| 0.38708056 |
| 0.38776506 |
| 0.38822017 |
| 0.38890238 |
| 0.38890238 |
| 0.38919553 |
| 0.3896099 |
| 0.38974392 |
| 0.38994502 |
| 0.39016491 |
| 0.39016491 |
| 0.39016491 |
| 0.3902758 |
| 0.39116065 |
| 0.39117156 |
| 0.39117156 |
| 0.39184615 |
| 0.39261818 |
| 0.39261818 |
| 0.39273898 |
| 0.39311357 |
| 0.39311357 |
| 0.39312487 |
| 0.39329701 |
| 0.39427681 |
| 0.39435409 |
| 0.39487522 |
| 0.39487522 |
| 0.39495993 |
| 0.39498142 |

0.39498142
0.39498142
0.39554255
0.39595902
0.39621975
0.39625199
0.39644101
0.39719551
0.39720764
0.39736807
0.39842747
0.39842747
0.39899918
0.39908741
0.39915745
0.39926298
0.39959648
0.40006208
0.40012498
0.40024332
0.40073546
0.40078972
0.40105365
0.40173372
0.40228948
0.40261165
0.40261165
0.40301903
0.40312765
0.40396599
0.40399653
0.40403497
0.40405512
0.40426432
0.40426432
0.40426432
0.40426432
0.40461254
0.40461254
0.40465134
0.40465134

| |
|---|
| 0.40494724 |
| 0.40521737 |
| 0.40539493 |
| 0.40559182 |
| 0.4056757 |
| 0.40601531 |
| 0.40601531 |
| 0.40667958 |
| 0.4067958 |
| 0.40693559 |
| 0.40693559 |
| 0.40693559 |
| 0.40693559 |
| 0.40705304 |
| 0.40710354 |
| 0.4073131 |
| 0.40750898 |
| 0.40835819 |
| 0.40858405 |
| 0.40891024 |
| 0.40920765 |
| 0.40931653 |
| 0.40993742 |
| 0.41010561 |
| 0.41015767 |
| 0.41030592 |
| 0.41041143 |
| 0.41041311 |
| 0.41056172 |
| 0.41067772 |
| 0.41067772 |
| 0.41166939 |
| 0.41166939 |
| 0.41202329 |
| 0.41224871 |
| 0.41344925 |
| 0.41488142 |
| 0.41531945 |
| 0.41589115 |
| 0.41589115 |
| 0.41630649 |

| |
|---|
| 0.41649587 |
| 0.41653356 |
| 0.41689914 |
| 0.41720432 |
| 0.41777435 |
| 0.41777435 |
| 0.41796314 |
| 0.41796314 |
| 0.41798951 |
| 0.41803743 |
| 0.41804187 |
| 0.41804997 |
| 0.41869052 |
| 0.41886453 |
| 0.41917945 |
| 0.41937217 |
| 0.41944728 |
| 0.41989487 |
| 0.42032801 |
| 0.42033735 |
| 0.42033735 |
| 0.42033735 |
| 0.42106244 |
| 0.4210732 |
| 0.42117209 |
| 0.42158922 |
| 0.4221268 |
| 0.42232025 |
| 0.42267037 |
| 0.42270719 |
| 0.42307485 |
| 0.423416 |
| 0.42379732 |
| 0.42467647 |
| 0.42523524 |
| 0.42523524 |
| 0.42523524 |
| 0.42545674 |
| 0.42566393 |
| 0.42568665 |
| 0.42599617 |

| |
|---|
| 0.42611629 |
| 0.4263016 |
| 0.42655039 |
| 0.4265629 |
| 0.42680291 |
| 0.42680291 |
| 0.42717604 |
| 0.42765594 |
| 0.42765594 |
| 0.42843337 |
| 0.42855004 |
| 0.42855004 |
| 0.42926839 |
| 0.42972888 |
| 0.42979865 |
| 0.43003132 |
| 0.43068167 |
| 0.43077964 |
| 0.43127139 |
| 0.43127139 |
| 0.43143435 |
| 0.43174237 |
| 0.43228069 |
| 0.43265525 |
| 0.43265525 |
| 0.43265525 |
| 0.43293704 |
| 0.43294736 |
| 0.43304751 |
| 0.43384857 |
| 0.43394678 |
| 0.43437431 |
| 0.43458809 |
| 0.43598949 |
| 0.43598949 |
| 0.43614382 |
| 0.4365767 |
| 0.43659354 |
| 0.43659354 |
| 0.43660228 |
| 0.43660228 |

0.43665593
0.43680181
0.43682652
0.43683561
0.43683561
0.43683561
0.43683561
0.43683561
0.43688417
0.43732942
0.43756163
0.43794976
0.43803791
0.43819012
0.43827975
0.43848602
0.43964283
0.43976214
0.43977187
0.43987129
0.44023037
0.44027068
0.44187013
0.44193221
0.44247432
0.44289158
0.44297155
0.44318416
0.44320698
0.44375577
0.44396358
0.44396358
0.44396358
0.44433593
0.44448117
0.44460519
0.44462568
0.44475261
0.44475261
0.44553288
0.44574792

| |
|---|
| 0.44627655 |
| 0.44635018 |
| 0.44653486 |
| 0.44670054 |
| 0.44697031 |
| 0.44818178 |
| 0.44818178 |
| 0.44861218 |
| 0.44881566 |
| 0.44933812 |
| 0.44948742 |
| 0.4502028 |
| 0.45048612 |
| 0.45048612 |
| 0.45048612 |
| 0.45048612 |
| 0.45048612 |
| 0.45204588 |
| 0.45209579 |
| 0.45209619 |
| 0.45234195 |
| 0.4529286 |
| 0.45321582 |
| 0.45341285 |
| 0.45347369 |
| 0.45347369 |
| 0.45381671 |
| 0.45381671 |
| 0.45423182 |
| 0.45439901 |
| 0.45545738 |
| 0.45592658 |
| 0.45598889 |
| 0.45644041 |
| 0.45650301 |
| 0.45666068 |
| 0.45666068 |
| 0.45688074 |
| 0.45711597 |
| 0.45711597 |
| 0.45751897 |

| |
|---|
| 0.45751897 |
| 0.45759177 |
| 0.45766092 |
| 0.45788302 |
| 0.45803621 |
| 0.45809021 |
| 0.45809021 |
| 0.45858111 |
| 0.4588427 |
| 0.45957172 |
| 0.46074699 |
| 0.46135145 |
| 0.46203692 |
| 0.46234643 |
| 0.46260181 |
| 0.46269582 |
| 0.46269582 |
| 0.46275365 |
| 0.46281399 |
| 0.46288027 |
| 0.46320691 |
| 0.46367055 |
| 0.46377191 |
| 0.46517248 |
| 0.46517248 |
| 0.46517248 |
| 0.46529927 |
| 0.46539235 |
| 0.46578852 |
| 0.46596811 |
| 0.46642881 |
| 0.46669226 |
| 0.46698747 |
| 0.46737153 |
| 0.46737153 |
| 0.46758268 |
| 0.46759571 |
| 0.46812723 |
| 0.46829664 |
| 0.46829664 |
| 0.46831748 |

| |
|---|
| 0.46887005 |
| 0.46902284 |
| 0.46964161 |
| 0.46974142 |
| 0.47009259 |
| 0.47037883 |
| 0.47077678 |
| 0.47077678 |
| 0.47197197 |
| 0.47303274 |
| 0.47308786 |
| 0.47308786 |
| 0.47326696 |
| 0.4732876 |
| 0.47356396 |
| 0.47356419 |
| 0.47362676 |
| 0.47397996 |
| 0.47401558 |
| 0.47401558 |
| 0.47457303 |
| 0.47529014 |
| 0.47552809 |
| 0.47568003 |
| 0.47568003 |
| 0.47568003 |
| 0.47573576 |
| 0.47585278 |
| 0.47635531 |
| 0.47655071 |
| 0.47724105 |
| 0.47742012 |
| 0.47779675 |
| 0.47783815 |
| 0.47801074 |
| 0.47812547 |
| 0.47848161 |
| 0.4787002 |
| 0.4787078 |
| 0.47882063 |
| 0.47948859 |

| |
|---|
| 0.47963307 |
| 0.47964708 |
| 0.47964708 |
| 0.47964708 |
| 0.47964708 |
| 0.4797287 |
| 0.4799266 |
| 0.48065145 |
| 0.4807251 |
| 0.48078864 |
| 0.48160446 |
| 0.48199125 |
| 0.48267228 |
| 0.48313029 |
| 0.48313029 |
| 0.48363218 |
| 0.48363643 |
| 0.48363643 |
| 0.48456706 |
| 0.48471493 |
| 0.48541945 |
| 0.48541945 |
| 0.48561435 |
| 0.48567434 |
| 0.48567434 |
| 0.48581416 |
| 0.48581548 |
| 0.4861541 |
| 0.48634223 |
| 0.48639359 |
| 0.48709181 |
| 0.48709181 |
| 0.48730794 |
| 0.4874435 |
| 0.4874435 |
| 0.4874435 |
| 0.4874435 |
| 0.48820724 |
| 0.48841224 |
| 0.48899602 |
| 0.48899602 |

| |
|---|
| 0.49035513 |
| 0.49261675 |
| 0.49301991 |
| 0.49347181 |
| 0.49375427 |
| 0.494102 |
| 0.49415995 |
| 0.49466409 |
| 0.49531858 |
| 0.49589693 |
| 0.49709139 |
| 0.49716651 |
| 0.49716651 |
| 0.49730829 |
| 0.49730829 |
| 0.49730829 |
| 0.49744358 |
| 0.4974558 |
| 0.49772784 |
| 0.49777613 |
| 0.49815482 |
| 0.4982428 |
| 0.49860824 |
| 0.49890328 |
| 0.49931476 |
| 0.49965657 |
| 0.49977532 |
| 0.49982584 |
| 0.50008466 |
| 0.50010097 |
| 0.50053519 |
| 0.50102463 |
| 0.50166547 |
| 0.50176405 |
| 0.50185419 |
| 0.5022809 |
| 0.50229929 |
| 0.50234153 |
| 0.50240281 |
| 0.50240281 |
| 0.5024691 |

| |
|---|
| 0.50284766 |
| 0.50295048 |
| 0.50295048 |
| 0.50295048 |
| 0.50295755 |
| 0.50305635 |
| 0.50335397 |
| 0.5035174 |
| 0.50363716 |
| 0.50434734 |
| 0.5047136 |
| 0.5047208 |
| 0.5047591 |
| 0.50499546 |
| 0.50513219 |
| 0.50513219 |
| 0.50533383 |
| 0.50579404 |
| 0.50609191 |
| 0.50617005 |
| 0.50633861 |
| 0.5068491 |
| 0.50689592 |
| 0.50692262 |
| 0.50708455 |
| 0.50719563 |
| 0.50731441 |
| 0.50737469 |
| 0.50771043 |
| 0.50771043 |
| 0.50950441 |
| 0.50959607 |
| 0.50959697 |
| 0.50959697 |
| 0.51002627 |
| 0.510231 |
| 0.5103396 |
| 0.51047695 |
| 0.51052045 |
| 0.51124421 |
| 0.51174779 |

| |
|---|
| 0.51223984 |
| 0.51236071 |
| 0.51293809 |
| 0.51297075 |
| 0.51314291 |
| 0.51314291 |
| 0.51414072 |
| 0.51414072 |
| 0.51449671 |
| 0.51462503 |
| 0.51515091 |
| 0.51516311 |
| 0.51608851 |
| 0.5163337 |
| 0.5163337 |
| 0.51662816 |
| 0.51706101 |
| 0.51790818 |
| 0.51821153 |
| 0.51821153 |
| 0.51821153 |
| 0.51845599 |
| 0.51845599 |
| 0.51857437 |
| 0.51885657 |
| 0.51885657 |
| 0.51984112 |
| 0.51984112 |
| 0.51991967 |
| 0.52033636 |
| 0.52033636 |
| 0.52033636 |
| 0.52046296 |
| 0.52046296 |
| 0.52046296 |
| 0.52046296 |
| 0.52046296 |
| 0.52083171 |
| 0.5210726 |
| 0.52160163 |
| 0.52160163 |

| |
|---|
| 0.52165255 |
| 0.52166205 |
| 0.52231453 |
| 0.52235279 |
| 0.52262054 |
| 0.52346626 |
| 0.52346626 |
| 0.52346626 |
| 0.52361371 |
| 0.52389349 |
| 0.52389349 |
| 0.52389349 |
| 0.52389349 |
| 0.52389349 |
| 0.52389349 |
| 0.52412299 |
| 0.52429368 |
| 0.52445947 |
| 0.52515662 |
| 0.52524473 |
| 0.52568257 |
| 0.52568257 |
| 0.52586361 |
| 0.52591644 |
| 0.52668758 |
| 0.52726086 |
| 0.52740525 |
| 0.52927967 |
| 0.52951222 |
| 0.52969456 |
| 0.53025978 |
| 0.53040857 |
| 0.53081619 |
| 0.53120042 |
| 0.53143539 |
| 0.53149555 |
| 0.53221071 |
| 0.53267463 |
| 0.53267463 |
| 0.53267463 |
| 0.53299147 |

| |
|---|
| 0.53299147 |
| 0.53299147 |
| 0.53299147 |
| 0.53299147 |
| 0.53344734 |
| 0.53344734 |
| 0.53431421 |
| 0.53527285 |
| 0.53562846 |
| 0.53570074 |
| 0.53581981 |
| 0.53588183 |
| 0.53591921 |
| 0.53609452 |
| 0.53623417 |
| 0.53682543 |
| 0.53693591 |
| 0.53695355 |
| 0.53714621 |
| 0.53719898 |
| 0.53730678 |
| 0.53730678 |
| 0.53730678 |
| 0.53745186 |
| 0.53790074 |
| 0.53791752 |
| 0.53791752 |
| 0.53834994 |
| 0.53834994 |
| 0.53843456 |
| 0.53843456 |
| 0.53845823 |
| 0.53845823 |
| 0.53886333 |
| 0.53886333 |
| 0.53886333 |
| 0.53886333 |
| 0.53889588 |
| 0.53889588 |
| 0.53906219 |
| 0.53927659 |

0.53930127
0.53994526
0.53994526
0.54009342
0.54009342
0.54036427
0.54113486
0.54161756
0.54165682
0.54229562
0.5425678
0.54257922
0.54263056
0.54263056
0.54317543
0.54330754
0.5442545
0.5445568
0.54536547
0.54536547
0.54536547
0.54588126
0.54670191
0.54671786
0.54700375
0.54706042
0.54708754
0.5478463
0.5479268
0.5479268
0.5479335
0.54861749
0.54896107
0.54898533
0.54910914
0.5494235
0.54958038
0.54958038
0.54965813
0.55031808
0.5503588

| |
|---|
| 0.55107119 |
| 0.55151067 |
| 0.55226275 |
| 0.5523496 |
| 0.55310056 |
| 0.55390876 |
| 0.5541827 |
| 0.55429471 |
| 0.55452774 |
| 0.55478713 |
| 0.55507956 |
| 0.55510767 |
| 0.55590823 |
| 0.55634107 |
| 0.55747332 |
| 0.55801898 |
| 0.55819371 |
| 0.55819495 |
| 0.55820932 |
| 0.55820932 |
| 0.55820932 |
| 0.5582178 |
| 0.55826829 |
| 0.55826829 |
| 0.55826829 |
| 0.55826829 |
| 0.55827632 |
| 0.55848935 |
| 0.55914144 |
| 0.55922806 |
| 0.55945097 |
| 0.55973722 |
| 0.55973722 |
| 0.55973722 |
| 0.55973722 |
| 0.55973722 |
| 0.55999483 |
| 0.56108437 |
| 0.56139491 |
| 0.56157858 |
| 0.56220884 |

| |
|---|
| 0.56220884 |
| 0.56248607 |
| 0.56253453 |
| 0.56279435 |
| 0.56299104 |
| 0.56347547 |
| 0.56351018 |
| 0.56404631 |
| 0.56404631 |
| 0.56419971 |
| 0.56519102 |
| 0.56620586 |
| 0.56661567 |
| 0.56661567 |
| 0.56712091 |
| 0.56712091 |
| 0.56747132 |
| 0.56781159 |
| 0.5683778 |
| 0.5683778 |
| 0.5683778 |
| 0.5683778 |
| 0.56859227 |
| 0.568634 |
| 0.56863449 |
| 0.56870407 |
| 0.56899949 |
| 0.56903927 |
| 0.56903927 |
| 0.56903927 |
| 0.56935598 |
| 0.56937185 |
| 0.569391 |
| 0.569391 |
| 0.5706194 |
| 0.5708144 |
| 0.57123297 |
| 0.57123297 |
| 0.57123297 |
| 0.57182404 |
| 0.57231671 |

| |
|---|
| 0.57231671 |
| 0.5729387 |
| 0.57314598 |
| 0.57314598 |
| 0.57406737 |
| 0.57471946 |
| 0.57471946 |
| 0.57513034 |
| 0.57513034 |
| 0.57513034 |
| 0.57592706 |
| 0.57592706 |
| 0.57592706 |
| 0.57621765 |
| 0.57636057 |
| 0.57636057 |
| 0.57696075 |
| 0.57732869 |
| 0.57762157 |
| 0.57779927 |
| 0.5785287 |
| 0.57880958 |
| 0.57880958 |
| 0.57884817 |
| 0.57931718 |
| 0.57931718 |
| 0.5800626 |
| 0.5800626 |
| 0.58021959 |
| 0.58021959 |
| 0.58027567 |
| 0.58062676 |
| 0.58102852 |
| 0.58124454 |
| 0.58203101 |
| 0.58296775 |
| 0.58299855 |
| 0.58336205 |
| 0.58371393 |
| 0.58475553 |
| 0.58518719 |

| |
|---|
| 0.58518719 |
| 0.58536215 |
| 0.58558855 |
| 0.58601745 |
| 0.58618824 |
| 0.58654917 |
| 0.58655242 |
| 0.58655242 |
| 0.5868269 |
| 0.58759797 |
| 0.58823379 |
| 0.58922698 |
| 0.58942122 |
| 0.59029365 |
| 0.59044923 |
| 0.59055268 |
| 0.59089799 |
| 0.591134 |
| 0.59134763 |
| 0.59134763 |
| 0.59176859 |
| 0.59184174 |
| 0.59194327 |
| 0.59247292 |
| 0.5931708 |
| 0.59330679 |
| 0.593389 |
| 0.59417926 |
| 0.59426374 |
| 0.59446886 |
| 0.59485091 |
| 0.59489282 |
| 0.59489282 |
| 0.59498891 |
| 0.59558262 |
| 0.59583592 |
| 0.59609031 |
| 0.59716788 |
| 0.59716788 |
| 0.59719668 |
| 0.59751852 |

| |
|---|
| 0.59761779 |
| 0.59782357 |
| 0.59789078 |
| 0.59805593 |
| 0.59833138 |
| 0.59838178 |
| 0.59838178 |
| 0.59854675 |
| 0.59870591 |
| 0.59870591 |
| 0.59912744 |
| 0.59949413 |
| 0.60031219 |
| 0.60034964 |
| 0.60195387 |
| 0.60236995 |
| 0.60236995 |
| 0.60249071 |
| 0.60283923 |
| 0.60283923 |
| 0.60298377 |
| 0.60298377 |
| 0.60341248 |
| 0.60402161 |
| 0.6041694 |
| 0.60425843 |
| 0.60441062 |
| 0.60488014 |
| 0.60521535 |
| 0.60527304 |
| 0.60562004 |
| 0.60585948 |
| 0.60589437 |
| 0.60593898 |
| 0.60627797 |
| 0.60640405 |
| 0.60640988 |
| 0.60640988 |
| 0.60687041 |
| 0.60687041 |
| 0.60687041 |

| |
|---|
| 0.60731357 |
| 0.60731357 |
| 0.60830939 |
| 0.60862808 |
| 0.60897036 |
| 0.60929077 |
| 0.60940695 |
| 0.60949988 |
| 0.60995821 |
| 0.61000931 |
| 0.61011682 |
| 0.61075514 |
| 0.61075514 |
| 0.61090495 |
| 0.61094727 |
| 0.61167627 |
| 0.61223084 |
| 0.61264601 |
| 0.61264601 |
| 0.61279648 |
| 0.61315782 |
| 0.61315782 |
| 0.61315782 |
| 0.61315782 |
| 0.61315782 |
| 0.61315782 |
| 0.61390838 |
| 0.61435693 |
| 0.61450496 |
| 0.61481552 |
| 0.61498447 |
| 0.61499954 |
| 0.61537437 |
| 0.61687081 |
| 0.61797481 |
| 0.61804 |
| 0.61812173 |
| 0.61812173 |
| 0.61887257 |
| 0.61935443 |
| 0.61935443 |

| |
|---|
| 0.61935512 |
| 0.61935512 |
| 0.61935512 |
| 0.61935512 |
| 0.61952808 |
| 0.62020783 |
| 0.62028686 |
| 0.62052299 |
| 0.62065272 |
| 0.62066558 |
| 0.6206931 |
| 0.6206931 |
| 0.62075865 |
| 0.62117089 |
| 0.62177958 |
| 0.62177958 |
| 0.62184002 |
| 0.62186241 |
| 0.6228061 |
| 0.6228061 |
| 0.62305605 |
| 0.62469633 |
| 0.62484578 |
| 0.62515475 |
| 0.62560678 |
| 0.62606197 |
| 0.62606197 |
| 0.62614464 |
| 0.62653645 |
| 0.62721862 |
| 0.62743079 |
| 0.62743079 |
| 0.62752441 |
| 0.62795699 |
| 0.62831908 |
| 0.62856455 |
| 0.62862281 |
| 0.62866628 |
| 0.62894503 |
| 0.62924493 |
| 0.63001773 |

| |
|---|
| 0.63002272 |
| 0.63045701 |
| 0.63065395 |
| 0.63106103 |
| 0.63141255 |
| 0.63141255 |
| 0.63176654 |
| 0.63176654 |
| 0.63192209 |
| 0.63241022 |
| 0.63256552 |
| 0.6335342 |
| 0.63357301 |
| 0.63392865 |
| 0.63407248 |
| 0.63498848 |
| 0.63556885 |
| 0.63556885 |
| 0.63556885 |
| 0.63556885 |
| 0.63557352 |
| 0.63650022 |
| 0.63724316 |
| 0.63726264 |
| 0.63726264 |
| 0.63741914 |
| 0.63759544 |
| 0.63788344 |
| 0.63831099 |
| 0.63881005 |
| 0.63974621 |
| 0.63998594 |
| 0.64008806 |
| 0.64062106 |
| 0.64062106 |
| 0.64062106 |
| 0.64065919 |
| 0.641057 |
| 0.64125301 |
| 0.64143225 |
| 0.64178104 |

| |
|---|
| 0.64189713 |
| 0.64213784 |
| 0.64220367 |
| 0.64227817 |
| 0.64316735 |
| 0.64371311 |
| 0.64371311 |
| 0.64388737 |
| 0.64413069 |
| 0.64468532 |
| 0.64468532 |
| 0.64468532 |
| 0.64472022 |
| 0.64561461 |
| 0.64589525 |
| 0.64589525 |
| 0.64622817 |
| 0.64685236 |
| 0.64685236 |
| 0.64696728 |
| 0.6469914 |
| 0.64750866 |
| 0.64797924 |
| 0.64819275 |
| 0.64832998 |
| 0.64855871 |
| 0.64861699 |
| 0.64894634 |
| 0.64989157 |
| 0.65012384 |
| 0.65020572 |
| 0.65020572 |
| 0.65020572 |
| 0.65058896 |
| 0.65069809 |
| 0.65135816 |
| 0.65139362 |
| 0.65154305 |
| 0.65156403 |
| 0.65156403 |
| 0.65157647 |

| |
|---|
| 0.65207718 |
| 0.65213872 |
| 0.65217976 |
| 0.65250134 |
| 0.65255899 |
| 0.65302776 |
| 0.65318548 |
| 0.65346247 |
| 0.65378209 |
| 0.6543296 |
| 0.6543296 |
| 0.65470049 |
| 0.65476288 |
| 0.65479148 |
| 0.65546704 |
| 0.65552135 |
| 0.65561317 |
| 0.65561317 |
| 0.65561317 |
| 0.65561317 |
| 0.65561317 |
| 0.65754496 |
| 0.65881044 |
| 0.6589738 |
| 0.6589738 |
| 0.65924804 |
| 0.65995853 |
| 0.66011077 |
| 0.66015479 |
| 0.66059565 |
| 0.66059565 |
| 0.6610208 |
| 0.66113295 |
| 0.66148893 |
| 0.66162725 |
| 0.66162725 |
| 0.66162725 |
| 0.6617037 |
| 0.6617037 |
| 0.66240423 |
| 0.66249315 |

| |
|---|
| 0.66321717 |
| 0.66358102 |
| 0.66387689 |
| 0.66414127 |
| 0.6643917 |
| 0.664585 |
| 0.6654319 |
| 0.6654319 |
| 0.66551934 |
| 0.6656854 |
| 0.6657081 |
| 0.66580909 |
| 0.66580909 |
| 0.66580909 |
| 0.66615285 |
| 0.66619336 |
| 0.66652821 |
| 0.66654059 |
| 0.66726026 |
| 0.66726026 |
| 0.66726026 |
| 0.66726026 |
| 0.66726026 |
| 0.66726026 |
| 0.66726026 |
| 0.66726026 |
| 0.66822703 |
| 0.66824138 |
| 0.66824138 |
| 0.66878813 |
| 0.6688426 |
| 0.66916408 |
| 0.669174 |
| 0.669174 |
| 0.66966577 |
| 0.66979474 |
| 0.67000952 |
| 0.67005964 |
| 0.67159333 |
| 0.67170865 |
| 0.67207376 |

| |
|---|
| 0.67226309 |
| 0.67339927 |
| 0.67339927 |
| 0.67360314 |
| 0.67430411 |
| 0.67431045 |
| 0.67431173 |
| 0.67465234 |
| 0.67465234 |
| 0.67465234 |
| 0.67496329 |
| 0.67570812 |
| 0.67643102 |
| 0.67643102 |
| 0.6765782 |
| 0.67681091 |
| 0.67740642 |
| 0.67753196 |
| 0.67755481 |
| 0.67755481 |
| 0.67812235 |
| 0.67828958 |
| 0.67923173 |
| 0.67943361 |
| 0.68010262 |
| 0.68010262 |
| 0.68042816 |
| 0.68138782 |
| 0.6814778 |
| 0.68151834 |
| 0.6818448 |
| 0.68224059 |
| 0.68273576 |
| 0.68340719 |
| 0.68389556 |
| 0.68424153 |
| 0.68461318 |
| 0.68508243 |
| 0.68574282 |
| 0.68578847 |
| 0.68587956 |

| |
|---|
| 0.68594688 |
| 0.68595036 |
| 0.68641036 |
| 0.68669238 |
| 0.68712178 |
| 0.68712178 |
| 0.68712178 |
| 0.68748128 |
| 0.68753269 |
| 0.68760774 |
| 0.68798218 |
| 0.6880413 |
| 0.6880413 |
| 0.68864002 |
| 0.68864002 |
| 0.68879255 |
| 0.68904047 |
| 0.68959683 |
| 0.69063935 |
| 0.69085345 |
| 0.69124238 |
| 0.69138071 |
| 0.69138071 |
| 0.69138071 |
| 0.69180804 |
| 0.69194175 |
| 0.69203042 |
| 0.69267744 |
| 0.69267744 |
| 0.69287721 |
| 0.69287721 |
| 0.69383166 |
| 0.69393394 |
| 0.69393394 |
| 0.69393394 |
| 0.69393394 |
| 0.6939665 |
| 0.69478691 |
| 0.69572052 |
| 0.69573499 |
| 0.69575733 |

| |
|---|
| 0.69575733 |
| 0.69575733 |
| 0.69575733 |
| 0.69613595 |
| 0.69635071 |
| 0.69649522 |
| 0.69653468 |
| 0.69662627 |
| 0.69757294 |
| 0.69763438 |
| 0.69763539 |
| 0.69798583 |
| 0.69800997 |
| 0.69814366 |
| 0.69833244 |
| 0.69841507 |
| 0.69864337 |
| 0.69869578 |
| 0.69875533 |
| 0.69898428 |
| 0.699654 |
| 0.699654 |
| 0.70003318 |
| 0.70037325 |
| 0.70056913 |
| 0.70056913 |
| 0.70056913 |
| 0.70084023 |
| 0.70102741 |
| 0.70143749 |
| 0.70167862 |
| 0.70249622 |
| 0.70249622 |
| 0.70252489 |
| 0.70252489 |
| 0.70296267 |
| 0.70313841 |
| 0.70334576 |
| 0.70334576 |
| 0.70340217 |
| 0.70340217 |

| |
|---|
| 0.70340217 |
| 0.70340217 |
| 0.70340217 |
| 0.70340217 |
| 0.70342499 |
| 0.70357427 |
| 0.70357427 |
| 0.70391838 |
| 0.70400791 |
| 0.7041201 |
| 0.70467208 |
| 0.704797 |
| 0.70525765 |
| 0.70531838 |
| 0.70629104 |
| 0.70662726 |
| 0.70669978 |
| 0.70669978 |
| 0.70669978 |
| 0.70707625 |
| 0.70742434 |
| 0.70742434 |
| 0.7077473 |
| 0.7077473 |
| 0.70799764 |
| 0.7082629 |
| 0.70831799 |
| 0.70843031 |
| 0.70876066 |
| 0.70912991 |
| 0.71016045 |
| 0.71016045 |
| 0.71016045 |
| 0.71026598 |
| 0.71026598 |
| 0.71096181 |
| 0.71117892 |
| 0.71170987 |
| 0.71170987 |
| 0.71201401 |
| 0.71253317 |

| |
|---|
| 0.71253317 |
| 0.71318566 |
| 0.71325371 |
| 0.71341928 |
| 0.71341928 |
| 0.71402769 |
| 0.71407916 |
| 0.71410365 |
| 0.71410365 |
| 0.71428461 |
| 0.71436599 |
| 0.71461722 |
| 0.71461722 |
| 0.71461722 |
| 0.71465433 |
| 0.71465433 |
| 0.71465433 |
| 0.71484041 |
| 0.71516178 |
| 0.71516178 |
| 0.71536684 |
| 0.71572119 |
| 0.71572166 |
| 0.71670272 |
| 0.71710823 |
| 0.71714328 |
| 0.71721815 |
| 0.71754 |
| 0.71757426 |
| 0.71767159 |
| 0.71804003 |
| 0.7180841 |
| 0.7180841 |
| 0.71829781 |
| 0.71829781 |
| 0.71907228 |
| 0.72029878 |
| 0.72029878 |
| 0.72110766 |
| 0.72200791 |
| 0.72217743 |

| |
|---|
| 0.72387765 |
| 0.72387765 |
| 0.72387889 |
| 0.72387889 |
| 0.72433544 |
| 0.72436426 |
| 0.7256575 |
| 0.72674849 |
| 0.72677597 |
| 0.72677597 |
| 0.72710496 |
| 0.72734262 |
| 0.72771449 |
| 0.72771449 |
| 0.72771449 |
| 0.72776189 |
| 0.72865684 |
| 0.7287001 |
| 0.72875755 |
| 0.72910191 |
| 0.72936488 |
| 0.729791 |
| 0.729791 |
| 0.73026632 |
| 0.73044096 |
| 0.73044096 |
| 0.73074391 |
| 0.73104156 |
| 0.73112963 |
| 0.73152858 |
| 0.73154973 |
| 0.73183203 |
| 0.73218889 |
| 0.73240003 |
| 0.73240003 |
| 0.73240003 |
| 0.73242287 |
| 0.73245334 |
| 0.73332193 |
| 0.73371695 |
| 0.73371695 |

| |
|---|
| 0.73371695 |
| 0.73371695 |
| 0.73401964 |
| 0.73473492 |
| 0.73498518 |
| 0.73558778 |
| 0.73558778 |
| 0.73570352 |
| 0.73573048 |
| 0.73598531 |
| 0.73599218 |
| 0.73614141 |
| 0.73669857 |
| 0.73722087 |
| 0.73722087 |
| 0.73750334 |
| 0.73750334 |
| 0.73750334 |
| 0.73750334 |
| 0.73766852 |
| 0.73771368 |
| 0.73786888 |
| 0.73879966 |
| 0.7397402 |
| 0.74025269 |
| 0.74054808 |
| 0.74152558 |
| 0.74152558 |
| 0.74152558 |
| 0.74160121 |
| 0.7426901 |
| 0.7426901 |
| 0.74269454 |
| 0.74281205 |
| 0.74326174 |
| 0.74381796 |
| 0.74400021 |
| 0.7440111 |
| 0.7440111 |
| 0.74437726 |
| 0.74458459 |

| |
|---|
| 0.74561395 |
| 0.74561395 |
| 0.74561395 |
| 0.74575172 |
| 0.74618107 |
| 0.74687237 |
| 0.74754174 |
| 0.7479085 |
| 0.7479085 |
| 0.7479085 |
| 0.74873368 |
| 0.74873368 |
| 0.74935708 |
| 0.74946977 |
| 0.74963526 |
| 0.74968812 |
| 0.75041386 |
| 0.75080057 |
| 0.75093122 |
| 0.75135671 |
| 0.75150053 |
| 0.75155861 |
| 0.75172602 |
| 0.75174158 |
| 0.75200302 |
| 0.75203379 |
| 0.75203379 |
| 0.75251293 |
| 0.7532966 |
| 0.75347949 |
| 0.7537923 |
| 0.75405066 |
| 0.75417772 |
| 0.75417772 |
| 0.7543691 |
| 0.75455342 |
| 0.75500923 |
| 0.75550317 |
| 0.75610767 |
| 0.75631978 |
| 0.7564402 |

| |
|---|
| 0.75674664 |
| 0.75752114 |
| 0.75759042 |
| 0.75772347 |
| 0.75772347 |
| 0.75806228 |
| 0.75824284 |
| 0.75825902 |
| 0.75825902 |
| 0.7583365 |
| 0.7583772 |
| 0.7583772 |
| 0.75870566 |
| 0.75910281 |
| 0.75957799 |
| 0.75957799 |
| 0.7601561 |
| 0.76030481 |
| 0.76040216 |
| 0.76047925 |
| 0.76047925 |
| 0.76056427 |
| 0.76056427 |
| 0.76056427 |
| 0.76075201 |
| 0.76093115 |
| 0.76125417 |
| 0.76132331 |
| 0.76132331 |
| 0.76132331 |
| 0.76132331 |
| 0.76132713 |
| 0.76181344 |
| 0.7620283 |
| 0.76210116 |
| 0.76237918 |
| 0.76280746 |
| 0.76280746 |
| 0.76329107 |
| 0.76329107 |
| 0.76341161 |

| |
|---|
| 0.76341161 |
| 0.76353984 |
| 0.76380951 |
| 0.76400343 |
| 0.76426859 |
| 0.76426859 |
| 0.76492767 |
| 0.76522612 |
| 0.76531582 |
| 0.76608013 |
| 0.76608013 |
| 0.76612359 |
| 0.76612359 |
| 0.76612359 |
| 0.76612359 |
| 0.76612359 |
| 0.76666543 |
| 0.76688288 |
| 0.76688288 |
| 0.76721958 |
| 0.7675235 |
| 0.7675235 |
| 0.76788521 |
| 0.76801555 |
| 0.76801555 |
| 0.76801555 |
| 0.76803409 |
| 0.76834777 |
| 0.76851352 |
| 0.76854777 |
| 0.76887473 |
| 0.76892645 |
| 0.76893356 |
| 0.76893356 |
| 0.76908762 |
| 0.76934057 |
| 0.76972645 |
| 0.76972645 |
| 0.76984091 |
| 0.77146891 |
| 0.77148497 |

0.77159002
0.77159804
0.77213538
0.77272462
0.77326758
0.77373006
0.77373006
0.77378924
0.77395014
0.77395014
0.77446175
0.77464956
0.77482971
0.77484325
0.77557998
0.77563214
0.77595632
0.77621681
0.77653348
0.77683041
0.77762015
0.77764994
0.77764994
0.77764994
0.77764994
0.77764994
0.77772704
0.77772704
0.77830749
0.77831657
0.78060595
0.78060595
0.78073255
0.78077727
0.78134813
0.78173021
0.78191742
0.78191742
0.78191742
0.78191742
0.78195071

| |
|---|
| 0.78195588 |
| 0.78195588 |
| 0.78230961 |
| 0.78261013 |
| 0.78263215 |
| 0.78323482 |
| 0.78328778 |
| 0.78360085 |
| 0.78386782 |
| 0.78389977 |
| 0.78389977 |
| 0.78416668 |
| 0.78417972 |
| 0.78417972 |
| 0.7847835 |
| 0.78494325 |
| 0.78497425 |
| 0.78533106 |
| 0.78573885 |
| 0.78573885 |
| 0.78584544 |
| 0.78584544 |
| 0.78584544 |
| 0.78584544 |
| 0.78601618 |
| 0.78601618 |
| 0.78642054 |
| 0.78684468 |
| 0.78684468 |
| 0.78685815 |
| 0.78819238 |
| 0.78819238 |
| 0.78823347 |
| 0.7884591 |
| 0.7884591 |
| 0.78898448 |
| 0.78912148 |
| 0.78938809 |
| 0.78940626 |
| 0.78947235 |
| 0.78947235 |

| |
|---|
| 0.78947235 |
| 0.78947235 |
| 0.78963317 |
| 0.78975513 |
| 0.79017667 |
| 0.79057624 |
| 0.79057624 |
| 0.79057624 |
| 0.79093525 |
| 0.79104859 |
| 0.79117297 |
| 0.79131009 |
| 0.79171458 |
| 0.79173828 |
| 0.79269114 |
| 0.7935478 |
| 0.79430831 |
| 0.79463201 |
| 0.79463201 |
| 0.79470019 |
| 0.79492684 |
| 0.79501889 |
| 0.79536901 |
| 0.79560722 |
| 0.7962052 |
| 0.79686459 |
| 0.79686459 |
| 0.79692633 |
| 0.79817839 |
| 0.7987248 |
| 0.79891078 |
| 0.79918396 |
| 0.79918916 |
| 0.801969 |
| 0.801969 |
| 0.80200225 |
| 0.80220044 |
| 0.80254984 |
| 0.80413551 |
| 0.80416213 |
| 0.80416213 |

0.80455905
0.80480183
0.80502303
0.80554333
0.80554333
0.80609784
0.80729995
0.80729997
0.80729997
0.80734046
0.80735254
0.80735254
0.80882566
0.80898736
0.80953821
0.80969731
0.80979171
0.81029493
0.81038786
0.81038786
0.81038786
0.81099166
0.81099166
0.81183163
0.81211906
0.81218858
0.81242944
0.81286837
0.81350171
0.8136807
0.81369793
0.81453448
0.81497011
0.81517337
0.81524423
0.81546568
0.81546568
0.81552957
0.81601245
0.81619776
0.81731161

| |
|---|
| 0.81731161 |
| 0.81732864 |
| 0.81764291 |
| 0.81764291 |
| 0.81764291 |
| 0.81764291 |
| 0.81791962 |
| 0.8180632 |
| 0.81844782 |
| 0.81853588 |
| 0.81918188 |
| 0.81925934 |
| 0.81925934 |
| 0.81987048 |
| 0.8200087 |
| 0.8200087 |
| 0.8200087 |
| 0.8200087 |
| 0.8200087 |
| 0.82013433 |
| 0.82038785 |
| 0.82119684 |
| 0.82128679 |
| 0.82128679 |
| 0.82149085 |
| 0.82184992 |
| 0.82184992 |
| 0.82205861 |
| 0.82235174 |
| 0.82236222 |
| 0.82240168 |
| 0.82272345 |
| 0.82395466 |
| 0.82395466 |
| 0.82434849 |
| 0.82435972 |
| 0.82448893 |
| 0.82489712 |
| 0.82493497 |
| 0.82553694 |
| 0.82670801 |

| |
|---|
| 0.82670801 |
| 0.82670801 |
| 0.82690855 |
| 0.82742999 |
| 0.82742999 |
| 0.82742999 |
| 0.82754941 |
| 0.82754941 |
| 0.82757515 |
| 0.82812109 |
| 0.82812109 |
| 0.82839528 |
| 0.82845753 |
| 0.82847418 |
| 0.82994238 |
| 0.83007661 |
| 0.83036838 |
| 0.83036838 |
| 0.83040272 |
| 0.83156512 |
| 0.83172579 |
| 0.83183594 |
| 0.83183594 |
| 0.83193058 |
| 0.83327428 |
| 0.83337001 |
| 0.83399345 |
| 0.83412263 |
| 0.83412263 |
| 0.83447662 |
| 0.83447662 |
| 0.83523614 |
| 0.83651032 |
| 0.83738046 |
| 0.83748231 |
| 0.83748231 |
| 0.83754091 |
| 0.83754091 |
| 0.83754091 |
| 0.83754091 |
| 0.83842241 |

| |
|---|
| 0.84018492 |
| 0.84067138 |
| 0.84067138 |
| 0.84067138 |
| 0.84067138 |
| 0.841035 |
| 0.84119041 |
| 0.84126018 |
| 0.84280247 |
| 0.84300613 |
| 0.84300613 |
| 0.84328113 |
| 0.84344713 |
| 0.84353938 |
| 0.84374224 |
| 0.84416221 |
| 0.84426638 |
| 0.84473637 |
| 0.8451616 |
| 0.84518965 |
| 0.84518965 |
| 0.84518965 |
| 0.84580106 |
| 0.84585912 |
| 0.84623038 |
| 0.8470435 |
| 0.84742064 |
| 0.84750228 |
| 0.84758955 |
| 0.84846694 |
| 0.84846694 |
| 0.84848385 |
| 0.84850001 |
| 0.84850001 |
| 0.84876112 |
| 0.84876304 |
| 0.84876304 |
| 0.84876304 |
| 0.84903432 |
| 0.84980237 |
| 0.84980237 |

| |
|---|
| 0.84982686 |
| 0.84982686 |
| 0.84982686 |
| 0.84982686 |
| 0.84992896 |
| 0.84995602 |
| 0.84995602 |
| 0.85060259 |
| 0.85072362 |
| 0.85282705 |
| 0.85289922 |
| 0.85307117 |
| 0.85307117 |
| 0.85329491 |
| 0.8536306 |
| 0.8536306 |
| 0.85366138 |
| 0.85408643 |
| 0.85408643 |
| 0.85408643 |
| 0.85408643 |
| 0.85487047 |
| 0.8554409 |
| 0.85551711 |
| 0.85593275 |
| 0.85620908 |
| 0.85620908 |
| 0.85620908 |
| 0.8571141 |
| 0.8571141 |
| 0.8571141 |
| 0.8571141 |
| 0.8571141 |
| 0.85734268 |
| 0.85734268 |
| 0.85734438 |
| 0.85751616 |
| 0.85769484 |
| 0.85805268 |
| 0.85896005 |
| 0.85997371 |

| |
|---|
| 0.86008669 |
| 0.86008669 |
| 0.86023973 |
| 0.86023973 |
| 0.86057759 |
| 0.86093018 |
| 0.86127785 |
| 0.86127785 |
| 0.86129567 |
| 0.86160487 |
| 0.86229906 |
| 0.86318546 |
| 0.86318546 |
| 0.86386845 |
| 0.86390045 |
| 0.86461943 |
| 0.86472095 |
| 0.8647857 |
| 0.86573334 |
| 0.86590256 |
| 0.86596413 |
| 0.86655468 |
| 0.86655468 |
| 0.86665994 |
| 0.86750062 |
| 0.86837109 |
| 0.86921377 |
| 0.86921377 |
| 0.86921377 |
| 0.86921377 |
| 0.86965356 |
| 0.87001864 |
| 0.87001864 |
| 0.87009198 |
| 0.87140963 |
| 0.87140963 |
| 0.87143159 |
| 0.87143159 |
| 0.87157415 |
| 0.87157872 |
| 0.87157872 |

| |
|---|
| 0.87170279 |
| 0.8718434 |
| 0.87187478 |
| 0.8722786 |
| 0.87260034 |
| 0.87260034 |
| 0.87304773 |
| 0.8738689 |
| 0.87412945 |
| 0.8746099 |
| 0.87477183 |
| 0.87596624 |
| 0.87596624 |
| 0.87643589 |
| 0.87656155 |
| 0.87680282 |
| 0.87719893 |
| 0.87719893 |
| 0.8775613 |
| 0.8775613 |
| 0.87794555 |
| 0.87835978 |
| 0.87856367 |
| 0.87993198 |
| 0.87993198 |
| 0.88098443 |
| 0.88141206 |
| 0.88192366 |
| 0.88231737 |
| 0.88267164 |
| 0.88339126 |
| 0.88349739 |
| 0.88393808 |
| 0.88512587 |
| 0.88538651 |
| 0.88563253 |
| 0.8856494 |
| 0.8856494 |
| 0.88638744 |
| 0.88654571 |
| 0.88654571 |

| |
|---|
| 0.88708005 |
| 0.88714761 |
| 0.88765866 |
| 0.88767799 |
| 0.88795012 |
| 0.88795012 |
| 0.88795012 |
| 0.88842284 |
| 0.88880446 |
| 0.8896718 |
| 0.8896718 |
| 0.89014437 |
| 0.89014437 |
| 0.89016954 |
| 0.89080282 |
| 0.89244326 |
| 0.89255661 |
| 0.89349396 |
| 0.89381038 |
| 0.89528159 |
| 0.89563212 |
| 0.89563813 |
| 0.89570033 |
| 0.89626481 |
| 0.89626481 |
| 0.89713137 |
| 0.89763475 |
| 0.89781287 |
| 0.89781287 |
| 0.89781287 |
| 0.89795701 |
| 0.89801923 |
| 0.8987807 |
| 0.89900846 |
| 0.89996422 |
| 0.90071092 |
| 0.90071092 |
| 0.90071092 |
| 0.90071092 |
| 0.90071092 |
| 0.90071092 |

| |
|---|
| 0.90071092 |
| 0.90071092 |
| 0.90071092 |
| 0.90071092 |
| 0.90103979 |
| 0.90133051 |
| 0.90142961 |
| 0.90159586 |
| 0.90185819 |
| 0.90206743 |
| 0.90267956 |
| 0.90279844 |
| 0.90279844 |
| 0.9029913 |
| 0.9029913 |
| 0.9029913 |
| 0.90315857 |
| 0.90315857 |
| 0.90338279 |
| 0.9034229 |
| 0.90349644 |
| 0.90369563 |
| 0.90369563 |
| 0.9041297 |
| 0.90421948 |
| 0.9049657 |
| 0.9049657 |
| 0.90514685 |
| 0.90551425 |
| 0.90584098 |
| 0.90588154 |
| 0.90588154 |
| 0.90625119 |
| 0.90674319 |
| 0.90702517 |
| 0.9071951 |
| 0.90726511 |
| 0.90791835 |
| 0.90812195 |
| 0.90843027 |
| 0.90881229 |

| |
|---|
| 0.90948687 |
| 0.91016299 |
| 0.91055248 |
| 0.91055278 |
| 0.91055278 |
| 0.91078126 |
| 0.91100778 |
| 0.91106499 |
| 0.91107596 |
| 0.91107596 |
| 0.91108061 |
| 0.91112144 |
| 0.91134604 |
| 0.91134604 |
| 0.91168945 |
| 0.91196129 |
| 0.91206577 |
| 0.91206577 |
| 0.91217027 |
| 0.91217027 |
| 0.91224617 |
| 0.91262834 |
| 0.912703 |
| 0.91274516 |
| 0.91275721 |
| 0.91275721 |
| 0.91306431 |
| 0.91306431 |
| 0.91306431 |
| 0.91306431 |
| 0.91306431 |
| 0.91313936 |
| 0.91351292 |
| 0.91351292 |
| 0.91367877 |
| 0.91500578 |
| 0.91537425 |
| 0.91537425 |
| 0.91537425 |
| 0.91537425 |
| 0.91537425 |

| |
|---|
| 0.9156343 |
| 0.9159453 |
| 0.91667844 |
| 0.91704221 |
| 0.91769379 |
| 0.91797391 |
| 0.91818605 |
| 0.9186104 |
| 0.9186104 |
| 0.9191784 |
| 0.91929334 |
| 0.92039856 |
| 0.92044227 |
| 0.92044465 |
| 0.92047861 |
| 0.92050405 |
| 0.92050405 |
| 0.92116358 |
| 0.92156583 |
| 0.92158103 |
| 0.92199944 |
| 0.92204174 |
| 0.92204174 |
| 0.92252284 |
| 0.92252284 |
| 0.92359759 |
| 0.92359759 |
| 0.92359759 |
| 0.92359759 |
| 0.92390229 |
| 0.92411095 |
| 0.92469118 |
| 0.92481428 |
| 0.92566681 |
| 0.92566681 |
| 0.92610479 |
| 0.92729446 |
| 0.92746574 |
| 0.92762277 |
| 0.92803947 |
| 0.92817286 |

| |
|---|
| 0.92817286 |
| 0.92817286 |
| 0.92850202 |
| 0.92907381 |
| 0.92960369 |
| 0.92960369 |
| 0.9296158 |
| 0.93052899 |
| 0.93057426 |
| 0.9306171 |
| 0.93066529 |
| 0.93091729 |
| 0.93267392 |
| 0.93304786 |
| 0.93304786 |
| 0.93336553 |
| 0.93336553 |
| 0.9334642 |
| 0.93437285 |
| 0.93437285 |
| 0.93437285 |
| 0.93437285 |
| 0.93452738 |
| 0.93457191 |
| 0.93457191 |
| 0.93457191 |
| 0.93457191 |
| 0.93457191 |
| 0.93543375 |
| 0.93598764 |
| 0.93665067 |
| 0.93665067 |
| 0.93858768 |
| 0.93877378 |
| 0.93947211 |
| 0.93947211 |
| 0.93972457 |
| 0.94004008 |
| 0.94023594 |
| 0.94042613 |
| 0.94042613 |

0.94042613
0.94080418
0.94106009
0.94106009
0.94109804
0.94121887
0.94159343
0.94203912
0.94215034
0.94215034
0.94267732
0.94322225
0.94322225
0.94323625
0.94392548
0.94392548
0.94393302
0.94456222
0.94507718
0.94522737
0.94535011
0.94539347
0.94539347
0.94568157
0.94572287
0.94572287
0.94631719
0.94631719
0.94636485
0.94657041
0.94657041
0.94657041
0.94694784
0.94694784
0.94745702
0.94753195
0.94765736
0.94765736
0.94765736
0.94851144
0.94883128

| |
|---|
| 0.94916053 |
| 0.94959099 |
| 0.94959099 |
| 0.94959099 |
| 0.94959099 |
| 0.9498375 |
| 0.95021932 |
| 0.95021932 |
| 0.95021932 |
| 0.95072849 |
| 0.95111544 |
| 0.9514829 |
| 0.95159293 |
| 0.95211788 |
| 0.95211788 |
| 0.95226775 |
| 0.95318385 |
| 0.95355242 |
| 0.95355242 |
| 0.95370227 |
| 0.9537049 |
| 0.95448168 |
| 0.9545571 |
| 0.95460486 |
| 0.95460486 |
| 0.955165 |
| 0.95528212 |
| 0.95546053 |
| 0.95573931 |
| 0.95578744 |
| 0.95595698 |
| 0.95595698 |
| 0.95599795 |
| 0.95636866 |
| 0.95637457 |
| 0.95678984 |
| 0.95734008 |
| 0.95734008 |
| 0.95769448 |
| 0.95769448 |
| 0.95860803 |

| |
|---|
| 0.95886456 |
| 0.95888235 |
| 0.95888235 |
| 0.95914392 |
| 0.95914392 |
| 0.95937465 |
| 0.95971506 |
| 0.95971506 |
| 0.95971506 |
| 0.95971506 |
| 0.95990273 |
| 0.96006162 |
| 0.9601566 |
| 0.96026162 |
| 0.96053615 |
| 0.96154688 |
| 0.96156469 |
| 0.96156469 |
| 0.96156469 |
| 0.96156469 |
| 0.96178457 |
| 0.96306123 |
| 0.96492525 |
| 0.96519871 |
| 0.96578352 |
| 0.96585706 |
| 0.96619737 |
| 0.96619737 |
| 0.96619737 |
| 0.96629248 |
| 0.96658906 |
| 0.96704686 |
| 0.96704686 |
| 0.9673977 |
| 0.96772555 |
| 0.96888704 |
| 0.96896978 |
| 0.96896978 |
| 0.97010564 |
| 0.97059072 |
| 0.97059072 |

|  |
| --- |
| 0.97070798 |
| 0.97070798 |
| 0.97078856 |
| 0.97137064 |
| 0.97176314 |
| 0.97176314 |
| 0.97176314 |
| 0.97176314 |
| 0.97176314 |
| 0.97209416 |
| 0.97210312 |
| 0.97276185 |
| 0.9728677 |
| 0.97309754 |
| 0.97342752 |
| 0.97413234 |
| 0.97418554 |
| 0.97435471 |
| 0.97449069 |
| 0.97449069 |
| 0.97531242 |
| 0.97531242 |
| 0.97531242 |
| 0.97531242 |
| 0.97531242 |
| 0.97531242 |
| 0.97531242 |
| 0.97531242 |
| 0.97531242 |
| 0.97606688 |
| 0.97606688 |
| 0.97606688 |
| 0.977129 |
| 0.97745549 |
| 0.97745549 |
| 0.97745549 |
| 0.97745549 |
| 0.97747708 |
| 0.97755749 |
| 0.97755749 |
| 0.97788665 |

| |
|---|
| 0.97805225 |
| 0.97805225 |
| 0.97820888 |
| 0.97975354 |
| 0.97975354 |
| 0.97979069 |
| 0.97979069 |
| 0.97979069 |
| 0.98008307 |
| 0.98017725 |
| 0.98026738 |
| 0.98026738 |
| 0.9805061 |
| 0.9805061 |
| 0.98050949 |
| 0.98050949 |
| 0.98050949 |
| 0.98050949 |
| 0.98050949 |
| 0.98051222 |
| 0.98051222 |
| 0.98051222 |
| 0.98051222 |
| 0.98089134 |
| 0.98089134 |
| 0.98089134 |
| 0.98102633 |
| 0.98108079 |
| 0.98127256 |
| 0.98145255 |
| 0.98180632 |
| 0.98259897 |
| 0.98259897 |
| 0.98261843 |
| 0.98263125 |
| 0.98292481 |
| 0.9833241 |
| 0.9833241 |
| 0.98500789 |
| 0.98531905 |
| 0.98531905 |

| |
|---|
| 0.98531905 |
| 0.98533802 |
| 0.98533802 |
| 0.98549131 |
| 0.98559399 |
| 0.98629177 |
| 0.98629177 |
| 0.98631021 |
| 0.98651353 |
| 0.98651353 |
| 0.98729477 |
| 0.98729789 |
| 0.98729789 |
| 0.98761415 |
| 0.98774032 |
| 0.98802867 |
| 0.98902396 |
| 0.98935076 |
| 0.98965117 |
| 0.99010328 |
| 0.99010933 |
| 0.99075789 |
| 0.99092071 |
| 0.99106378 |
| 0.99165725 |
| 0.99178783 |
| 0.99179008 |
| 0.99179008 |
| 0.99196909 |
| 0.99223357 |
| 0.99367645 |
| 0.99400605 |
| 0.99400605 |
| 0.99468758 |
| 0.99540002 |
| 0.99540002 |
| 0.99540002 |
| 0.99561742 |
| 0.99605186 |
| 0.99621876 |
| 0.99621876 |

| |
|---|
| 0.996372 |
| 0.9965514 |
| 0.99656549 |
| 0.99656549 |
| 0.9970105 |
| 0.9970105 |
| 0.99711904 |
| 0.99711904 |
| 0.9972006 |
| 0.99778432 |
| 0.99825043 |
| 0.99837867 |
| 0.99861162 |
| 0.99878864 |
| 0.99953294 |
| 0.99953294 |
| 0.99957591 |
| 0.99957591 |
| 0.99957591 |
| 0.99963991 |
| 0.99972153 |
| NA |
| NA |
| NA |
| NA |
| NA |
| NA |
| NA |
| NA |
| NA |
| NA |
| NA |
| NA |
| NA |
| NA |
| NA |
| NA |
| NA |
| NA |
| NA |
| NA |

NA
NA
NA
NA
NA
NA
NA
NA
NA
NA
NA
NA
NA
NA
NA
NA
NA
NA
NA
NA
NA
NA
NA
NA
NA
NA
NA
NA
NA
NA
NA
NA
NA
NA
NA
NA
NA
NA
NA
NA
NA

NA
NA
NA
NA
NA
NA
NA
NA
NA
NA
NA
NA
NA
NA
NA
NA
NA
NA
NA
NA
NA
NA
NA
NA
NA
NA
NA
NA
NA
NA
NA
NA
NA
NA
NA
NA
NA
NA
NA
NA
NA

NA
NA
NA
NA
NA
NA
NA
NA
NA
NA
NA
NA
NA
NA
NA
NA
NA
NA
NA
NA
NA
NA
NA
NA
NA
NA
NA
NA
NA
NA
NA
NA
NA
NA
NA
NA
NA
NA
NA
NA
NA

NA
NA
NA
NA
NA
NA
NA
NA
NA
NA
NA
NA
NA
NA
NA
NA
NA
NA
NA
NA
NA
NA
NA
NA
NA
NA
NA
NA
NA
NA
NA
NA
NA
NA
NA
NA
NA
NA
NA
NA
NA

NA
NA
NA
NA
NA
NA
NA
NA
NA
NA
NA
NA
NA
NA
NA
NA
NA
NA
NA
NA
NA
NA
NA
NA
NA
NA
NA
NA
NA
NA
NA
NA
NA
NA
NA
NA
NA
NA
NA
NA
NA

NA
NA
NA
NA
NA
NA
NA
NA
NA
NA
NA
NA
NA
NA
NA
NA
NA
NA
NA
NA
NA
NA
NA
NA
NA
NA
NA
NA
NA
NA
NA
NA
NA
NA
NA
NA
NA
NA
NA
NA
NA

NA
NA
NA
NA
NA
NA
NA
NA
NA
NA
NA
NA
NA
NA
NA
NA
NA
NA
NA
NA
NA
NA
NA
NA
NA
NA
NA
NA
NA
NA
NA
NA
NA
NA
NA
NA
NA
NA
NA
NA
NA

NA
NA
NA
NA
NA
NA
NA
NA
NA
NA
NA
NA
NA
NA
NA
NA
NA
NA
NA
NA
NA
NA
NA
NA
NA
NA
NA
NA
NA
NA
NA
NA
NA
NA
NA
NA
NA
NA
NA
NA
NA

NA
NA
